# Structural homology screens reveal poxvirus-encoded proteins impacting inflammasome-mediated defenses

**DOI:** 10.1101/2023.02.26.529821

**Authors:** Ian N. Boys, Alex G. Johnson, Meghan Quinlan, Philip J. Kranzusch, Nels C. Elde

**Affiliations:** Department of Human Genetics, University of Utah, Salt Lake City, Utah, 84112 USA; Department of Microbiology, Harvard Medical School, Boston, MA, 02115, USA; Department of Cancer Immunology and Virology, Dana-Farber Cancer Institute, Boston, MA, 02115, USA; Howard Hughes Medical Institute, Chevy Chase, Maryland, 20815, USA

## Abstract

Viruses acquire host genes via horizontal gene transfer and can express them to manipulate host biology during infections. Some viral and host homologs retain sequence identity, but evolutionary divergence can obscure host origins. We used structural modeling to compare vaccinia virus proteins with metazoan proteomes. We identified vaccinia *A47L* as a homolog of gasdermins, the executioners of pyroptosis. An X-ray crystal structure of A47 confirmed this homology and cell-based assays revealed that A47 inhibits pyroptosis. We also identified vaccinia *C1L* as the product of a cryptic gene fusion event coupling a Bcl-2 related fold with a pyrin domain. C1 associates with components of the inflammasome, a cytosolic innate immune sensor involved in pyroptosis, yet paradoxically enhances inflammasome activity, suggesting a benefit to poxvirus replication in some circumstances. Our findings demonstrate the potential of structural homology screens to reveal genes that viruses capture from hosts and repurpose to benefit viral fitness.

## Introduction

DNA viruses are particularly prominent “gene hunters” that capture a variety of host genes and express them in ways that alter host biology to promote viral replication ^2,3^. While the mechanisms facilitating gene capture are beginning to become clear ^4,5^, the identity and function of many host-derived viral genes remain unknown. In some cases, sequence-based comparisons clearly reveal identity between cellular and viral genes, and many crucial discoveries in cell biology and virology originated from such comparisons ^6^. However, in many instances rapid evolution and sequence divergence obscure homology with host genes.

While protein sequences diverge over time, critical structural elements are often preserved. Structural comparison is thus useful for inferring the origins and functions of viral proteins. For example, previous studies used structural homology searches to uncover hidden homology of viral proteins with receptor ligands ^7^, and other instances of mimicry have been proposed in homology searches ^8^. Such screens, however, are hindered by the relative dearth of structural data for viral proteins. Of the over 5,000,000 viral proteins in the UniProt database, fewer than 4,000 have associated experimental structural data (UniProt, January 2023). Recent advances in structural modeling such as implemented in AlphaFold2 ^9^ have the potential to bridge this gap, and the utility of this breakthrough in identifying cryptic homology is already being appreciated. For example, *ab initio* modeling has been used to broadly identify pathogen mimics of host proteins ^10^, to identify evolutionary connections among pathogen effectors ^11^, to expand an understanding of immunoglobulin gene family evolution ^12^, and to provide insight into the distant cellular origins of structural proteins found in viruses ^13^. In this study, we use AlphaFold to enable searches for hidden homology in viral proteomes and test its ability to inform the mechanistic study of host-pathogen interfaces.

We screened the vaccinia virus proteome for cryptic homology with metazoan proteins. From these comparisons, we identified and characterized two vaccinia proteins that influence host immunity. One, vaccinia virus *A47L*, encodes a protein predicted to have extensive structural overlap with gasdermin proteins, which we confirmed by solving a crystal structure of A47. A second gene, *C1L*, encodes a unique pyrin-Bcl-2 domain fusion. In functional studies, we provide mechanistic insight into the ability of A47 and C1 to modulate inflammasome-based immune responses.

Inflammasomes, and their ability to promote pyroptotic cell death, are important components of the host response to many viruses ^14^, including poxviruses ^15^. Previous studies have shown that poxviruses encode multiple inflammasome inhibitors ^16^, and by identifying two additional poxvirus effectors that modify inflammasome-mediated host defenses, our study broadens our understanding of how crucial the poxvirus-inflammasome interface is during infection. Importantly, these findings highlight the emerging power of structural modeling to detect cryptic homology among proteins and reveal new mechanisms for virus manipulation of host biology.

## Results

### Model-based homology screens uncover relationships between vaccinia and metazoan proteins

We performed a structural homology screen for vaccinia virus, a model virus and vaccine strain related to Mpox virus and the eradicated variola virus, the causative agent of smallpox. Vaccinia virus encodes over 200 proteins, many of which originated as host genes that were captured by horizontal transfer. While many links between vaccinia genes and host genes are evident from sequence identity, the origins of much of its proteome remain unclear ^17^. We modeled the vaccinia virus proteome using AlphaFold2 and performed homology searches using two orthogonal search tools. First, we used TM-align ^18^, a sensitive homology search tool that has been widely adopted for structural homology screens. We also used FATCAT ^19^, an alignment tool that permits rotations at flexible regions in structural alignments to enable searches with relaxed stringency where models may have ambiguities. We modeled all 257 predicted ORFs found in vaccinia virus strain Copenhagen and performed homology searches using AlphaFold-modeled proteomes of representative metazoans (Figure 1A).

**Figure 1:**
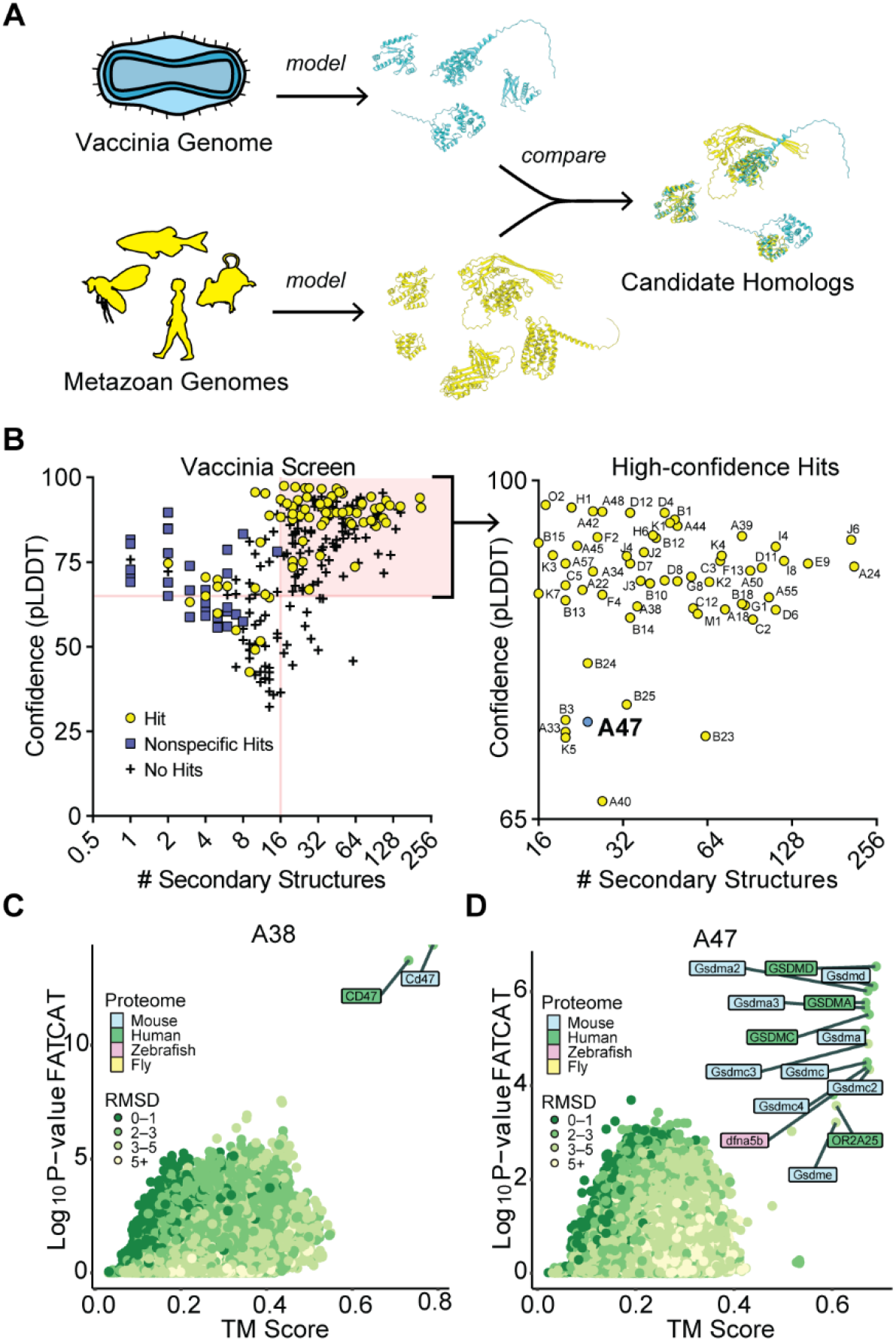
Structure-based homology screens reveal candidate homologs of vaccinia proteins. A) Homology screening pipeline. See Methods for details. B) Screen results. Right: high confidence hits (those with model confidence (pLDDT) > 65 and number of secondary structures > 15 for which FATCAT and TM-align converged) are indicated. C) FATCAT and TM-align results for A38, the poxvirus CD47 homolog. D) FATCAT and TM-align results for A47.

Focusing on proteins with homology revealed by both TM-align (TM score greater than 0.6) ^20^ and FATCAT (P-value cutoff of 0.01), we observed that small proteins (~100 amino acids or fewer) had a high proportion of nonspecific hits in our homology searches. In such instances, TM-align and FATCAT searches either produced divergent results or identified large groups of unrelated proteins as hits (Figure S1A). Seemingly nonhomologous proteins often have similar structural motifs^21^, and while the evolutionary relationships among such features are worth considering, it is difficult to distinguish between convergence or descent from a shared sequence. Many proteins with ambiguous homology were short, comprising few structural motifs. For example, A orf I, a predicted 73-residue vaccinia protein, has 6 predicted structural motifs. Homology searches identified many putative hits (Figure S1B), the strongest being p53-inducible protein 11. However, a comparison of the modeled structures of these proteins (Figure S1C) revealed that A orf I only matches a small portion of the top hit. In addition, phylogenetic analyses (Figure S1D) failed to resolve which of the widely divergent “hits” is biologically relevant. While such results may be informative in searches for distant origins of poxvirus proteins, this ambiguity led us to exclude these nonspecific hits from our screens.

By removing proteins with fewer than 16 secondary structure motifs we eliminated most nonspecific results (Figure S1E). Our search yielded 60 proteins for which structural homologs could be confidently predicted (Figure 1B, Table S1). We identified known homologs of host proteins, such as A38, the vaccinia homolog of CD47 ^22^ (Figure 1C). Other conserved folds, such as that of the helicase A18, the resolvase A22, the methyltransferase D12, and other components of the poxvirus transcriptional machinery, were identified, in line with previous studies ^17^ (Table S1).

Alongside well-characterized homologs of host proteins, our screen identified vaccinia A47 protein as a structural homolog of the regulatory domain of gasdermins, the protein executioners of pyroptotic cell death (Figure 1D). Pyroptosis is an inflammatory form of programmed cell death that serves to limit pathogen spread and is defined by the activity of the gasdermin protein family ^23^. Gasdermins are two-lobed molecules comprising a lipophilic N-terminal domain (NTD) and an autoinhibitory C-terminal domain (CTD). During canonical inflammasome signaling, diverse microbe-associated stimuli activate supramolecular assembly of inflammasomes and the activation of inflammatory caspases. Active inflammatory caspases subsequently cleave the interdomain linker of gasdermins, releasing the lipophilic NTD to oligomerize in membranes and form pores that induce lytic cell death and enable release of the cytokines IL-18 and IL-1β. Several viruses are known to antagonize inflammasome activity at multiple steps ^24^, including by targeting GSDMD for inactivating cleavage ^25^ and by blocking its cleavage by caspases ^26^. However, poxvirus-encoded inhibitors of gasdermin proteins have not been experimentally studied.

Since previous studies have demonstrated that the gasdermin D CTD can inhibit gasdermin function *in trans* ^27^, we hypothesized that a viral copy of a gasdermin CTD could inhibit pyroptotic cell death by mimicking its normal regulatory function.

### A crystal structure of a poxvirus A47 paralog confirms homology with gasdermins

To definitively determine whether A47 is a virus-encoded gasdermin, we determined a crystal structural of the eptesipoxvirus A47 ortholog at 1.46 Å resolution (Figure 2A). Eptesipox gasdermin has marked structural homology with diverse metazoan gasdermin CTDs and the predicted AlphaFold model of vaccinia A47 (Figures S2A–C). Eptesipox gasdermin and the gasdermin CTD share the distinctive α-helical bundle fold, comprising eight α-helices and three antiparallel β-strands (Figure 2A) ^28,29^. Besides the homologous regions, eptesipox gasdermin has an additional α-helix off its N-termini that is not present in metazoan gasdermins (Figure 2A, Figure S2D). Structural superposition demonstrates a potential steric clash of this helix with the gasdermin NTD, suggesting that at least some homologs of A47 may not directly inhibit pore formation by directly binding and blocking gasdermin NTDs (Figure 2A). These results confirm the presence of distinct viral gasdermin paralogs that are widely conserved among diverse poxviruses.

**Figure 2:**
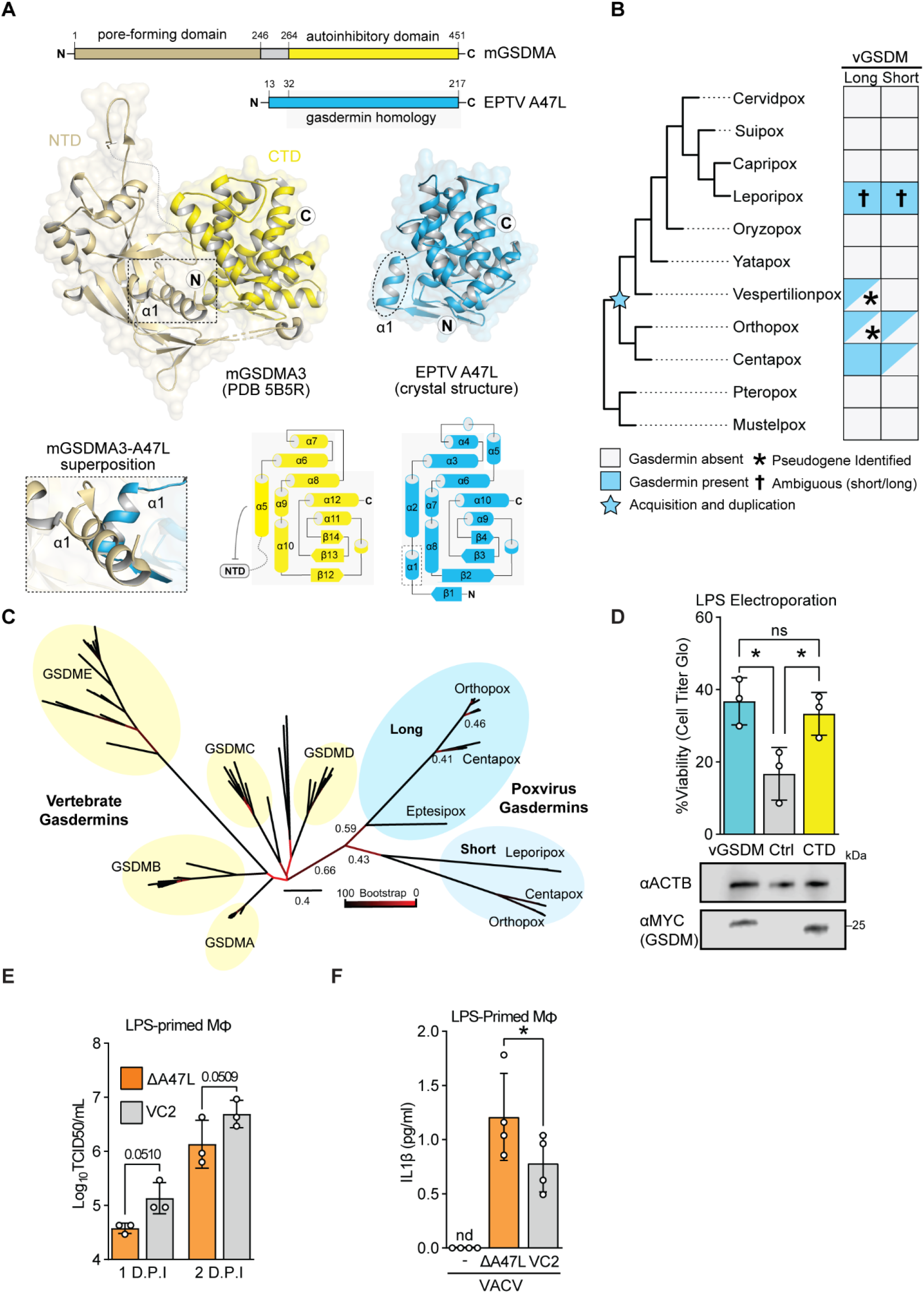
A poxvirus homolog of gasdermins is an inhibitor of pyroptosis. A) A crystal structure of Eptesipox virus (EPTV) gasdermin reveals homology to mammalian gasdermins. Top, domain organization of mouse GSDMA3 (mGSDMA3) and EPTV gasdermin indicating the pore-forming N-terminal domain (NTD) and the autoinhibitory C-terminal domain (CTD). Middle, crystal structures of full-length mGSDMA3 (PDB ID 5B5R) and EPTV gasdermin (this study). The N- and C-termini are indicated with the circled letters N and C, and the dashed box and oval indicate the first α-helix of each structure. Bottom left, superposition of mGSDMA3 and EPTV gasdermin predicts a steric clash between the α1 helices of each protein. Bottom right, topology diagrams of the mGSDMA3 CTD and EPTV gasdermin indicate a conserved region of gasdermin homology. B) Overview of probable gasdermin gain/loss events, based on phylogenetic and synteny analyses in B and C, respectively. Gasdermins were identified by a combination of sequence and structure-based approaches; see Methods for details. C) Maximum likelihood tree of select vertebrate gasdermin regulatory domains and poxvirus gasdermins. Clusters of vertebrate gasdermins are labeled based on human gasdermins. Bootstrap values from 100 replicates are indicated for select branches. Scale: AA substitutions per site. D) Top: HeLa cells expressing GSDMD-CTD, vaccina GSDM, or a vector control were electroporated with LPS to activate the non-canonical inflammasome. Viability as assessed by ATP levels relative to mock are indicated. n = 3 biological replicates. One-way ANOVA with Tukey’s multiple comparison test. Bottom: Alongside one replicate, levels of transfected gasdermin proteins in mock-electroporated wells were assessed by western blotting. E) Vaccinia replication in LPS-primed murine macrophages was measured by TCID50. n = 3 biological replicates. 2-way ANOVA with Fisher’s LSD test. ANOVA statistic for viral strain across timepoints: *p* = 0.0118. F) LPS-primed murine macrophages were infected with wild-type (VC2) or A47-deficient vaccinia virus and IL-1β was quantified by ELISA one day post-infection. n = 3 biological replicates. Paired t-test.

### A diverse family of poxvirus gasdermin homologs

Given predicted structural overlap between A47 and gasdermins, we performed complementary searches for viral gasdermins in poxvirus proteomes using psi-BLAST ^30^, a pox-specific local BLAST database (see Methods), and HMMER ^31^. These sensitive queries revealed poxvirus gasdermins in ortho- and centapox viruses consistent with a recent survey ^17^ and additional hits in vespertilion-, and lepori-poxviruses. These findings are consistent with the acquisition of the poxvirus gasdermin gene from an ancient host followed by extensive sequence divergence that obscured a host origin (Figure 2B).

Intriguingly, some poxvirus genomes in the orthopox and centapox clades have two copies of viral gasdermin, one “long” form, shared with vaccinia virus, and another somewhat-truncated “short” form. These gasdermins cluster phylogenetically (Figure 2C) and are proximally located in poxvirus genomes (Figure S2E), implying that they are descended from a single acquisition and early duplication. Whether present-day leporipox gasdermins are descended from the “long” or “short” gasdermins is ambiguous, as they are syntenic with the “long” gasdermin in ortho- and centa-poxviruses (Figure S2E), but cluster with the “short” gasdermin phylogenetically (Figure 2C). It is possible that leporipox gasdermins convergently evolved to include truncations like those in the “short” gasdermins found in other poxviruses, or that additional recombination events led to a rearrangement of the genes present at this locus. The persistence of both paralogs in several genomes in the orthopox and centapox clades further supports the idea that viral gasdermins are beneficial to poxviruses and may indicate that they serve distinct roles in poxvirus biology.

### Vaccinia A47 interferes with pyroptosis

To test whether A47 is an inhibitor of gasdermin-mediated pyroptosis, we used an established HeLa cell-based assay ^32^ to assess gasdermin-dependent pyroptosis. In this assay, electroporated LPS is sensed by the noncanonical inflammasome, promoting gasdermin D cleavage via activated caspase 4/5, resulting in pyroptosis. Importantly, expression of the regulatory domain of gasdermin D prevents LPS-induced cell death in this assay ^27^. We found that expression of vaccinia virus gasdermin similarly rescues cells from LPS-induced cell death (Figure 2D), consistent with a model in which poxviruses repurposed this homolog of a gasdermin regulatory domain as an inhibitor of pyroptosis.

We next assessed the role of poxvirus gasdermin during vaccinia infection by generating A47L-deficient virus (Figure S2F). Consistent with previous reports ^33^, A47L-deficient vaccinia virus replicated to wild-type levels in permissive cells such as BHKs (Figure S3G). However, we found that A47L-deficient virus was attenuated in a murine macrophage line (Figure 2E, ANOVA p = 0.0118) and promoted higher levels of IL-1β release during infection (Figure 2F). Together these findings provide complementary evidence that the ancient acquisition of a gasdermin gene by poxviruses led to evolution of viral gasdermins, such as A47L, that suppress pyroptosis during infection. A47 adds to a collection of poxvirus proteins that interfere with inflammasome activity ^24^. Among them are a soluble IL-1β binding protein ^34^, an NLRP1 inhibitor ^35^, and serine protease inhibitors implicated in regulating caspase activity ^36^. The diversity of means by which poxviruses target this immune pathway highlights its importance in host responses to poxvirus infections.

### Domain-specific homology searches reveal a cryptic pyrin domain-Bcl-2 fusion protein

While analyzing host gene capture events (Figure 1B, Table S1), an interesting class of genes encoding potential domain fusions became apparent. These were instances where TM-align and FATCAT results failed to converge and FATCAT, which allows for flexible alignments, identified homology with multiple distinct protein families. A notable example is vaccinia C1, which was previously described as a member of the poxvirus A46/Bcl-2-domain protein family ^37^. FATCAT identified limited overlap with Bcl-2 family members such as MCL1 and at the same time indicated robust homology with pyrin-domain containing proteins (Figure 3A). Consistent with it being the product of a fusion event, the model of C1 contains two globular protein domains attached by a flexible linker, and domain-specific searches (Figures 3B-C) demonstrated that C1 is comprised of an N-terminal pyrin and a C-terminal Bcl-2 domain. We used DALI ^38^ to search the protein data bank (PDB) for solved structures similar to these two domains and found that the pyrin domain exhibits similarity to structures of known pyrin-domain containing proteins while the Bcl-2 domain is most similar to Bcl-2 domain-containing proteins (Figure S3A). A structural comparison of the AlphaFold model of C1 protein to hits for each domain revealed marked homology, consistent with results from our model-based homology screens (Figure 3D).

**Figure 3:**
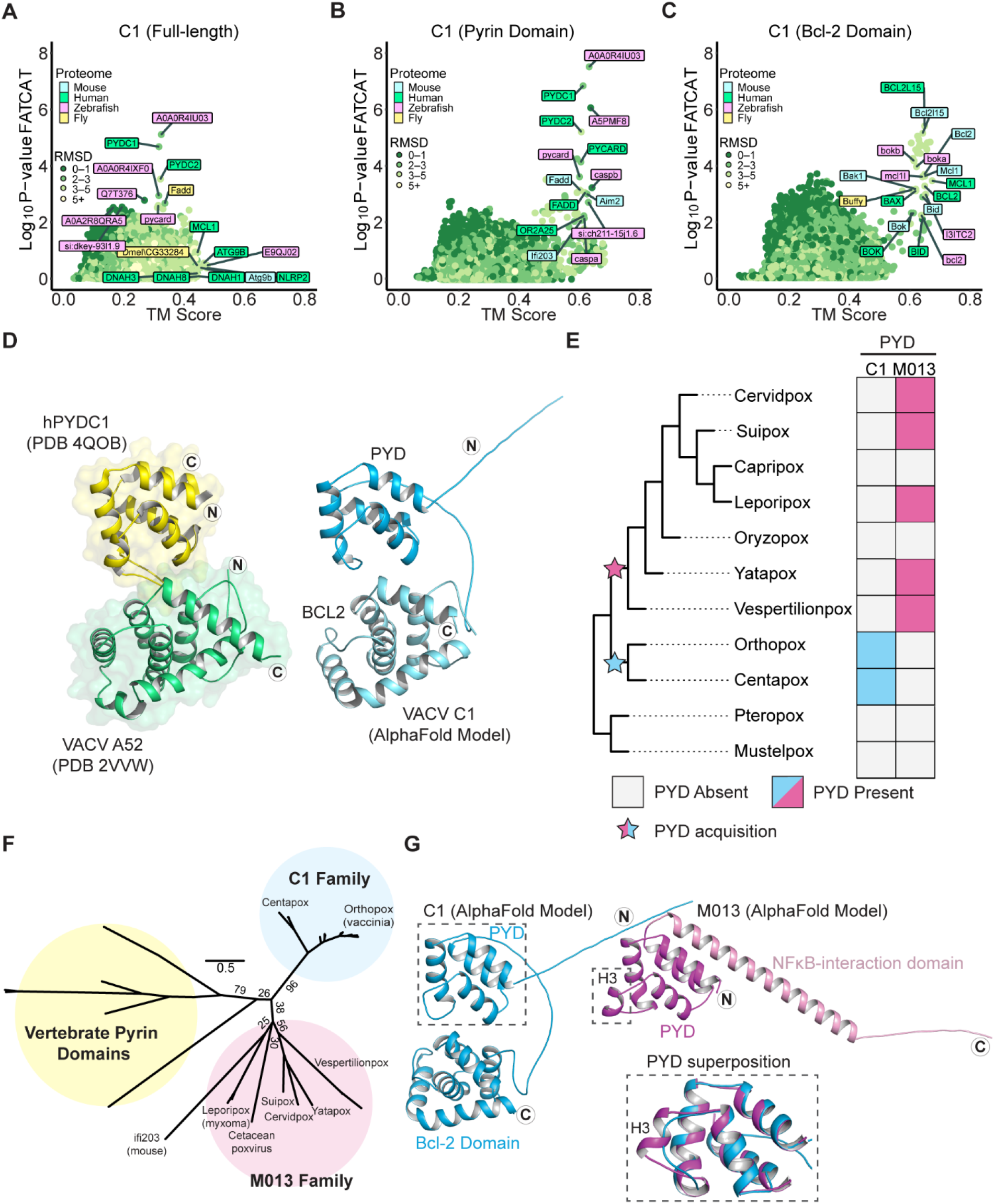
Vaccinia C1L encodes a unique pyrin-Bcl-2 fusion protein. A) FATCAT and TM-align results for full-length C1 protein. Top results for each individual search tool are labeled. B) FATCAT and TM-align results for the N-terminal (1-107) residues of C1 protein. Only a subset of hits is labeled. C) FATCAT and TM-align results for the C-terminal (108-224) residues of C1 protein. Only a subset of hits is labeled. D) C1 protein is a PYRIN-Bcl-2 fusion. Left: crystal structures of human PYDC1 (PDB 4QOB) and vaccinia virus A52 (PDB 2VVW). Right: AlphaFold model of vaccinia C1 protein. The N- and C-termini are indicated with the circled letters N and C. E) Reconstructed gain/loss tree highlighting C1 and M013 pyrin domains-containing proteins F) Maximum likelihood phylogenetic tree of vertebrate pyrin domains from Figure 4B alongside the M013 and C1 families. Bootstrap values from 1000 replicates are indicated for select branches. Scale: AA substitutions per site. G) AlphaFold models of vaccinia virus C1 protein and myxoma virus M013 protein. The N- and C-termini are indicated with circled letters N and C. Inset: superposition of the C1 and M013 pyrin domains showing the predicted absence of helix 3 in C1 protein.

Members of the poxvirus Bcl-2 domain family have been variously implicated as immunomodulatory proteins that act through multiple mechanisms ^37^, though no immunomodulatory function has been ascribed to C1 protein. Indeed, the only prior mechanistic study of C1 protein showed that unlike many other poxvirus Bcl-2 family proteins, C1 is not anti-apoptotic and does not bind or inhibit the pore-forming Bcl-2 protein BAX ^39^. Pyrin domains are protein-protein interaction domains that mediate many of the steps involved in inflammasome activation and pyroptosis. The pyrin domain-containing protein family is comprised of the pattern recognition receptors NLRPs (NOD-like receptor proteins), the DNA sensors AIM2 and IFI16, the adapter protein ASC, and pyrin-only proteins (POPs) which serve as regulators of inflammasome signaling ^40^. Notably, no proteins containing both a pyrin and a Bcl-2 domain have been described to date, highlighting the unusual composition of this poxvirus protein.

### C1 protein is a member of a unique family of viral pyrin domains

We sought to determine whether C1 protein is the result of *de novo* gene acquisition events, or if it is a divergent ortholog of a known poxvirus pyrin protein, myxoma virus M013, which might have acquired a Bcl-2 domain. To date, M013 is the only pyrin domain containing protein described in poxviruses. M013 regulates both NF-κB signaling and inflammasome activity through distinct mechanisms ^41 42^ and is a major contributor to myxoma virus pathogenesis ^43^. We searched poxvirus genomes for pyrin domains using both M013 and C1 proteins as seeds for psi-BLAST ^30^ and HMMER ^31^ searches, and only identified orthologs of C1 in clades of orthopox and centapox viruses (Figure 3E). As previously described, orthologs of M013 were found in clade II poxviruses and were absent in orthopoxviruses ^44^ (Figure 3E). We also identified distantly related homologs of M013 in the bat-infecting vespertilionpoxviruses eptesipox virus and hypsugopox virus, as well as in a cetacean poxvirus (Table S2).

Phylogenetic analysis of poxvirus pyrin domain-containing proteins and select vertebrate pyrin domains supports a model in which C1 and M013 resulted from independent gene acquisition events (Figure 3F). Furthermore, synteny analysis revealed that the genes encoding C1 and M013 are located in distinct regions of poxvirus genomes, also consistent with independent acquisitions of genes encoding these pyrin domains (Figure S3B). Finally, superimposing models of C1 and M013L revealed structural differences between the two proteins (Figure 3G), which are also apparent from comparisons of amino-acid sequences (Figure S3C). These data suggest that poxviruses acquired pyrin domains independently on at least two occasions, highlighting the potential of viral proteins containing pyrin domains to impact poxvirus infections.

### Vaccinia C1 protein promotes ASC-dependent inflammasome activation

The unique composition of C1 raises questions about its function during infections. Previous studies demonstrated that myxoma M013 protein regulates host immunity by interfering with NF-κB-dependent signaling ^41^. We assessed whether C1 or its individual domains had similar functions to M013. In an assay measuring NF-κB signaling following TNFα treatment, C1 protein did not phenocopy M013, suggesting that these proteins serve distinct functions (Figure 4A), a result consistent with our phylogenetic analyses suggesting that C1 protein is not an ortholog of M013. Curiously, the pyrin domain of C1 alone enhanced cellular responses to TNFα, which potentially implicates the Bcl-2 domain for a role in preventing aberrant activation of cellular inflammatory pathways by C1 during infection.

**Figure 4:**
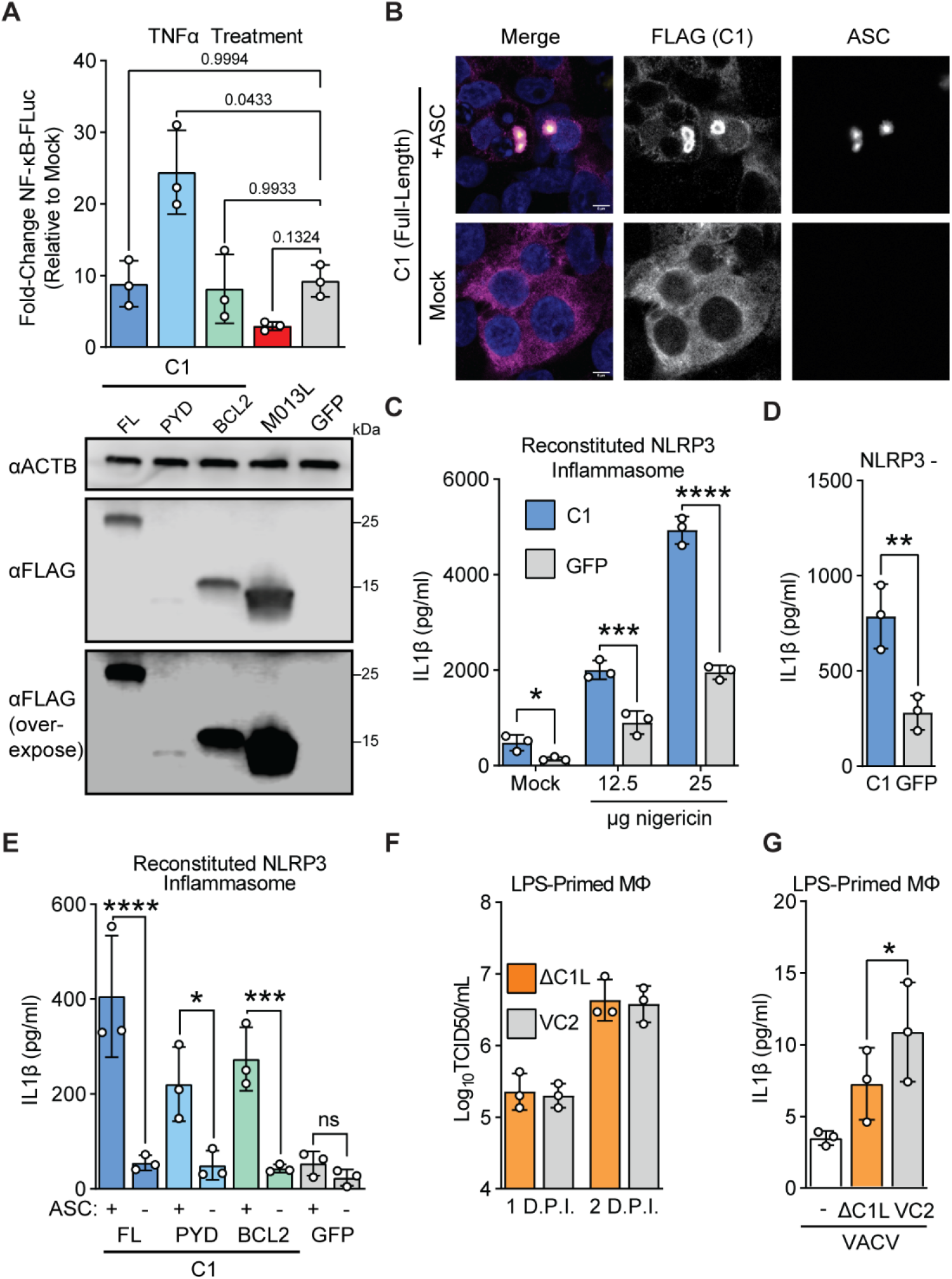
C1 protein promotes ASC-dependent inflammasome activation. A) Top: 293T cells were co-transfected with the indicated plasmids, a plasmid encoding firefly luciferase driven by an NF-κB promoter, and a constitutive renilla luciferase plasmid as a transfection control. 24 hours post-transfection, cells were treated with 1ng/mL TNFα, and promoter activity was assessed by luciferase assay 6-hours post transfection. n = 3 biological replicates. RM one-way ANOVA with Dunnett’s correction. Bottom: In parallel, cells were transfected with equal amounts of the indicated plasmids and protein expression was determined by western blotting. B) 293T cells were co-transfected with a plasmid encoding FLAG-C1 and either ASC-GFP ^1^ or an empty vector. 24 hours post-transfection, cells were fixed and stained for FLAG (C1) and subsequently imaged. Representative image from n=3 independent experiments. Scalebar: 5μm. C) 293T cells were co-transfected with plasmids encoding the NLRP3 inflammasome components NLRP3, ASC, NEK7, Caspase 1, and prolL-1β, in addition to either C1 or a GFP control. Following nigericin stimulation, IL-1β concentrations in supernatants were quantified by ELISA. n = 3 biological replicates. Two-way ANOVA with Šídák’s multiple comparisons test. D) 293T cells were co-transfected as in (d) but with NLRP3 replaced by an empty vector. Cells were not stimulated with nigericin. IL-1β concentrations in supernatants were quantified by ELISA. n = 3 biological replicates. Paired t-test. E) 293T cells were co-transfected as in (d) but with ASC replaced by an empty vector where indicated. Cells were not stimulated with nigericin. IL-1β concentrations in supernatants were quantified by ELISA. n = 3 biological replicates. Two-way ANOVA with Šídák’s multiple comparisons test. F) Vaccinia replication (inoculum MOI of 1) in LPS-primed murine macrophages was measured by TCID50. G) LPS-primed murine macrophages were infected with wild-type (VC2) or C1L-deficient vaccinia virus at an MOI of 10 and IL-1β was quantified by ELISA four hours post-infection. n = 3 biological replicates. Paired t-test.

To further characterize C1 function, we considered whether it interacts with ASC, a critical pyrin-domain containing protein, also known as pycard (<u>py</u>rin and <u>CARD</u>-domain containing protein). ASC is an adapter protein that links innate immune sensors and caspases for multiple inflammasomes including AIM2 and NLRP3, two inflammasomes implicated in counteracting vaccinia virus infections^16^. We found that ectopically-expressed C1 protein associates with ASC by forming a concentrated shell peripheral to ASC specks (Figures 4B and S4B-C). Surprisingly, the individual pyrin and Bcl-2 domains of C1 can separately associate with ASC (Figures S4B-C), suggesting multiple means of interfacing with ASC.

Because C1 appears to interact extensively with ASC, we tested whether C1 protein can modulate inflammasome signaling using a reconstituted NLRP3 inflammasome system ^45^. Unexpectedly, expression of C1 protein enhanced IL-1β release in the presence or absence of the NLRP3 agonist nigericin, indicating that C1 protein directly promotes ASC-dependent caspase activation (Figure 4C). This increase in inflammasome activity was NLRP3-independent (Figure 4D) and relied upon ASC (Figure 4E). We found that both domains of C1 protein could promote ASC-dependent IL-1β processing (Figure 4E), consistent with our observations that individual domains of C1 protein can associate with ASC (Figures S4B-C).

The apparent promotion of inflammasome activation by C1 protein seems at odds with the diverse range of inflammasome inhibitors present in poxviruses, so we next asked whether C1 protein impacted vaccinia virus replication or inflammasome activity during infection. We generated C1-deficient vaccinia virus (Figure S4D) and found that it replicated at levels comparable to wild-type virus in BHK-21J cells (Figure S4E) and in macrophages (Figures 4F and S4F). However, we found decreased IL-1β secretion early during infections with C1-deficient vaccinia (Figure 4G), consistent with a pro-inflammatory role also observed from cell-based reporter assays. Together these findings suggest that C1 protein enhances inflammasome signaling, adding a twist to poxvirus regulation of inflammasomes during infections as compared to a collection of inhibitory mechanisms.

## Discussion

In the present study, we used *ab initio* structural modeling to search for cryptic homology between host and virus proteins. Our screen revealed two vaccinia virus proteins that modulate inflammasome activity and pyroptosis, underscoring the importance of this immune pathway in viral infections. One, a homolog of gasdermins (A47), is an inhibitor of pyroptosis. Another, a unique pyrin-Bcl-2 fusion protein (C1), is counterintuitively pro-inflammatory, driving ASC-dependent inflammasome activity.

Homology with gasdermin proteins guided our mechanistic studies of A47. A previous study ^33^ established that A47 has no impact on viral replication in permissive cells such as BHK-21J cells, which is consistent with our findings using the same cell line (Figure S4E). The involvement of gasdermins in pyroptosis led us to explore the role of A47 in macrophages, where we found that A47 contributes to viral replication (Figure 2E) and suppression of cytokine release (Figure 2F). It is notable that the A47-dependent phenotypes are subtle, perhaps due to redundant mechanisms by which vaccinia virus inhibits inflammasome activity and pyroptosis ^24^. While combinatorial knockouts of A47 and other anti-pyroptotic proteins may be informative, we note the potential for homology screens to reveal overlapping functions and inform mechanistic study.

Our data are consistent with at least two models of A47-mediated inhibition of pyroptosis. In one, A47 could occupy the binding pocket of caspases, acting as a competitive inhibitor or promoting their deactivation and degradation ^46,47^, and the crystal structure of eptesipox gasdermin suggests this may be the case for this protein (Figure 2A). In a second model, a viral gasdermin could serve as a “molecular sponge” that sequesters pore-forming domains of cellular gasdermins post-cleavage, suppressing pore formation. Indeed, the presence of two viral gasdermin paralogs in some poxvirus genomes (Figures 2B-C and S2E) suggests that they may have diverged to inhibit pyroptosis through multiple mechanisms, perhaps with individual paralogs specializing for unique inhibitory roles. It is also possible that orthologs within the individual clades of viral gasdermins adopted distinct functions as poxviruses adapted to infect different hosts. Future mechanistic studies will shed light on how different viral gasdermins work to benefit viral infection.

Several gene fusions in poxvirus genomes have been described, though in many cases the contribution of individual domains to the function of hybrid proteins have not been explored ^17^. Our study identified a cryptic pyrin domain in the poxvirus Bcl-2 family protein C1 (Figure 3). To our knowledge, this is the only known example of a pyrin-Bcl-2 fusion protein, and the combination of these domains by poxviruses presents a unique opportunity to explore new facets of the biology of both domains. Curiously, the pyrin and Bcl-2 domains of C1 generally are redundant in function, with a notable difference being the ability of the pyrin domain to enhance NF-κB signaling following stimulation with TNFα (Figure 4A). We suspect that this “masking” is not the only function unique to the individual domains of C1 protein and anticipate that future genetic and biochemical studies of this protein will expand our understanding of the complexities of inflammasome activity in poxvirus infection.

Unlike an inhibitor of cell death such as A47, a pro-inflammatory protein is difficult to reconcile with a beneficial role for a virus. The persistence of C1 in diverse poxvirus genomes (Figure 3), however, argues that it somehow promotes viral replication. One possibility is that pro-inflammatory signals promoted by C1 protein allow vaccinia to recruit and infect cells such as macrophages, as infected macrophages may promote vaccinia dissemination ^48^. However, previous experimental infections of mice showed that macrophages are a critical component of the immune response to vaccinia infection ^49^. Future *in vivo* pathogenesis studies of C1-deficient vaccinia would help better define the role of this protein, as well as inflammation and macrophages, during vaccinia infection.

Identification of a novel domain organization in C1 protein raises the prospect that diverse gene fusion events are more common in viruses than currently appreciated. Freed from constraints in host genomes, individual domains and combinations thereof may diverge to acquire new functions, with viral genomes serving as a testbed of evolutionary innovation. Our study joins a growing body of work demonstrating the transformative power of structural modeling in identifying evolutionary connections ^11,50,51^ and informing mechanistic study ^12^. The ability to detect distant homology is especially important for hybrid proteins such as C1, where global homology searches may be inconclusive (Figures 3A-C). Tools such as Foldseek ^52^ and the AlphaFold-based domain identification tool DPAM ^53^ are beginning to address these limitations and will empower future studies of structural homology, with continuing computational advances unlocking new biology. Pipelines like ours have great potential to reveal cryptic captured genes in classes of viruses beyond poxviruses, such as the emerging class of giant viruses that can encode thousands of genes, deepening our understanding of how viruses manipulate their hosts during infection.

## Acknowledgements

We acknowledge the University of Utah CHPC for their assistance in implementing AlphaFold and FATCAT. We thank Jamie Gagnon and Katie Deets for critical manuscript feedback. We thank members of the Elde lab for helpful discussions. We particularly thank Katie Deets for discussions on inflammasome biology, and Zoë Hilbert for helpful discussions regarding evolutionary analyses. We additionally thank Patrick Mitchell for helpful discussions. We thank James Eaglesham and Kacie McCarty for insights and assistance on A47 structure determination. This work was supported by grants to N.C.E (NIH R35GM134936) and by the Pew Biomedical Scholars Program (P.J.K.), the Burroughs Wellcome Fund PATH award (P.J.K.), and the Mathers Foundation (P.J.K.). This work was additionally supported by two Life Science Research Foundation postdoctoral fellowships of the Open Philanthropy Project (I.N.B. and A.G.J.). Any opinions, findings, and conclusions or recommendations expressed in this material are those of the authors(s) and do not necessarily reflect the views of funding agencies.

## Author Contributions

Conceptualization, I.N.B and N.C.E.; Investigation, I.N.B and A.G.J.; Resources, I.N.B, A.G.J., M.Q.; Writing, original draft: I.N.B.; Writing, review and Editing: I.N.B., N.C.E., A.G.J., and P.J.K.; Visualization, I.N.B. and A.G.J; Funding Acquisition, I.N.B., A.G.J., P.J.K., and N.C.E.

## Methods

### Cell culture and viruses

BHK-21J (ATCC) and 293T (a kind gift from Wes Sundquist, University of Utah) and HeLa (a kind gift from Adam Geballe, Fred Hutchinson Cancer Research Center) cells were cultured in DMEM supplemented with 10% FBS. Immortalized mouse bone marrow derived macrophages (a kind gift from Sunny Shin, University of Pennsylvania) were cultured in RPMI supplemented with 10% FBS. All cells were cultured at 37°C in 5% CO_2_. Vaccinia strain Copenhagen (VC2) ^54^ was propagated in BHK cells by passaging.

### Viral infections

BHK or iB6 cells were seeded at 200,000 cells per well in a 12-well plate one day prior to infection. Virus was added in a minimal volume of media (RPMI or DMEM + 1%FBS, 400ul), after which 600ul of the appropriate complete (10% FBS) media was added per well. Virus was harvested by freezing plates at −80, thawing, and sonicating the entire contents of the well in a Qsonica Q500 cup-horn sonicator (50% amplitude, 2s on/2s off, 60s processing time). Lysates were cleared by centrifugation at >14,000xg for 3-5 minutes.

For viral stocks, virus was subsequently aliquot and stored at −80 prior to determination of titer. Titers of experimental samples were immediately determined by TCID50.

### TCID50 assays

Cells seeded at 20-30,000 cells per well on 96w plates. Viral stocks were serially-diluted in DMEM supplemented with 1% FBS, and cells were infected with 50ul of inoculum for 2 hours, after which 100ul of DMEM supplemented with 10% FBS was added per well. Cells were scored for cytopathic effect (CPE) four to five days post-infection. For all experimental assays, samples were coded and randomized prior to TCID50 infections to ensure that the investigator was unbiased in assessing for CPE. TCID50 values were calculated by the method of Spearman and Kärber.

### Recombinant vaccinia production

To generate recombinant vaccinia, BHK-21J cells were seeded on 6-well plates at 400,000. The next day, cells were infected at MOI of 2-5 in a 200ul of DMEM supplemented with 1% FBS. One hour post-infection, cells were transfected with 2ug of a homology donor plasmid containing a monomeric ultrastable GFP (muGFP) ^55^ marker using Lipofectamine 3000 (3.5ul of lipofectamine per microgram of plasmid, 2ul of p3000 reagent per microgram). Media was changed four to six hours post-transfection. The next day, virus was harvested as indicated above.

Recombinant virus was subsequently purified by 2-3 rounds of limiting dilution assays on 96 well plates, followed by two rounds of plaque purification on 6w plates. Purity of recombinant virus was verified by PCR on viral stocks, as described below. At each stage, recombinant virus was identified by GFP expression.

ΔA47L virus was generated by fully deleting the A47L ORF, replacing it with muGFP. ΔC1L virus was generated by introducing an in-frame stop codon and partial deletion of the C1L ORF to avoid disrupting N1L, which partially overlaps with C1L.

### Isolation of viral genomic DNA

One volume of viral stock (typically 50ul) was mixed with one volume of digestion buffer (0.9% NP50, 0.9% Tween-20, 20mM Tris-HCL pH 8.3, 3mM MgCl2, 100mM KCL). The mixture was supplemented with 1/20 volume of Proteinase K at 20mg/mL, and the reaction was incubated at 40C for 45 minutes. The completed reaction was transferred to a fresh tube and heat-killed by incubation at 95C for 10 minutes. This crude genomic DNA prep was used as input (1/10 reaction volume) for PCR analysis of viral DNA.

### PCR genotyping of viral stocks

For all recombinant viruses, four PCRs were performed. Primers flanking the 5’ and 3’ regions of the insertion region were paired with primers specific to either the GFP insertion or the native sequence. Recombinant virus was only considered to be pure if both recombinant PCRs were positive and both native PCRs were negative. Both water and wild-type VC2 DNA was included as controls in all assays.

### LPS electroporation assay

HeLa cells were plated one day prior to transfection at 250,000 cells per well on a 6w plate. The next day, cells were transfected with 2ug of the indicated constructs using Lipofectamine 3000 (2ul Lipofectamine 3000 per microgram of DNA and 2ul of p3000 reagent per microgram of DNA). Media was changed to fresh DMEM supplemented with 10% FBS six hours post-transfection. Two days after transfection, cells were electroporated with one microgram of LPS per million cells (or mock) using a Neon transfection system (1300V, 30ms pulse width, 1 pulse) (ThermoFisher). Cells were returned to pre-warmed DMEM supplemented with 10% FBS in 96-well plates for cell titer glo viability assays.

### NF-κB reporter assay

One day prior to transfection, 293T cells were seeded at a density of 50,000 cells per well on opaque 96 well plates. Cells were transfected with 200ng of the indicated pEF construct, 50ng of pNF<u>κ</u>Bpro-FLuc (A kind gift of Neal Alto, UT Southwestern Medical Center), and 10ng pRL-TK (Promega) per well using Lipofectamine 3000 (0.7ul lipofectamine per microgram of plasmid DNA, 2ul of p3000 reagent per microgram of DNA). 24 hours post-transfection, media was changed to 75ul of DMEM supplemented with 10% FBS containing TNFα (R&D Systems 210-TA) in either a dilution series or at approximately two times the EC50 within this assay, 1ng/mL, or a vehicle control. Six hours after treatment, plates were sealed and stored at −80°C until assay. Promoter activity (firefly luciferase) was determined by Dual-Glo luciferase assay (Promega) following manufacturer specifications, using Renilla signal as a transfection control. Luciferase readings were performed using either a BioTek Synergy HT or a BioTek H1 plate reader.

### Reconstituted NLRP3 inflammasome assay

The murine NLRP3 inflammasome was reconstituted in 293T cells per an established protocol ^45^. Briefly, 200,000 293T cells were seeded in a 24 well plate 18-24 hours prior to transfection. Cells were co-transfected with pCMV-pro-IL1β-C-Flag (200ng), pcDNA3-N-Flag-NLRP3 (200ng), pcDNA3-N-Flag-ASC (20ng), pcDNA3-N-Flag-Caspase-1 (100ng), pcDNA3-N-HA-NEK7 (200ng), and the indicated pEF (300ng) construct. For some assays, an empty CMV plasmid was used in place of indicated inflammasome components.

24 hours post-transfection, media was changed to 250ul of DMEM + 10% FBS. For some assays, cells were stimulated with nigericin (12.5 or 25 μg per well) diluted in ETOH or a vehicle control one hour prior to harvest. Six hours post-media change, supernatant was collected, cleared by centrifugation, and stored at −80 prior to downstream assays.

### ELISA

For all ELISA assays, supernatants were clarified by centrifugation at >14,000xg for 3-5 minutes. Cleared supernatants were frozen at −80C prior to assay. Supernatants were either assayed without dilution (most viral infections) or at an appropriate dilution in NS dilution buffer (Abcam).

Murine IL-1β was assayed using SimpleStep ELISA (Abcam ab197742). Briefly, samples and antibodies (detection and capture) were added to a pre-bound plate. Plates were incubated at room temperature for one hour with shaking, after which the plate was washed three times and TMB solution was added. Following 15-20 minutes of development, stop solution was added to all wells and OD450 absorbance was read on a BioTek HT plate reader. Concentrations of IL-1β were determined by comparison with a standard curve. All standards and experimental samples were assayed in technical duplicates.

### Immunofluorescence

Cells were fixed with 4% PFA in PBS. Cells were washed with PBS, then permeabilized with 0.2% Triton-X 100. Cells were blocked with 5% goat serum in PBS for at least 30 minutes. Primary antibody was added in blocking solution and incubated for 1-2 hours. Cells were washed 3x with PBS, after which secondary antibody was added in 3% BSA and incubated for 30 minutes. Cells were washed 3x with PBS, and then mounted using Fluormount-G (Southern Biotech). Imaging was performed on a Zeiss LSM980. Images were processed in ImageJ. When made, linear adjustments were applied evenly across all samples within each experiment.

### CellTiter-Glo viability assays

For most viability assays, cells were seeded at 50,000 cells per well in 96 well plates. At the experimental endpoint, one volume of CellTiter-Glo (Promega) assay reagent was added to each well. Cells were incubated at room temperature for 10 minutes with shaking, after which luminescence was read on a BioTek Synergy HT plate reader.

### Western blotting

Unless otherwise noted, cells were lysed directly in 1x SDS loading buffer (10% glycerol, 5% BME, 62.5mM TRIS-HCl pH 6.8, 2% SDS, and BPB), boiled, and sonicated (Qsonica Q500). Samples were run on “Any kD” acrylamide gels (Bio-Rad) and transferred to PVDF membranes using a Trans-Blot SD (Bio-Rad) or a Trans-Blot Turbo (Bio-Rad). Blots were blocked in 5% dry milk/TBS-T for 30 minutes to an hour at RT or overnight at 4C. Primary antibodies were diluted in 5% dry milk/TBS-T and added for 1 to 2 hours at RT or overnight at 4C. Blots were washed four times in TBS-T before addition of HRP-conjugated secondary antibody in 5% milk for thirty minutes. Blots were washed four times in TBS-T prior to detection with either ProSignal Dura (Prometheus) or Clarity ECL (Bio-Rad) substrate and exposure to radiography film or imaging on a C-Digit imager (LiCOR).

### Cloning

#### Overexpression clones

DNA encoding the A47L, C1L, M013L, and GSDMD sequences were synthesized (IDT, GeneWiz) and cloned into expression constructs by digest-based cloning. C1L truncations were cloned by PCR and restriction digest. See annotated primer table (Table S3) for details.

The A47L homology donor was synthesized and cloned into pUC19 (Genscript). The C1L homology donor was generated by PCR using VC2 DNA as a template and sequential restriction digest-based cloning for the 3’ and 5’ homology arms. See annotated primer table (Table S3) for details.

### Phylogenetic analyses

Multiple sequence alignments were generated using MUSCLE as implemented in MEGA 11 ^56^. For gasdermins (Figure 2C), the evolutionary history was inferred by using the Maximum Likelihood method and JTT matrix-based model in MEGA 11. Initial tree(s) for the heuristic search were obtained automatically by applying Neighbor-Join and BioNJ algorithms to a matrix of pairwise distances estimated using the JTT model, and then selecting the topology with superior log likelihood value. For pyrin-domain containing proteins (Figure 3F), IQ-TREE ^57^ was used to infer phylogeny. The best-fitting tree used the LG+G4 substitution model and had a log-likelihood of −4277.840.

### Identification of viral homologs of vaccinia screen hits

NCBI BLAST ^30^ and HMMER ^31^ were used to identify homologs of vaccinia A47L and C1L. Alignments of all previously-identified A47 and C1 homologs were used for iterative HMMER searches until no new homologs were identified.

### Local poxvirus BLAST database searches

A local installation of BLAST+ (version 2.13.0) ^58^ was used to generate a BLAST database comprised of representative poxvirus genomes (see Table S2). All identified poxvirus gasdermins were as search queries for BLASTN and discontinuous megablast searches to identify additional poxvirus proteins.

### Homology search pipeline

We used existing tools to perform deep homology searches for the vaccinia virus proteome. First, AlphaFold2 (v2.1.2) ^9^ was used as implemented at the University of Utah Center for High Performance Computing (CHPC) to generate models for all vaccinia virus proteins. Models with the overall greatest confidence (pLDDT) were used for homology searches.

Model organism (human, drosophila, mouse, and zebrafish) AlphaFold v2 proteomes were downloaded from the EBI AlphaFold Database to generate a target database for homology searches. TM-align searches were performed on a local Linux system (see file S5), while FATCAT searches were performed on a high-performance cluster at the University of Utah CHPC.

FATCAT and TM-align results were processed and cleaned using BASH scripts (see file S5). STRIDE ^59^ was used to tabulate the number of secondary structures in protein models for filtering. Cleaned data were further processed and visualized using R ^60^ (see file S5). Additional R packages ggplot2 ^61^, ggrepel ^62^, and ggnewscale ^63^ were used to visualize data.

### Protein expression and purification

Poxvirus homologs of vaccinia *A47L* were codon-optimized for *E. coli* expression, synthesized as gene fragments (IDT), and cloned into a custom pET vector encoding an N-terminal 6×His-hSUMO2 solubility tag. Cloning was performed by Gibson assembly and N-terminal truncation variants were subcloned by PCR. All plasmids were verified by Sanger sequencing. Plasmids were transformed into BL21 CodonPlus(DE3)-RIL *E. coli*, grown as starter cultures in MDG media (0.5% glucose, 25 mM Na2HPO4, 25 mM KH2PO4, 50 mM NH4Cl, 5 mM Na2SO4, 2 mM MgSO4, 0.25% aspartic acid, 100 mg mL^-1^ ampicillin, 34 mg mL^-1^ chloramphenicol, and trace metals), and expressed in M9ZB media (0.5% glycerol, 1% Cas-amino Acids, 47.8 mM Na2HPO4, 22 mM KH2PO4, 18.7 mM NH4Cl, 85.6 mM NaCl, 2 mM MgSO4, 100 mg mL^-1^ ampicillin, 34 mg mL^-1^ chloramphenicol, and trace metals). M9ZB cultures were grown at 37°C with 230 RPM shaking to an OD600 of ~2.5. Protein expression was induced by cooling cultures on ice for 20 min and then supplementing cultures with addition of 0.5 mM IPTG before incubation overnight at 16°C with shaking at 230 RPM. For each A47L variant, 2× 1 L cultures were expressed, pelleted, flash frozen in liquid nitrogen, and stored at −80°C prior to purification.

All protein purification steps were performed at 4°C using buffers containing 20 mM HEPES-KOH (pH 7.5). Expression cell pellets were thawed and lysed by sonication in buffer containing 400 mM NaCl and 1 mM DTT, clarified by centrifugation and glass wool, bound to NiNTA agarose beads (QIAGEN), washed with buffer containing 1 M NaCl and 1 mM DTT, and eluted with buffer containing 400 mM NaCl, 300 mM imidazole (pH 7.5), and 1 mM DTT. The SUMO2 tag was cleaved by the addition of recombinant hSENP2 protease (D364–L589, M497A) to the NiNTA elution with overnight dialysis in buffer containing 125–250 mM KCl and 1 mM DTT. The resulting dialyzed protein was purified by size-exclusion chromatography using a 16/600 Superdex 75 column (Cytiva) pre-equilibrated with buffer containing 250 mM KCl and 1 mM TCEP. Sized proteins were concentrated to >50 mg mL^-1^ using 10 kDa molecular weight cut-off concentrators (Millipore), flash frozen on liquid nitrogen, and stored at −80°C. Selenomethionine(SeMet)-substituted proteins were prepared in M9 media and purified in buffers containing 1 mM TCEP in place of DTT.

### Protein crystallization and structure determination

Of the homologs and variants tested, Eptesipox virus (EPTV) gasdermin (Genbank accession ASK51347) was found to have optimal yield and solubility. Eptesipox gasdermin crystals were grown by the hanging-drop vapor diffusion method at 18°C using NeXtal crystallization screens and optimized with EasyXtal 15-well trays (NeXtal) and the Slice pH screen (Hampton Research). Proteins for crystallization were thawed from −80°C stocks on ice and diluted to concentrations of 20 mg mL^-1^ protein and 20 mM HEPES-KOH (pH 7.5), 60 mM KCl, and 1 mM TCEP. 15-well crystal trays were set with 2 μL drops containing diluted protein and reservoir at a 1:1 ratio over wells with 350 μL reservoir. Crystals were grown for at least 3 days, incubated in reservoir solution without additional cryoprotectant, and harvested by flash freezing in liquid nitrogen. Native EPTV gasdermin (amino acids 12-217) crystals grew in 100 mM ADA (pH 6.4) and 40% PEG-200. SeMet-substituted EPTV gasdermin (amino acids 18-217) crystals grew in 100 mM DL-malic acid (pH 5.3) and 40% PEG-200. All crystals were cryoprotected in reservoir solution. X-ray diffraction data were acquired using Northeastern Collaborative Access Team beamlines 24-ID-C and 24-ID-E (P30 GM124165), and used a Pilatus detector (S10RR029205), an Eiger detector (S10OD021527) and the Argonne National Laboratory Advanced Photon Source (DE-AC02-06CH11357).

Data were processed with XDS and AIMLESS using the SSRL autoxds script (A. Gonzalez) ^64^. All structures were phased with anomalous data from SeMet-substituted crystals using Phenix Autosol version 1.19 ^65 66^. Atomic models were built in Coot ^67^ and refined in Phenix using native diffraction data. Statistics were analyzed as described in Supplemental Table 4 ^68 69 70^. Structure data were deposited in the Protein Data Bank (PDB ID 8GBE). Structure figures were generated using PyMOL version 2.4.0 (Schrödinger, LLC).

## Supplemental Figures

**Figure S1.**
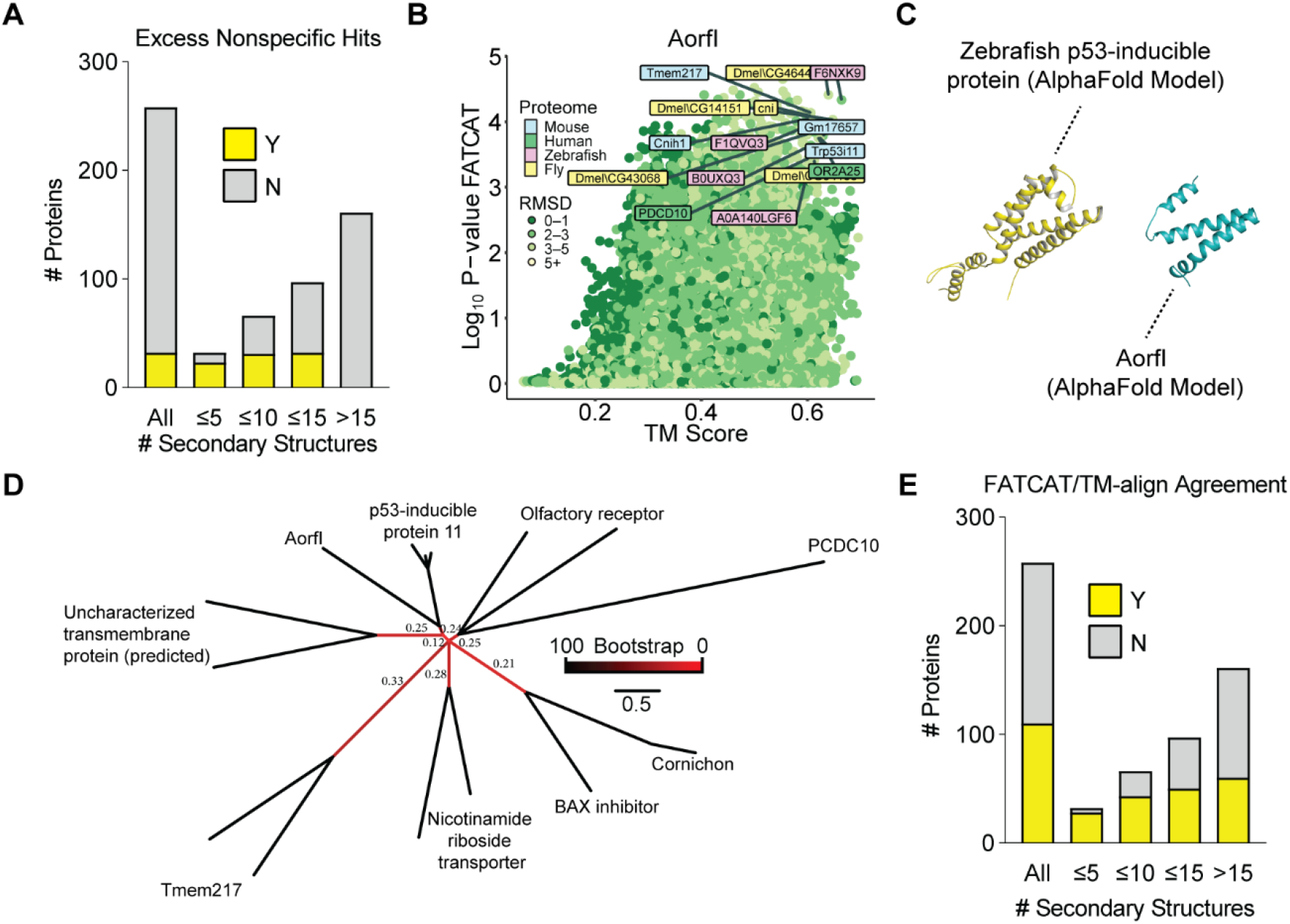
Related to Figure 1. a. Tabulation of nonspecific hits by number of protein secondary structures as determined by STRIDE. b. FATCAT and TM-align results for A orf I. Only a subset of the 100+ “hits” are indicated. c. Comparison of AlphaFold models of A orf I and danio p53-inducible protein 11. d. Maximum likelihood tree of the hits indicated in Figure S1A. Select bootstrap values from 100 replicates are indicated. e. Tabulation of proteins for which TM-align and FATCAT converged by number of protein secondary structures determined by STRIDE.

**Figure S2.**
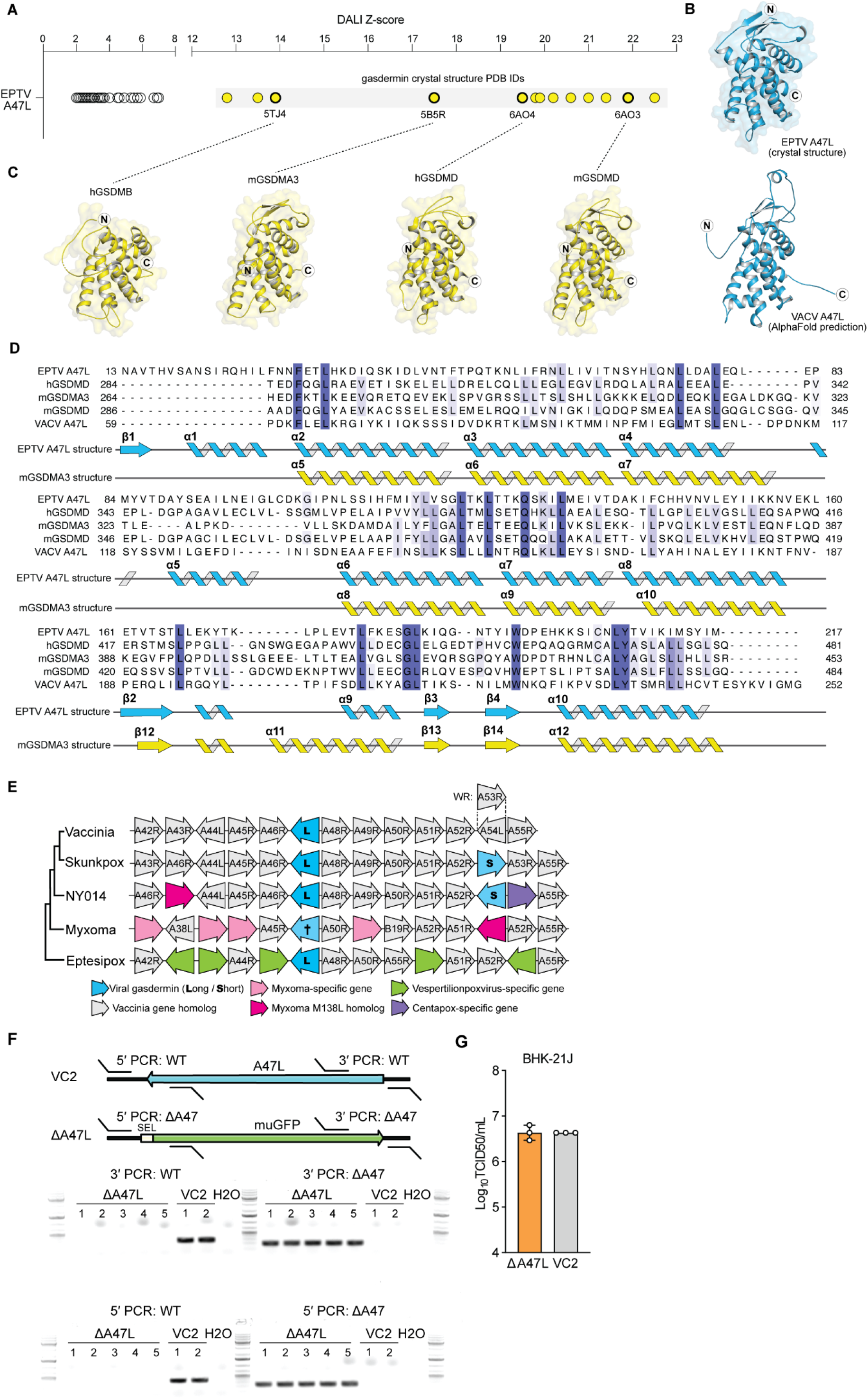
Related to Figure 2. a. DALI Z-scores from searching EPTV gasdermin structure against the full Protein Data Bank (PDB). Results for multiple chains were manually removed. PDB entries that represent gasdermin crystal structures in the PDB were highlighted yellow, and select PDB entries and the structures of four select structures are show in panel c. b. Experimental and predicted structures of poxvirus gasdermin proteins reveal gasdermin homology. Top, crystal structure of the Eptesipox virus (EPTV) gasdermin. Bottom, AlphaFold predicted structure of vaccinia virus (VACV) gasdermin. The N- and C-termini are indicated with the circled letters N and C. c. Representative published crystal structures of human (h) or mouse (m) gasdermins from the top EPTV A47L DALI hits shown in panel a. Shown structures are of the hGSDMB CTD (PDB ID 5TJ4), mGSDMA3 (PDB ID 5B5R, residues 264-453), hGSDMD CTD (PDB ID 6AO4), and mGSDMD (PDB ID 6AO3). The N- and C-termini are indicated with the circled letters N and C. d. Structure-based sequence alignment of the EPTV gasdermin crystal structure and structures from panels b and c. The hGSDMB structure was excluded due to large gaps in alignment, as were the first 58 residues of the VACV gasdermin AlphaFold predicted structure. The secondary structure of mGSDMA3 is based on the numbering scheme used previously ^28^. e. Synteny analysis of viral gasdermins. When applicable, genes are labeled based on their vaccinia homologs. †: The evolutionary history of leporipox gasdermin is ambiguous (see main text). f. Top: schematic of A47 locus for VC2 and ΔA47L recombinant virus with PCR genotyping primers indicated. Bottom: PCR genotyping of recombinant ΔA47L virus. VC2 (wild-type) is included as a control. ΔA47L clonal stock 1 was used for experiments. g. BHK-21J cells were infected with wild-type (VC2) or A47L-deficient vaccinia virus at an MOI of 1 and viral titer was quantified by TCID50 24 hours post-infection. n = 3 biological replicates.

**Figure S3.**
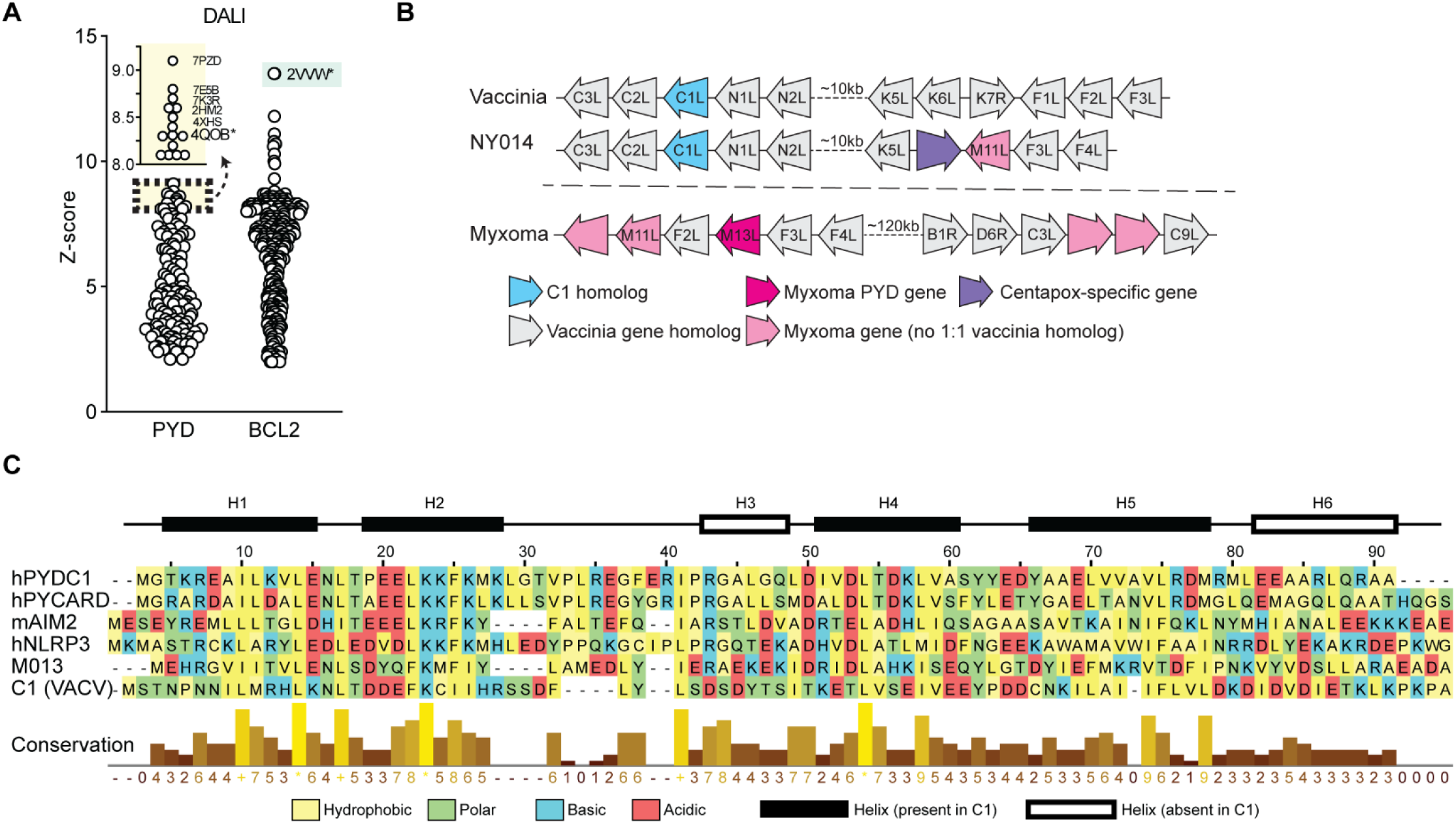
Related to Figure 3. a. DALI Z-scores from searching C1 subdomain AlphaFold models against the full Protein Data Bank (PDB). b. Synteny analysis of C1 and M013 families. When applicable, genes are labeled based on their vaccinia homologs. c. Amino acid alignment of select pyrin domains. Helices denoted are those present in ASC ^71^.

**Figure S4.**
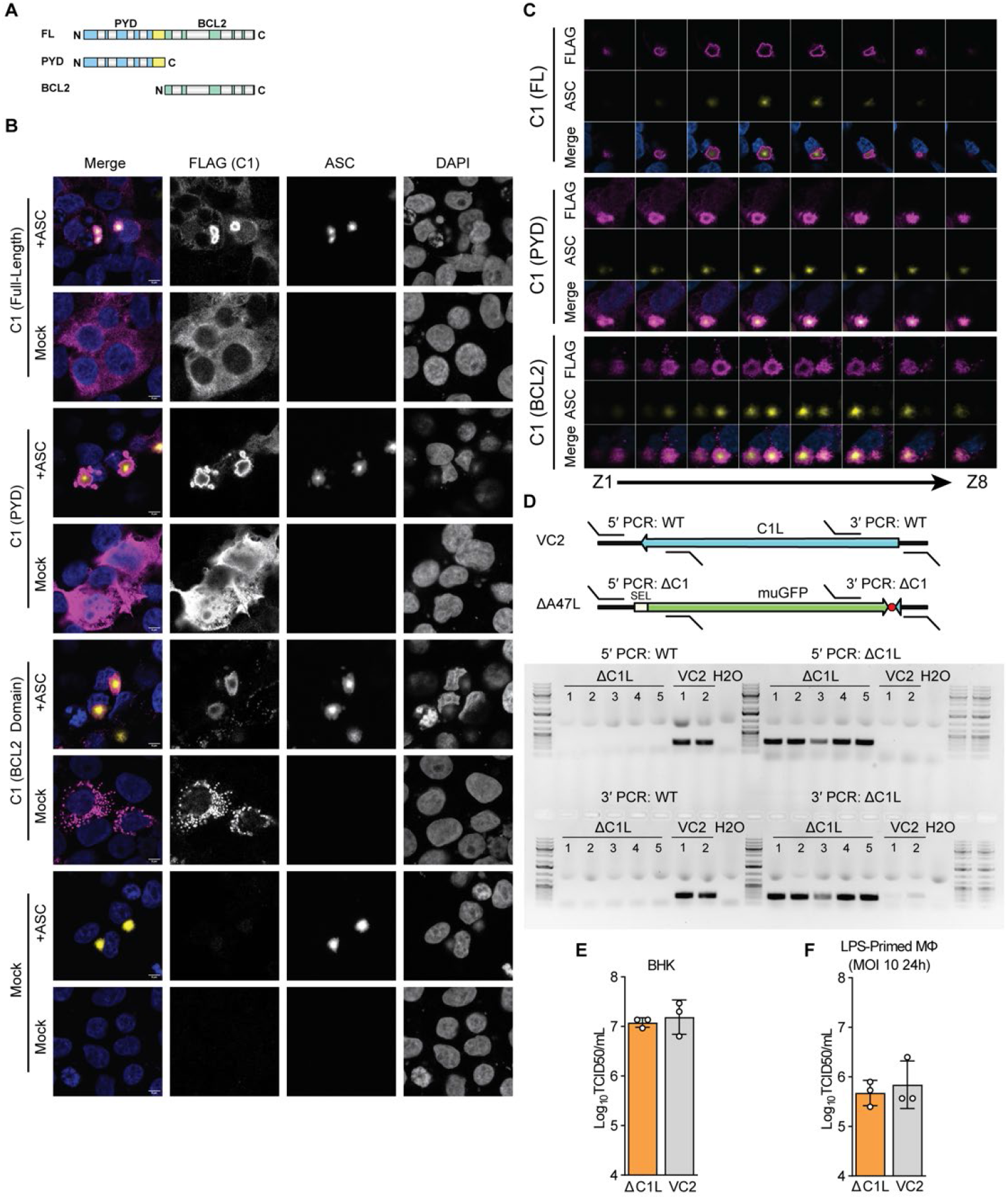
Related to Figure 4. a. C1 truncations used in this study. Helices (grey boxes) and the PYD-Bcl-2 linker (yellow) are indicated. b. 293T cells were co-transfected with the indicated plasmids and either ASC-GFP ^1^ or an empty vector. 24 hours post-transfection, cells were fixed and stained for FLAG (C1) and subsequently imaged. Representative image from n=3 independent experiments. Images for full-length C1 protein are the same as in Figure 4B. Scalebar: 5μm. c. Z stacks of ASC specks in 293T cells co-transfected with ASC-GFP and the indicated FLAG-tagged C1-expressing constructs. Z stacks are from the same experiment shown in Figure 4B. d. Top: schematic of C1L locus for VC2 and ΔC1L recombinant virus with PCR genotyping primers indicated. Red octagon represents the stop codon introduced in-frame to preserve the coding sequence of N1L (see methods). Bottom: PCR genotyping of recombinant ΔC1L virus. VC2 (wild-type) is included as a control. ΔC1L clonal stock 2 was used for experiments. e. BHK-21J cells were infected with wild-type (VC2) or C1L-deficient vaccinia virus at an MOI of 1 and viral titer was quantified by TCID50 24 hours post-infection. n = 3 biological replicates. f. LPS-primed murine macrophages were infected with wild-type (VC2) or C1L-deficient vaccinia virus at an MOI of 10 and viral titer was quantified by TCID50 24 hours post-infection. n = 3 biological replicates. Samples are from the same experiment as Figure 4G.

## List of Supplemental Files

Table S1: Vaccinia homology screen results

Table S2: Accession numbers used in this study

Table S3: Oligos and recombinant DNA used or generated in this study

Table S4: Crystallographic statistics

File S1: All vaccinia virus AlphaFold models

File S2: All vaccinia screen results (images)

File S3: Alignments and trees used for phylogenetic analyses

File S4: Scripts used in this study

