## Supplemental Materials for "Structural homology screens reveal poxvirus-encoded proteins impacting inflammasome-mediated defenses"

### Vaccinia screen results summary.

**Converged** = TM-align & FATCAT agreement **Excess Hits** = Large quantity of nonspec
**Proteins with <16 secondary structural motifs** (as determined by STRIDE) are highlighte
**%ID** was calculated based on a clustal omega alignment of the indicated uniprot identifi
**Some proteins with hits** have other hits that are extrememly close to the annotated hit.

| Uniprot ID | Orf Name | pLDDT | Length (AA) | ssMotifs | Converged | Excess Hits |
|---|---|---|---|---|---|---|
| P21113 | 7kDaAtIP | 68.74233 | 65 | **9** | N | |
| P20982 | A1 | 87.2308 | 150 | 22 | N | |
| P20642 | A10 | 77.11623 | 891 | 122 | N | |
| P20988 | A11 | 56.09341 | 318 | 25 | N | |
| P20989 | A12 | 62.00416 | 192 | 19 | N | |
| P20990 | A13 | 70.54972 | 70 | **4** | Y | |
| P20991 | A14 | 69.03298 | 90 | **2** | Y | Y |
| p0ck28 | A14p5 | 89.63571 | 53 | **2** | Y | Y |
| P20992 | A15 | 89.0949 | 94 | **15** | N | |
| P20993 | A16 | 81.98536 | 378 | 64 | N | |
| P68592 | A17 | 71.33835 | 203 | 22 | N | |
| p20534 | A18 | 86.7 | 493 | 74 | Y | |
| P20994 | A19 | 66.21313 | 77 | **15** | N | |
| P68318 | A2 | 77.66306 | 224 | 23 | N | |
| P20995 | A20 | 90.89355 | 426 | 58 | N | |
| P20996 | A21 | 85.92326 | 117 | 19 | N | |
| P20997 | A22 | 88.70113 | 176 | 23 | Y | |
| P20998 | A23 | 91.58571 | 382 | 60 | N | |
| P68694 | A24 | 91.11326 | 1164 | 213 | Y | |
| P21114 | A26 | 77.11563 | 322 | 34 | N | |
| p20535 | A27 | 78.05599 | 110 | **15** | Y | Y |
| P68632 | A28 | 81.33939 | 146 | 25 | N | |
| P21087 | A29 | 95.74789 | 305 | 51 | N | |
| p0ck20 | A2p5 | 69.70142 | 76 | **7** | N | |
| P20643 | A3 | 77.41739 | 644 | 97 | N | |
| P68595 | A30 | 78.62309 | 77 | **6** | Y | Y |
| P21096 | A31 | 80.39635 | 124 | 24 | N | |
| P68614 | A32 | 89.23297 | 270 | 42 | N | |
| P68616 | A33 | 74.04233 | 185 | 20 | Y | |
| P21057 | A34 | 90.60423 | 168 | 25 | Y | |
| P21058 | A35 | 88.00402 | 176 | 38 | N | |
| P68618 | A36 | 50.56892 | 221 | 18 | N | |
| P21060 | A37 | 87.80005 | 263 | 50 | N | |
| P21061 | A38 | 87.01837 | 277 | 36 | Y | |
| P21062 | A39 | 94.25722 | 403 | 85 | Y | |
| P20983 | A4 | 52.38249 | 281 | 28 | N | |
| P21063 | A40 | 66.87082 | 168 | 27 | Y | |
| P21064 | A41 | 89.74088 | 219 | 36 | N | |
| P68695 | A42 | 96.85205 | 133 | 25 | Y | |
| P21065 | A43 | 67.30829 | 194 | 24 | N | |
| P21097 | A44 | 95.30341 | 346 | 50 | Y | |

| | | | | | | |
|---|---|---|---|---|---|---|
| P21132 | A45 | 93.24252 | 125 | 22 | Y | |
| P21066 | A46 | 71.92699 | 214 | 37 | N | |
| P21067 | A47 | 75.06809 | 244 | 24 | Y | |
| P68693 | A48 | 96.80824 | 204 | 27 | Y | |
| P21068 | A49 | 91.49097 | 162 | 18 | N | |
| P68610 | A5 | 76.81659 | 164 | 17 | N | |
| p20492 | A50 | 90.70989 | 552 | 91 | Y | |
| P21069 | A51 | 83.25104 | 334 | 56 | N | |
| P21070 | A52 | 88.23601 | 190 | 17 | N | |
| P21071 | A53 | 66.26683 | 103 | 20 | N | |
| P68620 | A54 | 58.26713 | 90 | **5** | N | Y |
| P21073 | A55 | 87.92701 | 564 | 106 | Y | |
| P20978 | A56 | 64.19411 | 315 | 35 | N | |
| P21074 | A57 | 91.44574 | 151 | 20 | Y | |
| P20985 | A6 | 91.34894 | 372 | 30 | N | |
| p20635 | A7 | 84.76429 | 710 | 134 | N | |
| P20986 | A8 | 79.39829 | 288 | 35 | N | |
| P20987 | A9 | 65.02727 | 99 | **4** | Y | |
| p20510 | AorfA | 60.24153 | 119 | 26 | n | |
| p20511 | AorfB | 75.7881 | 86 | **1** | N | Y |
| p20512 | AorfC | 40.40304 | 128 | **12** | N | |
| p20513 | AorfD | 49.14148 | 88 | **10** | Y | |
| p20514 | AorfE | 42.43184 | 166 | **13** | N | |
| p20515 | AorfF | 65.75827 | 75 | **12** | N | |
| p20516 | AorfG | 59.32088 | 75 | **12** | N | |
| p20517 | AorfH | 36.84384 | 128 | **13** | N | |
| p20518 | AorfI | 55.97135 | 73 | **6** | Y | Y |
| p20519 | AorfJ | 48.84006 | 73 | **9** | N | |
| p20520 | AorfK | 39.61527 | 70 | **11** | N | |
| P68624 | AorfL | 54.92525 | 88 | **7** | Y | |
| P68626 | AorfM | 58.78473 | 76 | **3** | Y | Y |
| p20523 | AorfN | 70.84175 | 70 | **1** | Y | Y |
| p20524 | AorfO | 40.7194 | 83 | **12** | N | |
| p20525 | AorfP | 35.78624 | 89 | **14** | N | |
| p20526 | AorfQ | 44.01094 | 77 | **7** | N | |
| p20527 | AorfR | 61.5128 | 66 | **4** | Y | Y |
| P68628 | AorfS | 57.68266 | 67 | **11** | N | |
| P68630 | AorfT | 52.02503 | 81 | **8** | N | |
| p20530 | AorfU | 63.18684 | 72 | **3** | Y | |
| p20505 | B1 | 95.97757 | 300 | 49 | Y | |
| P68687 | B10 | 89.35938 | 166 | 40 | Y | |
| P21007 | B11 | 69.78397 | 88 | **5** | Y | |
| P21098 | B12 | 94.05945 | 283 | 42 | Y | |
| P20841 | B13 | 87.6357 | 116 | 20 | Y | |
| P20842 | B14 | 85.83112 | 222 | 34 | Y | |
| P21089 | B15 | 93.55426 | 149 | 16 | Y | |
| P21116 | B16 | 81.66437 | 30 | **1** | Y | Y |
| P21075 | B17 | 45.73089 | 340 | 61 | N | |

| | | | | | | |
|---|---|---|---|---|---|---|
| P21076 | B18 | 87.10394 | 574 | 87 | Y | |
| P21077 | B19 | 93.77266 | 353 | 70 | N | |
| P20999 | B2 | 91.24447 | 219 | 42 | N | |
| P21078 | B20 | 49.01911 | 127 | 32 | N | |
| P21099 | B21 | 64.58082 | 91 | **13** | Y | |
| P21100 | B22 | 80.37308 | 181 | 21 | N | |
| P21101 | B23 | 73.59851 | 386 | 63 | Y | |
| P21102 | B24 | 81.14964 | 150 | 24 | Y | |
| P21103 | B25 | 76.88415 | 259 | 33 | Y | |
| P21104 | B26 | 67.3626 | 103 | **10** | Y | |
| P21105 | B27 | 77.64603 | 113 | **15** | N | |
| P21106 | B28 | 66.49728 | 122 | 24 | N | |
| P21000 | B3 | 75.25853 | 124 | 20 | Y | |
| P21001 | B4 | 74.79937 | 558 | 67 | N | |
| P21115 | B5 | 85.58255 | 317 | 62 | N | |
| P68443 | B6 | 80.86823 | 173 | **11** | Y | |
| P68445 | B7 | 69.05378 | 182 | 32 | N | |
| P21004 | B8 | 92.88861 | 272 | 45 | N | |
| P21005 | B9 | 81.06937 | 77 | **9** | N | |
| P68473 | BorfA | 46.51293 | 107 | **9** | N | |
| p20542 | BorfB | 59.96655 | 66 | **9** | N | |
| p20543 | BorfC | 36.44316 | 95 | 16 | N | |
| P68475 | BorfD | 59.85997 | 71 | **5** | Y | |
| P68477 | BorfE | 40.35753 | 91 | **13** | N | |
| p20546 | BorfF | 50.60222 | 75 | **10** | N | |
| p20547 | BorfG | 65.03501 | 91 | **2** | Y | Y |
| p20548 | BorfH | 43.73027 | 70 | **13** | N | |
| p20549 | BorfI | 56.48816 | 67 | **10** | N | |
| P21036 | C1 | 82.9674 | 224 | 20 | N | |
| P21043 | C10 | 85.73095 | 331 | 53 | N | |
| p20494 | c11 | 72.89068 | 142 | 24 | N | |
| p20531 | C12 | 86.81473 | 353 | 57 | Y | |
| P21044 | C13 | 64.16698 | 65 | **9** | N | |
| P21045 | C14 | 81.71396 | 82 | **12** | N | |
| P21037 | C2 | 85.64696 | 512 | 93 | Y | |
| P21090 | C23 | 88.57905 | 244 | 40 | N | |
| P68639 | C3 | 90.06353 | 263 | 56 | Y | |
| P21038 | C4 | 81.96497 | 316 | 51 | N | |
| P21039 | C5 | 89.17651 | 204 | 20 | Y | |
| P21040 | C6 | 92.6986 | 151 | 16 | N | |
| P68599 | C7 | 93.20519 | 150 | 29 | N | |
| P21041 | C8 | 83.61364 | 184 | 28 | N | |
| P21042 | C9 | 86.34565 | 634 | 85 | N | |
| P21119 | CorfA | 50.19871 | 69 | **12** | N | |
| P21120 | CorfB | 60.48731 | 72 | **6** | Y | Y |
| P21121 | CorfC | 71.63799 | 82 | **6** | Y | Y |
| P21122 | CorfD | 66.64765 | 70 | **3** | Y | Y |
| P21123 | CorfE | 84.93081 | 65 | **2** | Y | Y |

| | | | | | | |
|---|---|---|---|---|---|---|
| P20979 | D1 | 90.48484 | 844 | 140 | N | |
| P21012 | D10 | 94.27866 | 248 | 34 | N | |
| P20637 | D11 | 91.01643 | 631 | 100 | Y | |
| P20980 | D12 | 96.66837 | 287 | 34 | Y | |
| P68441 | D13 | 95.34617 | 551 | 105 | N | |
| P21008 | D2 | 84.00116 | 146 | 21 | N | |
| P21009 | D3 | 87.61361 | 237 | 38 | N | |
| p20536 | D4 | 96.6796 | 218 | 45 | Y | |
| P21010 | D5 | 89.58724 | 785 | 147 | N | |
| p20634 | D6 | 86.64423 | 637 | 112 | Y | |
| P21034 | D7 | 91.4259 | 161 | 34 | Y | |
| p20508 | D8 | 89.6699 | 304 | 45 | Y | |
| P21011 | D9 | 91.26687 | 213 | 32 | N | |
| p20550 | DorfA | 59.06901 | 84 | **4** | Y | Y |
| p20551 | DorfB | 52.66891 | 80 | **15** | N | |
| p20552 | DorfC | 41.73074 | 69 | **10** | N | |
| P68479 | DorfD | 55.33893 | 72 | **8** | N | |
| P68481 | DorfE | 48.15648 | 80 | 17 | N | |
| P68483 | DorfF | 65.46633 | 69 | **12** | Y | |
| P68485 | DorfG | 55.62048 | 90 | **5** | Y | Y |
| P68487 | DorfH | 69.0556 | 61 | **1** | Y | Y |
| p20553 | DorfI | 72.22042 | 74 | **2** | N | |
| P21079 | E1 | 94.07629 | 479 | 70 | N | |
| P21050 | E10 | 95.66748 | 95 | **10** | Y | |
| P68449 | E11 | 84.83324 | 129 | 22 | N | |
| P21080 | E2 | 89.72825 | 737 | 95 | N | |
| P21081 | E3 | 78.20192 | 190 | 27 | N | |
| P21082 | E4 | 68.36226 | 259 | 29 | N | |
| P21046 | E5 | 74.01381 | 341 | 34 | N | |
| P21047 | E6 | 84.78101 | 567 | 62 | N | |
| P68447 | E7 | 44.55575 | 166 | 20 | N | |
| P21049 | E8 | 80.53922 | 273 | 42 | N | |
| p20509 | E9 | 91.44555 | 1006 | 146 | Y | |
| P21085 | EorfA | 49.13843 | 71 | **9** | n | |
| p20554 | EorfB | 42.55432 | 106 | **9** | Y | |
| p20555 | EorfC | 42.52227 | 70 | **11** | N | |
| p20556 | EorfD | 53.83275 | 66 | **15** | N | |
| p20557 | EorfE | 63.04056 | 65 | **4** | Y | Y |
| p20558 | EorfF | 51.66814 | 115 | **11** | Y | |
| P68450 | F1 | 68.00448 | 226 | 21 | N | |
| P21095 | F10 | 89.65752 | 439 | 65 | N | |
| P21052 | F11 | 79.95733 | 354 | 45 | N | |
| P21053 | F12 | 88.68075 | 635 | 95 | N | |
| P20638 | F13 | 91.70991 | 372 | 71 | Y | |
| P21019 | F14 | 65.32265 | 73 | **6** | N | |
| P21020 | F15 | 81.8579 | 158 | 37 | N | |
| P21021 | F16 | 84.14321 | 231 | 25 | N | |
| P68454 | F17 | 70.68404 | 101 | 21 | N | |

| | | | | | |
|---|---|---|---|---|---|
| P68634 | F2 | 94.15347 | 147 | 26 | Y | |
| P21013 | F3 | 89.12424 | 480 | 94 | N | |
| p20493 | F4 | 88.21558 | 319 | 27 | Y | |
| P21014 | F5 | 70.75118 | 321 | 51 | N | |
| P68601 | F6 | 66.58151 | 74 | **10** | N | |
| P21016 | F7 | 50.84788 | 92 | **9** | N | |
| P21017 | F8 | 68.05785 | 65 | **5** | N | |
| P68452 | F9 | 83.26356 | 212 | 21 | N | |
| p20559 | ForfA | 59.44491 | 67 | **5** | Y | Y |
| p20560 | ForfB | 65.59204 | 78 | **7** | N | |
| p20561 | ForfC | 59.57557 | 96 | **7** | Y | Y |
| p20562 | ForfD | 67.73666 | 85 | **5** | Y | |
| p20563 | ForfE | 58.68662 | 71 | **5** | Y | Y |
| p20564 | ForfF | 66.70897 | 74 | **4** | Y | Y |
| P21022 | G1 | 87.26159 | 591 | 85 | Y | |
| P68457 | G2 | 93.47739 | 220 | 30 | N | |
| P68459 | G3 | 84.01472 | 111 | **14** | N | |
| P68461 | G4 | 91.5997 | 124 | 22 | N | |
| P21026 | G5 | 93.24123 | 434 | 65 | N | |
| P68315 | G5p5 | 77.45979 | 63 | **10** | N | |
| P21027 | G6 | 90.72357 | 165 | 36 | N | |
| P21028 | G7 | 64.33256 | 371 | 33 | N | |
| P68612 | G8 | 89.59898 | 260 | 50 | Y | |
| P21030 | G9 | 86.37393 | 340 | 51 | N | |
| p20565 | GorfA | 32.24285 | 132 | **13** | N | |
| p20566 | GorfB | 72.70645 | 72 | **1** | Y | Y |
| P20495 | H1 | 97.21163 | 171 | 21 | Y | |
| p20496 | H2 | 84.26683 | 189 | 28 | N | |
| p20497 | H3 | 86.44234 | 324 | 50 | N | |
| P68439 | H4 | 71.77599 | 795 | 107 | N | |
| p20538 | H5 | 65.45242 | 203 | **10** | N | |
| P68697 | H6 | 94.36452 | 314 | 41 | Y | |
| p20539 | H7 | 93.55347 | 146 | 16 | N | |
| p20567 | HorfA | 77.6054 | 71 | **2** | Y | Y |
| p20498 | I1 | 83.96295 | 312 | 50 | N | |
| P68604 | I2 | 67.87248 | 73 | **6** | Y | |
| p20499 | I3 | 81.56212 | 269 | 45 | N | |
| P20503 | I4 | 93.1913 | 771 | 112 | Y | |
| p20500 | I5 | 83.24622 | 79 | **8** | Y | Y |
| P68463 | I6 | 85.76244 | 382 | 60 | N | |
| p20501 | I7 | 91.60765 | 423 | 55 | N | |
| p20502 | I8 | 91.6925 | 676 | 120 | Y | |
| p20568 | IorfA | 57.44336 | 77 | **8** | Y | Y |
| P21032 | J1 | 73.72111 | 153 | **11** | N | |
| P68564 | J2 | 92.61234 | 177 | 38 | Y | |
| P21033 | J3 | 89.63018 | 333 | 37 | Y | |
| P68608 | J4 | 92.21356 | 185 | 33 | Y | |
| P21083 | J5 | 80.47185 | 133 | 20 | N | |

| | | | | | | |
|---|---|---|---|---|---|---|
| p20504 | J6 | 93.89595 | 1286 | 208 | Y | |
| p20632 | K1 | 95.64797 | 284 | 47 | Y | |
| p20532 | K2 | 89.54373 | 369 | 65 | Y | |
| P20639 | K3 | 92.30271 | 88 | 18 | Y | |
| p20537 | K4 | 92.23112 | 424 | 72 | Y | |
| P21084 | K5 | 73.44188 | 136 | 20 | Y | |
| P68465 | K6 | 88.78416 | 81 | **13** | Y | |
| P68467 | K7 | 88.34052 | 149 | 16 | Y | |
| p20569 | KorfA | 44.21513 | 81 | **12** | N | |
| p20570 | KorfB | 61.64135 | 79 | **6** | Y | Y |
| p20540 | L1 | 78.59391 | 250 | 34 | N | |
| P20843 | L2 | 74.66685 | 87 | **2** | Y | |
| P21031 | L3 | 79.56406 | 350 | 43 | N | |
| P20981 | L4 | 79.44113 | 251 | 45 | N | |
| P68622 | L5 | 77.47556 | 128 | 18 | N | |
| P20640 | M1 | 86.22016 | 472 | 59 | Y | |
| P21092 | M2 | 75.51065 | 220 | 39 | N | |
| P21054 | N1 | 95.31486 | 117 | **12** | Y | |
| P20641 | N2 | 75.03608 | 175 | **13** | N | |
| P21093 | O1 | 82.19695 | 666 | 77 | N | |
| P68690 | O2 | 97.4889 | 108 | 17 | Y | |
| p0ck22 | O3 | 80.35554 | 35 | **1** | Y | Y |
| P68489 | U7kDa | 73.8132 | 62 | **3** | Y | Y |
| P68491 | U8p7kDa | 84.77117 | 72 | **2** | Y | Y |

ific (such as groups of unrelated proteins, not a large protein family) observed. This commonly overlaps with va
d in yellow and excluded as hits. Some select proteins in this category were still further annotated and may be o
ers
While these are often orthologs of the same gene/protein, they may be of interest. See supplemental figures.

| Hit | Hit ID | %ID (Top Hit) |
|---|---|---|
| Helicases (top: HELLS) | Q9NRZ9 | 20.55 |
| Mitochondrial resolvase tefm | Q0P4D6 | 21.34 |
| Polymerases (Polr2b) | Q8CFI7 | 24.44 |
| Klrh1 | Q3UU75 | 23.87 |
| Klrb1c | P27814 | 20.75 |
| CD47 | Q61735 | 27.08 |
| Semaphorin Sema7a | Q9QUR8 | 27.23 |
| Klra1 | P20937 | 24.2 |
| Profilin (Pfn2) | A0A0R4IPL1 | 27.27 |
| Steroid dehydrogenases (Top: Hsd3b2) | P26149 | 32.46 |

| | | |
|---|---|---|
| sod1 | O73872 | 18.55 |
| GSDMD | P57764 | 20.09 |
| DTYMK | P23919 | 18.01 |
| Ligases (Lig3) | P97386 | 55.43 |
| TNFRSF1B | P20333 | 24 |
| Kelch proteins (diablo is top) | Q9VUU5 | 23.49 |
| Guk1 | Q64520 | 46.36 |
| Serinc1/drosophila ND-15 | | |
| PSME1 | Q06323 | 19.72 |
| K7DYB1 | | |
| Kinases (top: VRK1) | Q99986 | 40.6 |
| Kelch proteins (top: Danio kelch-like protein 12) | A0A2R8Q6R0 | 28.57 |
| **ambiguous** | | |
| Kinases (top: VRK1) | Q99986 | 33.33 |
| Serpins (top: Fly Spn42Dc) | A1Z6R4 | 33.04 *B13/B14 are fragment |
| Serpins (top: Serpina3j) | D3Z451 | 29.03 |
| Bcl2l2 | P70345 | 13.86 |

| | | |
|---|---|---|
| Ankyrin repeat and SOCS box-containing 15a | A0A2R8PZ70 | 20.53 |
| Below threshhold: IL1r2 | | |
| Ankrd37 | | |
| ASB4 | Q9Y574 | 20 |
| Ankyrin proteins (top: ANKRD22) | Q5VYY1 | 22.76 |
| Ankyrin proteins (top: Danio Ankyrin repeat and SOCS box pr | Q7T386 | 23.67 |
| ambiguous - ankyrin or protein phosphatase 1? | | |
| Slfn1 | Q3UU73 | 25.21 |
| Buffy | Q8T8Y5 | 20.96 |
| F3/IFNGR1 (below threshold) | | |
| Tmem91, KANTR | | |
| Strong pyrin domain hits w/ FATCAT alone, some BCL2 domains with TM-align. See main text. | | |
| Serpins (top: Serpinb1c) | Q5SV42 | 34.2 |
| Keap1 | Q14145 | 22.2 |
| C4BPA | P04003 | 37.45 |
| Kelch proteins( top: KLHL8) | Q9P2G9 | 21.03 |
| below thresshold: bcl2l15 | | |

| | | | |
|---|---|---|---|
| just under thresshold: Nudt2 | | | |
| Chd3, HELLS, smarcad1 - helicases | O16102 | 19.44 | |
| Cmtr1, 2 (methyltransferase) | Q9W4N2 | 19.79 | |
| UNG1 (danio) | A2BIQ2 | 19.81 | |
| smarcad1b (danio) | E7F1C4 | 21.98 | **look back |
| POL2RG | P62487 | 17.39 | |
| Ca5b | Q9QZA0 | 33.33 | |
| just under thresshold: NudT15, 16, etc | | | |
| | A1A750, weak | | |
| lots, ambiguous | | | |
| lots, ambiguous | | | |
| GFER | | | |
| PolD1 | P54358 | 24.26 | |
| TMEM18 | | | |
| CHIC1 | Q5VXU3 | 18.18 | |
| PLD4 | Q96BZ4 | 25.62 | |

| | | |
|---|---|---|
| Dut | Q9CQ43 | 59.86 |
| Rrm2b | Q6PEE3 | 75.24 |
| Ifitm10? | Q8BR26 | 17.11 |
| Ide | P22817 | 17.9 |

just below threshhold: FEN1/XRN2/XRN1 nucleases
just below threshhold: polr2l
just below threshhold: Plaat1, tmem222 - previously shown to have similarity to bacterial peptidase, NLPC/P6

| | | |
|---|---|---|
| PCNA | Q9VIT0 | 20.4 |
| Phosphatases (top: Danio tyrosine phosphatase Q0P4E3) | Q0P4E3 | 27.03 |
| top1 | E7FE49 | 19.59 |
| mt-co3 | | |
| Rrm1 | P07742 | 76.01 |

ambiguous, maybe odorant receptors/olfactory receptors?

| | | |
|---|---|---|
| helicases (top: DHX35) | Q9H5Z1 | 20.5 |
| TK1 | P04184 | 66.67 |
| cmtr2 | Q9UAS6 | 19.75 |
| POLR2E | P19388 | 21.88 |

| | | |
|---|---|---|
| POLR2A | P24928 | 24.07 |
| Ankyrin proteins (top: Nfkbid) | Q2TB02 | 19.03 |
| Serpins (top: SERPINA3 | P01011 | 24.59 |
| polr2g | P62488 | 20.48 |
| PLD3 | O35405 | 47.73 |
| Mgll | O35678 | 32.35 |
| Mgll | | |
| BCL2L10 | Q9Z0F3 | 18.8 |
| ambiguous | | |
| ASB2, other ankyrin | Q96Q27 | 23.94 |
| MCL1 | | |
| Glrx | P35754 | 46.23 |

ccinia proteins with few structural motifs.
f interest.

s of a Serpin present as a single ORF in vaccinia strain WR

family, Plaat is a phospholipase/acyltransferase within this family

**Synteny Analysis**

| Genome | Accession |
|---|---|
| Vaccinia virus | NC_006998 |
| Skunkpox virus | NC_031038 |
| NY014 poxvirus | NC_035469.1 |
| Myxoma virus | NC_001132.2 |
| Eptesipox virus | NC_035460.1 |

**Poxvirus BLAST Database**

| Genome | Accession |
|---|---|
| Cotia virus | HQ646181.2 |
| Saltwater crocodilepox | MK903858.1 |
| Deerpox virus | AY689437.1 |
| Fowlpox virus | MH706124.1 |
| Eastern grey kangaroopox virus | MF661791.2 |
| Lumpy skin disease virus | NC_003027.1 |
| Molluscum contagiosum virus | MH320554.1 |
| Orf virus | MG716417.1 |
| Pteropox virus | KU980965.1 |
| Sea otter poxvirus | MH427217.1 |
| Sheeppox virus | MN072631.1 |
| Squirrel poxvirus | HE601899.1 |
| Swinepox virus | MZ682626.1 |
| Yaba monkey tumor virus | AY386371.1 |

**Viral gasdermins**

| Protein | Accession |
|---|---|
| Vaccinia virus gasdermin (COP) | M35027.1:c153690-152956 |
| Rabbitpox virus gasdermin | AY484669.1:c159516-158782 |
| Horsepox virus gasdermin | KY349117.1:c166272-165538 |
| Buffalopox virus gasdermin | MG599038.1:c157906-157148 |
| Vaccinia virus gasdermin (WR) | AY243312.1:c154675-153917 |
| Cowpox virus gasdermin | KC813500.1:c153312-152578 |
| Orthopoxvirus abatino gasdermin | NC 055231.1:c163765-163031 |
| Ectromelia virus gasdermin | MN912466.1:c156293-155559 |
| Akhmeta virus gasdermin | MH607143.1:c167386-166652 |
| Taterapox virus gasdermin | DQ437594.1:c153307-152573 |
| Variola virus gasdermin (17CE) | KY358055.1:c145385-144651 |
| Variola virus gasdermin (1947) | DQ441432.1:c146026-145292 |
| Variola virus gasdermin (1969) | DQ441437.1:c146904-146170 |
| Alaskapox virus gasdermin | MN240300.1:c166555-165827 |
| Skunkpox virus gasdermin | KU749310.1:c167631-166873 |
| Volepox virus gasdermin | KU749311.1:c169057-168281 |
| Raccoonpox virus gasdermin | KU749309.1:c145369-144644 |
| Camelpox virus gasdermin (CMS, truncated) | AY009089.1:c154402-154034 |
| Camelpox virus gasdermin (truncated) | MK910851.1:c154204-153836 |
| NY014 poxvirus gasdermin | MF001305.1:c159640-158771 |
| Murmansk poxxvirus gasdermin | MF001304.1:c160603-159728 |
| Yoka poxvirus gasdermin | HQ849551.1:c134784-134035 |
| Skunkpox virus gasdermin (short) | KU749310.1:172437-172985 |
| Racconpox virus gasdermin (short) | KU749309.1:150174-150722 |
| Volepox virus gasdermin (short) | KU749311.1:173861-174445 |
| Murmansk poxxvirus gasdermin (short) | MF001304.1:c165763-165224 |
| NY014 poxvirus gasdermin (short) | MF001305.1:c164808-164269 |
| Myxoma virus gasdermin | NP_051846.1 |
| Rabbit fibroma virus gasdermin | NP_052021.1 |
| Eptesipox virus gasdermin | YP_009408097.1 |

**Vertebrate gasdermins**

| Protein | Accession |
|---|---|
| Human gasdermin a | NP_835465.2 |
| Chicken gasdermin a | NP_001026532.1 |
| Bos taurus gasdermin a | NP_001071558.1 |
| Macaca mulatta gasdermin a | XP_014975037.1 |
| Pan troglodytes gasdermin a | XP_001171222.1 |
| Desmodus rotundus gasdermin a | XP_024429158.1 |
| Rousettus aegyptiacus gasdermin a | XP_016020091.2 |
| Pteropus alecto gasdermin a | XP_006924750.1 |
| Halichoerus grypus gasdermin a | XP_035935879.1 |
| Octodon degus gasdermin a | XP_004633937.1 |
| Rhinatrema bivittatum gasdermin a | XP_029428328.1 |
| Amblyraja radiata gasdermin a | XP_032891479.1 |
| Mouse gasdermin a3 | Q5Y4Y6 |
| Mouse gasdermin a3 | Q9EST1 |

| | |
|---|---|
| Mouse gasdermin a2 | Q32M21 |
| Human gasdermin b | NP_001159430.1 |
| Bos taurus gasdermin b | NP_001231146.1 |
| Pan troglodytes gasdermin b | XP_009430521.1 |
| Sarcophilus harrisii gasdermin b | XP_023358160.1 |
| Pteropus alecto gasdermin b | XP_024897515.1 |
| Rousettus aegyptiacus gasdermin b | XP_036078993.1 |
| Desmodus rotundus gasdermin b | XP_024429114.1 |
| Macaca mulatta gasdermin b | XP_014975035.1 |
| Halichoerus grypus gasdermin b | XP_035935913.1 |
| Human gasdermin c | NP_113603.1 |
| Macaca mulatta gasdermin c | XP_28708741.1 |
| Bos taurus gasdermin c | NP_001039469.1 |
| Sarcophilus harrisii gasdermin c | XP_31803207.1 |
| Rousettus aegyptiacus gasdermin c | XP_036088919.1 |
| Desmodus rotundus gasdermin c | XP_024427836.1 |
| Felis catus gasdermin c | XP_019678683.1 |
| Pan troglodytes gasdermin c | XP_001153860.1 |
| Mus pahari gasdermin c | XP_021073018.1 |
| Human gasdermin d | NP_079012.3 |
| Mouse gasdermin d | NP_081236.1 |
| Bos taurus gasdermin d | NP_001039625.1 |
| Macaca mulatta gasdermin d | XP_015001615.2 |
| Pteropus alecto gasdermin d | XP_006913014.1 |
| Rousettus aegyptiacus gasdermin d | XP_036089037.1 |
| Desmodus rotundus gasdermin d | XP_024428237.1 |
| Pan troglodytes gasdermin d | XP_009454389.2 |
| Felis catus gasdermin d | XP_023104116.1 |
| Human gasdermin e | NP_001120925.1 |
| Mouse gasdermin e | NP_061239.1 |
| Chicken gasdermin e | NP_001006361.1 |
| Bos taurus gasdermin e | NP_001180041.1 |
| Xenopus tropicalis gasdermin e | XP_002933445.2 |
| Macaca mulatta gasdermin e | XP_001096213.2 |
| Pteropus alecto gasdermin e | XP_006912063.1 |
| Sarcophilus harrisii gasdermin e | XP_031796450.1 |
| Chelonia mydas gasdermin e | XP_027689547.1 |
| Rana temporaria gasdermin e | XP_040208817.1 |
| Chrysemys picta bellii gasdermin e | XP_023962559.1 |
| Rhinatrema bivittatum gasdermin e | XP_029444983.1 |
| Octodon degus gasdermin e | XP_004626528.1 |
| Mus pahari gasdermin e | XP_021046698.1 |
| Pygosceli adeliae gasdermin e | XP_009324004.1 |
| Desmodus rotundus gasdermin e | XP_024418892.1 |
| Rousettus aegyptiacus gasdermin e | XP_016015046.2 |
| Amblyraja radiata gasdermin e | XP_032903802.1 |
| Pan troglodytes gasdermin e | XP_001159685.3 |

**Omitted from tree (severe truncation)**

| | |
|---|---|
| Hypsugopox virus gasdermin (pseudogene) | MK860688.1:c133573-132651 |

**Vertebrate pyrin domain containing proteins**

| Protein | Accession |
|---|---|
| POP1 (Human) | NP_690865.1 |
| ASC (Human) | NP_660183.1 |
| Danio PYD protein | NP_001104231.1 |
| ASC (Danio) | Q9I9N6.1 |
| PYDC2 (Human) | NP_001076777.1 |
| AIM2 (Mouse) | NP_001013801.2 |
| IFI203 (Mouse) | BAE43039.1 |
| Danio CASPb | NP_690840.2 |
| Danio CASPa | NP_571580.1 |

**M013L family**

| Protein | Accession |
|---|---|
| M013L | NP_051727.1 |
| S013L | NP_051902.1 |
| Deerpox M013L homolog | YP_227401.1 |
| Yaba M013L homolog | NP_073403.1 |
| Tanapox M013L homolog | ABQ43490.1 |
| Swinepox m013L homolog | NP_570174.1 |

**C1L family**

| Protein | Accession |
|---|---|
| Vaccinia C1 protein | P21036.1 |
| Buffalopox C1 homolog | QCY53926.1 |
| Cowpox C1 homolog | AGY97559.1 |
| Rabbitpox C1 homolog | AAS49732.1 |
| Horsepox C1 homolog | YP_010509239.1 |
| Orthopox abatino C1 homolog | YP_010085697.1 |
| Akhmeta virus C1 homolog | YP_010085480.1 |
| Variola B22L (C1 homolog) | AAA69361.1 |
| Monkeypox C1 homolog | YP_010377017.1 |
| Alaskapox C1 homolog | QED21215.1 |
| Volepox C1 homolog | YP_009281774.1 |
| Skunkpox C1 homolog | YP_009282720.1 |
| Raccoonpox C1 homolog | YP_009143336.1 |
| NY014 poxvirus C1 homolog | YP_009408409.1 |
| Murmansk poxvirus C1 homolog | YP_009408207.1 |
| Yoka poxvirus C1 homolog | YP_004821365.1 |

**Cloning**

| OLIGO |
|---|
| pCMV_hsGSDMD_N_R |
| pCMV_hsGSDMD_C_F |
| pCMV_hsGSDMD_C_R |
| pEF-Flag_F |
| pEF-C1_R |
| C1_PYD_r |
| C1_bcl2_fwd1 |
| C1_bcl2_fwd2 |
| cop_donor_c1_5pF |
| cop_donor_c1_5pR |
| cop_donor_c1_3pF |
| cop_donor_c1_3pR |

**PCR Genotyping for Recombinant Virus**

| OLIGO |
|---|
| a47_5p_flank_PCRfwd |
| a47_5p_flank_PCRrev_rec |
| a47_5p_flank_PCRrev_wt |
| a47_3p_flank_PCRfwd_rec |
| a47_3p_flank_PCRfwd_wt |
| a47_3p_flank_PCRrev |
| c1_5p_flank_PCRfwd |
| c1_5p_flank_PCRrev_wt |
| c1_3p_flank_PCRfwd_wt |
| c1_3p_flank_PCRrev |

| **SEQUENCE** |
| --- |
| CGTCAActcgagctaATCTGTCAAGAAATTGTGGAG |
| gacctctagaaccATGGAACAAAAACTCATCTCAGAAGAGGATCTGggttctGGGGTCCCCGCTGAAG |
| CGTCAACTCGAGctaATGGGGTTCTTGGCTAAGACCGGAC |
| gaatCCCGGGaccatggactacaaagacg |
| CCACTAgcggccgcctaTTTGCTCATAAGTTCCTTC |
| CATAgcggccgcctaGAGTATAGCAAATCTGACTGCTG |
| gacgacaagGGTGGAGGTGGTACCatgGACAAAATGACCGAGGATATTAAGC |
| gacctctagaaccatggactacaaagacgatgacgacaagGGTGGAGG |
| **gctc**aagcttttcatcctttgataattttccaaattc |
| gctaCAATTGttttttataaaaaatattgaatatataaacctttac |
| gcttCTCGAGttatttatttttcaccatatagatcaatcattagatc |
| gctaGAGCTCttcaaatgtttcattaactacgtg |

| **SEQUENCE** |
| --- |
| cgattaatgccccacgagac |
| **GTGACCGTTTACGTCGCC** |
| **gagcttagtccgtttgtccac** |
| **GAGGACGTGACGGCTG** |
| ctgagaggtcaatacctaactcc |
| **cgtctacatgttctgccgtaac** |
| cggagacgccgtaataatttcc |
| **cctacgataggacacgtaaaagc** |
| **catcatctgtaagatttttgagatgtc** |
| **cgctaagcgaatgatcgc** |

| NOTES |
| --- |
| PCR off of gBlock for full-length GSDMD |
| PCR off of gBlock for full-length GSDMD |
| PCR off of gBlock for full-length GSDMD |
| For all FLAG-tagged constructs (C1) |
| For C1 full length and C1 BCL2 domain |
| used with pEF-Flag_F |
| first PCR for BCL2 domain cloning: used with with pEF_C1_R |
| second PCR for BCL2 domain cloning: used with pEF_C1_R, had to add a third PCR round (pEF_C1_ |
| Used VC2 DNA as template to paste into cut pUC19_muGFP_A47_donor vector |
| Used VC2 DNA as template to paste into cut pUC19_muGFP_A47_donor vector |
| Used VC2 DNA as template to paste into cut pUC19_muGFP_A47_donor vector |
| Used VC2 DNA as template to paste into cut pUC19_muGFP_A47_donor vector |

| NOTES |
| --- |
| expect:250bp |
| expect: 380bp |
| expect: 287bp |
| expect: 191 |
| TM 57, for recomb: use with a47_5p_flank_PCRrev_rec: expect 270bp product, for WT use with c1_5 |
| TM56 |
| TM 54 |
| TM 55, for rec use with a47_5p_flank_PCRrev_wt, expect 220bp product, for WT expect 240 (with c1_ |

R) to add correct cut site due to mistake in cloning design

p_flank_PCRfwd, expect 240bp product

_5p_flank_PCRrev_wt)

| | |
|---|---|
| pCMV_vvGSDM | This Study |
| pCMV_GSDMD_CTD | This Study |
| pEF1α_C1 | This Study |
| pEF1α_C1_PYD | This Study |
| pEF1α_C1_BCL2 | This Study |
| pUC19_muGFP_Donor_A47 | This Study |
| pUC19_muGFP_Donor_C1 | This Study |
| pET_SUMO_eptvGSDM | This Study |

pLEX-MCS-ASC-GFP was a gift from Christian Stehlik (Addgene, catalog number: 73957)
pCMV-pro-IL1β-C-Flag was a kind gift from Bruce Beutler (Addgene, catalog number: 75131)
pcDNA3-N-Flag-NLRP3 was a kind gift from Bruce Beutler (Addgene, catalog number: 75127)
pcDNA3-N-Flag-ASC1 was a kind gift from Bruce Beutler (Addgene, catalog number: 75134)
pcDNA3-N-Flag-Caspase-1 was a kind gift from Bruce Beutler (Addgene, catalog number: 75128)
pcDNA3-N-HA-NEK7 was a kind gift from Bruce Beutler (Addgene, catalog number: 75142)

**Table S1. Crystallographic statistics, Related to Figure 2 and Figure NN**

| | EPTV A47L | EPTV A47L (SeMet) |
|---|---|---|
| **Data Collection** | | |
| Resolution (Å)[a] | 48.65–1.46 (1.48–1.46) | 48.39–1.54 (1.57–1.54) |
| Wavelength (Å) | 0.9792 | 0.9792 |
| Space group | C 2 2 $2_1$ | C 2 2 $2_1$ |
| Unit cell: a, b, c (Å) | 64.46 74.14 95.60 | 63.51, 74.71, 95.29 |
| Unit cell: α, β, γ (°) | 90.00 90.00 90.00 | 90.00, 90.00, 90.00 |
| Molecules per ASU | 1 | 1 |
| Total reflections | 543256 | 920245 |
| Unique reflections | 40114 | 33934 |
| Completeness (%)[a] | 100.0 (99.6) | 100.0 (99.9) |
| Multiplicity[a] | 13.5 (13.2) | 27.1 (23.4) |
| $I/\sigma I$[a] | 18.3 (1.6) | 15.7 (2.1) |
| CC(1/2)[b] (%)[a] | 99.9 (50.5) | 99.9 (88.8) |
| Rpim[c] (%)[a] | 1.9 (71.8) | 3.0 (68.9) |
| Sites | | 5 |
| **Refinement** | | |
| Resolution (Å) | 48.65–1.46 | |
| Free reflections | 40033 | |
| R-factor / R-free | 19.0 / 21.2 | |
| Bond distance (RMS Å) | 0.01 | |
| Bond angles (RMS °) | 1.10 | |
| **Structure/Stereochemistry** | | |
| No. atoms: protein | 1658 | |
| No. atoms: water | 85 | |
| Average B-factor: protein | 32.8 | |
| Average B-factor: water | 43.1 | |
| Ramachandran plot: favored | 99.01% | |
| Ramachandran plot: allowed | 0.99% | |
| Ramachandran plot: outliers | 0.00% | |
| Rotamer outliers | 0.52% | |
| MolProbity[d] score | 1.05 | |
| Protein Data Bank ID | **TBD** | |

[a] Highest resolution shell values in parentheses
[b] (Karplus and Diederichs, 2012)
[c] (Weiss, 2001)
[d] (Chen et al., 2010)

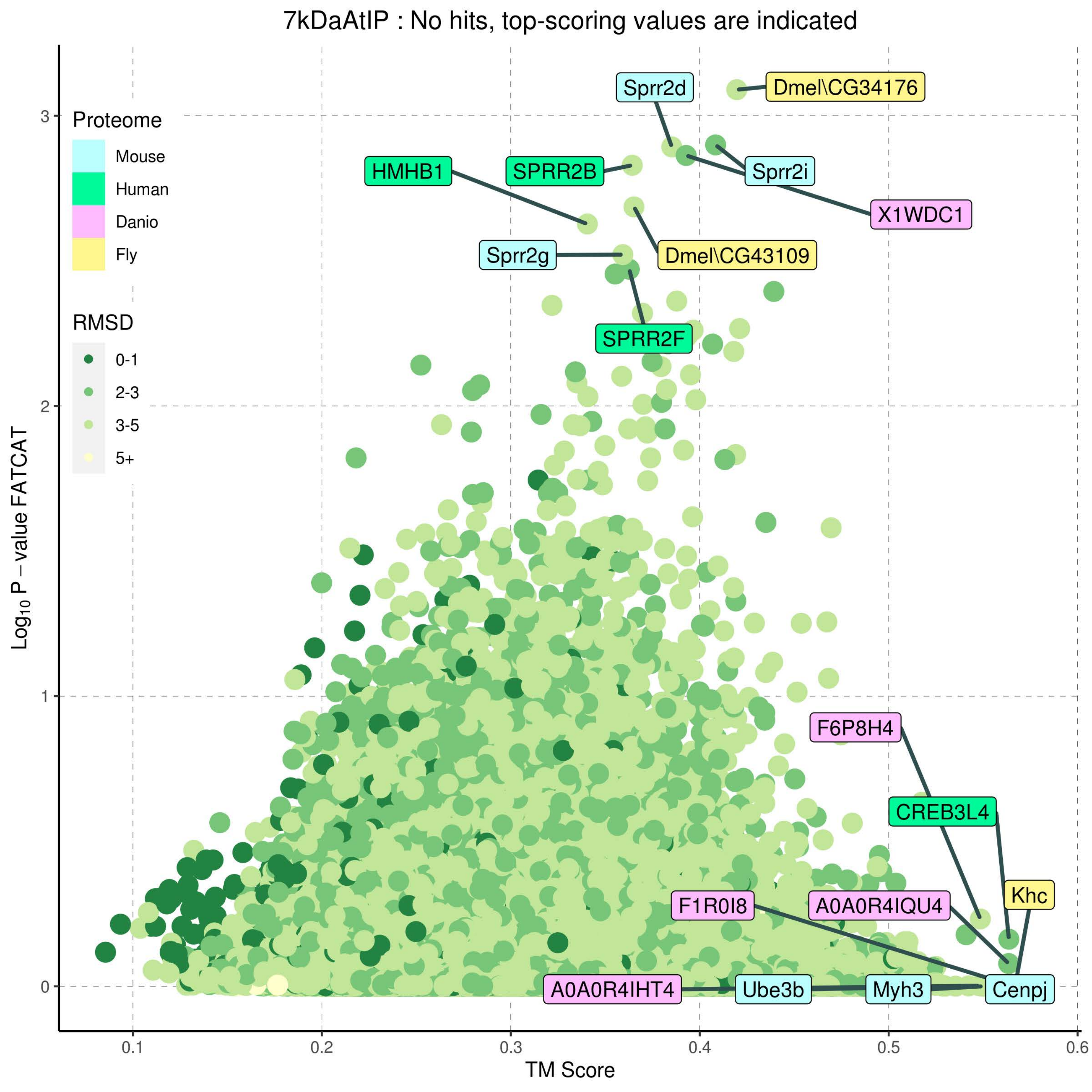

7kDaAtIP : No hits, top-scoring values are indicated
Proteome
Mouse
Human
Danio
Fly
RMSD
0-1
2-3
3-5
5+
Sprr2d
Dmel\CG34176
HMHB1
SPRR2B
Sprr2i
X1WDC1
Sprr2g
Dmel\CG43109
SPRR2F
F6P8H4
CREB3L4
Khc
F1R0I8
A0A0R4IQU4
A0A0R4IHT4
Ube3b
Myh3
Cenpj
Log10 P − value FATCAT
TM Score
0
1
2
3
0.1
0.2
0.3
0.4
0.5
0.6

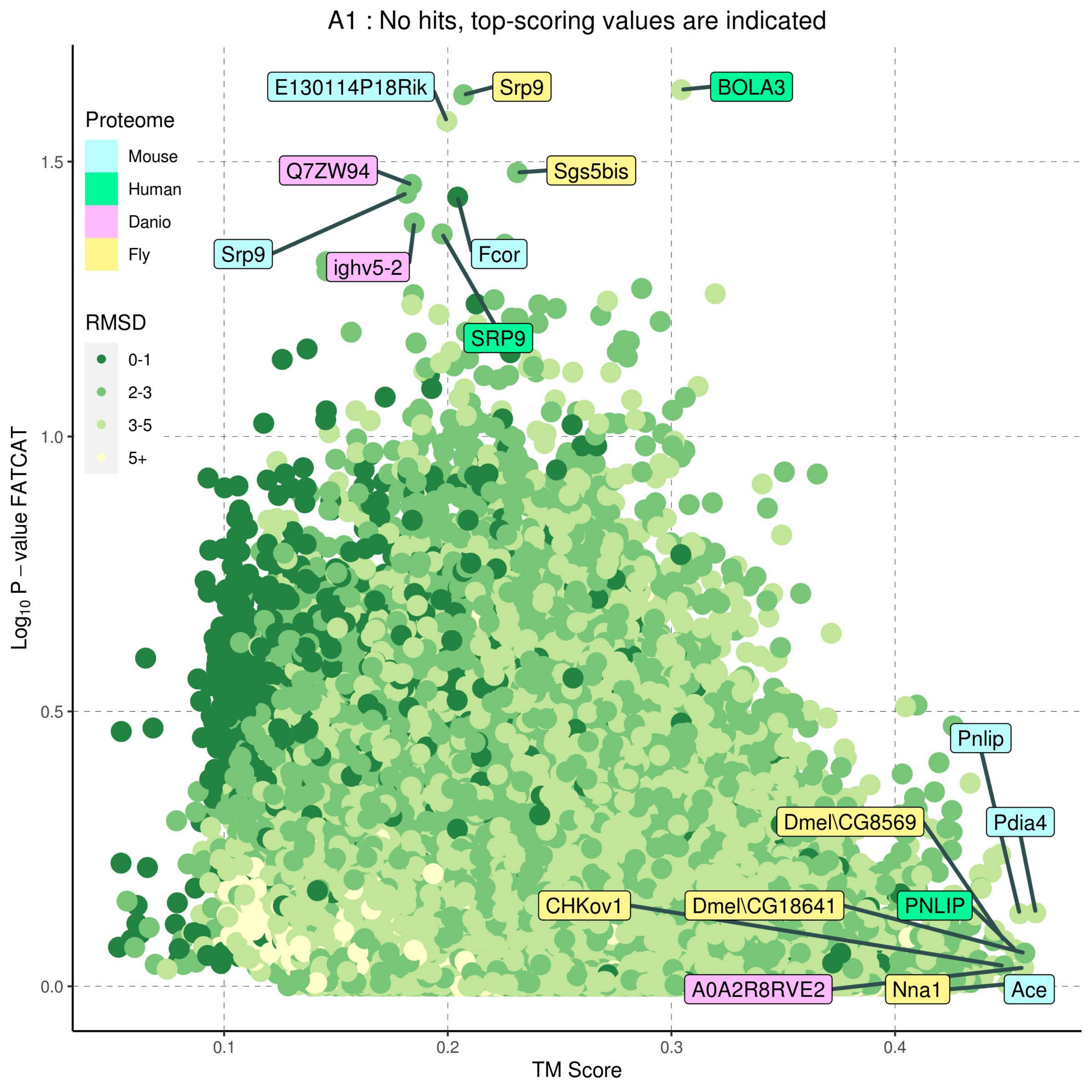

A1 : No hits, top-scoring values are indicated
Proteome
Mouse
Human
Danio
Fly
RMSD
0-1
2-3
3-5
5+
E130114P18Rik
Srp9
BOLA3
Q7ZW94
Sgs5bis
Srp9
ighv5-2
Fcor
SRP9
Pnlip
Pdia4
Dmel\CG8569
CHKov1
Dmel\CG18641
PNLIP
A0A2R8RVE2
Nna1
Ace
1.5
1.0
0.5
0.0
0.1
0.2
0.3
0.4
$Log_{10}$ P − value FATCAT
TM Score

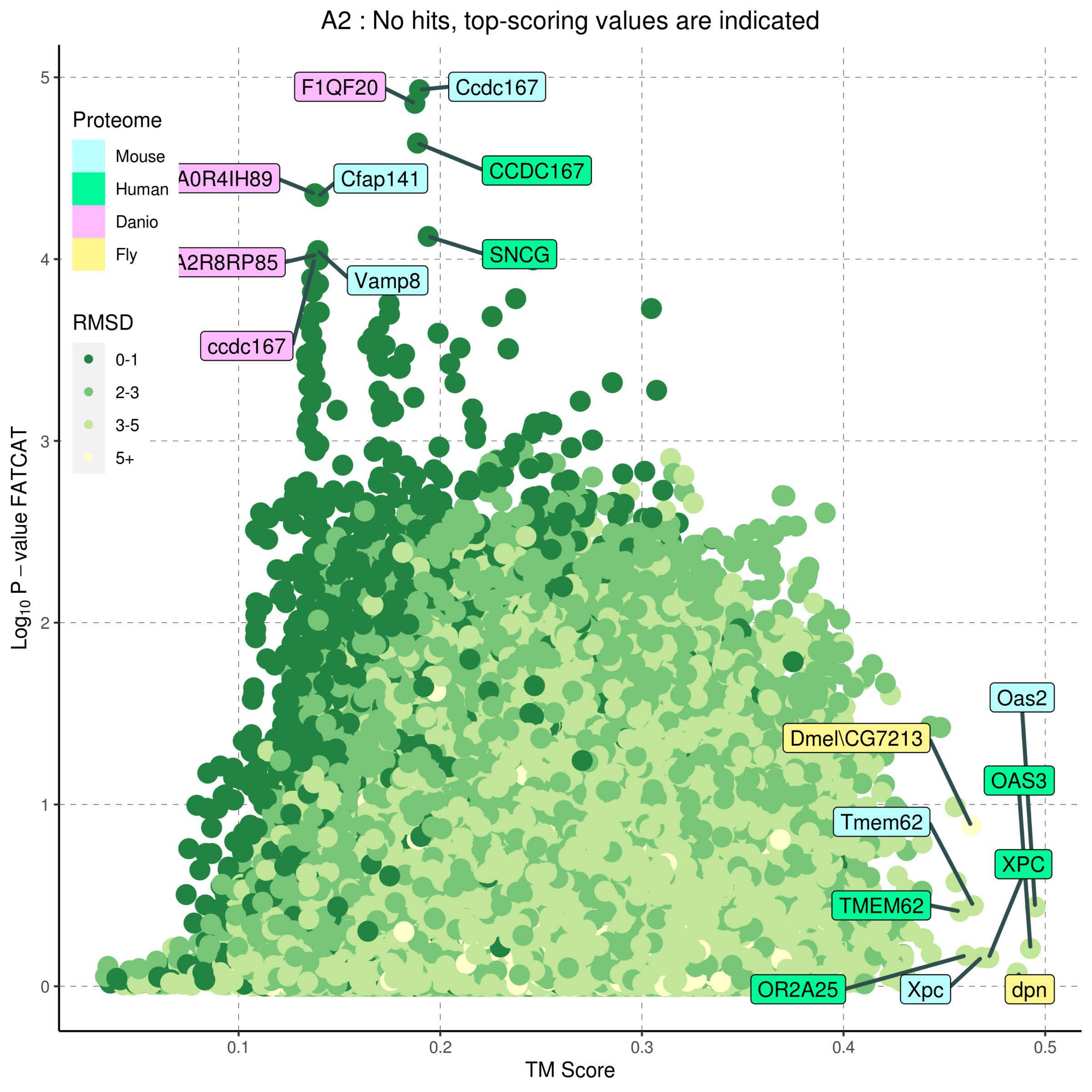
A2 : No hits, top-scoring values are indicated
Proteome
Mouse
Human
Danio
Fly
RMSD
0-1
2-3
3-5
5+
F1QF20
Ccdc167
CCDC167
A0R4IH89
Cfap141
SNCG
A2R8RP85
Vamp8
ccdc167
Oas2
Dmel\CG7213
OAS3
Tmem62
XPC
TMEM62
OR2A25
Xpc
dpn
Log10 P − value FATCAT
TM Score
0
1
2
3
4
5
0.1
0.2
0.3
0.4
0.5

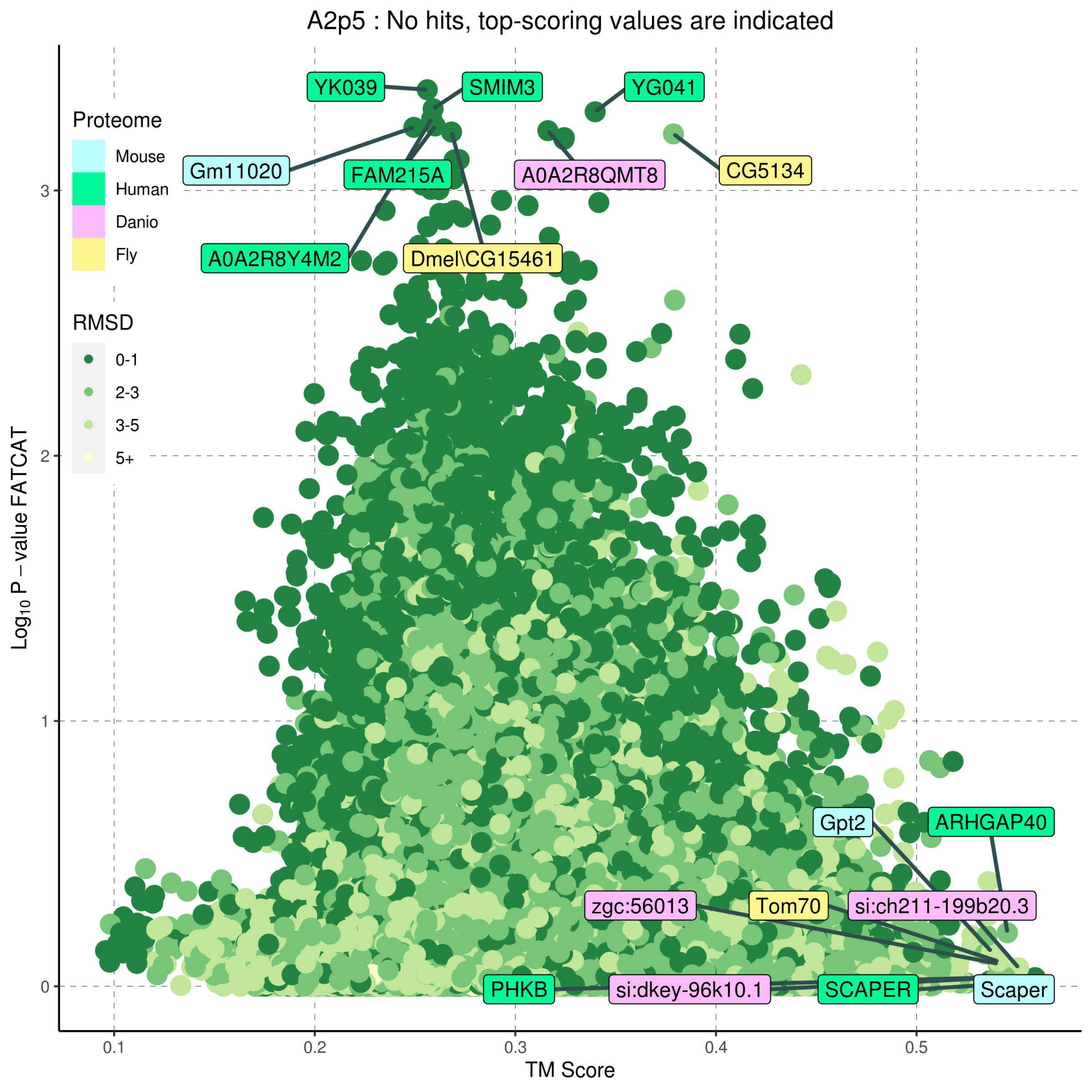

A2p5 : No hits, top-scoring values are indicated
Proteome
Mouse
Human
Danio
Fly
RMSD
0-1
2-3
3-5
5+
YK039
SMIM3
YG041
Gm11020
FAM215A
A0A2R8QMT8
CG5134
A0A2R8Y4M2
Dmel\CG15461
Gpt2
ARHGAP40
zgc:56013
Tom70
si:ch211-199b20.3
PHKB
si:dkey-96k10.1
SCAPER
Scaper
Log10 P – value FATCAT
TM Score
0
1
2
3
0.1
0.2
0.3
0.4
0.5

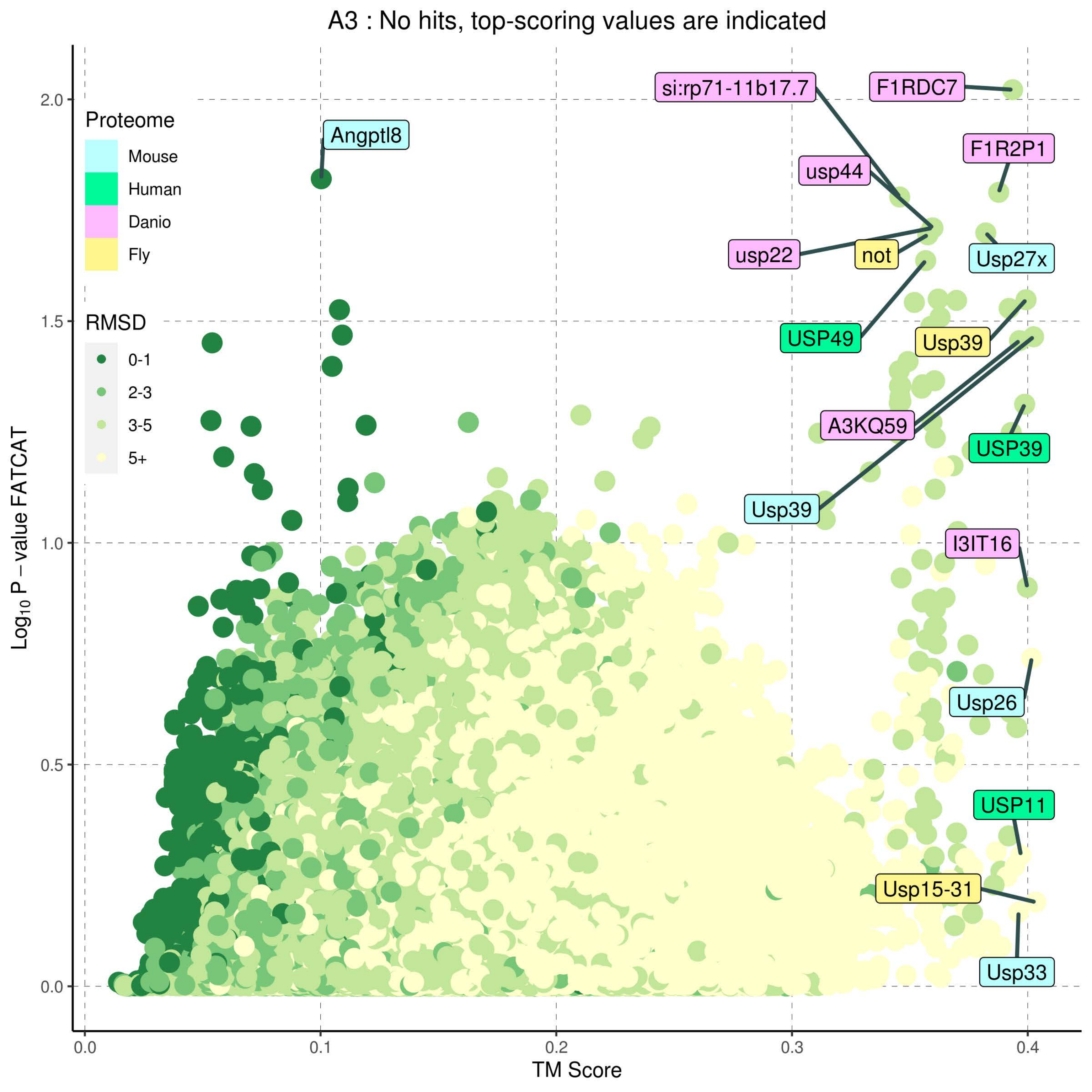
A3 : No hits, top-scoring values are indicated
Proteome
Mouse
Human
Danio
Fly
RMSD
0-1
2-3
3-5
5+
Angptl8
si:rp71-11b17.7
F1RDC7
F1R2P1
usp44
usp22
not
Usp27x
USP49
Usp39
A3KQ59
USP39
Usp39
I3IT16
Usp26
USP11
Usp15-31
Usp33
Log10 P – value FATCAT
TM Score
0.0
0.1
0.2
0.3
0.4
0.0
0.5
1.0
1.5
2.0

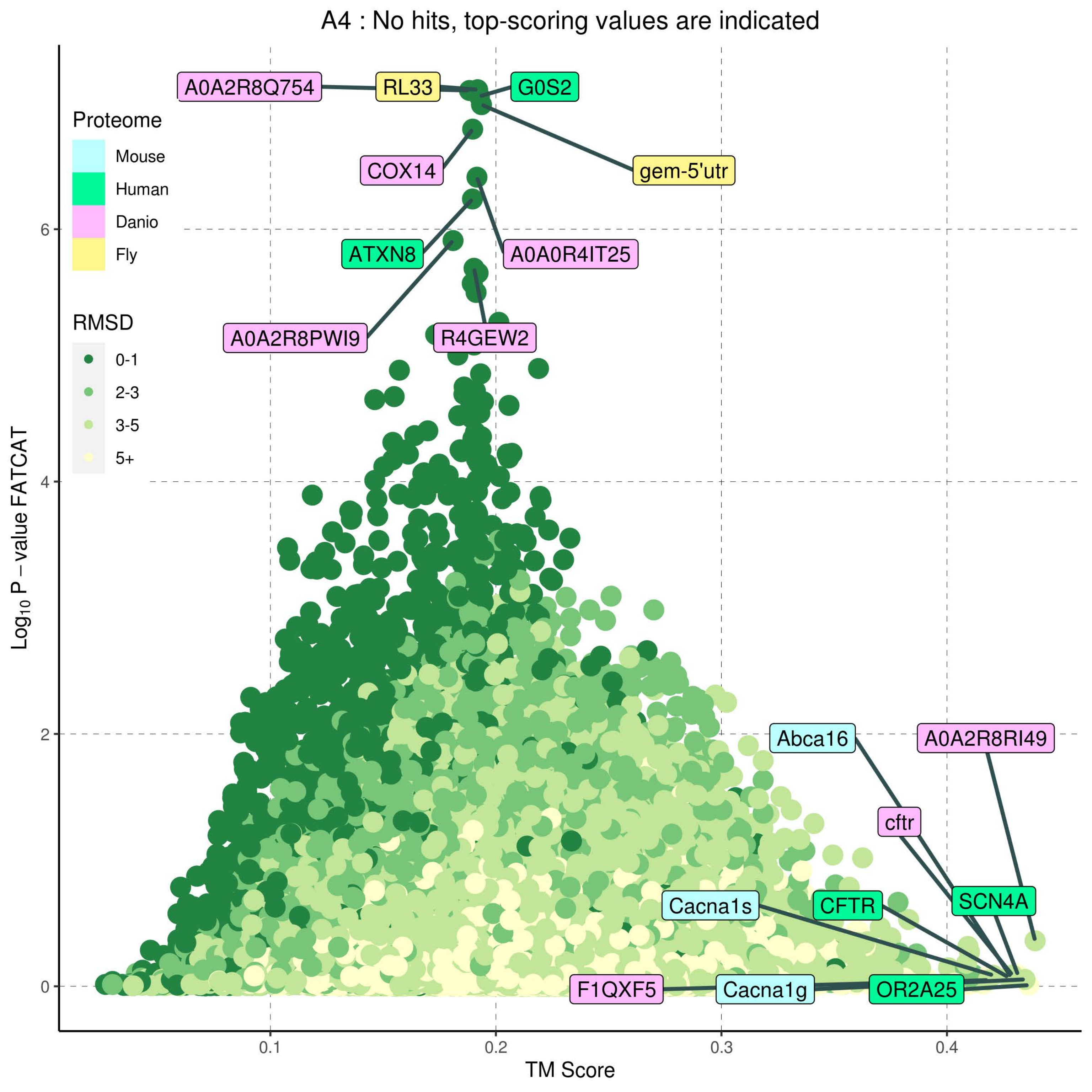
A4 : No hits, top-scoring values are indicated
Proteome
Mouse
Human
Danio
Fly
RMSD
0-1
2-3
3-5
5+
A0A2R8Q754
RL33
G0S2
COX14
gem-5'utr
ATXN8
A0A0R4IT25
A0A2R8PWI9
R4GEW2
Abca16
A0A2R8RI49
cftr
SCN4A
Cacna1s
CFTR
F1QXF5
Cacna1g
OR2A25
$Log_{10}$ P – value FATCAT
TM Score
0
2
4
6
0.1
0.2
0.3
0.4

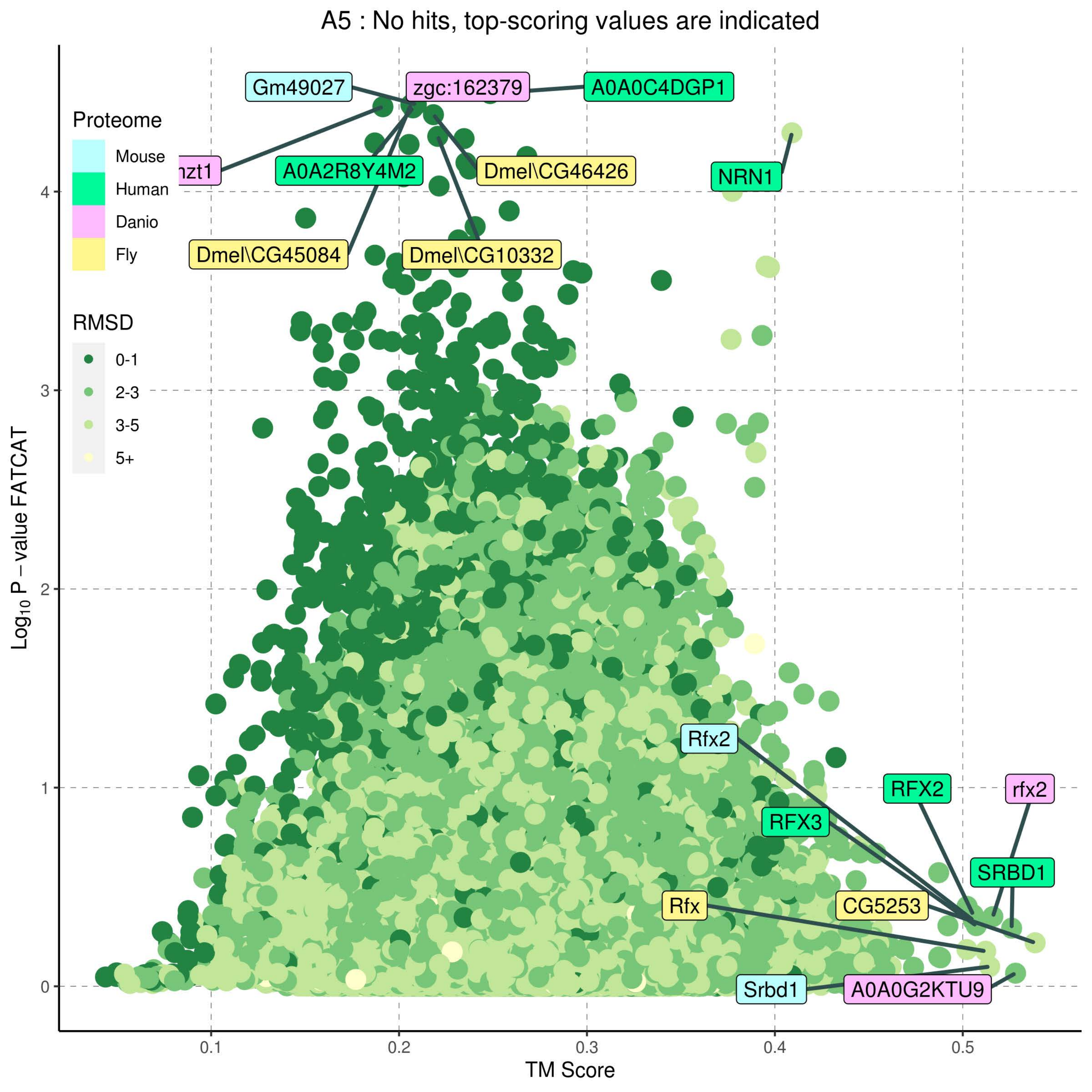

A5 : No hits, top-scoring values are indicated
Proteome
Mouse
Human
Danio
Fly
RMSD
0-1
2-3
3-5
5+
Gm49027
zgc:162379
A0A0C4DGP1
nzt1
A0A2R8Y4M2
Dmel\CG46426
NRN1
Dmel\CG45084
Dmel\CG10332
Rfx2
RFX2
rfx2
RFX3
SRBD1
Rfx
CG5253
Srbd1
A0A0G2KTU9
Log10 P – value FATCAT
TM Score
0
1
2
3
4
0.1
0.2
0.3
0.4
0.5

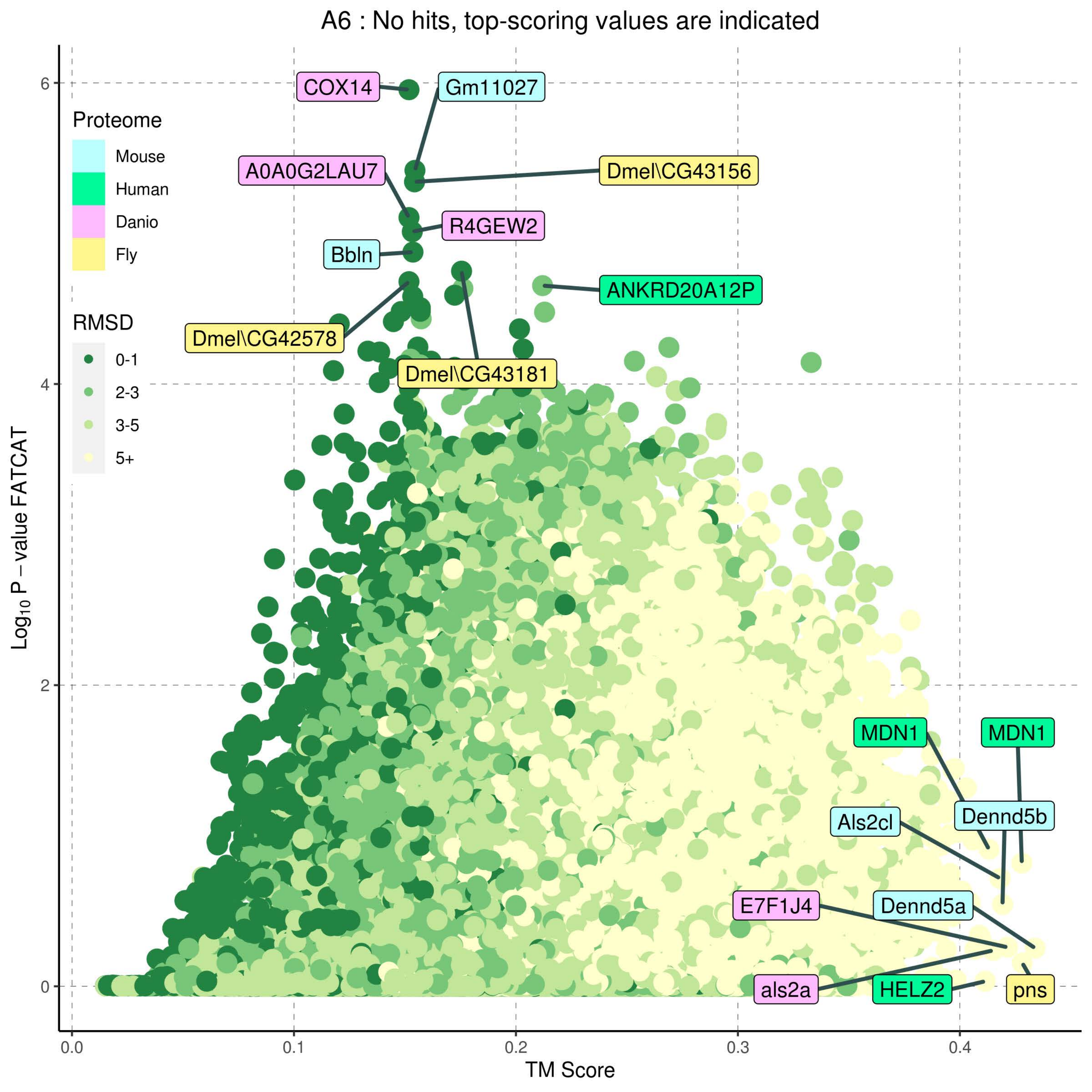

A6 : No hits, top-scoring values are indicated
Proteome
Mouse
Human
Danio
Fly
RMSD
0-1
2-3
3-5
5+
COX14
Gm11027
A0A0G2LAU7
Dmel\CG43156
R4GEW2
Bbln
ANKRD20A12P
Dmel\CG42578
Dmel\CG43181
MDN1
MDN1
Als2cl
Dennd5b
E7F1J4
Dennd5a
als2a
HELZ2
pns
0
2
4
6
0.0
0.1
0.2
0.3
0.4
Log10 P – value FATCAT
TM Score

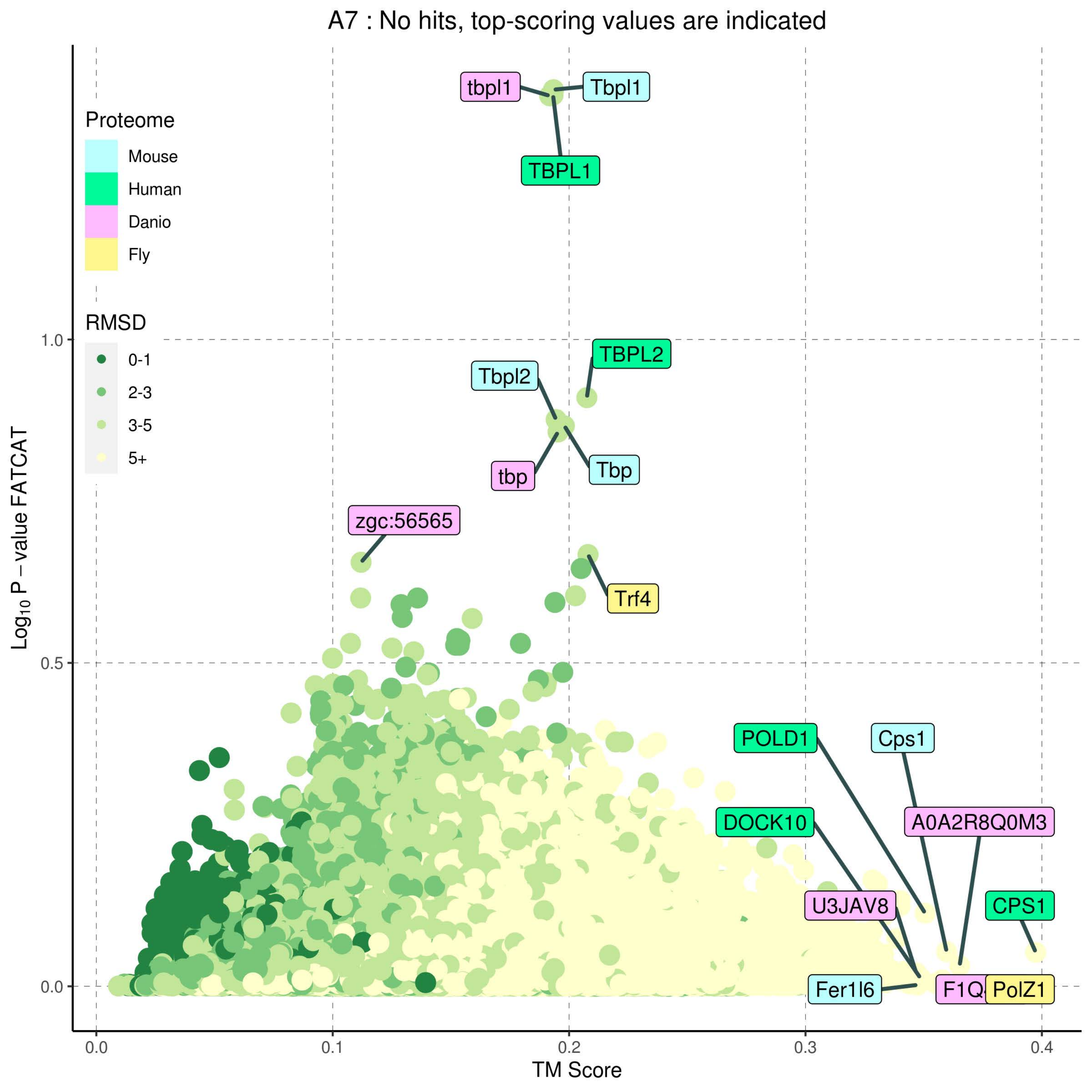
A7 : No hits, top-scoring values are indicated
Proteome
Mouse
Human
Danio
Fly
RMSD
0-1
2-3
3-5
5+
tbpl1
Tbpl1
TBPL1
TBPL2
Tbpl2
tbp
Tbp
zgc:56565
Trf4
POLD1
Cps1
DOCK10
A0A2R8Q0M3
U3JAV8
CPS1
Fer1l6
PolZ1
1.0
0.5
0.0
0.0
0.1
0.2
0.3
0.4
Log10 P − value FATCAT
TM Score

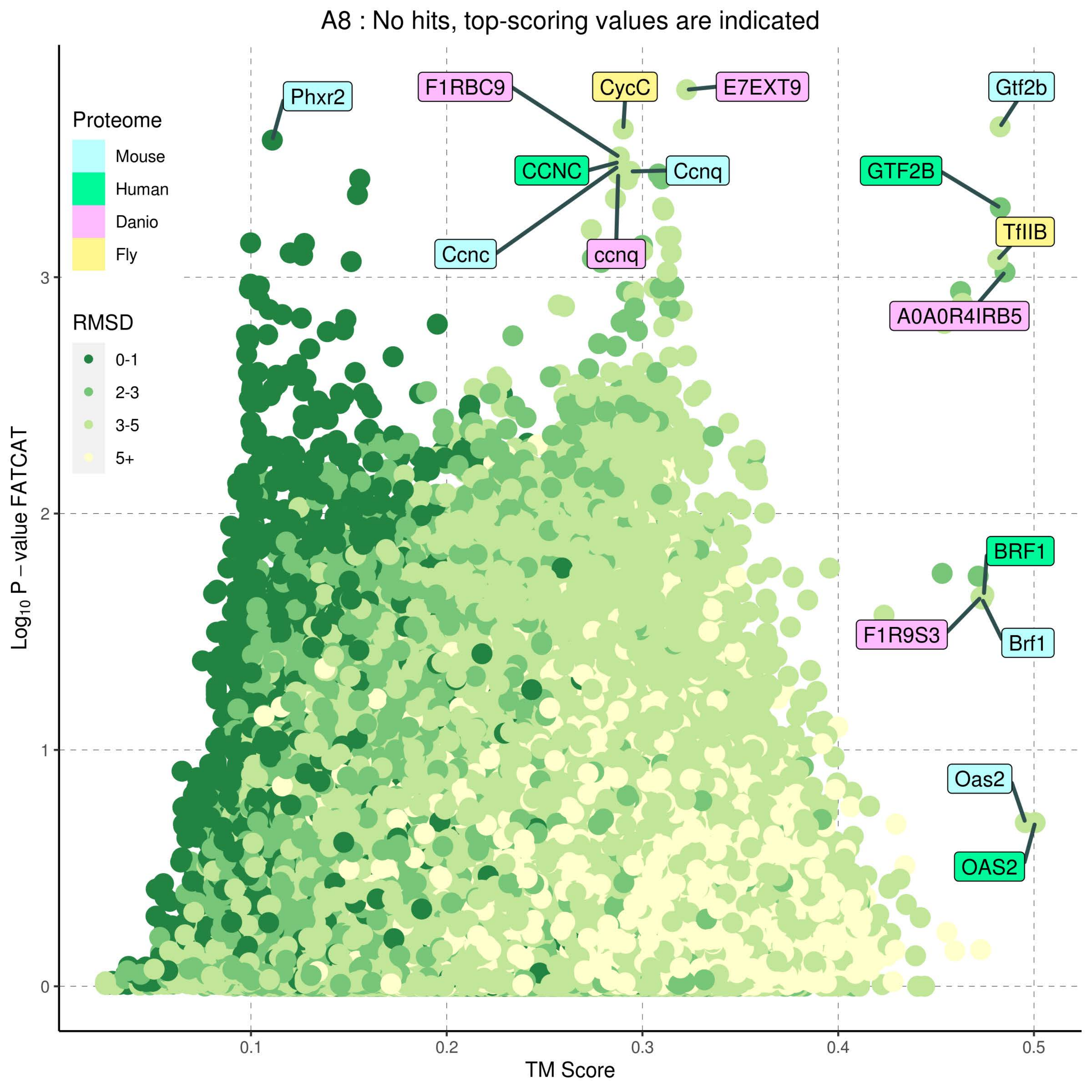
A8 : No hits, top-scoring values are indicated
Proteome
Mouse
Human
Danio
Fly
RMSD
0-1
2-3
3-5
5+
Phxr2
F1RBC9
CycC
E7EXT9
Gtf2b
CCNC
Ccnq
GTF2B
Ccnc
ccnq
TfIIB
A0A0R4IRB5
BRF1
F1R9S3
Brf1
Oas2
OAS2
0
1
2
3
0.1
0.2
0.3
0.4
0.5
TM Score
$Log_{10}$ P – value FATCAT

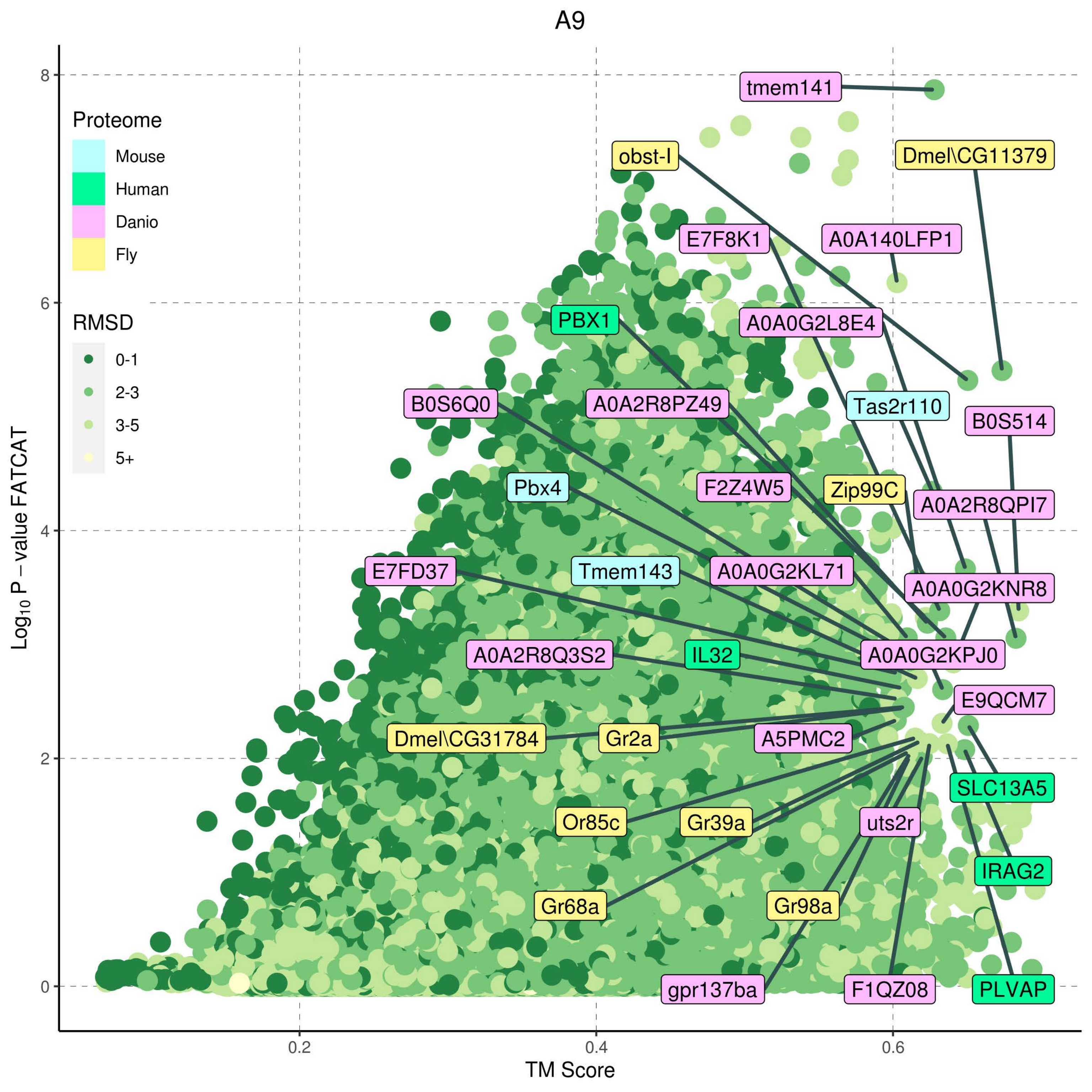

A9
Proteome
Mouse
Human
Danio
Fly
RMSD
0-1
2-3
3-5
5+
Log10 P − value FATCAT
TM Score
0
2
4
6
8
0.2
0.4
0.6
tmem141
obst-I
Dmel\CG11379
E7F8K1
A0A140LFP1
PBX1
A0A0G2L8E4
B0S6Q0
A0A2R8PZ49
Tas2r110
B0S514
Pbx4
F2Z4W5
Zip99C
A0A2R8QPI7
E7FD37
Tmem143
A0A0G2KL71
A0A0G2KNR8
A0A2R8Q3S2
IL32
A0A0G2KPJ0
E9QCM7
Dmel\CG31784
Gr2a
A5PMC2
SLC13A5
Or85c
Gr39a
uts2r
IRAG2
Gr68a
Gr98a
gpr137ba
F1QZ08
PLVAP

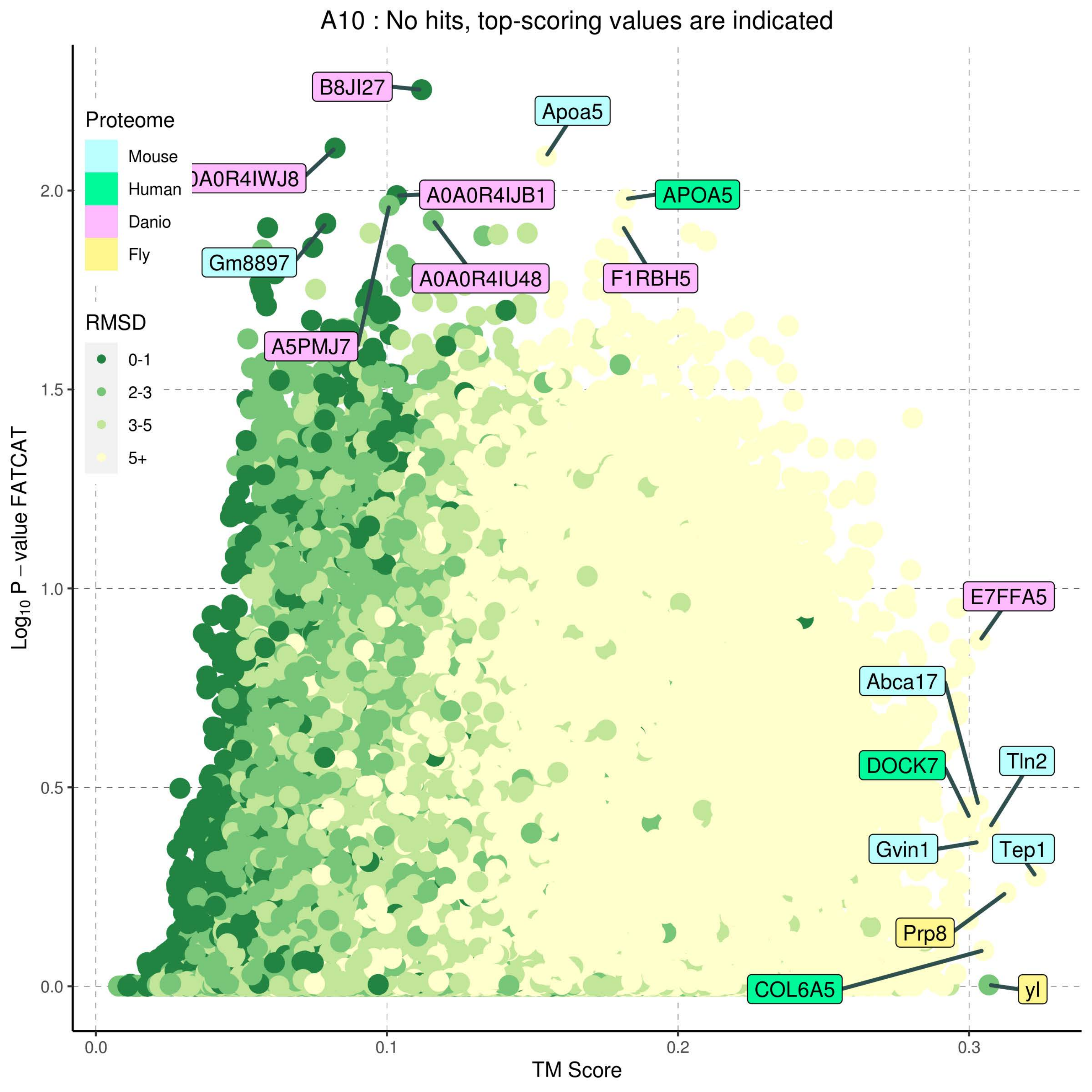
A10 : No hits, top-scoring values are indicated
Proteome
Mouse
Human
Danio
Fly
RMSD
0-1
2-3
3-5
5+
B8JI27
Apoa5
0A0R4IWJ8
A0A0R4IJB1
APOA5
Gm8897
A0A0R4IU48
F1RBH5
A5PMJ7
E7FFA5
Abca17
DOCK7
Tln2
Gvin1
Tep1
Prp8
COL6A5
yl
$Log_{10}$ P – value FATCAT
TM Score
0.0
0.5
1.0
1.5
2.0
0.0
0.1
0.2
0.3

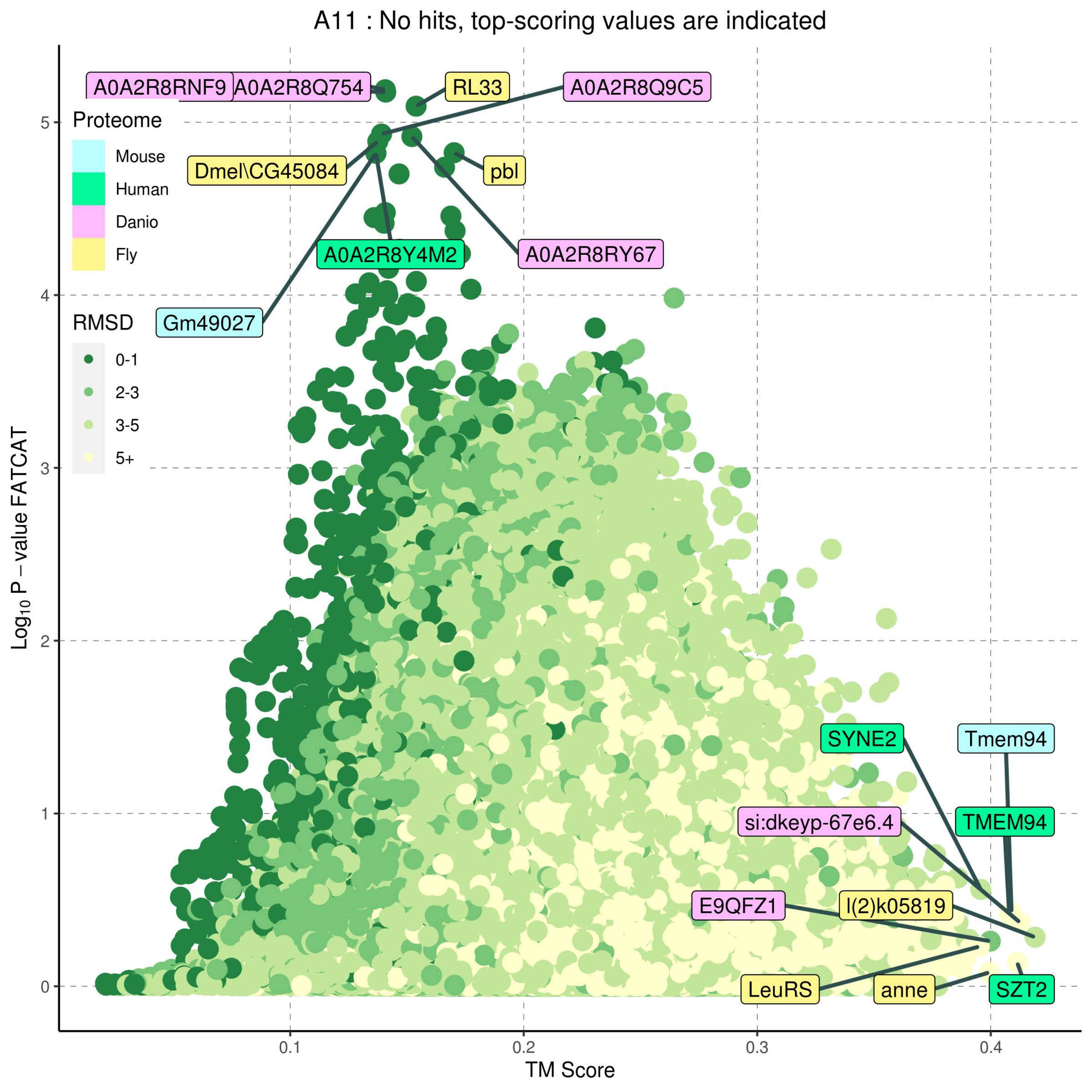

A11 : No hits, top-scoring values are indicated
Proteome
Mouse
Human
Danio
Fly
RMSD
0-1
2-3
3-5
5+
A0A2R8RNF9
A0A2R8Q754
RL33
A0A2R8Q9C5
Dmel\CG45084
pbl
A0A2R8Y4M2
A0A2R8RY67
Gm49027
SYNE2
Tmem94
si:dkeyp-67e6.4
TMEM94
E9QFZ1
l(2)k05819
LeuRS
anne
SZT2
Log10 P − value FATCAT
TM Score
0
1
2
3
4
5
0.1
0.2
0.3
0.4

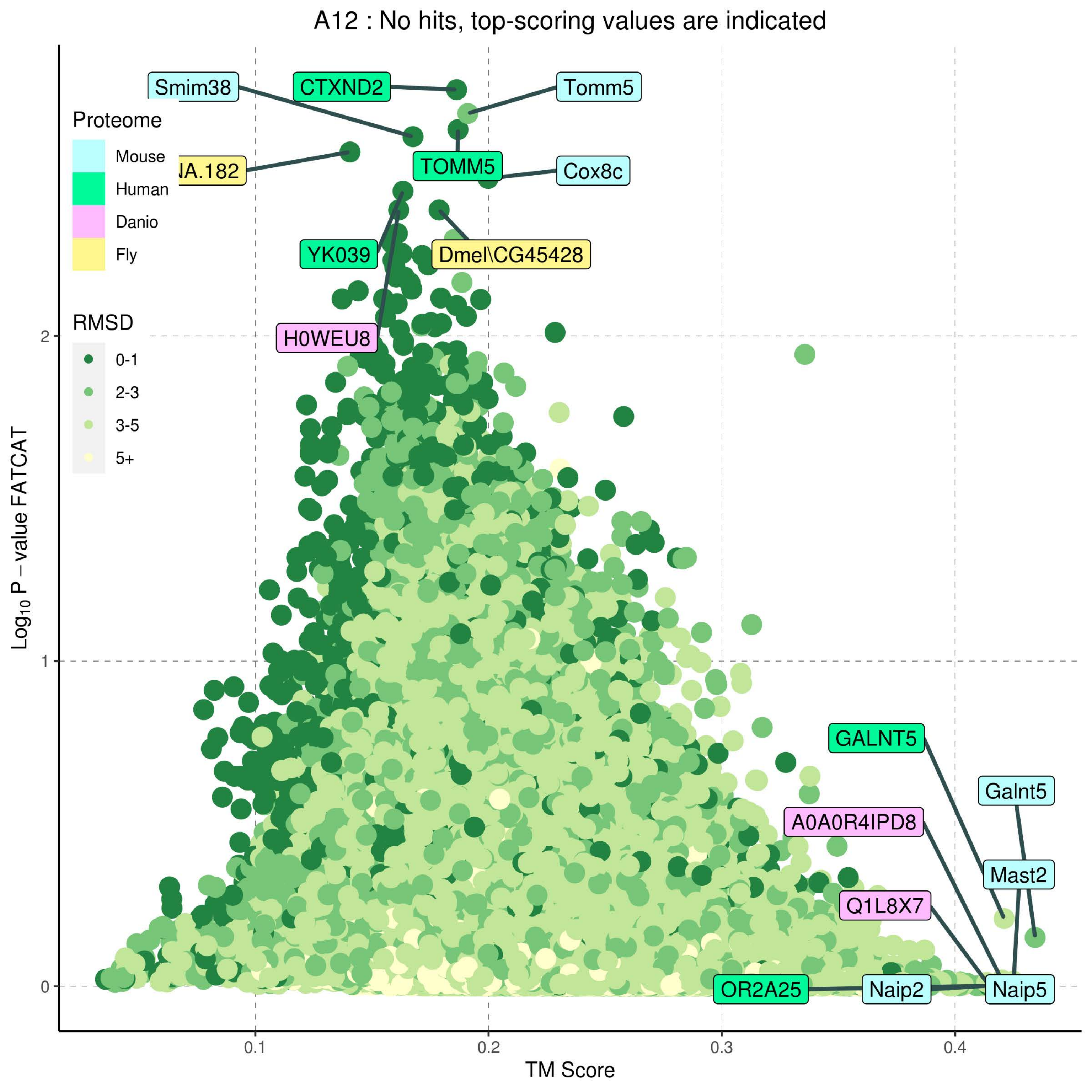

A12 : No hits, top-scoring values are indicated
Proteome
Mouse
Human
Danio
Fly
RMSD
0-1
2-3
3-5
5+
Smim38
CTXND2
Tomm5
NA.182
TOMM5
Cox8c
YK039
Dmel\CG45428
H0WEU8
GALNT5
Galnt5
A0A0R4IPD8
Mast2
Q1L8X7
OR2A25
Naip2
Naip5
Log10 P − value FATCAT
TM Score
0
1
2
0.1
0.2
0.3
0.4

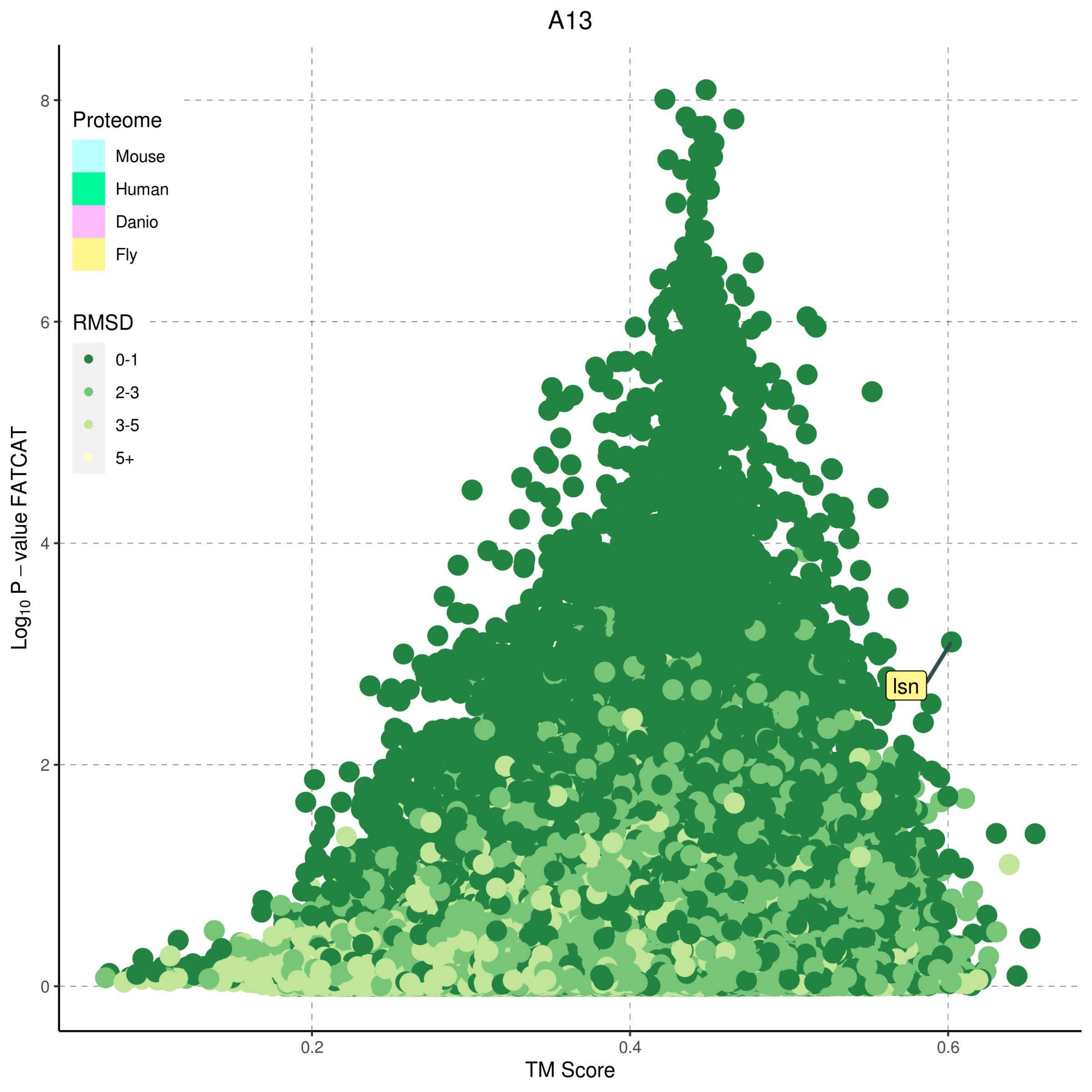
A13
Proteome
Mouse
Human
Danio
Fly
RMSD
0-1
2-3
3-5
5+
Isn
Log10 P − value FATCAT
TM Score
0
2
4
6
8
0.2
0.4
0.6

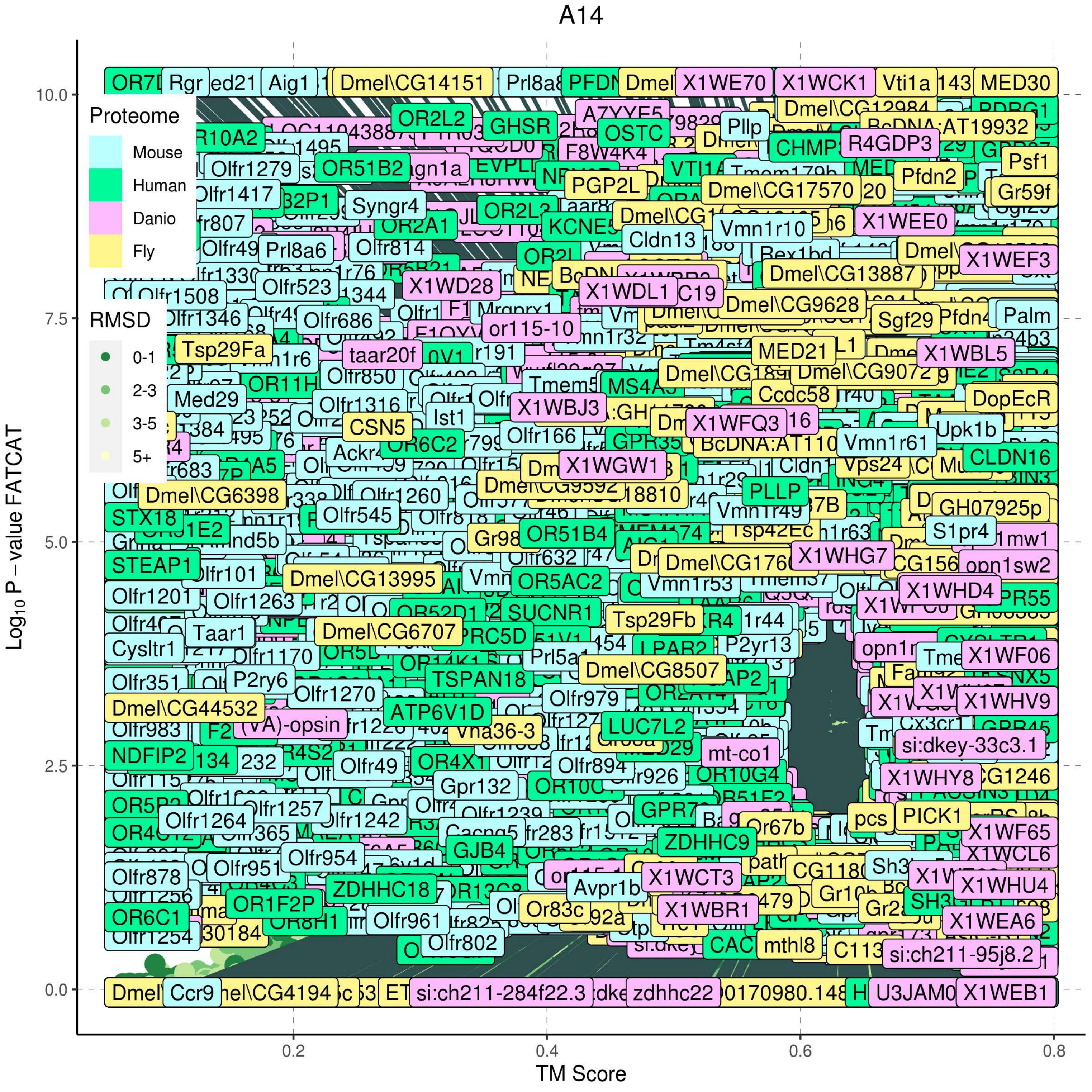
A14
Proteome
Mouse
Human
Danio
Fly
RMSD
0-1
2-3
3-5
5+
Log10 P − value FATCAT
10.0
7.5
5.0
2.5
0.0
TM Score
0.2
0.4
0.6
0.8

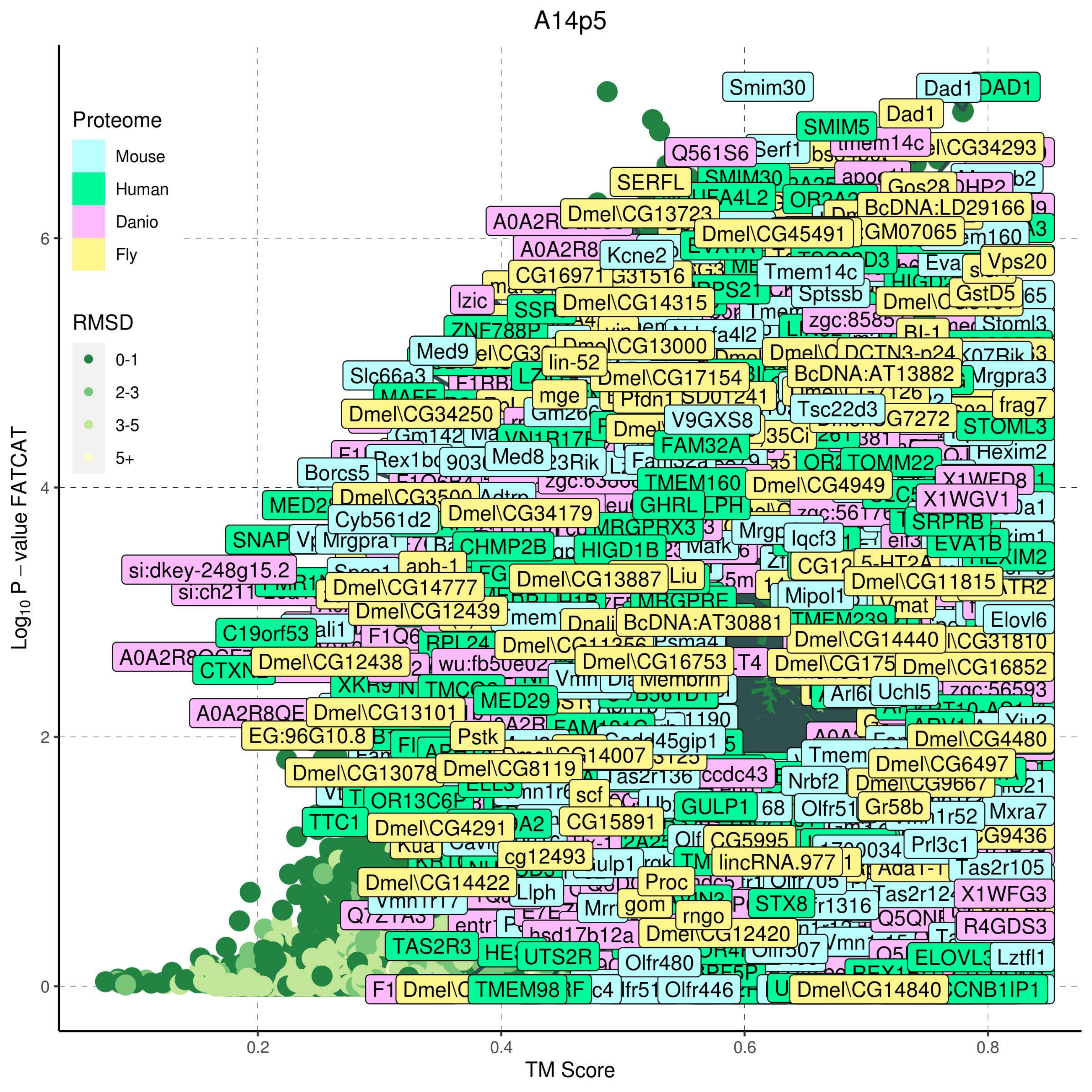

A14p5
Proteome
Mouse
Human
Danio
Fly
RMSD
0-1
2-3
3-5
5+
$Log_{10}$ P – value FATCAT
TM Score
0
2
4
6
0.2
0.4
0.6
0.8

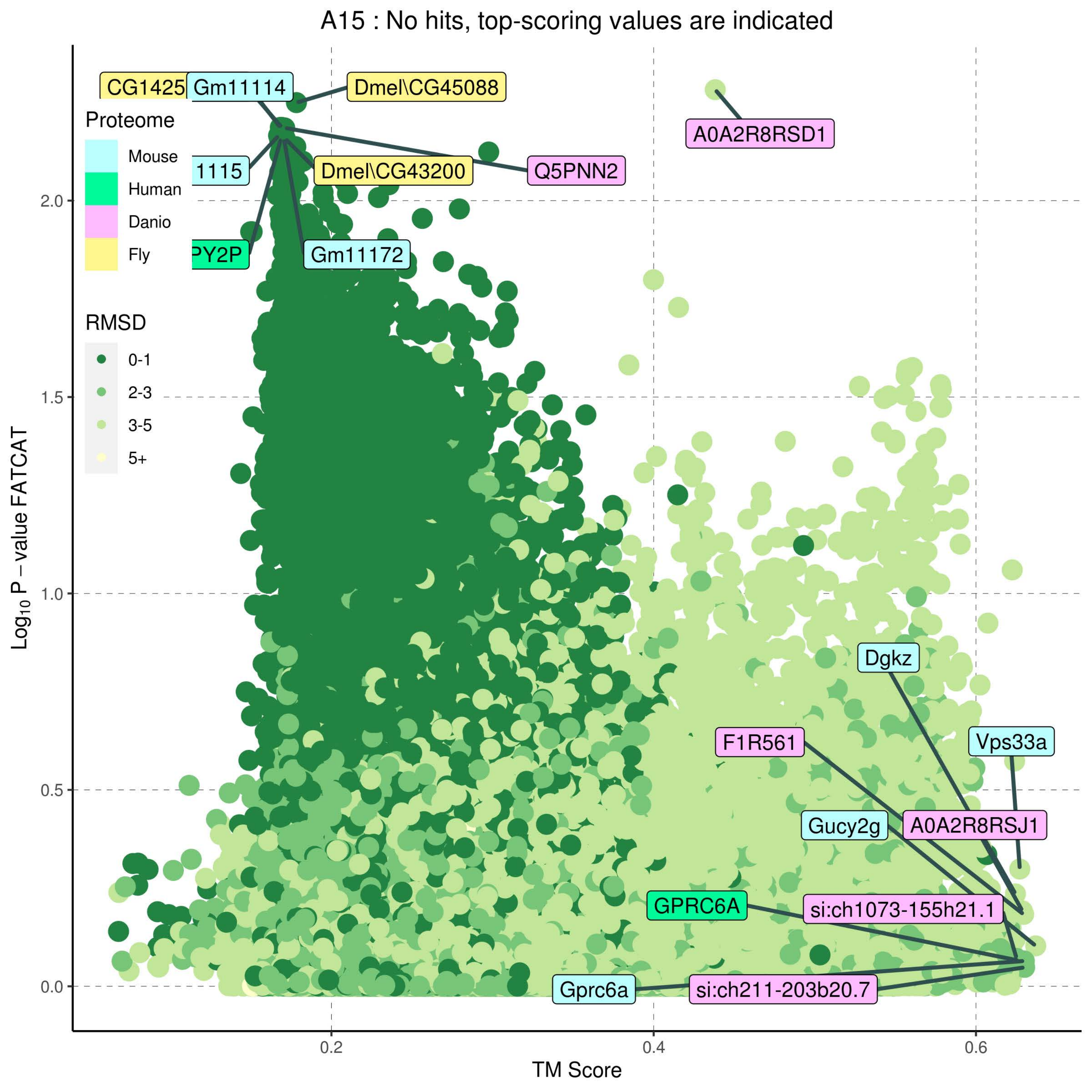

A15 : No hits, top-scoring values are indicated
Proteome
Mouse
Human
Danio
Fly
RMSD
0-1
2-3
3-5
5+
CG1425
Gm11114
Dmel\CG45088
A0A2R8RSD1
1115
Dmel\CG43200
Q5PNN2
PY2P
Gm11172
Dgkz
F1R561
Vps33a
Gucy2g
A0A2R8RSJ1
GPRC6A
si:ch1073-155h21.1
Gprc6a
si:ch211-203b20.7
2.0
1.5
1.0
0.5
0.0
0.2
0.4
0.6
$Log_{10}$ P – value FATCAT
TM Score

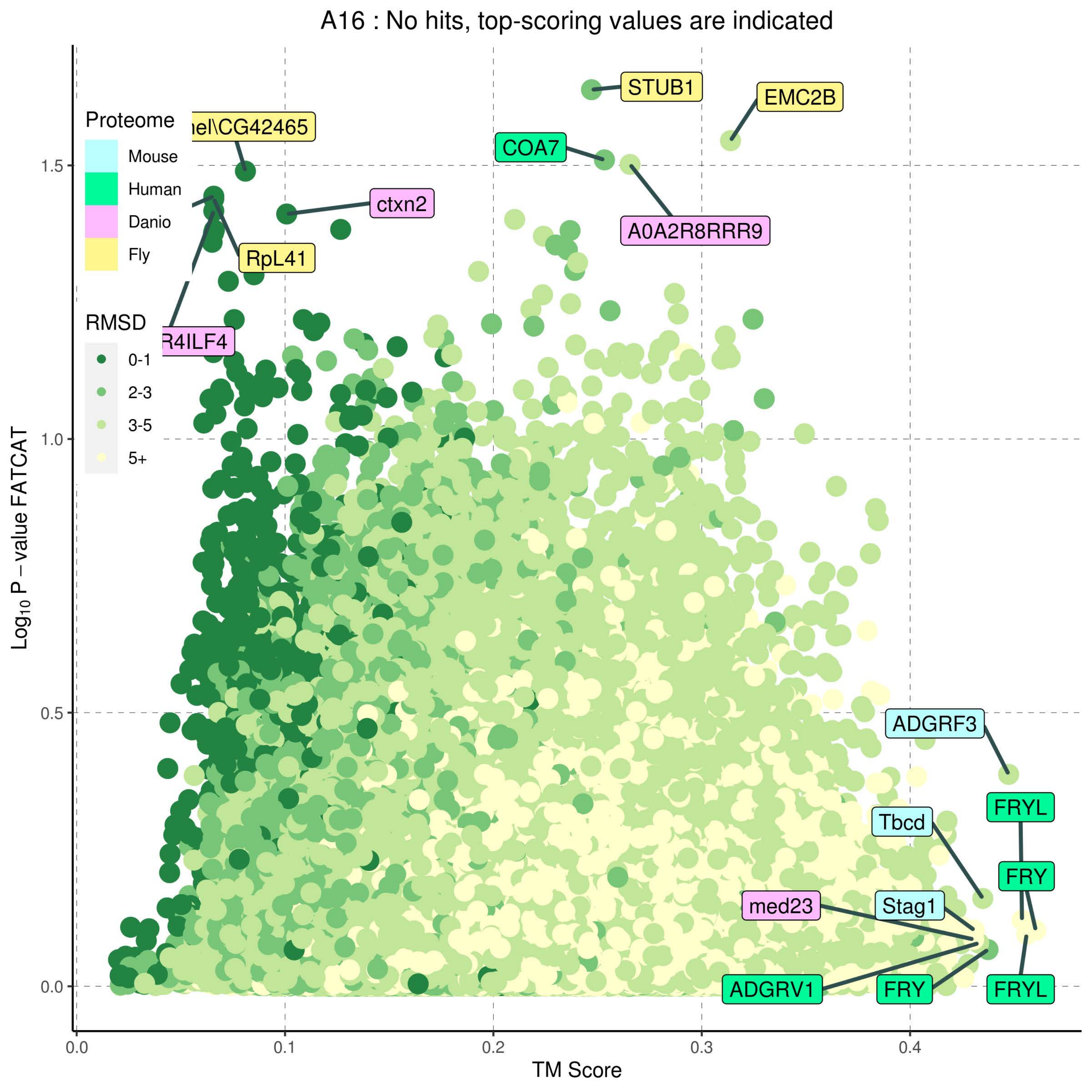

A16 : No hits, top-scoring values are indicated
Proteome
Mouse
Human
Danio
Fly
RMSD
0-1
2-3
3-5
5+
STUB1
EMC2B
el\CG42465
COA7
ctxn2
A0A2R8RRR9
RpL41
R4ILF4
ADGRF3
FRYL
Tbcd
FRY
med23
Stag1
ADGRV1
FRY
FRYL
Log10 P − value FATCAT
TM Score
0.0
0.5
1.0
1.5
0.0
0.1
0.2
0.3
0.4

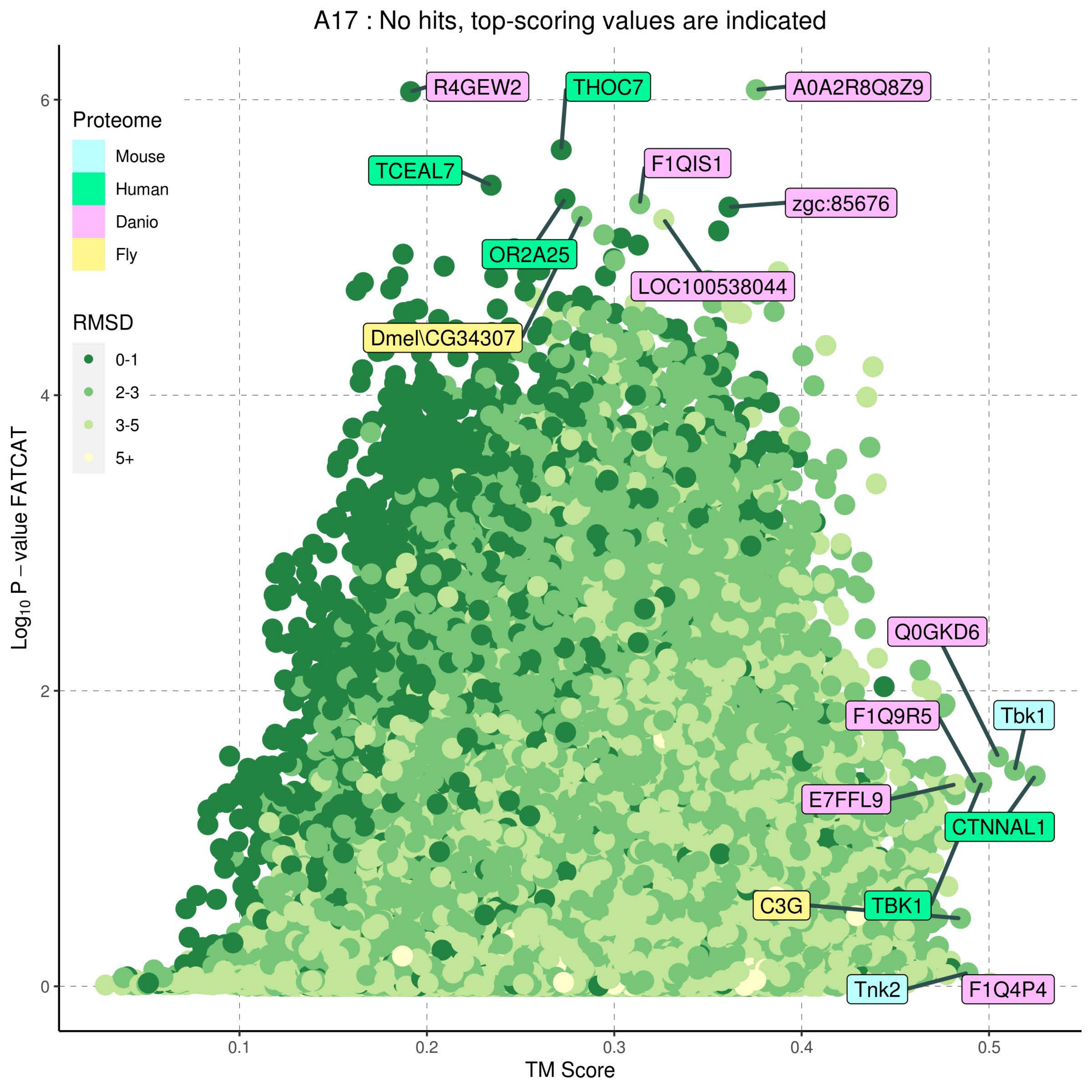
A17 : No hits, top-scoring values are indicated
Proteome
Mouse
Human
Danio
Fly
RMSD
0-1
2-3
3-5
5+
R4GEW2
THOC7
A0A2R8Q8Z9
TCEAL7
F1QIS1
zgc:85676
OR2A25
LOC100538044
Dmel\CG34307
Q0GKD6
F1Q9R5
Tbk1
E7FFL9
CTNNAL1
C3G
TBK1
Tnk2
F1Q4P4
Log10 P − value FATCAT
TM Score
0
2
4
6
0.1
0.2
0.3
0.4
0.5

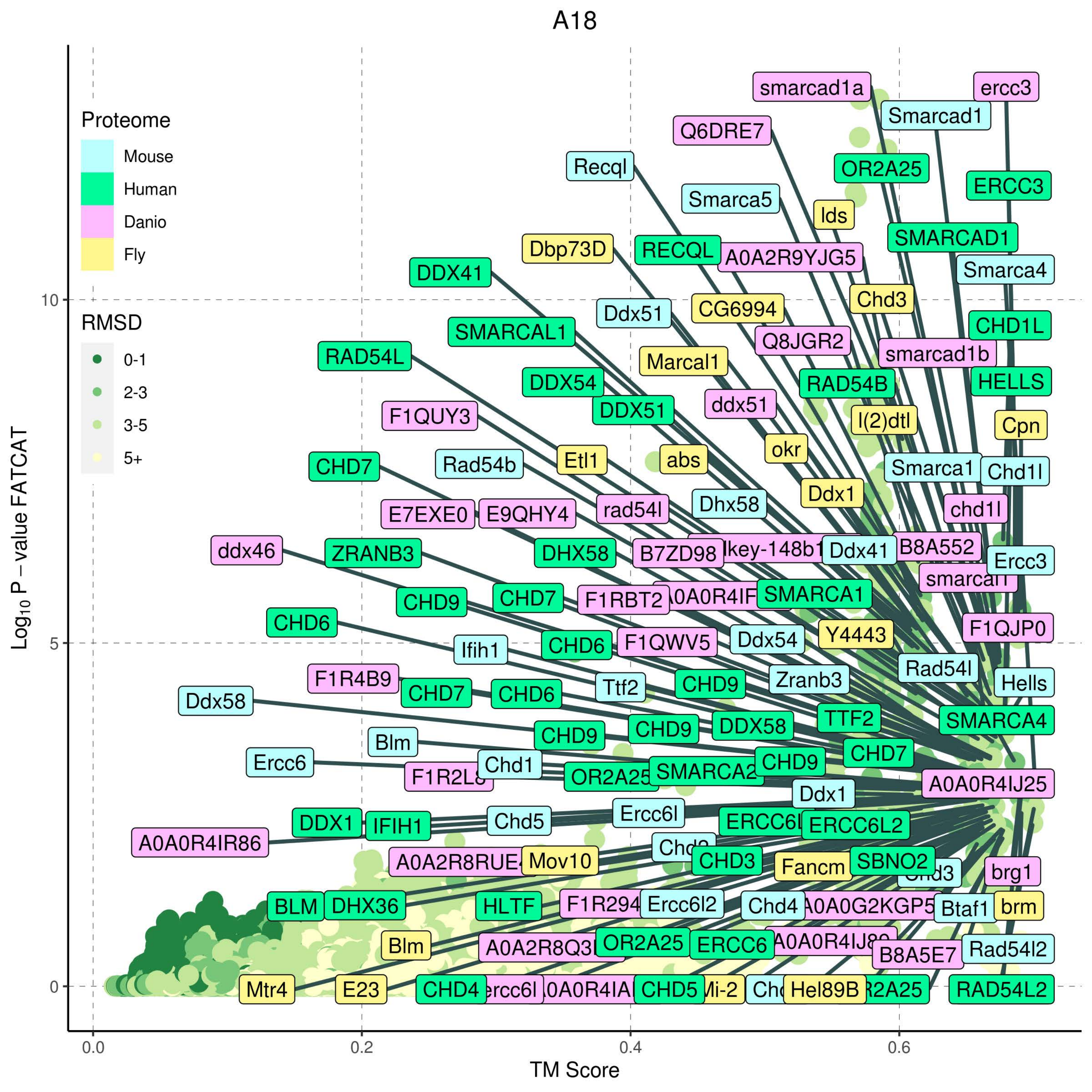

A18
Proteome
Mouse
Human
Danio
Fly
RMSD
0-1
2-3
3-5
5+
Log10 P − value FATCAT
TM Score
0.0
0.2
0.4
0.6
0
5
10
smarcad1a
ercc3
Smarcad1
Q6DRE7
Recql
OR2A25
ERCC3
Smarca5
lds
Dbp73D
RECQL
A0A2R9YJG5
SMARCAD1
DDX41
Smarca4
Chd3
Ddx51
CG6994
CHD1L
SMARCAL1
Q8JGR2
smarcad1b
RAD54L
Marcal1
DDX54
RAD54B
HELLS
ddx51
DDX51
F1QUY3
l(2)dtl
Cpn
okr
CHD7
Rad54b
Etl1
abs
Smarca1
Chd1l
Ddx1
Dhx58
E7EXE0
E9QHY4
rad54l
chd1l
ddx46
ZRANB3
DHX58
B7ZD98
Ddx41
B8A552
Ercc3
CHD9
CHD7
F1RBT2
SMARCA1
CHD6
F1QJP0
Ifih1
CHD6
F1QWV5
Ddx54
Y4443
Rad54l
F1R4B9
CHD7
CHD6
Ttf2
CHD9
Zranb3
Hells
Ddx58
TTF2
SMARCA4
CHD9
CHD9
DDX58
Blm
CHD7
Ercc6
Chd1
CHD9
A0A0R4IJ25
SMARCA2
Ddx1
Ercc6l
DDX1
IFIH1
Chd5
ERCC6L2
A0A0R4IR86
Mov10
CHD3
Fancm
SBNO2
brg1
BLM
DHX36
HLTF
Ercc6l2
Chd4
Btaf1
brm
Blm
OR2A25
ERCC6
B8A5E7
Rad54l2
Mtr4
E23
CHD4
CHD5
Hel89B
RAD54L2

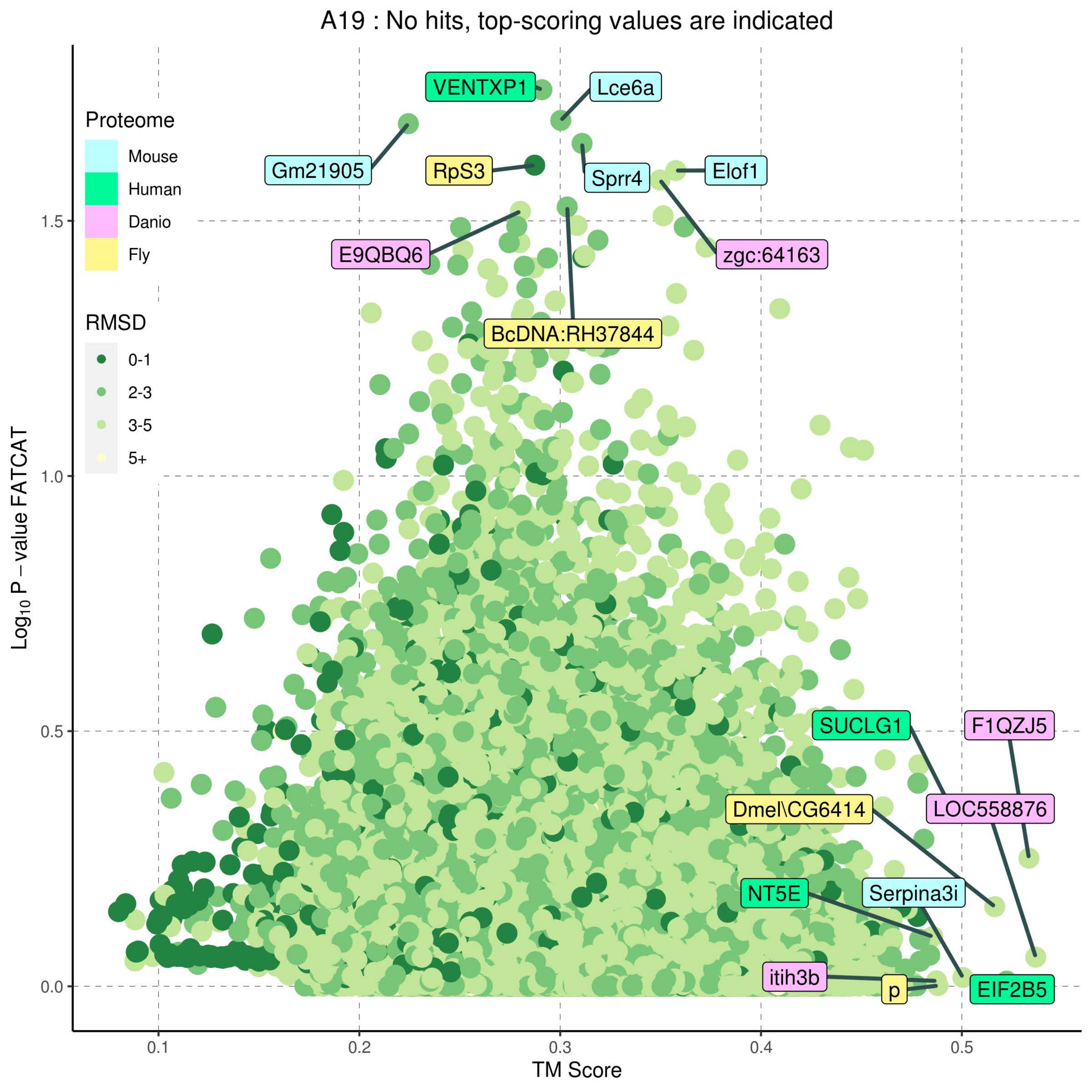

A19 : No hits, top-scoring values are indicated
Proteome
Mouse
Human
Danio
Fly
RMSD
0-1
2-3
3-5
5+
VENTXP1
Lce6a
Gm21905
RpS3
Sprr4
Elof1
E9QBQ6
zgc:64163
BcDNA:RH37844
SUCLG1
F1QZJ5
Dmel\CG6414
LOC558876
NT5E
Serpina3i
itih3b
p
EIF2B5
Log10 P − value FATCAT
TM Score
0.0
0.5
1.0
1.5
0.1
0.2
0.3
0.4
0.5

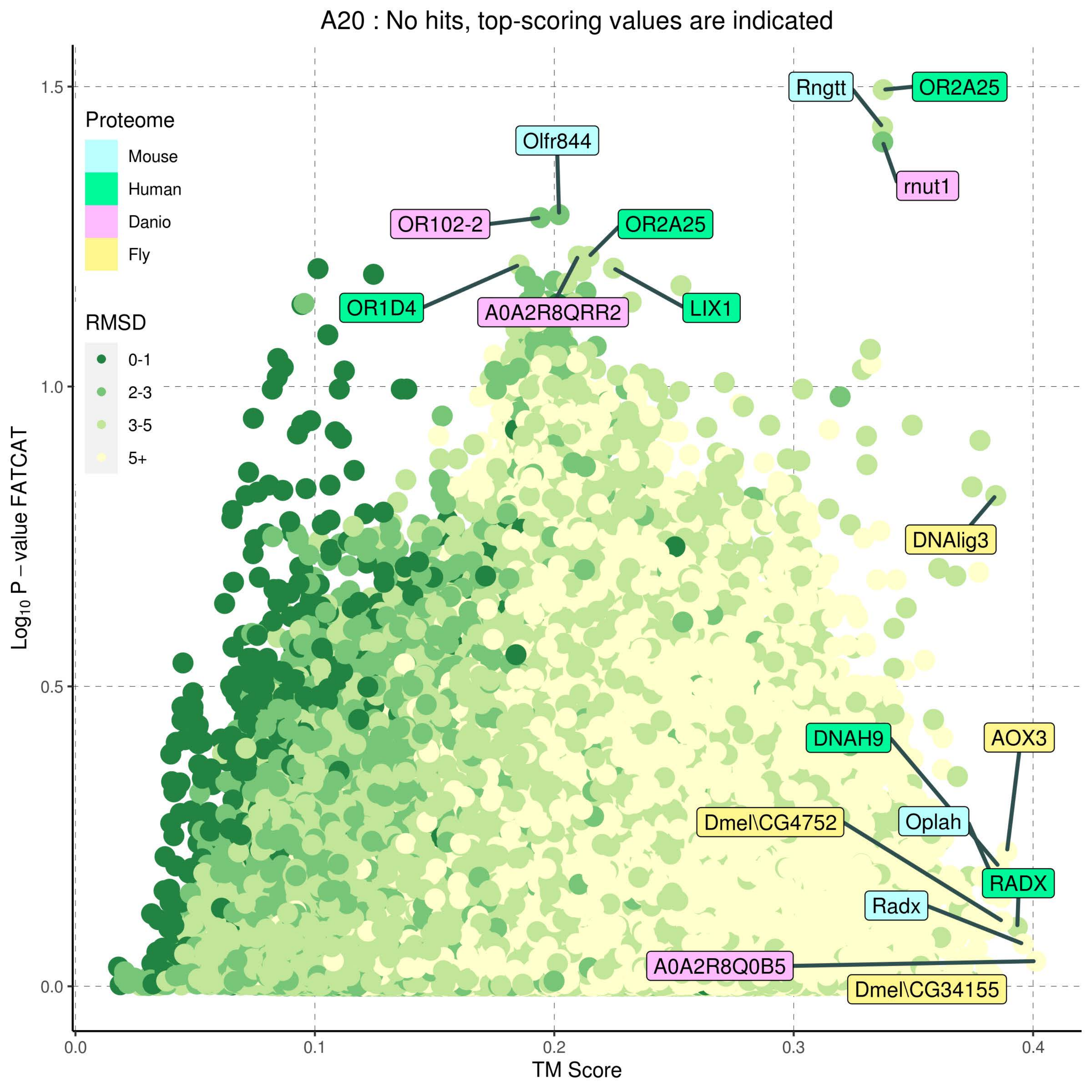
A20 : No hits, top-scoring values are indicated
Proteome
Mouse
Human
Danio
Fly
RMSD
0-1
2-3
3-5
5+
Rngtt
OR2A25
Olfr844
rnut1
OR102-2
OR2A25
OR1D4
A0A2R8QRR2
LIX1
DNAlig3
DNAH9
AOX3
Oplah
Dmel\CG4752
RADX
Radx
A0A2R8Q0B5
Dmel\CG34155
TM Score
$Log_{10}$ P − value FATCAT
0.0
0.1
0.2
0.3
0.4
0.0
0.5
1.0
1.5

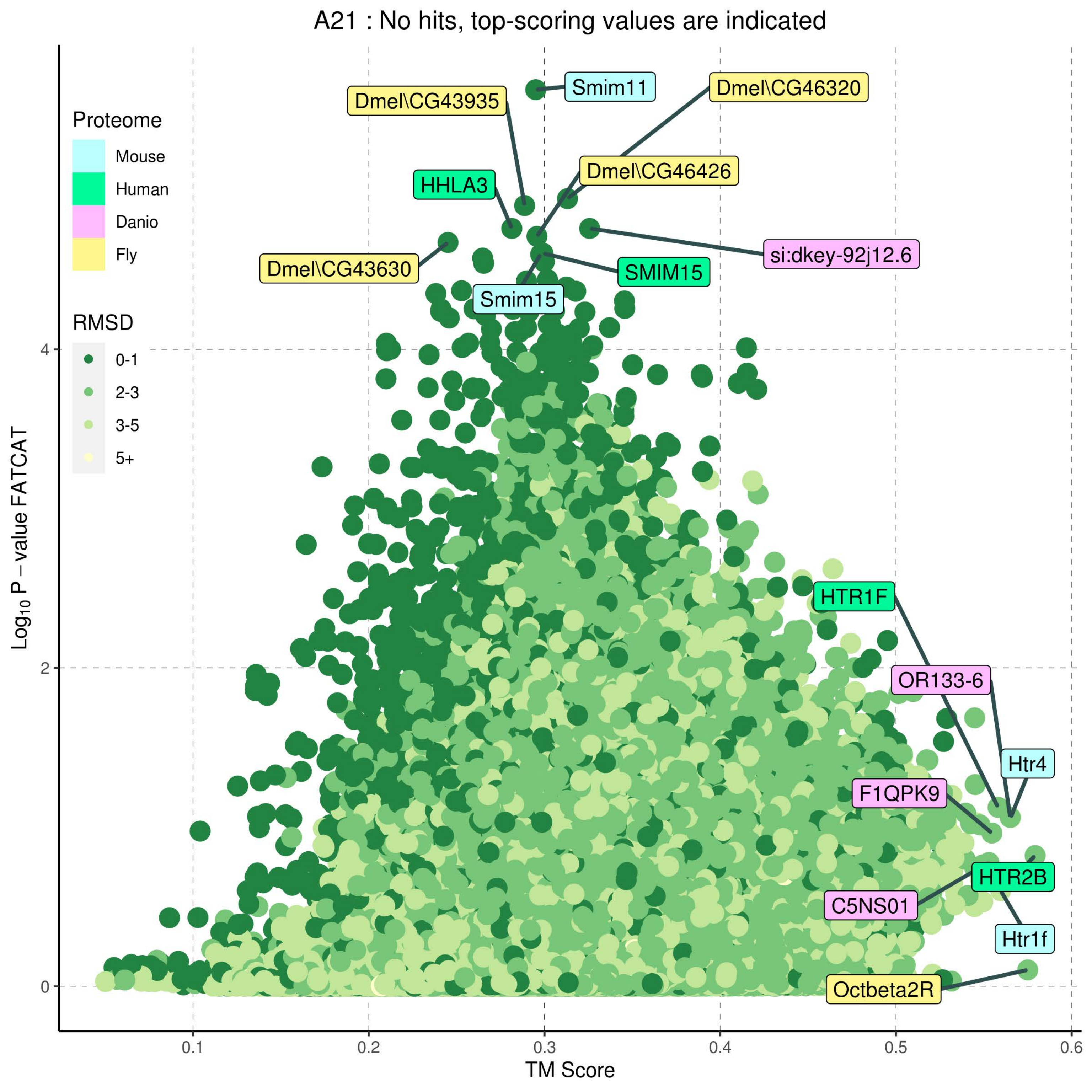

A21 : No hits, top-scoring values are indicated
Proteome
Mouse
Human
Danio
Fly
RMSD
0-1
2-3
3-5
5+
Smim11
Dmel\CG43935
Dmel\CG46320
Dmel\CG46426
HHLA3
si:dkey-92j12.6
Dmel\CG43630
SMIM15
Smim15
HTR1F
OR133-6
Htr4
F1QPK9
HTR2B
C5NS01
Htr1f
Octbeta2R
Log10 P − value FATCAT
TM Score
0
2
4
0.1
0.2
0.3
0.4
0.5
0.6

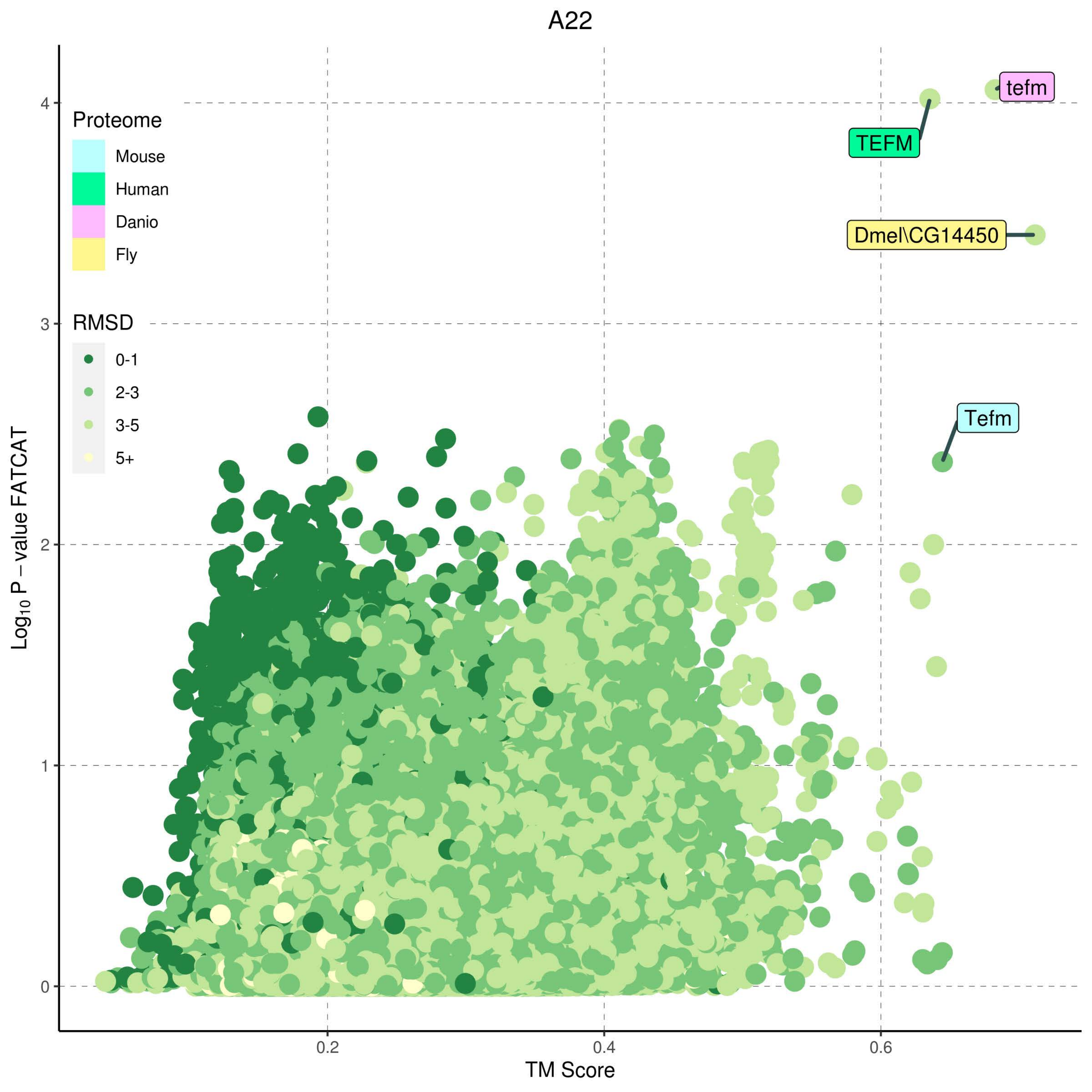

A22
Proteome
Mouse
Human
Danio
Fly
RMSD
0-1
2-3
3-5
5+
tefm
TEFM
Dmel\CG14450
Tefm
Log10 P − value FATCAT
TM Score
0
1
2
3
4
0.2
0.4
0.6

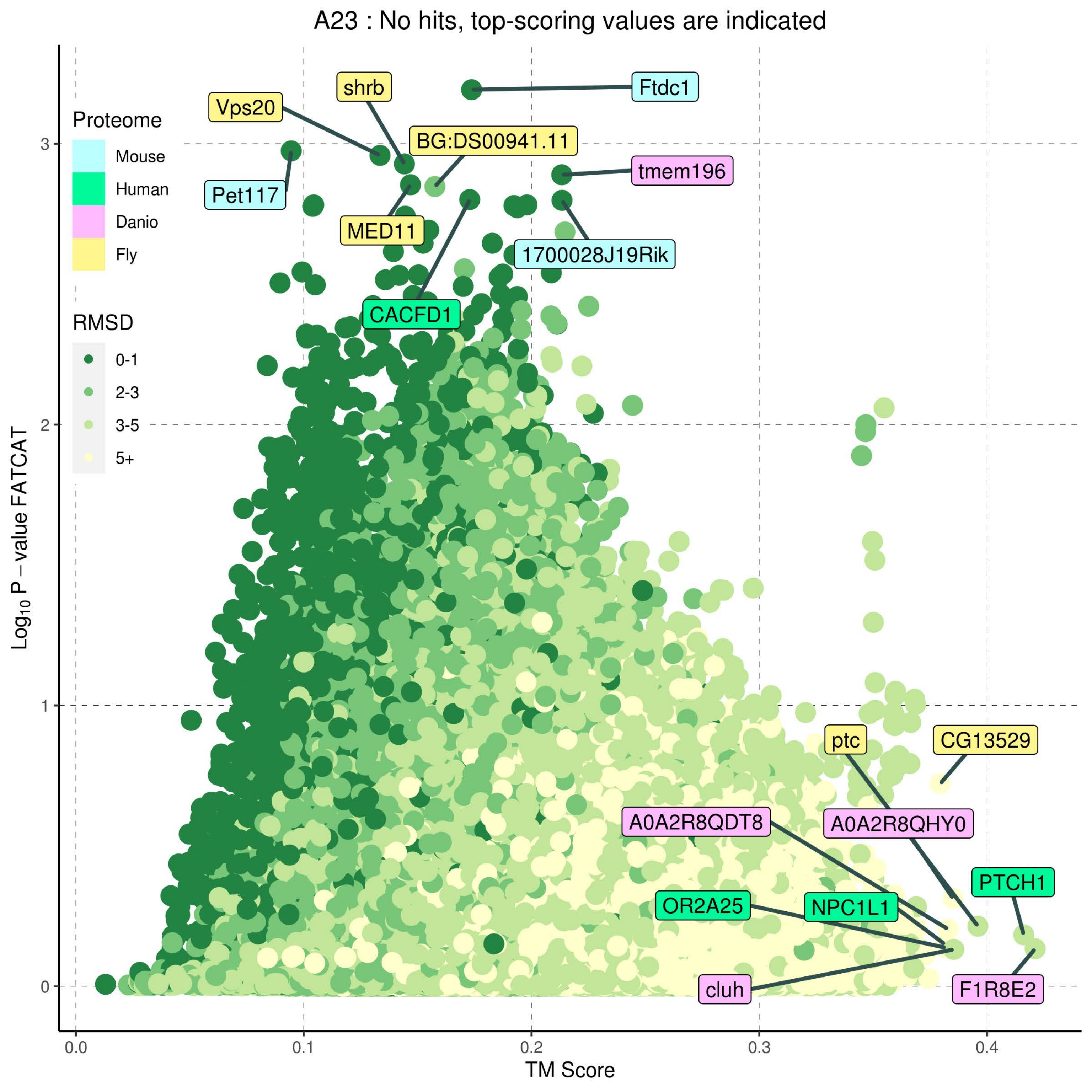
A23 : No hits, top-scoring values are indicated
Proteome
Mouse
Human
Danio
Fly
RMSD
0-1
2-3
3-5
5+
Ftdc1
shrb
Vps20
BG:DS00941.11
tmem196
Pet117
MED11
1700028J19Rik
CACFD1
ptc
CG13529
A0A2R8QDT8
A0A2R8QHY0
PTCH1
OR2A25
NPC1L1
cluh
F1R8E2
Log10 P − value FATCAT
TM Score
0.0
0.1
0.2
0.3
0.4
0
1
2
3

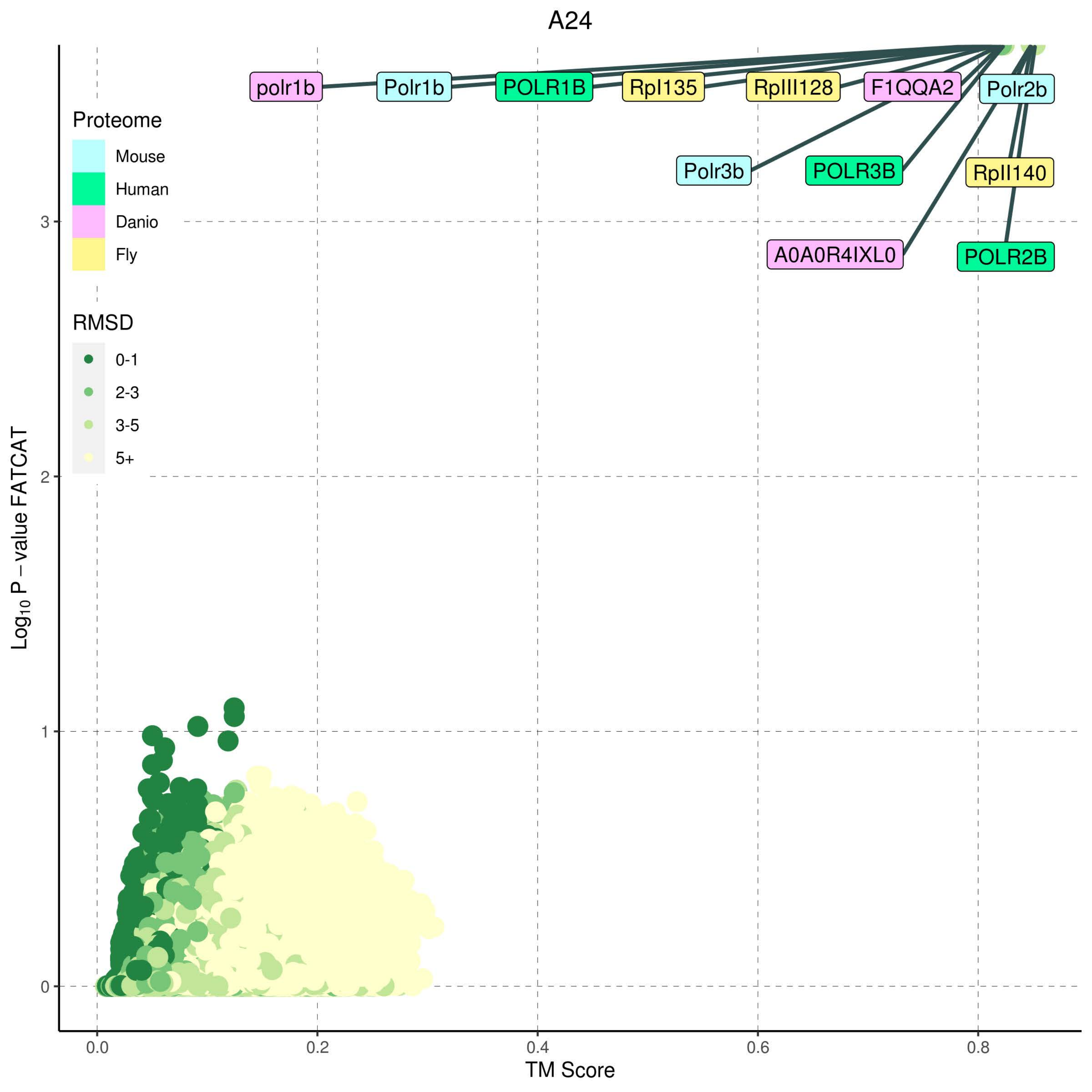
A24
polr1b
Polr1b
POLR1B
Rpl135
RpIII128
F1QQA2
Polr2b
Polr3b
POLR3B
RpII140
A0A0R4IXL0
POLR2B
Proteome
Mouse
Human
Danio
Fly
RMSD
0-1
2-3
3-5
5+
Log10 P − value FATCAT
0
1
2
3
0.0
0.2
0.4
0.6
0.8
TM Score

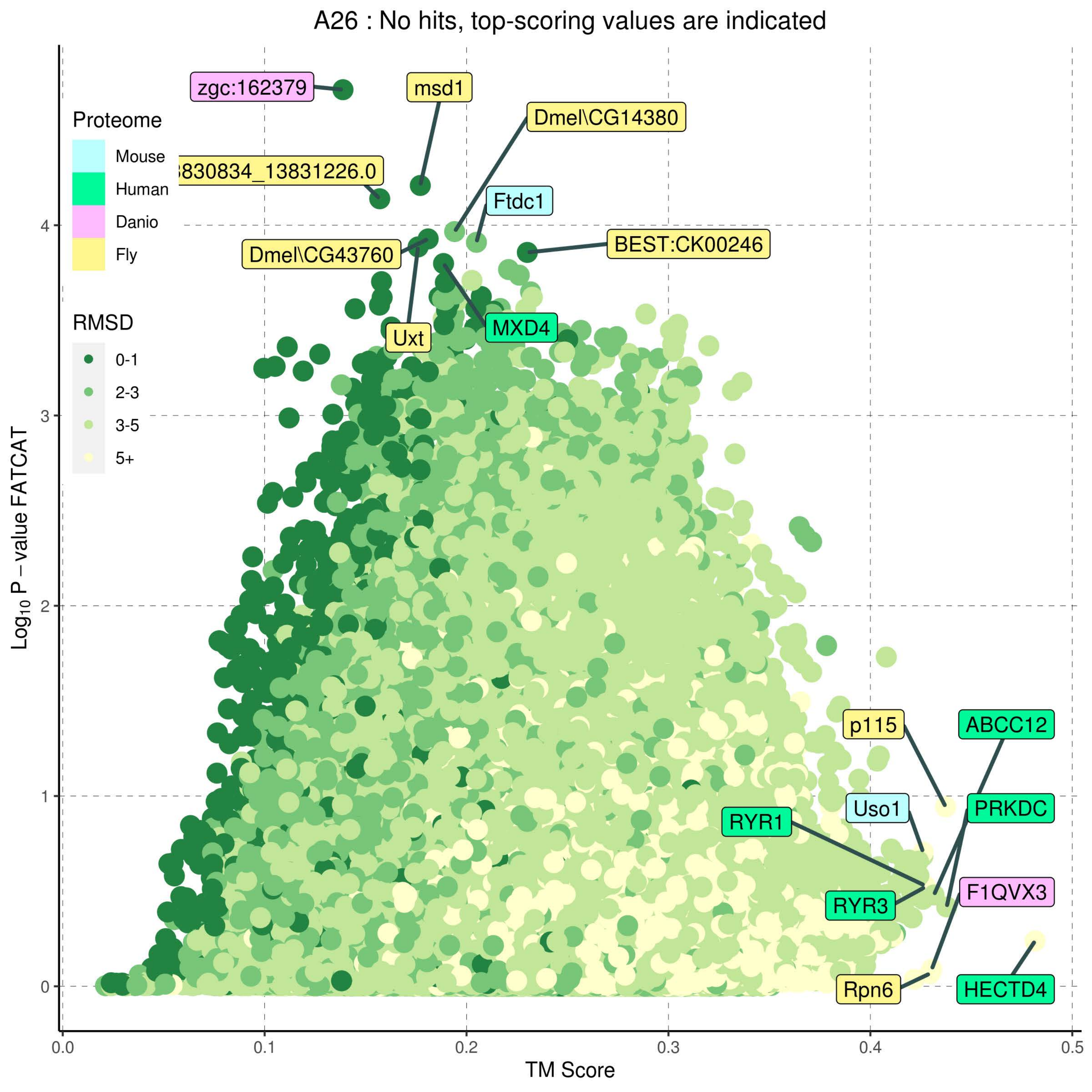
A26 : No hits, top-scoring values are indicated
Proteome
Mouse
Human
Danio
Fly
RMSD
0-1
2-3
3-5
5+
zgc:162379
msd1
Dmel\CG14380
830834_13831226.0
Ftdc1
Dmel\CG43760
BEST:CK00246
MXD4
Uxt
p115
ABCC12
Uso1
PRKDC
RYR1
RYR3
F1QVX3
Rpn6
HECTD4
Log10 P – value FATCAT
TM Score
0
1
2
3
4
0.0
0.1
0.2
0.3
0.4
0.5

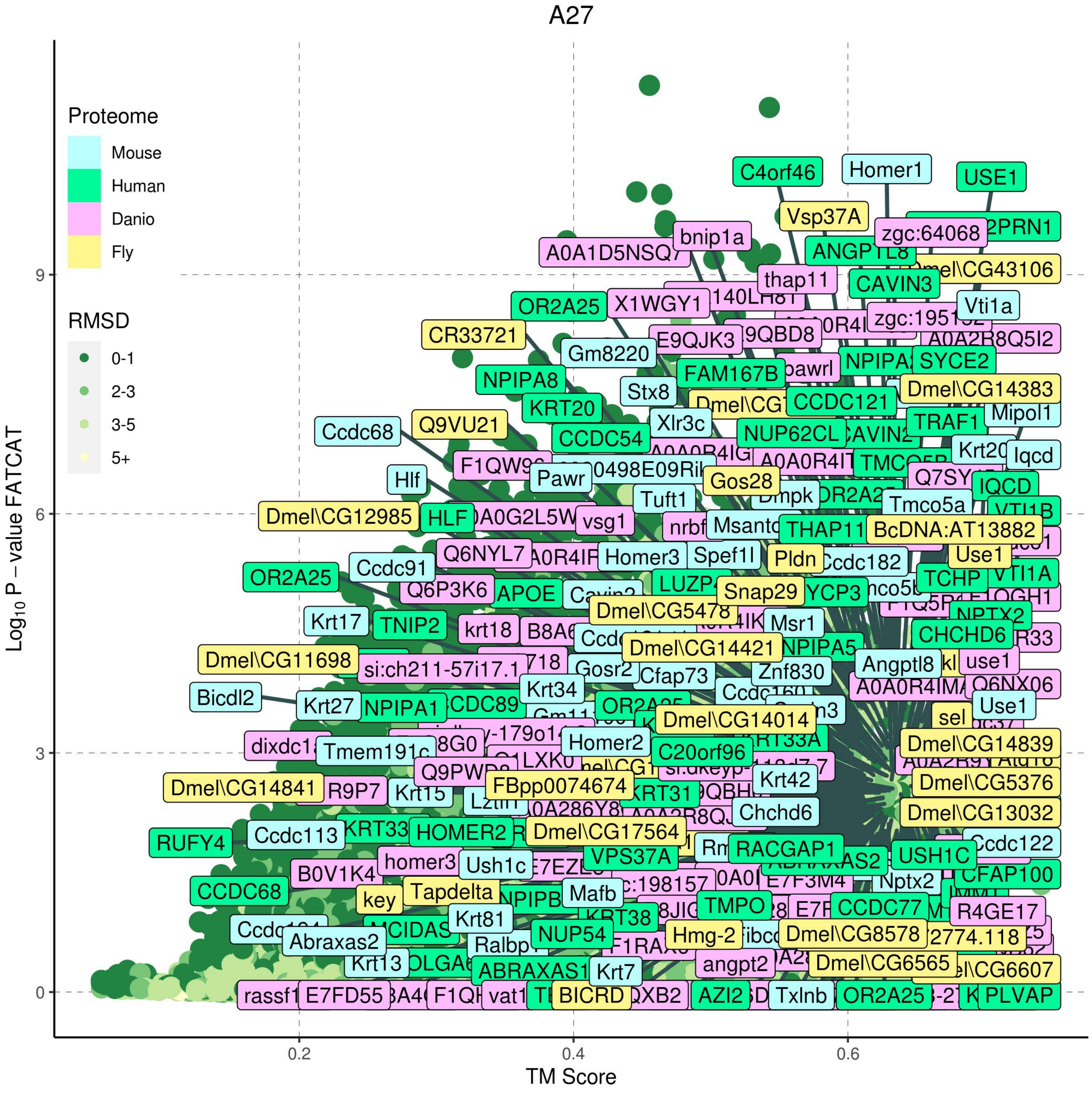

A27
Proteome
Mouse
Human
Danio
Fly
RMSD
0-1
2-3
3-5
5+
Log10 P − value FATCAT
TM Score
0
3
6
9
0.2
0.4
0.6
C4orf46
Homer1
USE1
Vsp37A
bnip1a
zgc:64068
A0A1D5NSQ7
thap11
CAVIN3
OR2A25
X1WGY1
Vti1a
CR33721
E9QJK3
A0A2R8Q5I2
Gm8220
NPIPA8
FAM167B
SYCE2
Stx8
Dmel\CG14383
KRT20
CCDC121
Mipol1
Ccdc68
Q9VU21
Xlr3c
NUP62CL
TRAF1
CCDC54
Iqcd
Pawr
Gos28
IQCD
Hlf
Tuft1
Tmco5a
Dmel\CG12985
HLF
vsg1
THAP11
BcDNA:AT13882
Q6NYL7
Homer3
Spef1l
Pldn
Use1
OR2A25
Ccdc91
Ccdc182
TCHP
VTI1A
Q6P3K6
APOE
Snap29
Dmel\CG5478
Krt17
TNIP2
krt18
Msr1
CHCHD6
Dmel\CG14421
Dmel\CG11698
si:ch211-57i17.1
Gosr2
Cfap73
Znf830
Angptl8
use1
Q6NX06
Bicdl2
Krt27
NPIPA1
Krt34
Dmel\CG14014
Use1
Homer2
C20orf96
Dmel\CG14839
Q9PWD2
FBpp0074674
Krt42
Dmel\CG5376
Dmel\CG14841
R9P7
Krt15
Chchd6
Dmel\CG13032
RUFY4
Ccdc113
KRT33
HOMER2
Dmel\CG17564
RACGAP1
Ccdc122
homer3
Ush1c
VPS37A
USH1C
B0V1K4
CCDC68
Tapdelta
Mafb
Nptx2
CFAP100
key
Krt81
TMPO
CCDC77
R4GE17
NUP54
Hmg-2
Dmel\CG8578
Abraxas2
Ralbp1
Krt13
ABRAXAS1
Krt7
angpt2
Dmel\CG6565
rassf1
E7FD55
BICRD
AZI2
Txlnb
OR2A25
PLVAP

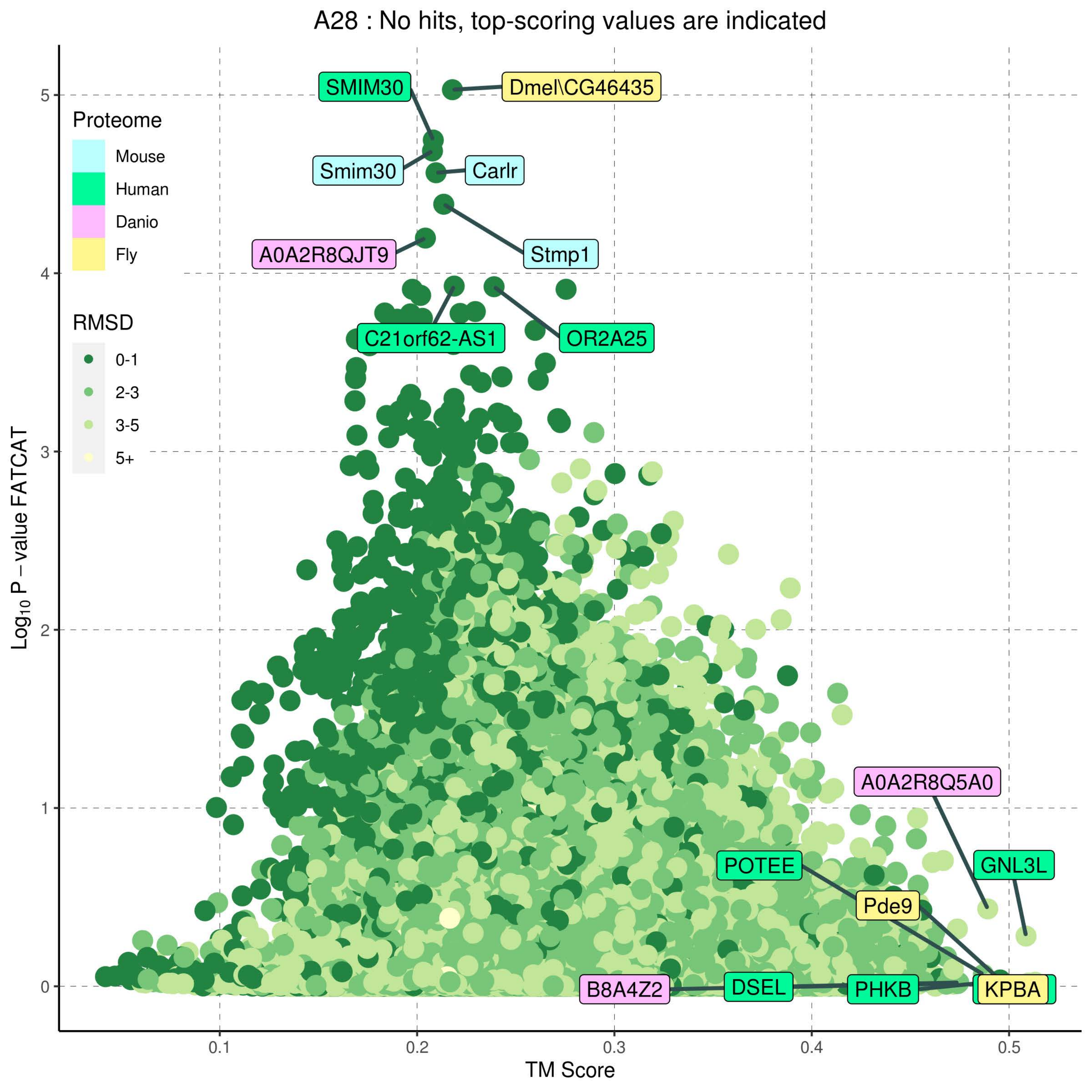
A28 : No hits, top-scoring values are indicated
Proteome
Mouse
Human
Danio
Fly
RMSD
0-1
2-3
3-5
5+
SMIM30
Dmel\CG46435
Smim30
Carlr
A0A2R8QJT9
Stmp1
C21orf62-AS1
OR2A25
A0A2R8Q5A0
POTEE
GNL3L
Pde9
B8A4Z2
DSEL
PHKB
KPBA
Log10 P − value FATCAT
TM Score
0
1
2
3
4
5
0.1
0.2
0.3
0.4
0.5

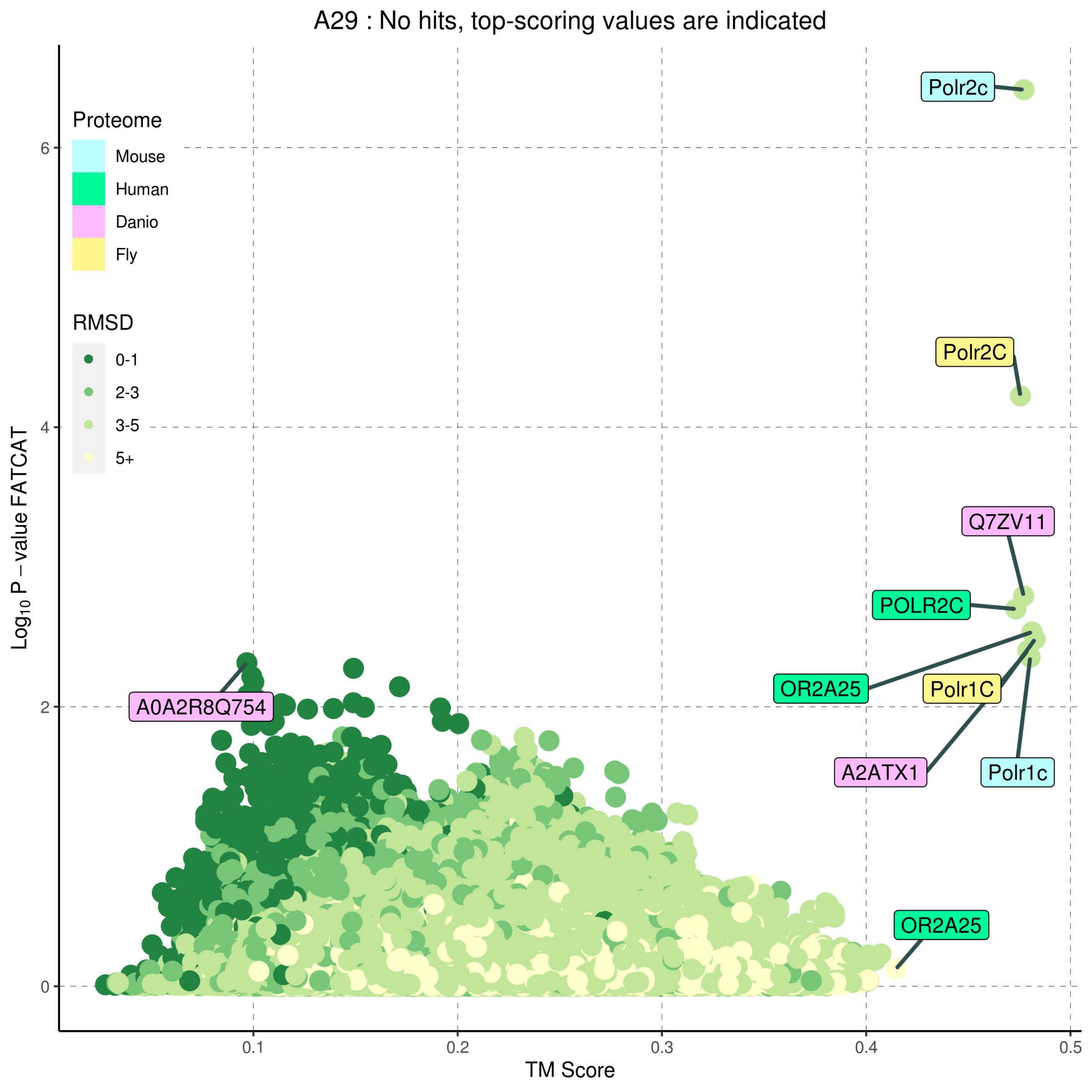
A29 : No hits, top-scoring values are indicated
Proteome
Mouse
Human
Danio
Fly
RMSD
0-1
2-3
3-5
5+
Polr2c
Polr2C
Q7ZV11
POLR2C
OR2A25
Polr1C
A2ATX1
Polr1c
A0A2R8Q754
OR2A25
0
2
4
6
0.1
0.2
0.3
0.4
0.5
TM Score
Log10 P − value FATCAT

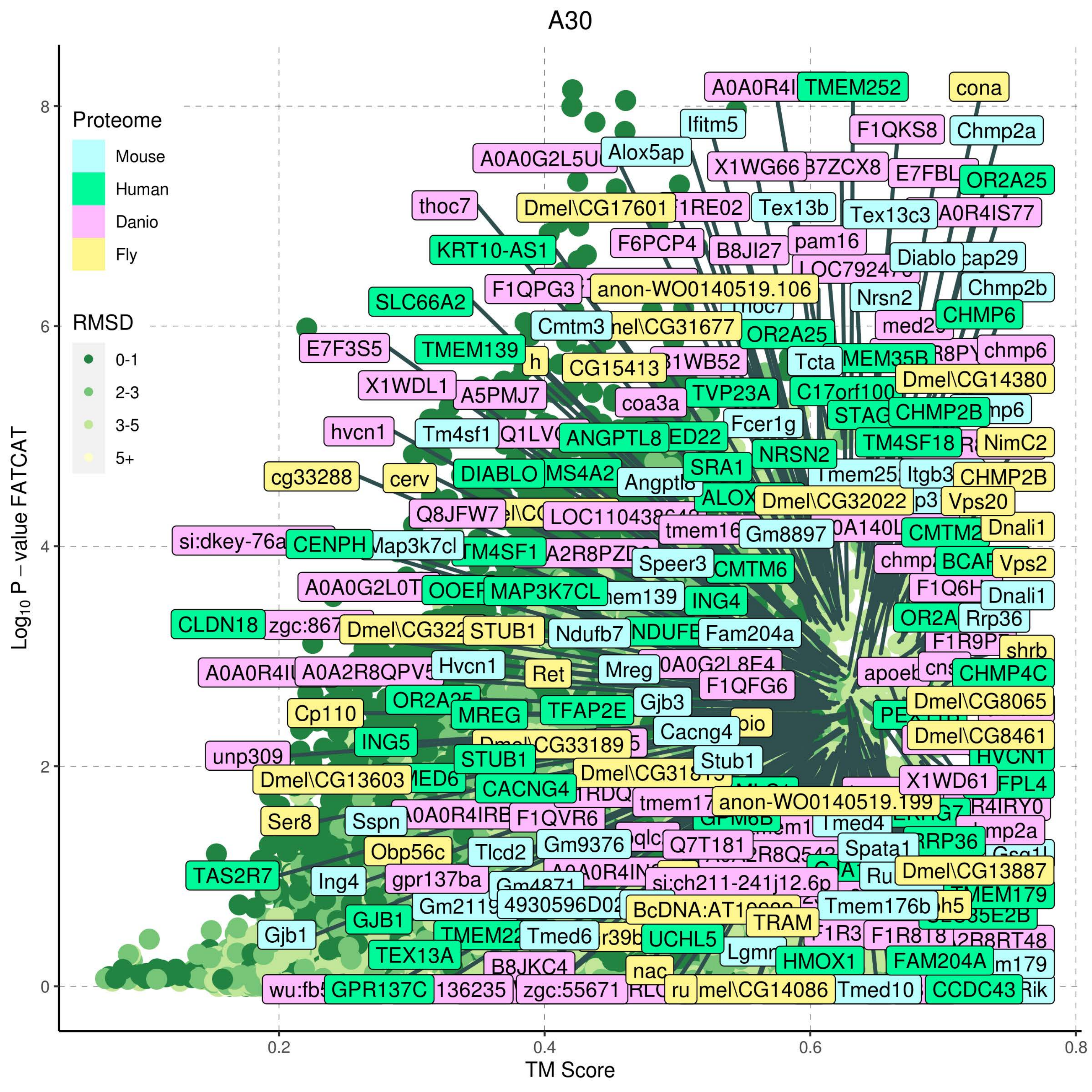

A30
Proteome
Mouse
Human
Danio
Fly
RMSD
0-1
2-3
3-5
5+
Log10 P – value FATCAT
TM Score
0
2
4
6
8
0.2
0.4
0.6
0.8

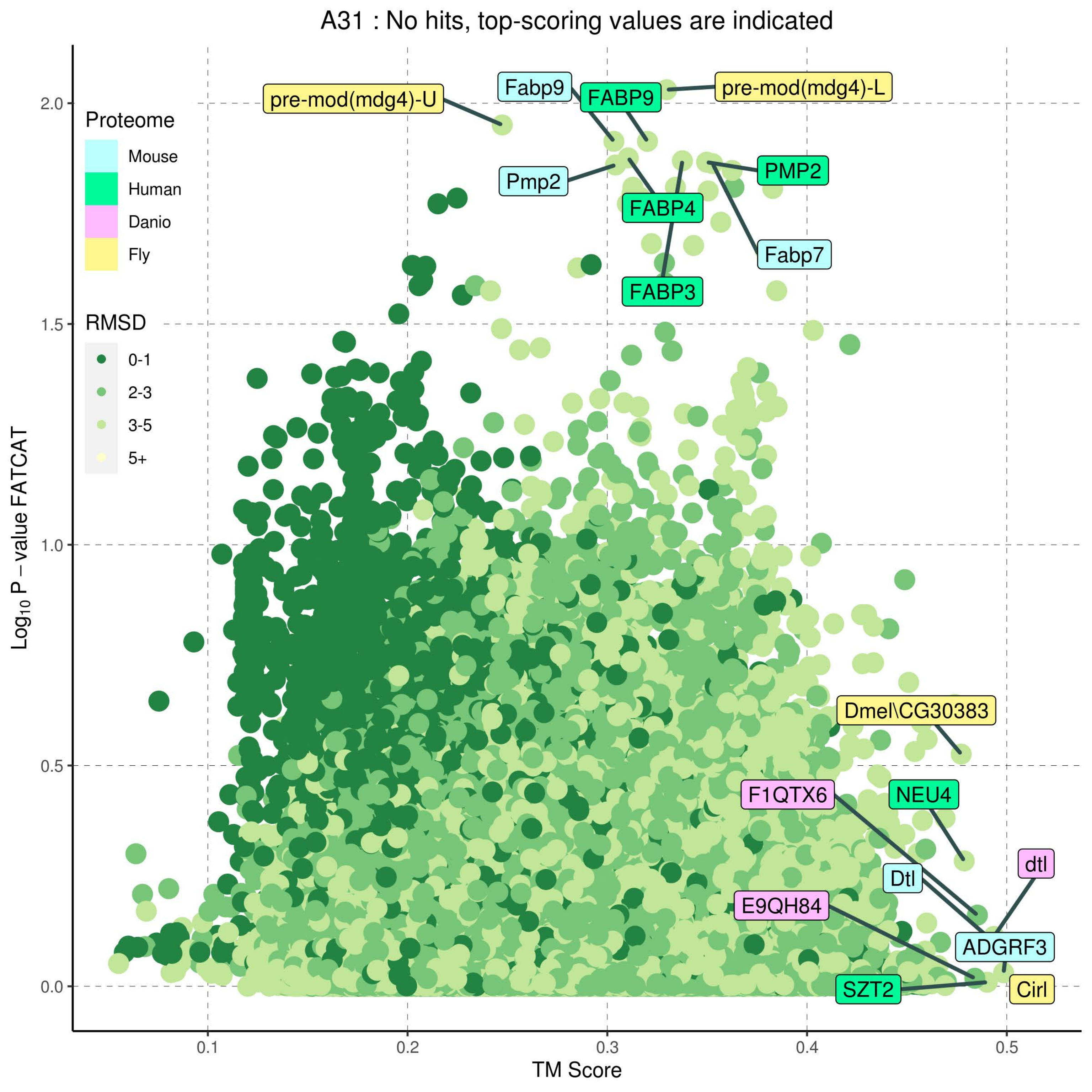

A31 : No hits, top-scoring values are indicated
Proteome
Mouse
Human
Danio
Fly
RMSD
0-1
2-3
3-5
5+
Log10 P – value FATCAT
TM Score
0.0
0.5
1.0
1.5
2.0
0.1
0.2
0.3
0.4
0.5
pre-mod(mdg4)-U
Fabp9
FABP9
pre-mod(mdg4)-L
Pmp2
PMP2
FABP4
Fabp7
FABP3
Dmel\CG30383
F1QTX6
NEU4
dtl
Dtl
E9QH84
ADGRF3
SZT2
Cirl

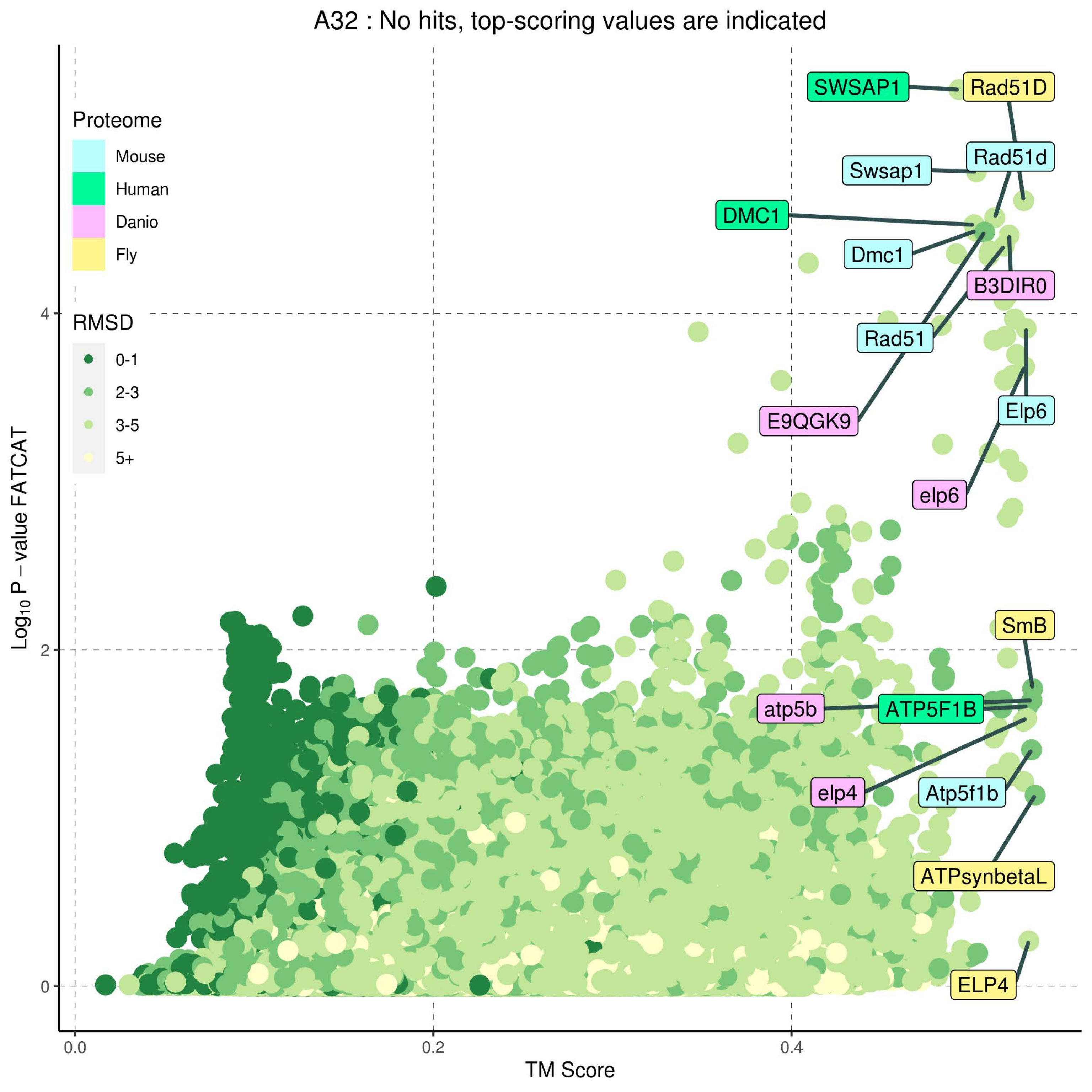

A32 : No hits, top-scoring values are indicated
Proteome
Mouse
Human
Danio
Fly
RMSD
0-1
2-3
3-5
5+
SWSAP1
Rad51D
Swsap1
Rad51d
DMC1
Dmc1
B3DIR0
Rad51
E9QGK9
Elp6
elp6
SmB
atp5b
ATP5F1B
elp4
Atp5f1b
ATPsynbetaL
ELP4
Log10 P − value FATCAT
TM Score
0.0
0.2
0.4
0
2
4

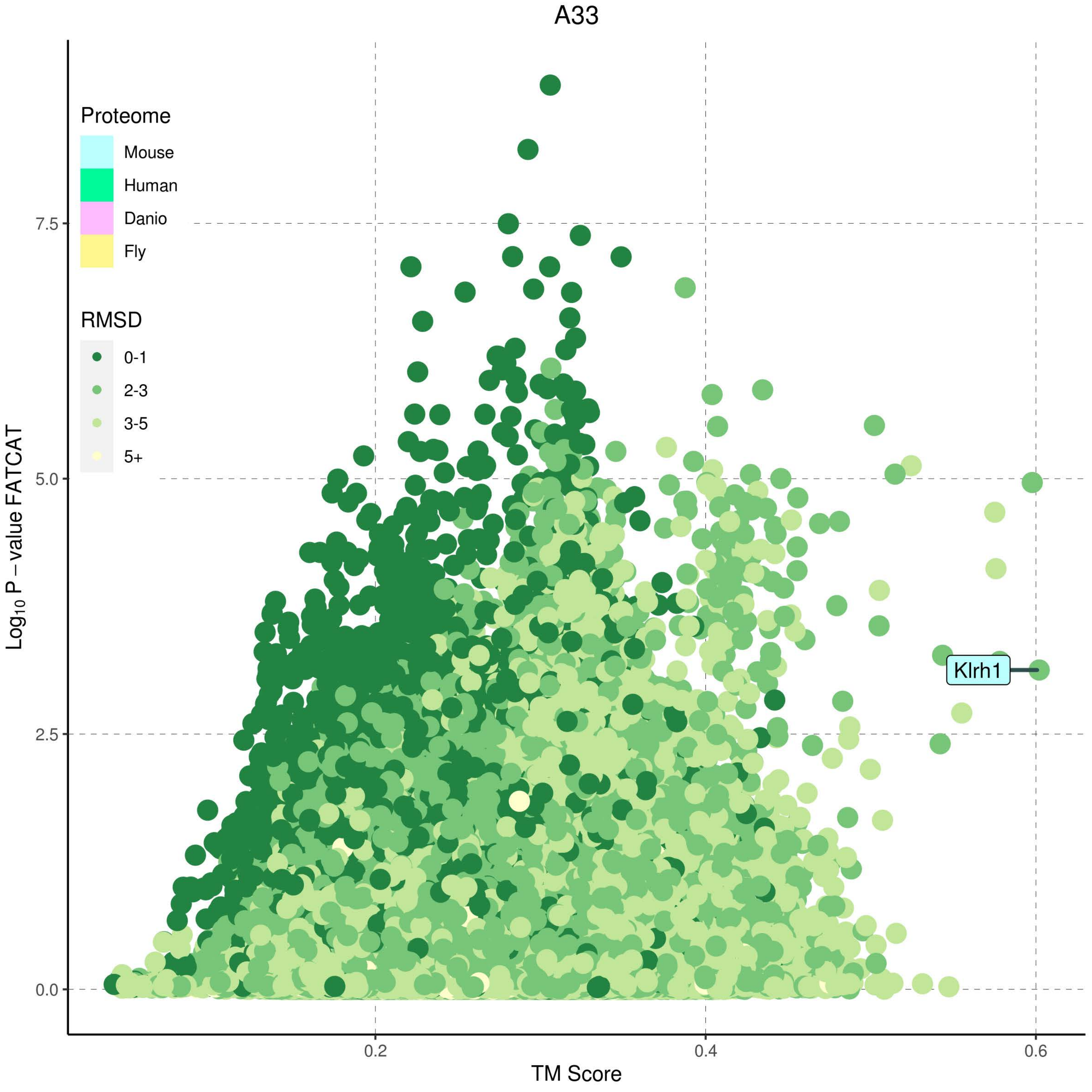
A33
Proteome
Mouse
Human
Danio
Fly
RMSD
0-1
2-3
3-5
5+
Log10 P − value FATCAT
7.5
5.0
2.5
0.0
0.2
0.4
0.6
TM Score
Klrh1

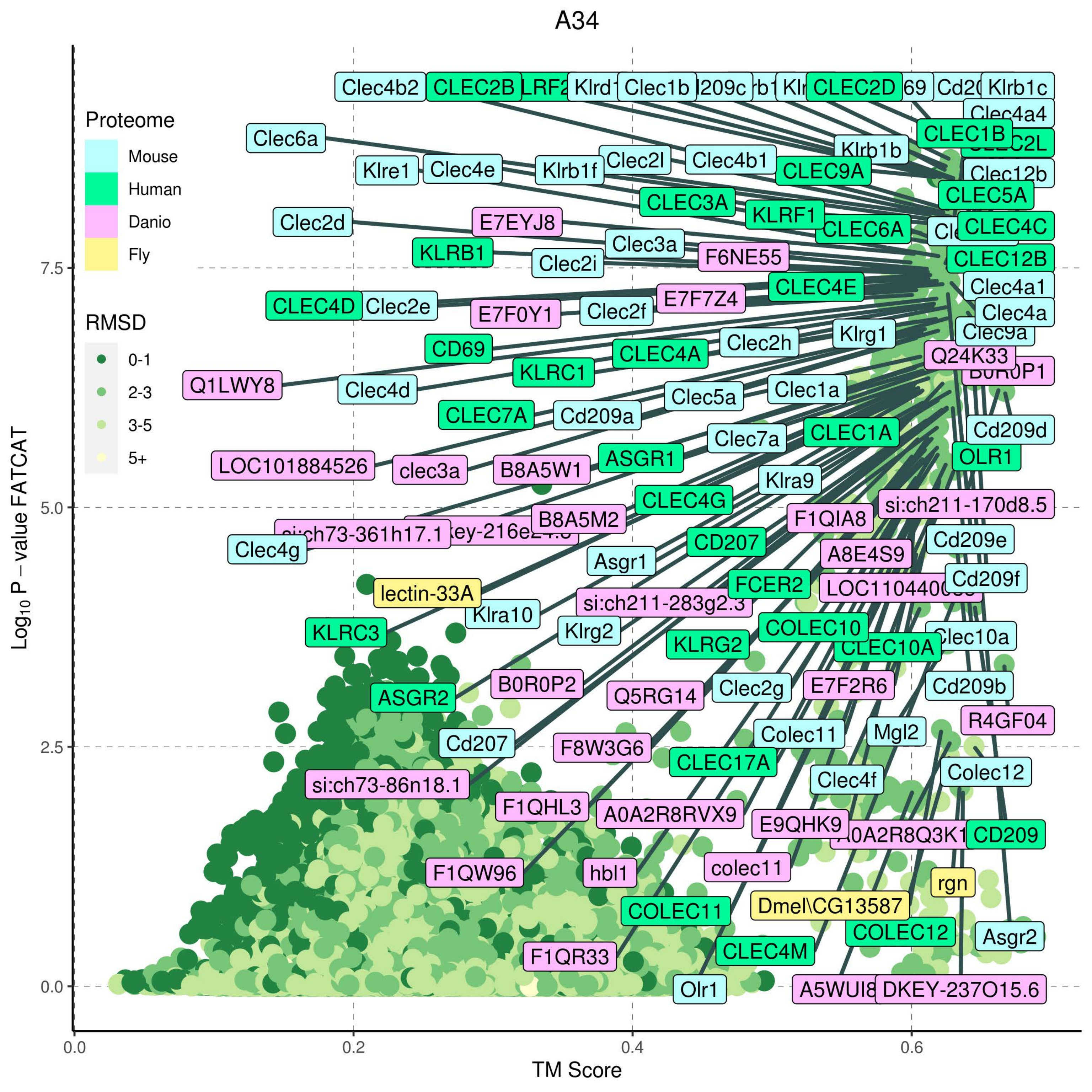

A34
Proteome
Mouse
Human
Danio
Fly
RMSD
0-1
2-3
3-5
5+
Log10 P − value FATCAT
TM Score
0.0
2.5
5.0
7.5
0.0
0.2
0.4
0.6
Clec4b2
CLEC2B
Clec1b
CLEC2D
Klrb1c
Clec4a4
CLEC1B
Clec6a
Klrb1b
Clec4b1
Clec2l
Klre1
Clec4e
Klrb1f
CLEC9A
Clec12b
CLEC5A
CLEC3A
KLRF1
Clec2d
E7EYJ8
CLEC6A
CLEC4C
Clec3a
KLRB1
F6NE55
Clec2i
CLEC12B
CLEC4E
Clec4a1
E7F7Z4
CLEC4D
Clec2e
E7F0Y1
Clec2f
Clec4a
Klrg1
Clec2h
CD69
CLEC4A
Q24K33
KLRC1
Q1LWY8
Clec4d
Clec1a
Clec5a
CLEC7A
Cd209a
Cd209d
Clec7a
CLEC1A
ASGR1
OLR1
LOC101884526
clec3a
B8A5W1
Klra9
CLEC4G
si:ch211-170d8.5
B8A5M2
F1QIA8
CD207
Cd209e
Clec4g
A8E4S9
Asgr1
Cd209f
FCER2
lectin-33A
si:ch211-283g2.3
Klra10
COLEC10
KLRC3
Klrg2
KLRG2
B0R0P2
Clec2g
E7F2R6
Cd209b
ASGR2
Q5RG14
R4GF04
Mgl2
Colec11
Cd207
F8W3G6
CLEC17A
Colec12
Clec4f
si:ch73-86n18.1
F1QHL3
A0A2R8RVX9
E9QHK9
CD209
F1QW96
hbl1
colec11
rgn
Dmel\CG13587
COLEC11
COLEC12
Asgr2
CLEC4M
F1QR33
Olr1
DKEY-237O15.6

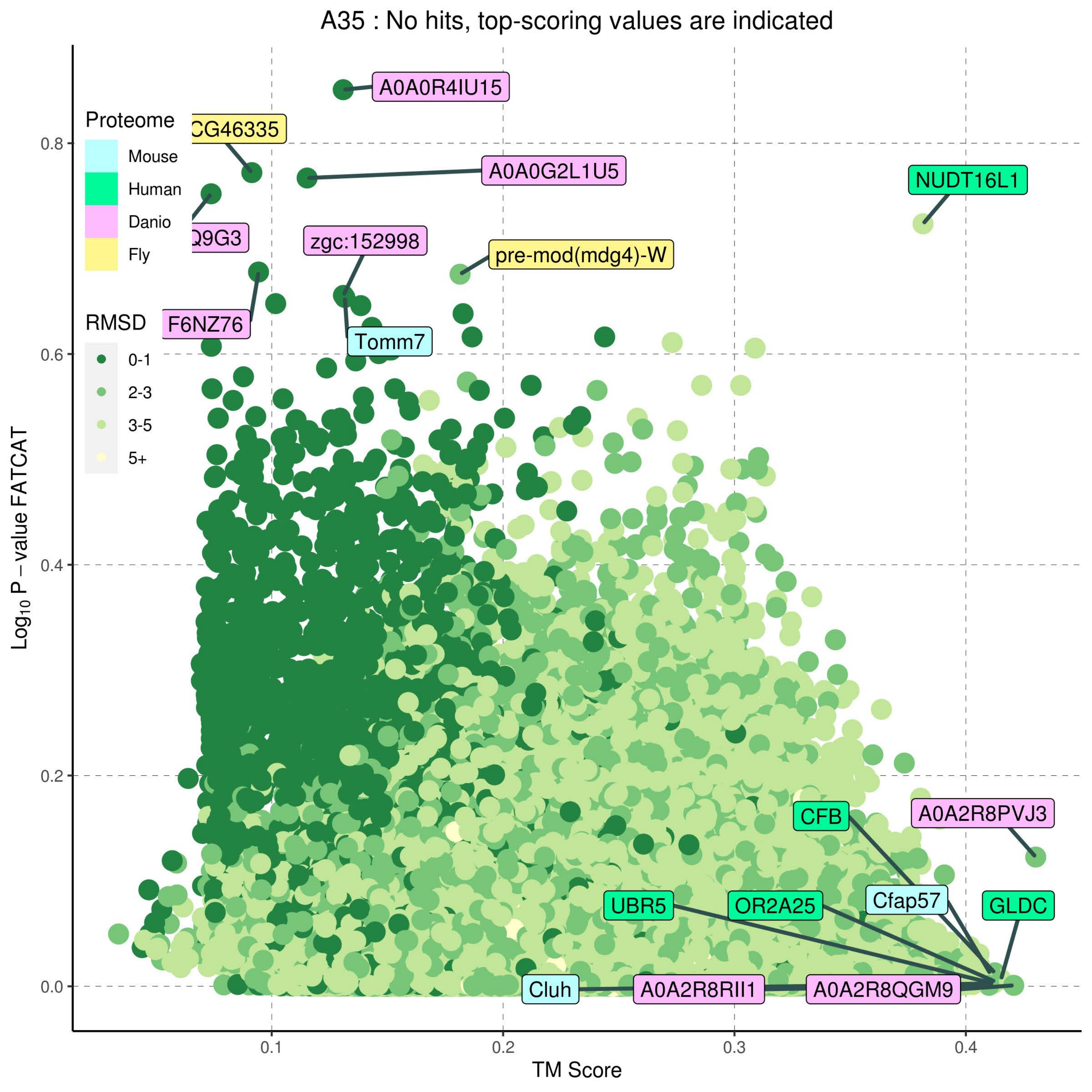
A35 : No hits, top-scoring values are indicated
Proteome
Mouse
Human
Danio
Fly
RMSD
0-1
2-3
3-5
5+
A0A0R4IU15
CG46335
A0A0G2L1U5
NUDT16L1
Q9G3
zgc:152998
pre-mod(mdg4)-W
F6NZ76
Tomm7
CFB
A0A2R8PVJ3
UBR5
OR2A25
Cfap57
GLDC
Cluh
A0A2R8RII1
A0A2R8QGM9
0.0
0.2
0.4
0.6
0.8
0.1
0.2
0.3
0.4
TM Score
Log10 P – value FATCAT

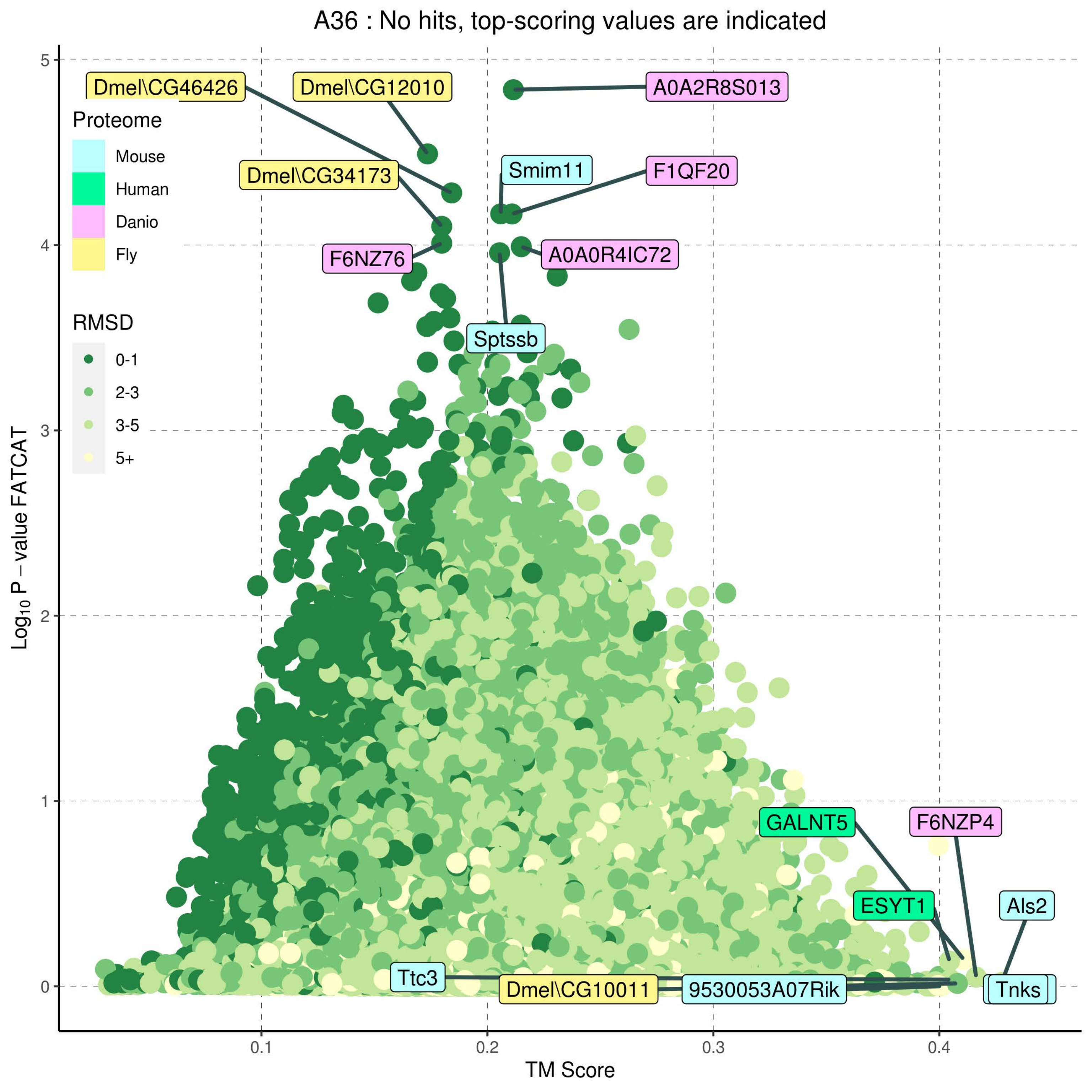
A36 : No hits, top-scoring values are indicated
Proteome
Mouse
Human
Danio
Fly
RMSD
0-1
2-3
3-5
5+
Dmel\CG46426
Dmel\CG12010
A0A2R8S013
Smim11
F1QF20
Dmel\CG34173
F6NZ76
A0A0R4IC72
Sptssb
GALNT5
F6NZP4
ESYT1
Als2
Ttc3
Dmel\CG10011
9530053A07Rik
Tnks
$Log_{10}$ P – value FATCAT
TM Score
0
1
2
3
4
5
0.1
0.2
0.3
0.4

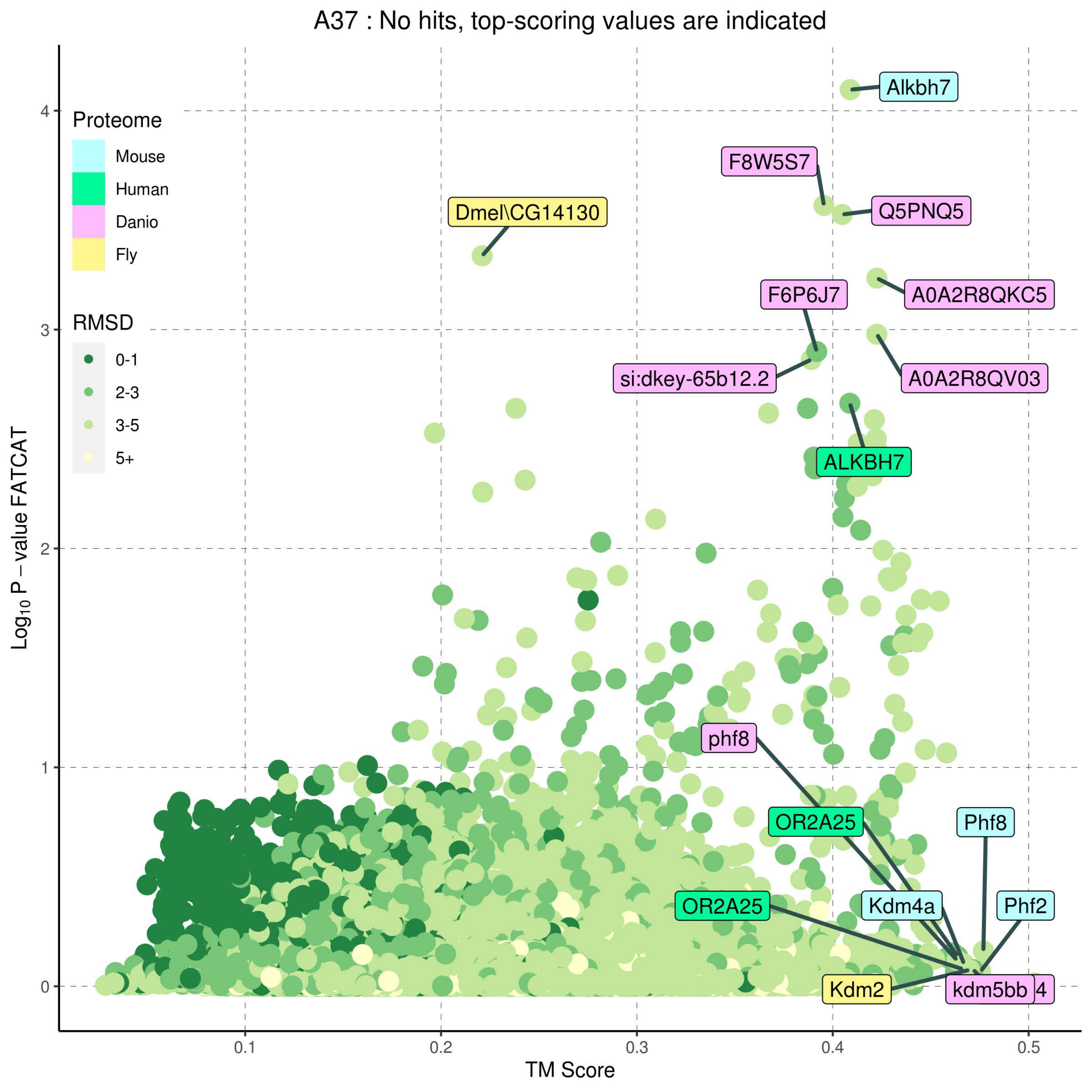
A37 : No hits, top-scoring values are indicated
Proteome
Mouse
Human
Danio
Fly
RMSD
0-1
2-3
3-5
5+
Alkbh7
F8W5S7
Q5PNQ5
Dmel\CG14130
A0A2R8QKC5
F6P6J7
si:dkey-65b12.2
A0A2R8QV03
ALKBH7
phf8
OR2A25
Phf8
OR2A25
Kdm4a
Phf2
Kdm2
kdm5bb
4
Log10 P − value FATCAT
TM Score
0
1
2
3
4
0.1
0.2
0.3
0.4
0.5

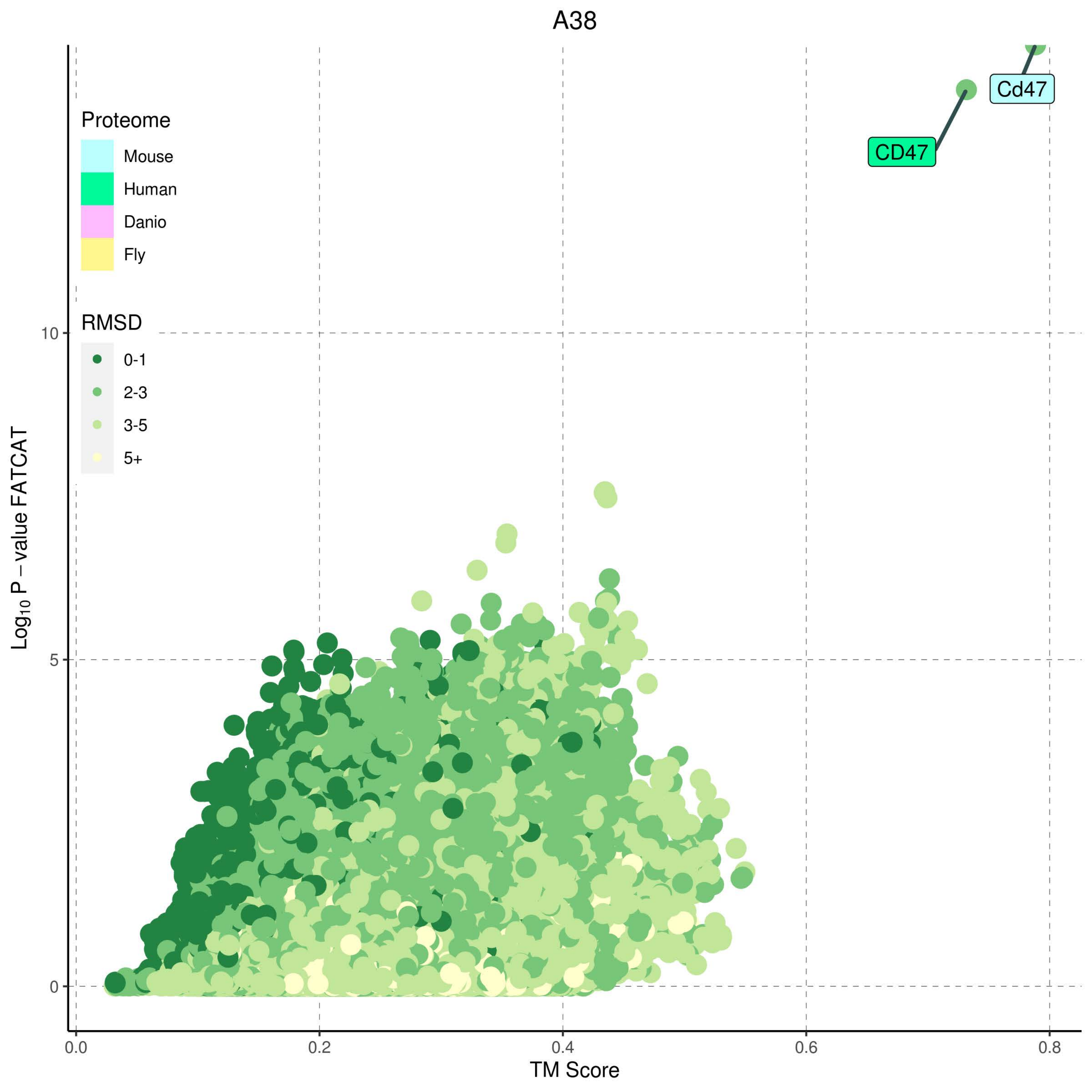
A38
Proteome
Mouse
Human
Danio
Fly
RMSD
0-1
2-3
3-5
5+
Cd47
CD47
10
5
0
Log10 P −value FATCAT
0.0
0.2
0.4
0.6
0.8
TM Score

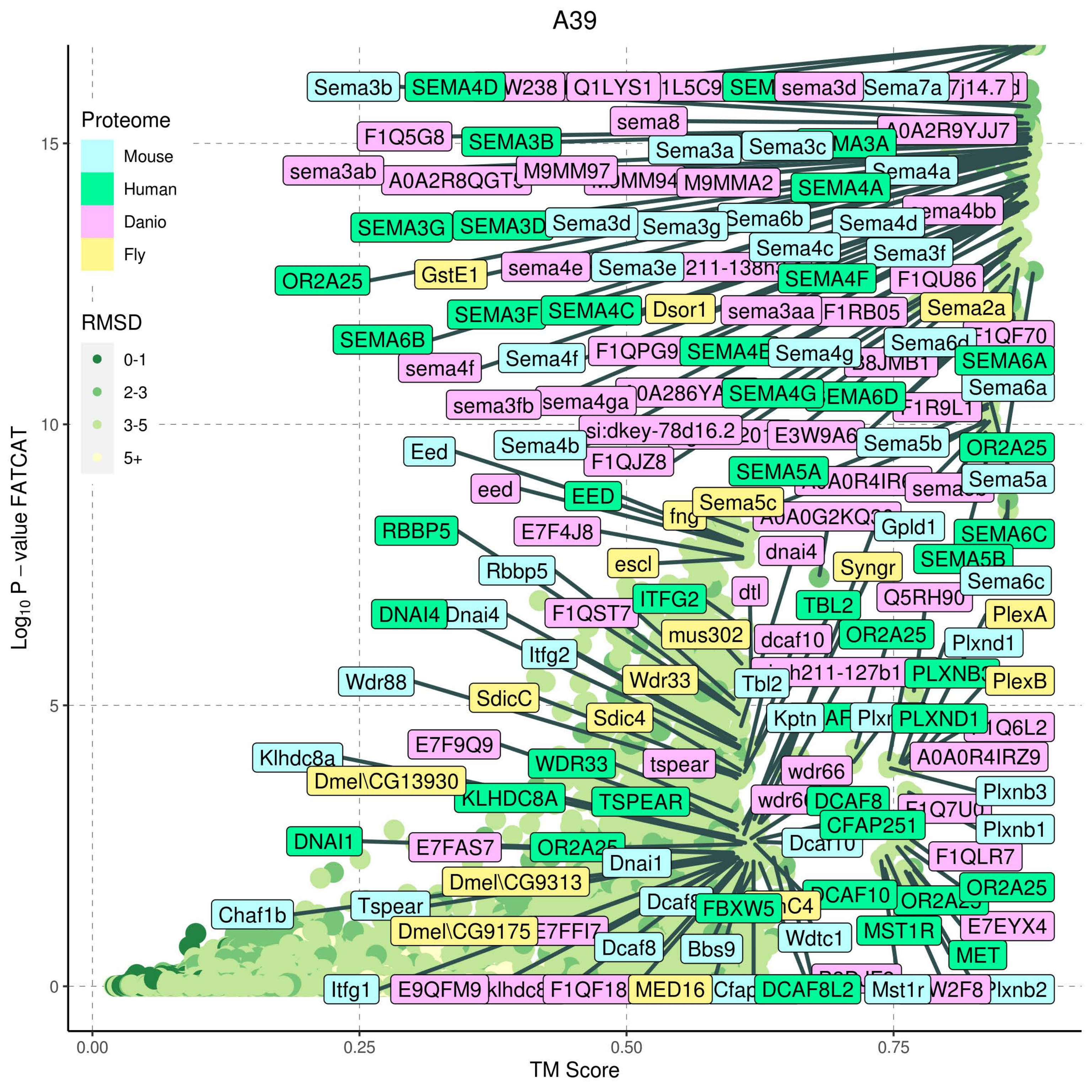

A39
Proteome
Mouse
Human
Danio
Fly
RMSD
0-1
2-3
3-5
5+
Log10 P − value FATCAT
TM Score
0.00
0.25
0.50
0.75
0
5
10
15
Sema3b
SEMA4D
Q1LYS1
sema3d
Sema7a
sema8
F1Q5G8
SEMA3B
A0A2R9YJJ7
Sema3a
Sema3c
sema3ab
M9MM97
M9MMA2
SEMA4A
Sema4a
SEMA3G
SEMA3D
Sema3d
Sema3g
Sema6b
Sema4d
Sema4c
Sema3f
OR2A25
GstE1
sema4e
Sema3e
SEMA4F
F1QU86
SEMA3F
SEMA4C
Dsor1
sema3aa
F1RB05
Sema2a
SEMA6B
sema4f
Sema4f
Sema4g
SEMA6A
Sema6a
sema3fb
sema4ga
SEMA4G
si:dkey-78d16.2
Sema5b
Eed
Sema4b
F1QJZ8
SEMA5A
Sema5a
eed
EED
Sema5c
Gpld1
SEMA6C
RBBP5
E7F4J8
dnai4
escl
Syngr
Rbbp5
SEMA5B
dtl
Sema6c
DNAI4
Dnai4
F1QST7
ITFG2
TBL2
PlexA
mus302
dcaf10
Plxnd1
Itfg2
Wdr88
SdicC
Wdr33
Tbl2
PlexB
Sdic4
Kptn
PLXND1
E7F9Q9
Klhdc8a
WDR33
tspear
wdr66
A0A0R4IRZ9
Dmel\CG13930
Plxnb3
KLHDC8A
TSPEAR
DCAF8
CFAP251
Plxnb1
DNAI1
E7FAS7
Dnai1
F1QLR7
Dmel\CG9313
DCAF10
Chaf1b
Tspear
FBXW5
Dmel\CG9175
MST1R
E7EYX4
Dcaf8
Bbs9
Wdtc1
MET
Itfg1
E9QFM9
MED16
DCAF8L2
Mst1r

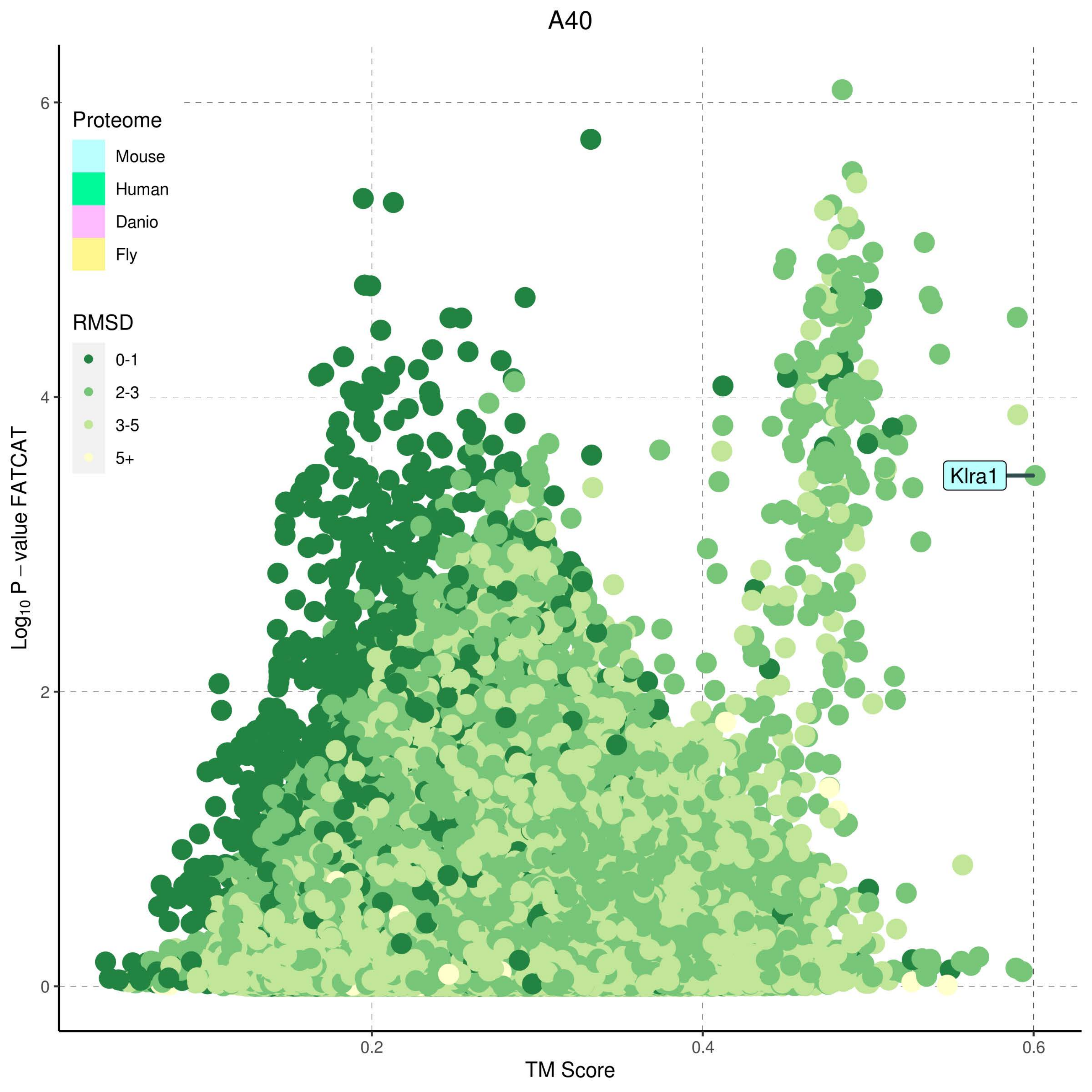
A40
Proteome
Mouse
Human
Danio
Fly
RMSD
0-1
2-3
3-5
5+
Klra1
Log10 P − value FATCAT
TM Score
0
2
4
6
0.2
0.4
0.6

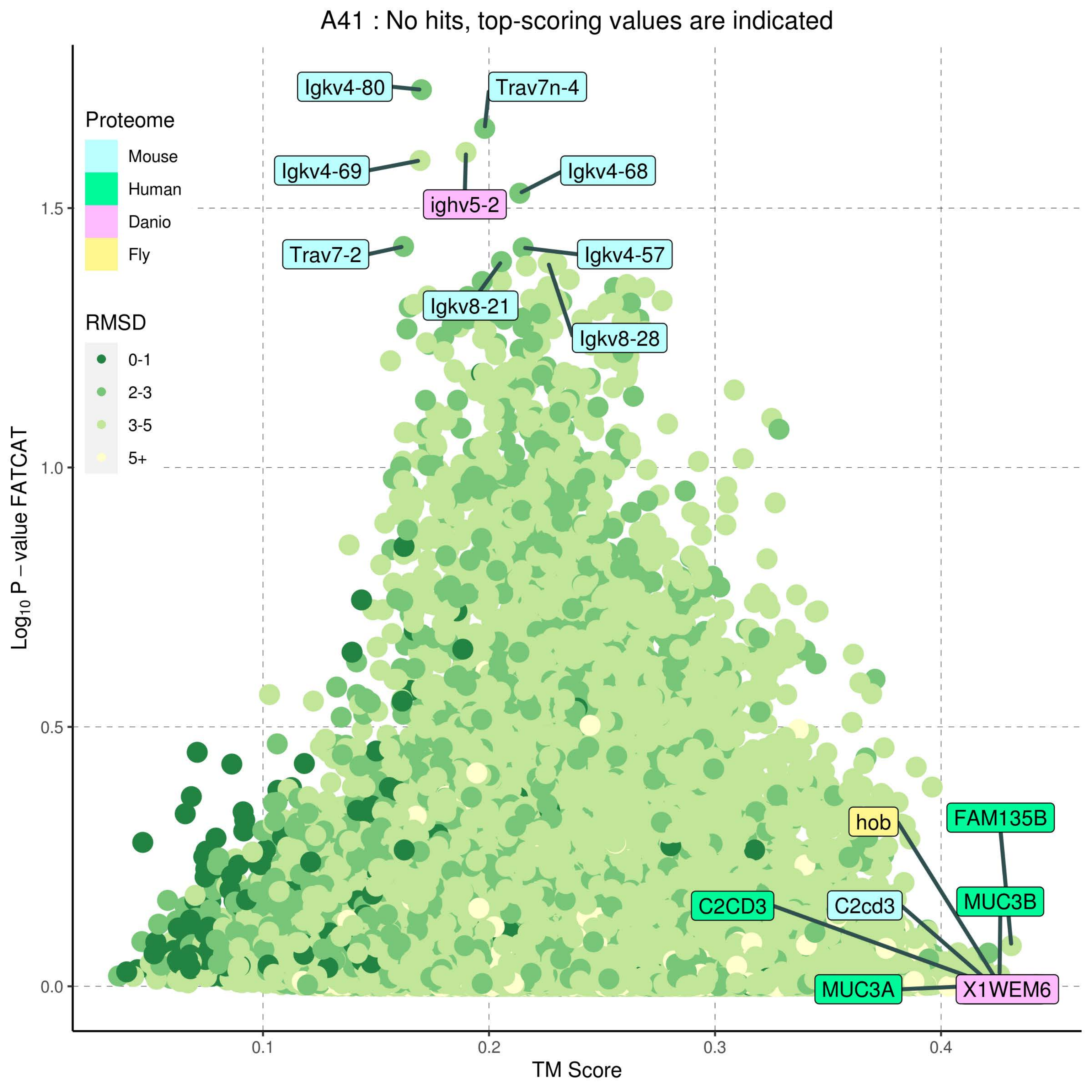

A41 : No hits, top-scoring values are indicated
Proteome
Mouse
Human
Danio
Fly
RMSD
0-1
2-3
3-5
5+
Igkv4-80
Trav7n-4
Igkv4-69
Igkv4-68
ighv5-2
Trav7-2
Igkv4-57
Igkv8-21
Igkv8-28
hob
FAM135B
C2CD3
C2cd3
MUC3B
MUC3A
X1WEM6
Log10 P – value FATCAT
TM Score
0.0
0.5
1.0
1.5
0.1
0.2
0.3
0.4

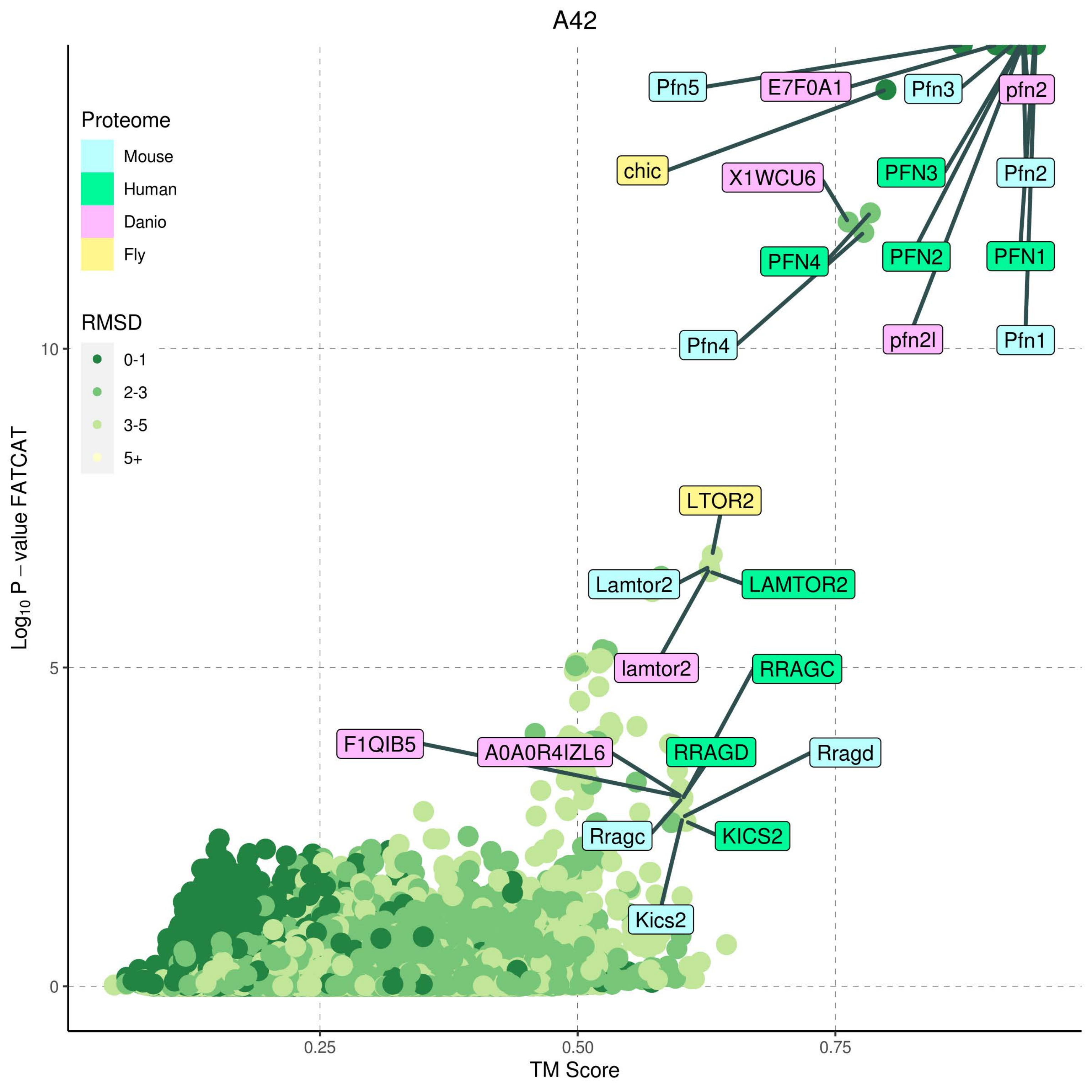
A42
Proteome
Mouse
Human
Danio
Fly
RMSD
0-1
2-3
3-5
5+
Pfn5
E7F0A1
Pfn3
pfn2
chic
X1WCU6
PFN3
Pfn2
PFN4
PFN2
PFN1
Pfn4
pfn2l
Pfn1
LTOR2
Lamtor2
LAMTOR2
lamtor2
RRAGC
F1QIB5
A0A0R4IZL6
RRAGD
Rragd
Rragc
KICS2
Kics2
10
5
0
0.25
0.50
0.75
TM Score
$Log_{10}$ P – value FATCAT

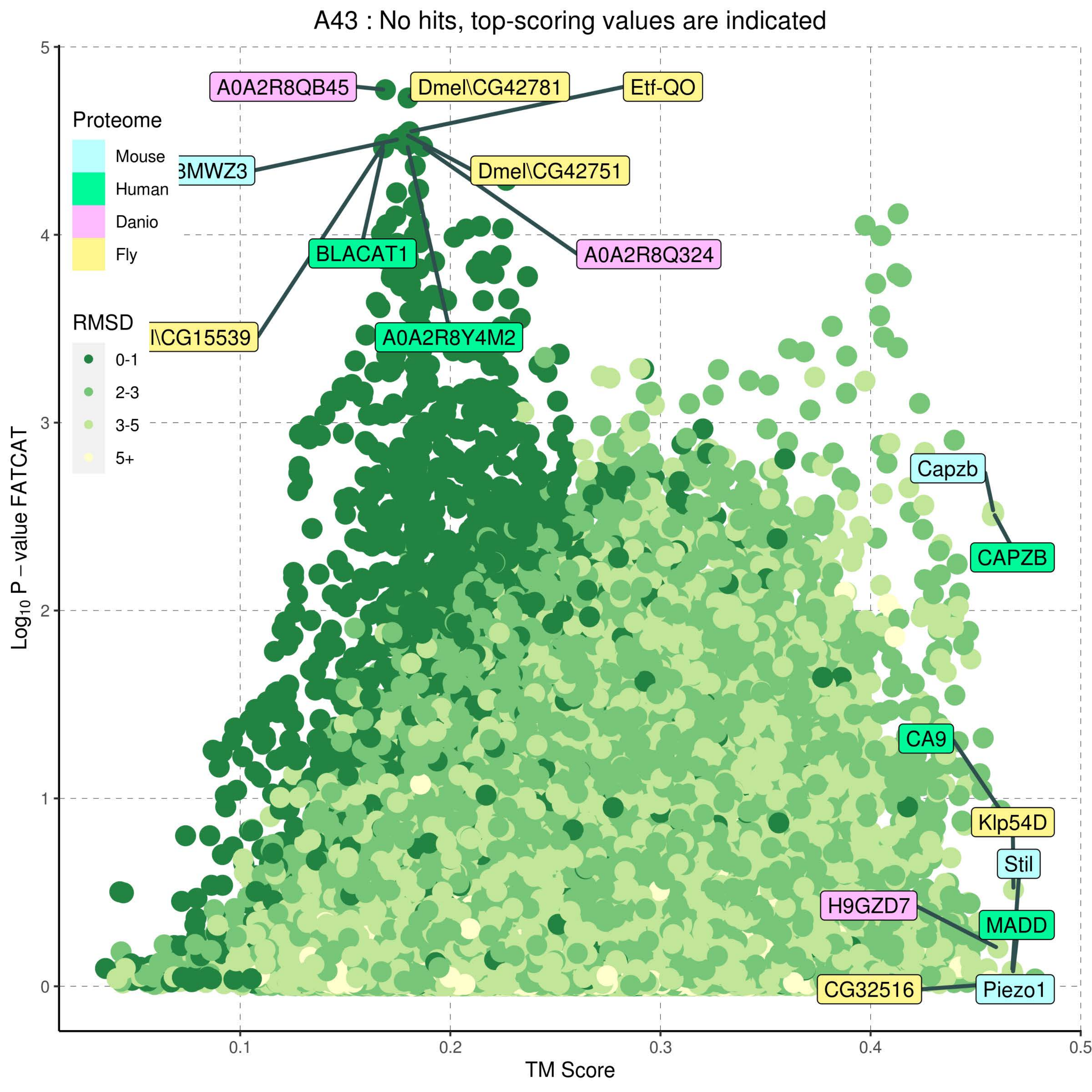
A43 : No hits, top-scoring values are indicated
Proteome
Mouse
Human
Danio
Fly
RMSD
0-1
2-3
3-5
5+
A0A2R8QB45
Dmel\CG42781
Etf-QO
3MWZ3
Dmel\CG42751
A0A2R8Q324
BLACAT1
l\CG15539
A0A2R8Y4M2
Capzb
CAPZB
CA9
Klp54D
Stil
H9GZD7
MADD
CG32516
Piezo1
Log10 P – value FATCAT
TM Score
0
1
2
3
4
5
0.1
0.2
0.3
0.4
0.5

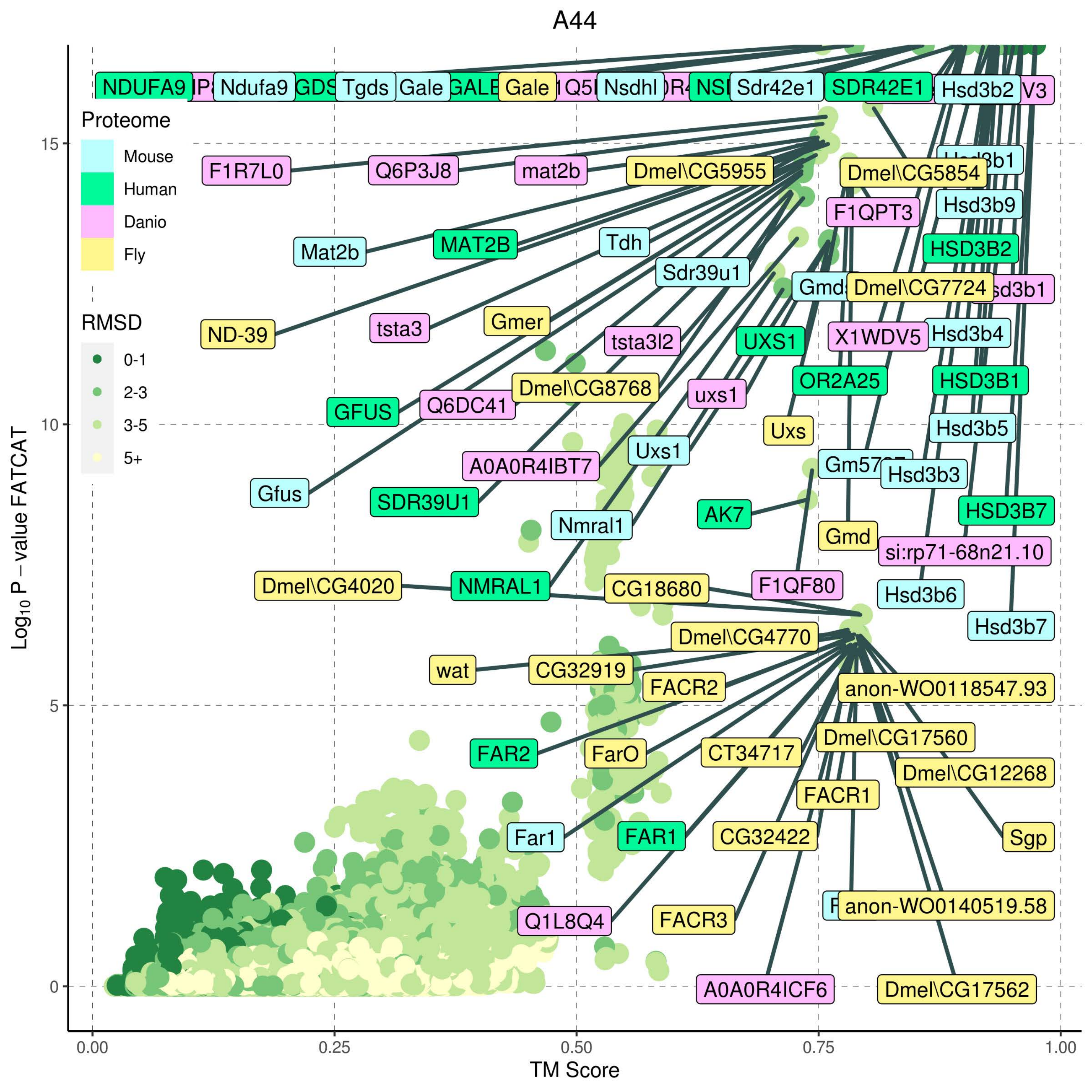

A44
Proteome
Mouse
Human
Danio
Fly
RMSD
0-1
2-3
3-5
5+
Log10 P – value FATCAT
TM Score
0.00
0.25
0.50
0.75
1.00
0
5
10
15
NDUFA9
Ndufa9
Tgds
Gale
Gale
Nsdhl
Sdr42e1
SDR42E1
Hsd3b2
F1R7L0
Q6P3J8
mat2b
Dmel\CG5955
Dmel\CG5854
Hsd3b9
F1QPT3
Mat2b
MAT2B
Tdh
HSD3B2
Sdr39u1
Dmel\CG7724
ND-39
tsta3
Gmer
tsta3l2
UXS1
X1WDV5
Hsd3b4
Dmel\CG8768
uxs1
OR2A25
HSD3B1
GFUS
Q6DC41
Uxs
Hsd3b5
Uxs1
A0A0R4IBT7
Hsd3b3
Gfus
SDR39U1
AK7
HSD3B7
Nmral1
Gmd
si:rp71-68n21.10
Dmel\CG4020
NMRAL1
CG18680
F1QF80
Hsd3b6
Hsd3b7
Dmel\CG4770
wat
CG32919
FACR2
anon-WO0118547.93
Dmel\CG17560
FAR2
FarO
CT34717
Dmel\CG12268
FACR1
Far1
FAR1
CG32422
Sgp
anon-WO0140519.58
Q1L8Q4
FACR3
A0A0R4ICF6
Dmel\CG17562

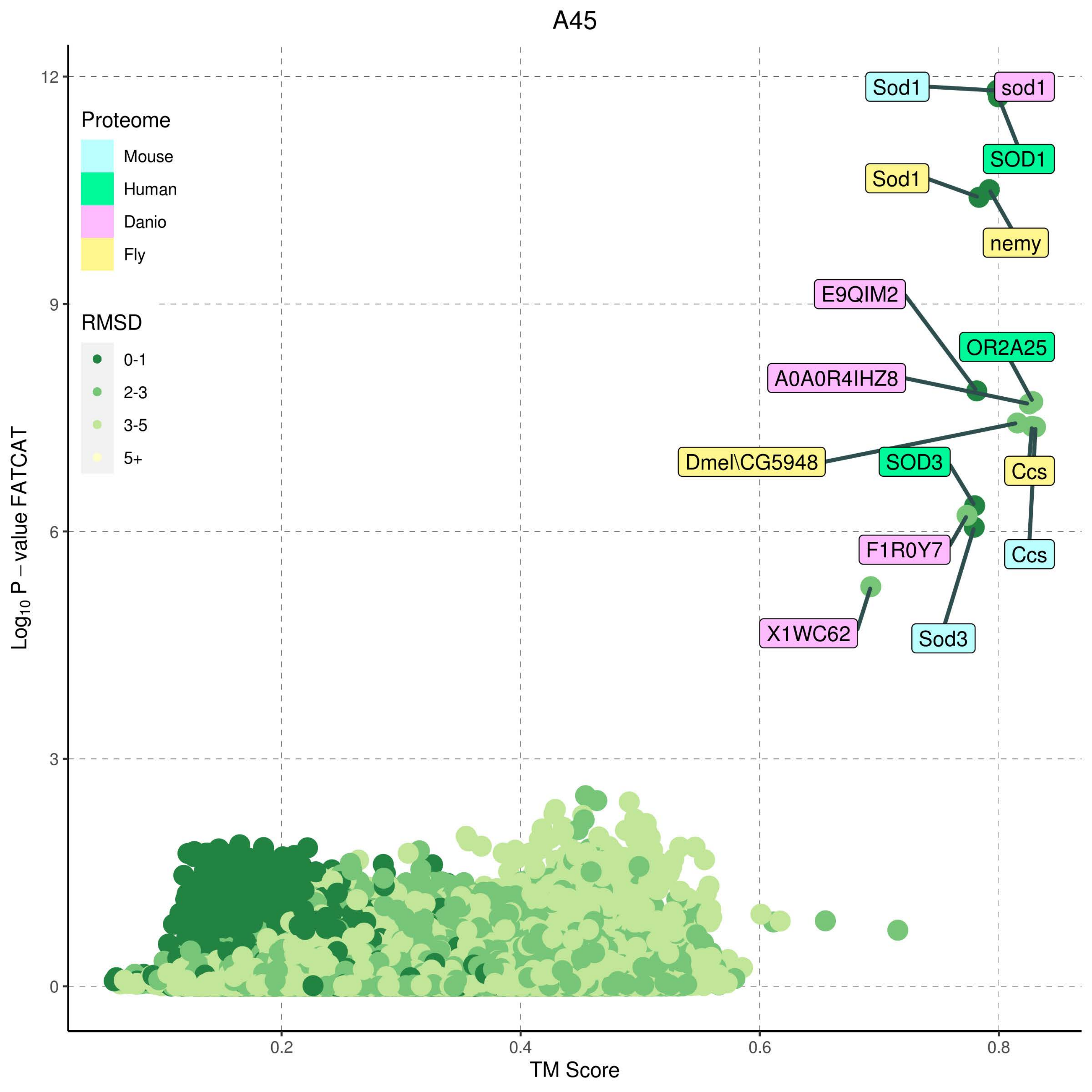
A45
Proteome
Mouse
Human
Danio
Fly
RMSD
0-1
2-3
3-5
5+
Sod1
sod1
SOD1
Sod1
nemy
E9QIM2
OR2A25
A0A0R4IHZ8
Dmel\CG5948
SOD3
Ccs
F1R0Y7
Ccs
X1WC62
Sod3
12
9
6
3
0
0.2
0.4
0.6
0.8
TM Score
$Log_{10}$ P – value FATCAT

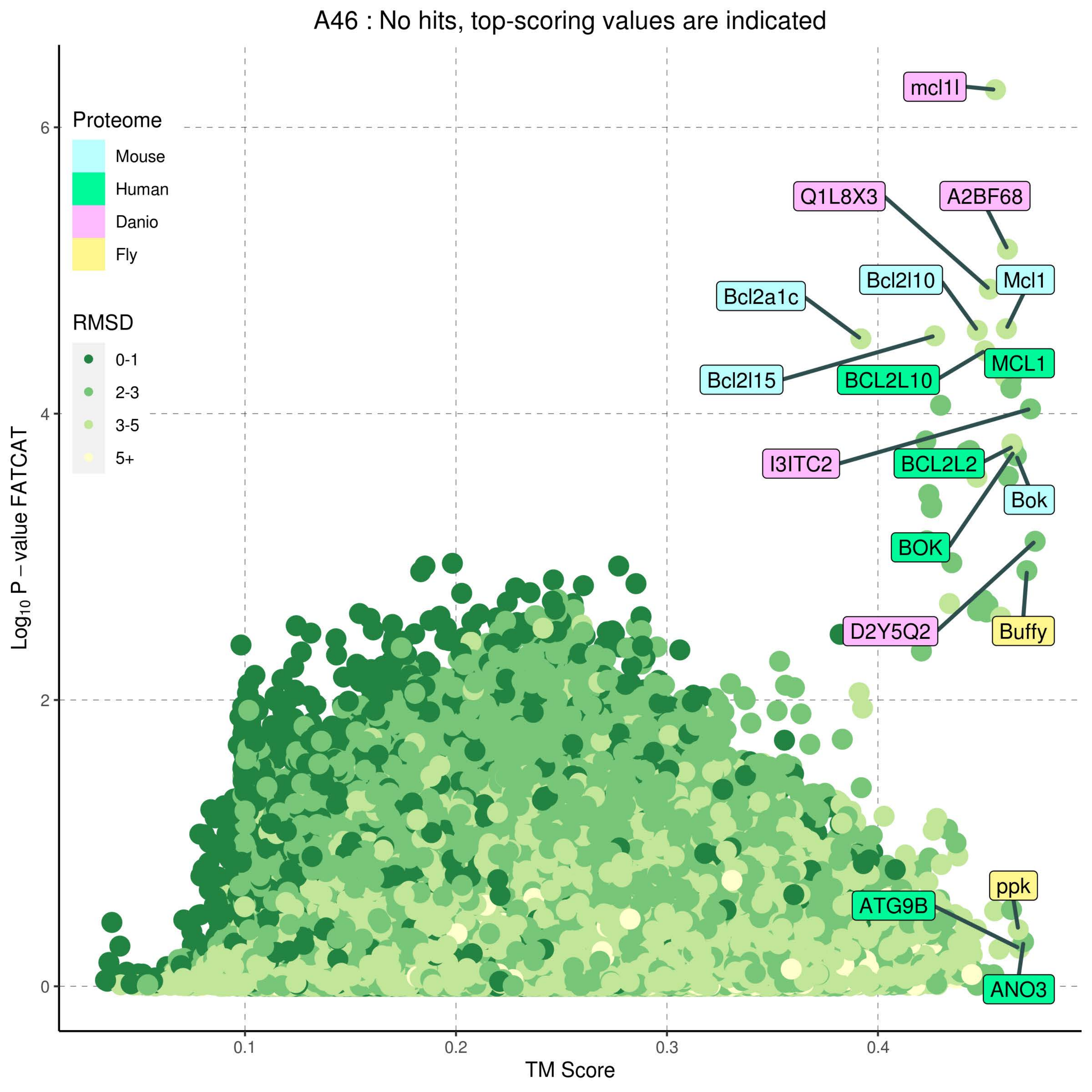

A46 : No hits, top-scoring values are indicated
Proteome
Mouse
Human
Danio
Fly
RMSD
0-1
2-3
3-5
5+
Log10 P − value FATCAT
TM Score
0.1
0.2
0.3
0.4
0
2
4
6
mcl1l
Q1L8X3
A2BF68
Bcl2l10
Mcl1
Bcl2a1c
Bcl2l15
BCL2L10
MCL1
I3ITC2
BCL2L2
Bok
BOK
D2Y5Q2
Buffy
ppk
ATG9B
ANO3

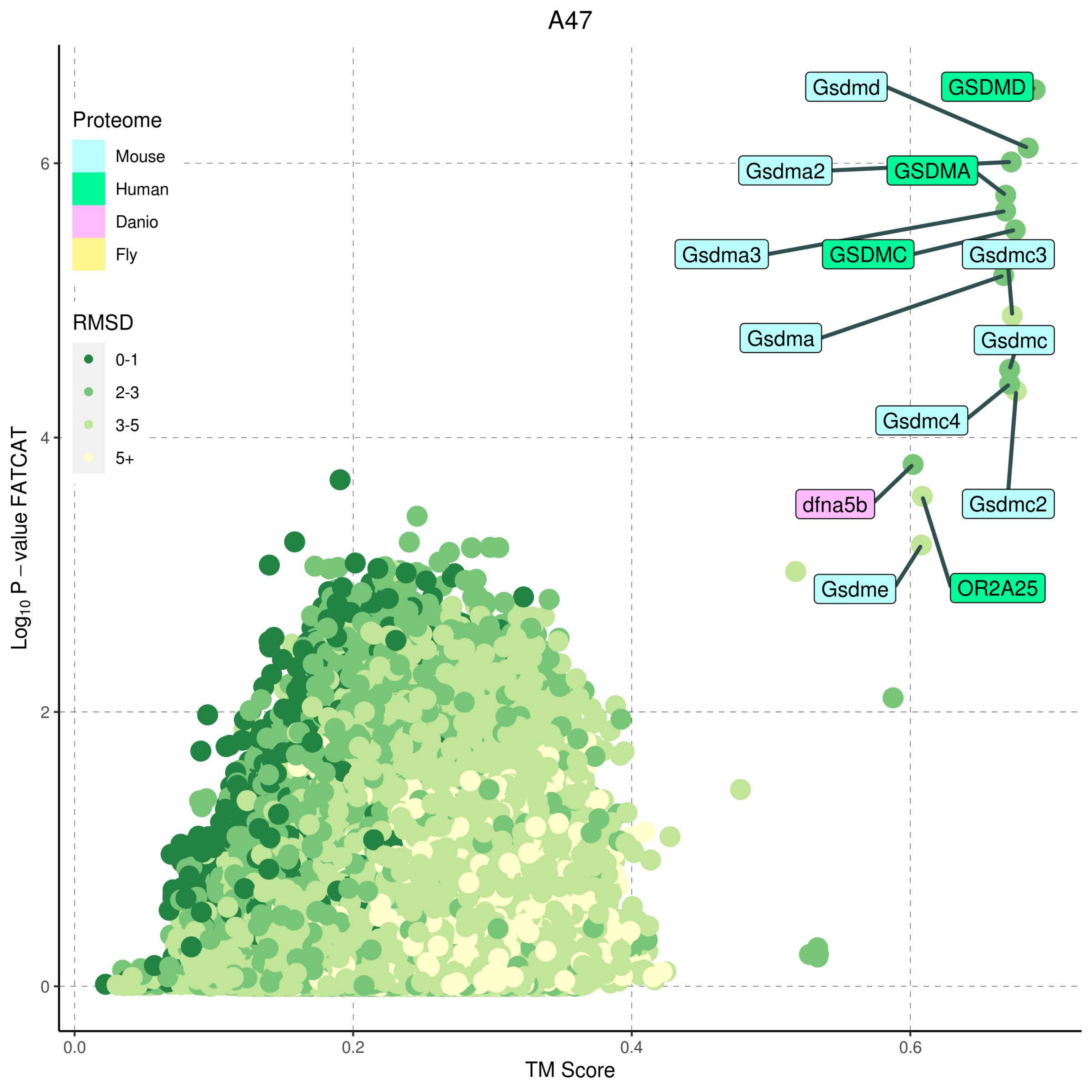

A47
Proteome
Mouse
Human
Danio
Fly
RMSD
0-1
2-3
3-5
5+
Gsdmd
GSDMD
Gsdma2
GSDMA
Gsdma3
GSDMC
Gsdmc3
Gsdma
Gsdmc
Gsdmc4
dfna5b
Gsdmc2
Gsdme
OR2A25
$Log_{10}$ P − value FATCAT
TM Score
0
2
4
6
0.0
0.2
0.4
0.6

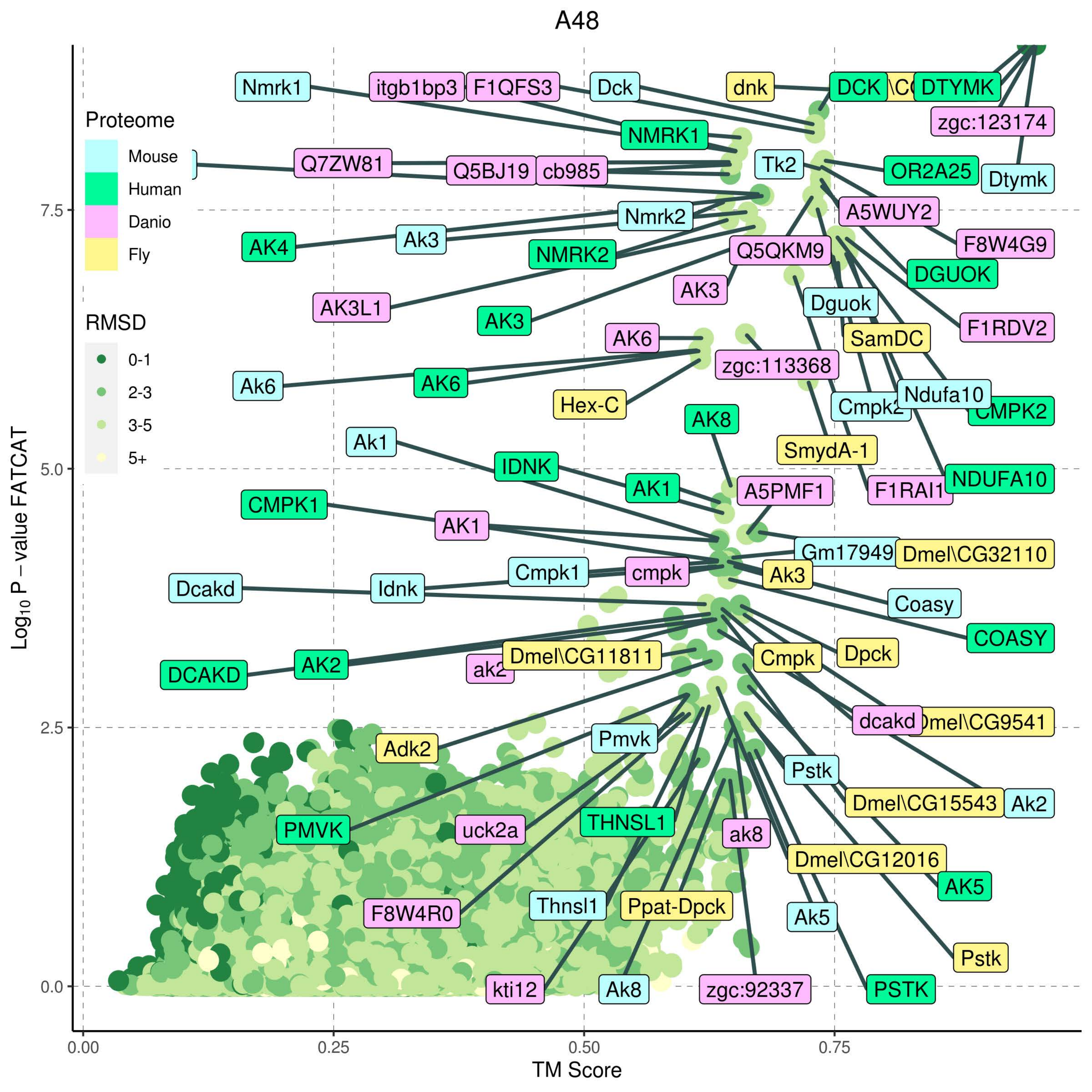

A48
Proteome
Mouse
Human
Danio
Fly
RMSD
0-1
2-3
3-5
5+
Log10 P − value FATCAT
TM Score
0.0
2.5
5.0
7.5
0.00
0.25
0.50
0.75
Nmrk1
itgb1bp3
F1QFS3
Dck
dnk
DCK
DTYMK
zgc:123174
NMRK1
Q7ZW81
Q5BJ19
cb985
Tk2
OR2A25
Dtymk
Nmrk2
A5WUY2
AK4
Ak3
NMRK2
Q5QKM9
F8W4G9
DGUOK
AK3
AK3L1
Dguok
AK3
F1RDV2
AK6
SamDC
zgc:113368
Ak6
AK6
Hex-C
Ndufa10
CMPK2
AK8
Ak1
SmydA-1
IDNK
NDUFA10
AK1
A5PMF1
F1RAI1
CMPK1
AK1
Gm17949
Dmel\CG32110
Cmpk1
cmpk
Ak3
Dcakd
Idnk
Coasy
COASY
Dmel\CG11811
Dpck
Cmpk
DCAKD
AK2
dcakd
Adk2
Pmvk
Pstk
Dmel\CG15543
Ak2
PMVK
uck2a
THNSL1
ak8
Dmel\CG12016
AK5
F8W4R0
Thnsl1
Ppat-Dpck
Ak5
Pstk
kti12
Ak8
zgc:92337
PSTK

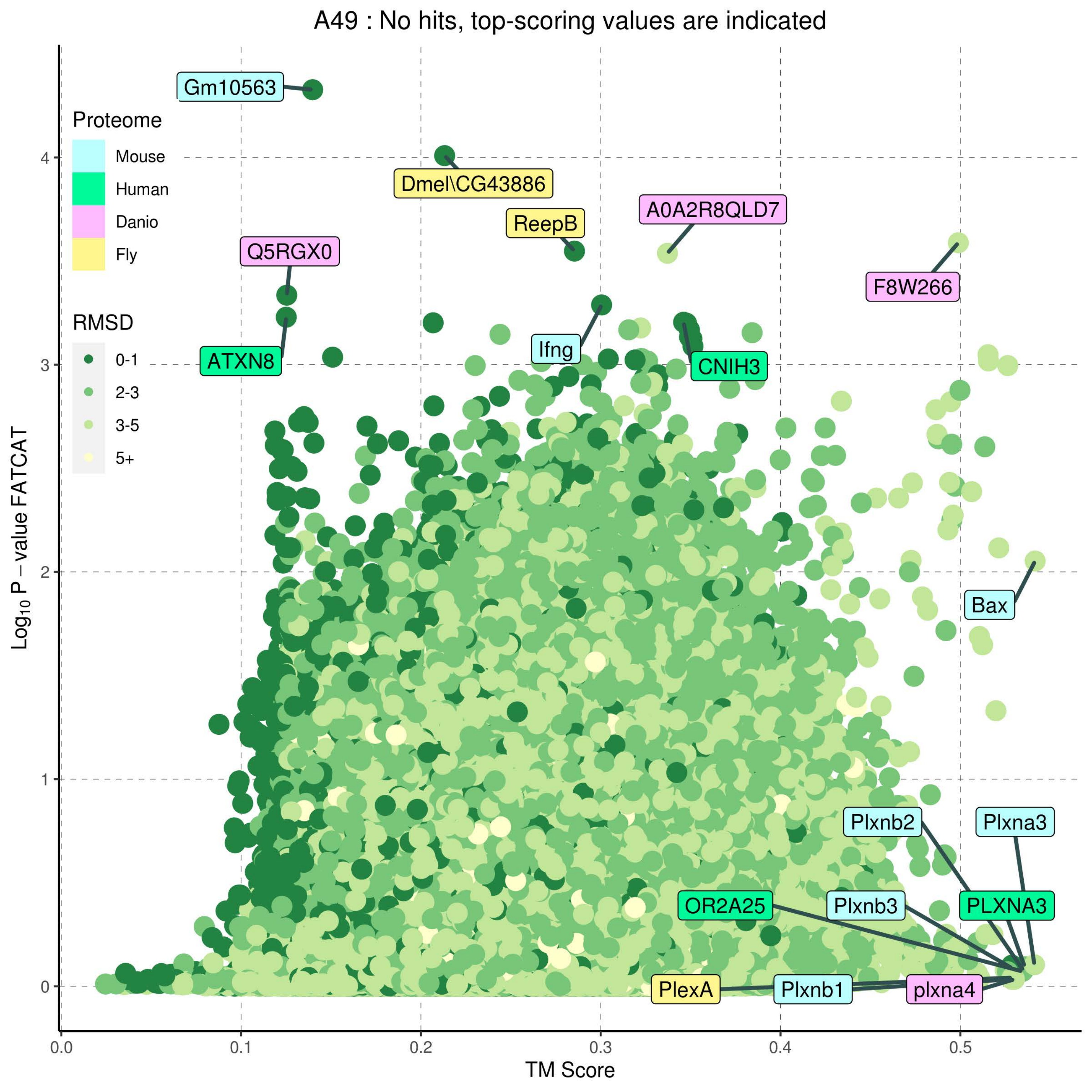

A49 : No hits, top-scoring values are indicated
Proteome
Mouse
Human
Danio
Fly
RMSD
0-1
2-3
3-5
5+
Gm10563
Dmel\CG43886
ReepB
A0A2R8QLD7
Q5RGX0
F8W266
ATXN8
Ifng
CNIH3
Bax
Plxnb2
Plxna3
OR2A25
Plxnb3
PLXNA3
PlexA
Plxnb1
plxna4
Log10 P − value FATCAT
TM Score
0.0
0.1
0.2
0.3
0.4
0.5
0
1
2
3
4

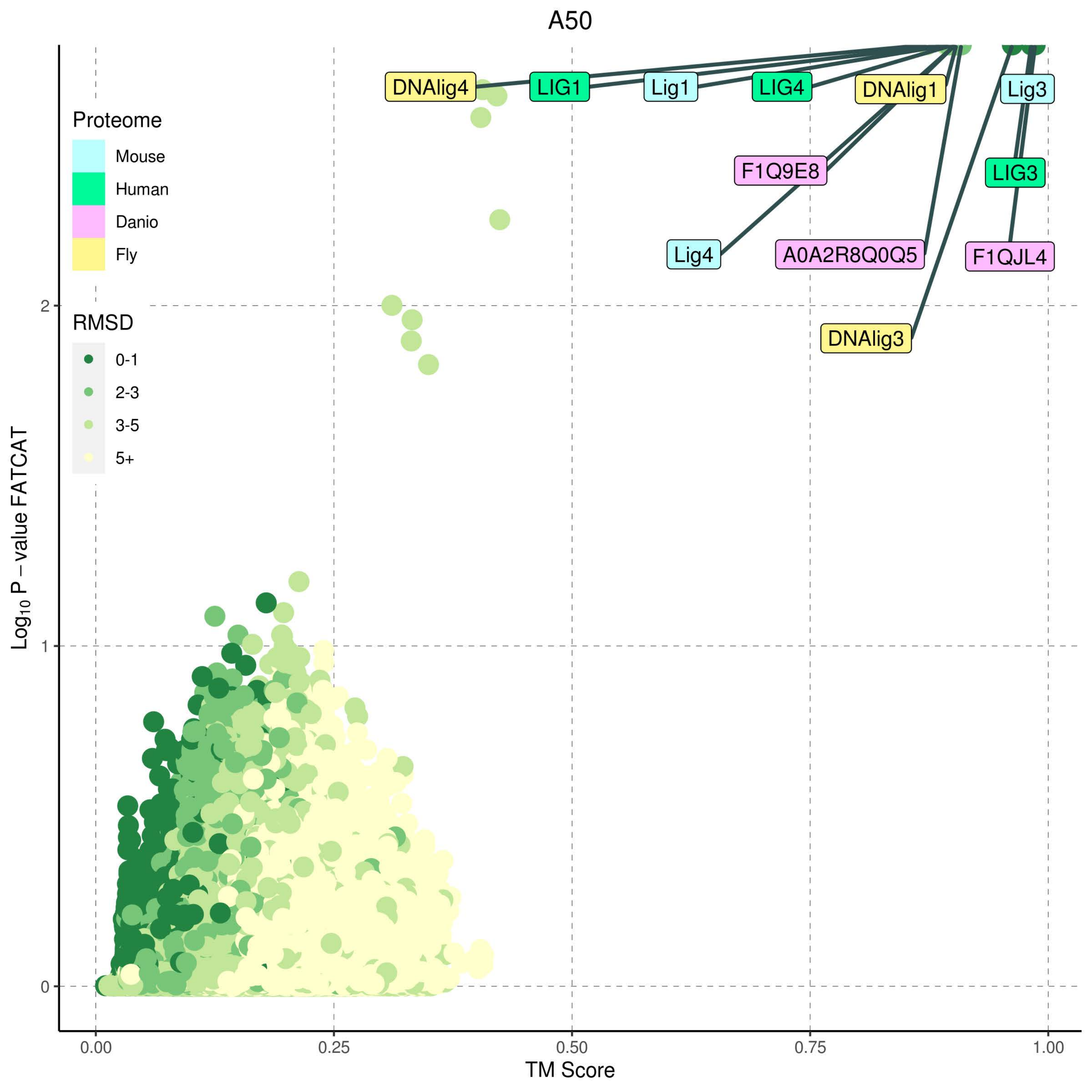
A50
Proteome
Mouse
Human
Danio
Fly
RMSD
0-1
2-3
3-5
5+
DNAlig4
LIG1
Lig1
LIG4
DNAlig1
Lig3
F1Q9E8
LIG3
Lig4
A0A2R8Q0Q5
F1QJL4
DNAlig3
$Log_{10}$ P – value FATCAT
2
1
0
0.00
0.25
0.50
0.75
1.00
TM Score

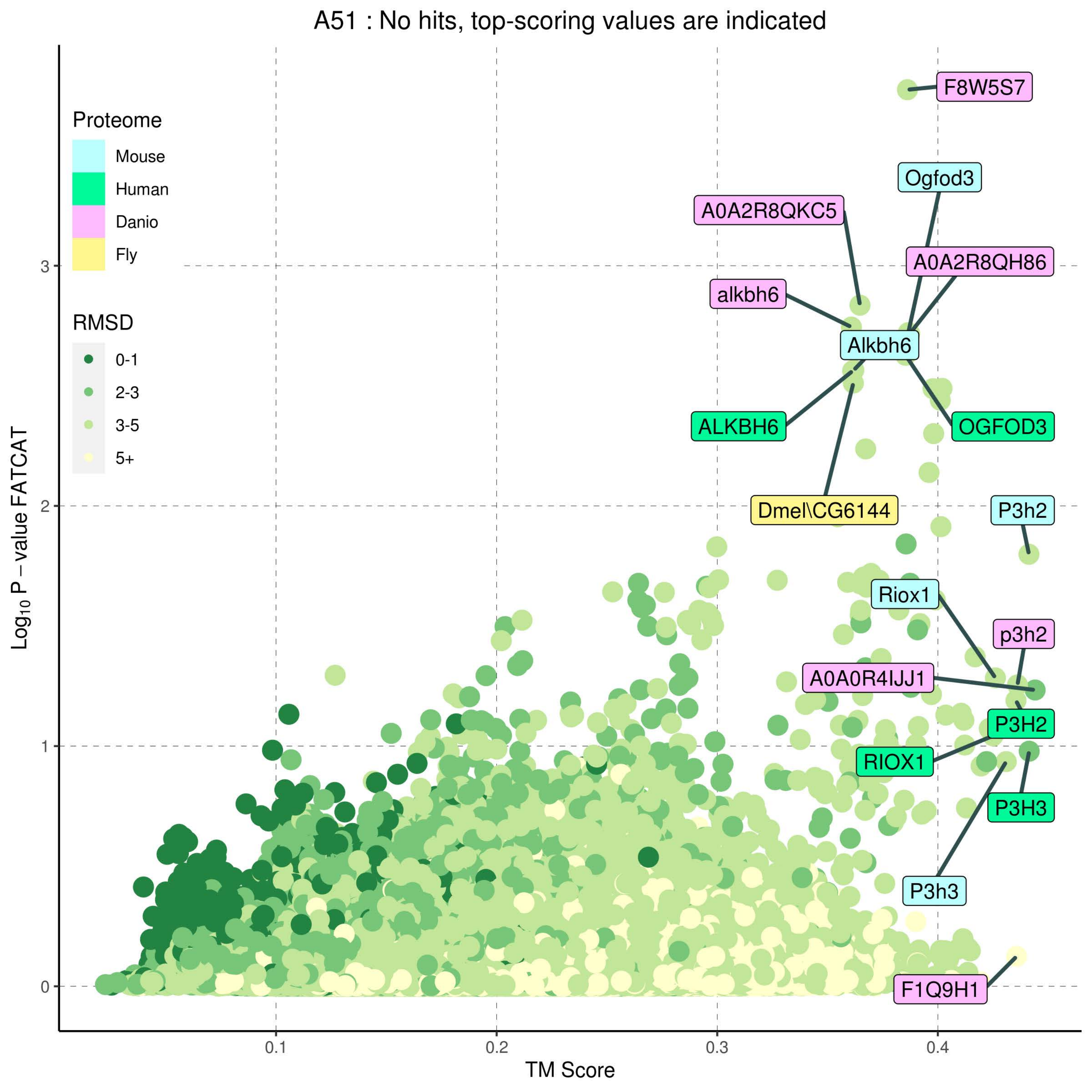
A51 : No hits, top-scoring values are indicated
Proteome
Mouse
Human
Danio
Fly
RMSD
0-1
2-3
3-5
5+
Log10 P − value FATCAT
TM Score
F8W5S7
Ogfod3
A0A2R8QKC5
A0A2R8QH86
alkbh6
Alkbh6
ALKBH6
OGFOD3
Dmel\CG6144
P3h2
Riox1
p3h2
A0A0R4IJJ1
P3H2
RIOX1
P3H3
P3h3
F1Q9H1

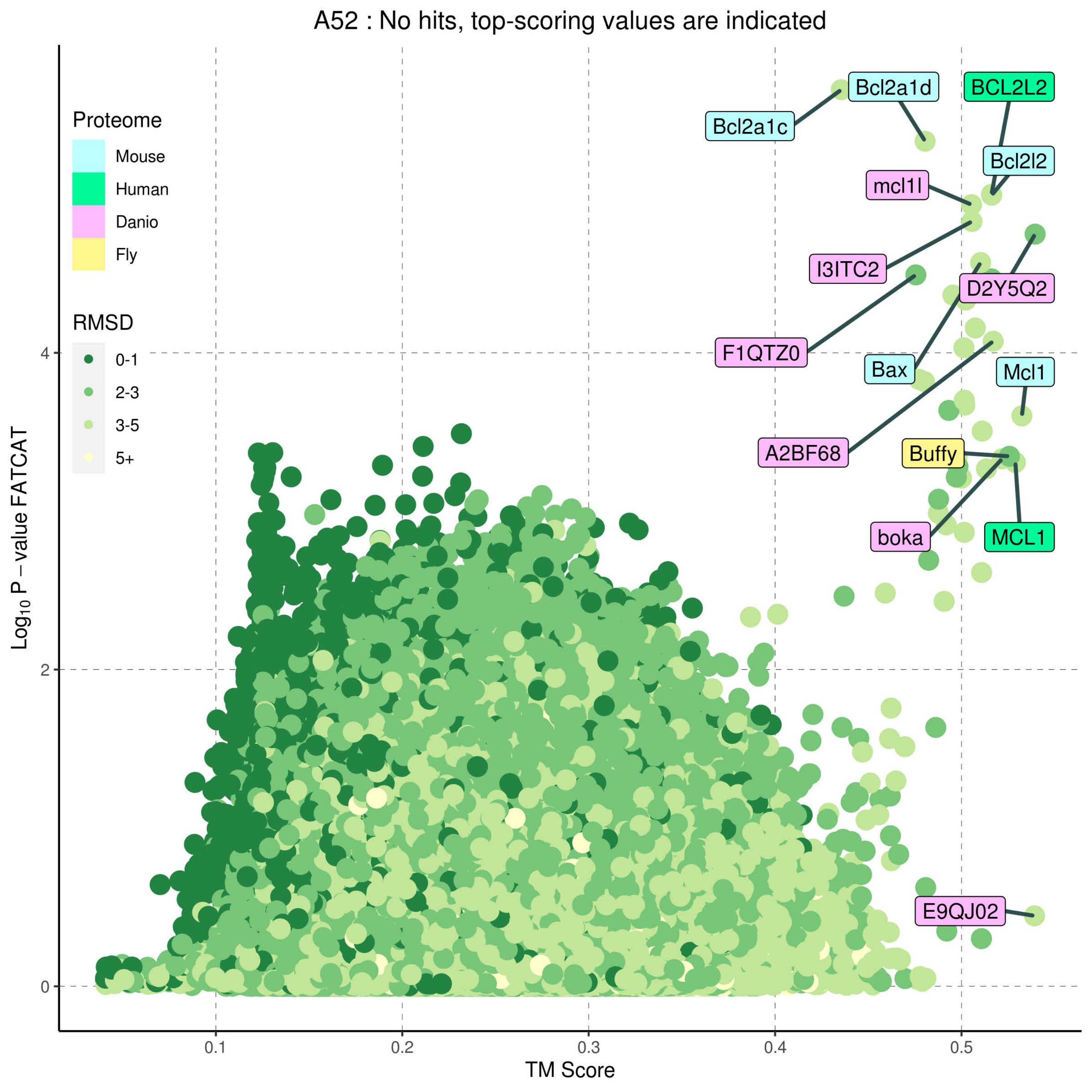
A52 : No hits, top-scoring values are indicated
Proteome
Mouse
Human
Danio
Fly
RMSD
0-1
2-3
3-5
5+
Bcl2a1d
BCL2L2
Bcl2a1c
Bcl2l2
mcl1l
I3ITC2
D2Y5Q2
F1QTZ0
Bax
Mcl1
A2BF68
Buffy
boka
MCL1
E9QJ02
Log10 P − value FATCAT
TM Score
0
2
4
0.1
0.2
0.3
0.4
0.5

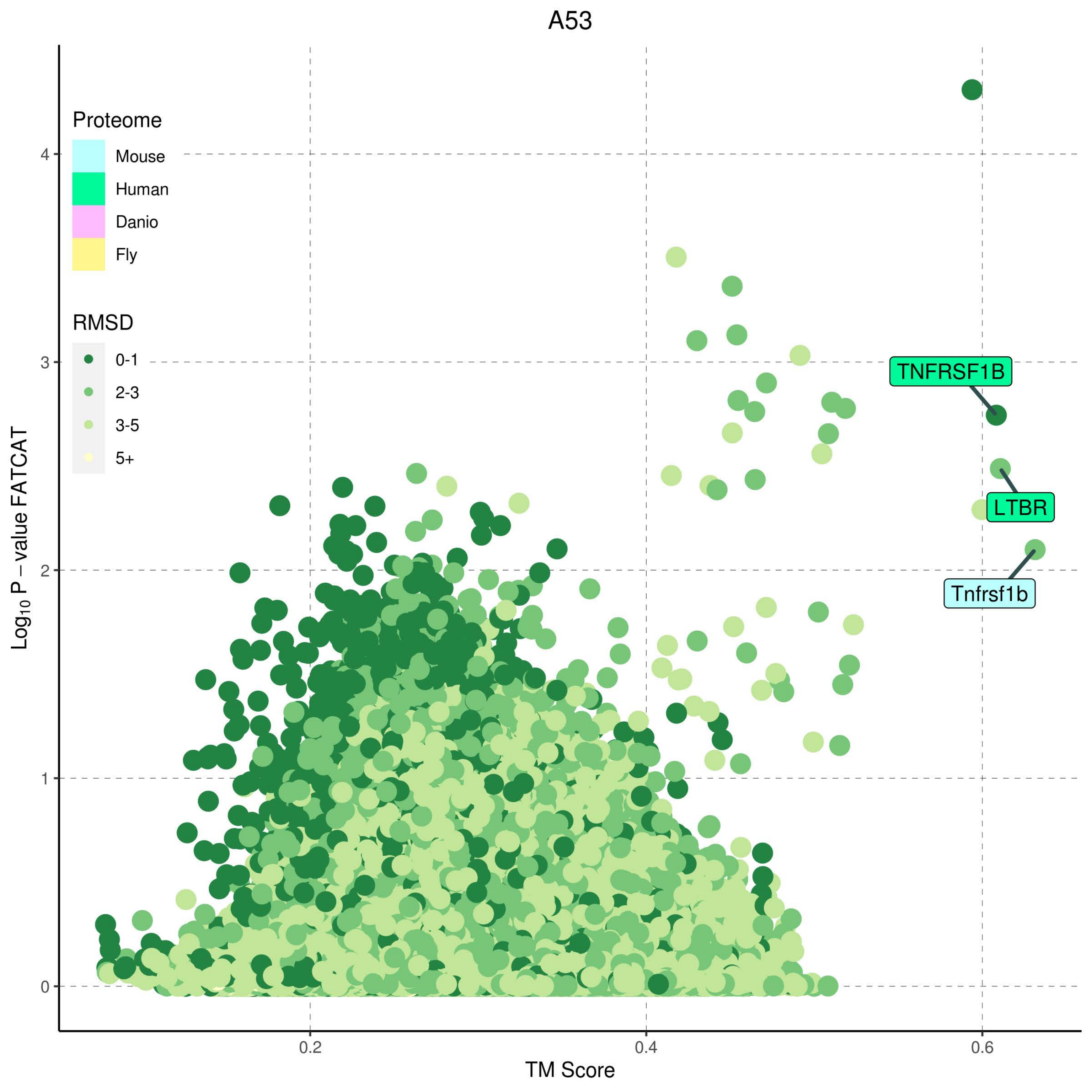

A53
Proteome
Mouse
Human
Danio
Fly
RMSD
0-1
2-3
3-5
5+
TNFRSF1B
LTBR
Tnfrsf1b
Log10 P – value FATCAT
TM Score
0
1
2
3
4
0.2
0.4
0.6

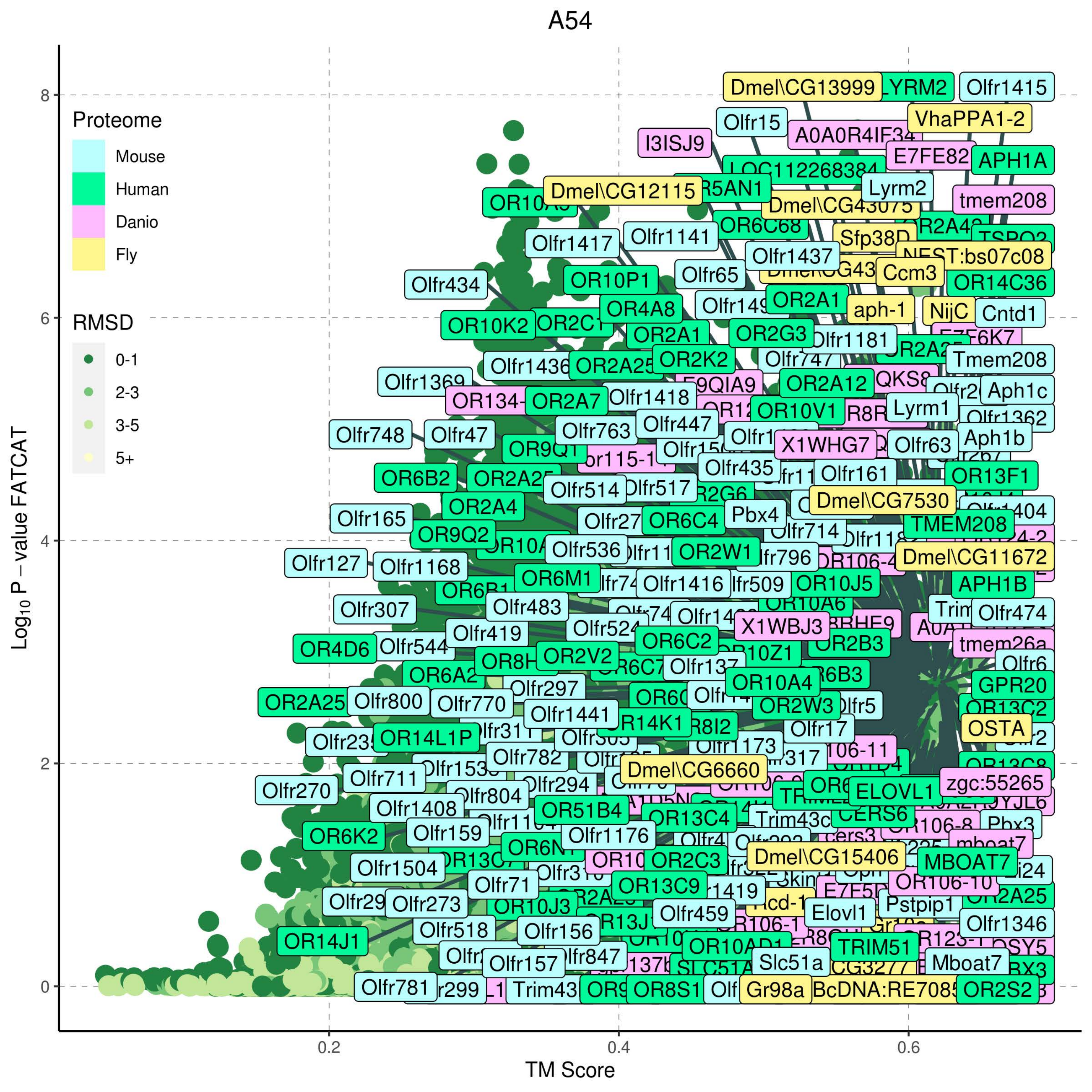
A54
Proteome
Mouse
Human
Danio
Fly
RMSD
0-1
2-3
3-5
5+
Log10 P − value FATCAT
TM Score
0
2
4
6
8
0.2
0.4
0.6
Dmel\CG13999
Olfr1415
Olfr15
VhaPPA1-2
I3ISJ9
E7FE82
APH1A
Dmel\CG12115
Lyrm2
tmem208
Olfr1417
Olfr1141
Olfr1437
Olfr434
Olfr65
Ccm3
OR14C36
OR10P1
OR4A8
aph-1
NijC
Cntd1
OR10K2
OR2A1
OR2G3
Olfr1181
Olfr1436
OR2K2
Tmem208
Olfr1369
OR2A12
Aph1c
OR2A7
Olfr1418
Lyrm1
OR10V1
Olfr748
Olfr47
Olfr763
Olfr447
X1WHG7
Olfr63
Aph1b
Olfr435
Olfr161
OR13F1
OR6B2
Olfr514
Dmel\CG7530
OR2A4
Pbx4
Olfr165
OR6C4
Olfr714
TMEM208
OR9Q2
Olfr536
OR2W1
Dmel\CG11672
Olfr127
Olfr1168
OR6M1
Olfr1416
OR10J5
APH1B
Olfr307
Olfr483
Olfr474
Olfr419
Olfr524
X1WBJ3
OR4D6
Olfr544
OR6C2
OR2B3
OR2V2
GPR20
Olfr297
OR10A4
OR2A25
Olfr800
Olfr770
OR2W3
Olfr1441
OSTA
OR14L1P
Olfr17
Olfr782
Dmel\CG6660
Olfr270
Olfr711
zgc:55265
Olfr804
Olfr294
ELOVL1
Olfr1408
OR51B4
OR13C4
CERS6
Olfr159
OR6K2
Olfr1176
Pbx3
Dmel\CG15406
MBOAT7
Olfr1504
OR2C3
Olfr71
OR13C9
Olfr273
OR10J3
Olfr459
Elovl1
Pstpip1
Olfr518
Olfr156
Olfr1346
OR14J1
TRIM51
Olfr157
Slc51a
Mboat7
Olfr781
Trim43
OR8S1
Gr98a
OR2S2

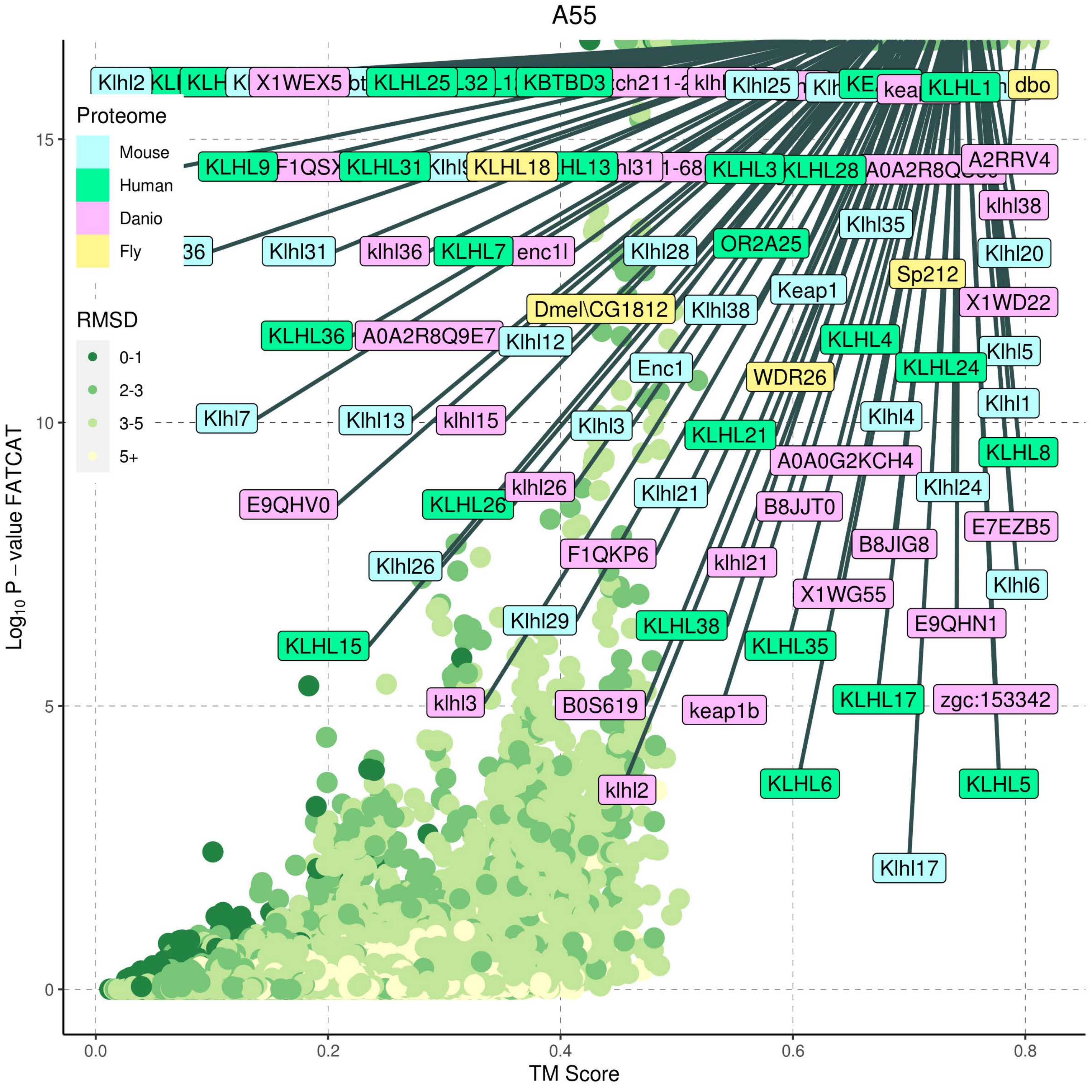

A55
Proteome
Mouse
Human
Danio
Fly
RMSD
0-1
2-3
3-5
5+
Log10 P – value FATCAT
TM Score
0.0
0.2
0.4
0.6
0.8
0
5
10
15
Klhl2
X1WEX5
KLHL25
KBTBD3
Klhl25
KLHL1
dbo
KLHL9
KLHL31
KLHL18
KLHL3
KLHL28
A2RRV4
klhl38
Klhl35
Klhl31
klhl36
KLHL7
enc1l
Klhl28
OR2A25
Klhl20
Sp212
Keap1
Dmel\CG1812
Klhl38
X1WD22
KLHL36
A0A2R8Q9E7
Klhl12
KLHL4
Klhl5
Enc1
WDR26
KLHL24
Klhl7
Klhl13
klhl15
Klhl3
Klhl4
Klhl1
KLHL21
A0A0G2KCH4
KLHL8
klhl26
Klhl21
Klhl24
E9QHV0
KLHL26
B8JJT0
E7EZB5
F1QKP6
B8JIG8
Klhl26
klhl21
X1WG55
Klhl6
Klhl29
KLHL38
E9QHN1
KLHL15
KLHL35
klhl3
B0S619
keap1b
KLHL17
zgc:153342
klhl2
KLHL6
KLHL5
Klhl17

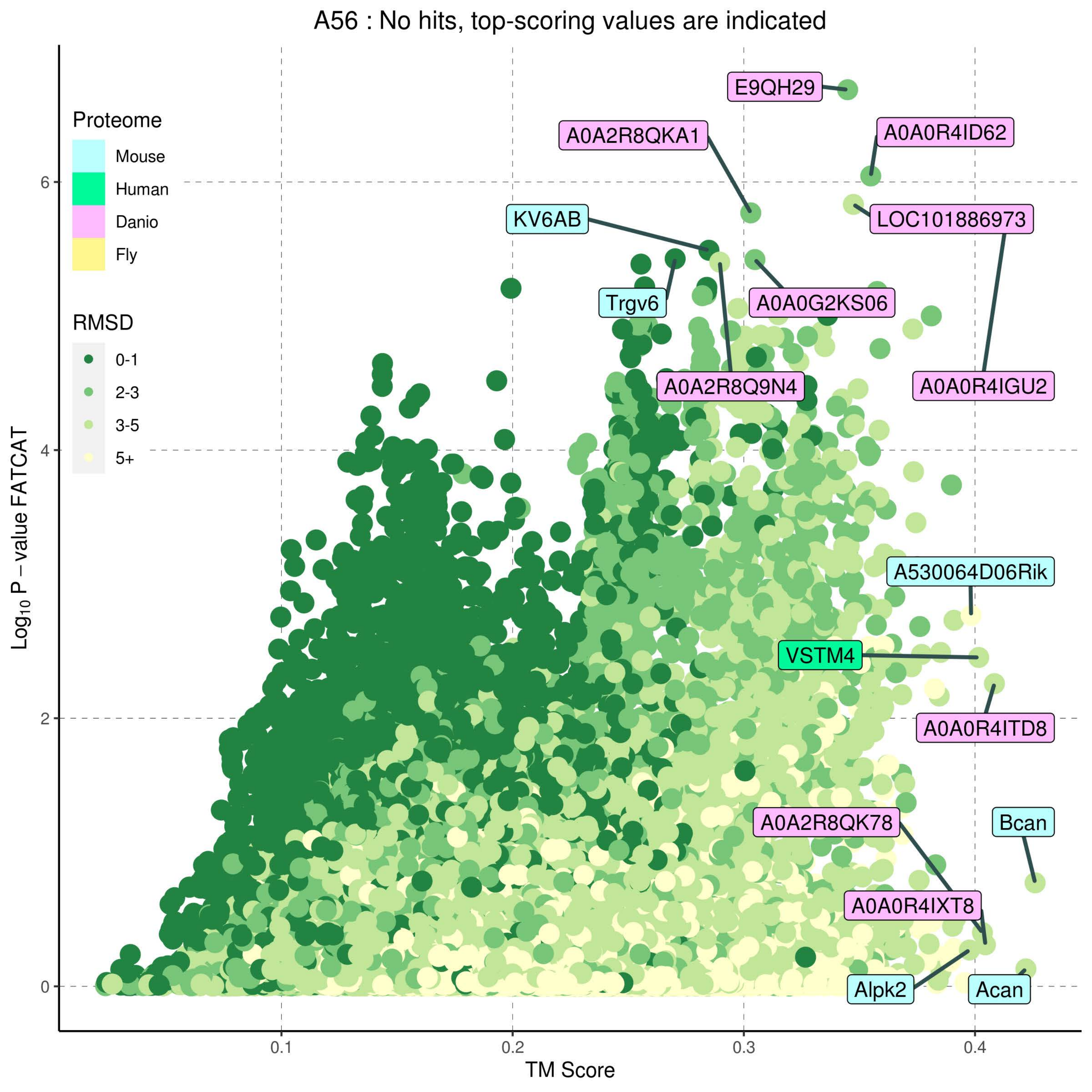

A56 : No hits, top-scoring values are indicated
Proteome
Mouse
Human
Danio
Fly
RMSD
0-1
2-3
3-5
5+
Log10 P − value FATCAT
TM Score
0
2
4
6
0.1
0.2
0.3
0.4
E9QH29
A0A2R8QKA1
A0A0R4ID62
KV6AB
LOC101886973
Trgv6
A0A0G2KS06
A0A2R8Q9N4
A0A0R4IGU2
A530064D06Rik
VSTM4
A0A0R4ITD8
A0A2R8QK78
Bcan
A0A0R4IXT8
Alpk2
Acan

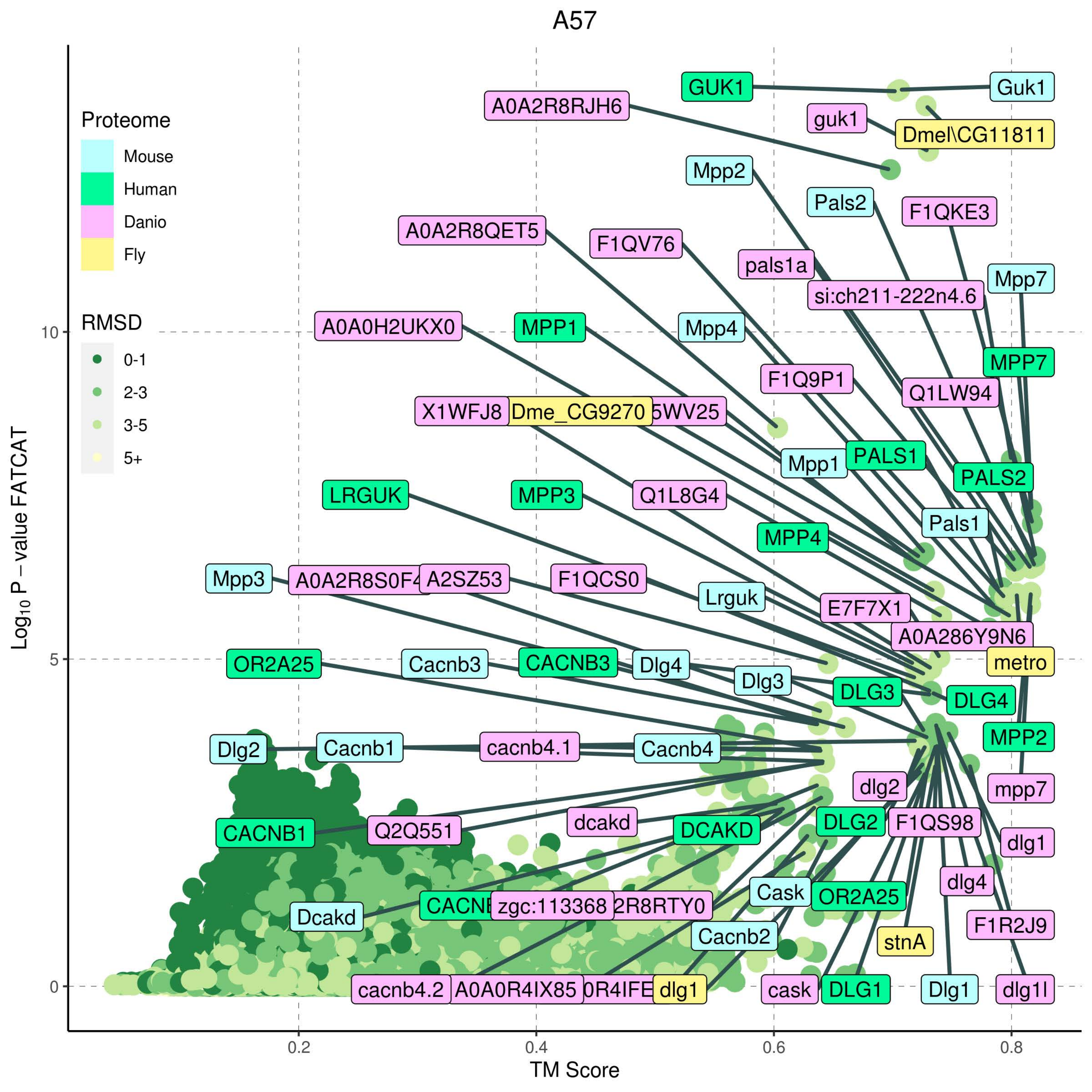

A57
Proteome
Mouse
Human
Danio
Fly
RMSD
0-1
2-3
3-5
5+
Log10 P − value FATCAT
TM Score
0
5
10
0.2
0.4
0.6
0.8
GUK1
Guk1
A0A2R8RJH6
guk1
Dmel\CG11811
Mpp2
Pals2
F1QKE3
A0A2R8QET5
F1QV76
pals1a
Mpp7
si:ch211-222n4.6
A0A0H2UKX0
MPP1
Mpp4
MPP7
F1Q9P1
Q1LW94
X1WFJ8
Dme_CG9270
Mpp1
PALS1
PALS2
LRGUK
MPP3
Q1L8G4
Pals1
MPP4
Mpp3
A2SZ53
F1QCS0
Lrguk
E7F7X1
A0A286Y9N6
OR2A25
Cacnb3
CACNB3
Dlg4
metro
Dlg3
DLG3
DLG4
MPP2
Dlg2
Cacnb1
cacnb4.1
Cacnb4
dlg2
mpp7
CACNB1
Q2Q551
dcakd
DCAKD
DLG2
F1QS98
dlg1
dlg4
Cask
OR2A25
Dcakd
zgc:113368
F1R2J9
Cacnb2
stnA
cacnb4.2
A0A0R4IX85
dlg1
cask
DLG1
Dlg1
dlg1l

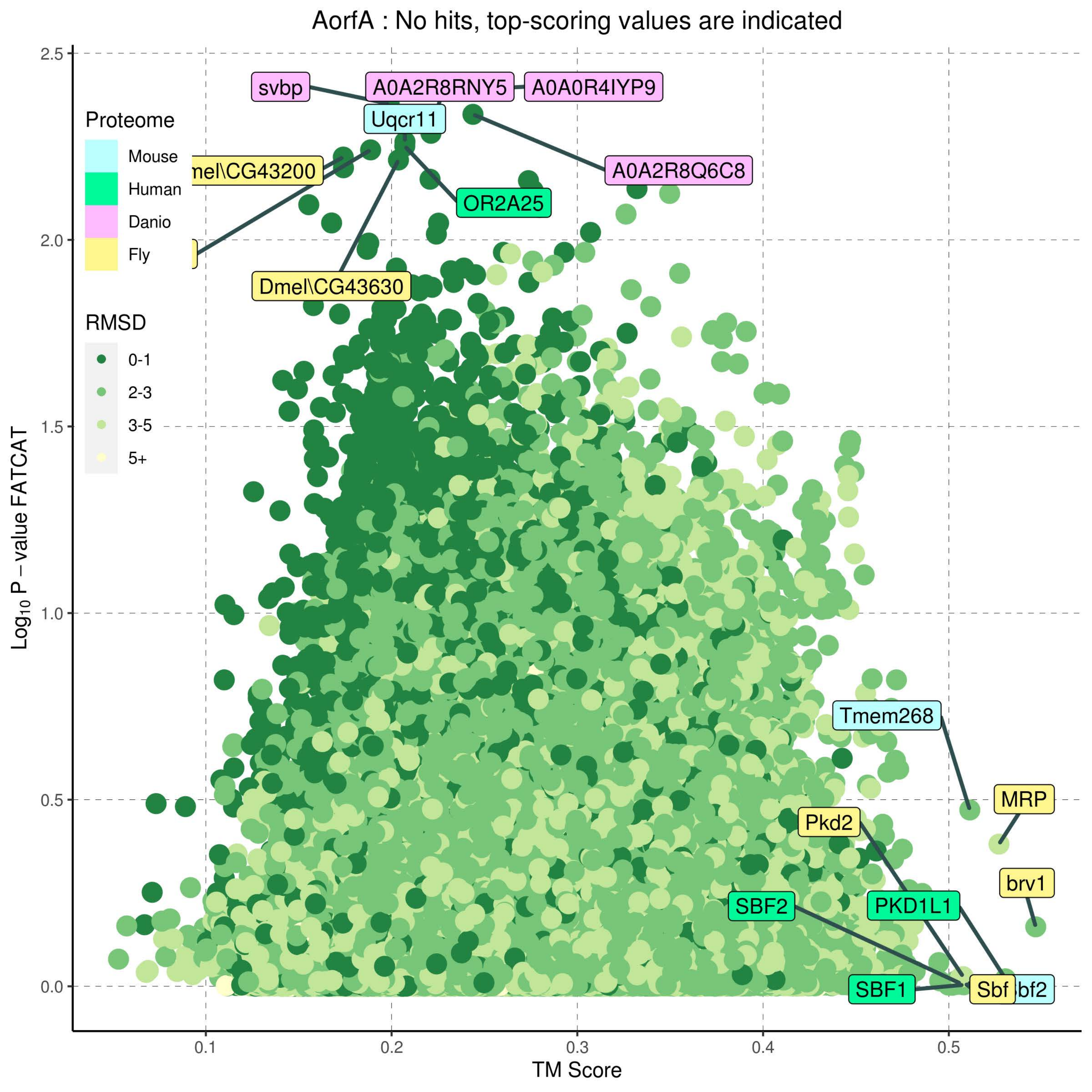

AorfA : No hits, top-scoring values are indicated
Proteome
Mouse
Human
Danio
Fly
RMSD
0-1
2-3
3-5
5+
svbp
A0A2R8RNY5
A0A0R4IYP9
Uqcr11
mel\CG43200
A0A2R8Q6C8
OR2A25
Dmel\CG43630
Tmem268
MRP
Pkd2
brv1
SBF2
PKD1L1
SBF1
Sbf
bf2
2.5
2.0
1.5
1.0
0.5
0.0
0.1
0.2
0.3
0.4
0.5
Log10 P – value FATCAT
TM Score

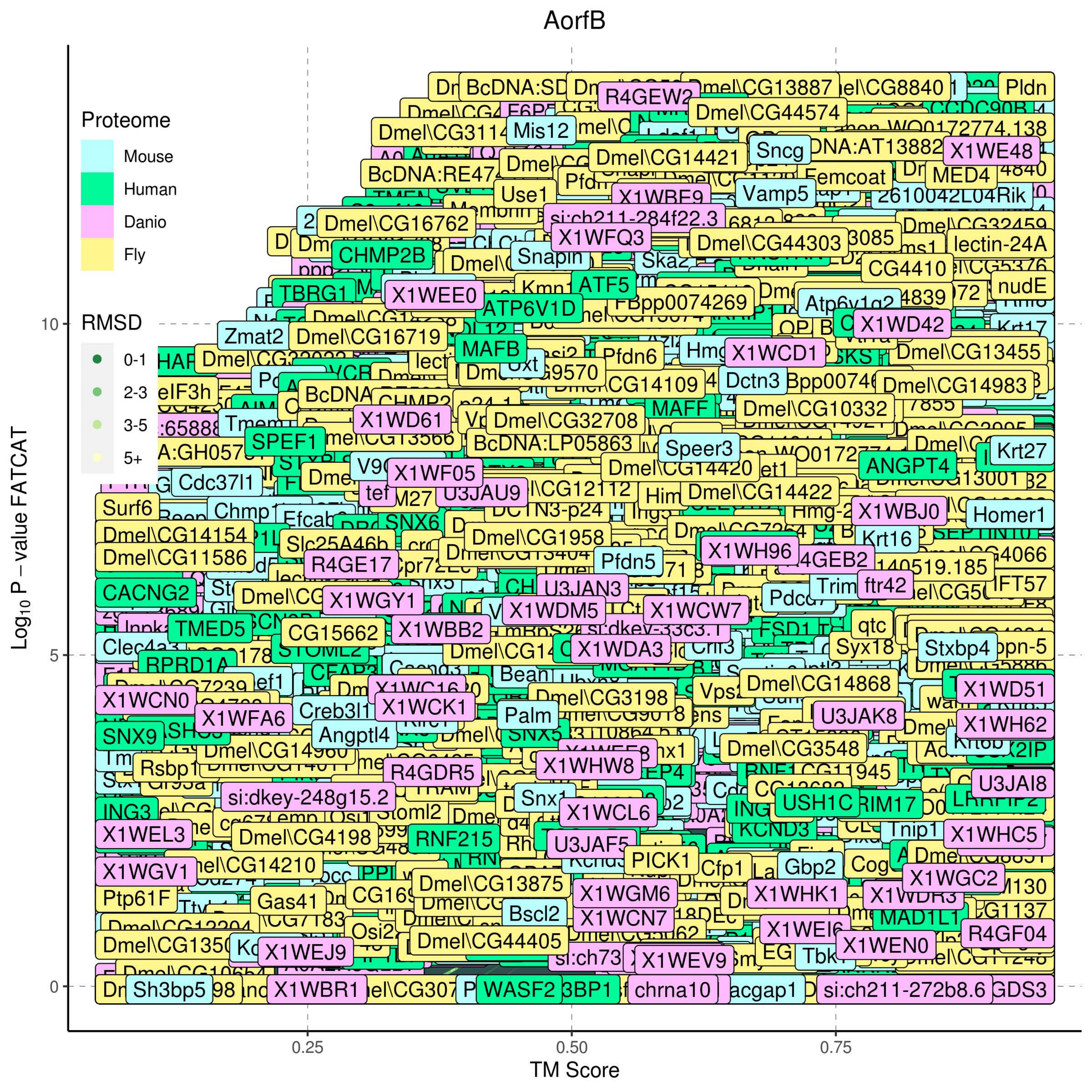
AorfB
Proteome
Mouse
Human
Danio
Fly
RMSD
0-1
2-3
3-5
5+
$Log_{10}$ P – value FATCAT
10
5
0
TM Score
0.25
0.50
0.75

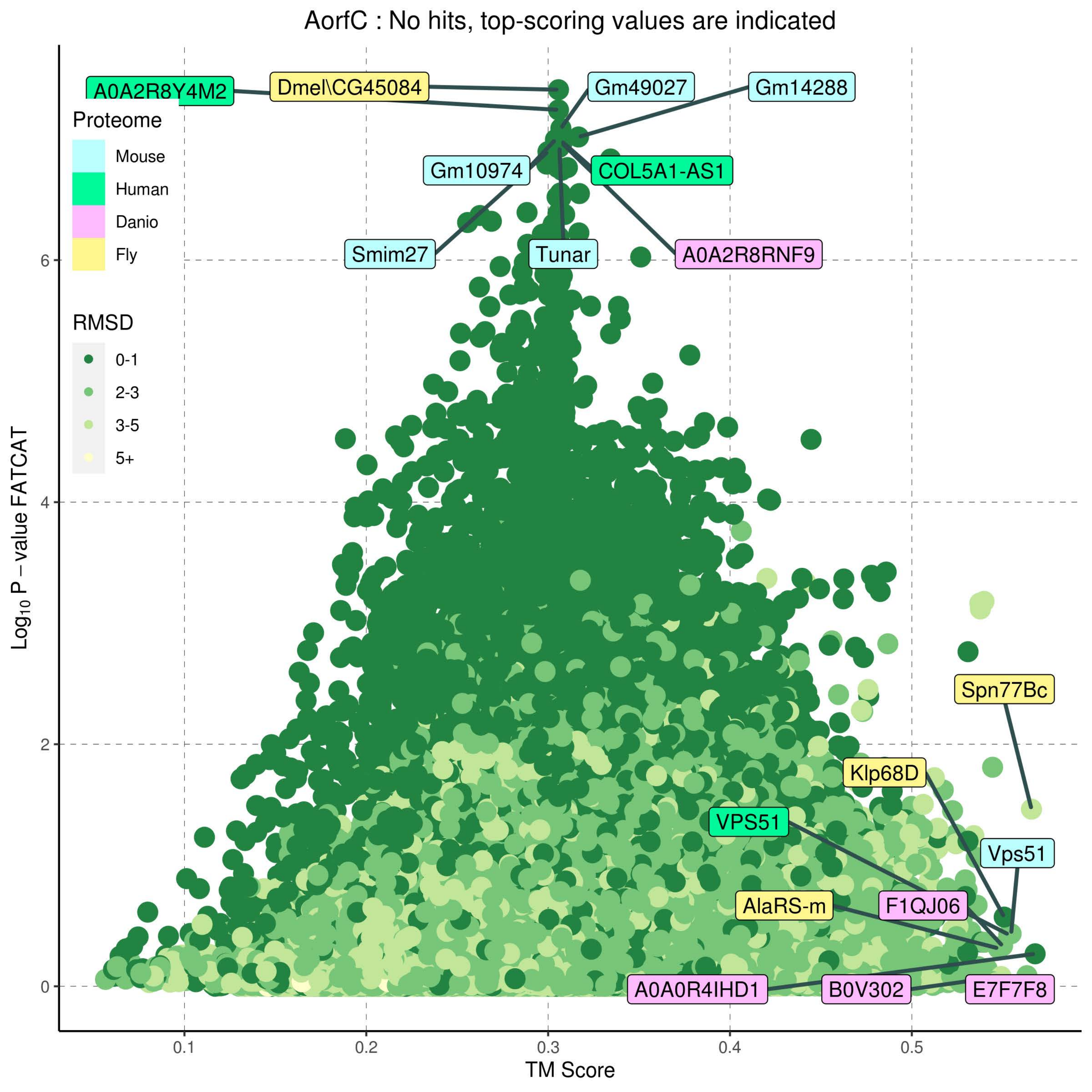
AorfC : No hits, top-scoring values are indicated
Proteome
Mouse
Human
Danio
Fly
RMSD
0-1
2-3
3-5
5+
A0A2R8Y4M2
Dmel\CG45084
Gm49027
Gm14288
Gm10974
COL5A1-AS1
Smim27
Tunar
A0A2R8RNF9
Spn77Bc
Klp68D
VPS51
Vps51
AlaRS-m
F1QJ06
A0A0R4IHD1
B0V302
E7F7F8
Log10 P – value FATCAT
TM Score
0
2
4
6
0.1
0.2
0.3
0.4
0.5

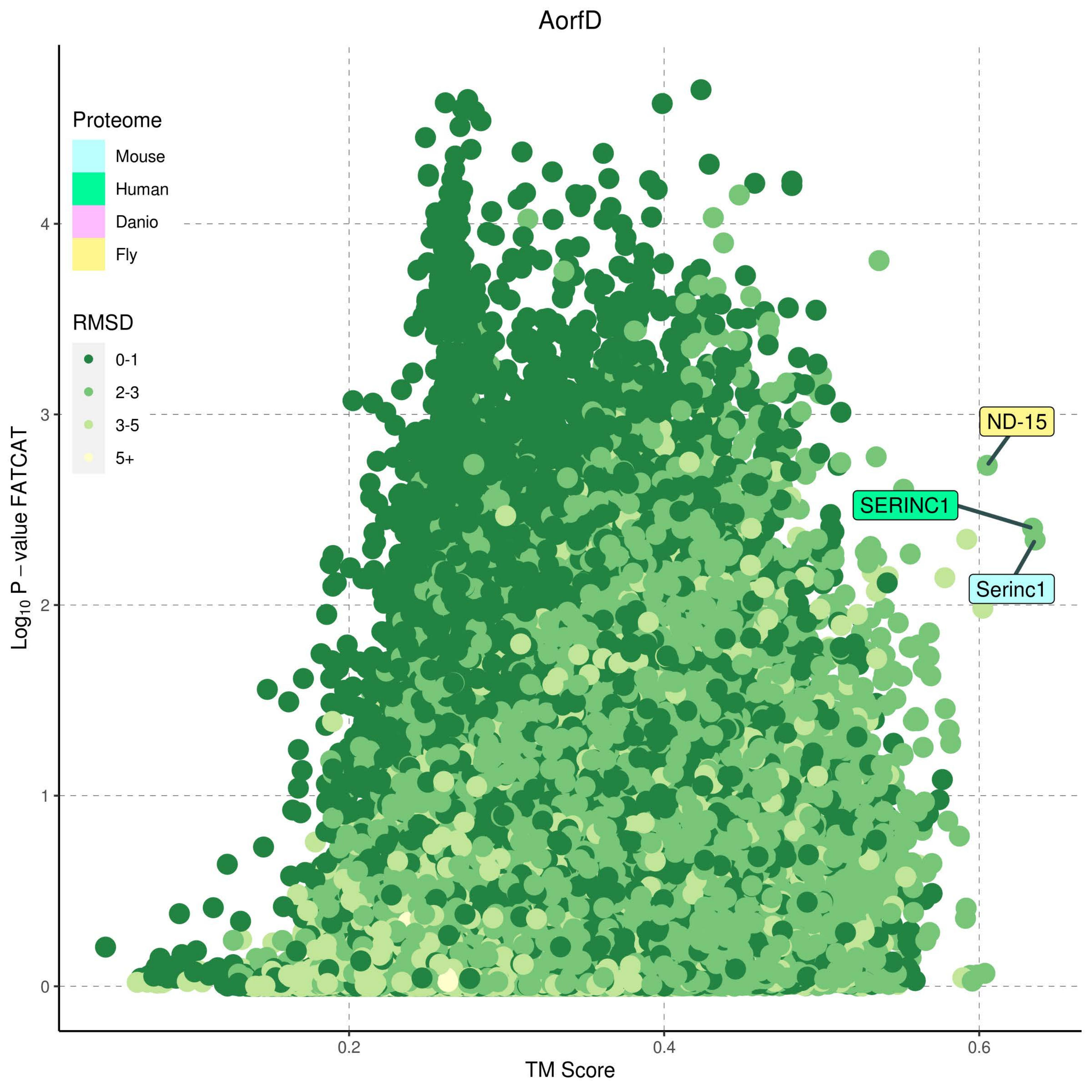
AorfD
Proteome
Mouse
Human
Danio
Fly
RMSD
0-1
2-3
3-5
5+
ND-15
SERINC1
Serinc1
Log10 P − value FATCAT
TM Score
0
1
2
3
4
0.2
0.4
0.6

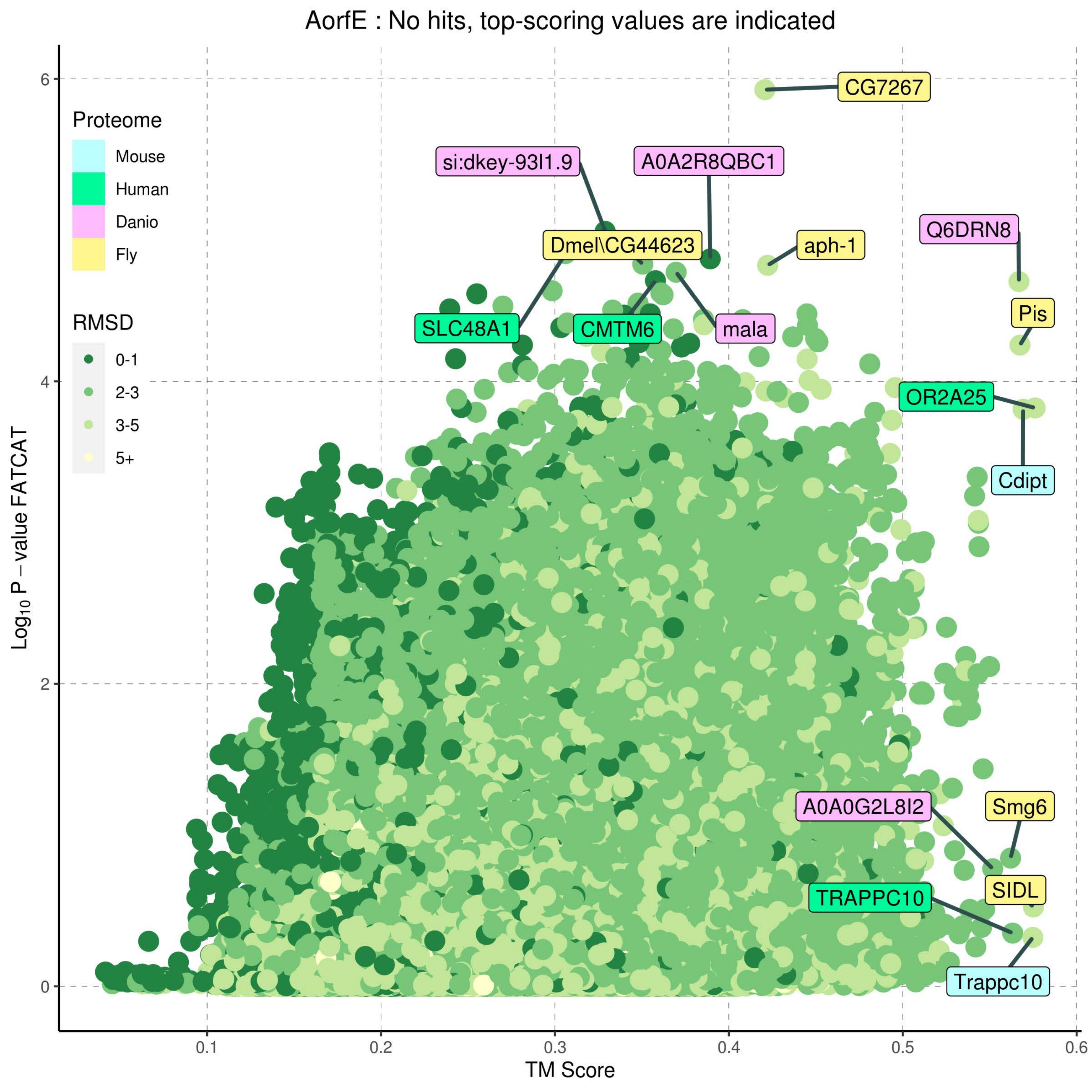
AorfE : No hits, top-scoring values are indicated
Proteome
Mouse
Human
Danio
Fly
RMSD
0-1
2-3
3-5
5+
CG7267
si:dkey-93l1.9
A0A2R8QBC1
Dmel\CG44623
aph-1
Q6DRN8
Pis
SLC48A1
CMTM6
mala
OR2A25
Cdipt
A0A0G2L8I2
Smg6
SIDL
TRAPPC10
Trappc10
Log10 P − value FATCAT
TM Score
0
2
4
6
0.1
0.2
0.3
0.4
0.5
0.6

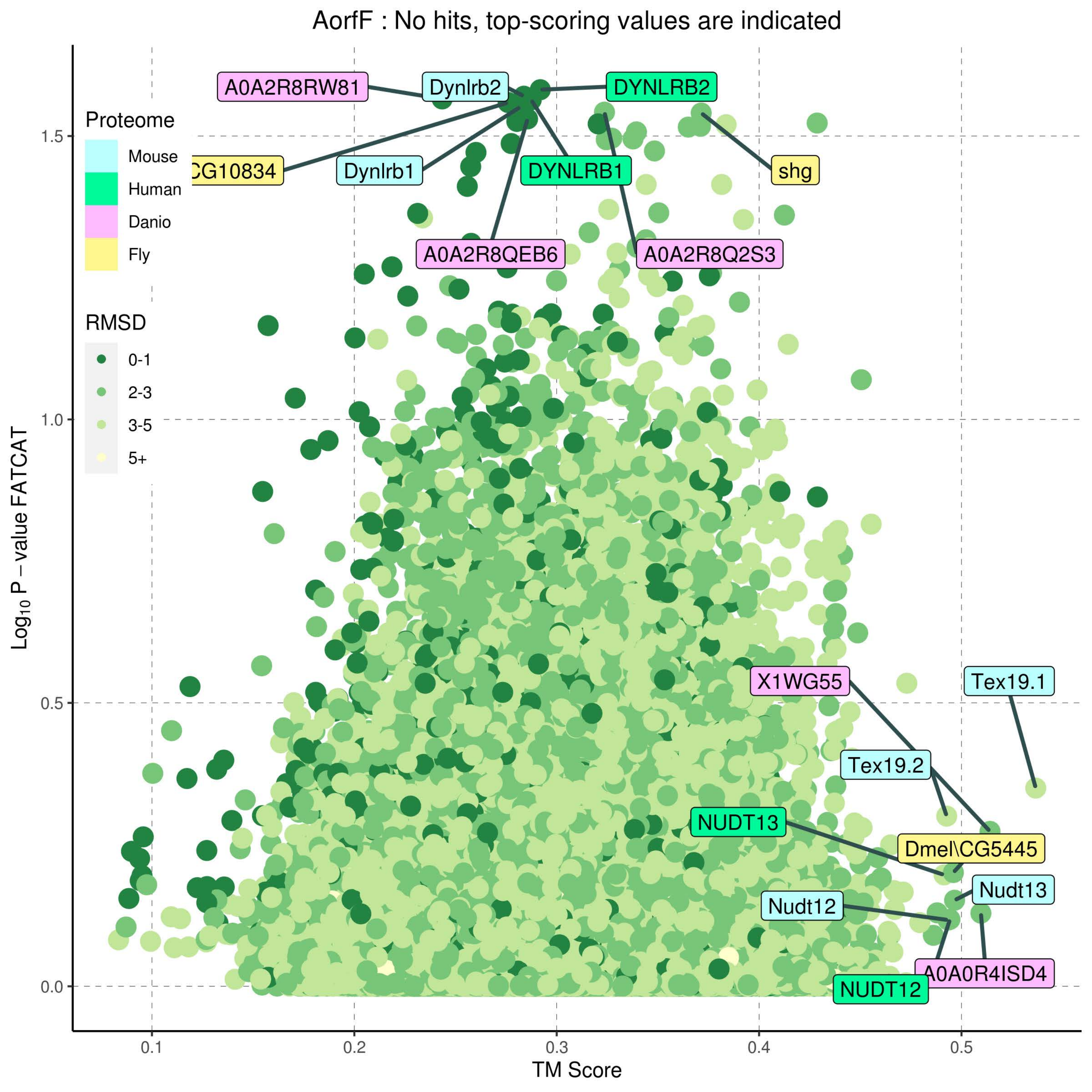

AorfF : No hits, top-scoring values are indicated
Proteome
Mouse
Human
Danio
Fly
RMSD
0-1
2-3
3-5
5+
A0A2R8RW81
Dynlrb2
DYNLRB2
CG10834
Dynlrb1
DYNLRB1
shg
A0A2R8QEB6
A0A2R8Q2S3
X1WG55
Tex19.1
Tex19.2
NUDT13
Dmel\CG5445
Nudt13
Nudt12
A0A0R4ISD4
NUDT12
1.5
1.0
0.5
0.0
0.1
0.2
0.3
0.4
0.5
Log10 P − value FATCAT
TM Score

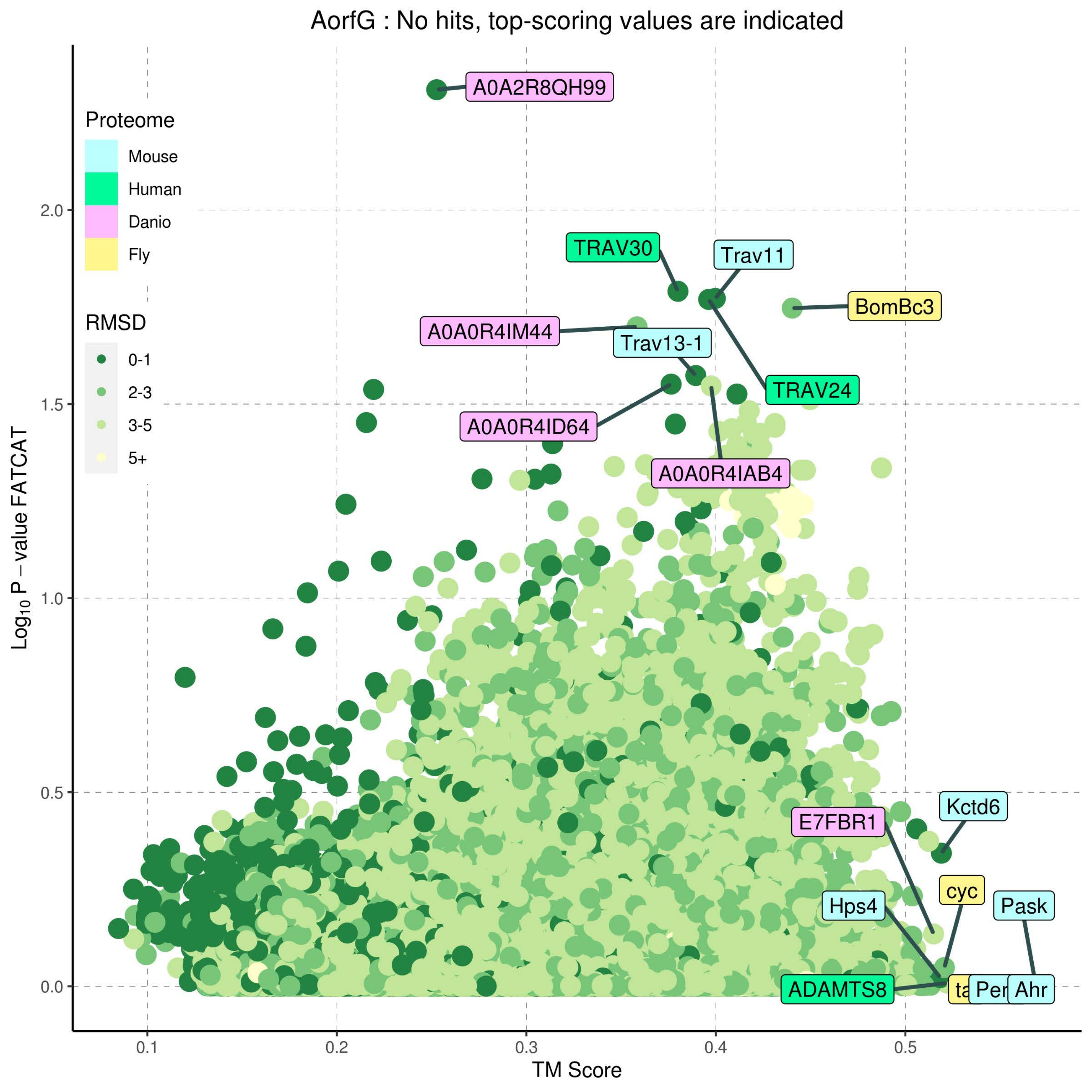
AorfG : No hits, top-scoring values are indicated
Proteome
Mouse
Human
Danio
Fly
RMSD
0-1
2-3
3-5
5+
A0A2R8QH99
TRAV30
Trav11
BomBc3
A0A0R4IM44
Trav13-1
TRAV24
A0A0R4ID64
A0A0R4IAB4
Kctd6
E7FBR1
cyc
Hps4
Pask
ADAMTS8
Per
Ahr
Log10 P − value FATCAT
TM Score
0.0
0.5
1.0
1.5
2.0
0.1
0.2
0.3
0.4
0.5

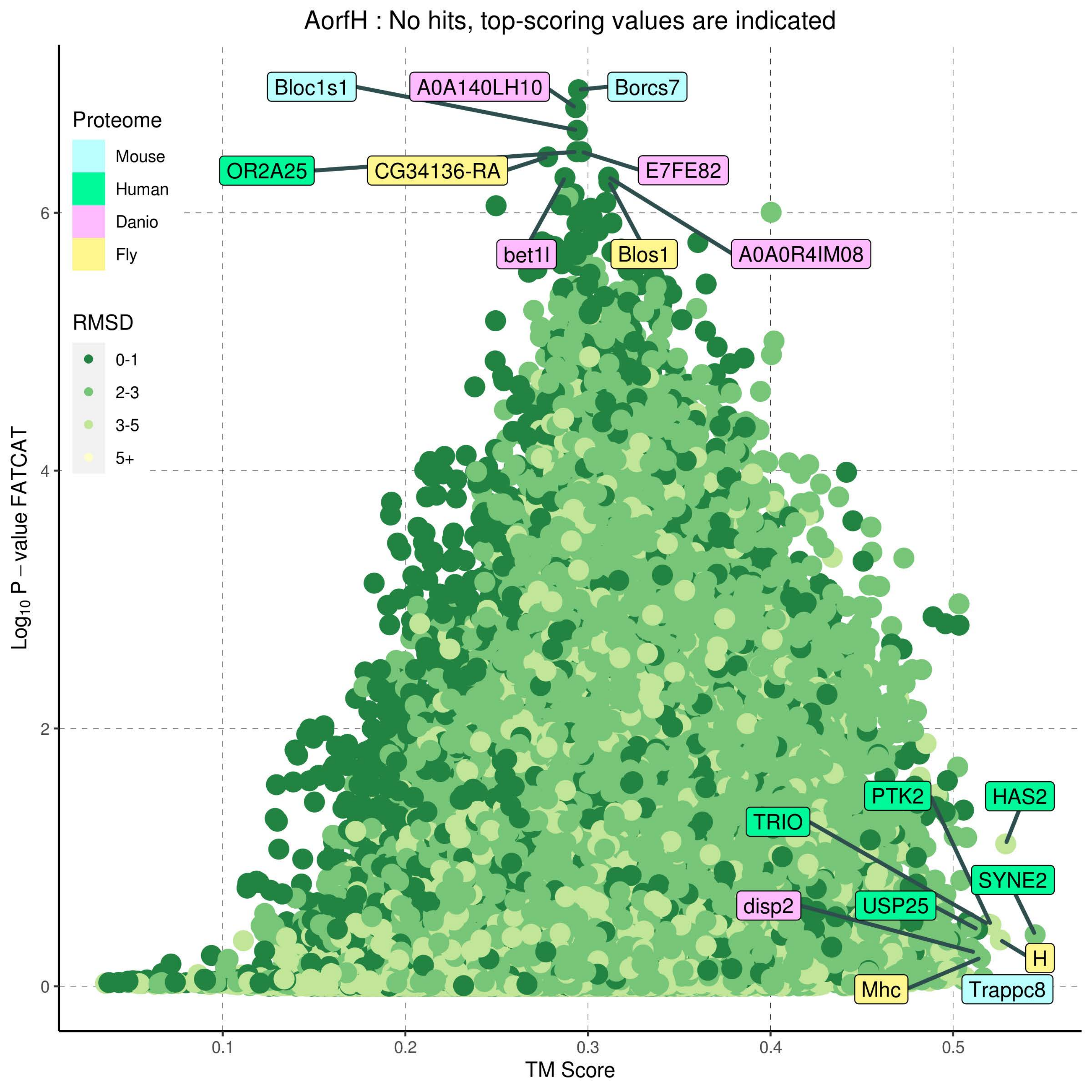
AorfH : No hits, top-scoring values are indicated
Proteome
Mouse
Human
Danio
Fly
RMSD
0-1
2-3
3-5
5+
Bloc1s1
A0A140LH10
Borcs7
OR2A25
CG34136-RA
E7FE82
bet1l
Blos1
A0A0R4IM08
PTK2
HAS2
TRIO
SYNE2
disp2
USP25
H
Mhc
Trappc8
Log10 P − value FATCAT
TM Score
0
2
4
6
0.1
0.2
0.3
0.4
0.5

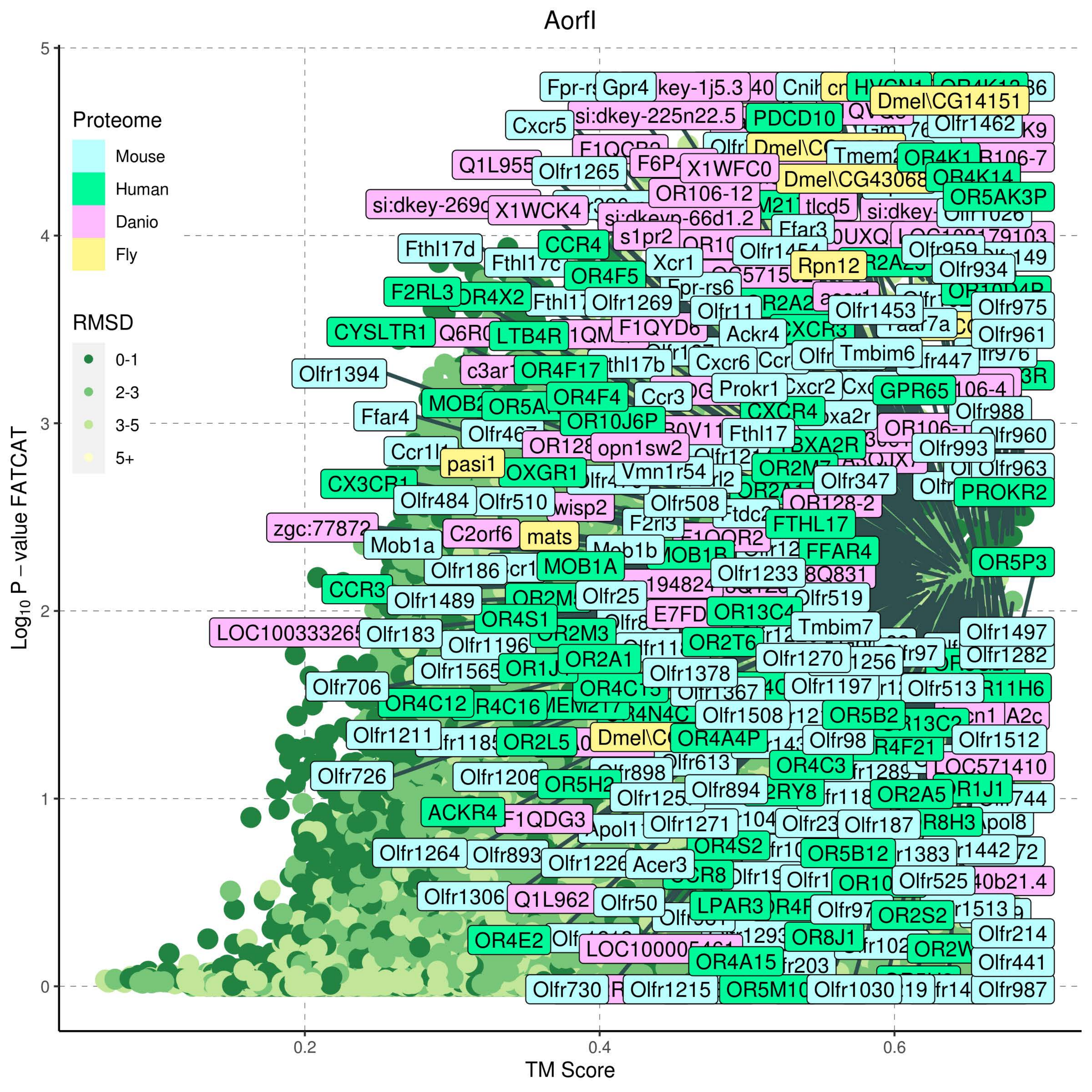

Aorfl
Proteome
Mouse
Human
Danio
Fly
RMSD
0-1
2-3
3-5
5+
Log10 P − value FATCAT
TM Score
0
1
2
3
4
5
0.2
0.4
0.6
Dmel\CG14151
Dmel\CG43068
PDCD10
OR5AK3P
Olfr1462
Olfr1265
Cxcr5
Gpr4
si:dkey-225n22.5
X1WFC0
X1WCK4
OR106-12
Fthl17d
CCR4
Xcr1
OR4F5
Rpn12
Olfr934
F2RL3
Olfr1269
Olfr1453
Olfr975
CYSLTR1
LTB4R
F1QYD6
Ackr4
Olfr961
Olfr1394
OR4F17
Cxcr6
Tmbim6
Prokr1
GPR65
Ffar4
OR4F4
Ccr3
OR10J6P
Fthl17
Olfr988
opn1sw2
pasi1
OXGR1
Vmn1r54
Olfr993
Olfr960
CX3CR1
Olfr484
Olfr510
wisp2
Olfr508
Olfr347
PROKR2
zgc:77872
C2orf6
mats
FTHL17
FFAR4
Mob1a
MOB1A
OR5P3
Olfr186
Olfr1233
CCR3
Olfr1489
Olfr25
Olfr519
OR13C4
OR4S1
OR2M3
Tmbim7
Olfr1497
Olfr183
Olfr1196
OR2T6
Olfr1270
Olfr1282
Olfr1565
OR2A1
Olfr1378
Olfr706
Olfr1197
Olfr513
OR4C12
Olfr1508
OR5B2
Olfr1211
OR2L5
OR4A4P
Olfr98
Olfr1512
Olfr726
Olfr1206
OR4C3
LOC571410
ACKR4
F1QDG3
Olfr894
OR2A5
Olfr1271
Olfr187
Olfr1264
Olfr893
Acer3
OR4S2
OR5B12
Olfr525
Olfr1306
Q1L962
Olfr50
LPAR3
OR2S2
OR4E2
Olfr214
OR8J1
OR4A15
Olfr441
Olfr730
Olfr1215
Olfr1030
Olfr987

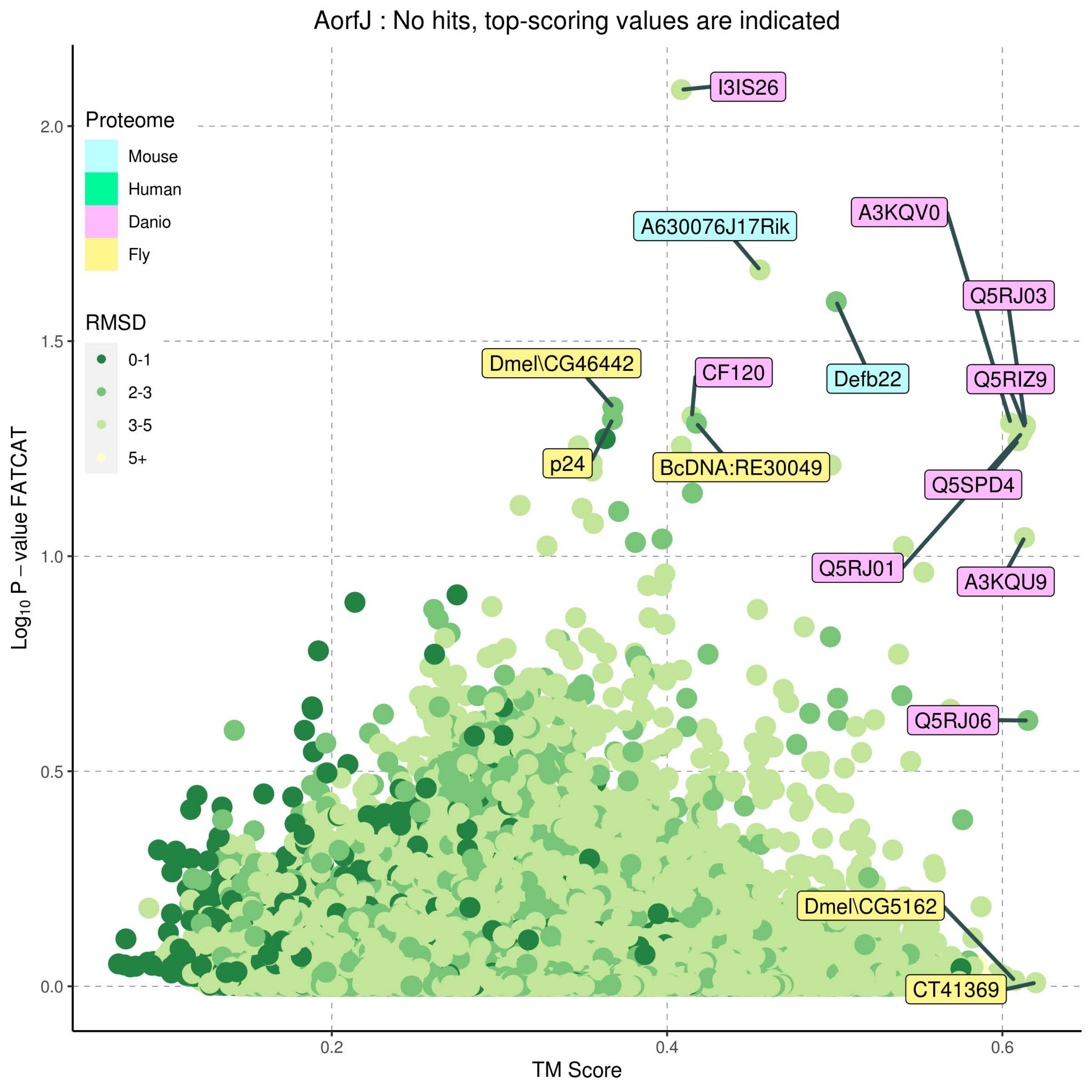
AorfJ : No hits, top-scoring values are indicated
Proteome
Mouse
Human
Danio
Fly
RMSD
0-1
2-3
3-5
5+
I3IS26
A630076J17Rik
A3KQV0
Q5RJ03
Defb22
Q5RIZ9
Dmel\CG46442
CF120
p24
BcDNA:RE30049
Q5SPD4
Q5RJ01
A3KQU9
Q5RJ06
Dmel\CG5162
CT41369
2.0
1.5
1.0
0.5
0.0
0.2
0.4
0.6
Log10 P − value FATCAT
TM Score

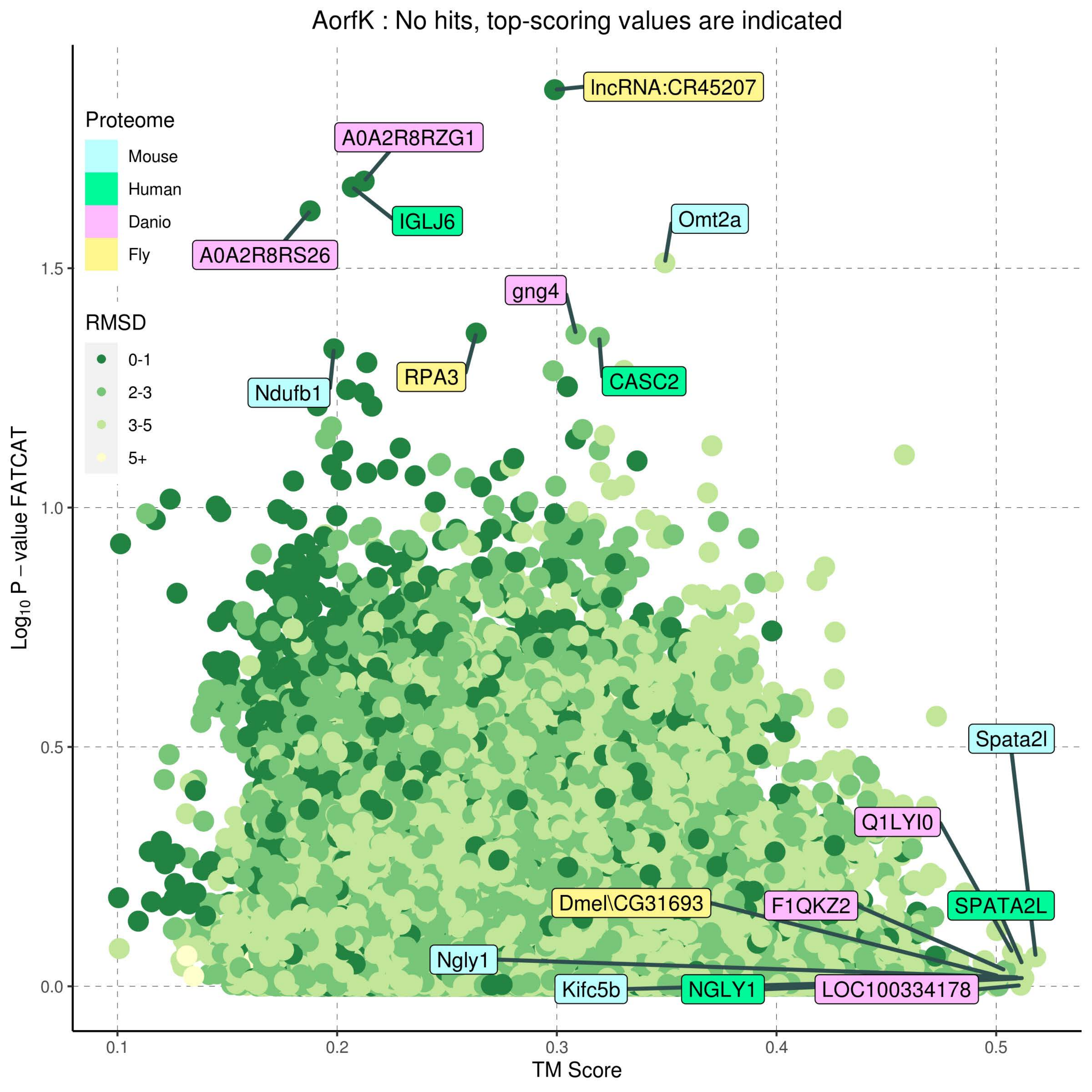
AorfK : No hits, top-scoring values are indicated
Proteome
Mouse
Human
Danio
Fly
RMSD
0-1
2-3
3-5
5+
lncRNA:CR45207
A0A2R8RZG1
IGLJ6
A0A2R8RS26
Omt2a
gng4
RPA3
CASC2
Ndufb1
Spata2l
Q1LYI0
Dmel\CG31693
F1QKZ2
SPATA2L
Ngly1
Kifc5b
NGLY1
LOC100334178
Log10 P − value FATCAT
TM Score
0.0
0.5
1.0
1.5
0.1
0.2
0.3
0.4
0.5

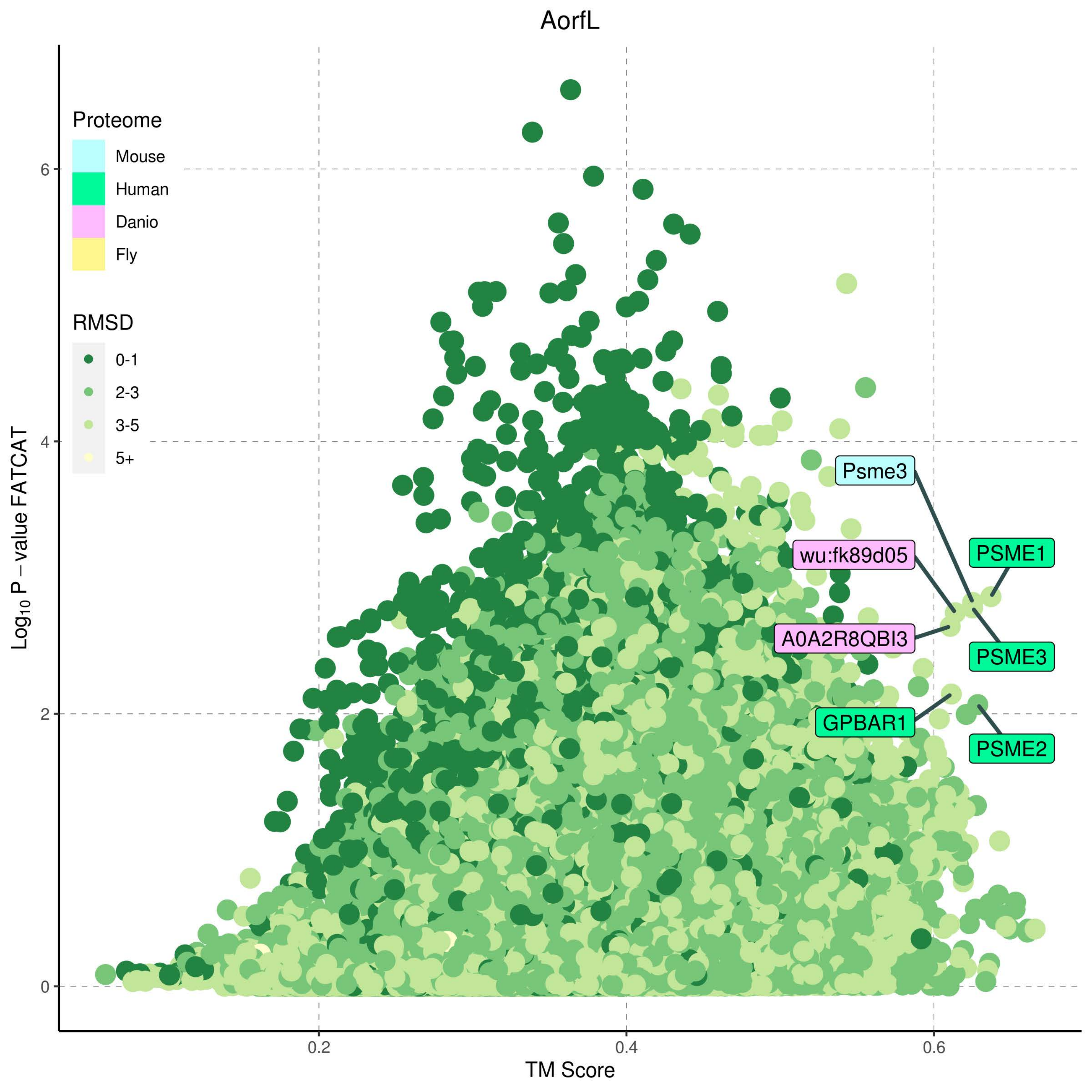

AorfL
Proteome
Mouse
Human
Danio
Fly
RMSD
0-1
2-3
3-5
5+
Psme3
wu:fk89d05
PSME1
A0A2R8QBI3
PSME3
GPBAR1
PSME2
Log10 P − value FATCAT
TM Score
0
2
4
6
0.2
0.4
0.6

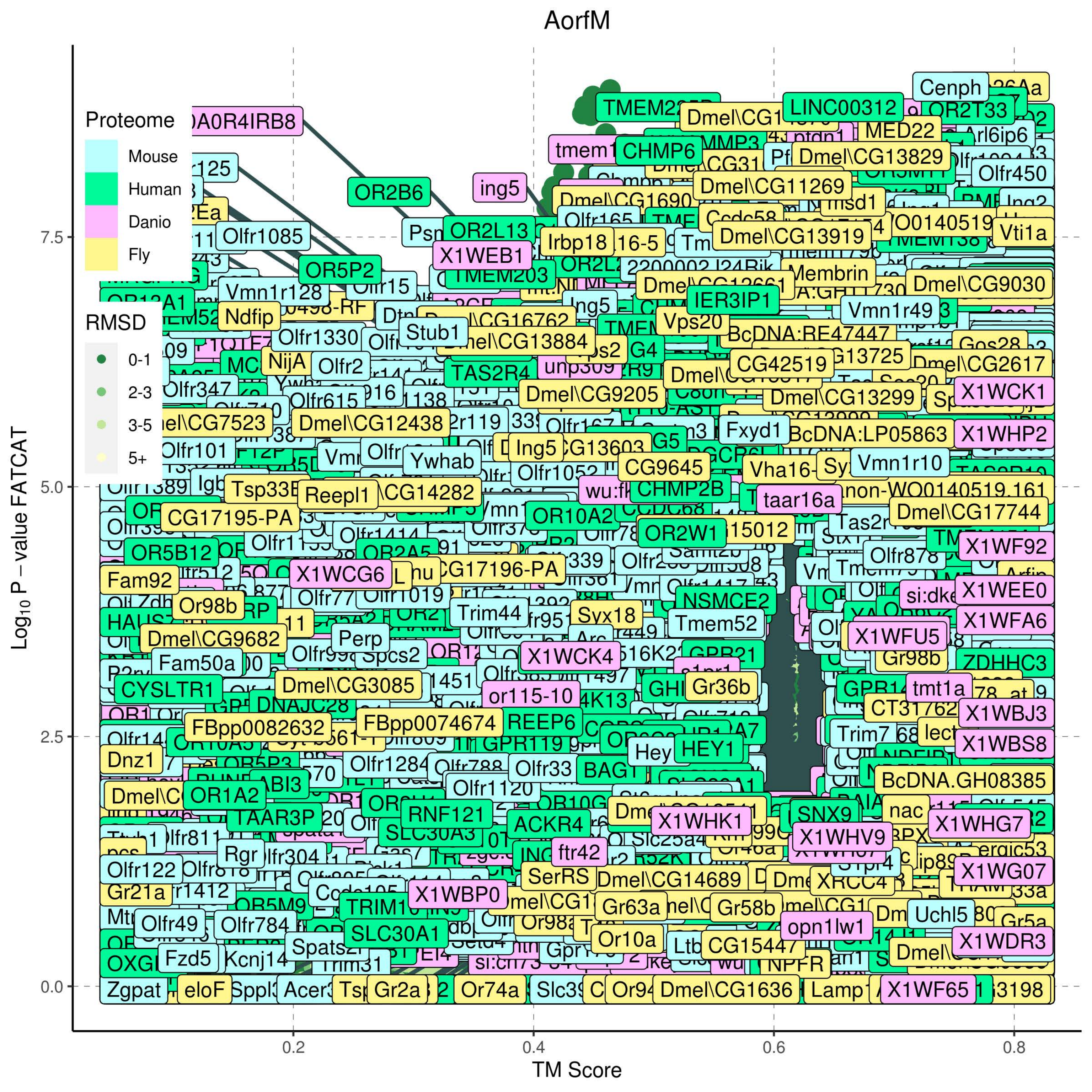

AorfM
Proteome
Mouse
Human
Danio
Fly
RMSD
0-1
2-3
3-5
5+
Log10 P − value FATCAT
7.5
5.0
2.5
0.0
0.2
0.4
0.6
0.8
TM Score

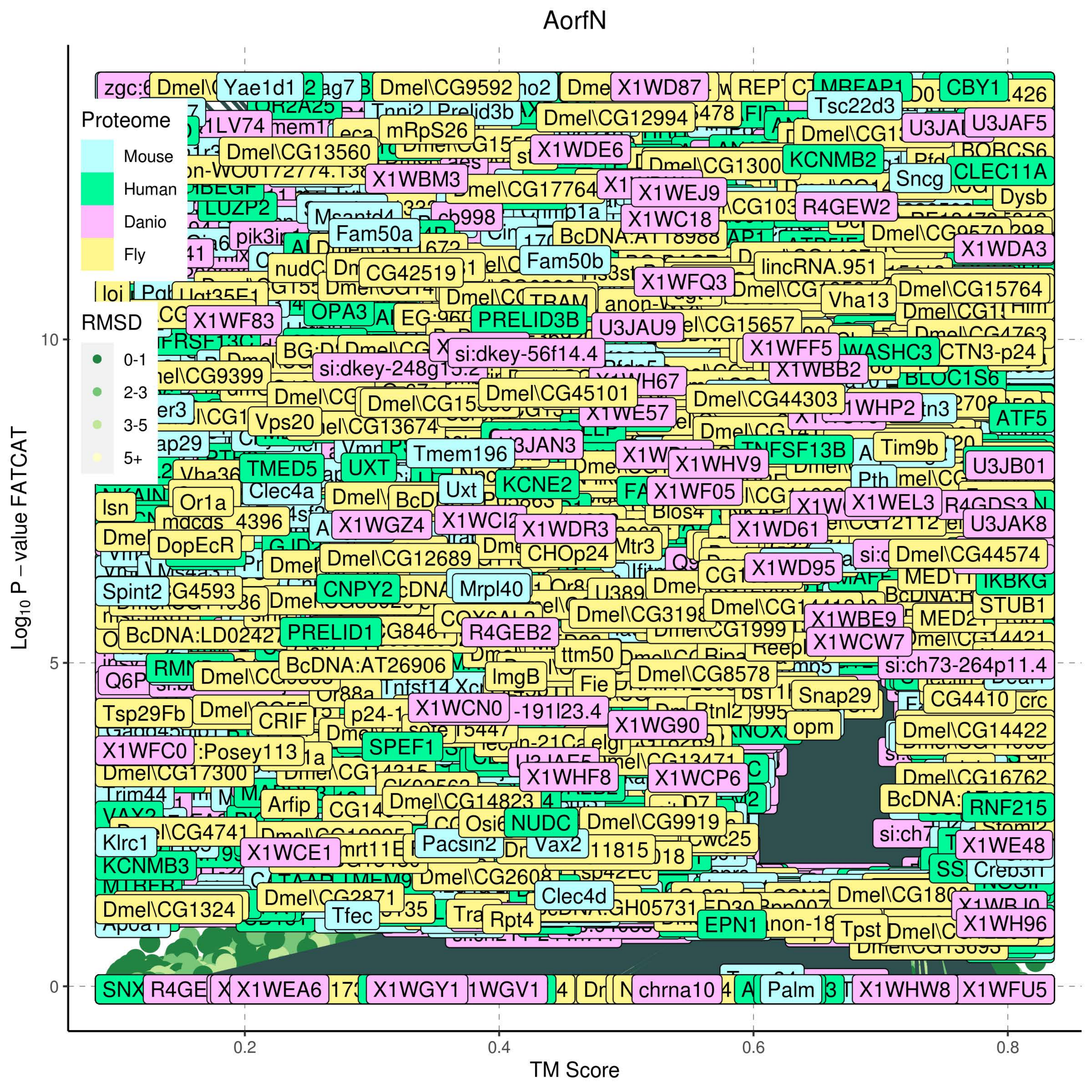

AorfN
Proteome
Mouse
Human
Danio
Fly
RMSD
0-1
2-3
3-5
5+
$Log_{10}$ P – value FATCAT
10
5
0
0.2
0.4
0.6
0.8
TM Score

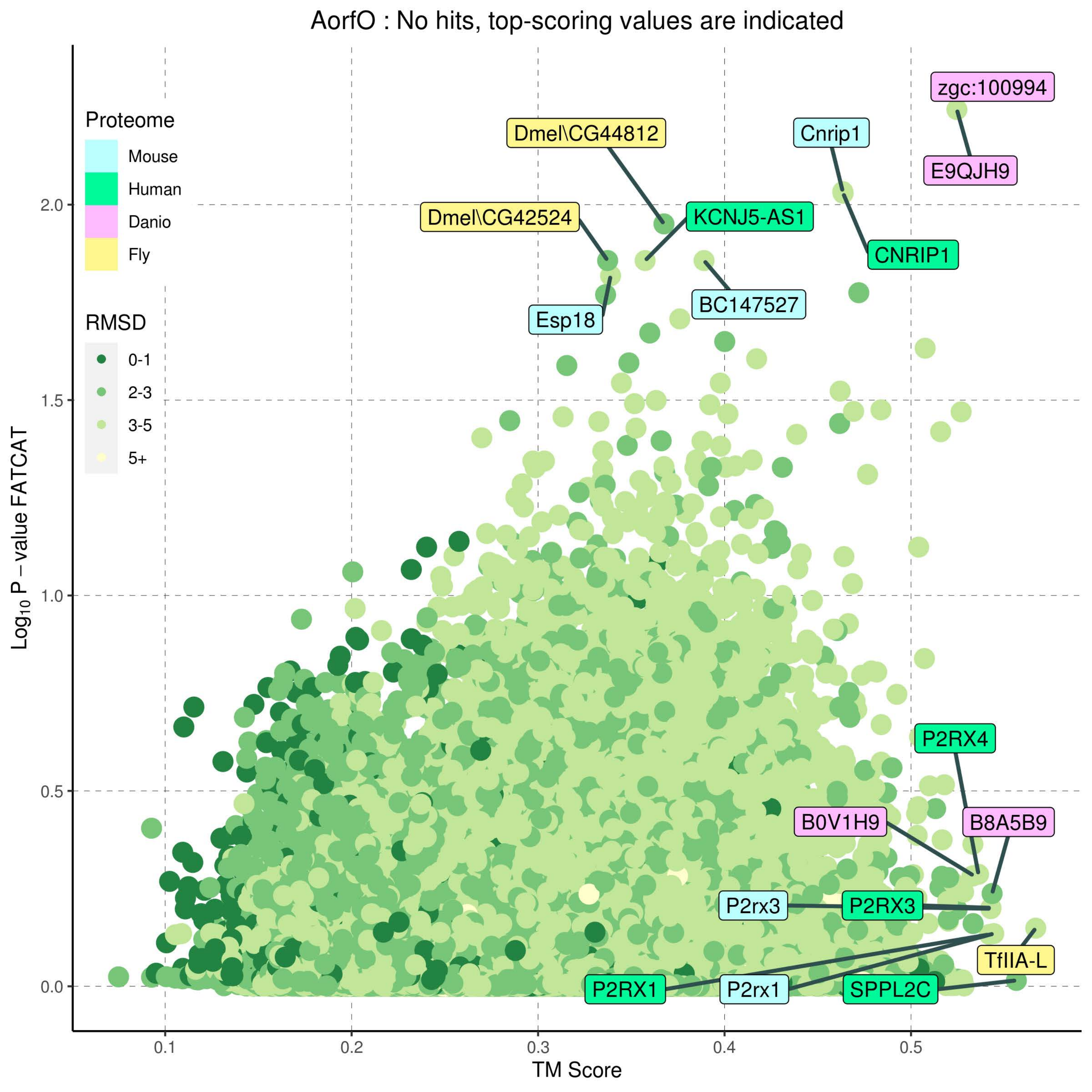

AorfO : No hits, top-scoring values are indicated
Proteome
Mouse
Human
Danio
Fly
RMSD
0-1
2-3
3-5
5+
zgc:100994
E9QJH9
Cnrip1
CNRIP1
Dmel\CG44812
KCNJ5-AS1
Dmel\CG42524
Esp18
BC147527
P2RX4
B0V1H9
B8A5B9
P2rx3
P2RX3
TfIIA-L
P2RX1
P2rx1
SPPL2C
Log10 P − value FATCAT
TM Score
0.0
0.5
1.0
1.5
2.0
0.1
0.2
0.3
0.4
0.5

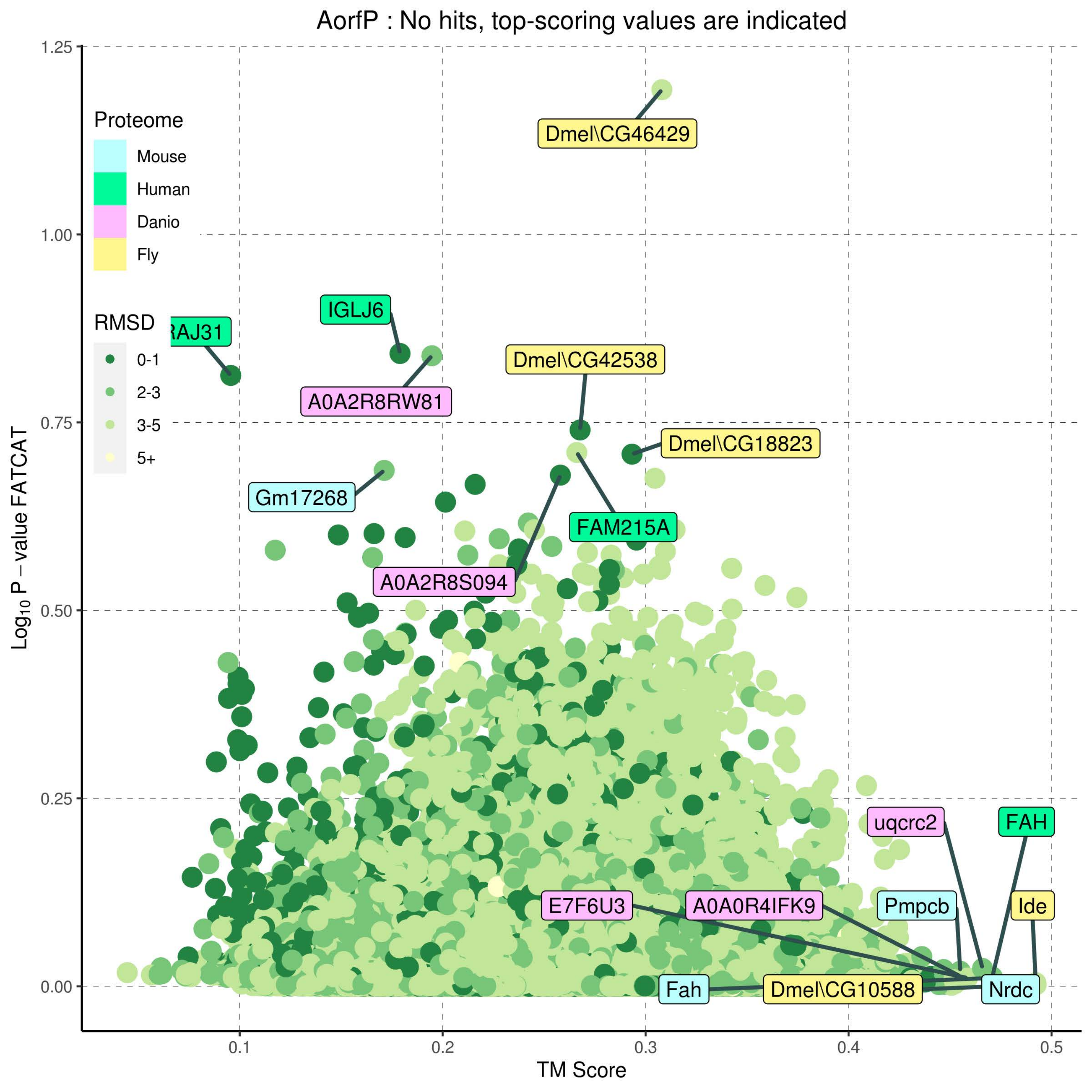

AorfP : No hits, top-scoring values are indicated
Proteome
Mouse
Human
Danio
Fly
RMSD
0-1
2-3
3-5
5+
Dmel\CG46429
RAJ31
IGLJ6
A0A2R8RW81
Dmel\CG42538
Dmel\CG18823
Gm17268
FAM215A
A0A2R8S094
uqcrc2
FAH
E7F6U3
A0A0R4IFK9
Pmpcb
Ide
Fah
Dmel\CG10588
Nrdc
1.25
1.00
0.75
0.50
0.25
0.00
0.1
0.2
0.3
0.4
0.5
TM Score
$Log_{10}$ P – value FATCAT

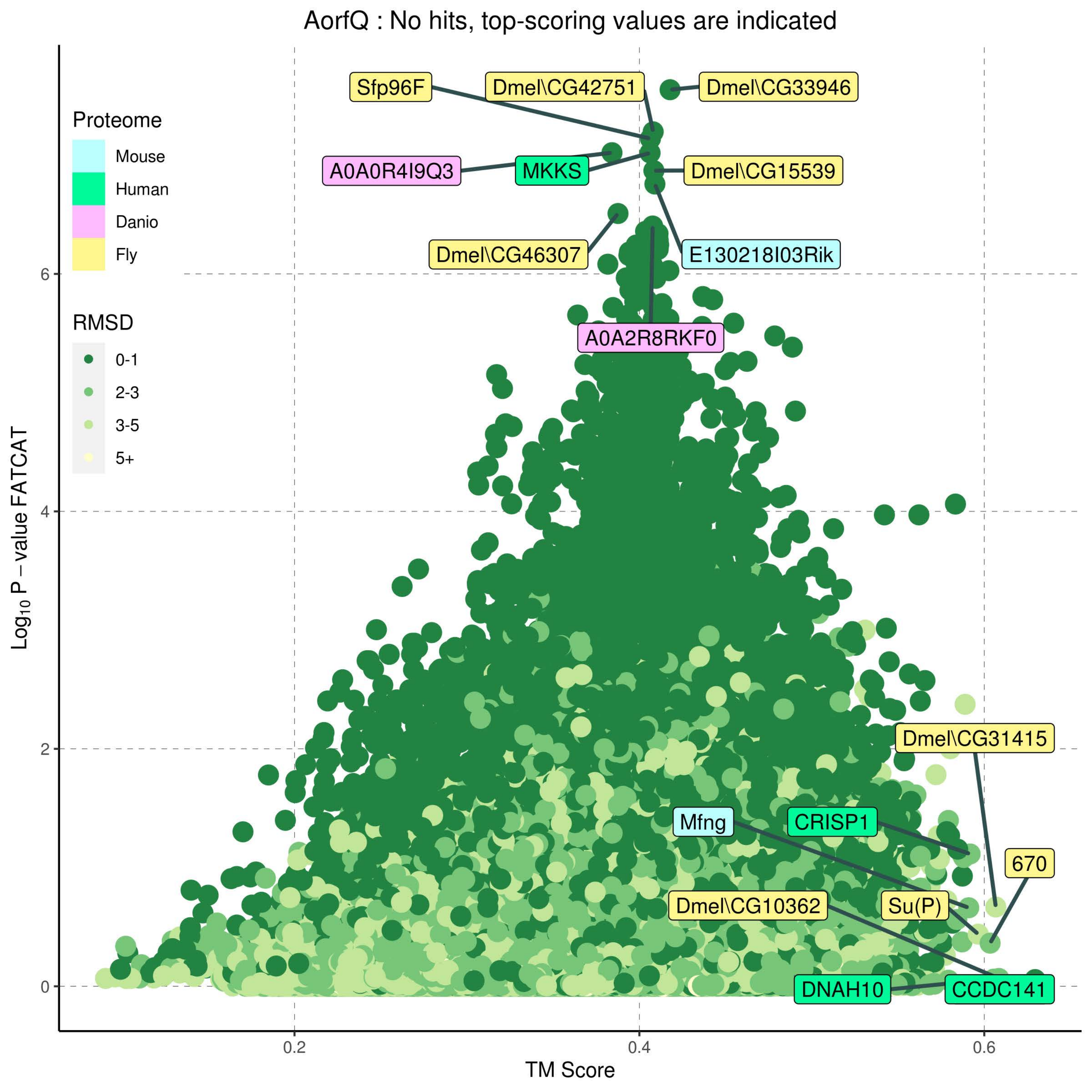

AorfQ : No hits, top-scoring values are indicated
Proteome
Mouse
Human
Danio
Fly
RMSD
0-1
2-3
3-5
5+
Sfp96F
Dmel\CG42751
Dmel\CG33946
A0A0R4I9Q3
MKKS
Dmel\CG15539
Dmel\CG46307
E130218I03Rik
A0A2R8RKF0
Dmel\CG31415
Mfng
CRISP1
670
Dmel\CG10362
Su(P)
DNAH10
CCDC141
Log10 P − value FATCAT
TM Score
0
2
4
6
0.2
0.4
0.6

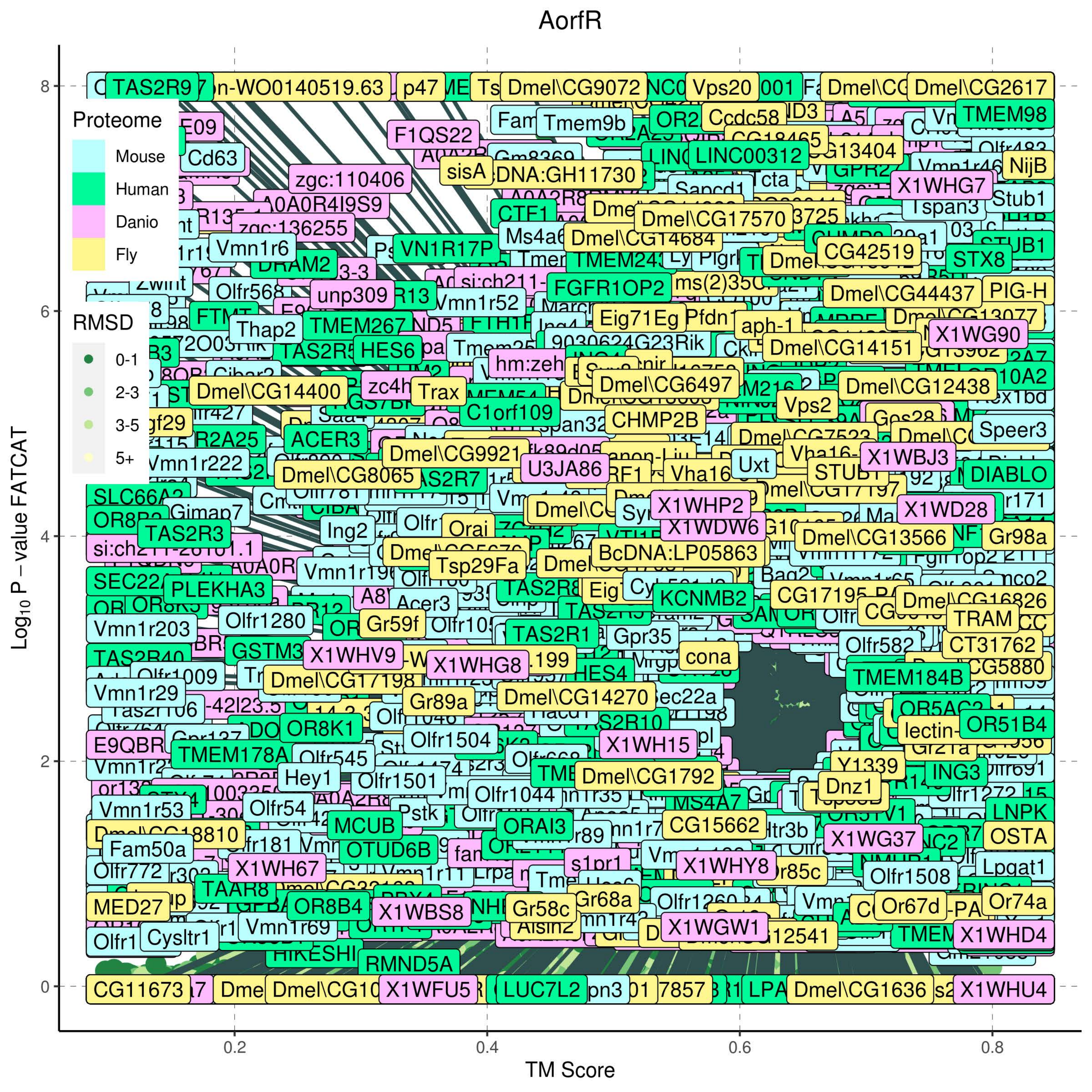

AorfR
Proteome
Mouse
Human
Danio
Fly
RMSD
0-1
2-3
3-5
5+
$Log_{10}$ P – value FATCAT
TM Score
0
2
4
6
8
0.2
0.4
0.6
0.8

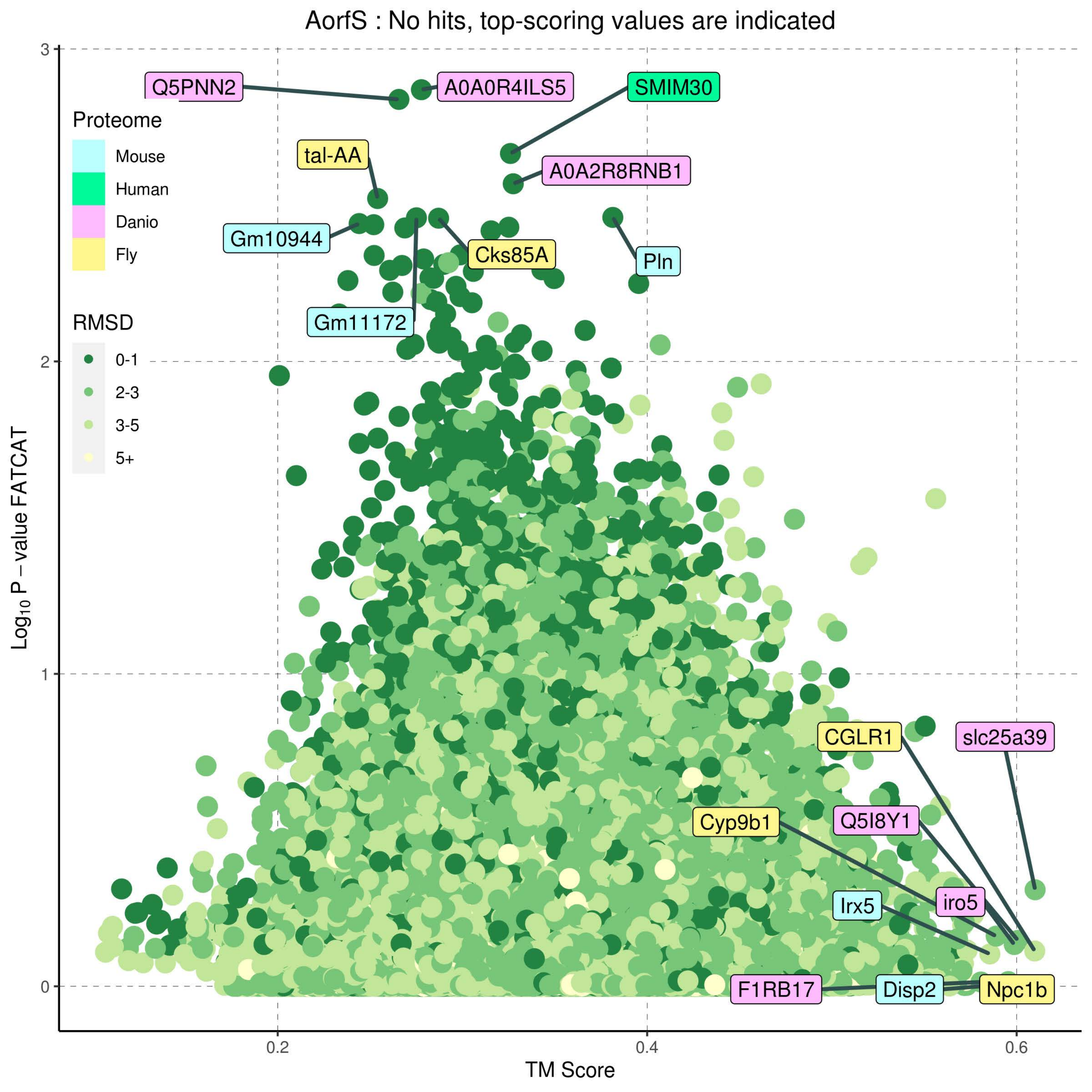

AorfS : No hits, top-scoring values are indicated
Proteome
Mouse
Human
Danio
Fly
RMSD
0-1
2-3
3-5
5+
Q5PNN2
A0A0R4ILS5
SMIM30
tal-AA
A0A2R8RNB1
Gm10944
Cks85A
Pln
Gm11172
CGLR1
slc25a39
Cyp9b1
Q5I8Y1
iro5
Irx5
F1RB17
Disp2
Npc1b
Log10 P – value FATCAT
TM Score
0
1
2
3
0.2
0.4
0.6

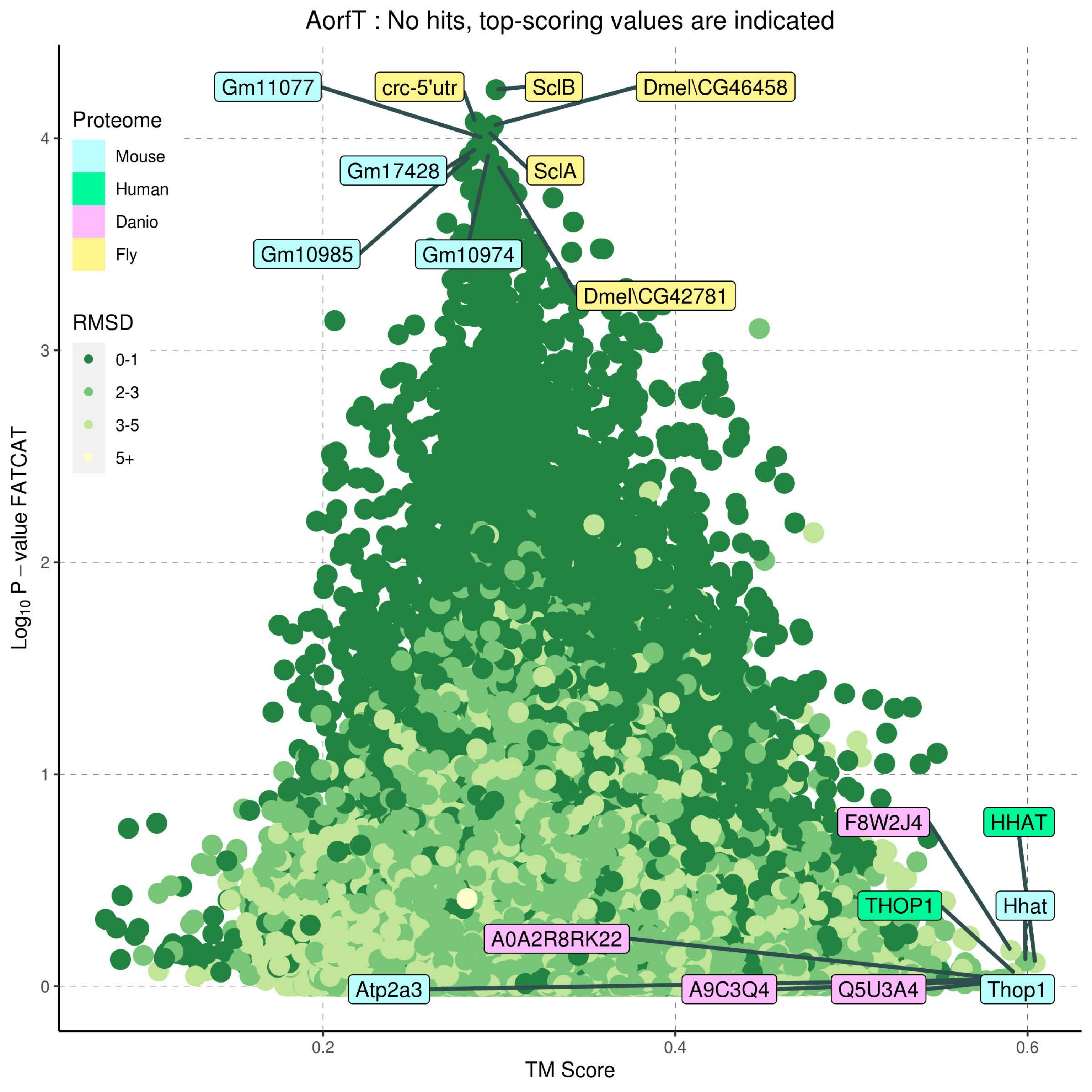
AorfT : No hits, top-scoring values are indicated
Proteome
Mouse
Human
Danio
Fly
RMSD
0-1
2-3
3-5
5+
Gm11077
crc-5'utr
SclB
Dmel\CG46458
Gm17428
SclA
Gm10985
Gm10974
Dmel\CG42781
F8W2J4
HHAT
THOP1
Hhat
A0A2R8RK22
Atp2a3
A9C3Q4
Q5U3A4
Thop1
$Log_{10}$ P – value FATCAT
TM Score
0
1
2
3
4
0.2
0.4
0.6

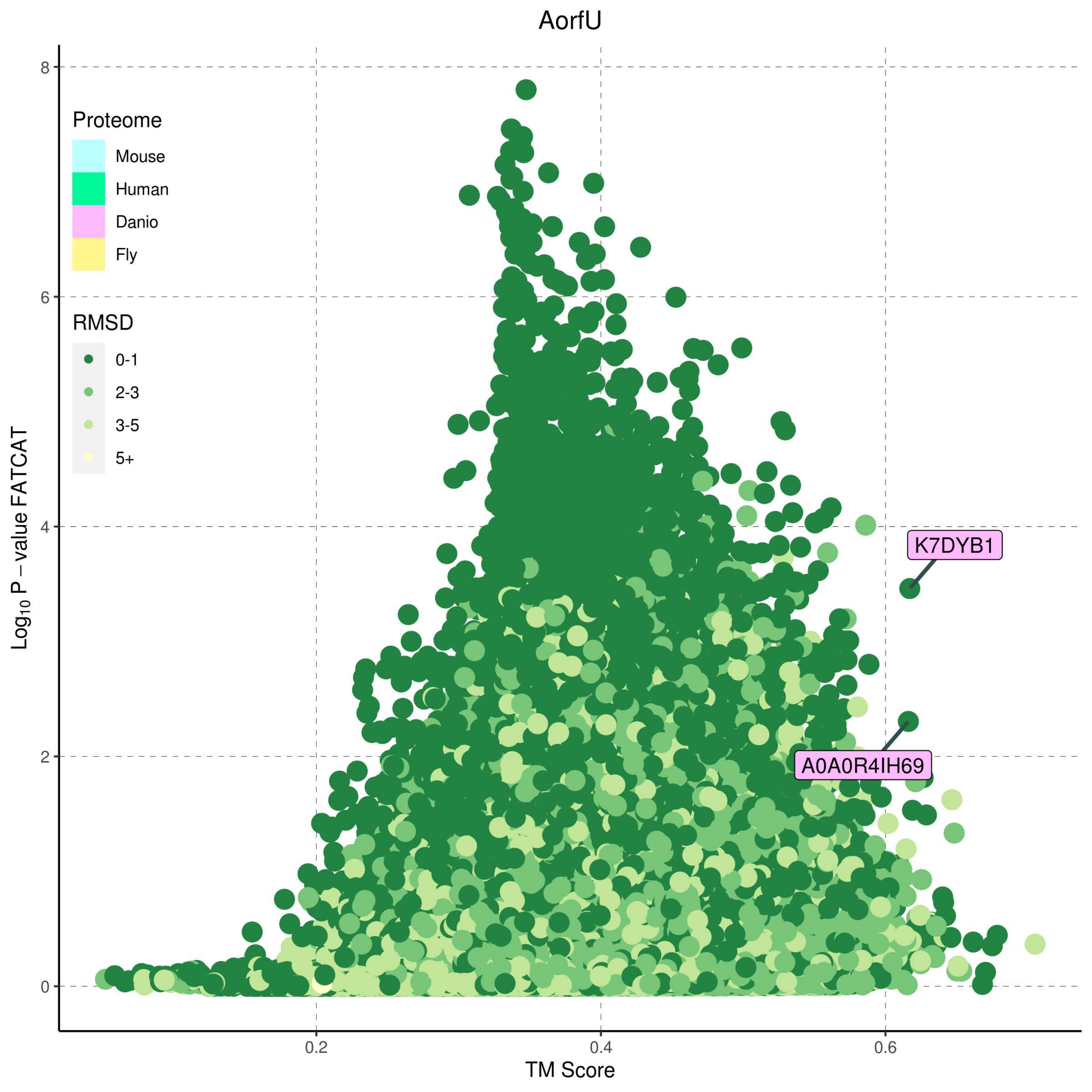
AorfU
Proteome
Mouse
Human
Danio
Fly
RMSD
0-1
2-3
3-5
5+
K7DYB1
A0A0R4IH69
Log10 P – value FATCAT
TM Score
0
2
4
6
8
0.2
0.4
0.6

B1

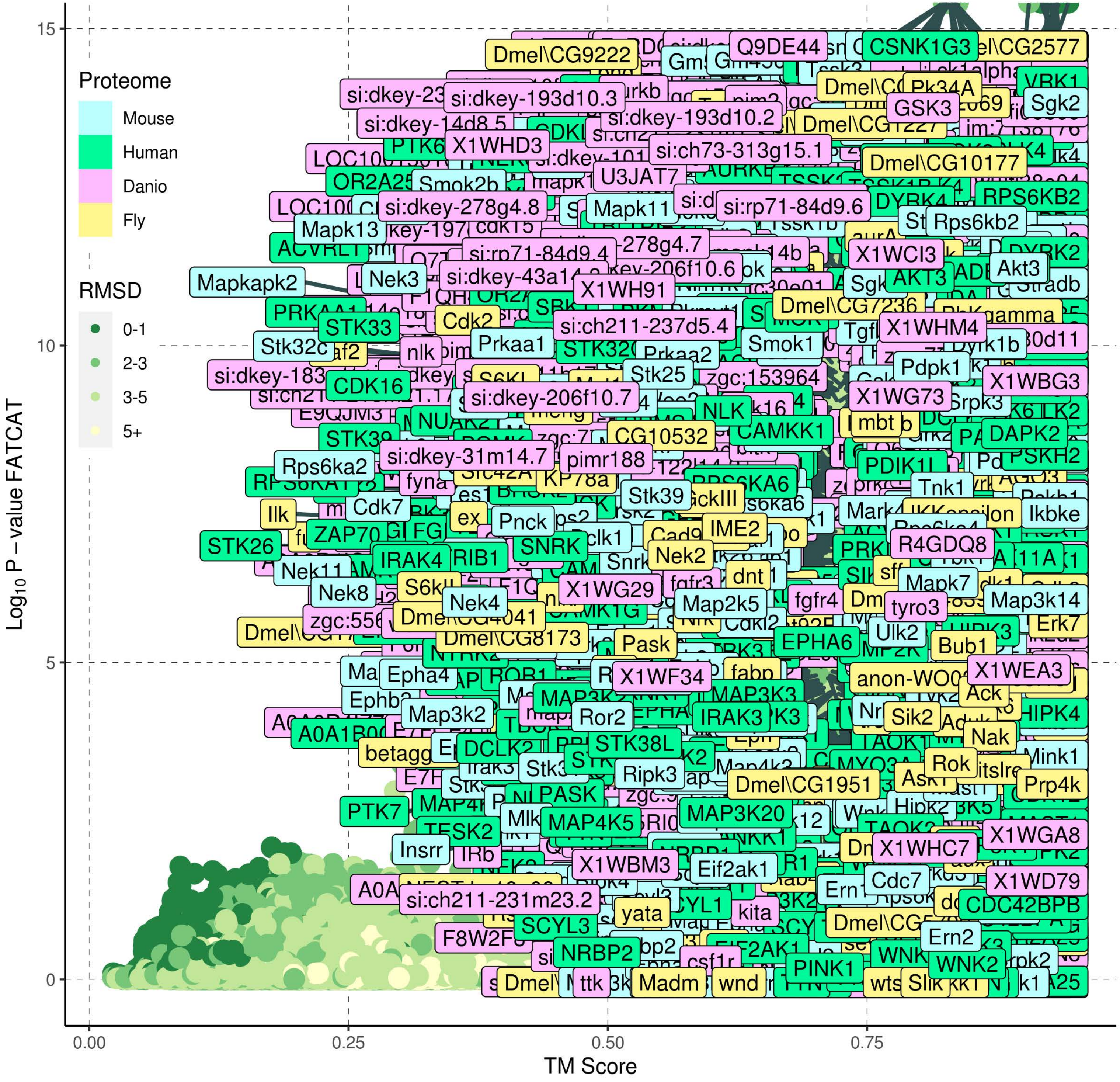
Proteome
Mouse
Human
Danio
Fly
RMSD
0-1
2-3
3-5
5+
Log10 P − value FATCAT
TM Score
0
5
10
15
0.00
0.25
0.50
0.75

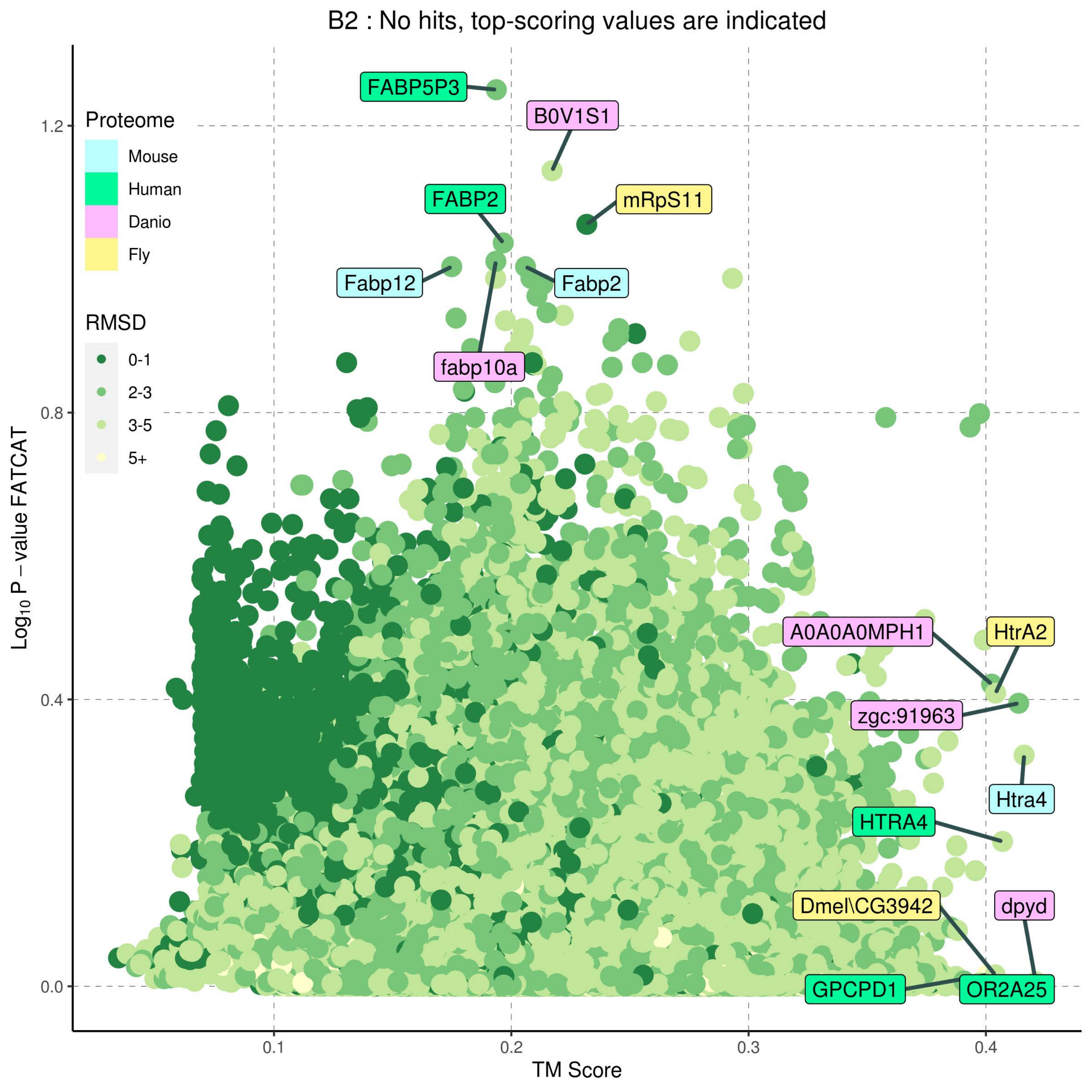

B2 : No hits, top-scoring values are indicated
Proteome
Mouse
Human
Danio
Fly
RMSD
0-1
2-3
3-5
5+
FABP5P3
B0V1S1
FABP2
mRpS11
Fabp12
Fabp2
fabp10a
A0A0A0MPH1
HtrA2
zgc:91963
Htra4
HTRA4
Dmel\CG3942
dpyd
GPCPD1
OR2A25
1.2
0.8
0.4
0.0
0.1
0.2
0.3
0.4
Log10 P − value FATCAT
TM Score

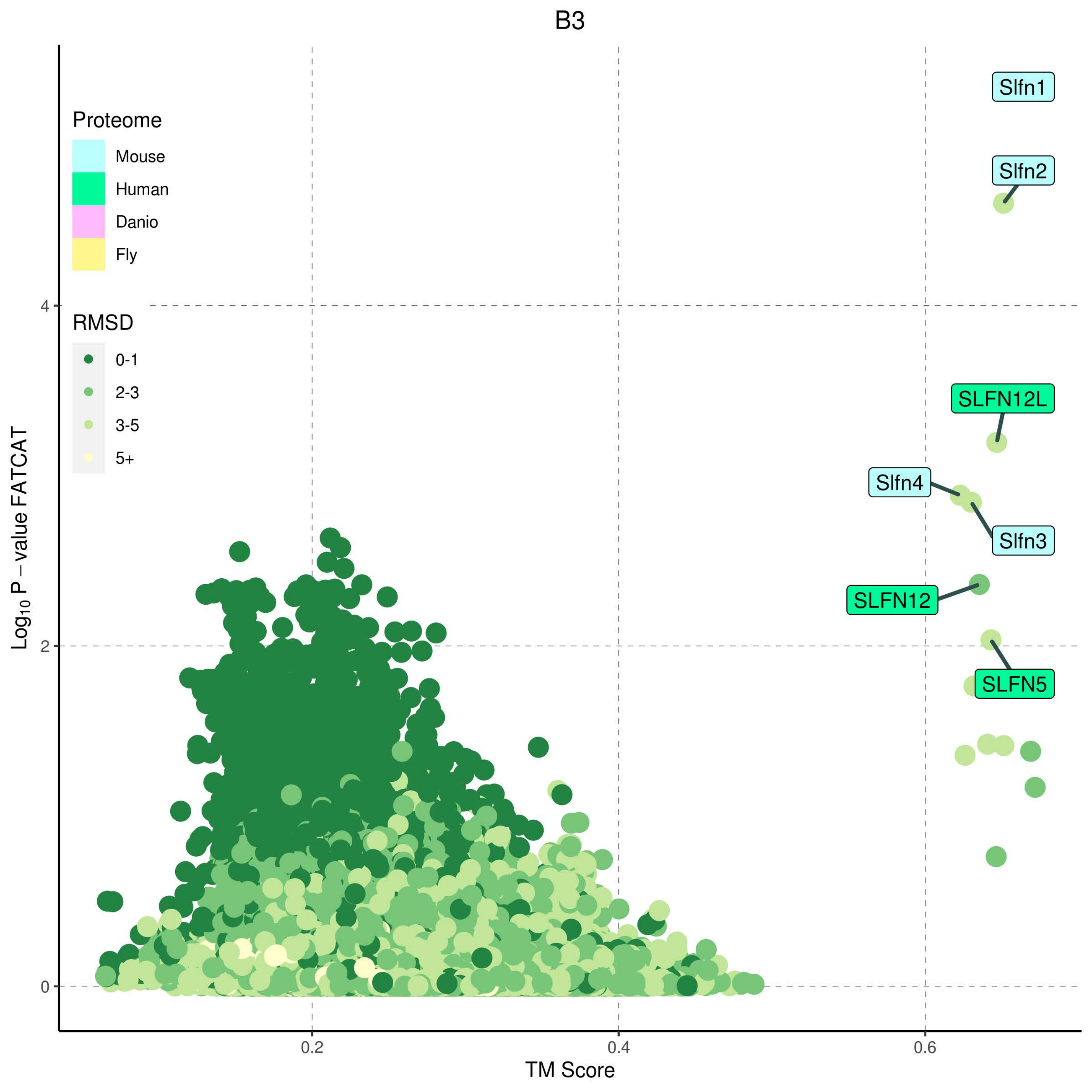
B3
Proteome
Mouse
Human
Danio
Fly
RMSD
0-1
2-3
3-5
5+
Slfn1
Slfn2
SLFN12L
Slfn4
Slfn3
SLFN12
SLFN5
$Log_{10}$ P − value FATCAT
TM Score
0
2
4
0.2
0.4
0.6

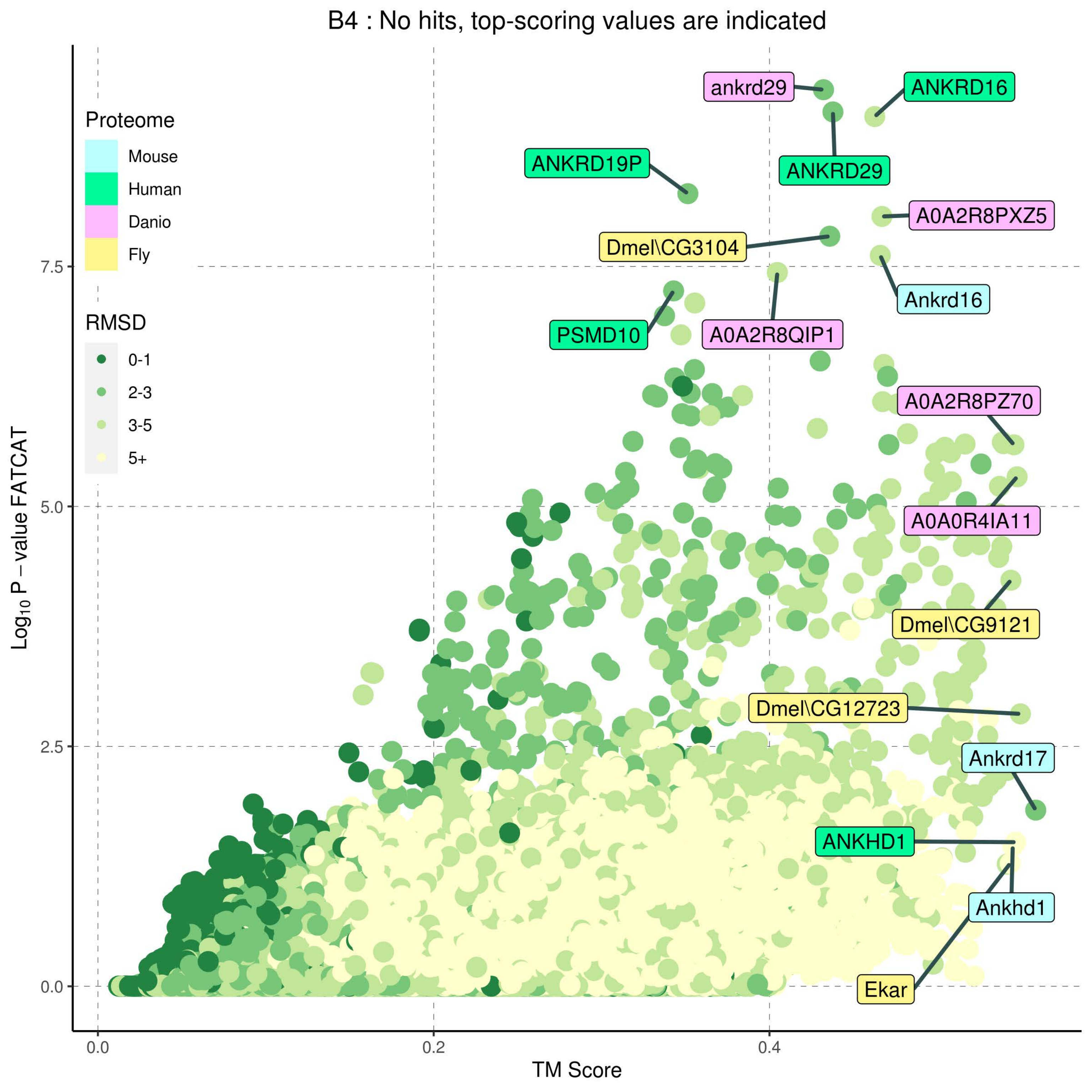
B4 : No hits, top-scoring values are indicated
Proteome
Mouse
Human
Danio
Fly
RMSD
0-1
2-3
3-5
5+
ankrd29
ANKRD16
ANKRD29
ANKRD19P
A0A2R8PXZ5
Dmel\CG3104
Ankrd16
PSMD10
A0A2R8QIP1
A0A2R8PZ70
A0A0R4IA11
Dmel\CG9121
Dmel\CG12723
Ankrd17
ANKHD1
Ankhd1
Ekar
Log10 P − value FATCAT
7.5
5.0
2.5
0.0
0.2
0.4
TM Score

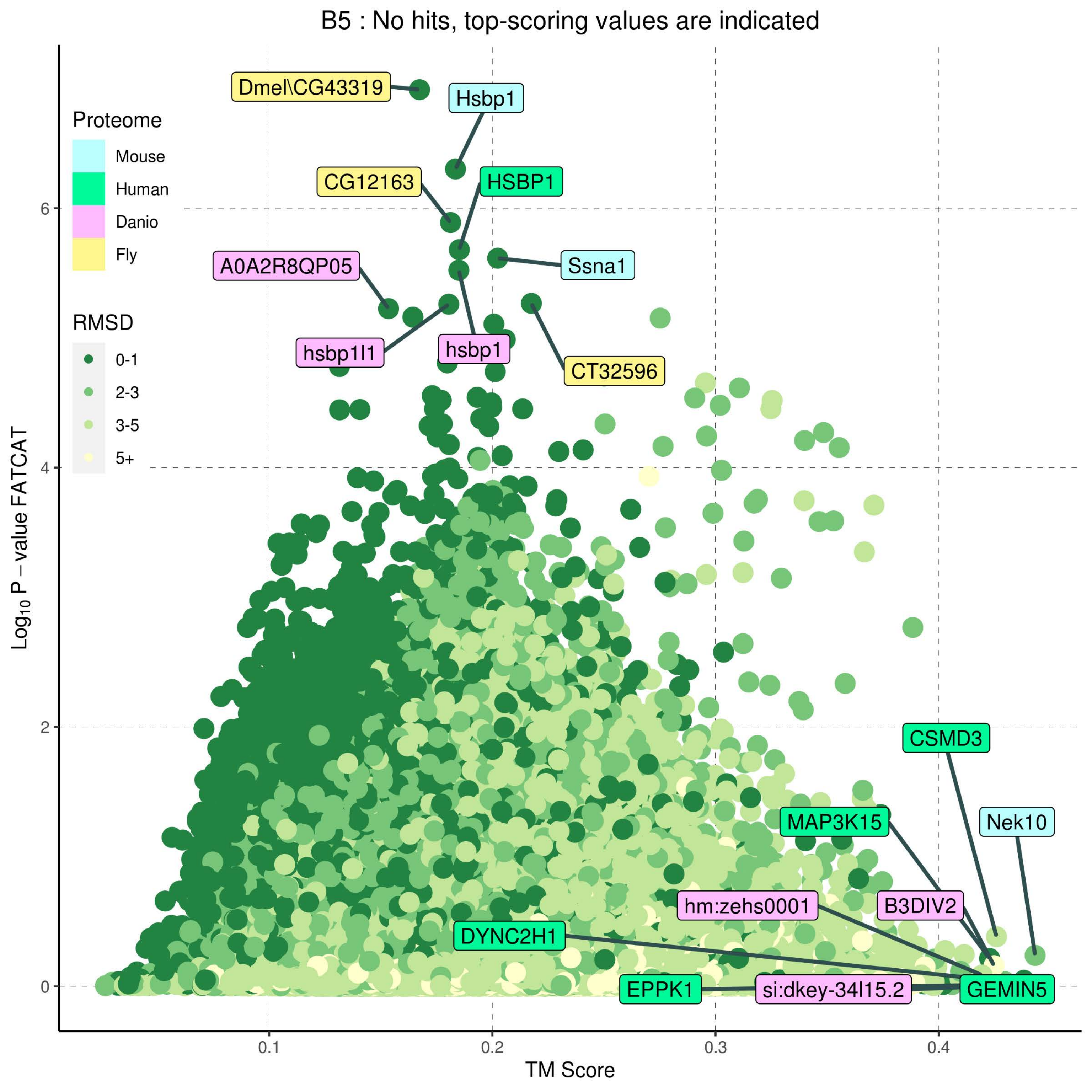
B5 : No hits, top-scoring values are indicated
Proteome
Mouse
Human
Danio
Fly
RMSD
0-1
2-3
3-5
5+
Dmel\CG43319
Hsbp1
CG12163
HSBP1
Ssna1
A0A2R8QP05
hsbp1l1
hsbp1
CT32596
CSMD3
MAP3K15
Nek10
hm:zehs0001
B3DIV2
DYNC2H1
EPPK1
si:dkey-34l15.2
GEMIN5
Log10 P – value FATCAT
TM Score
0
2
4
6
0.1
0.2
0.3
0.4

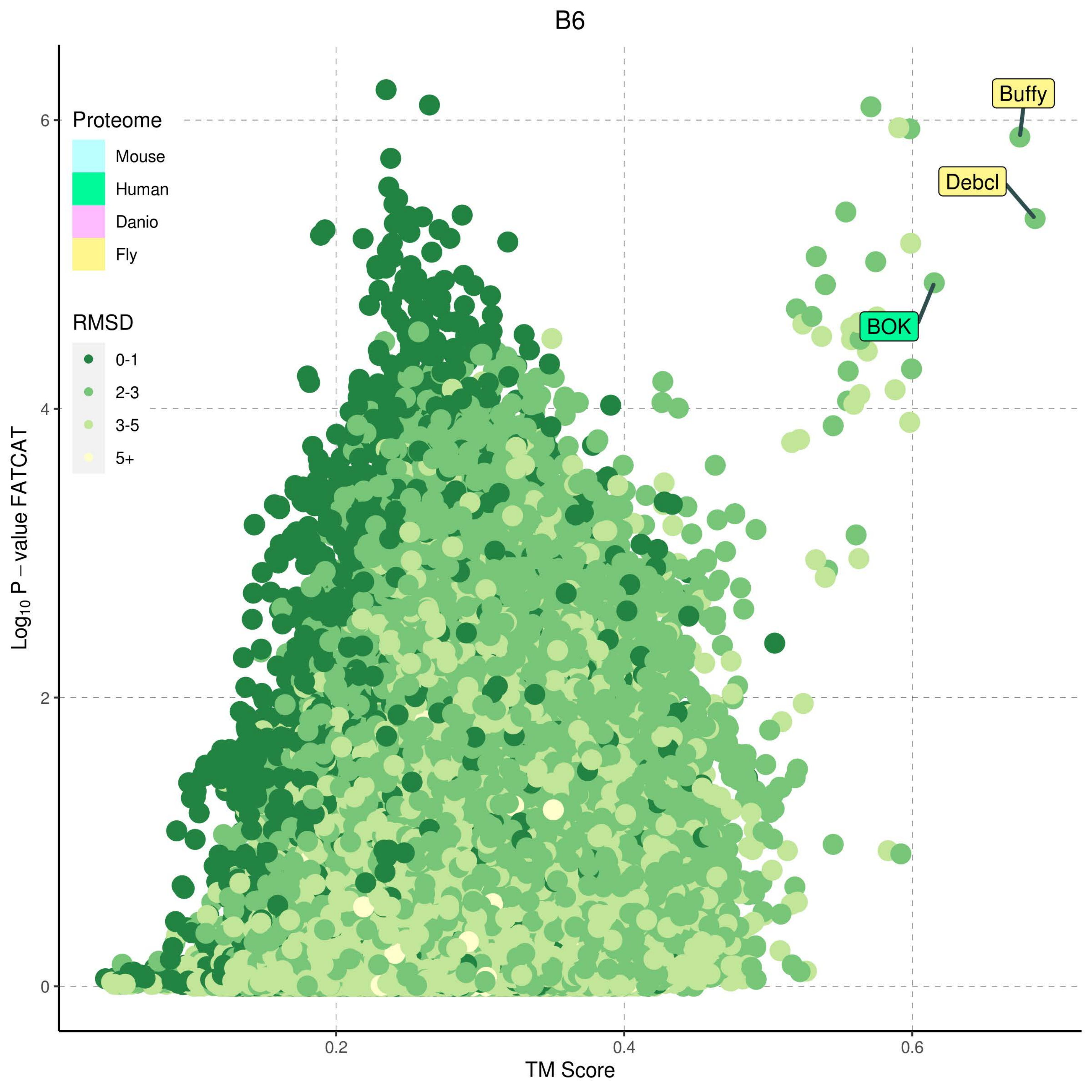
B6
Proteome
Mouse
Human
Danio
Fly
RMSD
0-1
2-3
3-5
5+
Buffy
Debcl
BOK
$\mathrm{Log}_{10}$ P − value FATCAT
TM Score
0
2
4
6
0.2
0.4
0.6

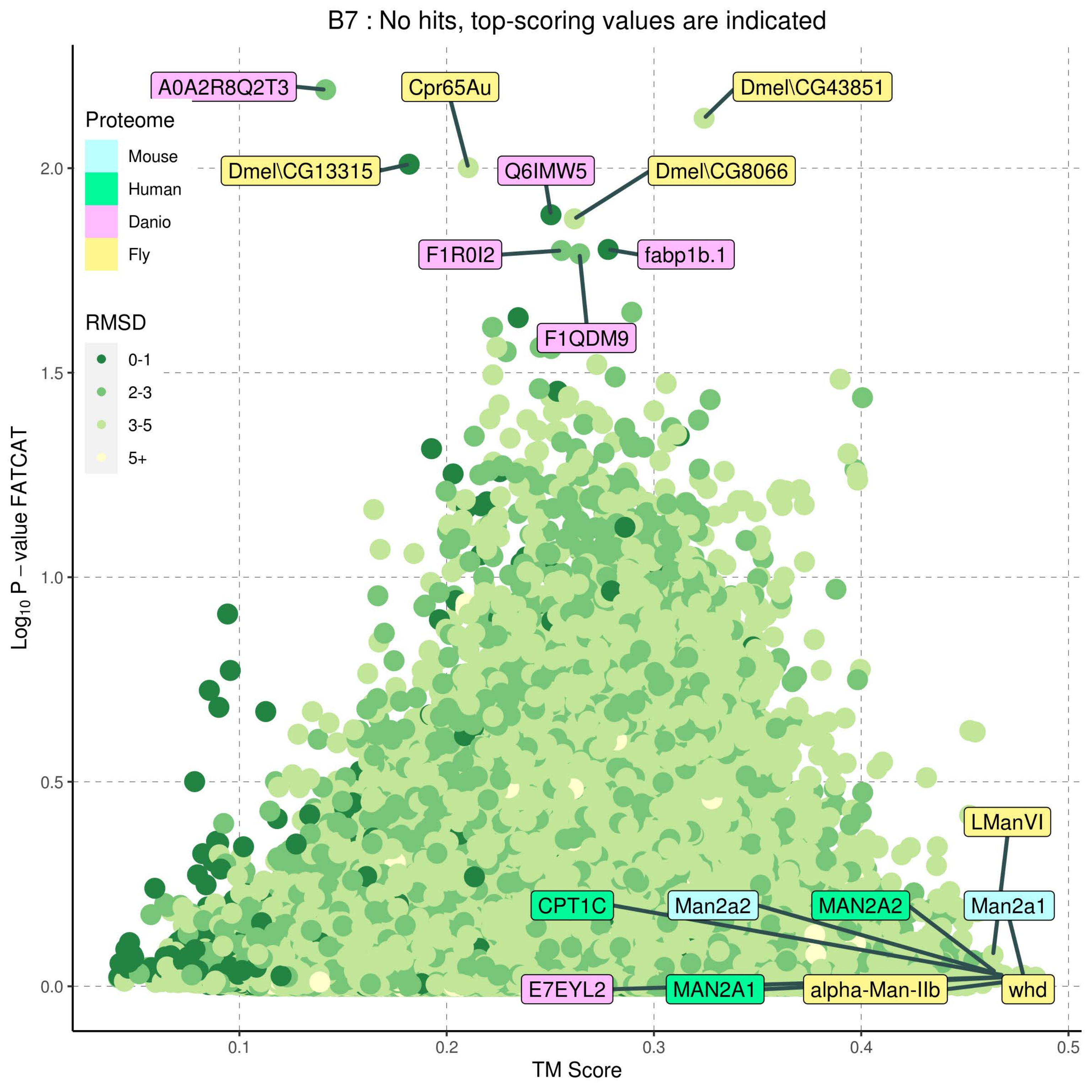

B7 : No hits, top-scoring values are indicated
Proteome
Mouse
Human
Danio
Fly
RMSD
0-1
2-3
3-5
5+
A0A2R8Q2T3
Cpr65Au
Dmel\CG43851
Dmel\CG13315
Q6IMW5
Dmel\CG8066
F1R0I2
fabp1b.1
F1QDM9
LManVI
CPT1C
Man2a2
MAN2A2
Man2a1
E7EYL2
MAN2A1
alpha-Man-IIb
whd
Log10 P − value FATCAT
TM Score
0.0
0.5
1.0
1.5
2.0
0.1
0.2
0.3
0.4
0.5

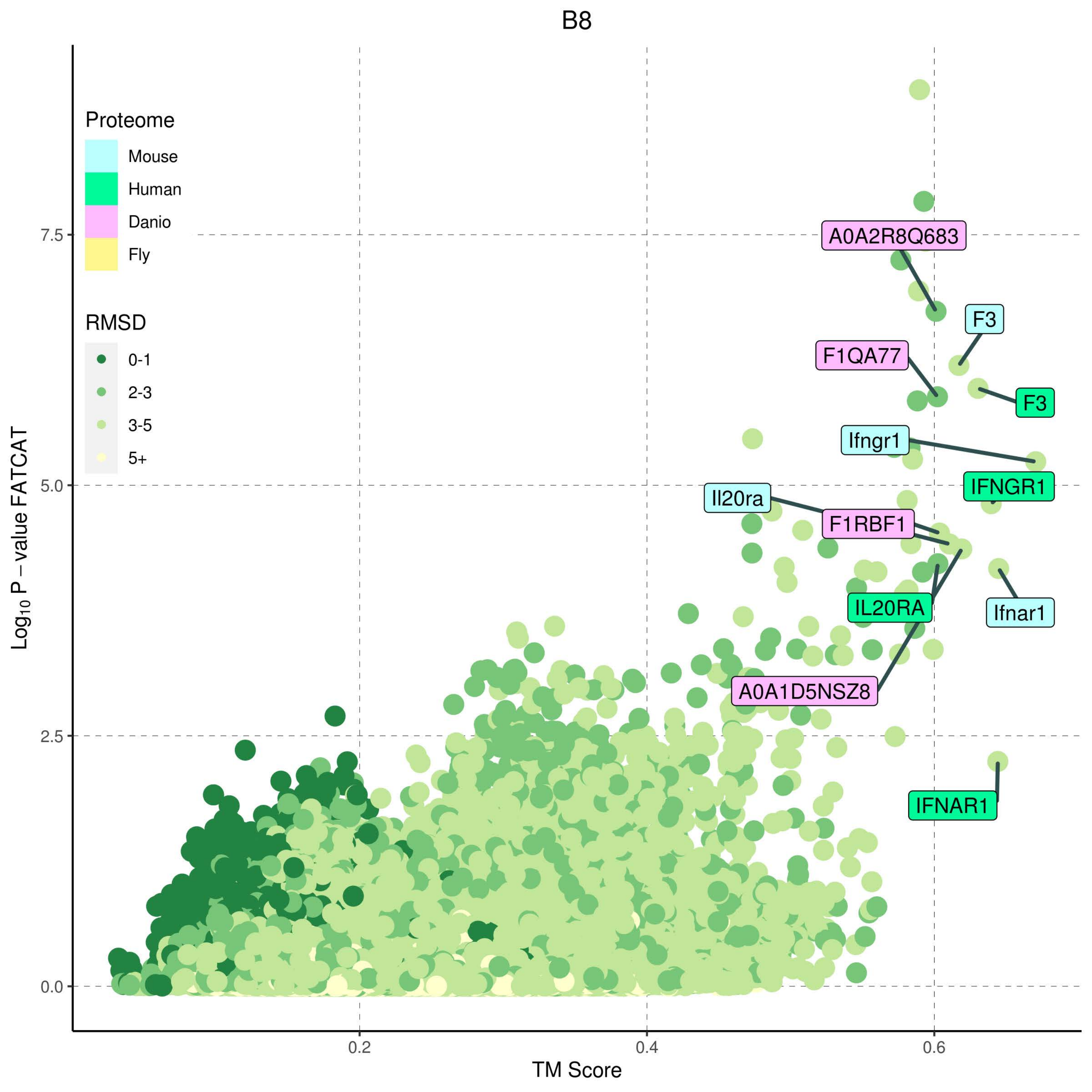
B8
Proteome
Mouse
Human
Danio
Fly
RMSD
0-1
2-3
3-5
5+
A0A2R8Q683
F3
F1QA77
F3
Ifngr1
IFNGR1
Il20ra
F1RBF1
IL20RA
Ifnar1
A0A1D5NSZ8
IFNAR1
7.5
5.0
2.5
0.0
0.2
0.4
0.6
Log10 P − value FATCAT
TM Score

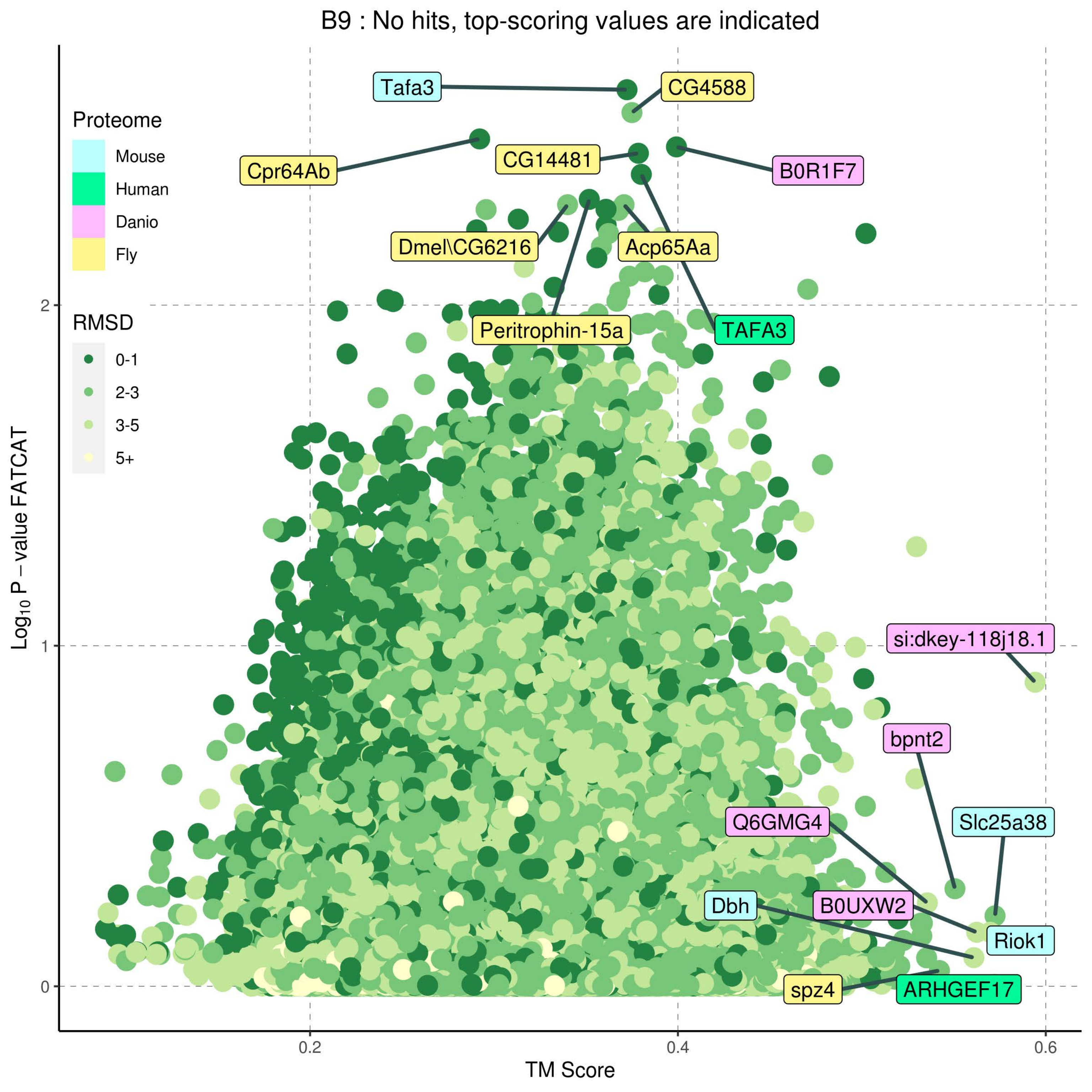
B9 : No hits, top-scoring values are indicated
Proteome
Mouse
Human
Danio
Fly
RMSD
0-1
2-3
3-5
5+
Tafa3
CG4588
Cpr64Ab
CG14481
B0R1F7
Dmel\CG6216
Acp65Aa
Peritrophin-15a
TAFA3
si:dkey-118j18.1
bpnt2
Q6GMG4
Slc25a38
Dbh
B0UXW2
Riok1
spz4
ARHGEF17
Log10 P − value FATCAT
TM Score
0
1
2
0.2
0.4
0.6

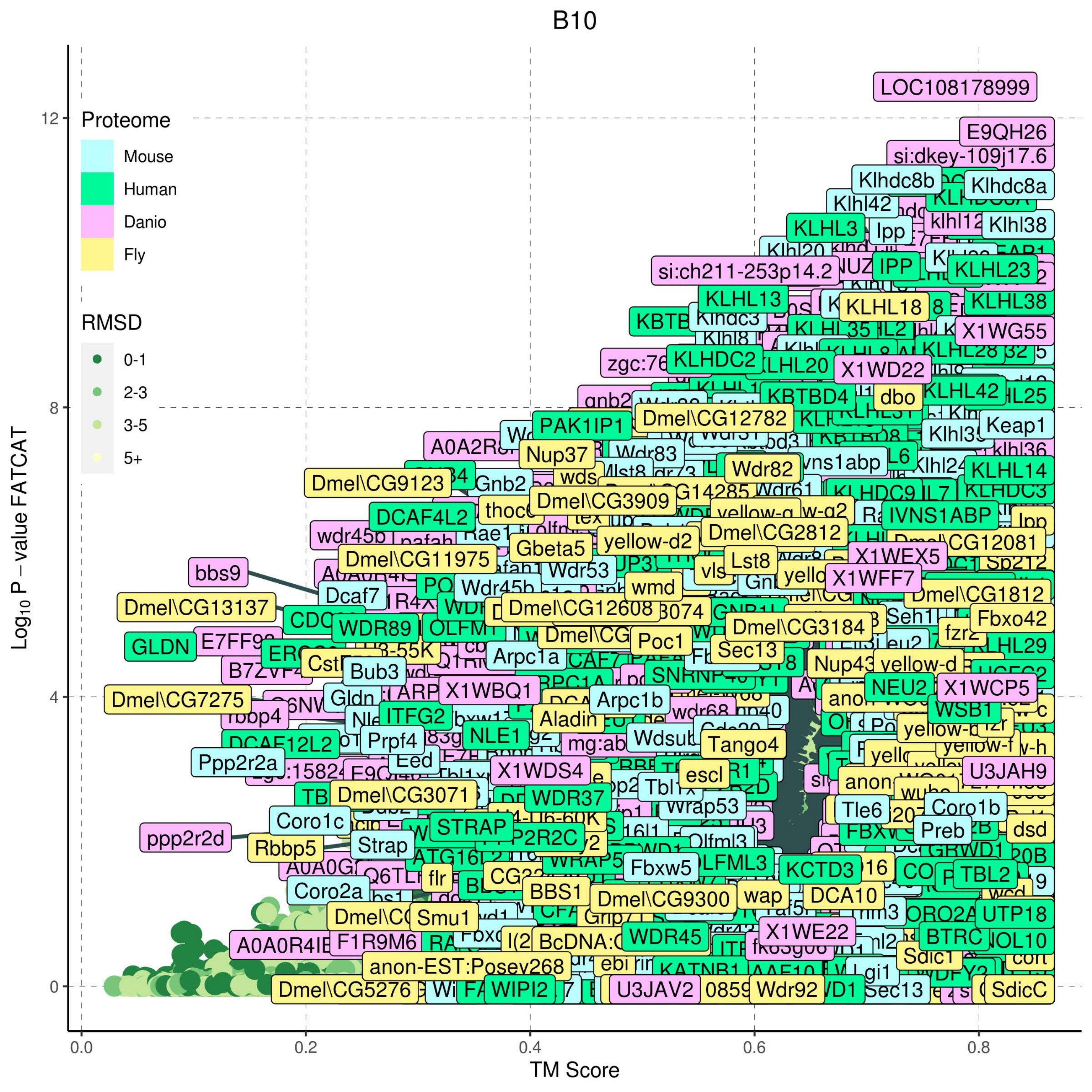

B10
Proteome
Mouse
Human
Danio
Fly
RMSD
0-1
2-3
3-5
5+
Log10 P − value FATCAT
TM Score
0.0
0.2
0.4
0.6
0.8
0
4
8
12
LOC108178999
E9QH26
si:dkey-109j17.6
Klhdc8b
Klhdc8a
Klhl42
KLHL3
Ipp
Klhl38
si:ch211-253p14.2
IPP
KLHL23
KLHL13
KLHL18
KLHL38
X1WG55
KLHDC2
KLHL20
X1WD22
KBTBD4
dbo
KLHL42
Keap1
PAK1IP1
Dmel\CG12782
Nup37
Wdr83
Wdr82
KLHL14
Dmel\CG9123
Gnb2
Dmel\CG3909
DCAF4L2
IVNS1ABP
Dmel\CG2812
yellow-d2
Gbeta5
Dmel\CG11975
X1WEX5
bbs9
Wdr53
Lst8
X1WFF7
Dcaf7
wmd
Dmel\CG1812
Dmel\CG13137
Dmel\CG12608
Fbxo42
WDR89
Dmel\CG3184
GLDN
Poc1
Arpc1a
Sec13
Bub3
Nup43
Dmel\CG7275
X1WBQ1
Arpc1b
NEU2
X1WCP5
ITFG2
Aladin
WSB1
NLE1
Prpf4
Tango4
Ppp2r2a
X1WDS4
escl
U3JAH9
Dmel\CG3071
WDR37
Tle6
Coro1b
Coro1c
STRAP
Preb
dsd
ppp2r2d
Rbbp5
Strap
Fbxw5
KCTD3
TBL2
BBS1
Dmel\CG9300
wap
DCA10
UTP18
Smu1
X1WE22
A0A0R4IE
F1R9M6
WDR45
BTRC
anon-EST:Posev268
Sdic1
Lgi1
Dmel\CG5276
WIPI2
U3JAV2
Wdr92
Sec13
SdicC

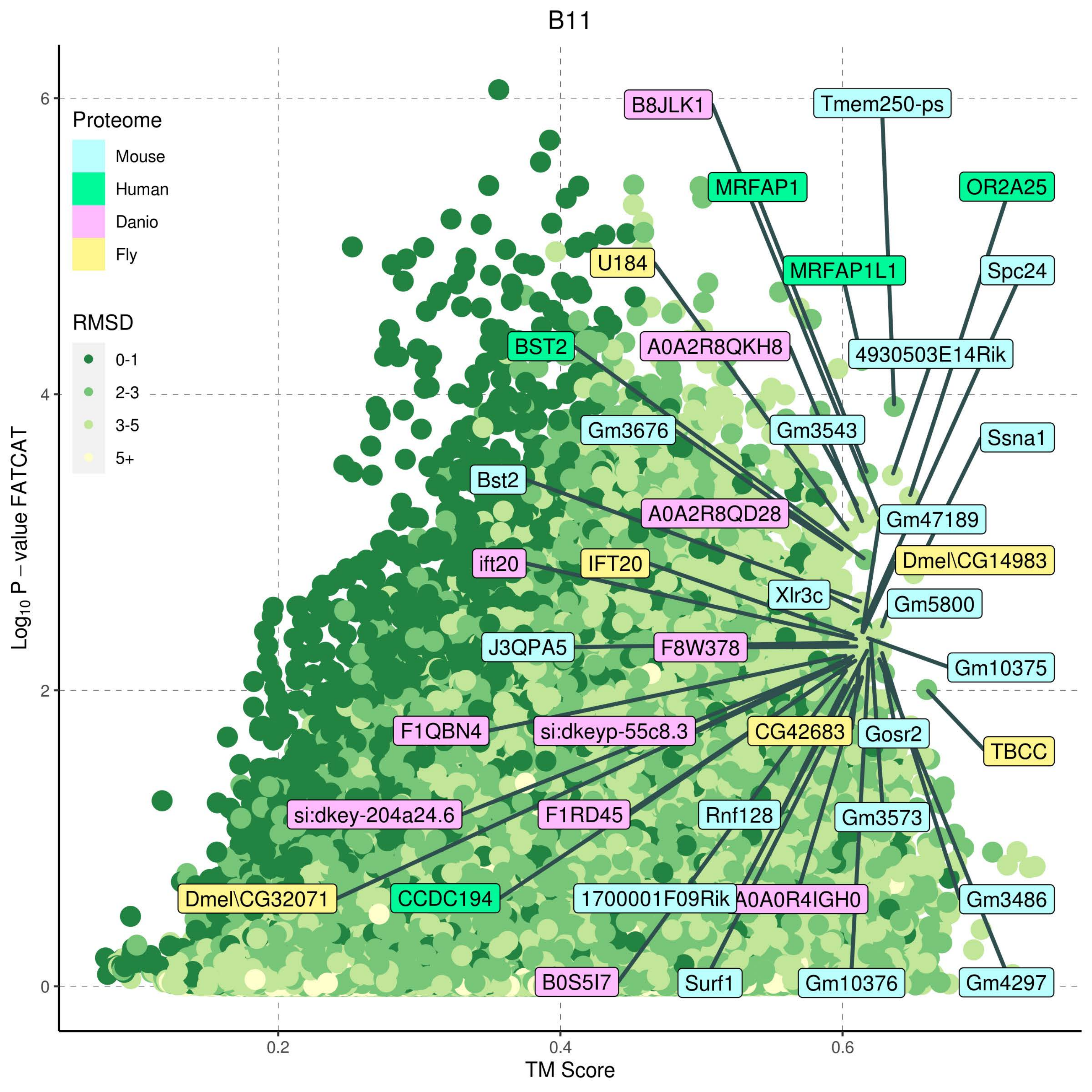

B11
Proteome
Mouse
Human
Danio
Fly
RMSD
0-1
2-3
3-5
5+
Log10 P – value FATCAT
TM Score
0
2
4
6
0.2
0.4
0.6
B8JLK1
Tmem250-ps
MRFAP1
OR2A25
MRFAP1L1
Spc24
U184
BST2
A0A2R8QKH8
4930503E14Rik
Gm3676
Gm3543
Ssna1
Bst2
A0A2R8QD28
Gm47189
ift20
IFT20
Dmel\CG14983
Xlr3c
Gm5800
J3QPA5
F8W378
Gm10375
F1QBN4
si:dkeyp-55c8.3
CG42683
Gosr2
TBCC
si:dkey-204a24.6
F1RD45
Rnf128
Gm3573
Dmel\CG32071
CCDC194
1700001F09Rik
A0A0R4IGH0
Gm3486
B0S5I7
Surf1
Gm10376
Gm4297

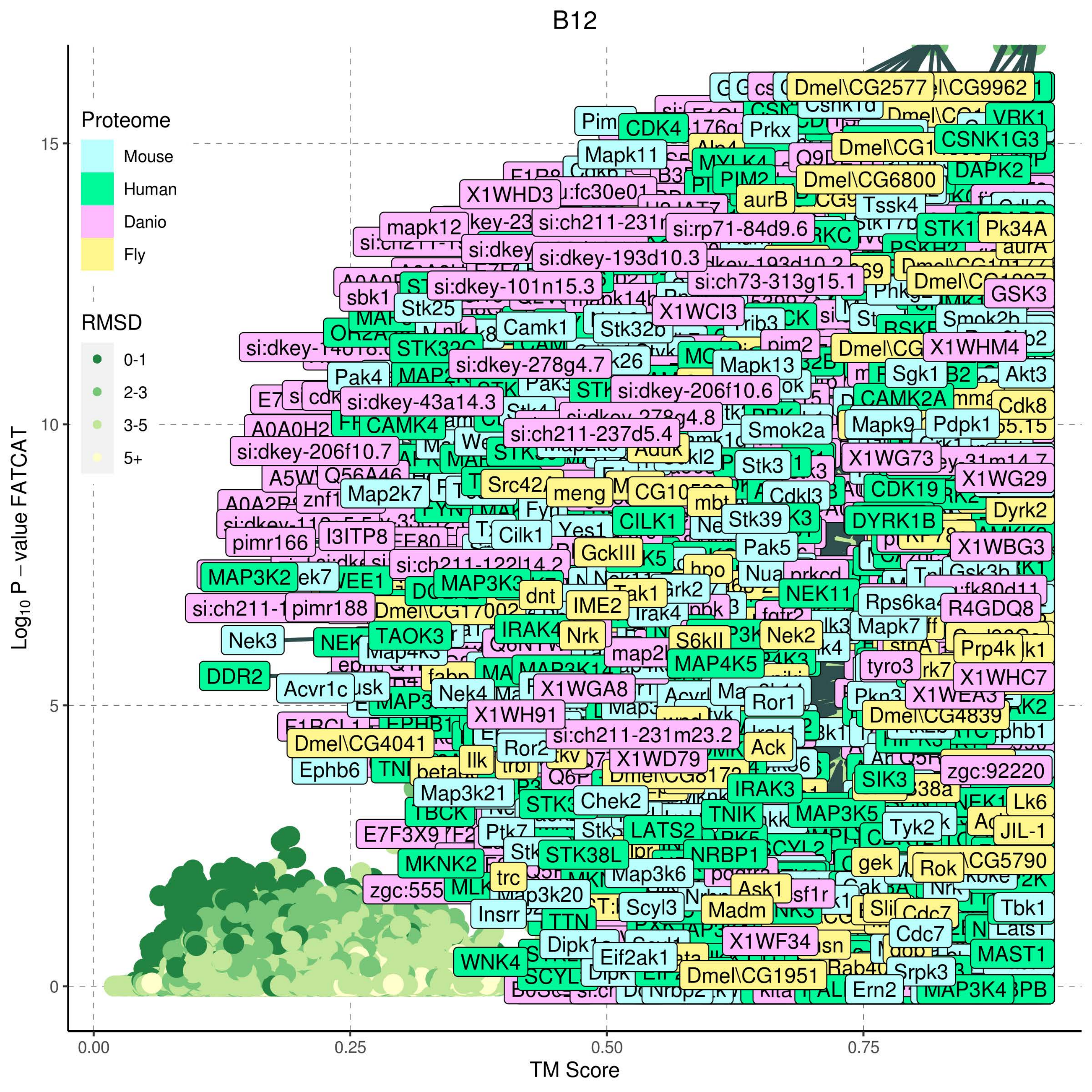

B12
Proteome
Mouse
Human
Danio
Fly
RMSD
0-1
2-3
3-5
5+
Log10 P − value FATCAT
TM Score
0.00
0.25
0.50
0.75
0
5
10
15
Dmel\CG2577
Pim
CDK4
Prkx
VRK1
CSNK1G3
Mapk11
DAPK2
PIM2
Dmel\CG6800
X1WHD3
aurB
Tssk4
mapk12
si:rp71-84d9.6
Pk34A
si:dkey-193d10.3
si:ch73-313g15.1
si:dkey-101n15.3
sbk1
Stk25
GSK3
X1WCI3
Camk1
Stk32b
si:dkey-278g4.7
X1WHM4
Pak4
Mapk13
Sgk1
Akt3
si:dkey-206f10.6
si:dkey-43a14.3
CAMK2A
Cdk8
CAMK4
Mapk9
Pdpk1
si:ch211-237d5.4
Smok2a
si:dkey-206f10.7
X1WG73
Stk3
X1WG29
Src42A
meng
CDK19
Map2k7
Dyrk2
DYRK1B
Stk39
pimr166
I3ITP8
Cilk1
Yes1
CILK1
X1WBG3
GckIII
Pak5
MAP3K2
hpo
dnt
IME2
NEK11
si:ch211-1
pimr188
Rps6ka
R4GDQ8
Nek3
TAOK3
IRAK4
Nrk
S6kII
Nek2
Mapk7
Prp4k
MAP4K5
tyro3
X1WHC7
DDR2
Acvr1c
Nek4
X1WGA8
Ror1
X1WH91
Dmel\CG4839
si:ch211-231m23.2
Dmel\CG4041
Ack
Ephb6
Ilk
Ror2
X1WD79
zgc:92220
SIK3
Map3k21
Chek2
IRAK3
Lk6
TNIK
MAP3K5
Tyk2
E7F3X9
LATS2
JIL-1
MKNK2
STK38L
NRBP1
gek
Rok
trc
Map3k6
zgc:555
Map3k20
Ask1
Insrr
Scyl3
Madm
Tbk1
Cdc7
Dipk1
X1WF34
WNK4
Eif2ak1
MAST1
Dmel\CG1951
Srpk3
Ern2
MAP3K4
Nrbp2

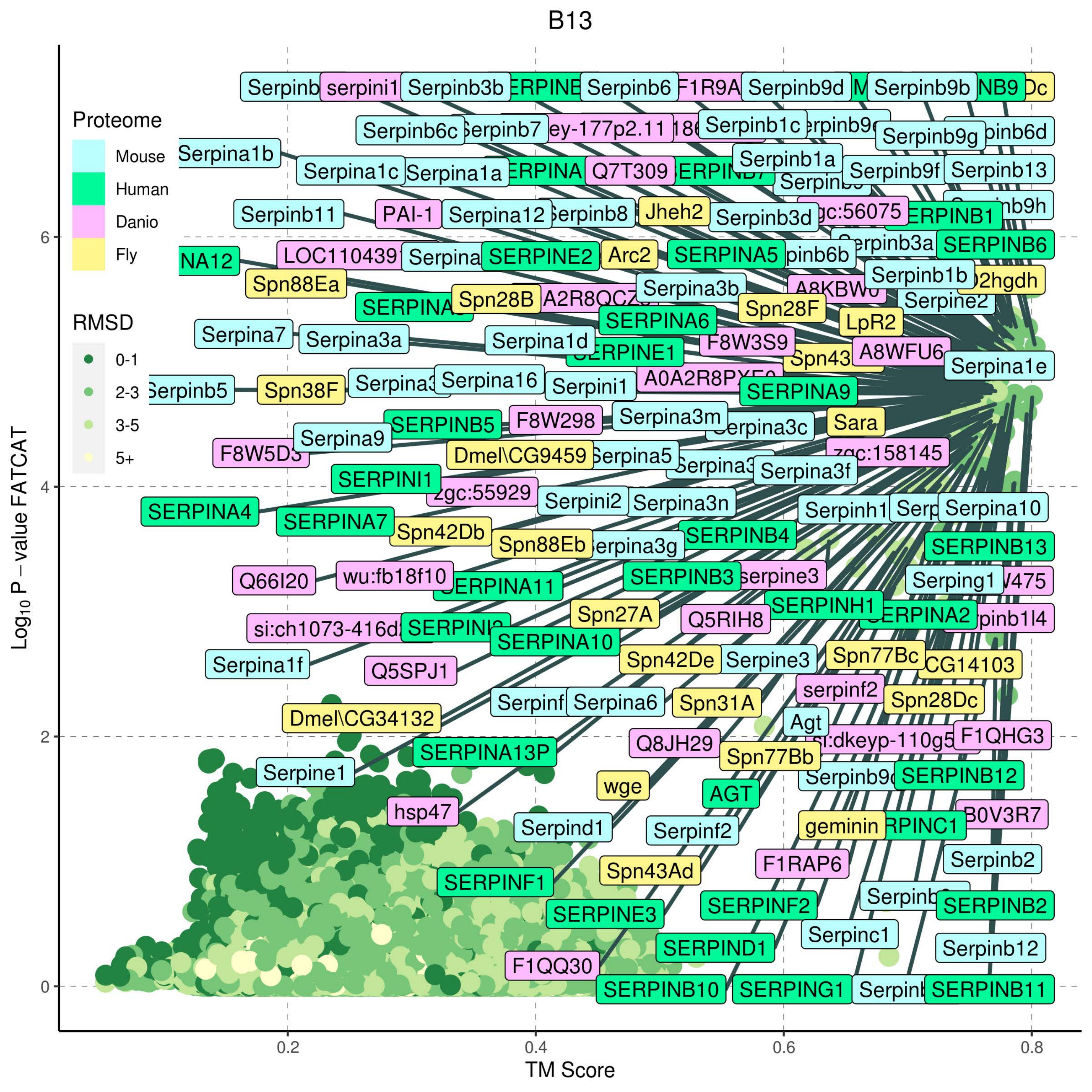
B13
Proteome
Mouse
Human
Danio
Fly
RMSD
0-1
2-3
3-5
5+
Log10 P − value FATCAT
TM Score
0
2
4
6
0.2
0.4
0.6
0.8
serpini1
Serpinb3b
Serpinb6
Serpinb9d
Serpinb9b
Serpinb6c
Serpinb1c
Serpinb9g
Serpina1b
Serpinb1a
Serpina1c
Serpina1a
Q7T309
Serpinb9f
Serpinb13
Serpinb11
PAI-1
Serpina12
Jheh2
Serpinb3d
SERPINE2
Arc2
SERPINA5
SERPINB6
Spn88Ea
Serpinb1b
Serpina3b
Serpine2
Spn28B
SERPINA6
Spn28F
LpR2
Serpina7
Serpina3a
Serpina1d
F8W3S9
A8WFU6
Serpina1e
Serpinb5
Spn38F
Serpina16
Serpini1
SERPINA9
SERPINB5
F8W298
Serpina3m
Sara
Serpina9
Dmel\CG9459
Serpina3f
SERPINI1
Serpini2
Serpina3n
Serpinh1
Serpina10
SERPINA4
SERPINA7
Spn42Db
Spn88Eb
SERPINB4
SERPINB13
Q66I20
wu:fb18f10
SERPINB3
serpine3
Serping1
SERPINH1
Spn27A
Q5RIH8
SERPINA10
Serpina1f
Q5SPJ1
Spn42De
Serpine3
Spn77Bc
CG14103
serpinf2
Serpina6
Spn31A
Spn28Dc
Dmel\CG34132
Agt
F1QHG3
Q8JH29
SERPINA13P
Spn77Bb
Serpine1
wge
AGT
SERPINB12
hsp47
Serpind1
Serpinf2
geminin
B0V3R7
Serpinb2
Spn43Ad
F1RAP6
SERPINF1
SERPINF2
SERPINB2
SERPINE3
Serpinc1
SERPIND1
Serpinb12
F1QQ30
SERPINB10
SERPING1
SERPINB11

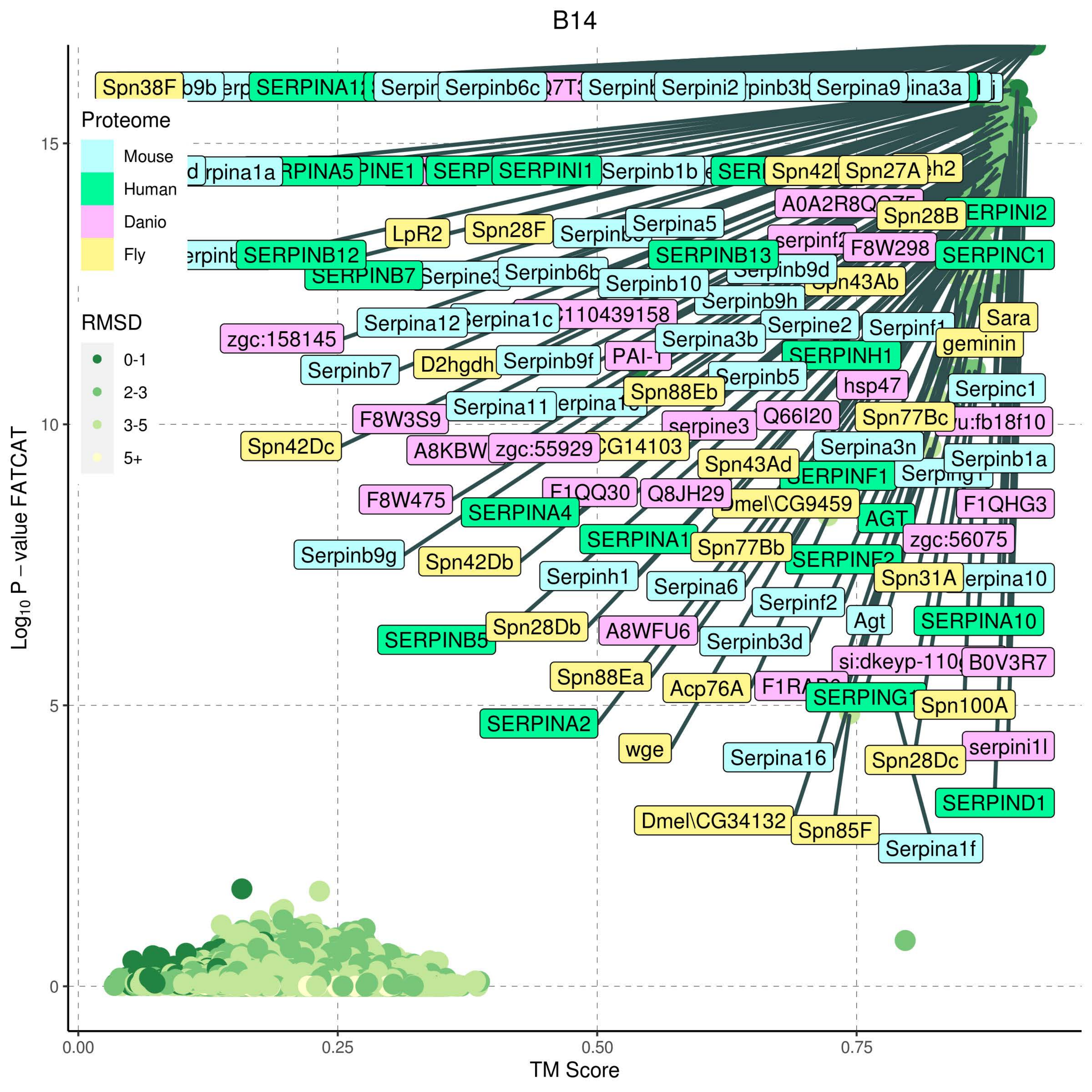

B14
Proteome
Mouse
Human
Danio
Fly
RMSD
0-1
2-3
3-5
5+
Spn38F
Serpinb6c
Serpini2
Serpina9
SERPINI1
Serpinb1b
Spn27A
LpR2
Spn28F
Serpina5
SERPINB12
SERPINB13
Spn28B
F8W298
SERPINC1
Serpinb10
Serpinb9h
Serpina12
zgc:158145
Serpine2
Sara
geminin
Serpina3b
Serpinb7
D2hgdh
Serpinb9f
SERPINH1
Serpinb5
hsp47
Serpinc1
Spn88Eb
Serpina11
F8W3S9
Q66I20
Spn77Bc
serpine3
Spn42Dc
zgc:55929
Serpina3n
Serpinb1a
Spn43Ad
SERPINF1
F8W475
Q8JH29
F1QHG3
SERPINA4
AGT
zgc:56075
Serpinb9g
Spn77Bb
Spn42Db
Serpinh1
Spn31A
Serpina6
Serpinf2
Agt
SERPINA10
SERPINB5
Spn28Db
A8WFU6
Serpinb3d
B0V3R7
Spn88Ea
Acp76A
Spn100A
SERPINA2
wge
serpini1l
Serpina16
Spn28Dc
SERPIND1
Dmel\CG34132
Spn85F
Serpina1f
$Log_{10}$ P – value FATCAT
0
5
10
15
0.00
0.25
0.50
0.75
TM Score

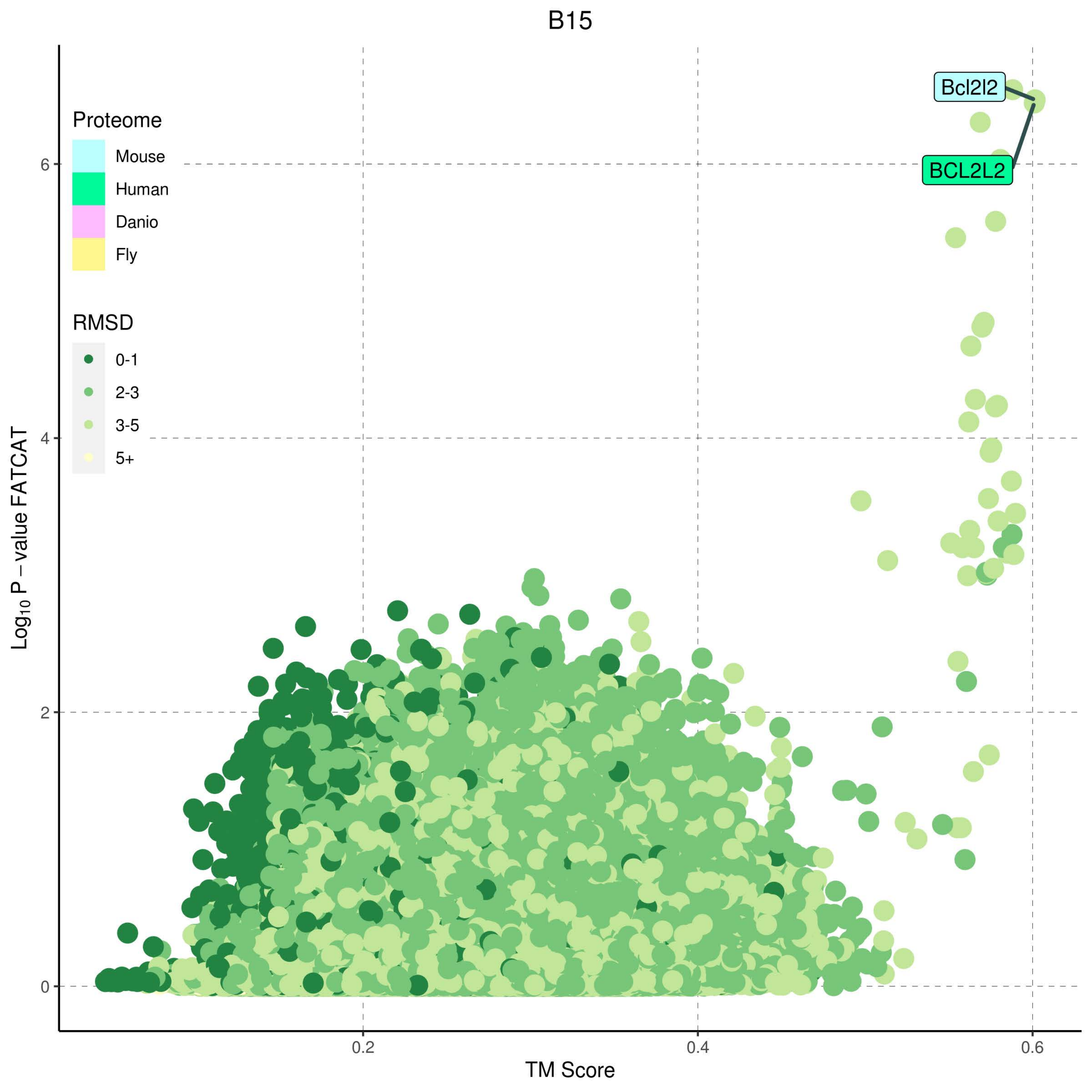
B15
Proteome
Mouse
Human
Danio
Fly
RMSD
0-1
2-3
3-5
5+
Bcl2l2
BCL2L2
Log10 P − value FATCAT
0
2
4
6
0.2
0.4
0.6
TM Score

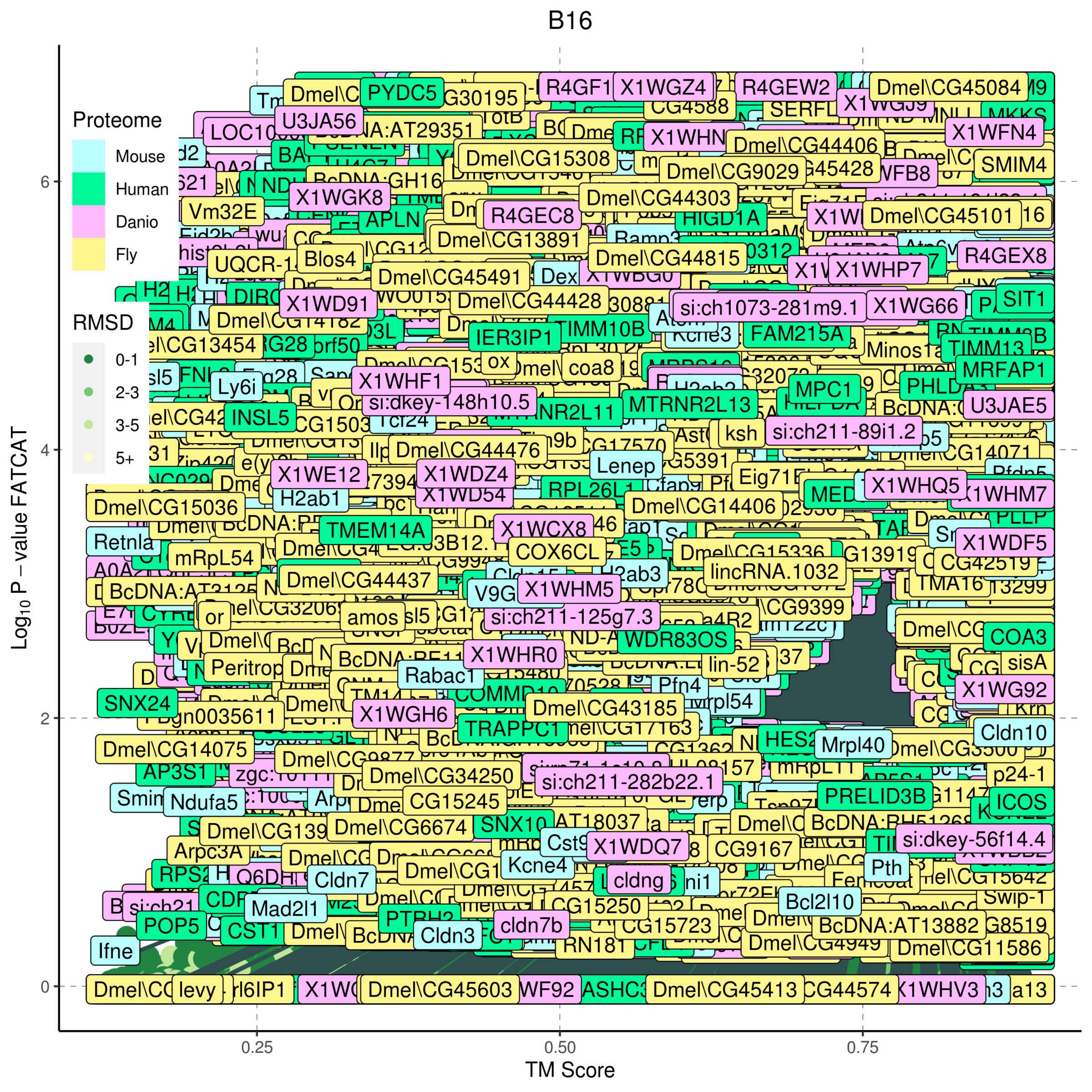
B16
Proteome
Mouse
Human
Danio
Fly
RMSD
0-1
2-3
3-5
5+
Log10 P − value FATCAT
TM Score
0
2
4
6
0.25
0.50
0.75
Dmel\CG45084
PYDC5
X1WGZ4
R4GEW2
U3JA56
X1WFN4
Dmel\CG44406
Dmel\CG15308
SMIM4
Dmel\CG9029
X1WGK8
Dmel\CG44303
Vm32E
APLN
R4GEC8
HIGD1A
Dmel\CG45101
Dmel\CG13891
Blos4
Dmel\CG44815
R4GEX8
Dmel\CG45491
X1WHP7
X1WD91
Dmel\CG44428
si:ch1073-281m9.1
X1WG66
SIT1
IER3IP1
TIMM10B
FAM215A
Minos1a
TIMM13
MRFAP1
Ly6i
X1WHF1
si:dkey-148h10.5
MTRNR2L13
MPC1
U3JAE5
INSL5
ksh
si:ch211-89i1.2
Dmel\CG44476
Lenep
X1WE12
X1WDZ4
H2ab1
Dmel\CG15036
Dmel\CG14406
X1WHQ5
Retnla
TMEM14A
X1WCX8
X1WDF5
mRpL54
COX6CL
Dmel\CG15336
CG42519
Dmel\CG44437
lincRNA.1032
X1WHM5
TMA16
amos
si:ch211-125g7.3
WDR83OS
COA3
X1WHR0
Rabac1
lin-52
sisA
Pfn4
X1WG92
SNX24
Dmel\CG43185
X1WGH6
TRAPPC1
Cldn10
Dmel\CG14075
HES2
Mrpl40
Dmel\CG34250
p24-1
si:ch211-282b22.1
Ndufa5
CG15245
PRELID3B
ICOS
Dmel\CG6674
SNX10
si:dkey-56f14.4
X1WDQ7
CG9167
Arpc3A
Kcne4
Cldn7
cldng
Pth
Bcl2l10
Mad2l1
CG15250
POP5
CST1
Cldn3
cldn7b
CG15723
BcDNA:AT13882
Dmel\CG4949
Ifne
Dmel\CG11586
levy
Dmel\CG45603
Dmel\CG45413
a13

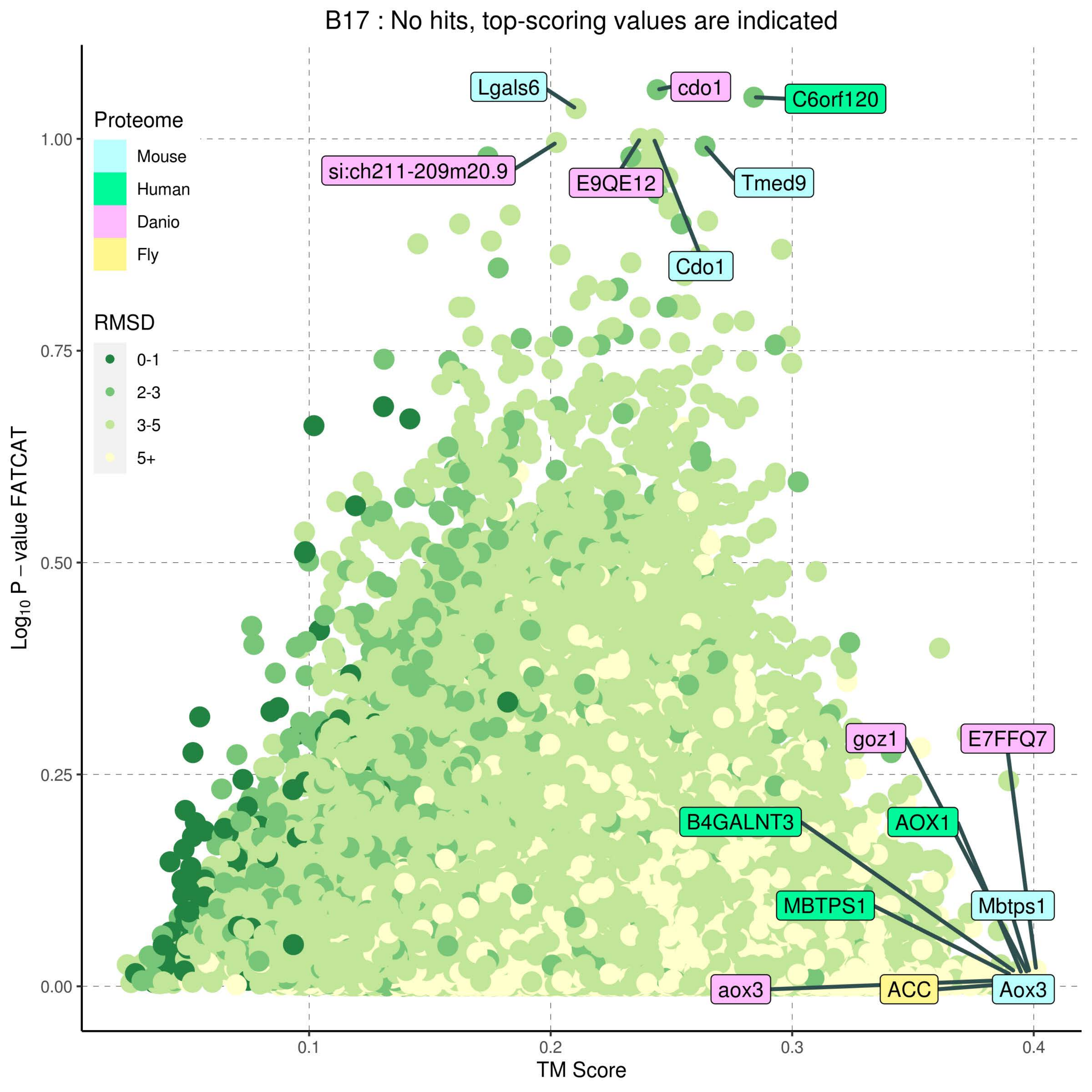
B17 : No hits, top-scoring values are indicated
Proteome
Mouse
Human
Danio
Fly
RMSD
0-1
2-3
3-5
5+
Lgals6
cdo1
C6orf120
si:ch211-209m20.9
E9QE12
Tmed9
Cdo1
goz1
E7FFQ7
B4GALNT3
AOX1
MBTPS1
Mbtps1
aox3
ACC
Aox3
1.00
0.75
0.50
0.25
0.00
0.1
0.2
0.3
0.4
Log10 P − value FATCAT
TM Score

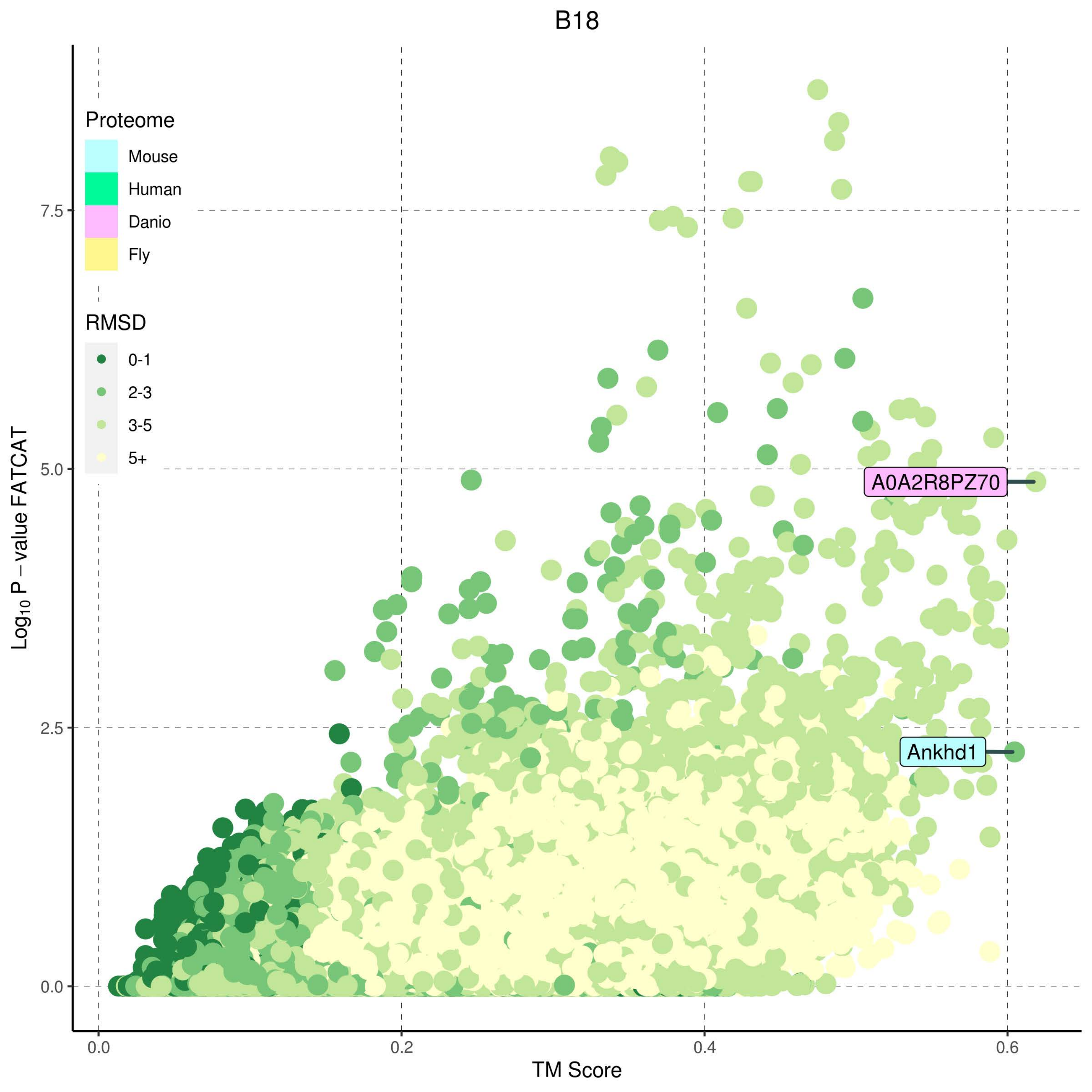
B18
Proteome
Mouse
Human
Danio
Fly
RMSD
0-1
2-3
3-5
5+
A0A2R8PZ70
Ankhd1
7.5
5.0
2.5
0.0
Log10 P − value FATCAT
0.0
0.2
0.4
0.6
TM Score

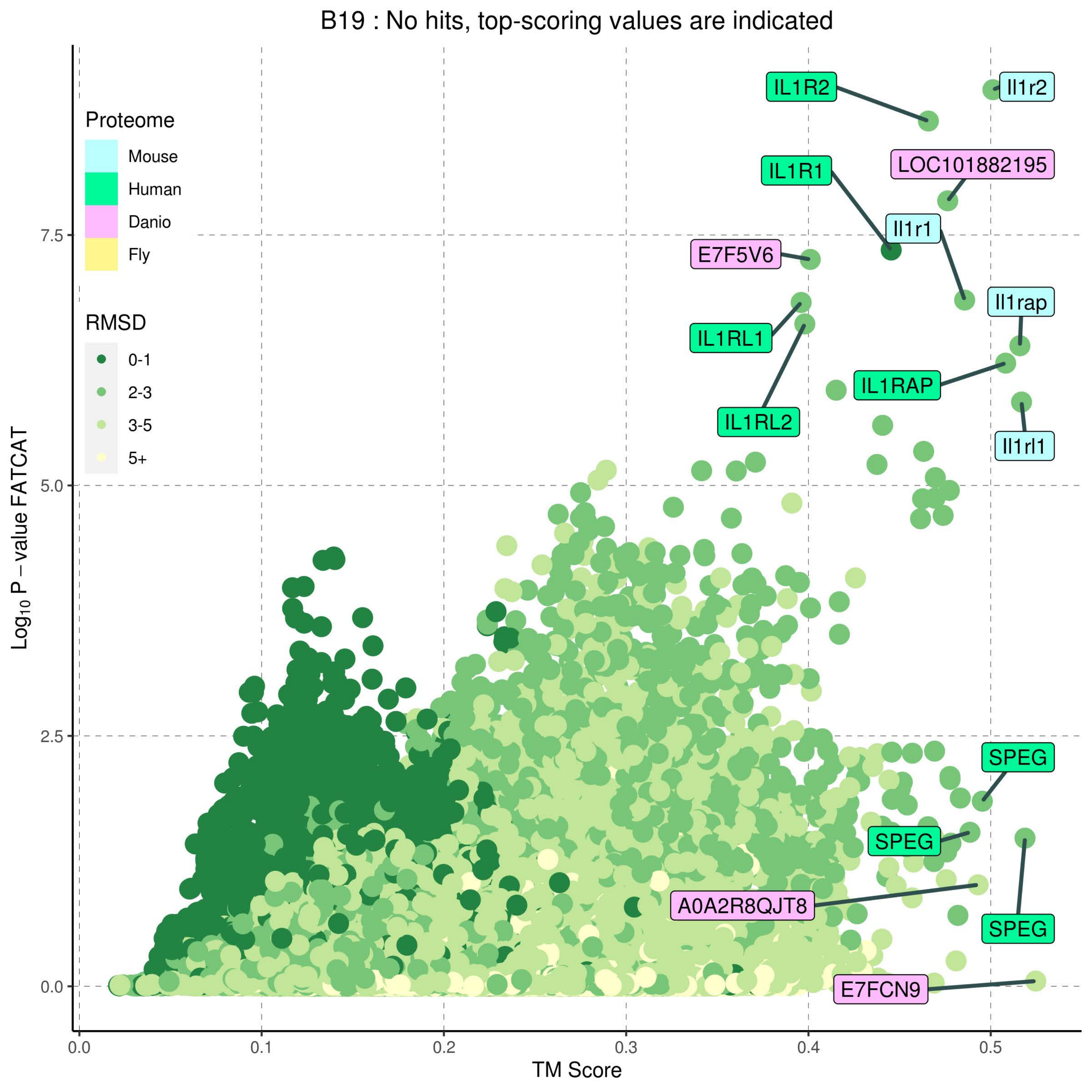

B19 : No hits, top-scoring values are indicated
Proteome
Mouse
Human
Danio
Fly
RMSD
0-1
2-3
3-5
5+
IL1R2
Il1r2
LOC101882195
IL1R1
Il1r1
E7F5V6
IL1RL1
IL1RL2
Il1rap
IL1RAP
Il1rl1
SPEG
SPEG
A0A2R8QJT8
SPEG
E7FCN9
7.5
5.0
2.5
0.0
0.0
0.1
0.2
0.3
0.4
0.5
TM Score
Log10 P − value FATCAT

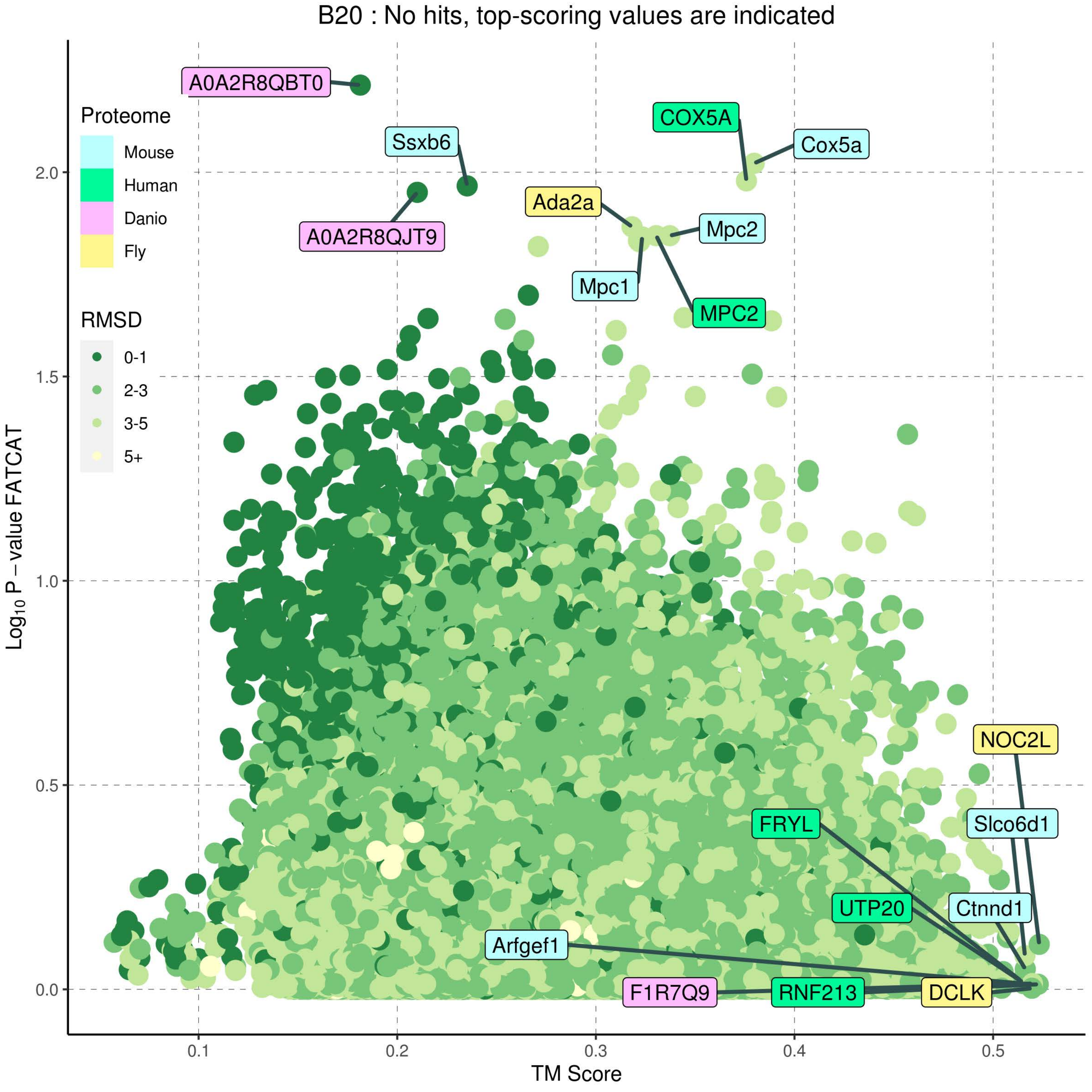

B20 : No hits, top-scoring values are indicated
Proteome
Mouse
Human
Danio
Fly
RMSD
0-1
2-3
3-5
5+
A0A2R8QBT0
Ssxb6
A0A2R8QJT9
COX5A
Cox5a
Ada2a
Mpc2
Mpc1
MPC2
NOC2L
Slco6d1
FRYL
UTP20
Ctnnd1
Arfgef1
F1R7Q9
RNF213
DCLK
2.0
1.5
1.0
0.5
0.0
0.1
0.2
0.3
0.4
0.5
Log10 P − value FATCAT
TM Score

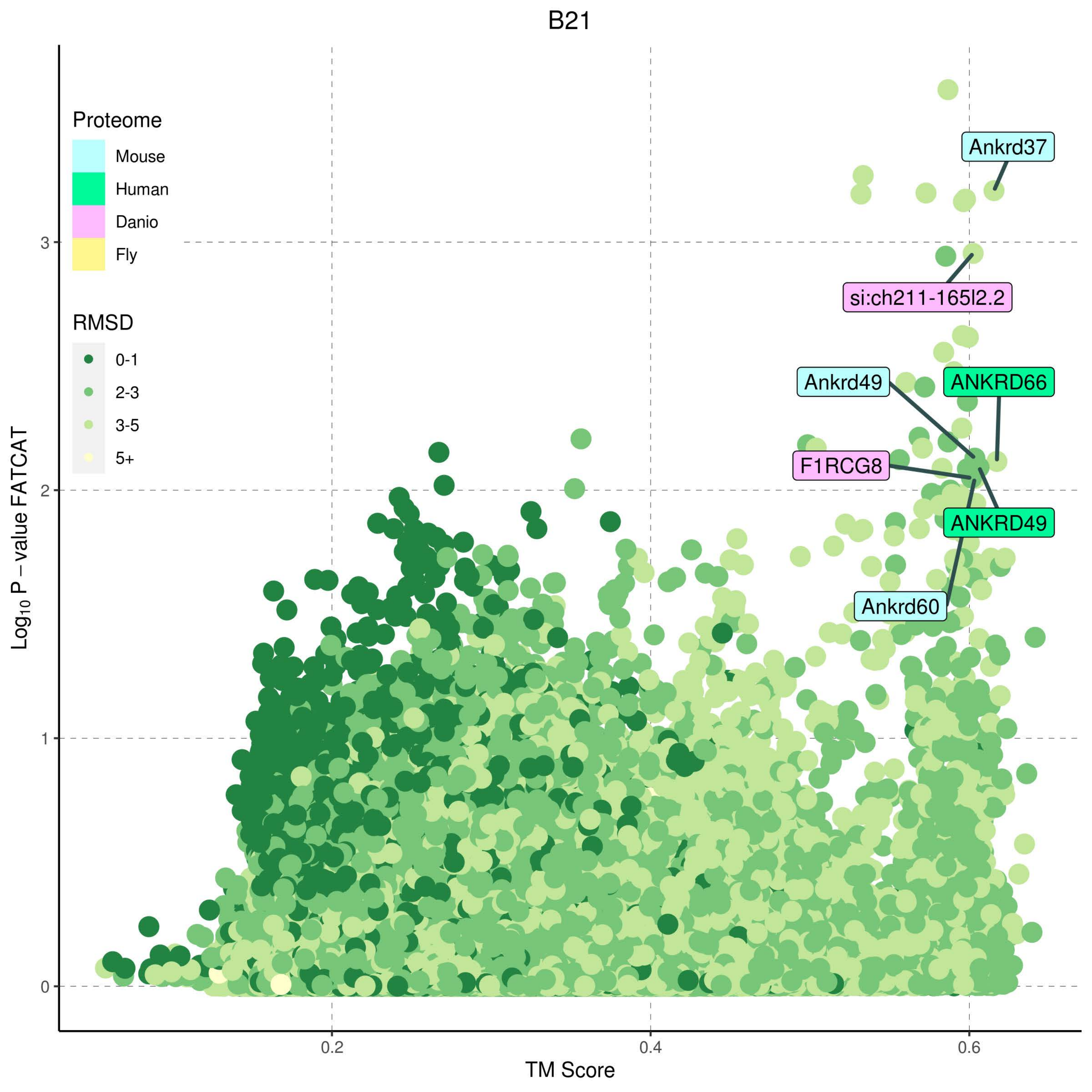

B21
Proteome
Mouse
Human
Danio
Fly
RMSD
0-1
2-3
3-5
5+
Ankrd37
si:ch211-165l2.2
Ankrd49
ANKRD66
F1RCG8
ANKRD49
Ankrd60
Log10 P – value FATCAT
TM Score
0
1
2
3
0.2
0.4
0.6

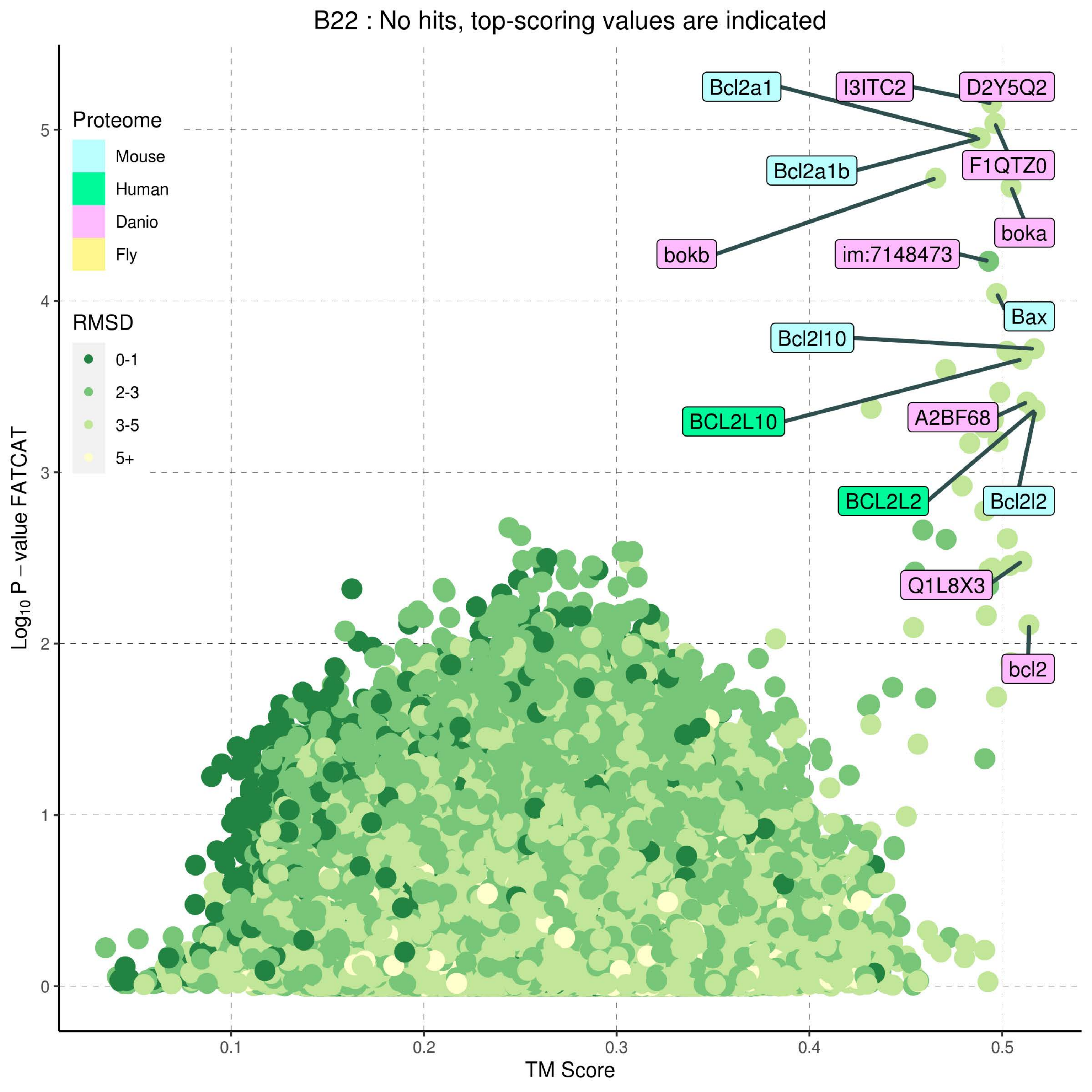
B22 : No hits, top-scoring values are indicated
Proteome
Mouse
Human
Danio
Fly
RMSD
0-1
2-3
3-5
5+
Bcl2a1
I3ITC2
D2Y5Q2
F1QTZ0
Bcl2a1b
boka
bokb
im:7148473
Bax
Bcl2l10
BCL2L10
A2BF68
BCL2L2
Bcl2l2
Q1L8X3
bcl2
Log10 P − value FATCAT
TM Score
0
1
2
3
4
5
0.1
0.2
0.3
0.4
0.5

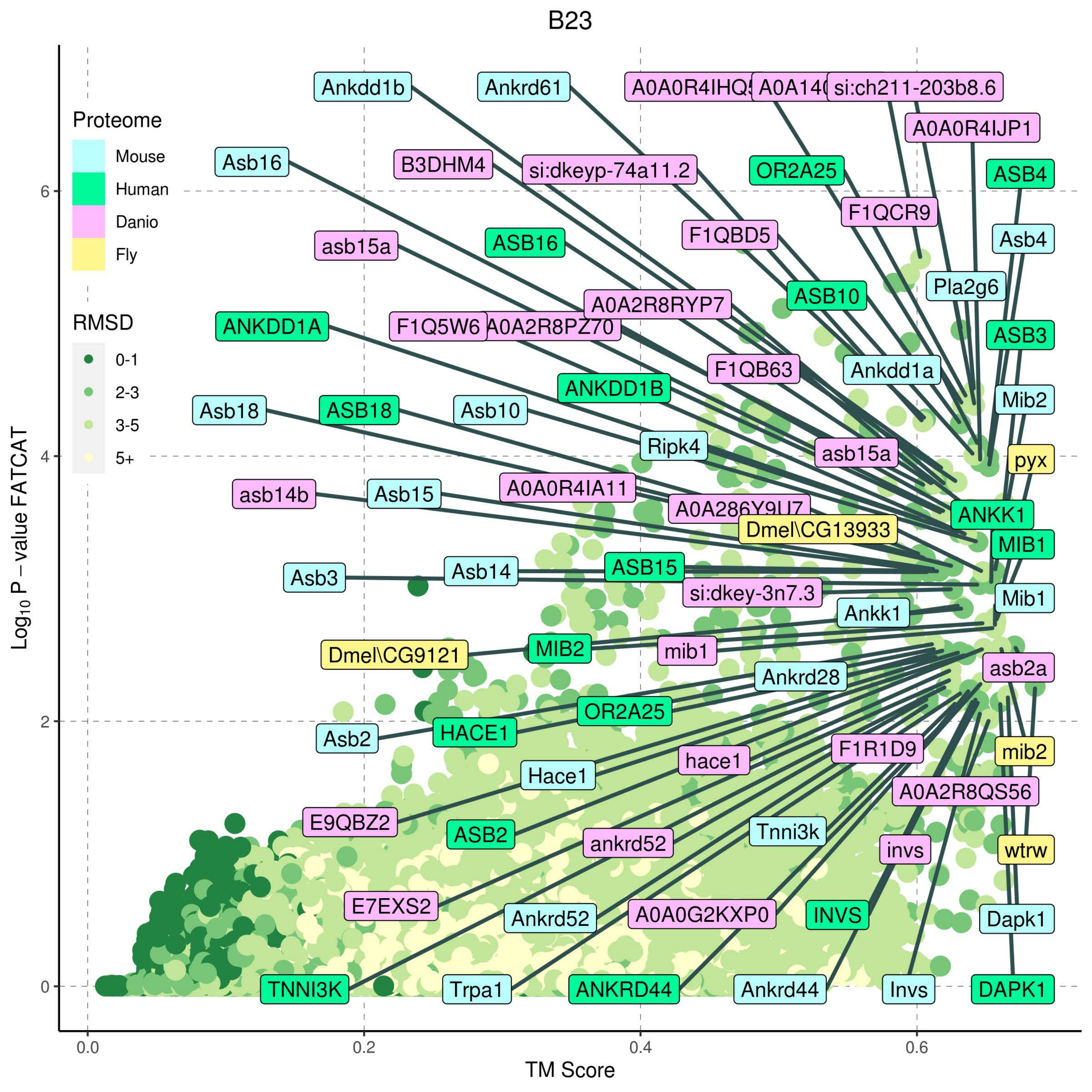
B23
Proteome
Mouse
Human
Danio
Fly
RMSD
0-1
2-3
3-5
5+
Ankdd1b
Ankrd61
A0A0R4IHQ5
si:ch211-203b8.6
A0A0R4IJP1
Asb16
B3DHM4
si:dkeyp-74a11.2
OR2A25
ASB4
F1QCR9
asb15a
ASB16
F1QBD5
Asb4
Pla2g6
ASB10
A0A2R8RYP7
ANKDD1A
F1Q5W6
A0A2R8PZ70
ASB3
F1QB63
Ankdd1a
ANKDD1B
Mib2
Asb18
ASB18
Asb10
Ripk4
asb15a
pyx
asb14b
Asb15
A0A0R4IA11
ANKK1
Dmel\CG13933
MIB1
Asb3
Asb14
ASB15
si:dkey-3n7.3
Mib1
Ankk1
Dmel\CG9121
MIB2
mib1
asb2a
Ankrd28
OR2A25
Asb2
HACE1
mib2
hace1
F1R1D9
Hace1
A0A2R8QS56
E9QBZ2
Tnni3k
ASB2
ankrd52
invs
wtrw
E7EXS2
Ankrd52
A0A0G2KXP0
INVS
Dapk1
TNNI3K
Trpa1
ANKRD44
Ankrd44
Invs
DAPK1
Log10 P - value FATCAT
TM Score
0
2
4
6
0.0
0.2
0.4
0.6

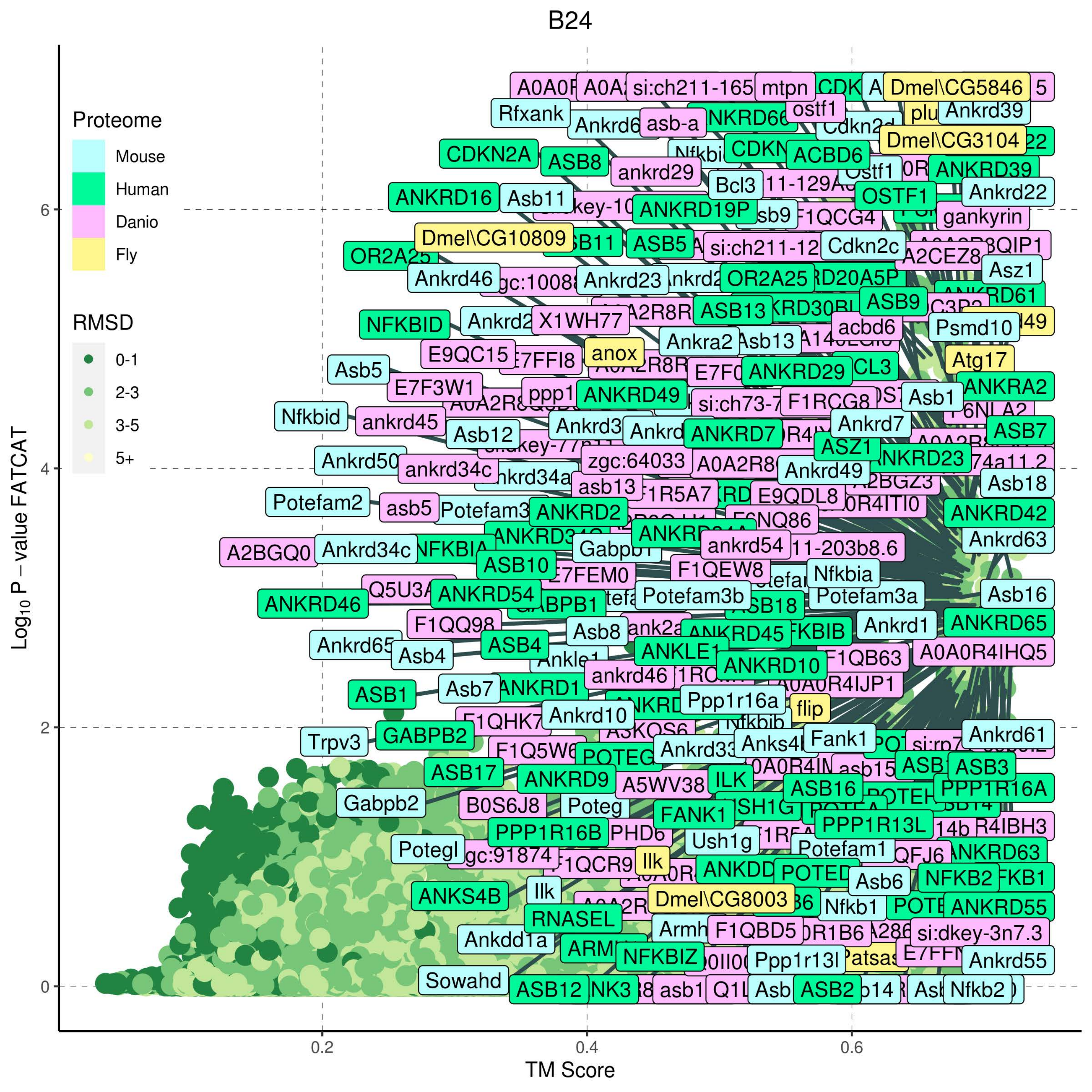

B24
Proteome
Mouse
Human
Danio
Fly
RMSD
0-1
2-3
3-5
5+
Log10 P − value FATCAT
TM Score
0
2
4
6
0.2
0.4
0.6
Dmel\CG5846
mtpn
Rfxank
asb-a
ostf1
Ankrd39
Dmel\CG3104
CDKN2A
ASB8
ACBD6
ankrd29
ANKRD39
Bcl3
ANKRD16
Asb11
ANKRD19P
OSTF1
Ankrd22
gankyrin
Dmel\CG10809
ASB5
Cdkn2c
A2CEZ8
Asz1
Ankrd46
Ankrd23
OR2A25
ASB9
ASB13
NFKBID
X1WH77
acbd6
Psmd10
anox
Ankra2
Asb13
E9QC15
Atg17
Asb5
ANKRD29
E7F3W1
ANKRA2
ANKRD49
F1RCG8
Asb1
Nfkbid
ankrd45
Ankrd7
Asb12
ANKRD7
ASB7
ASZ1
Ankrd50
zgc:64033
ankrd34c
Ankrd49
Asb18
asb13
E9QDL8
Potefam2
asb5
ANKRD2
ANKRD42
Ankrd63
A2BGQ0
Ankrd34c
ankrd54
ASB10
F1QEW8
Nfkbia
E7FEM0
Potefam3b
Potefam3a
ANKRD46
ANKRD54
Asb16
F1QQ98
Asb8
Ankrd1
ANKRD65
Ankrd65
Asb4
ASB4
ANKRD45
ANKLE1
A0A0R4IHQ5
ankrd46
ANKRD10
F1QB63
Asb7
ANKRD1
A0A0R4IJP1
ASB1
Ppp1r16a
flip
Ankrd10
Trpv3
GABPB2
Fank1
Ankrd61
ASB17
ANKRD9
A5WV38
ILK
ASB16
ASB3
PPP1R16A
Gabpb2
B0S6J8
Poteg
FANK1
PPP1R16B
PPP1R13L
Ush1g
Potegl
Potefam1
Ilk
ANKRD63
NFKB2
ANKS4B
Dmel\CG8003
Asb6
Nfkb1
ANKRD55
RNASEL
F1QBD5
Ankdd1a
si:dkey-3n7.3
NFKBIZ
Ppp1r13l
Ankrd55
Sowahd
ASB12
ASB2
Nfkb2

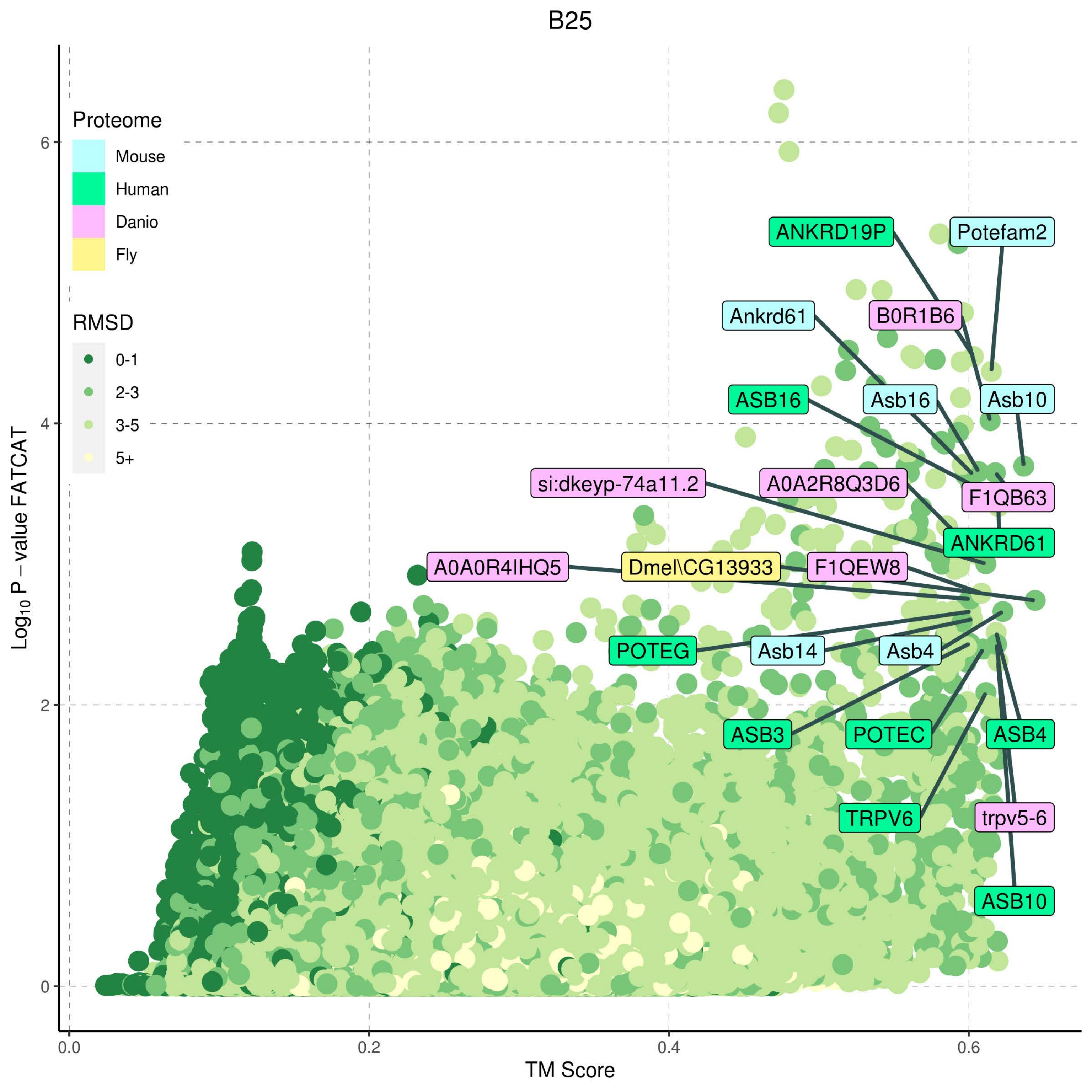

B25
Proteome
Mouse
Human
Danio
Fly
RMSD
0-1
2-3
3-5
5+
ANKRD19P
Potefam2
Ankrd61
B0R1B6
ASB16
Asb16
Asb10
si:dkeyp-74a11.2
A0A2R8Q3D6
F1QB63
ANKRD61
A0A0R4IHQ5
Dmel\CG13933
F1QEW8
POTEG
Asb14
Asb4
ASB3
POTEC
ASB4
TRPV6
trpv5-6
ASB10
Log10 P – value FATCAT
TM Score
0
2
4
6
0.0
0.2
0.4
0.6

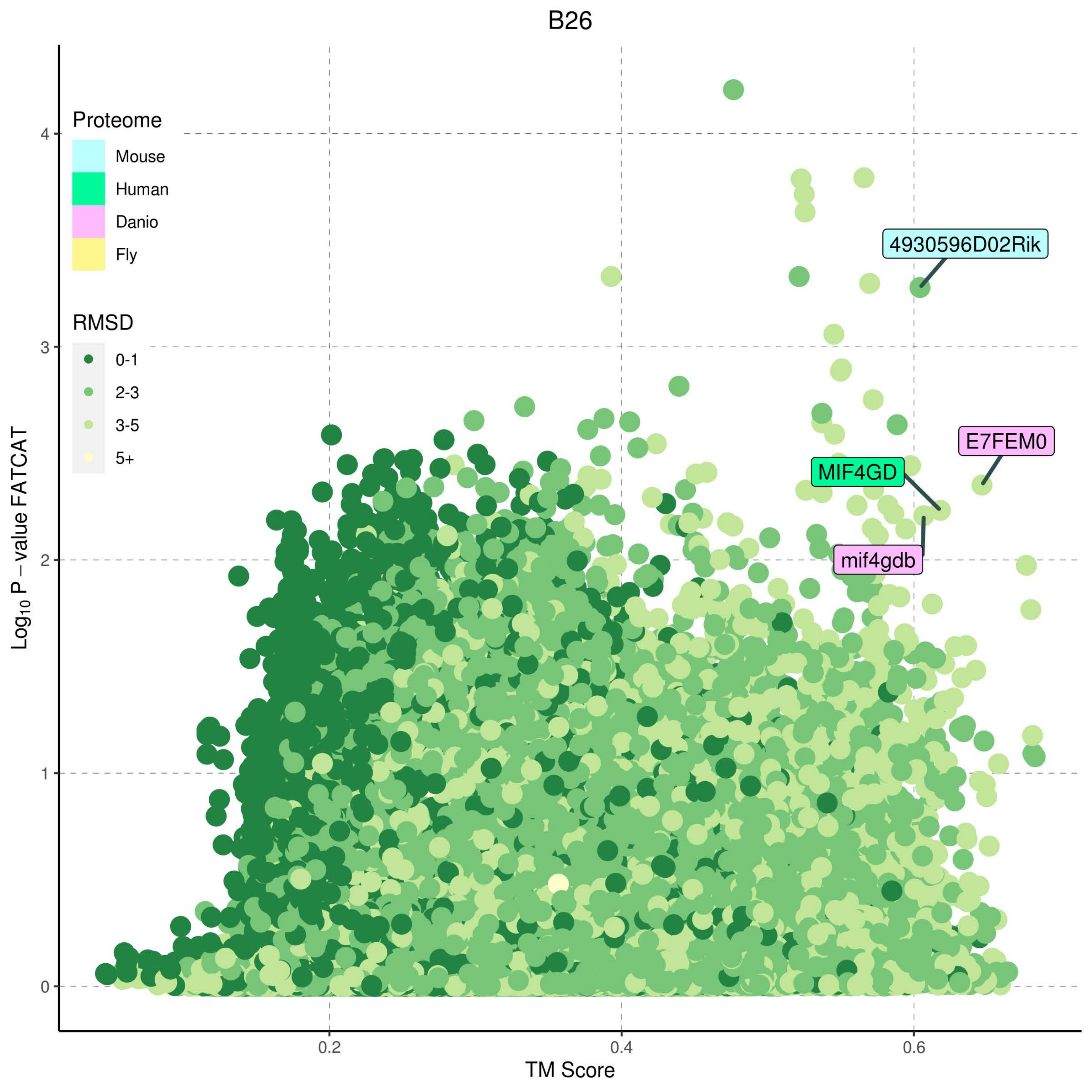
B26
Proteome
Mouse
Human
Danio
Fly
RMSD
0-1
2-3
3-5
5+
4930596D02Rik
E7FEM0
MIF4GD
mif4gdb
Log10 P − value FATCAT
TM Score
0
1
2
3
4
0.2
0.4
0.6

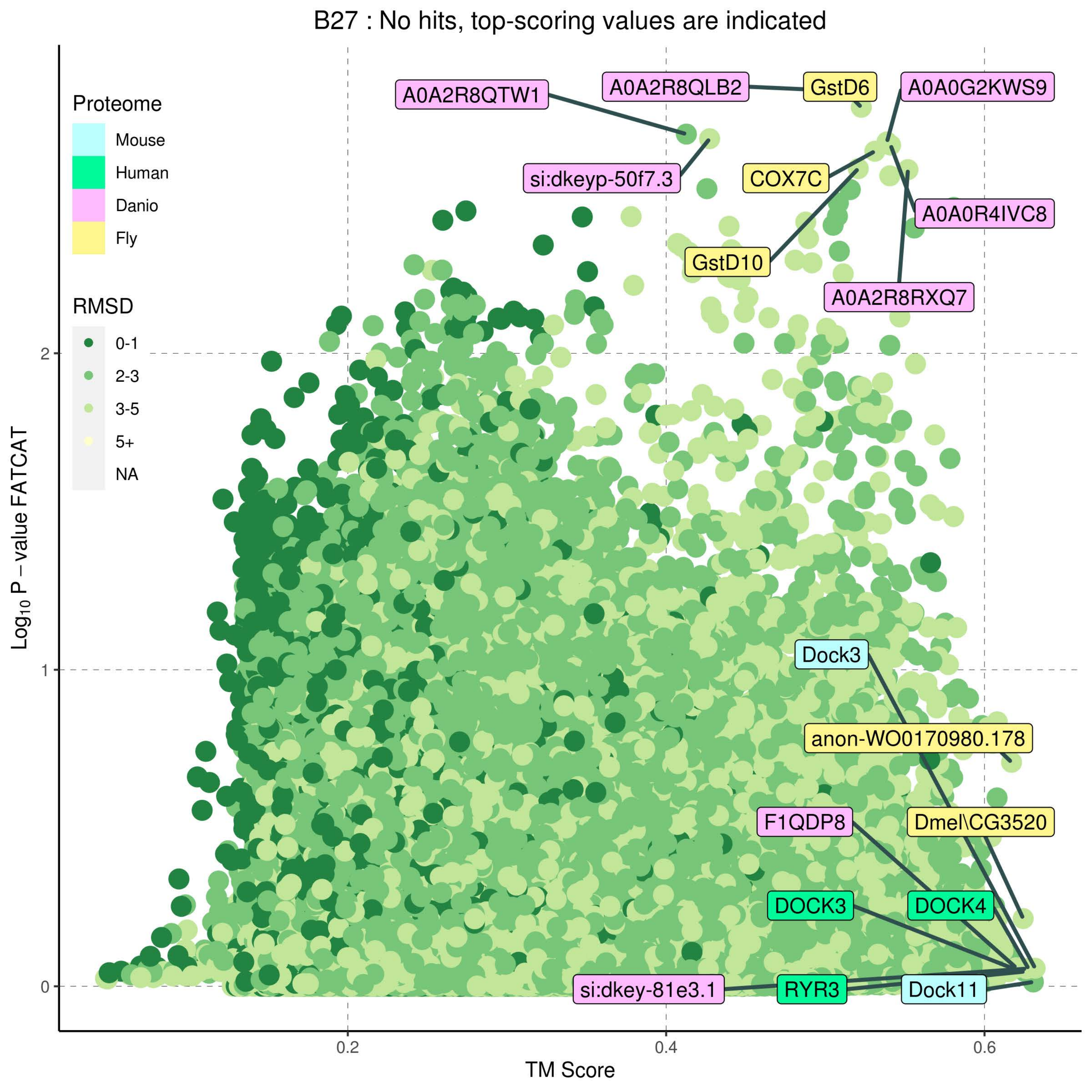
B27 : No hits, top-scoring values are indicated
Proteome
Mouse
Human
Danio
Fly
RMSD
0-1
2-3
3-5
5+
NA
Log10 P – value FATCAT
TM Score
0
1
2
0.2
0.4
0.6
A0A2R8QTW1
A0A2R8QLB2
GstD6
A0A0G2KWS9
si:dkeyp-50f7.3
COX7C
A0A0R4IVC8
GstD10
A0A2R8RXQ7
Dock3
anon-WO0170980.178
F1QDP8
Dmel\CG3520
DOCK3
DOCK4
si:dkey-81e3.1
RYR3
Dock11

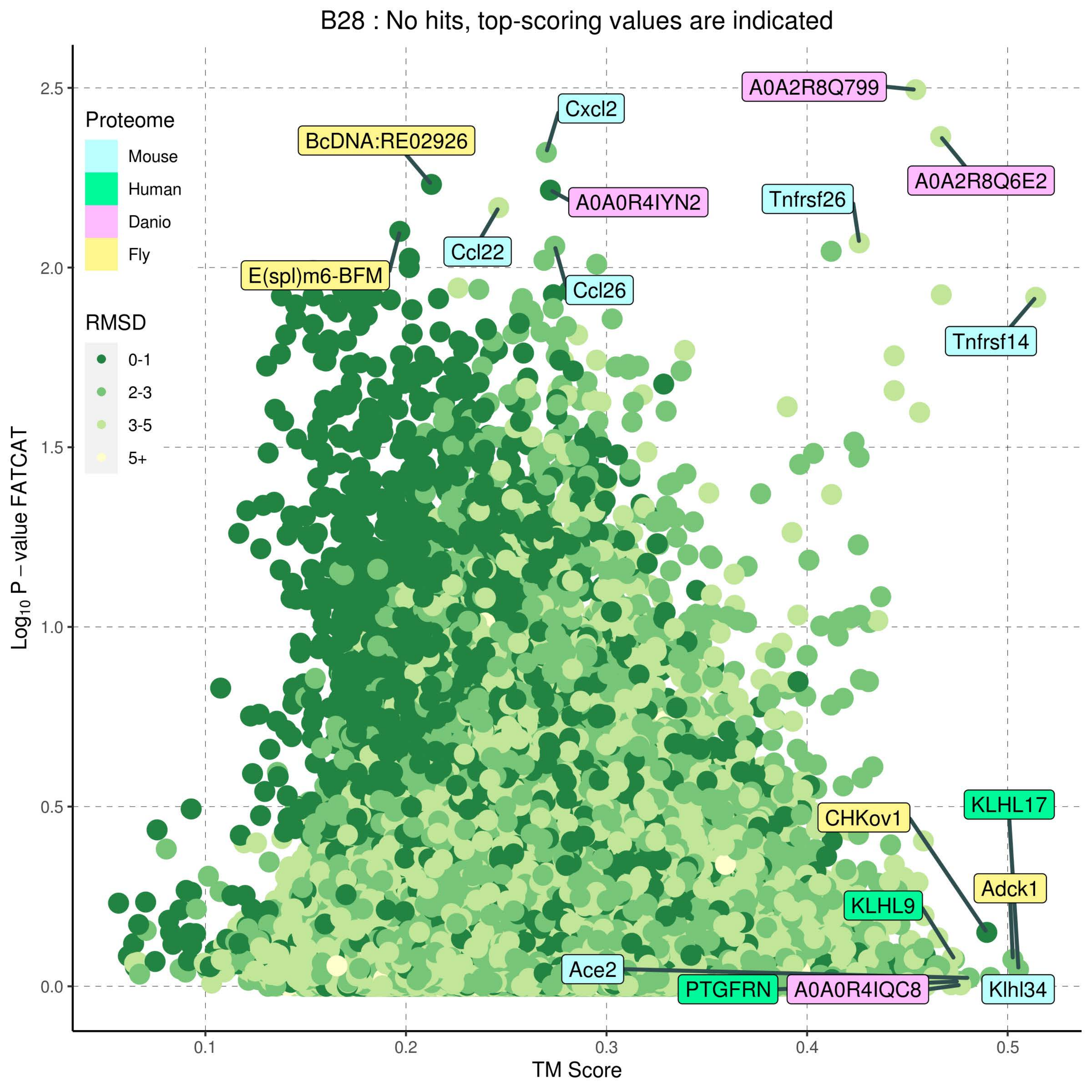

B28 : No hits, top-scoring values are indicated
Proteome
Mouse
Human
Danio
Fly
RMSD
0-1
2-3
3-5
5+
A0A2R8Q799
Cxcl2
BcDNA:RE02926
A0A2R8Q6E2
A0A0R4IYN2
Tnfrsf26
Ccl22
E(spl)m6-BFM
Ccl26
Tnfrsf14
KLHL17
CHKov1
Adck1
KLHL9
Ace2
PTGFRN
A0A0R4IQC8
Klhl34
Log10 P − value FATCAT
TM Score
2.5
2.0
1.5
1.0
0.5
0.0
0.1
0.2
0.3
0.4
0.5

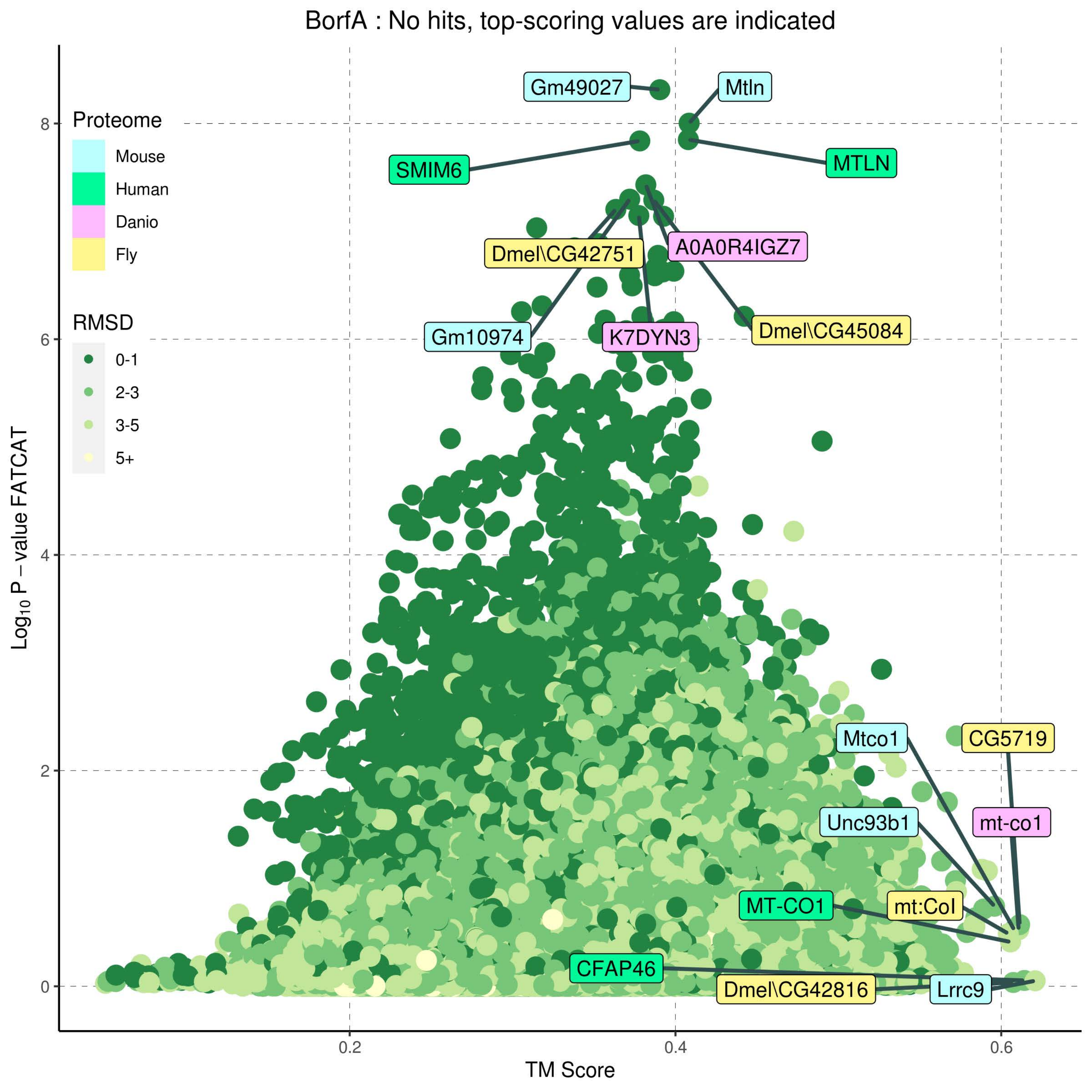
BorfA : No hits, top-scoring values are indicated
Proteome
Mouse
Human
Danio
Fly
RMSD
0-1
2-3
3-5
5+
Gm49027
Mtln
SMIM6
MTLN
Dmel\CG42751
A0A0R4IGZ7
Gm10974
K7DYN3
Dmel\CG45084
Mtco1
CG5719
Unc93b1
mt-co1
MT-CO1
mt:CoI
CFAP46
Dmel\CG42816
Lrrc9
0
2
4
6
8
0.2
0.4
0.6
TM Score
Log10 P – value FATCAT

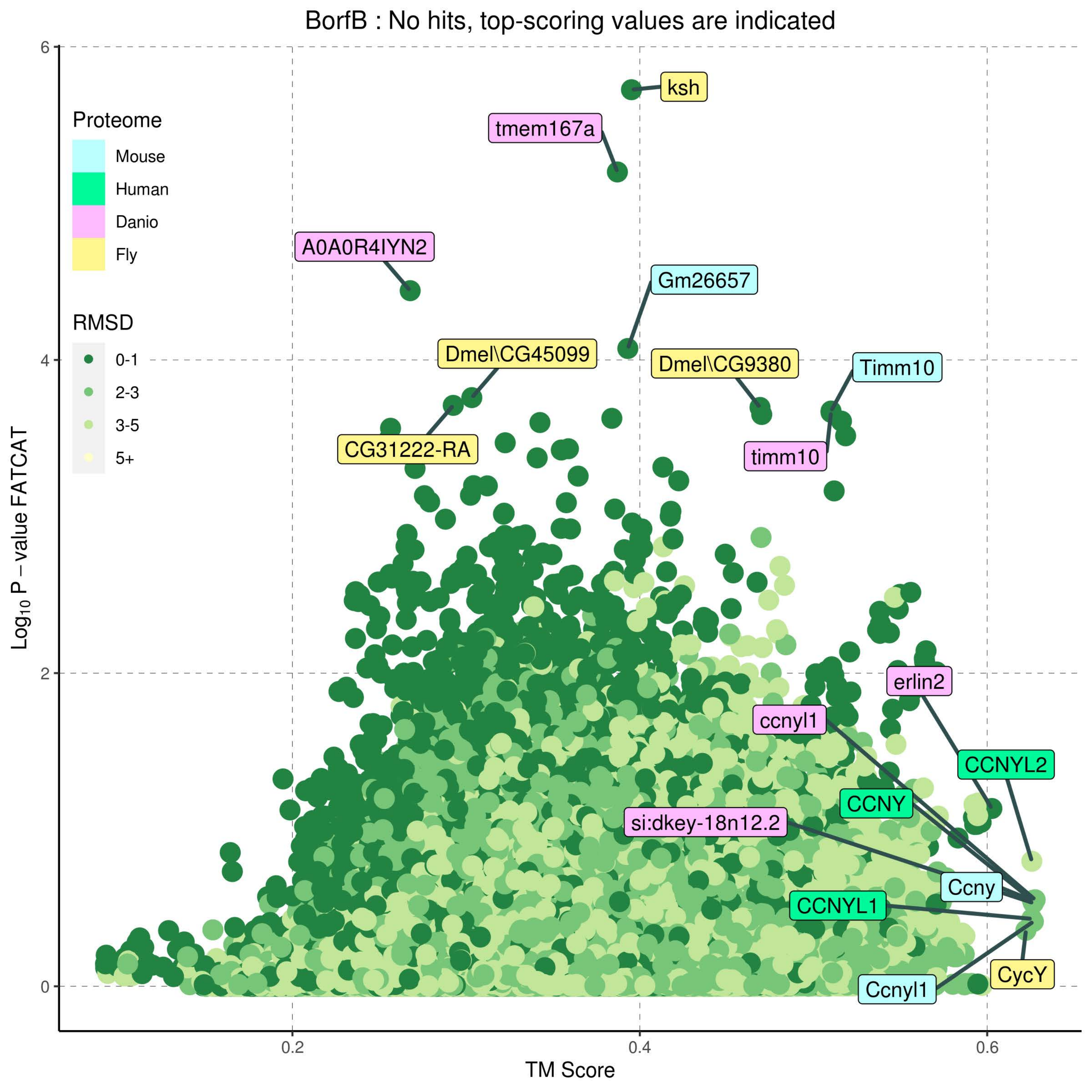

BorfB : No hits, top-scoring values are indicated
Proteome
Mouse
Human
Danio
Fly
RMSD
0-1
2-3
3-5
5+
ksh
tmem167a
A0A0R4IYN2
Gm26657
Dmel\CG45099
Dmel\CG9380
Timm10
CG31222-RA
timm10
erlin2
ccnyl1
CCNYL2
CCNY
si:dkey-18n12.2
Ccny
CCNYL1
CycY
Ccnyl1
Log10 P − value FATCAT
TM Score
0
2
4
6
0.2
0.4
0.6

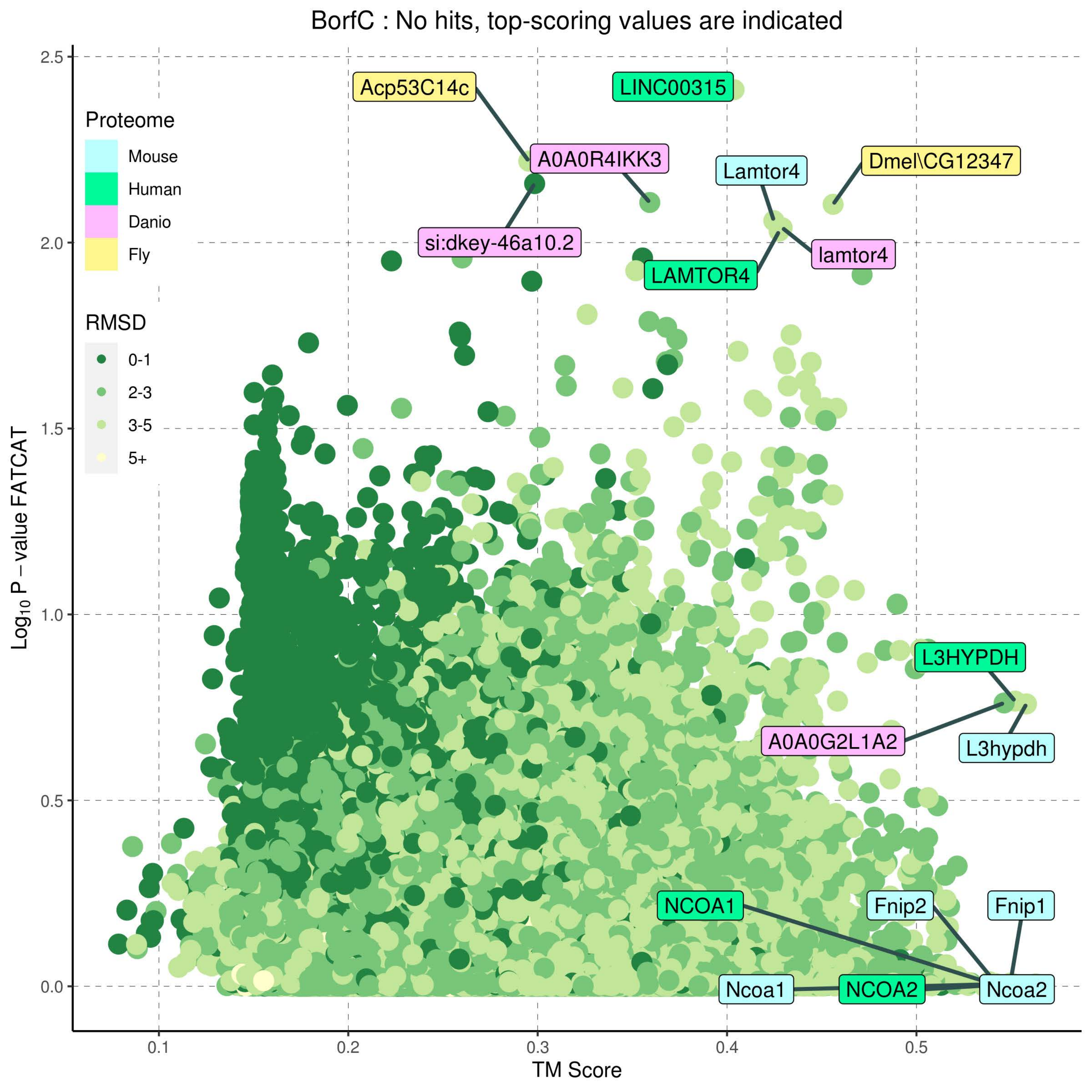

BorfC : No hits, top-scoring values are indicated
Proteome
Mouse
Human
Danio
Fly
RMSD
0-1
2-3
3-5
5+
Acp53C14c
LINC00315
A0A0R4IKK3
Lamtor4
Dmel\CG12347
si:dkey-46a10.2
lamtor4
LAMTOR4
L3HYPDH
A0A0G2L1A2
L3hypdh
NCOA1
Fnip2
Fnip1
Ncoa1
NCOA2
Ncoa2
2.5
2.0
1.5
1.0
0.5
0.0
0.1
0.2
0.3
0.4
0.5
TM Score
$Log_{10}$ P – value FATCAT

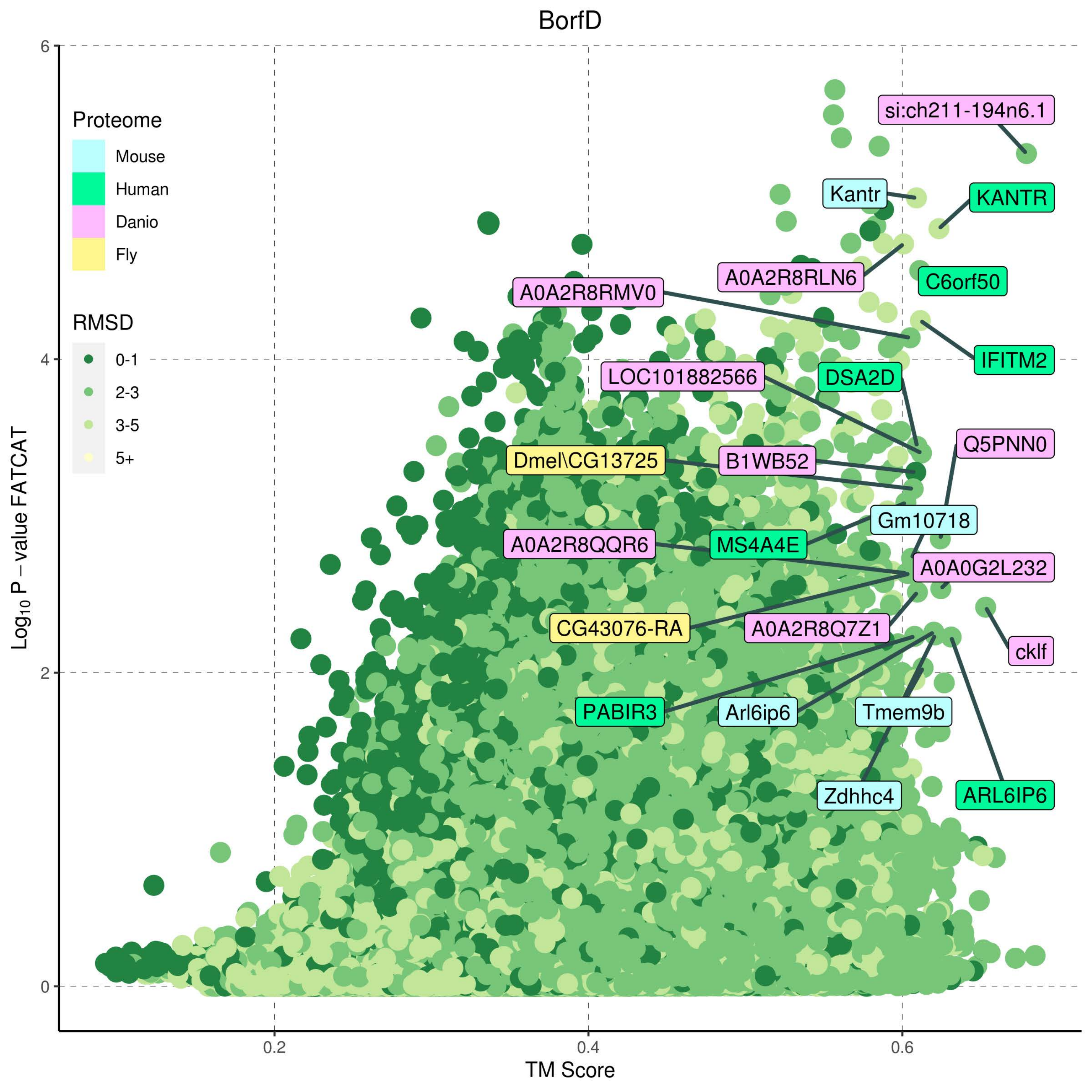

BorfD
Proteome
Mouse
Human
Danio
Fly
RMSD
0-1
2-3
3-5
5+
si:ch211-194n6.1
Kantr
KANTR
A0A2R8RLN6
C6orf50
A0A2R8RMV0
IFITM2
LOC101882566
DSA2D
Q5PNN0
Dmel\CG13725
B1WB52
Gm10718
A0A2R8QQR6
MS4A4E
A0A0G2L232
CG43076-RA
A0A2R8Q7Z1
cklf
PABIR3
Arl6ip6
Tmem9b
Zdhhc4
ARL6IP6
Log10 P – value FATCAT
TM Score
0
2
4
6
0.2
0.4
0.6

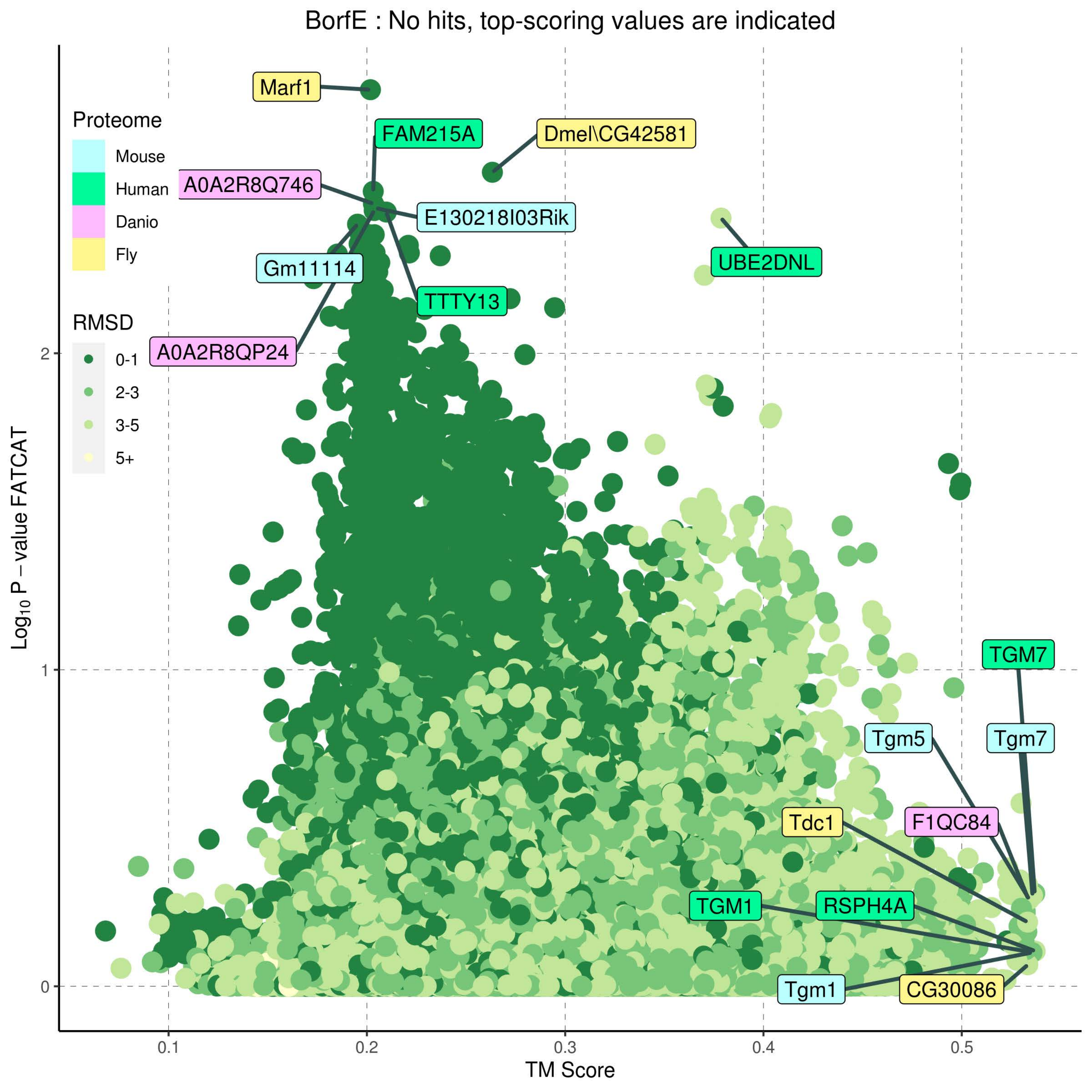
BorfE : No hits, top-scoring values are indicated
Proteome
Mouse
Human
Danio
Fly
RMSD
0-1
2-3
3-5
5+
Marf1
FAM215A
Dmel\CG42581
A0A2R8Q746
E130218I03Rik
Gm11114
UBE2DNL
TTTY13
A0A2R8QP24
TGM7
Tgm5
Tgm7
Tdc1
F1QC84
TGM1
RSPH4A
Tgm1
CG30086
Log10 P – value FATCAT
TM Score
0
1
2
0.1
0.2
0.3
0.4
0.5

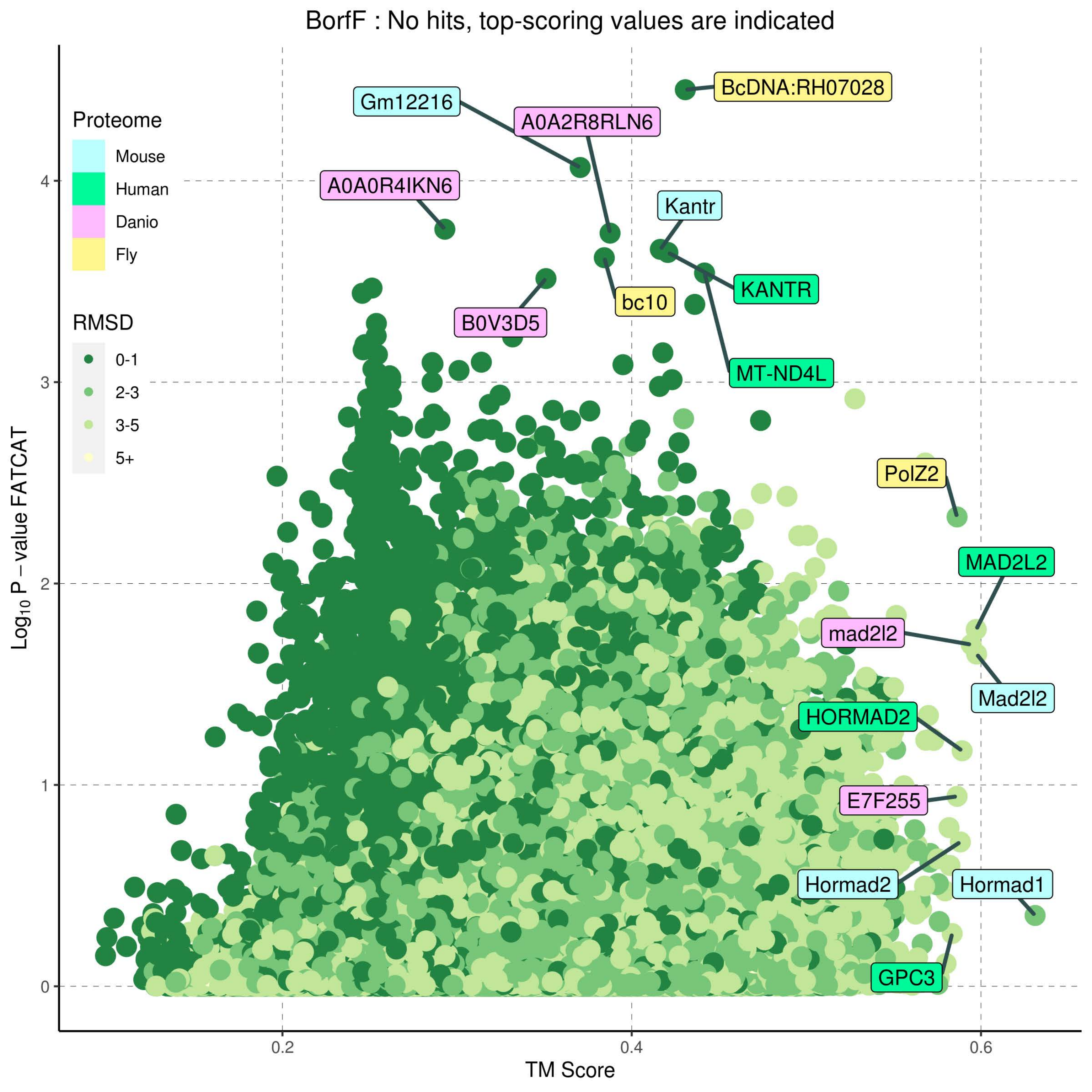
BorfF : No hits, top-scoring values are indicated
Proteome
Mouse
Human
Danio
Fly
RMSD
0-1
2-3
3-5
5+
Gm12216
A0A2R8RLN6
BcDNA:RH07028
A0A0R4IKN6
Kantr
KANTR
bc10
B0V3D5
MT-ND4L
PolZ2
MAD2L2
mad2l2
Mad2l2
HORMAD2
E7F255
Hormad2
Hormad1
GPC3
0
1
2
3
4
0.2
0.4
0.6
TM Score
Log10 P – value FATCAT

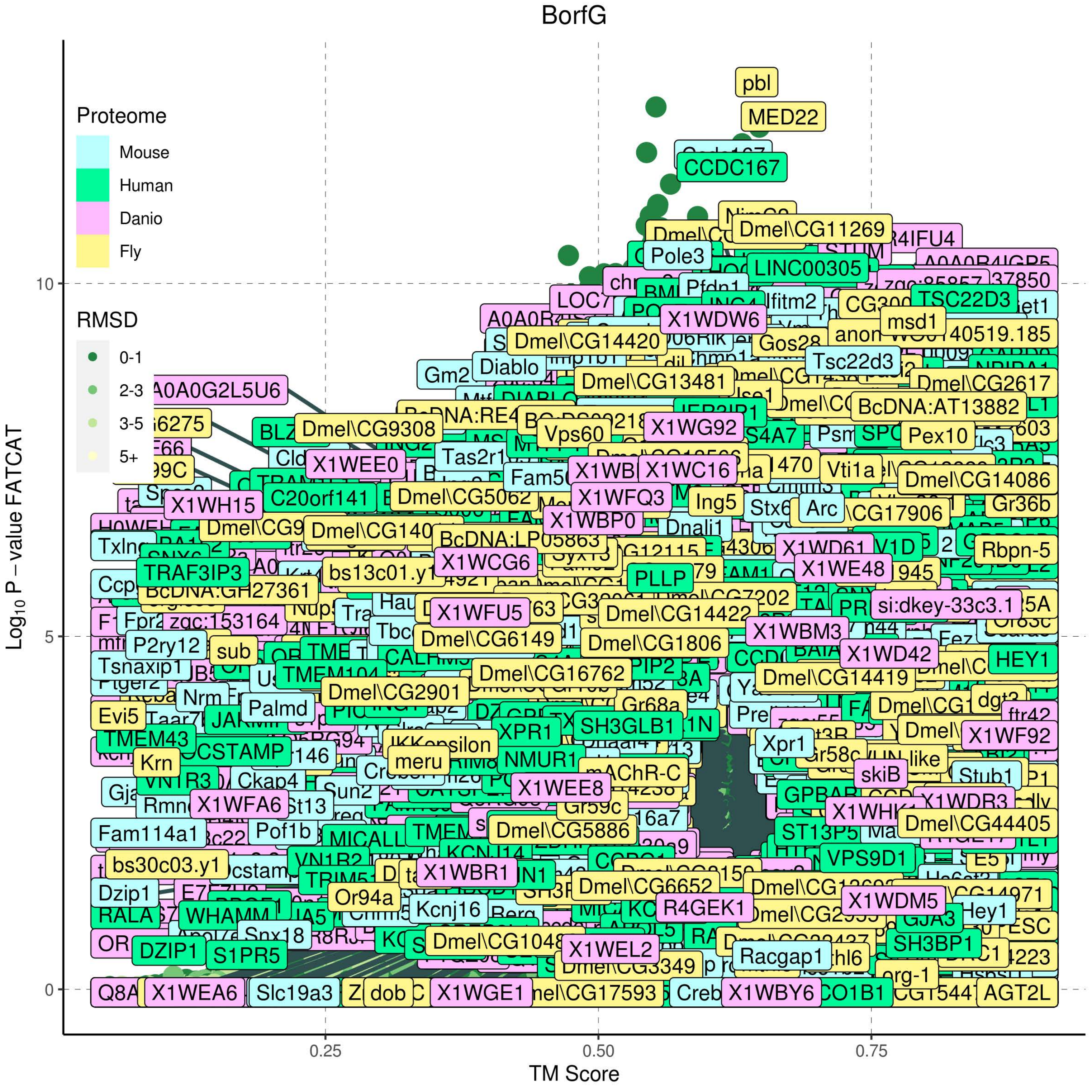
BorfG
Proteome
Mouse
Human
Danio
Fly
RMSD
0-1
2-3
3-5
5+
Log10 P − value FATCAT
TM Score
0
5
10
0.25
0.50
0.75
pbl
MED22
CCDC167
Dmel\CG11269
Pole3
LINC00305
Pfdn1
Ifitm2
TSC22D3
X1WDW6
msd1
Dmel\CG14420
Gos28
Tsc22d3
Diablo
Dmel\CG13481
Dmel\CG2617
BcDNA:AT13882
A0A0G2L5U6
Dmel\CG9308
X1WG92
Vps60
Pex10
X1WEE0
Tas2r1
X1WC16
Vti1a
Dmel\CG14086
C20orf141
Dmel\CG5062
X1WFQ3
Ing5
Arc
X1WH15
X1WBP0
Gr36b
Dnali1
BcDNA:LP05863
Rbpn-5
X1WCG6
TRAF3IP3
bs13c01.y1
PLLP
X1WE48
si:dkey-33c3.1
zgc:153164
X1WFU5
Dmel\CG14422
X1WBM3
P2ry12
sub
Dmel\CG6149
Dmel\CG1806
X1WD42
HEY1
Tsnaxip1
Dmel\CG16762
Dmel\CG14419
Dmel\CG2901
Palmd
Evi5
SH3GLB1
XPR1
TMEM43
Xpr1
X1WF92
Krn
meru
NMUR1
mAChR-C
skiB
Stub1
X1WEE8
Ckap4
X1WFA6
Gr59c
Dmel\CG44405
Fam114a1
Pof1b
Dmel\CG5886
ST13P5
bs30c03.y1
VPS9D1
X1WBR1
Dzip1
Or94a
Dmel\CG6652
R4GEK1
X1WDM5
Kcnj16
Hey1
RALA
Snx18
Dmel\CG1048
X1WEL2
SH3BP1
DZIP1
S1PR5
Racgap1
org-1
X1WEA6
Slc19a3
dob
X1WGE1
X1WBY6
AGT2L

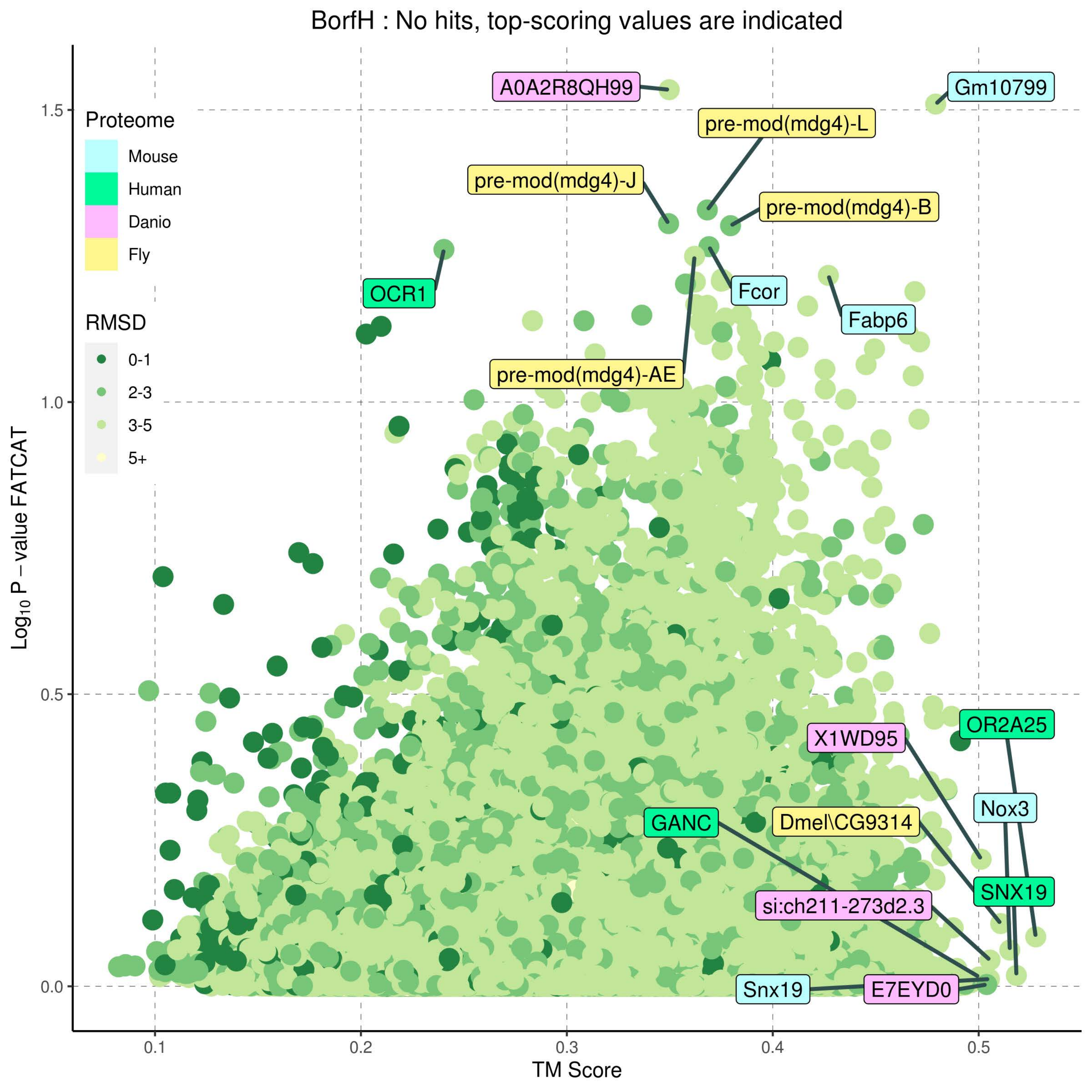
BorfH : No hits, top-scoring values are indicated
Proteome
Mouse
Human
Danio
Fly
RMSD
0-1
2-3
3-5
5+
A0A2R8QH99
Gm10799
pre-mod(mdg4)-L
pre-mod(mdg4)-J
pre-mod(mdg4)-B
OCR1
Fcor
Fabp6
pre-mod(mdg4)-AE
OR2A25
X1WD95
Nox3
GANC
Dmel\CG9314
SNX19
si:ch211-273d2.3
Snx19
E7EYD0
Log10 P - value FATCAT
TM Score
1.5
1.0
0.5
0.0
0.1
0.2
0.3
0.4
0.5

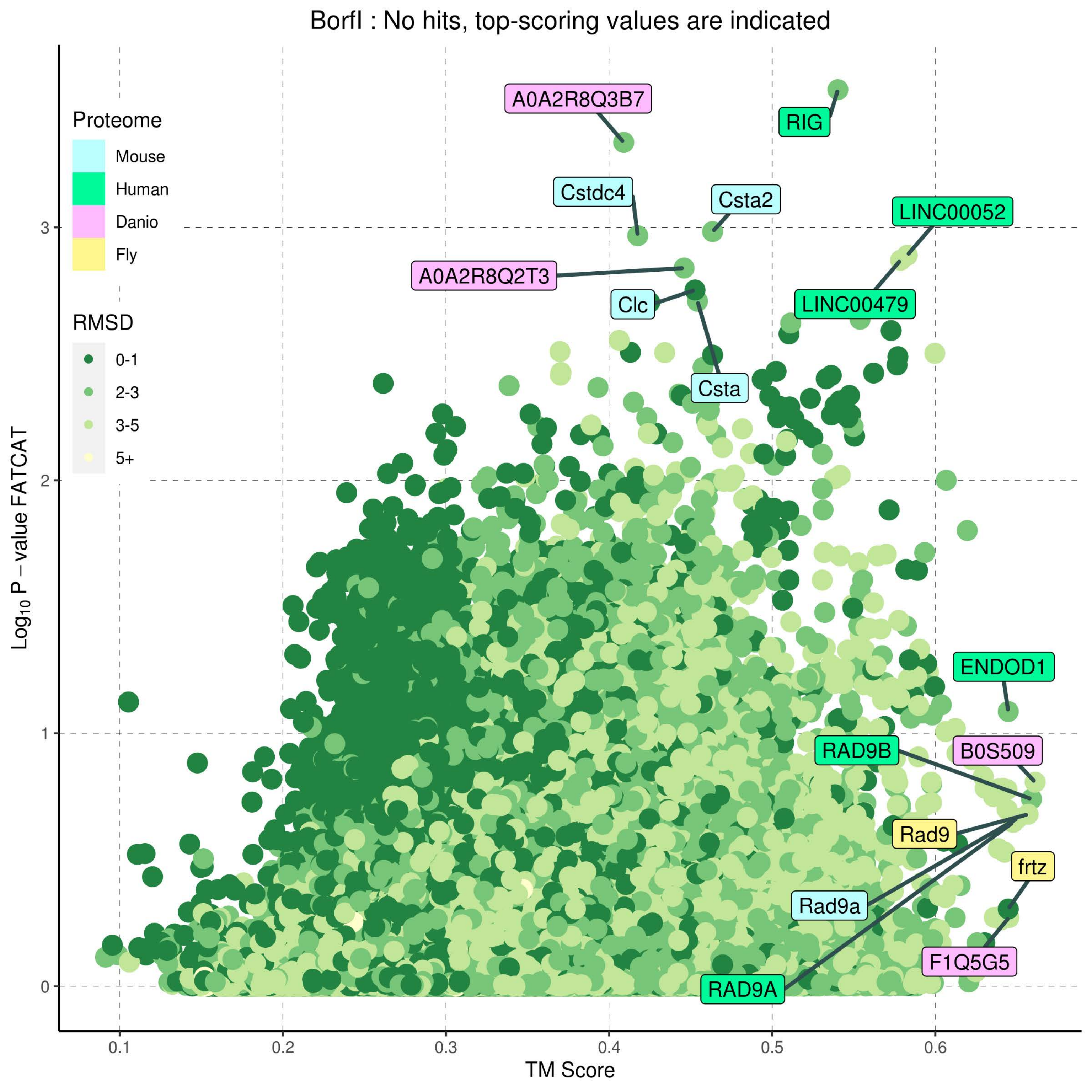

BorfI : No hits, top-scoring values are indicated
Proteome
Mouse
Human
Danio
Fly
RMSD
0-1
2-3
3-5
5+
A0A2R8Q3B7
RIG
Cstdc4
Csta2
LINC00052
A0A2R8Q2T3
Clc
LINC00479
Csta
ENDOD1
RAD9B
B0S509
Rad9
frtz
Rad9a
F1Q5G5
RAD9A
0
1
2
3
0.1
0.2
0.3
0.4
0.5
0.6
TM Score
Log10 P − value FATCAT

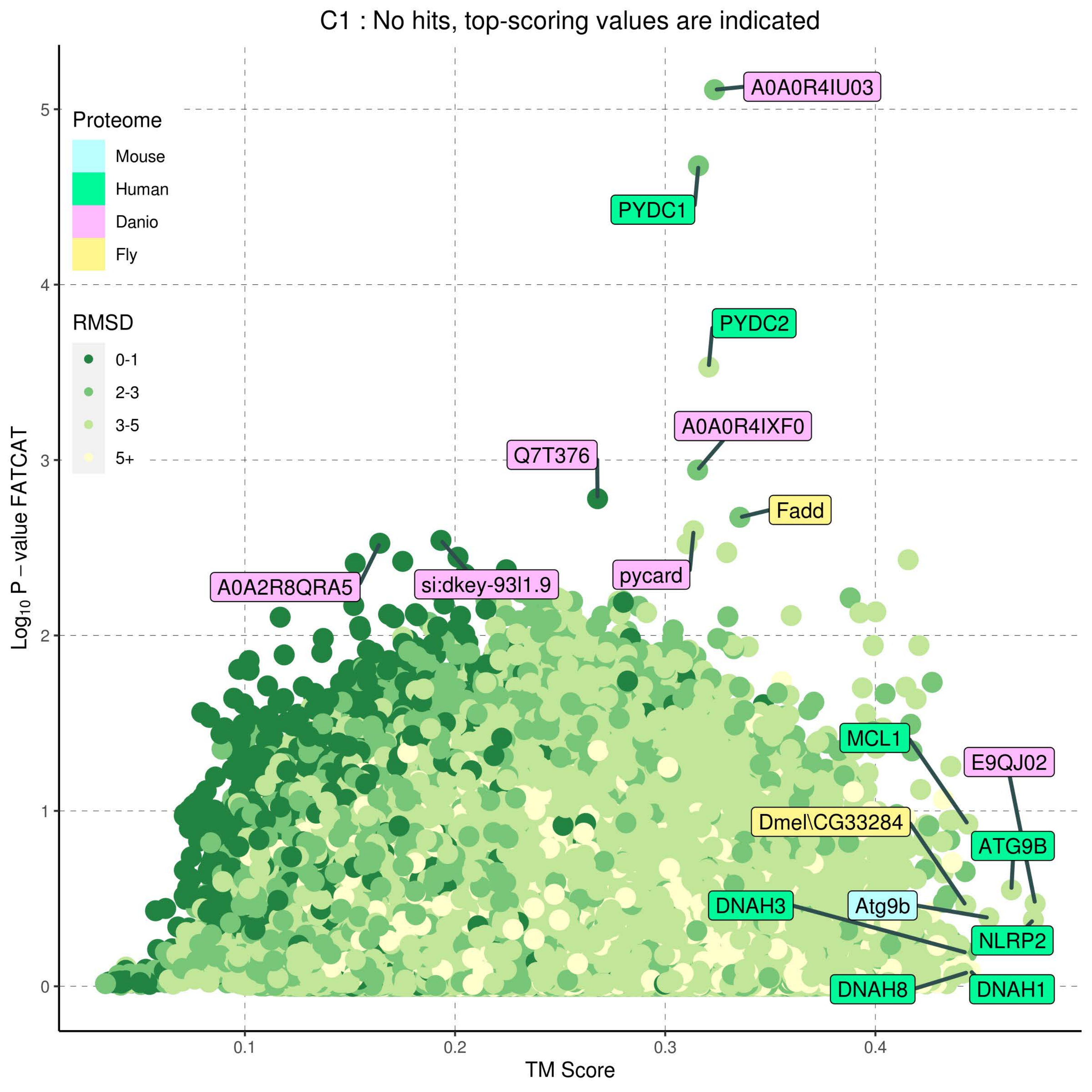
C1 : No hits, top-scoring values are indicated
Proteome
Mouse
Human
Danio
Fly
RMSD
0-1
2-3
3-5
5+
A0A0R4IU03
PYDC1
PYDC2
A0A0R4IXF0
Q7T376
Fadd
A0A2R8QRA5
si:dkey-93l1.9
pycard
MCL1
E9QJ02
Dmel\CG33284
ATG9B
DNAH3
Atg9b
NLRP2
DNAH8
DNAH1
Log10 P − value FATCAT
TM Score
0
1
2
3
4
5
0.1
0.2
0.3
0.4

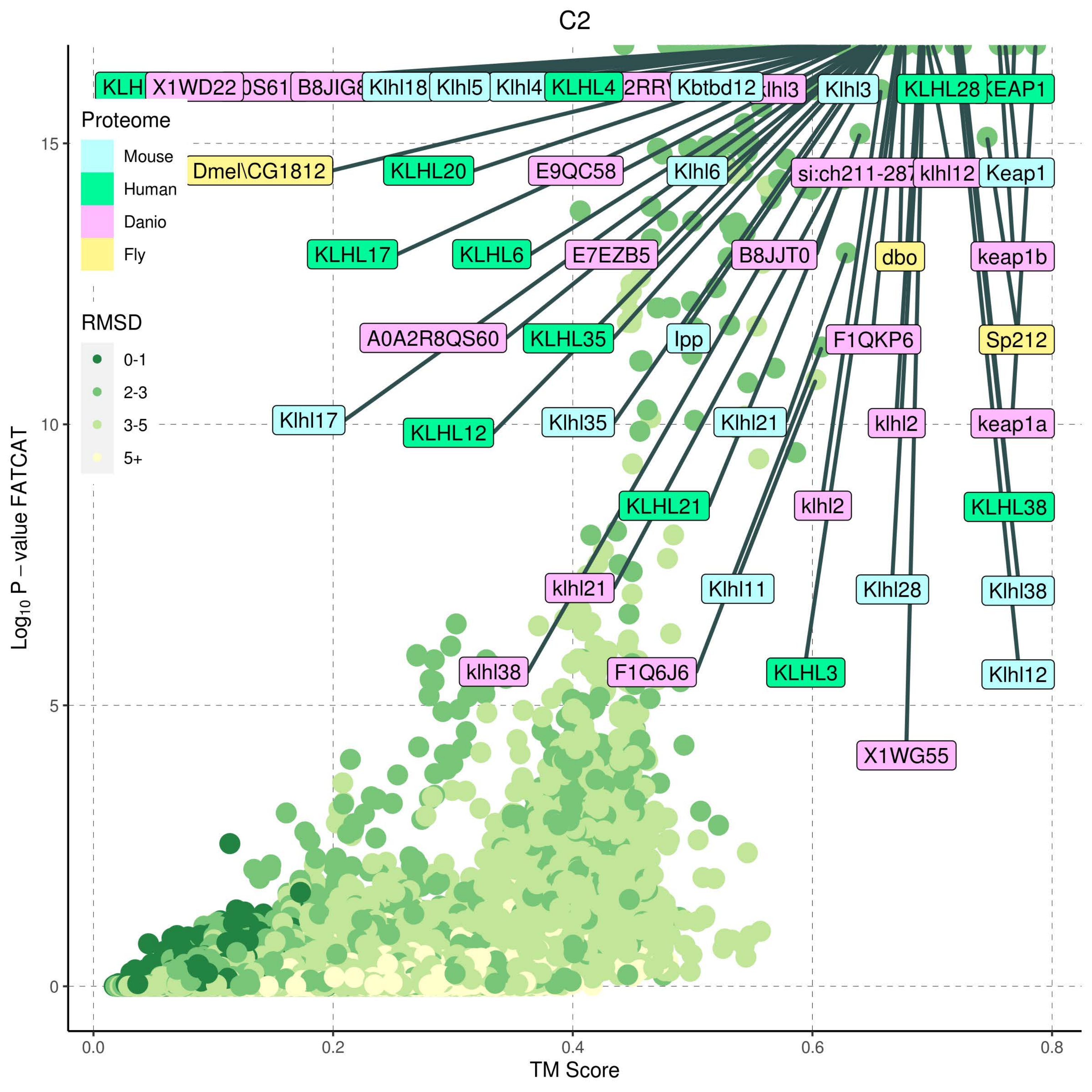

C2
Proteome
Mouse
Human
Danio
Fly
RMSD
0-1
2-3
3-5
5+
KLH
X1WD22
0S61
B8JIG8
Klhl18
Klhl5
Klhl4
KLHL4
2RRY
Kbtbd12
klhl3
Klhl3
KLHL28
KEAP1
Dmel\CG1812
KLHL20
E9QC58
Klhl6
si:ch211-287
klhl12
Keap1
KLHL17
KLHL6
E7EZB5
B8JJT0
dbo
keap1b
A0A2R8QS60
KLHL35
lpp
F1QKP6
Sp212
Klhl17
KLHL12
Klhl35
Klhl21
klhl2
keap1a
KLHL21
klhl2
KLHL38
klhl21
Klhl11
Klhl28
Klhl38
klhl38
F1Q6J6
KLHL3
Klhl12
X1WG55
Log10 P − value FATCAT
TM Score
0
5
10
15
0.0
0.2
0.4
0.6
0.8

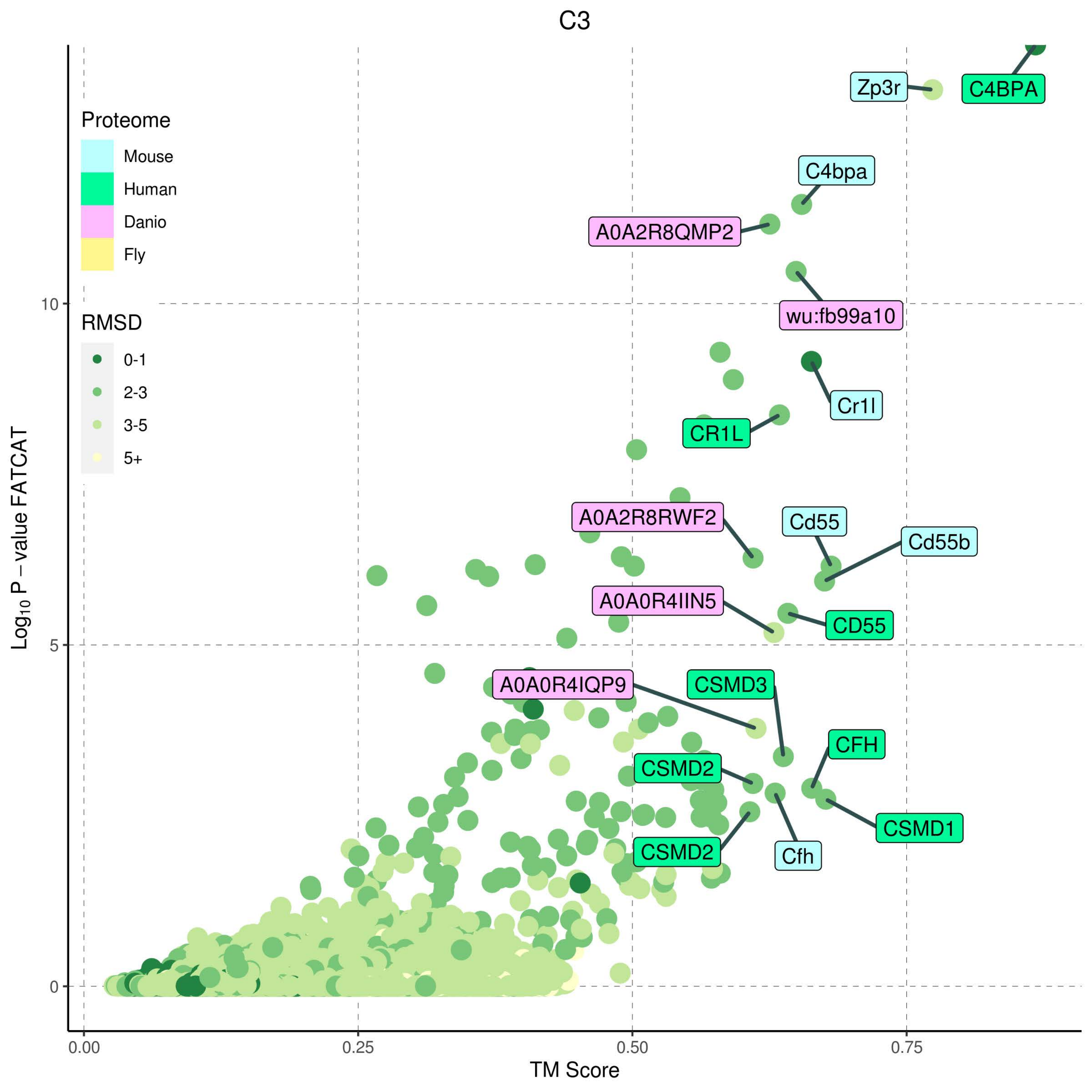

C3
Proteome
Mouse
Human
Danio
Fly
RMSD
0-1
2-3
3-5
5+
Zp3r
C4BPA
C4bpa
A0A2R8QMP2
wu:fb99a10
Cr1l
CR1L
A0A2R8RWF2
Cd55
Cd55b
A0A0R4IIN5
CD55
A0A0R4IQP9
CSMD3
CFH
CSMD2
CSMD1
CSMD2
Cfh
Log10 P - value FATCAT
10
5
0
0.00
0.25
0.50
0.75
TM Score

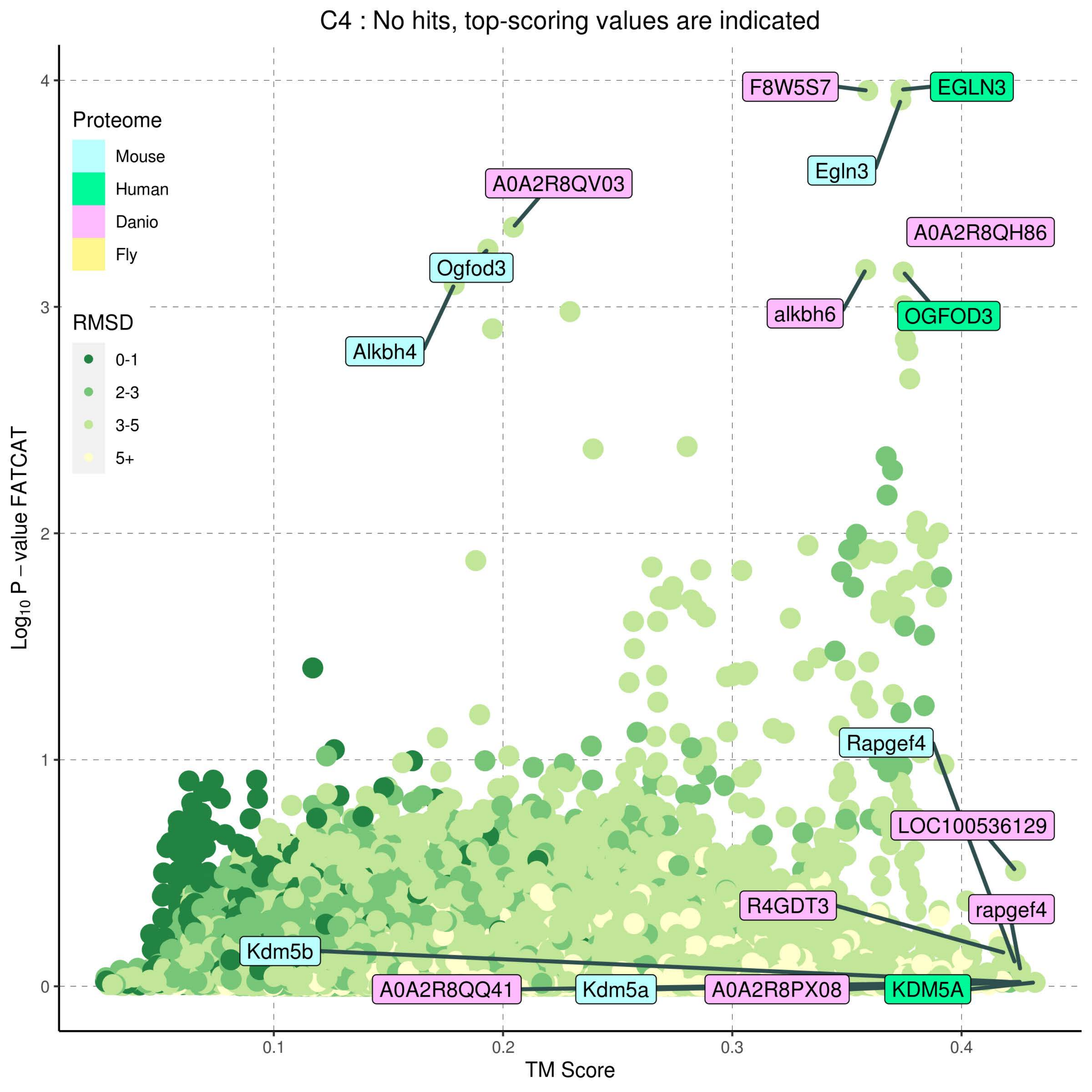

C4 : No hits, top-scoring values are indicated
Proteome
Mouse
Human
Danio
Fly
RMSD
0-1
2-3
3-5
5+
Log10 P – value FATCAT
TM Score
0
1
2
3
4
0.1
0.2
0.3
0.4
F8W5S7
EGLN3
Egln3
A0A2R8QV03
A0A2R8QH86
Ogfod3
alkbh6
OGFOD3
Alkbh4
Rapgef4
LOC100536129
R4GDT3
rapgef4
Kdm5b
A0A2R8QQ41
Kdm5a
A0A2R8PX08
KDM5A

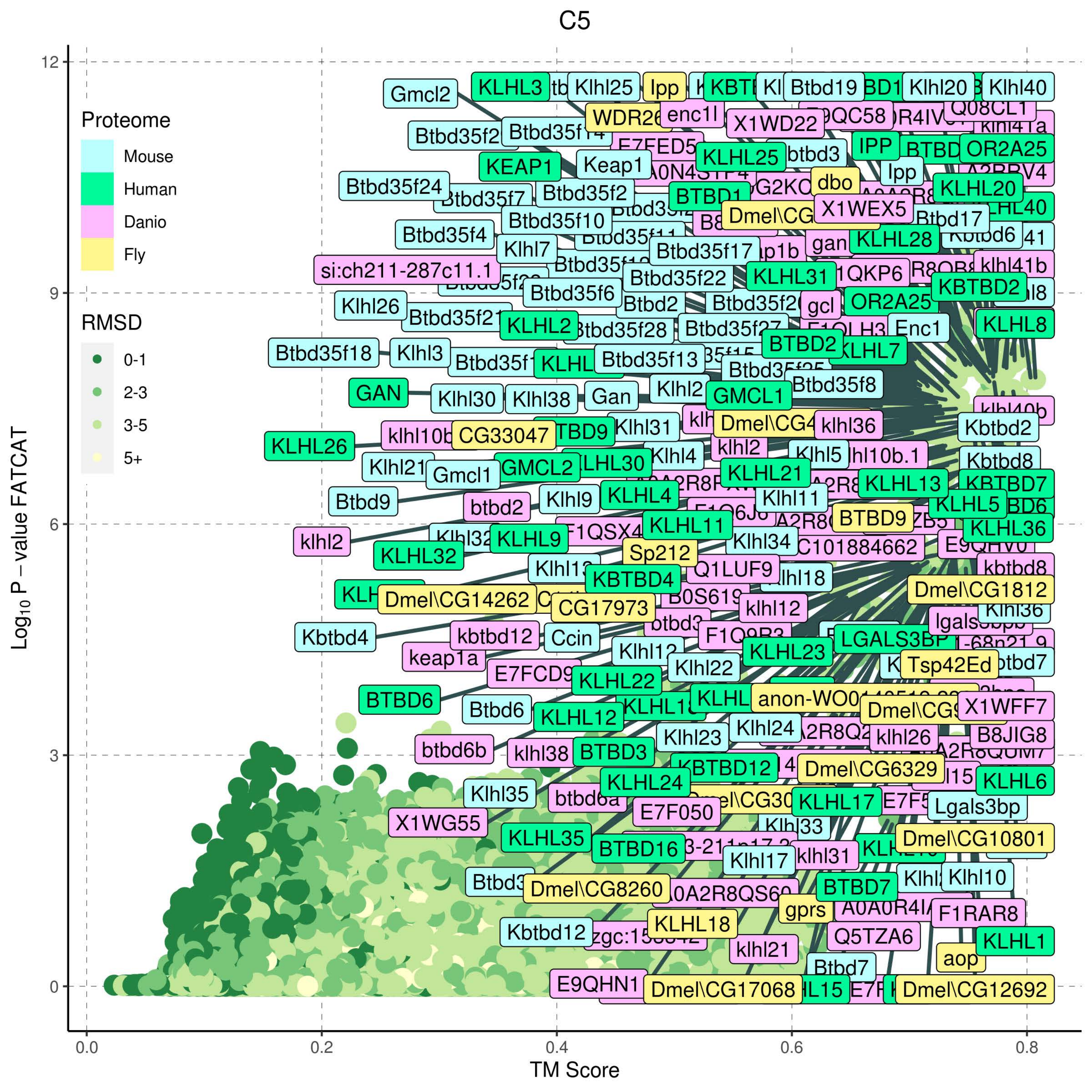
C5
Proteome
Mouse
Human
Danio
Fly
RMSD
0-1
2-3
3-5
5+
Log10 P – value FATCAT
TM Score
0.0
0.2
0.4
0.6
0.8
0
3
6
9
12
Gmcl2
KLHL3
Klhl25
Ipp
Btbd19
Klhl20
Klhl40
WDR26
enc1l
X1WD22
Q08CL1
Btbd35f2
Btbd35f14
KEAP1
Keap1
KLHL25
btbd3
IPP
OR2A25
Btbd35f24
Btbd35f7
Btbd35f2
BTBD1
dbo
KLHL20
X1WEX5
Btbd35f10
Btbd17
Btbd35f4
Klhl7
Btbd35f17
gan
KLHL28
si:ch211-287c11.1
Btbd35f22
KLHL31
klhl41b
KBTBD2
Btbd35f6
Klhl26
Btbd2
gcl
OR2A25
Btbd35f21
KLHL2
Btbd35f28
Enc1
KLHL8
Btbd35f18
Klhl3
BTBD2
KLHL7
Btbd35f13
Btbd35f8
GAN
Klhl30
Klhl38
Gan
Klhl2
GMCL1
klhl10b
CG33047
Klhl31
klhl36
Kbtbd2
KLHL26
klhl2
Klhl4
Klhl5
Klhl21
Gmcl1
GMCL2
KLHL30
KLHL21
KLHL13
Kbtbd8
KBTBD7
Btbd9
btbd2
Klhl9
KLHL4
Klhl11
KLHL5
BTBD9
KLHL36
klhl2
KLHL9
KLHL11
Klhl34
KLHL32
Sp212
Q1LUF9
kbtbd8
KBTBD4
Dmel\CG1812
Dmel\CG14262
CG17973
klhl12
Klhl36
Kbtbd4
kbtbd12
Ccin
LGALS3BP
keap1a
KLHL23
Klhl22
Tsp42Ed
KLHL22
BTBD6
Btbd6
KLHL12
X1WFF7
Klhl23
Klhl24
klhl26
B8JIG8
btbd6b
klhl38
BTBD3
KBTBD12
Dmel\CG6329
KLHL24
KLHL6
Klhl35
btbd6a
E7F050
KLHL17
Lgals3bp
X1WG55
Klhl33
KLHL35
BTBD16
Dmel\CG10801
Klhl17
klhl31
Dmel\CG8260
BTBD7
Klhl10
gprs
F1RAR8
KLHL18
Kbtbd12
Q5TZA6
klhl21
KLHL1
aop
Btbd7
E9QHN1
Dmel\CG17068
Dmel\CG12692

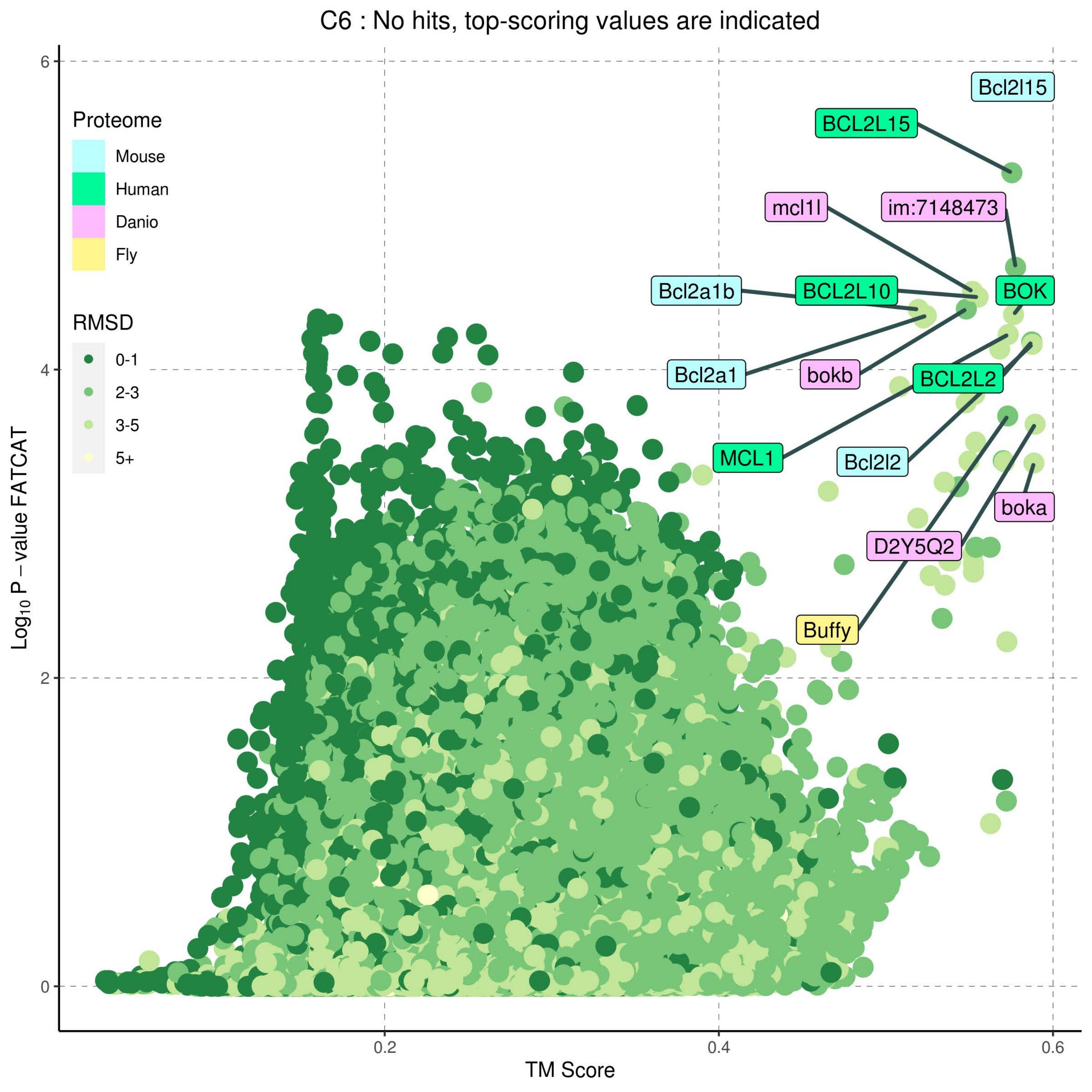

C6 : No hits, top-scoring values are indicated
Proteome
Mouse
Human
Danio
Fly
RMSD
0-1
2-3
3-5
5+
Log10 P − value FATCAT
TM Score
0
2
4
6
0.2
0.4
0.6
Bcl2l15
BCL2L15
mcl1l
im:7148473
Bcl2a1b
BCL2L10
BOK
Bcl2a1
bokb
BCL2L2
MCL1
Bcl2l2
boka
D2Y5Q2
Buffy

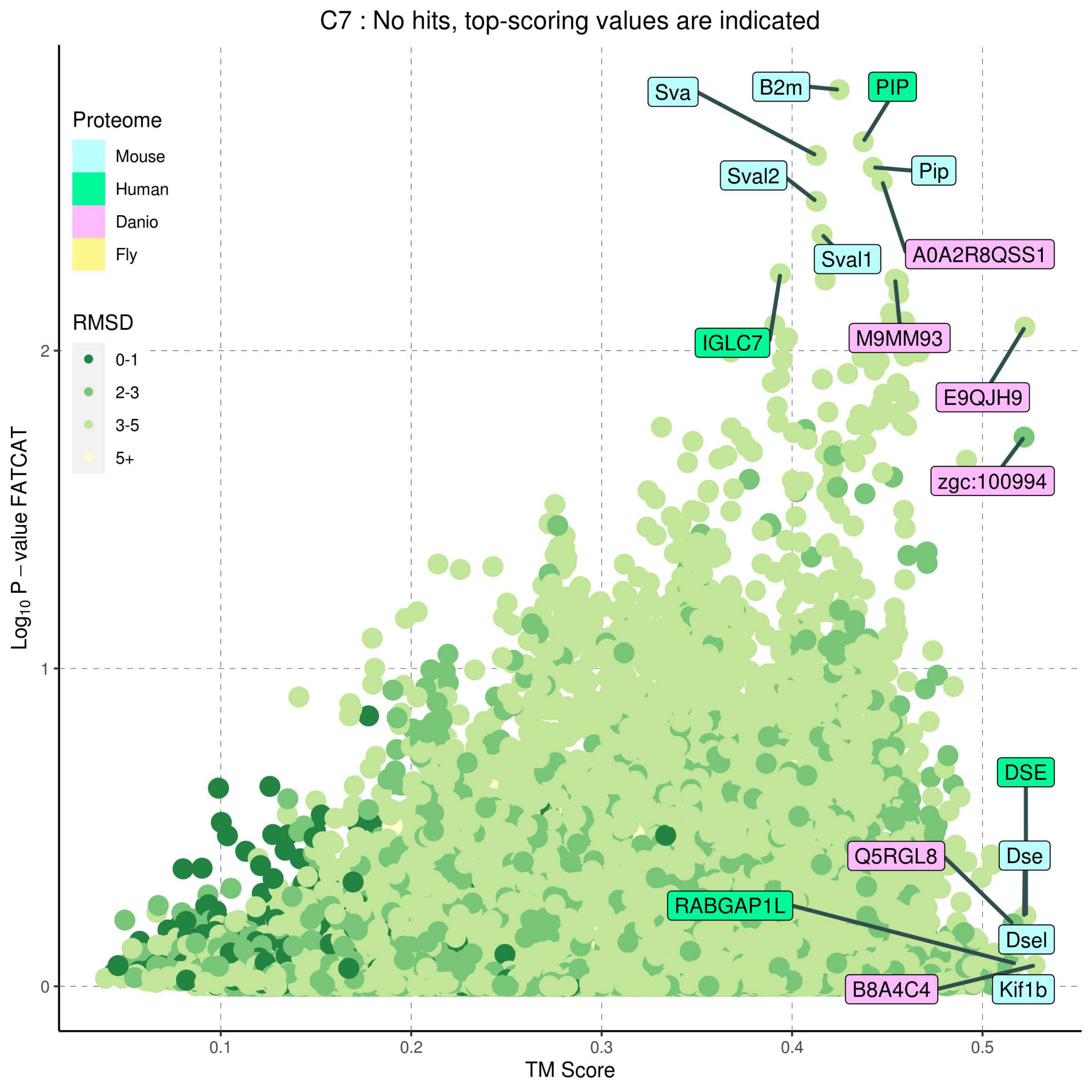
C7 : No hits, top-scoring values are indicated
Proteome
Mouse
Human
Danio
Fly
RMSD
0-1
2-3
3-5
5+
Log10 P – value FATCAT
TM Score
0
1
2
0.1
0.2
0.3
0.4
0.5
Sva
B2m
PIP
Pip
Sval2
Sval1
A0A2R8QSS1
IGLC7
M9MM93
E9QJH9
zgc:100994
DSE
Dse
Q5RGL8
RABGAP1L
Dsel
B8A4C4
Kif1b

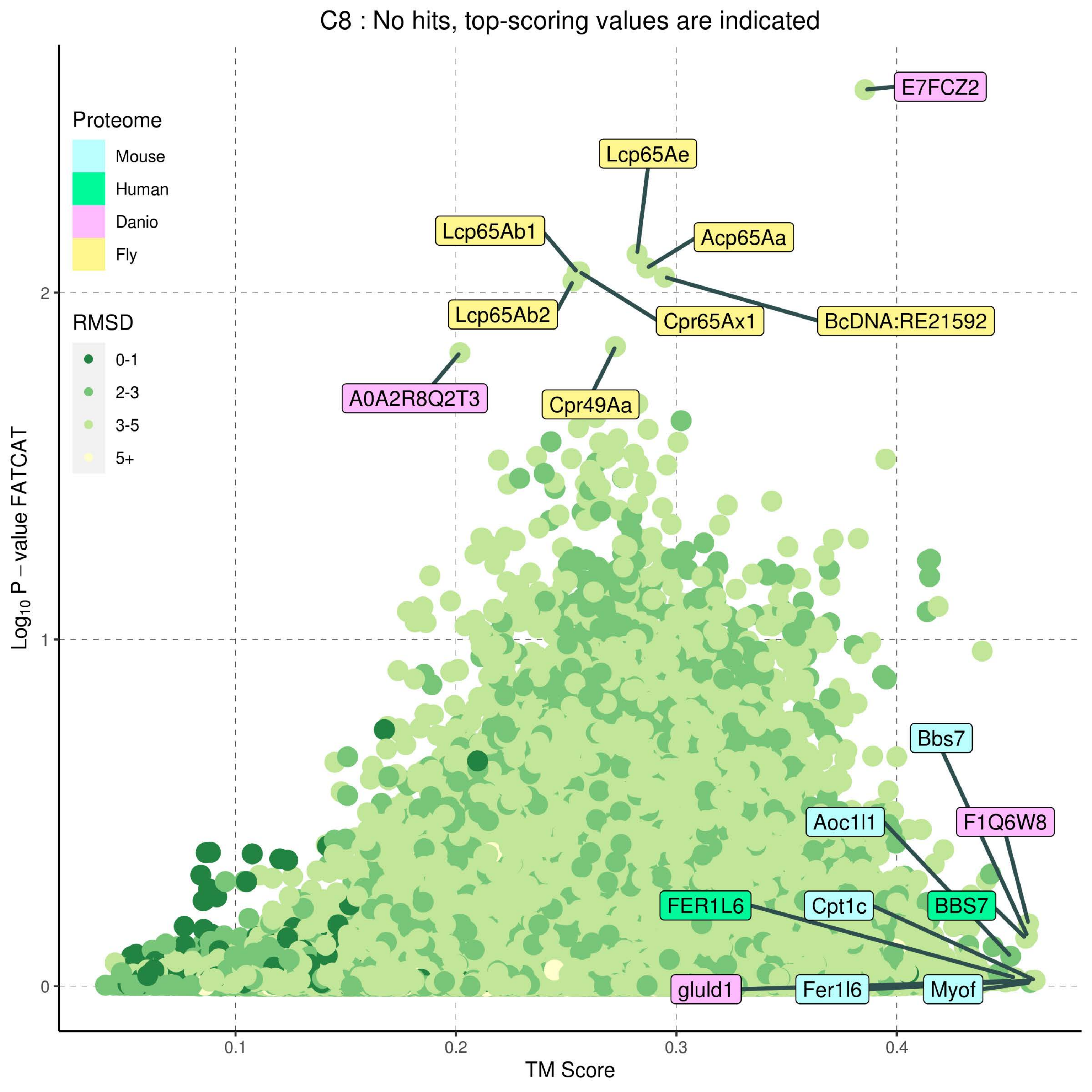

C8 : No hits, top-scoring values are indicated
Proteome
Mouse
Human
Danio
Fly
RMSD
0-1
2-3
3-5
5+
E7FCZ2
Lcp65Ae
Acp65Aa
Lcp65Ab1
Lcp65Ab2
Cpr65Ax1
BcDNA:RE21592
A0A2R8Q2T3
Cpr49Aa
Bbs7
F1Q6W8
Aoc1l1
FER1L6
Cpt1c
BBS7
gluld1
Fer1l6
Myof
Log10 P − value FATCAT
TM Score
0
1
2
0.1
0.2
0.3
0.4

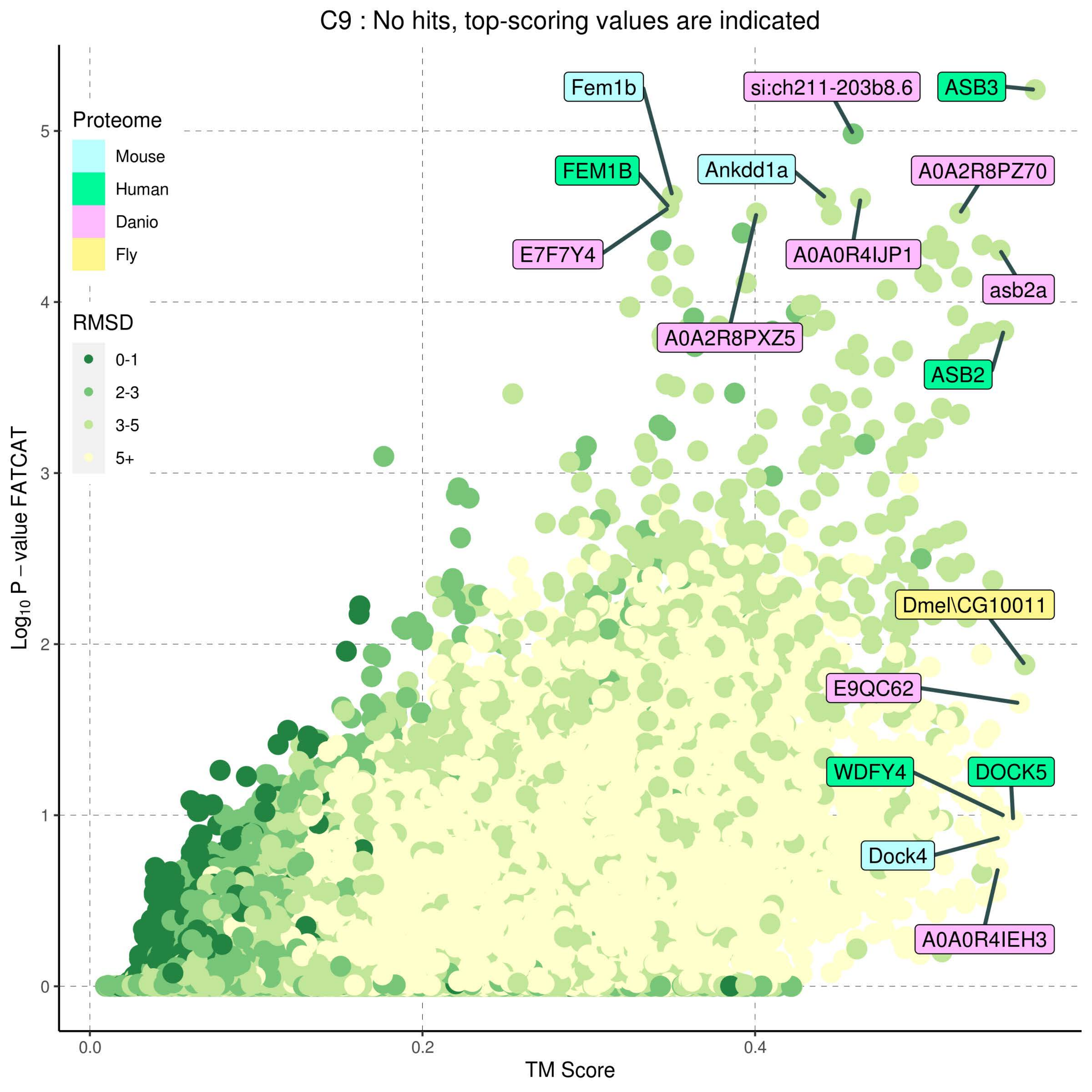
C9 : No hits, top-scoring values are indicated
Proteome
Mouse
Human
Danio
Fly
RMSD
0-1
2-3
3-5
5+
Fem1b
si:ch211-203b8.6
ASB3
FEM1B
Ankdd1a
A0A2R8PZ70
E7F7Y4
A0A0R4IJP1
asb2a
A0A2R8PXZ5
ASB2
Dmel\CG10011
E9QC62
WDFY4
DOCK5
Dock4
A0A0R4IEH3
Log10 P − value FATCAT
TM Score
0.0
0.2
0.4
0
1
2
3
4
5

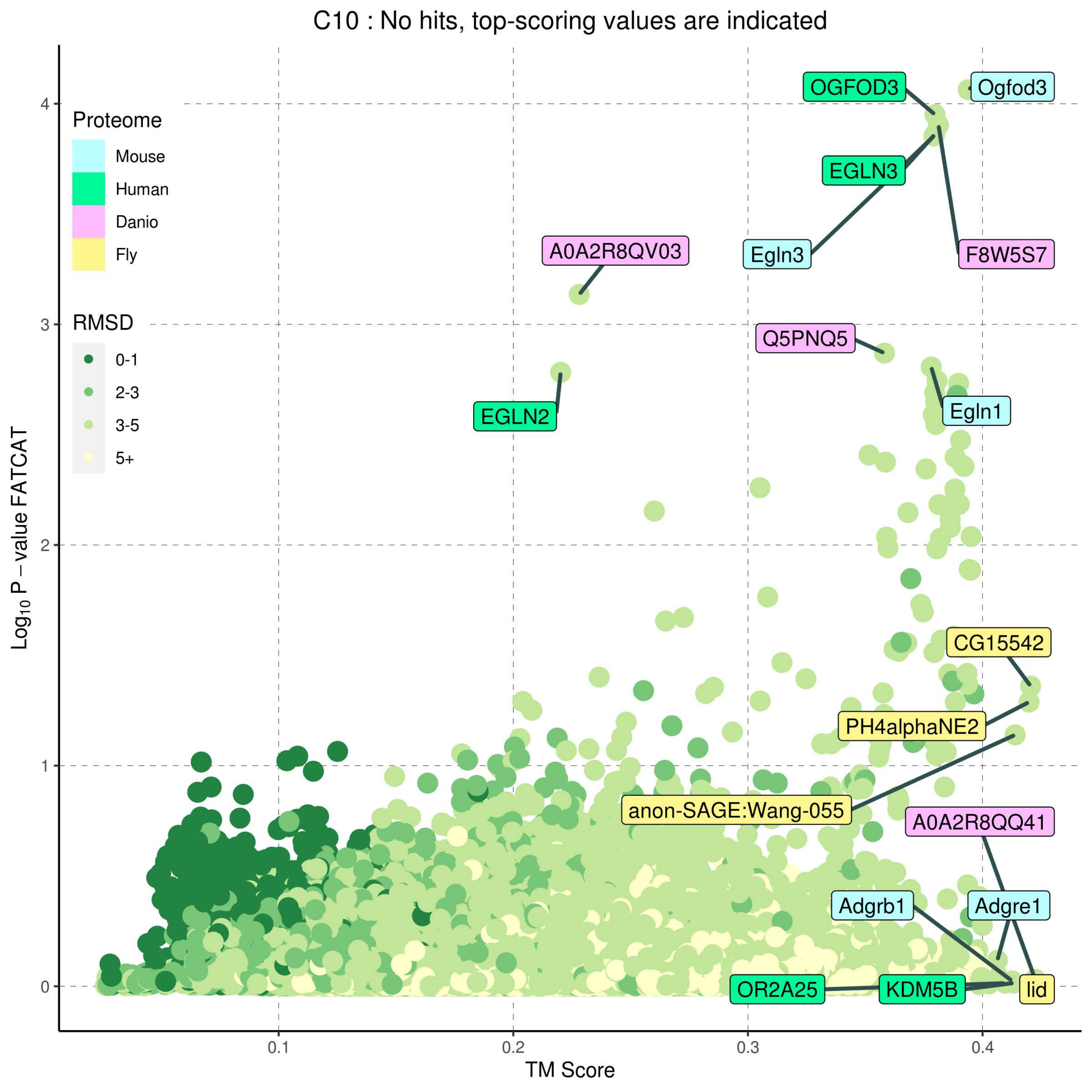
C10 : No hits, top-scoring values are indicated
Proteome
Mouse
Human
Danio
Fly
RMSD
0-1
2-3
3-5
5+
$Log_{10}$ P − value FATCAT
TM Score
0
1
2
3
4
0.1
0.2
0.3
0.4
OGFOD3
Ogfod3
EGLN3
Egln3
F8W5S7
A0A2R8QV03
Q5PNQ5
EGLN2
Egln1
CG15542
PH4alphaNE2
anon-SAGE:Wang-055
A0A2R8QQ41
Adgrb1
Adgre1
OR2A25
KDM5B
lid

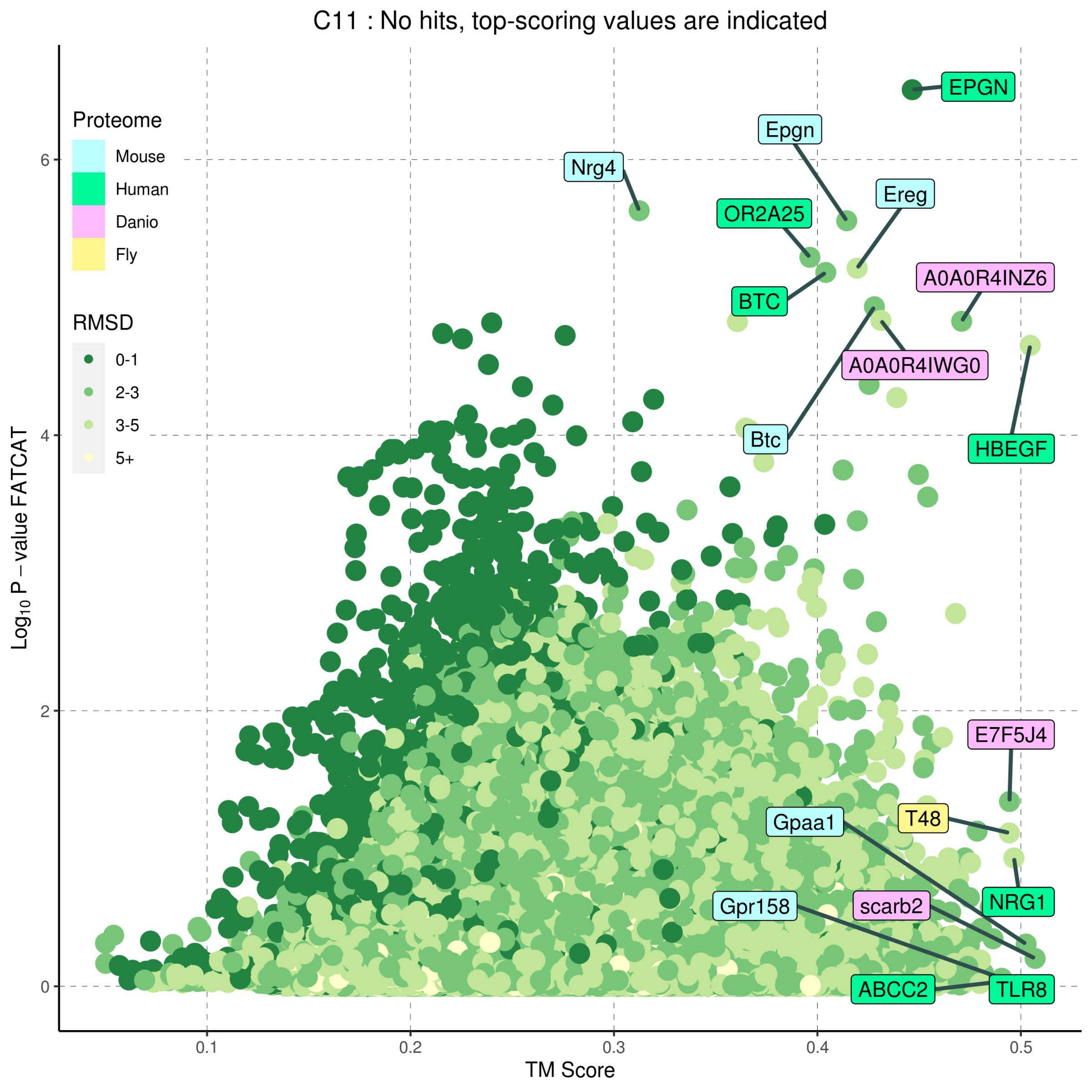
C11 : No hits, top-scoring values are indicated
Proteome
Mouse
Human
Danio
Fly
RMSD
0-1
2-3
3-5
5+
EPGN
Epgn
Nrg4
Ereg
OR2A25
BTC
A0A0R4INZ6
A0A0R4IWG0
Btc
HBEGF
E7F5J4
T48
Gpaa1
NRG1
scarb2
Gpr158
ABCC2
TLR8
$Log_{10}$ P − value FATCAT
TM Score
0
2
4
6
0.1
0.2
0.3
0.4
0.5

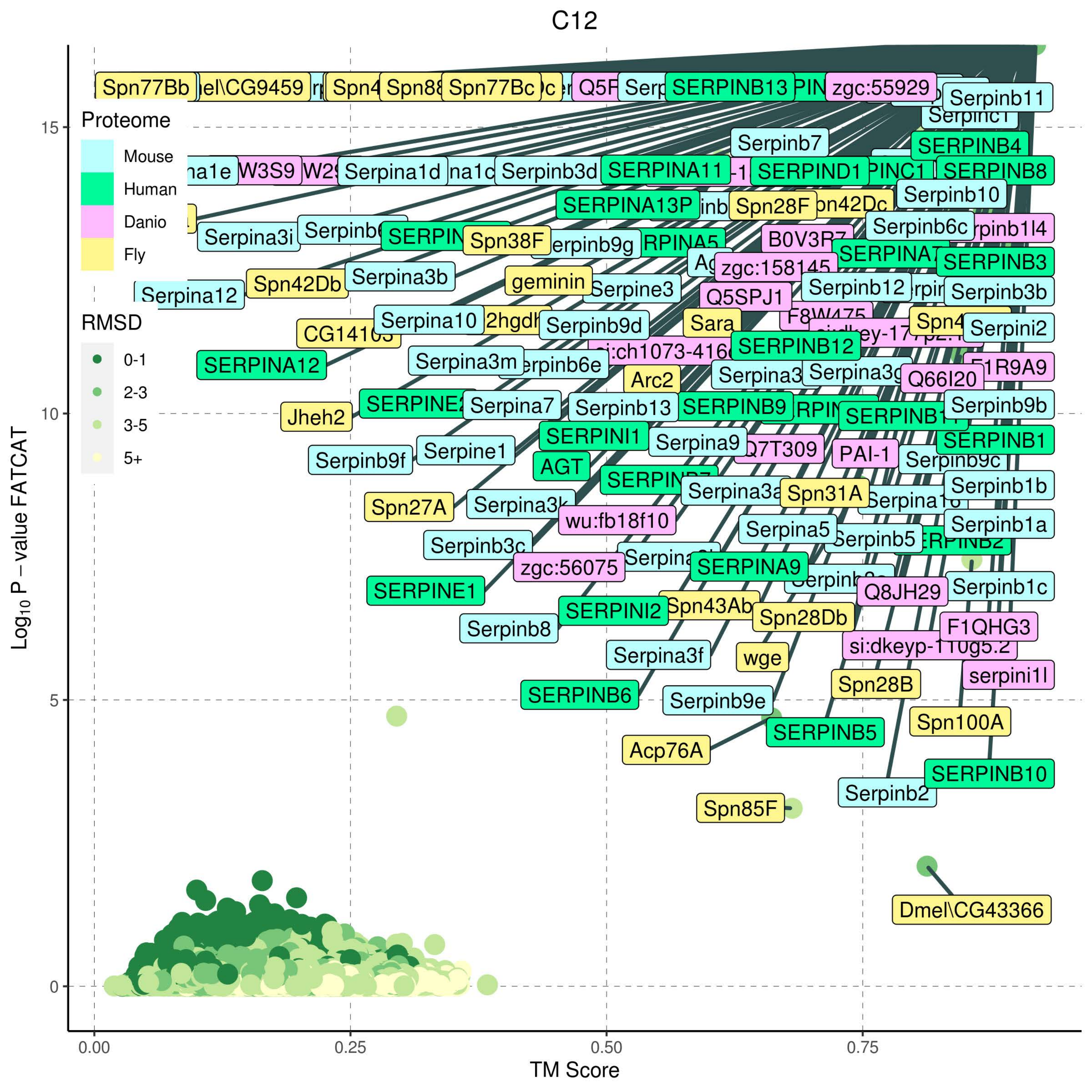

C12
Log10 P – value FATCAT
TM Score
0.00
0.25
0.50
0.75
0
5
10
15
Proteome
Mouse
Human
Danio
Fly
RMSD
0-1
2-3
3-5
5+
Spn77Bb
Spn77Bc
SERPINB13
zgc:55929
Serpinb11
Serpinb7
SERPINB4
Serpina1d
Serpinb3d
SERPINA11
SERPIND1
SERPINB8
Serpinb10
SERPINA13P
Spn28F
Serpina3i
Spn38F
Serpinb6c
SERPINB3
Spn42Db
Serpina3b
geminin
Serpine3
Serpinb12
Serpinb3b
Serpina12
Q5SPJ1
Serpina10
Serpinb9d
Sara
Serpini2
SERPINA12
SERPINB12
Serpina3m
Arc2
Q66I20
Jheh2
Serpina7
Serpinb13
SERPINB9
Serpinb9b
SERPINI1
Serpina9
SERPINB1
Serpine1
Serpinb9f
AGT
PAI-1
Serpinb1b
Spn27A
Spn31A
wu:fb18f10
Serpina5
Serpinb1a
Serpinb5
Serpinb3c
zgc:56075
SERPINA9
SERPINE1
Q8JH29
Serpinb1c
SERPINI2
Spn28Db
F1QHG3
Serpinb8
Serpina3f
wge
serpini1l
SERPINB6
Spn28B
Serpinb9e
Spn100A
SERPINB5
Acp76A
SERPINB10
Serpinb2
Spn85F
Dmel\CG43366

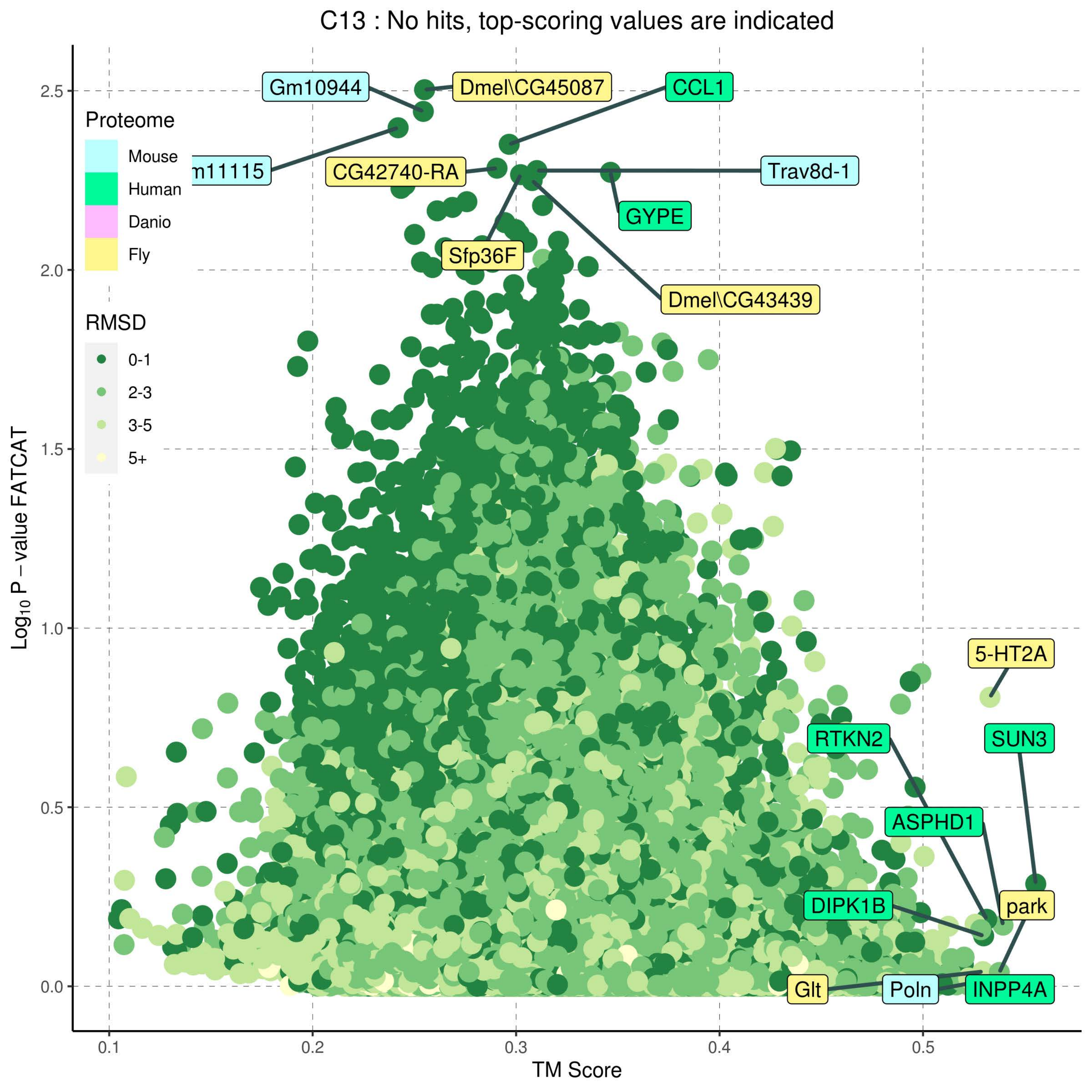
C13 : No hits, top-scoring values are indicated
Proteome
Mouse
Human
Danio
Fly
RMSD
0-1
2-3
3-5
5+
Gm10944
Dmel\CG45087
CCL1
m11115
CG42740-RA
Trav8d-1
GYPE
Sfp36F
Dmel\CG43439
5-HT2A
RTKN2
SUN3
ASPHD1
DIPK1B
park
Glt
Poln
INPP4A
Log10 P – value FATCAT
TM Score
0.0
0.5
1.0
1.5
2.0
2.5
0.1
0.2
0.3
0.4
0.5

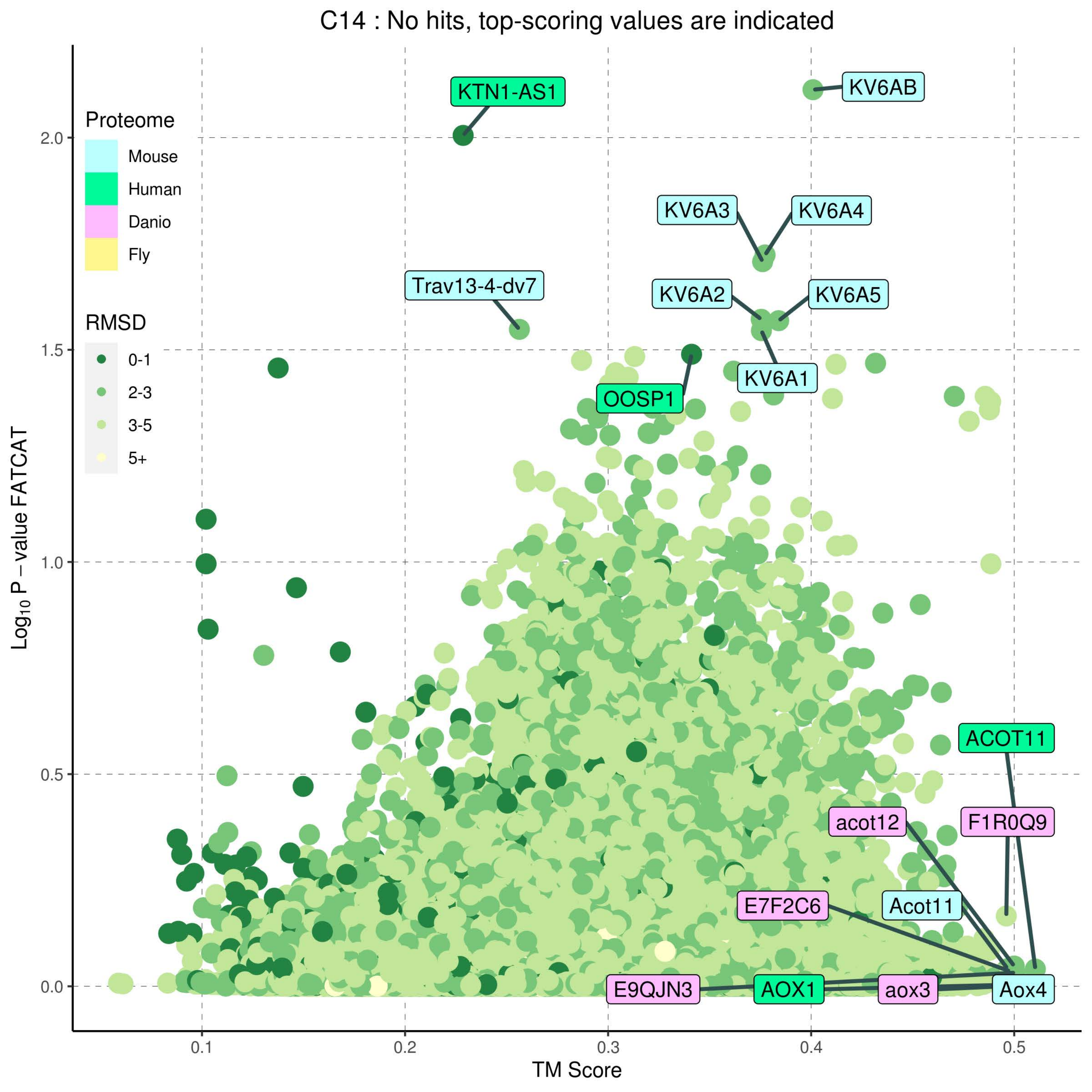
C14 : No hits, top-scoring values are indicated
Proteome
Mouse
Human
Danio
Fly
RMSD
0-1
2-3
3-5
5+
KTN1-AS1
KV6AB
KV6A3
KV6A4
Trav13-4-dv7
KV6A2
KV6A5
KV6A1
OOSP1
ACOT11
acot12
F1R0Q9
E7F2C6
Acot11
E9QJN3
AOX1
aox3
Aox4
Log10 P − value FATCAT
TM Score
0.0
0.5
1.0
1.5
2.0
0.1
0.2
0.3
0.4
0.5

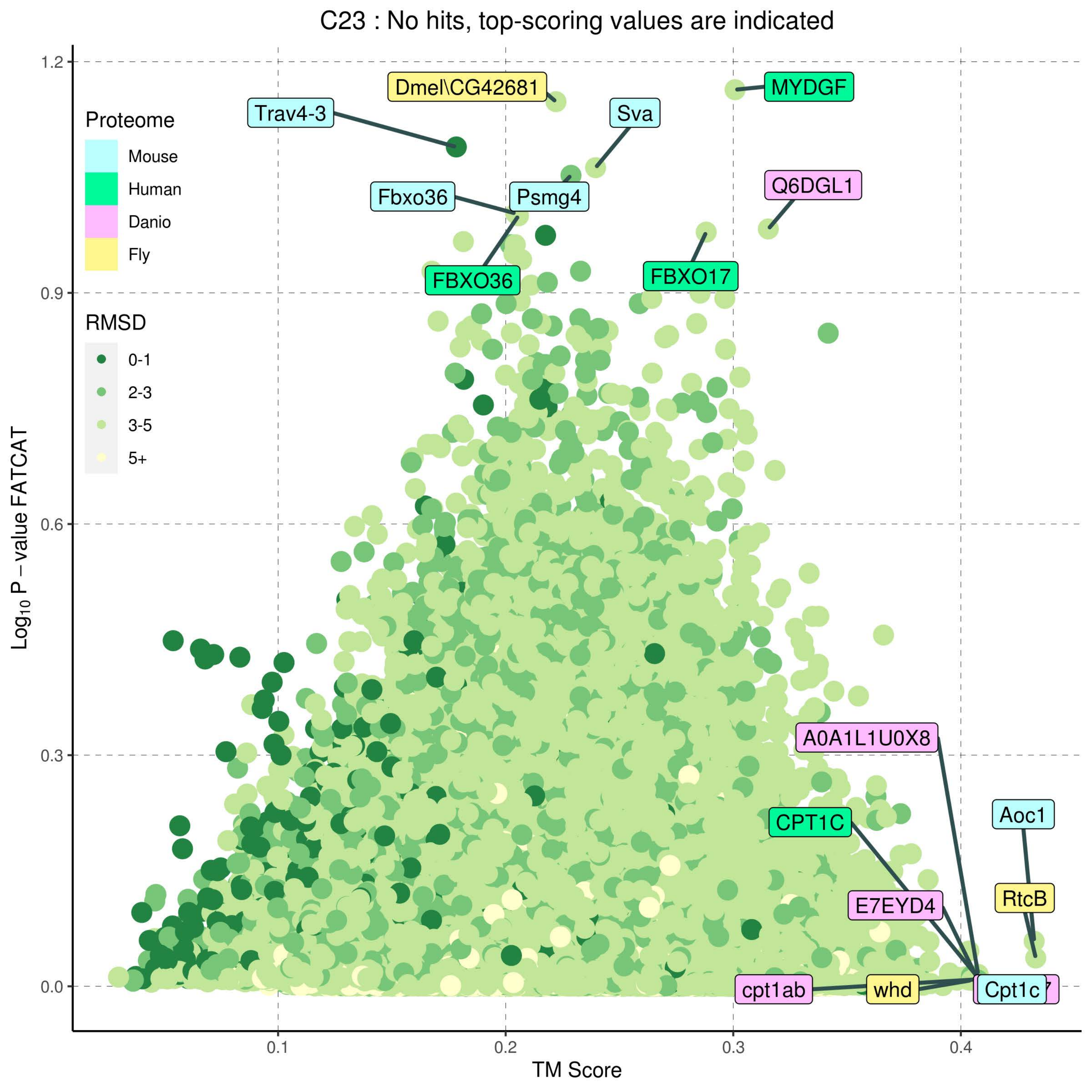

C23 : No hits, top-scoring values are indicated
Proteome
Mouse
Human
Danio
Fly
RMSD
0-1
2-3
3-5
5+
Log10 P – value FATCAT
TM Score
0.0
0.3
0.6
0.9
1.2
0.1
0.2
0.3
0.4
Dmel\CG42681
MYDGF
Trav4-3
Sva
Fbxo36
Psmg4
Q6DGL1
FBXO36
FBXO17
A0A1L1U0X8
Aoc1
CPT1C
RtcB
E7EYD4
cpt1ab
whd
Cpt1c

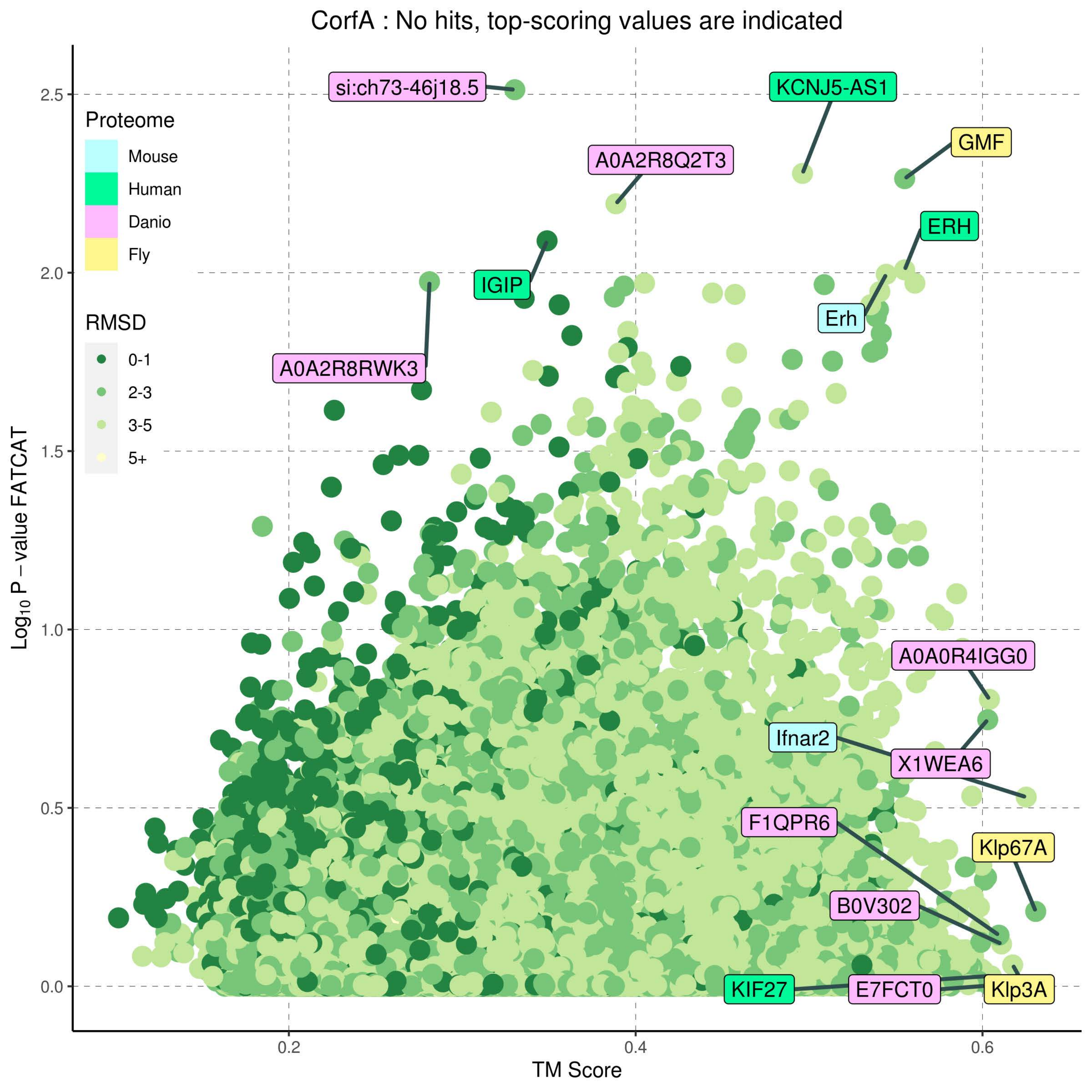

CorfA : No hits, top-scoring values are indicated
Proteome
Mouse
Human
Danio
Fly
RMSD
0-1
2-3
3-5
5+
si:ch73-46j18.5
KCNJ5-AS1
GMF
A0A2R8Q2T3
ERH
IGIP
Erh
A0A2R8RWK3
A0A0R4IGG0
Ifnar2
X1WEA6
F1QPR6
Klp67A
B0V302
KIF27
E7FCT0
Klp3A
Log10 P – value FATCAT
TM Score
2.5
2.0
1.5
1.0
0.5
0.0
0.2
0.4
0.6

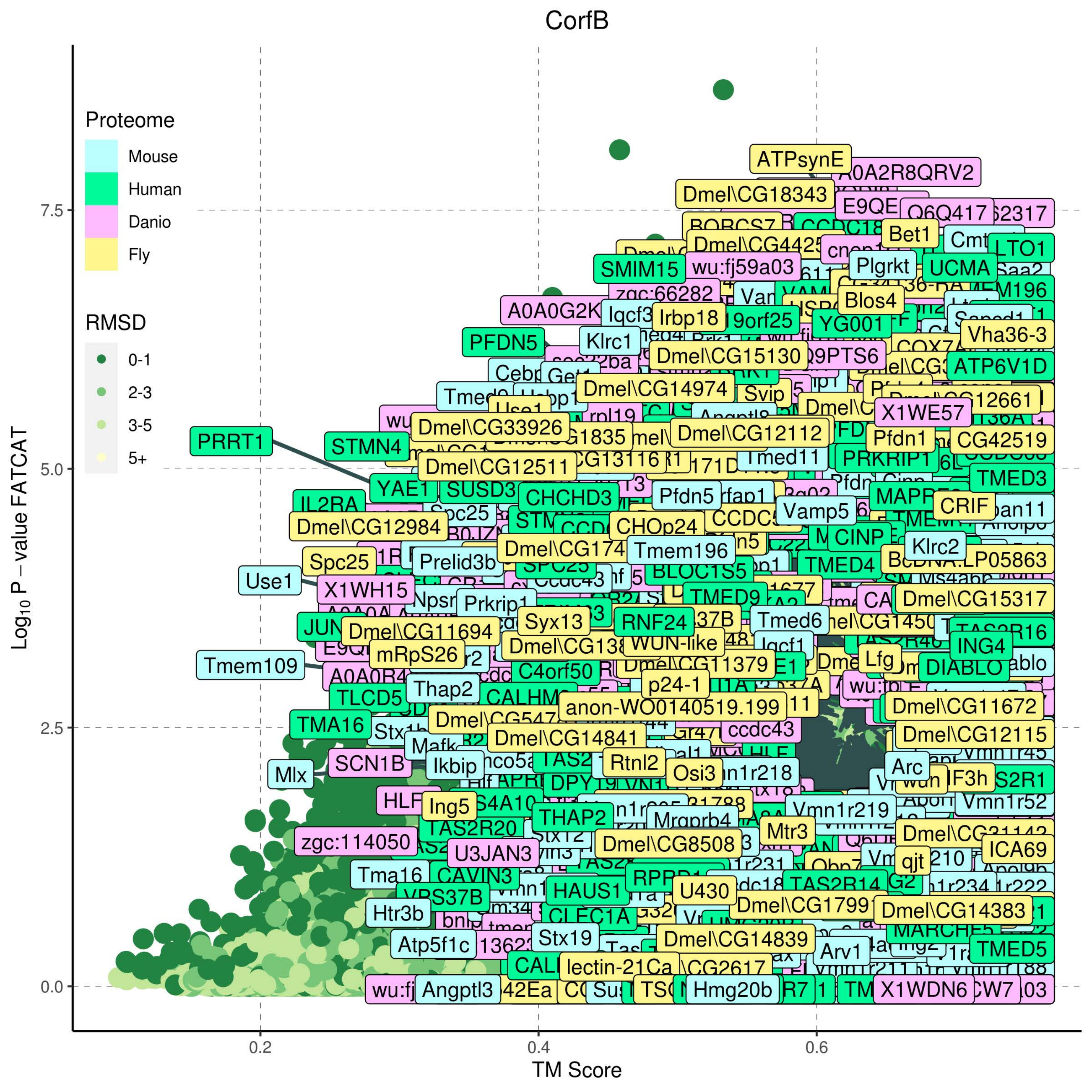
CorfB
Proteome
Mouse
Human
Danio
Fly
RMSD
0-1
2-3
3-5
5+
Log10 P − value FATCAT
TM Score
0.0
2.5
5.0
7.5
0.2
0.4
0.6
ATPsynE
A0A2R8QRV2
Dmel\CG18343
SMIM15
wu:fj59a03
zgc:66282
A0A0G2K
Iqcf3
Irbp18
PFDN5
Klrc1
Dmel\CG15130
Q9PTS6
Vha36-3
ATP6V1D
Dmel\CG14974
Svip
X1WE57
PRRT1
STMN4
Dmel\CG33926
Dmel\CG12112
Pfdn1
CG42519
Dmel\CG12511
Tmed11
PRKRIP1
TMED3
YAE1
CHCHD3
Pfdn5
Vamp5
CRIF
Dmel\CG12984
CHOp24
CINP
Klrc2
Spc25
Prelid3b
Tmem196
TMED4
Use1
X1WH15
BLOC1S5
Dmel\CG15317
Prkrip1
Syx13
RNF24
Tmed6
Dmel\CG11694
ING4
mRpS26
Tmem109
Lfg
DIABLO
C4orf50
p24-1
Thap2
anon-WO0140519.199
Dmel\CG11672
TMA16
Dmel\CG14841
ccdc43
Dmel\CG12115
SCN1B
Ikbip
Rtnl2
Osi3
Arc
Mlx
Ing5
Vmn1r219
Vmn1r52
THAP2
Mrgprb4
TAS2R20
Mtr3
zgc:114050
Dmel\CG8508
ICA69
U3JAN3
qjt
Tma16
CAVIN3
HAUS1
U430
TAS2R14
Htr3b
Dmel\CG14383
CLEC1A
Stx19
Dmel\CG14839
Atp5f1c
Arv1
TMED5
lectin-21Ca
Angptl3
Hmg20b
X1WDN6

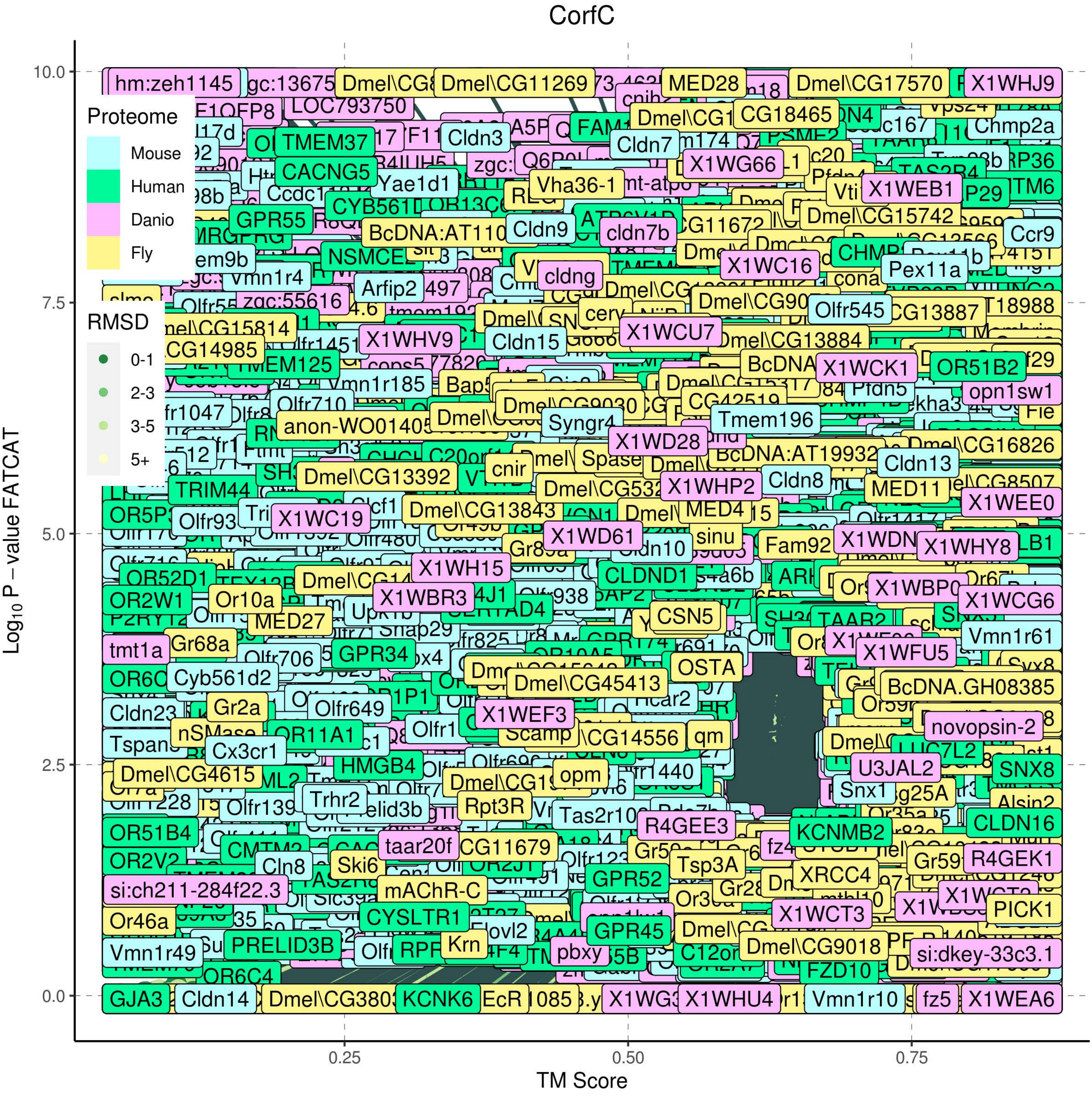
CorfC
Proteome
Mouse
Human
Danio
Fly
RMSD
0-1
2-3
3-5
5+
Log10 P – value FATCAT
TM Score
0.0
2.5
5.0
7.5
10.0
0.25
0.50
0.75

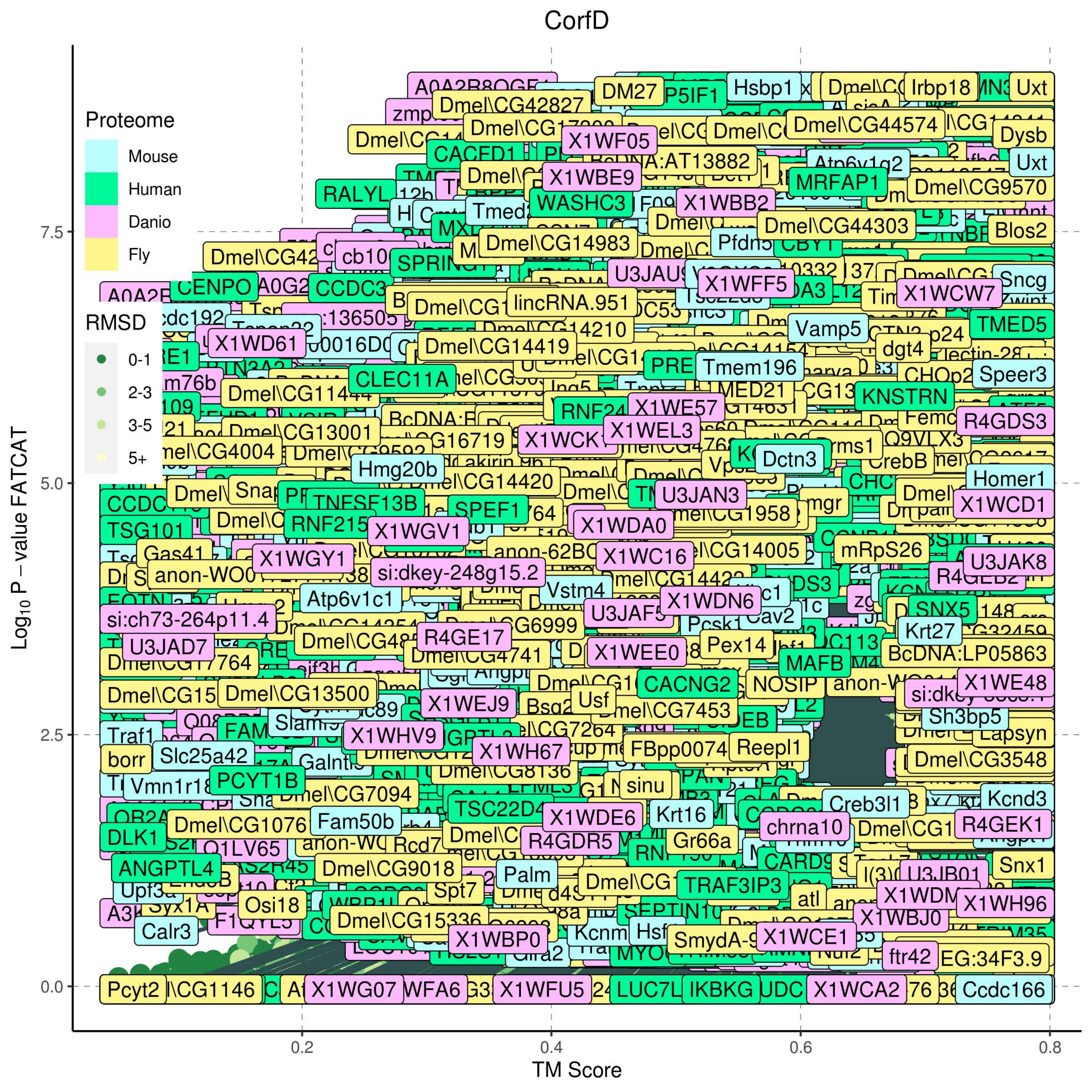
CorfD
Proteome
Mouse
Human
Danio
Fly
RMSD
0-1
2-3
3-5
5+
Log10 P – value FATCAT
TM Score
0.0
2.5
5.0
7.5
0.2
0.4
0.6
0.8
DM27
Hsbp1
Irbp18
Uxt
Dmel\CG42827
Dmel\CG44574
Dysb
X1WF05
CACFD1
X1WBE9
MRFAP1
Dmel\CG9570
RALYL
WASHC3
X1WBB2
Tmed2
Dmel\CG44303
Blos2
Dmel\CG14983
CENPO
X1WFF5
X1WCW7
Sncg
lincRNA.951
X1WD61
Dmel\CG14419
Vamp5
TMED5
dgt4
Tmem196
CHOp2
Speer3
CLEC11A
Dmel\CG11444
KNSTRN
X1WE57
R4GDS3
Dmel\CG13001
X1WEL3
Dctn3
CrebB
Hmg20b
Dmel\CG14420
Homer1
TNFSF13B
U3JAN3
X1WCD1
SPEF1
X1WDA0
TSG101
X1WGV1
RNF215
mRpS26
Gas41
X1WGY1
X1WC16
U3JAK8
si:dkey-248g15.2
Vstm4
Atp6v1c1
X1WDN6
si:ch73-264p11.4
SNX5
Krt27
R4GE17
U3JAD7
X1WEE0
Pex14
MAFB
Dmel\CG4741
BcDNA:LP05863
CACNG2
NOSIP
X1WE48
Dmel\CG13500
X1WEJ9
Usf
Dmel\CG7453
Sh3bp5
X1WHV9
X1WH67
Reepl1
Dmel\CG3548
borr
Slc25a42
PCYT1B
Dmel\CG8136
sinu
Dmel\CG7094
TSC22D4
Creb3l1
Kcnd3
Fam50b
X1WDE6
Krt16
chrna10
R4GEK1
DLK1
R4GDR5
Gr66a
Q1LV65
ANGPTL4
Dmel\CG9018
Snx1
Palm
TRAF3IP3
Spt7
X1WH96
Osi18
X1WBJ0
Calr3
Dmel\CG15336
X1WCE1
X1WBP0
ftr42
EG:34F3.9
Pcyt2
X1WG07
X1WFU5
IKBKG
X1WCA2
Ccdc166

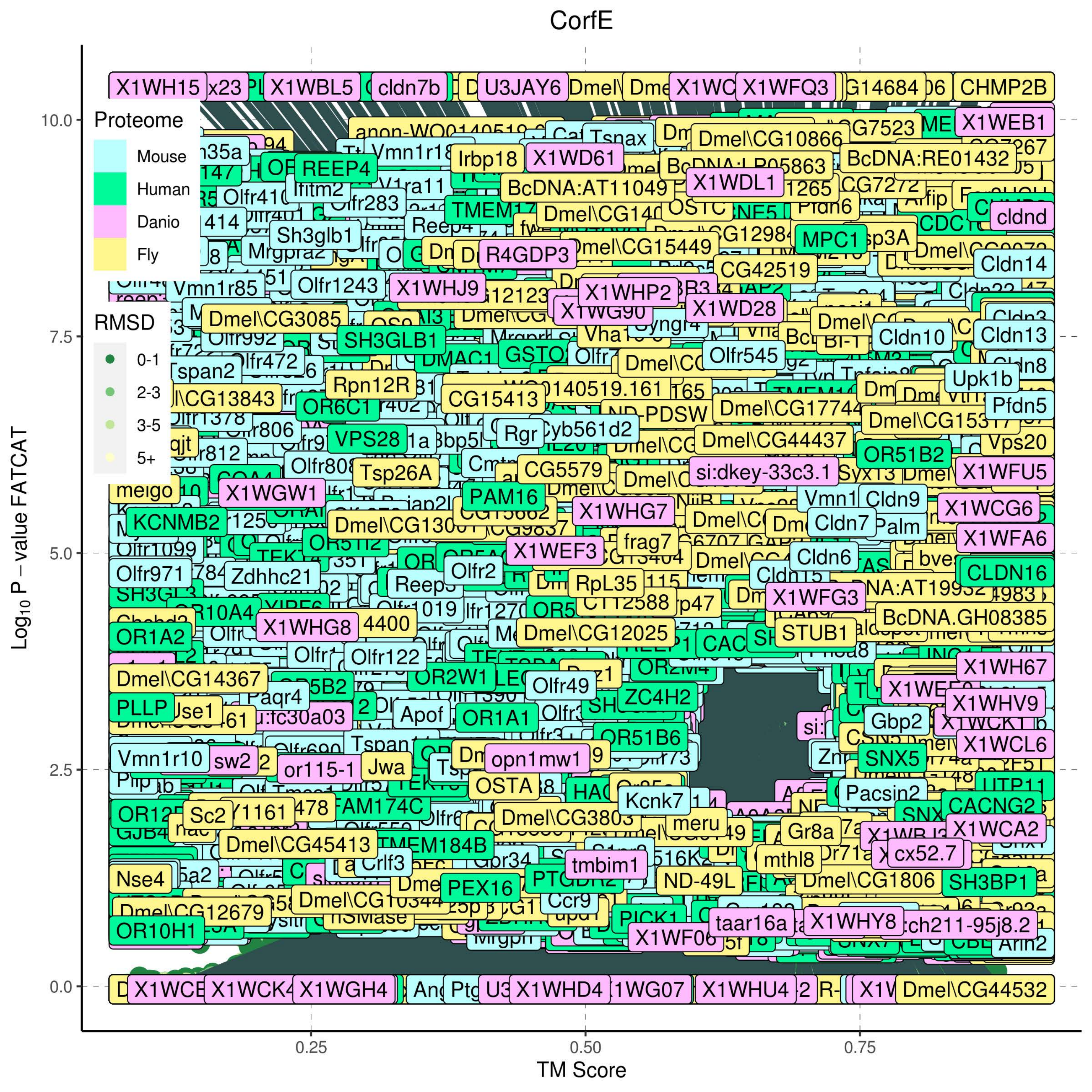
CorfE
Proteome
Mouse
Human
Danio
Fly
RMSD
0-1
2-3
3-5
5+
$\text{Log}_{10}$ P – value FATCAT
TM Score
0.0
2.5
5.0
7.5
10.0
0.25
0.50
0.75

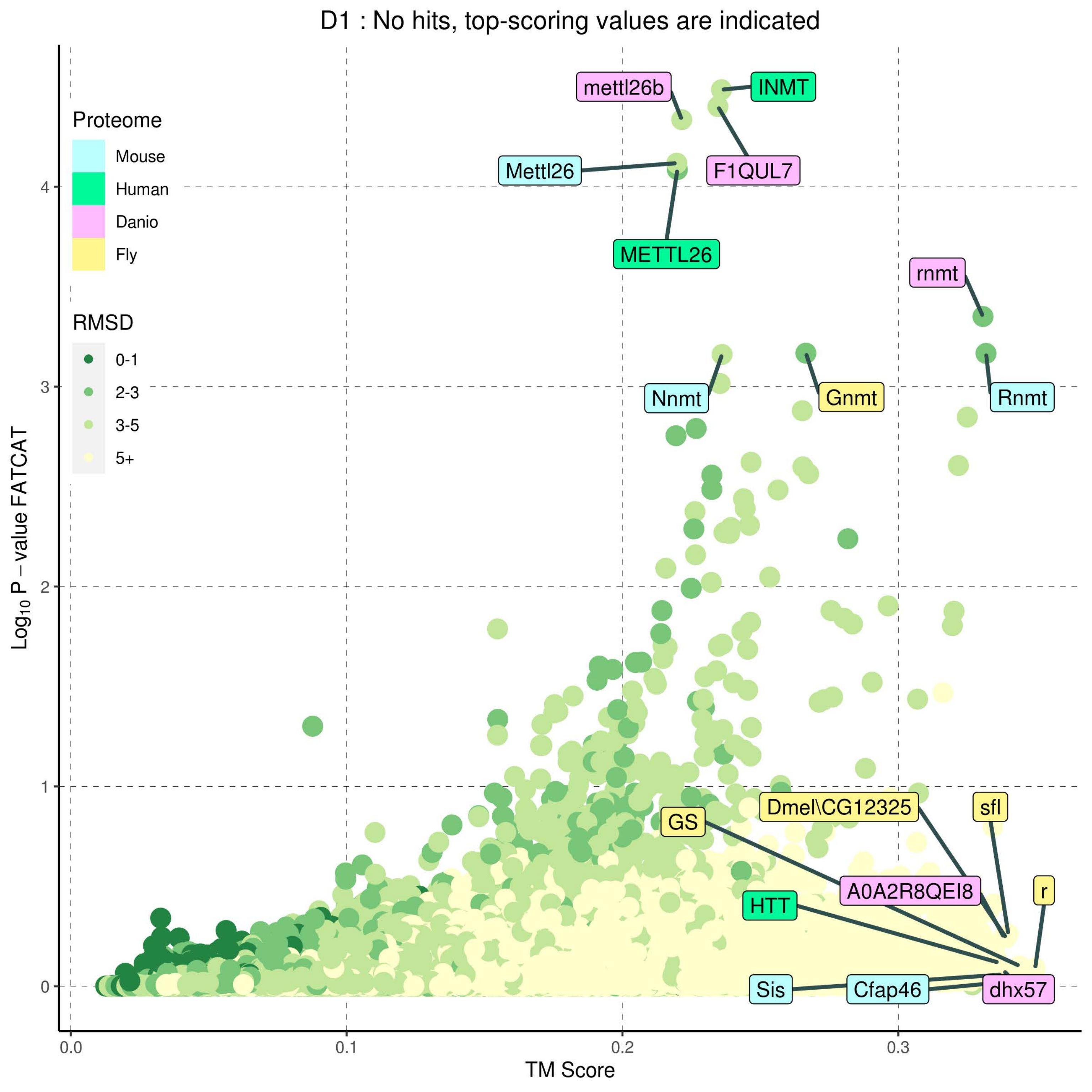
D1 : No hits, top-scoring values are indicated
Proteome
Mouse
Human
Danio
Fly
RMSD
0-1
2-3
3-5
5+
Log10 P − value FATCAT
TM Score
mettl26b
INMT
Mettl26
F1QUL7
METTL26
rnmt
Nnmt
Gnmt
Rnmt
GS
Dmel\CG12325
sfl
A0A2R8QEI8
r
HTT
Sis
Cfap46
dhx57

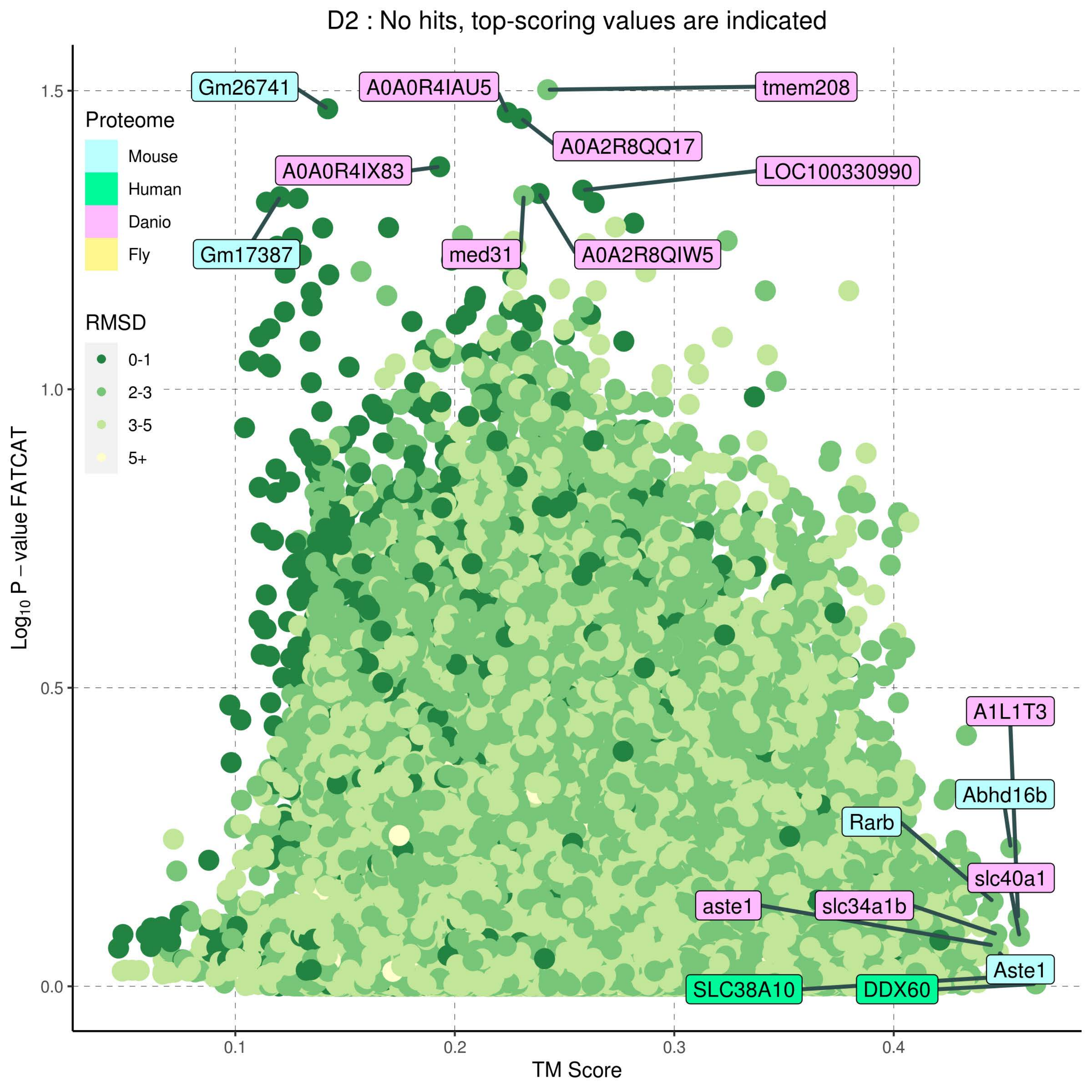
D2 : No hits, top-scoring values are indicated
Proteome
Mouse
Human
Danio
Fly
RMSD
0-1
2-3
3-5
5+
Gm26741
A0A0R4IAU5
tmem208
A0A2R8QQ17
A0A0R4IX83
LOC100330990
Gm17387
med31
A0A2R8QIW5
A1L1T3
Abhd16b
Rarb
slc40a1
aste1
slc34a1b
Aste1
SLC38A10
DDX60
1.5
1.0
0.5
0.0
0.1
0.2
0.3
0.4
TM Score
Log10 P – value FATCAT

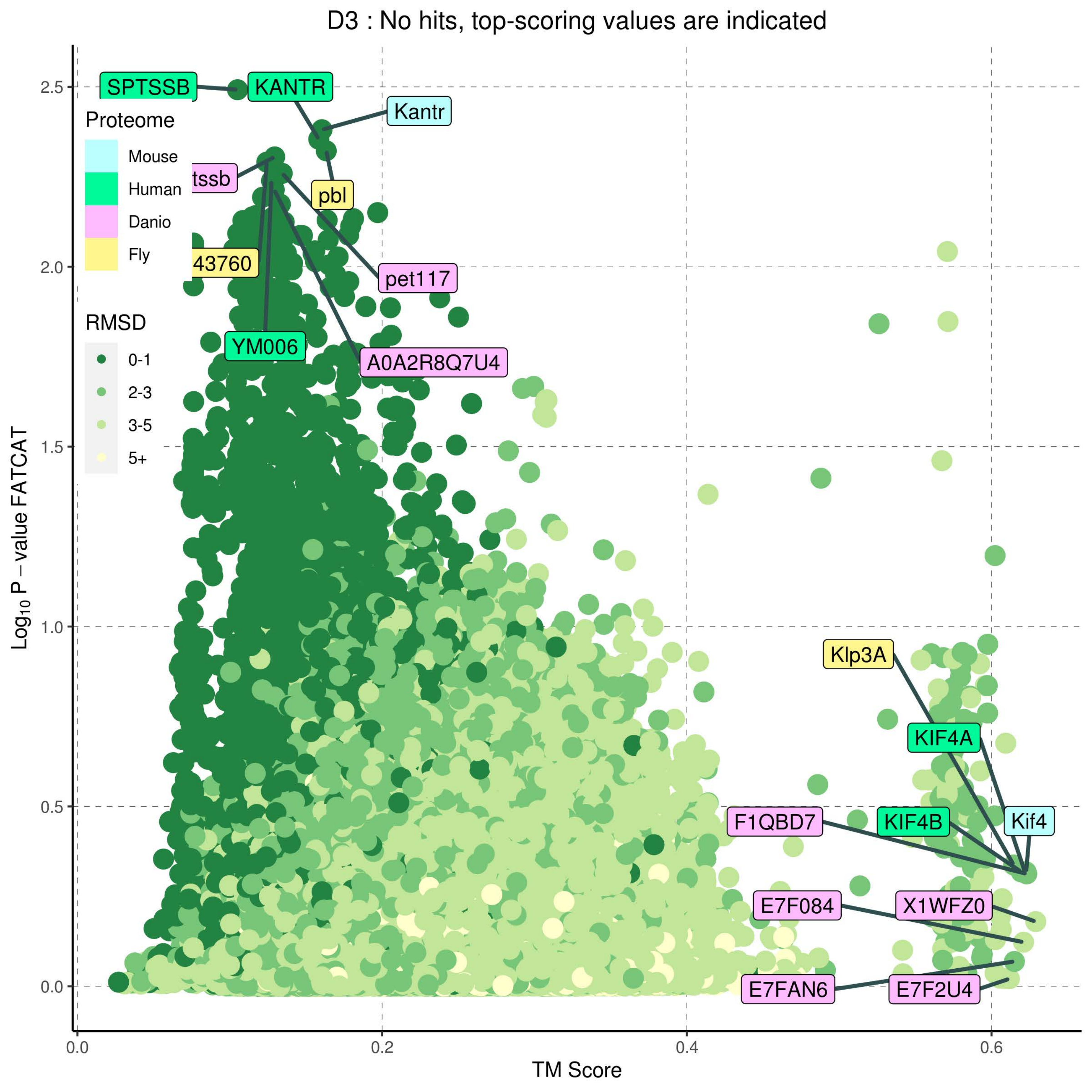

D3 : No hits, top-scoring values are indicated
Proteome
Mouse
Human
Danio
Fly
RMSD
0-1
2-3
3-5
5+
SPTSSB
KANTR
Kantr
tssb
pbl
43760
pet117
YM006
A0A2R8Q7U4
Klp3A
KIF4A
F1QBD7
KIF4B
Kif4
E7F084
X1WFZ0
E7FAN6
E7F2U4
$Log_{10}$ P − value FATCAT
TM Score
0.0
0.2
0.4
0.6
0.5
1.0
1.5
2.0
2.5

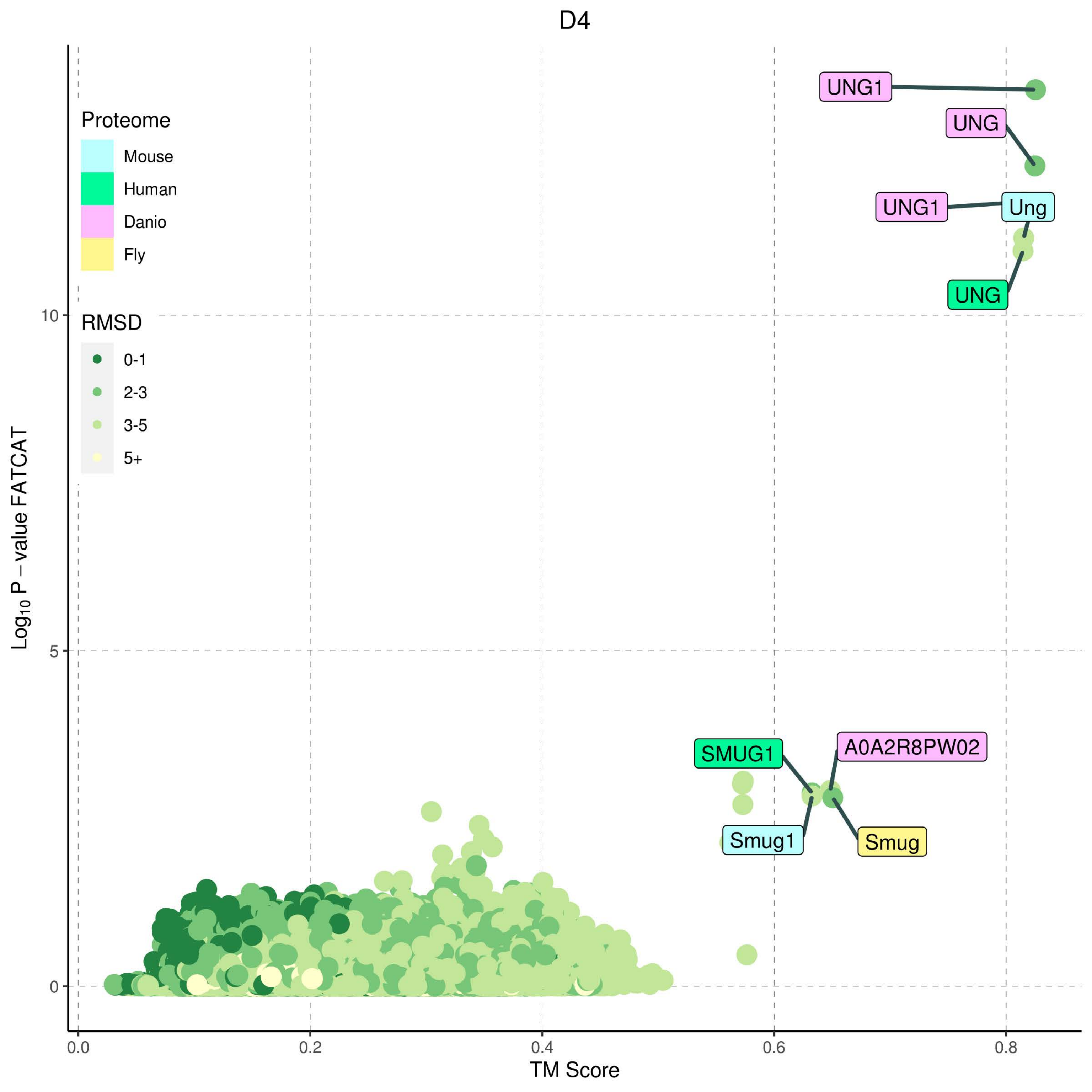

D4
Proteome
Mouse
Human
Danio
Fly
RMSD
0-1
2-3
3-5
5+
UNG1
UNG
UNG1
Ung
UNG
SMUG1
A0A2R8PW02
Smug1
Smug
Log10 P – value FATCAT
TM Score
0.0
0.2
0.4
0.6
0.8
0
5
10

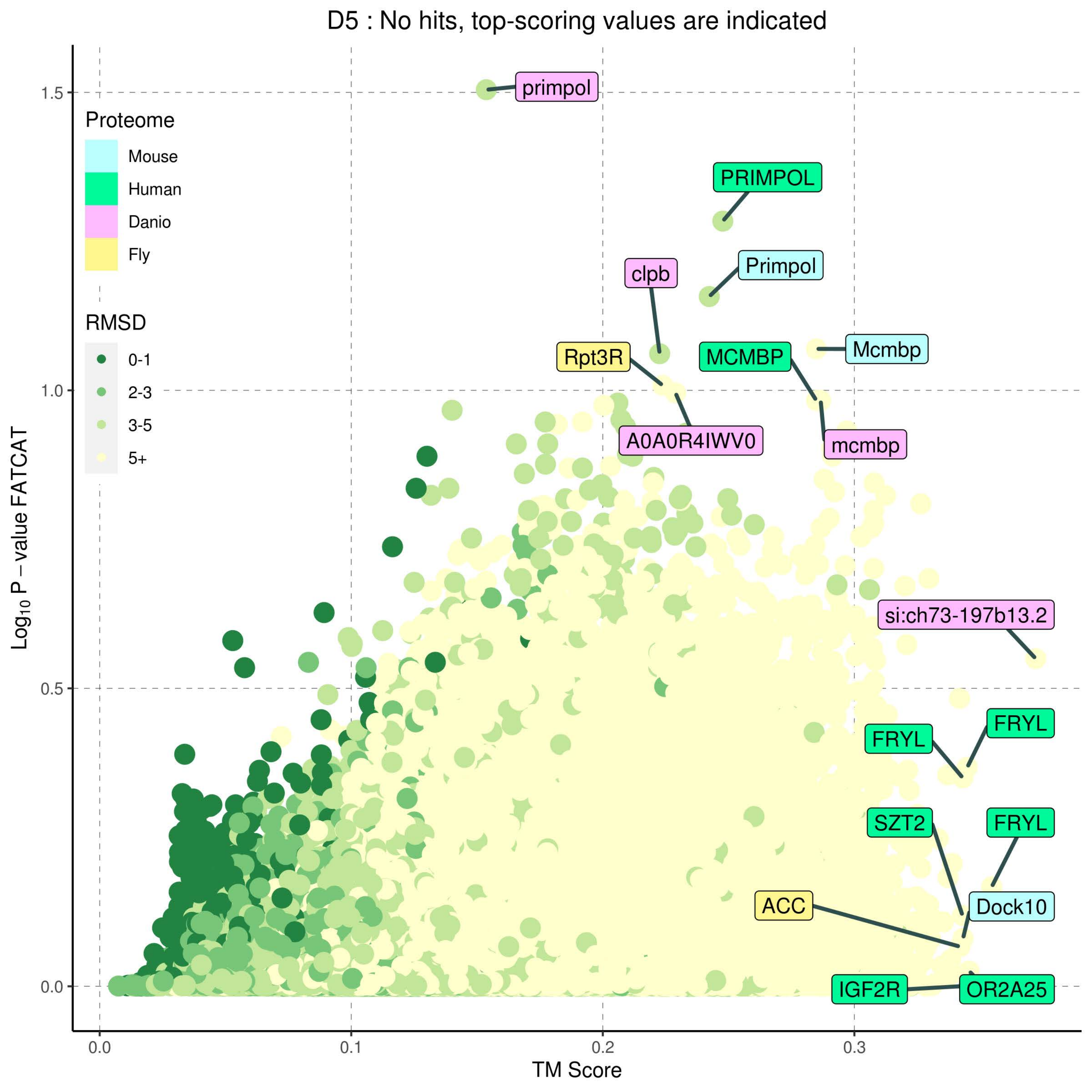
D5 : No hits, top-scoring values are indicated
Proteome
Mouse
Human
Danio
Fly
RMSD
0-1
2-3
3-5
5+
Log10 P − value FATCAT
TM Score
0.0
0.1
0.2
0.3
0.5
1.0
1.5
primpol
PRIMPOL
Primpol
clpb
Rpt3R
Mcmbp
MCMBP
A0A0R4IWV0
mcmbp
si:ch73-197b13.2
FRYL
FRYL
SZT2
FRYL
Dock10
ACC
IGF2R
OR2A25

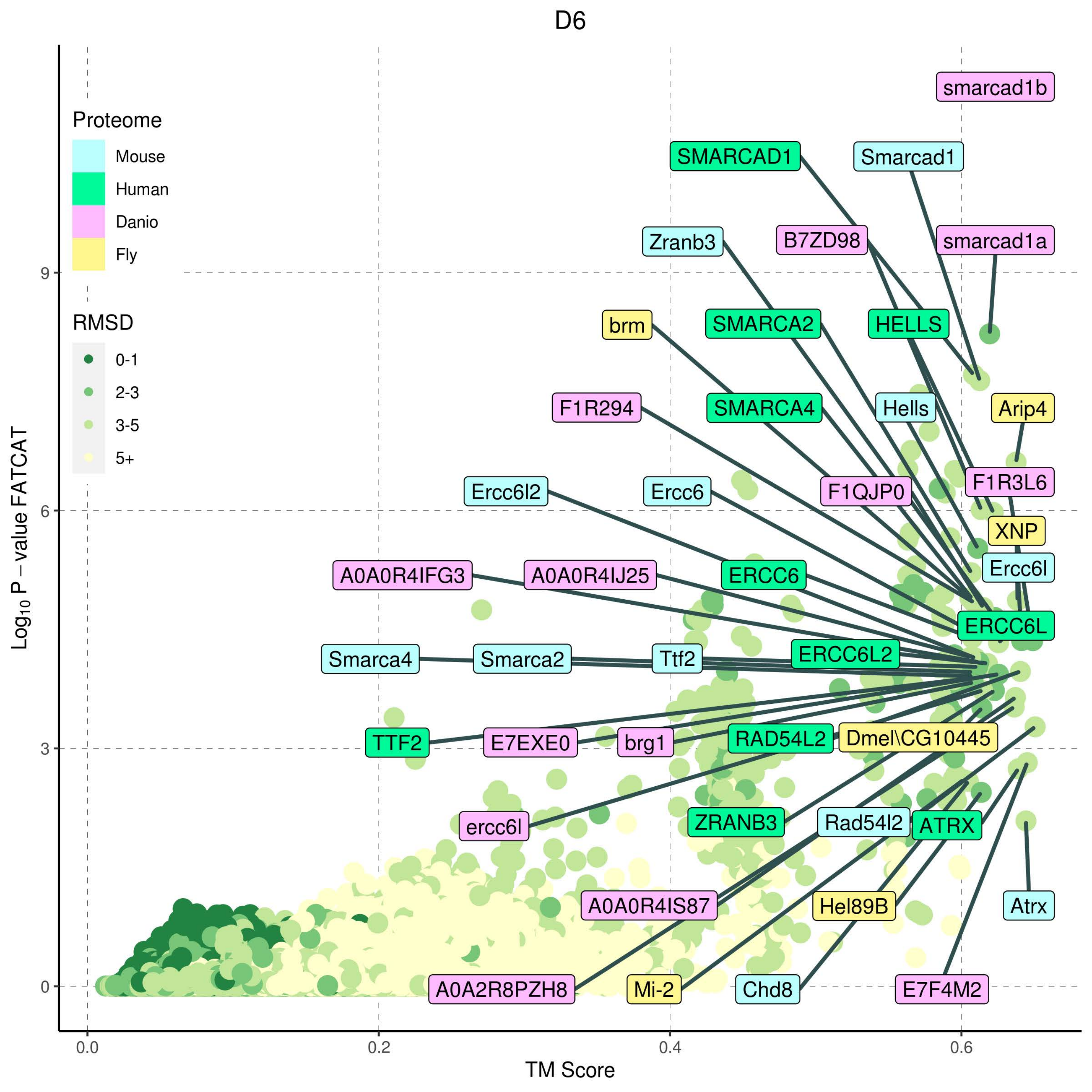
D6
Proteome
Mouse
Human
Danio
Fly
RMSD
0-1
2-3
3-5
5+
smarcad1b
SMARCAD1
Smarcad1
Zranb3
B7ZD98
smarcad1a
brm
SMARCA2
HELLS
F1R294
SMARCA4
Hells
Arip4
Ercc6l2
Ercc6
F1QJP0
F1R3L6
XNP
A0A0R4IFG3
A0A0R4IJ25
ERCC6
Ercc6l
ERCC6L
Smarca4
Smarca2
Ttf2
ERCC6L2
TTF2
E7EXE0
brg1
RAD54L2
Dmel\CG10445
ercc6l
ZRANB3
Rad54l2
ATRX
A0A0R4IS87
Hel89B
Atrx
A0A2R8PZH8
Mi-2
Chd8
E7F4M2
Log10 P − value FATCAT
TM Score
0
3
6
9
0.0
0.2
0.4
0.6

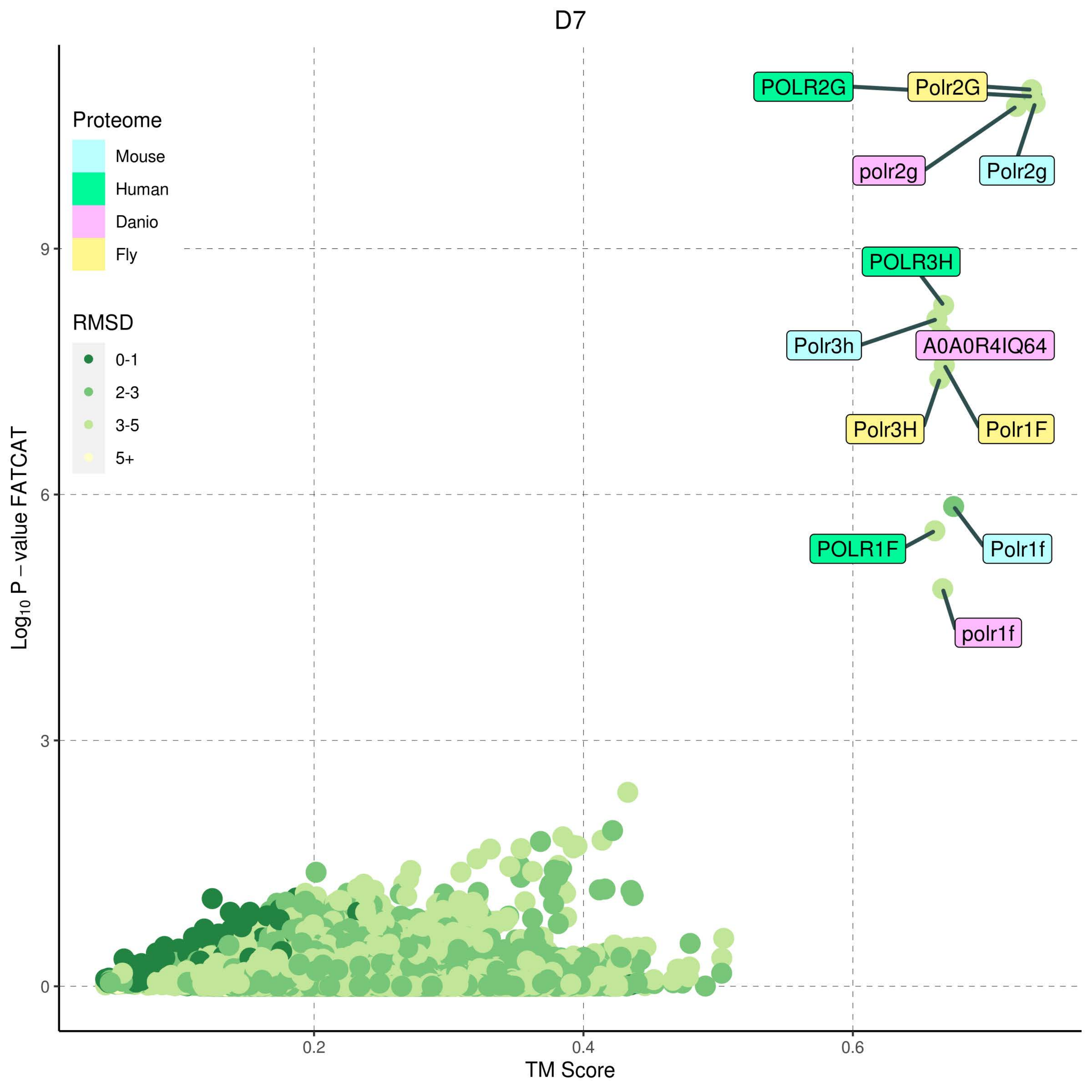
D7
Proteome
Mouse
Human
Danio
Fly
RMSD
0-1
2-3
3-5
5+
POLR2G
Polr2G
polr2g
Polr2g
POLR3H
Polr3h
A0A0R4IQ64
Polr3H
Polr1F
POLR1F
Polr1f
polr1f
Log10 P − value FATCAT
TM Score
0
3
6
9
0.2
0.4
0.6

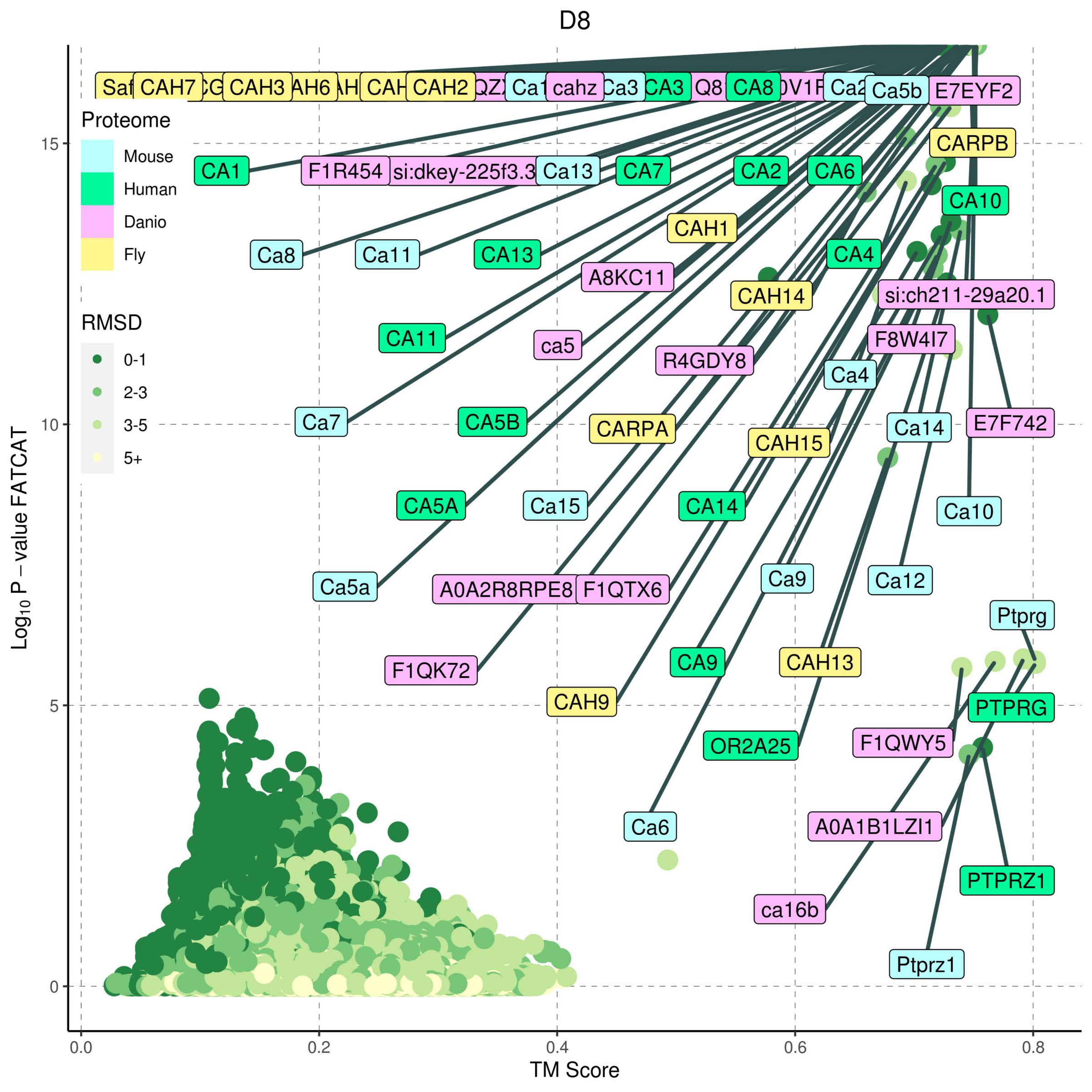
D8
Proteome
Mouse
Human
Danio
Fly
RMSD
0-1
2-3
3-5
5+
Log10 P – value FATCAT
TM Score
CAH7
CAH3
CAH2
cahz
Ca3
CA3
CA8
Ca5b
E7EYF2
CARPB
CA1
F1R454
si:dkey-225f3.3
Ca13
CA7
CA2
CA6
CA10
CAH1
Ca8
Ca11
CA13
CA4
A8KC11
CAH14
si:ch211-29a20.1
CA11
ca5
R4GDY8
F8W4I7
Ca4
Ca7
CA5B
CARPA
CAH15
Ca14
E7F742
CA5A
Ca15
CA14
Ca10
Ca5a
A0A2R8RPE8
F1QTX6
Ca9
Ca12
Ptprg
F1QK72
CA9
CAH13
CAH9
PTPRG
OR2A25
F1QWY5
Ca6
A0A1B1LZI1
PTPRZ1
ca16b
Ptprz1

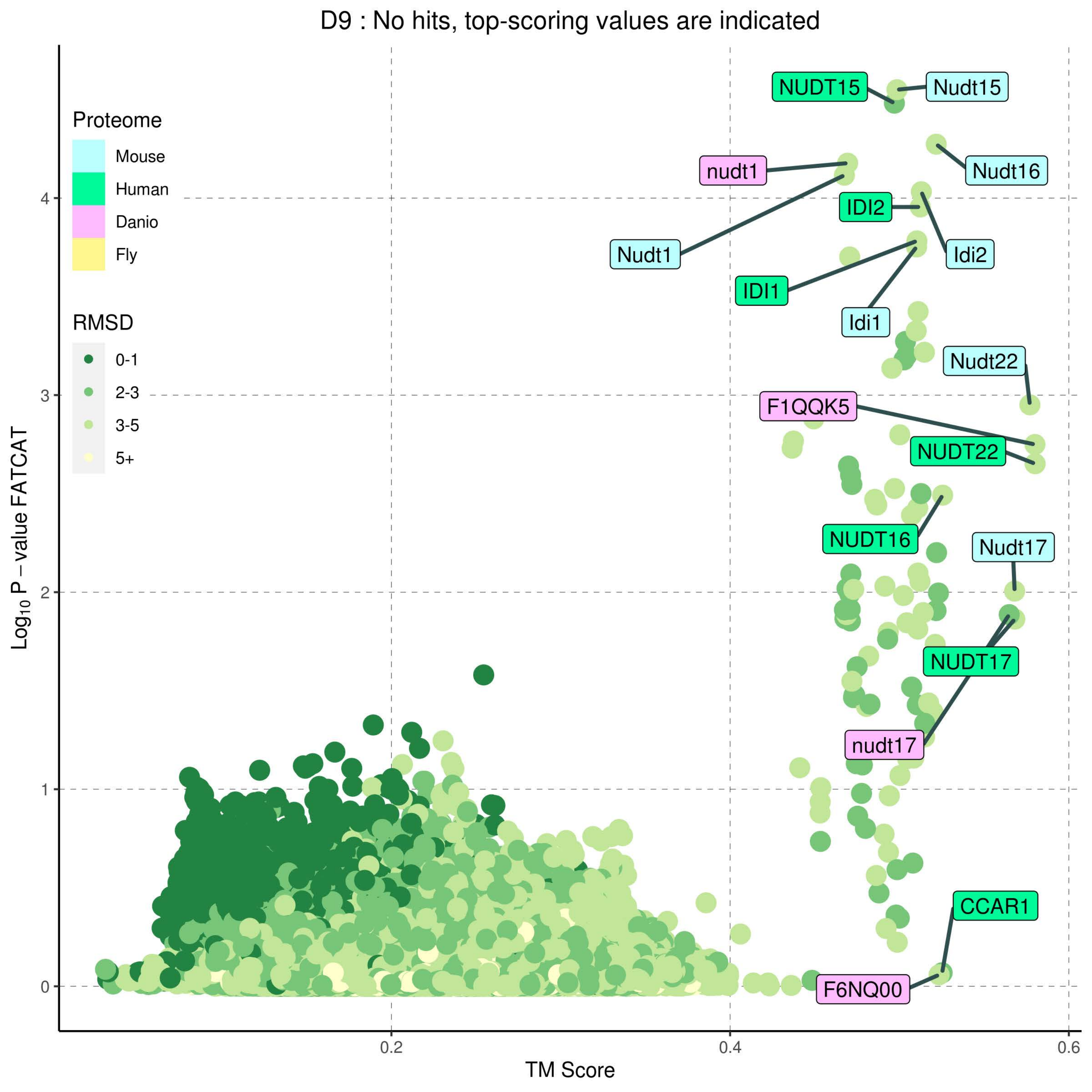
D9 : No hits, top-scoring values are indicated
Proteome
Mouse
Human
Danio
Fly
RMSD
0-1
2-3
3-5
5+
NUDT15
Nudt15
nudt1
Nudt16
IDI2
Nudt1
Idi2
IDI1
Idi1
Nudt22
F1QQK5
NUDT22
NUDT16
Nudt17
NUDT17
nudt17
CCAR1
F6NQ00
Log10 P – value FATCAT
TM Score
0
1
2
3
4
0.2
0.4
0.6

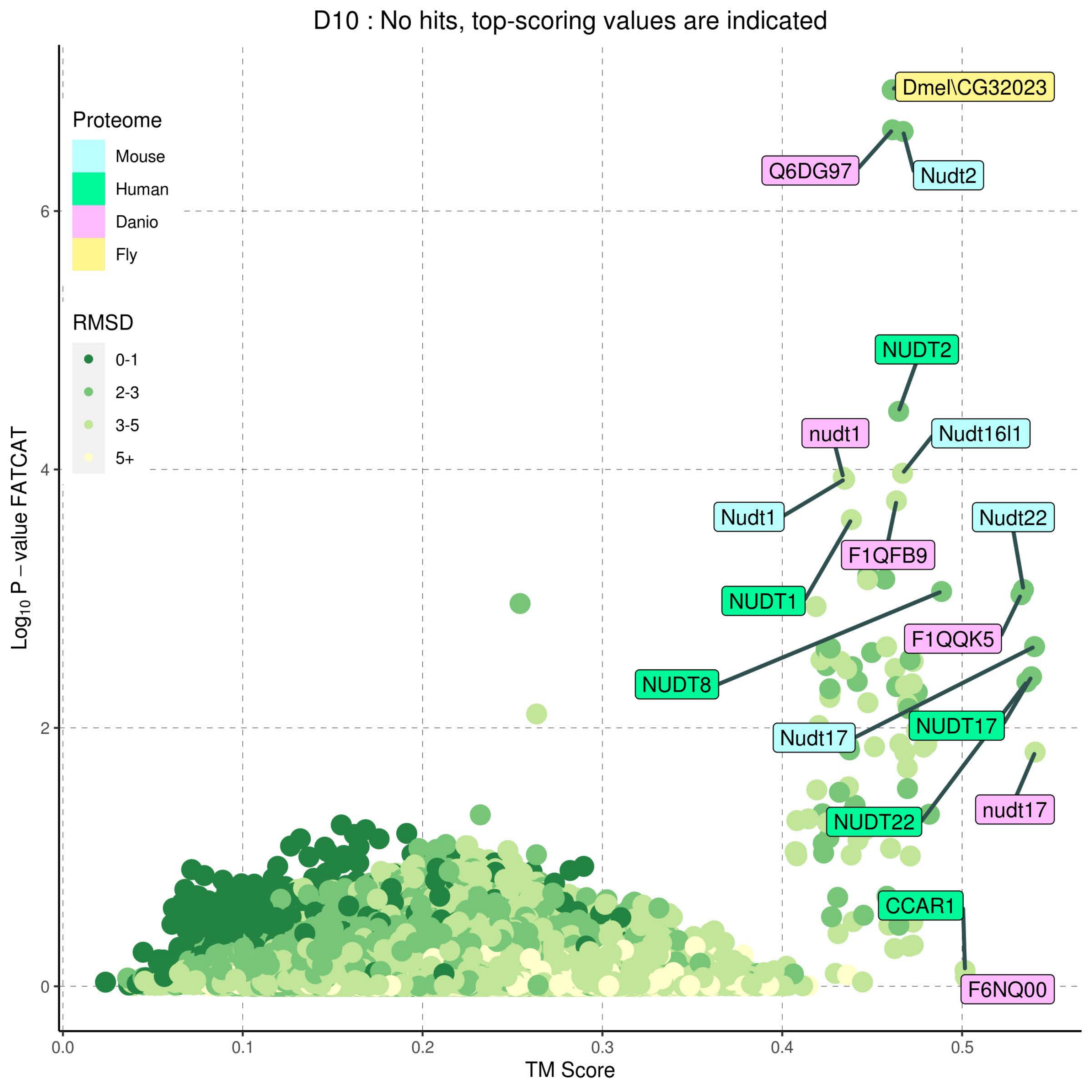

D10 : No hits, top-scoring values are indicated
Proteome
Mouse
Human
Danio
Fly
RMSD
0-1
2-3
3-5
5+
Log10 P − value FATCAT
TM Score
0
2
4
6
0.0
0.1
0.2
0.3
0.4
0.5
Dmel\CG32023
Q6DG97
Nudt2
NUDT2
nudt1
Nudt16l1
Nudt1
F1QFB9
Nudt22
NUDT1
F1QQK5
NUDT8
Nudt17
NUDT17
nudt17
NUDT22
CCAR1
F6NQ00

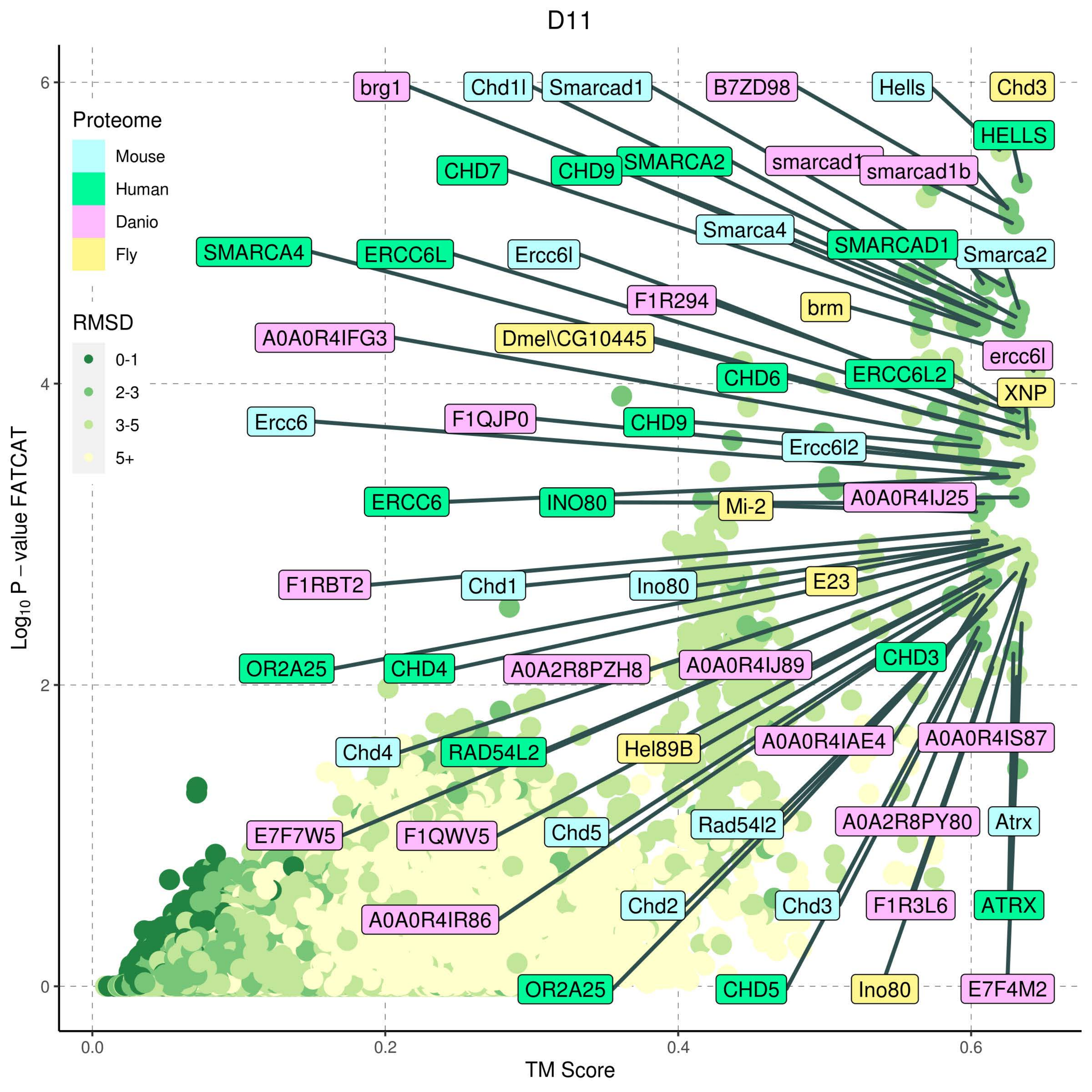

D11
Proteome
Mouse
Human
Danio
Fly
RMSD
0-1
2-3
3-5
5+
Log10 P − value FATCAT
TM Score
0.0
0.2
0.4
0.6
0
2
4
6
brg1
Chd1l
Smarcad1
B7ZD98
Hells
Chd3
HELLS
SMARCA2
smarcad1
smarcad1b
CHD7
CHD9
Smarca4
SMARCAD1
Smarca2
SMARCA4
ERCC6L
Ercc6l
F1R294
brm
A0A0R4IFG3
Dmel\CG10445
ercc6l
CHD6
ERCC6L2
XNP
Ercc6
F1QJP0
CHD9
Ercc6l2
ERCC6
INO80
Mi-2
A0A0R4IJ25
F1RBT2
Chd1
Ino80
E23
OR2A25
CHD4
A0A2R8PZH8
A0A0R4IJ89
CHD3
Chd4
RAD54L2
Hel89B
A0A0R4IAE4
A0A0R4IS87
E7F7W5
F1QWV5
Chd5
Rad54l2
A0A2R8PY80
Atrx
Chd2
Chd3
F1R3L6
ATRX
A0A0R4IR86
OR2A25
CHD5
Ino80
E7F4M2

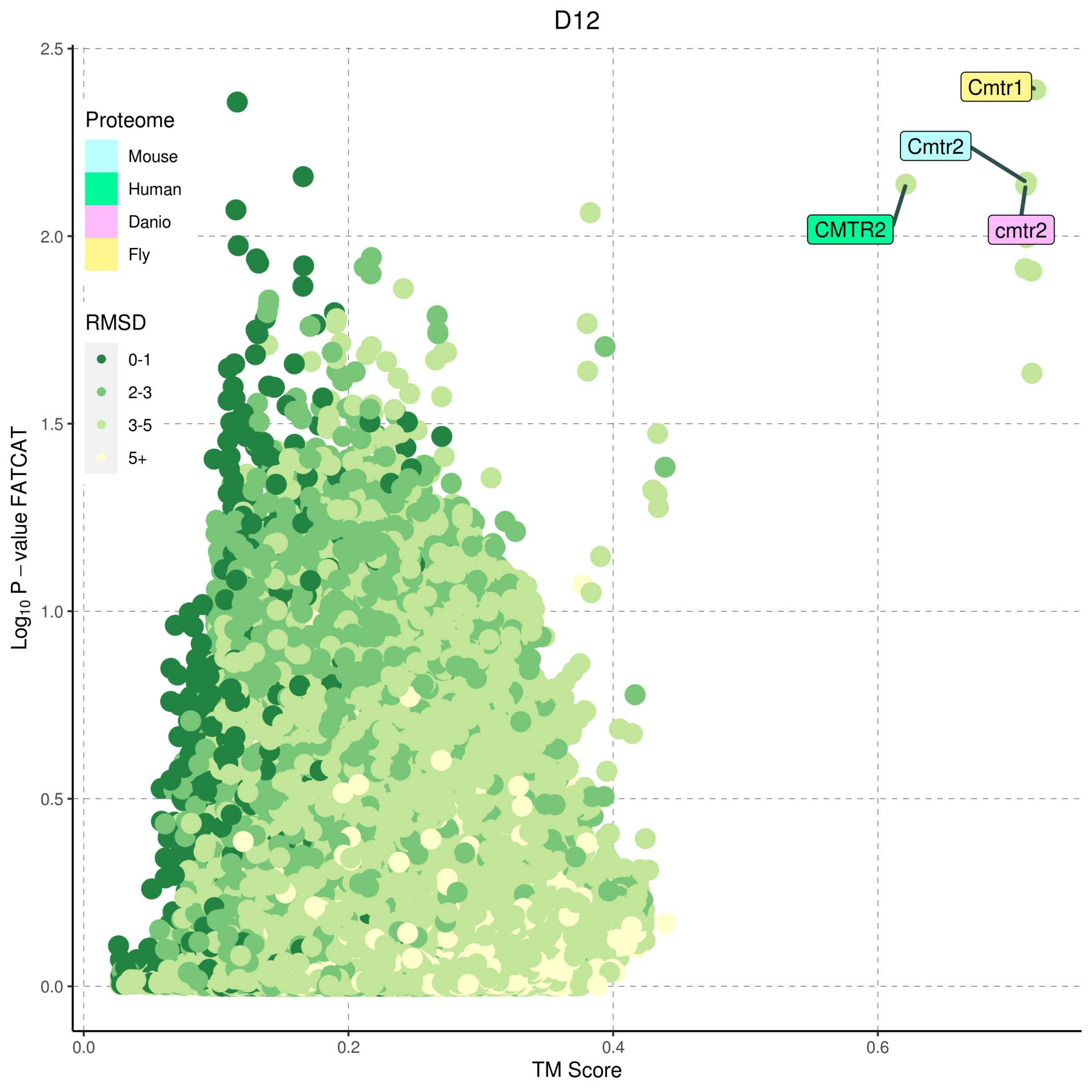
D12
Proteome
Mouse
Human
Danio
Fly
RMSD
0-1
2-3
3-5
5+
Cmtr1
Cmtr2
CMTR2
cmtr2
Log10 P − value FATCAT
TM Score
0.0
0.5
1.0
1.5
2.0
2.5
0.0
0.2
0.4
0.6

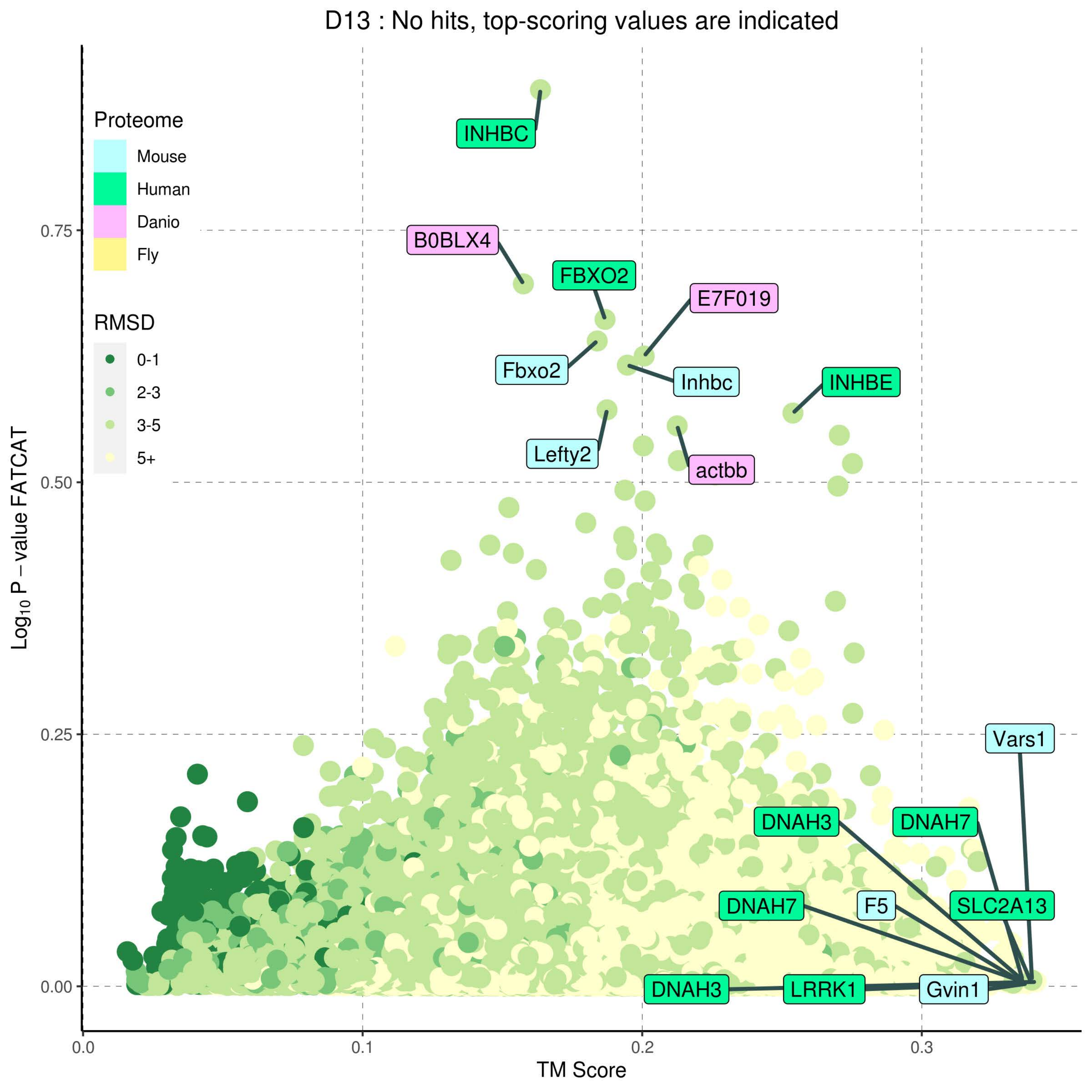
D13 : No hits, top-scoring values are indicated
Proteome
Mouse
Human
Danio
Fly
RMSD
0-1
2-3
3-5
5+
INHBC
B0BLX4
FBXO2
E7F019
Fbxo2
Inhbc
INHBE
Lefty2
actbb
Vars1
DNAH3
DNAH7
DNAH7
F5
SLC2A13
DNAH3
LRRK1
Gvin1
0.00
0.25
0.50
0.75
0.0
0.1
0.2
0.3
Log10 P − value FATCAT
TM Score

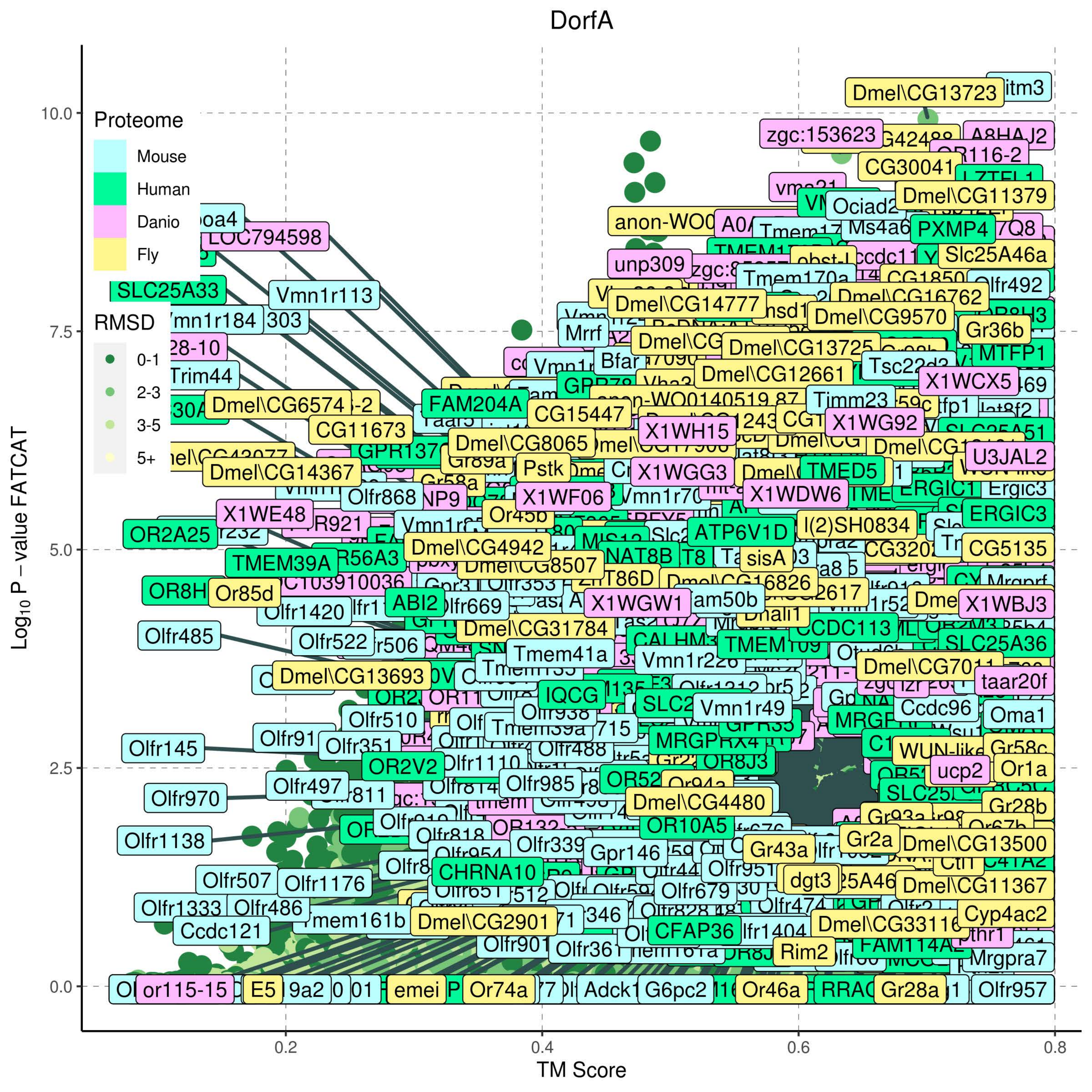
DorfA
Proteome
Mouse
Human
Danio
Fly
RMSD
0-1
2-3
3-5
5+
Log10 P – value FATCAT
10.0
7.5
5.0
2.5
0.0
TM Score
0.2
0.4
0.6
0.8

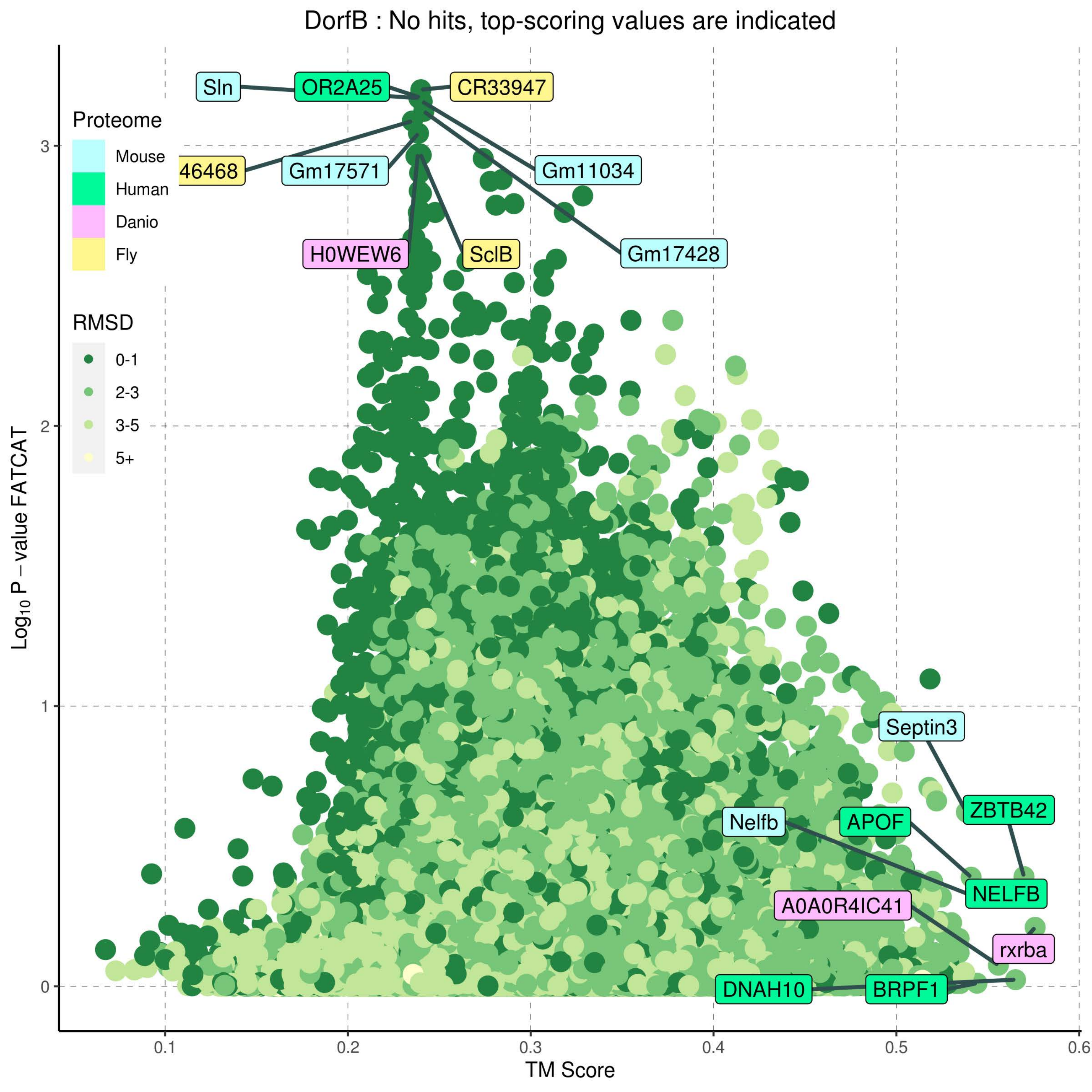

DorfB : No hits, top-scoring values are indicated
Proteome
Mouse
Human
Danio
Fly
RMSD
0-1
2-3
3-5
5+
Log10 P − value FATCAT
TM Score
0
1
2
3
0.1
0.2
0.3
0.4
0.5
0.6
Sln
OR2A25
CR33947
46468
Gm17571
Gm11034
H0WEW6
SclB
Gm17428
Septin3
ZBTB42
Nelfb
APOF
NELFB
A0A0R4IC41
rxrba
DNAH10
BRPF1

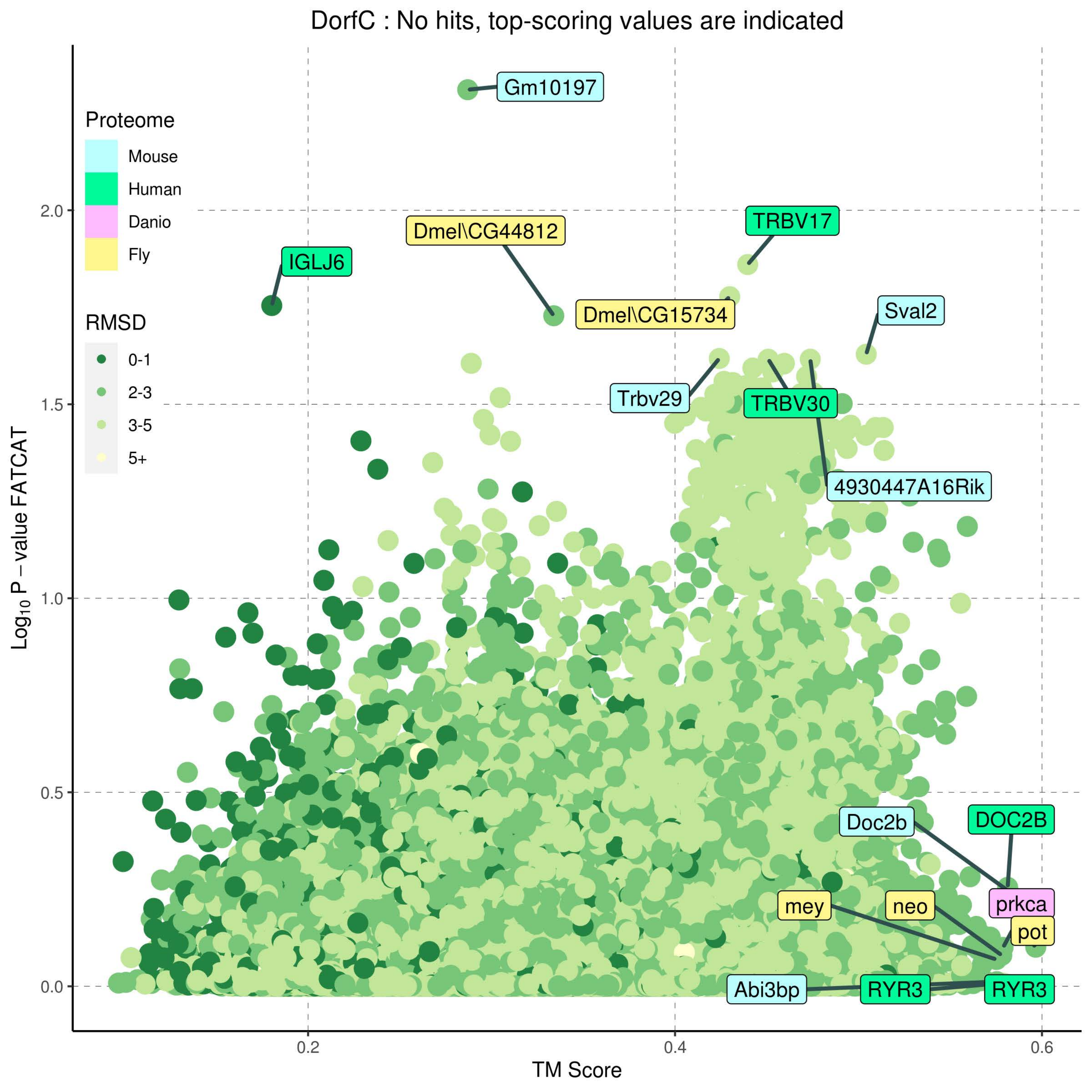
DorfC : No hits, top-scoring values are indicated
Proteome
Mouse
Human
Danio
Fly
RMSD
0-1
2-3
3-5
5+
Log10 P − value FATCAT
TM Score
0.0
0.5
1.0
1.5
2.0
0.2
0.4
0.6
Gm10197
Dmel\CG44812
IGLJ6
TRBV17
Dmel\CG15734
Sval2
Trbv29
TRBV30
4930447A16Rik
Doc2b
DOC2B
prkca
mey
neo
pot
Abi3bp
RYR3
RYR3

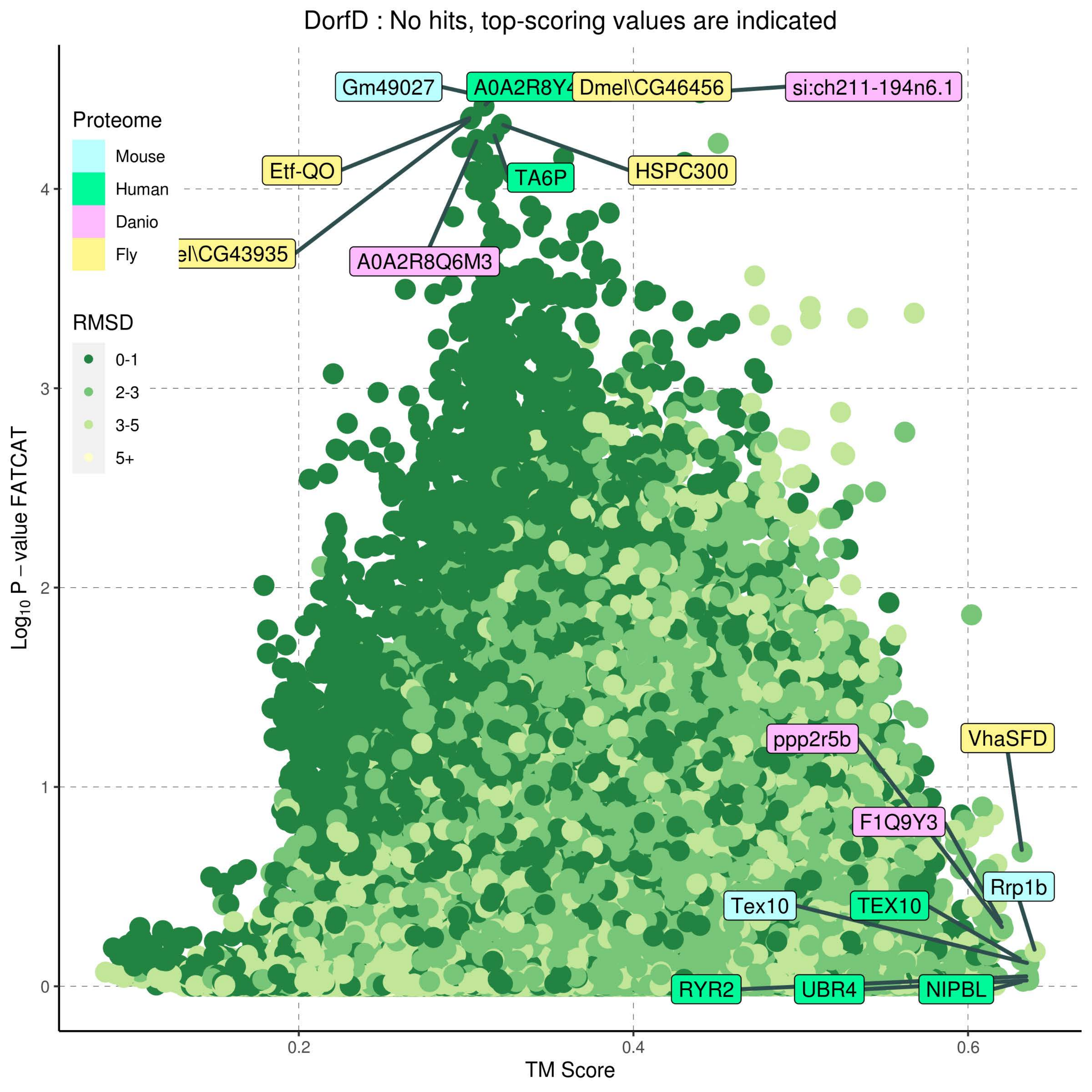

DorfD : No hits, top-scoring values are indicated
Proteome
Mouse
Human
Danio
Fly
RMSD
0-1
2-3
3-5
5+
Gm49027
A0A2R8Y4
Dmel\CG46456
si:ch211-194n6.1
Etf-QO
TA6P
HSPC300
el\CG43935
A0A2R8Q6M3
ppp2r5b
VhaSFD
F1Q9Y3
Rrp1b
Tex10
TEX10
RYR2
UBR4
NIPBL
Log10 P − value FATCAT
TM Score
0
1
2
3
4
0.2
0.4
0.6

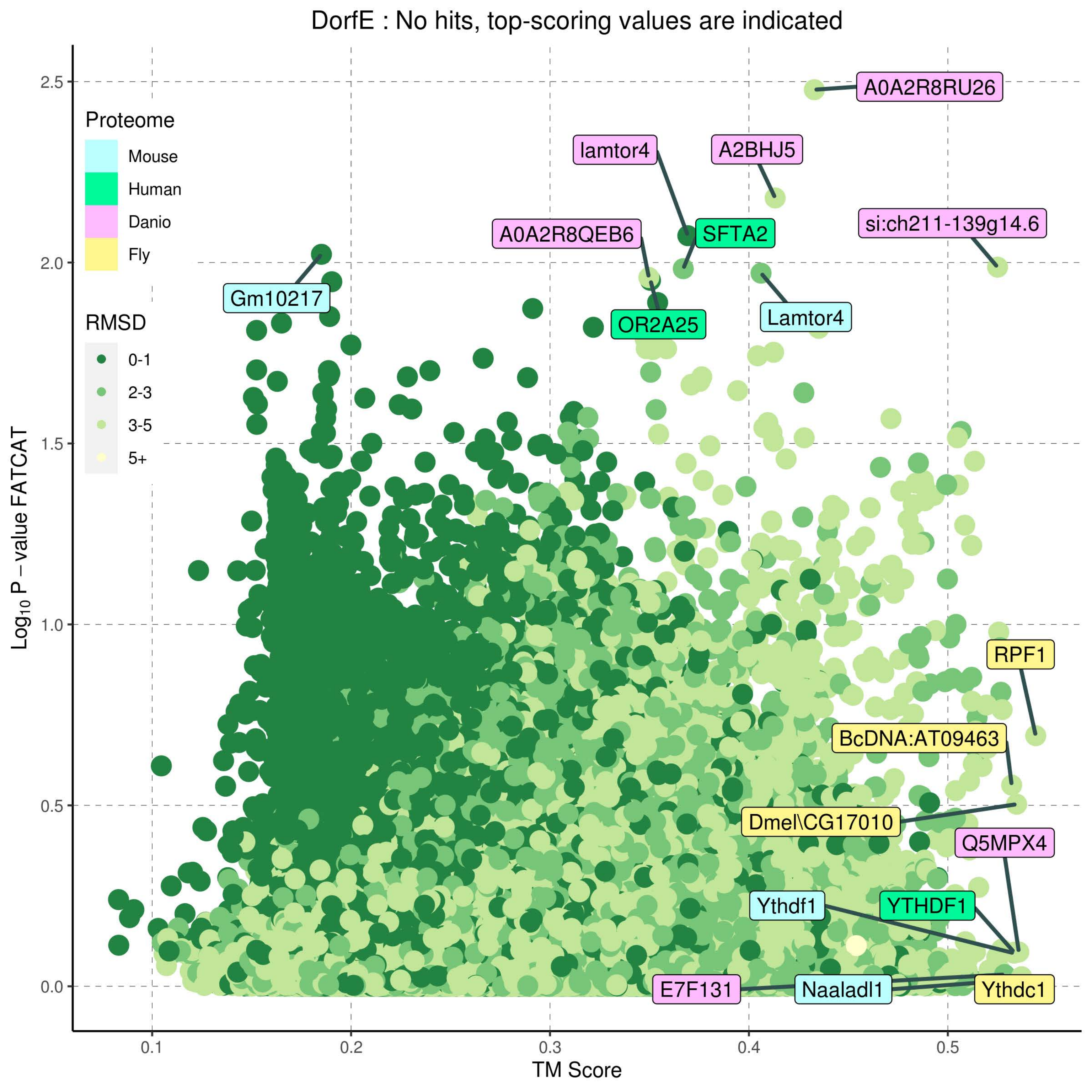
DorfE : No hits, top-scoring values are indicated
Proteome
Mouse
Human
Danio
Fly
RMSD
0-1
2-3
3-5
5+
A0A2R8RU26
lamtor4
A2BHJ5
A0A2R8QEB6
SFTA2
si:ch211-139g14.6
Gm10217
OR2A25
Lamtor4
RPF1
BcDNA:AT09463
Dmel\CG17010
Q5MPX4
Ythdf1
YTHDF1
E7F131
Naaladl1
Ythdc1
Log10 P − value FATCAT
TM Score
0.0
0.5
1.0
1.5
2.0
2.5
0.1
0.2
0.3
0.4
0.5

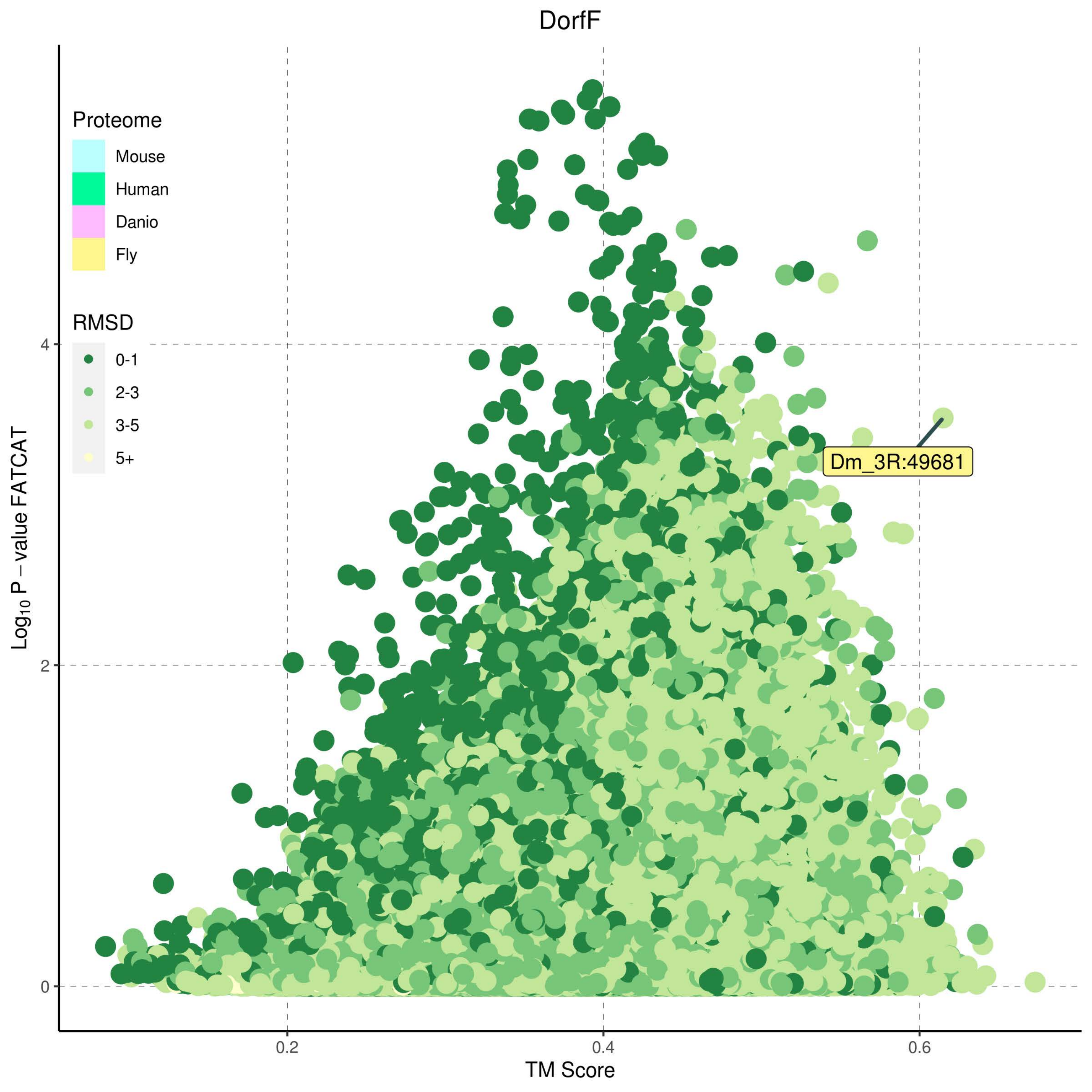

DorfF
Proteome
Mouse
Human
Danio
Fly
RMSD
0-1
2-3
3-5
5+
Dm_3R:49681
Log10 P − value FATCAT
TM Score
0
2
4
0.2
0.4
0.6

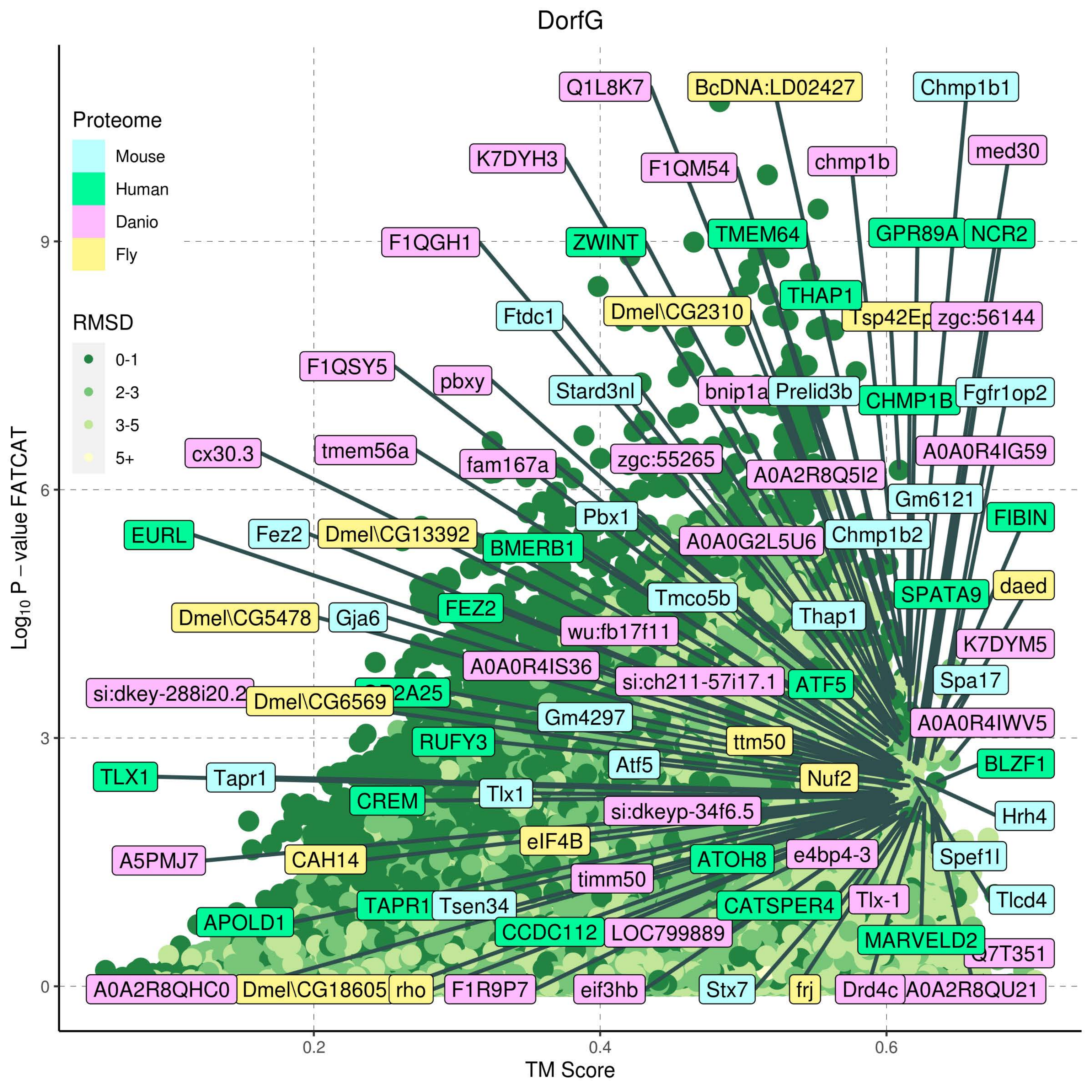

DorfG
Proteome
Mouse
Human
Danio
Fly
RMSD
0-1
2-3
3-5
5+
Log10 P – value FATCAT
TM Score
0
3
6
9
0.2
0.4
0.6
Q1L8K7
BcDNA:LD02427
Chmp1b1
K7DYH3
F1QM54
chmp1b
med30
F1QGH1
ZWINT
TMEM64
GPR89A
NCR2
THAP1
Ftdc1
Dmel\CG2310
zgc:56144
F1QSY5
pbxy
Stard3nl
bnip1a
Prelid3b
CHMP1B
Fgfr1op2
cx30.3
tmem56a
fam167a
zgc:55265
A0A2R8Q5I2
A0A0R4IG59
Gm6121
FIBIN
EURL
Fez2
Dmel\CG13392
BMERB1
Pbx1
A0A0G2L5U6
Chmp1b2
Tmco5b
daed
SPATA9
Dmel\CG5478
Gja6
FEZ2
wu:fb17f11
Thap1
K7DYM5
A0A0R4IS36
si:ch211-57i17.1
ATF5
Spa17
si:dkey-288i20.2
Dmel\CG6569
Gm4297
A0A0R4IWV5
RUFY3
ttm50
TLX1
Tapr1
Atf5
Nuf2
BLZF1
CREM
Tlx1
si:dkeyp-34f6.5
Hrh4
eIF4B
A5PMJ7
CAH14
ATOH8
e4bp4-3
Spef1l
timm50
TAPR1
Tsen34
CATSPER4
Tlx-1
Tlcd4
APOLD1
CCDC112
LOC799889
MARVELD2
A0A2R8QHC0
Dmel\CG18605
rho
F1R9P7
eif3hb
Stx7
frj
Drd4c
A0A2R8QU21

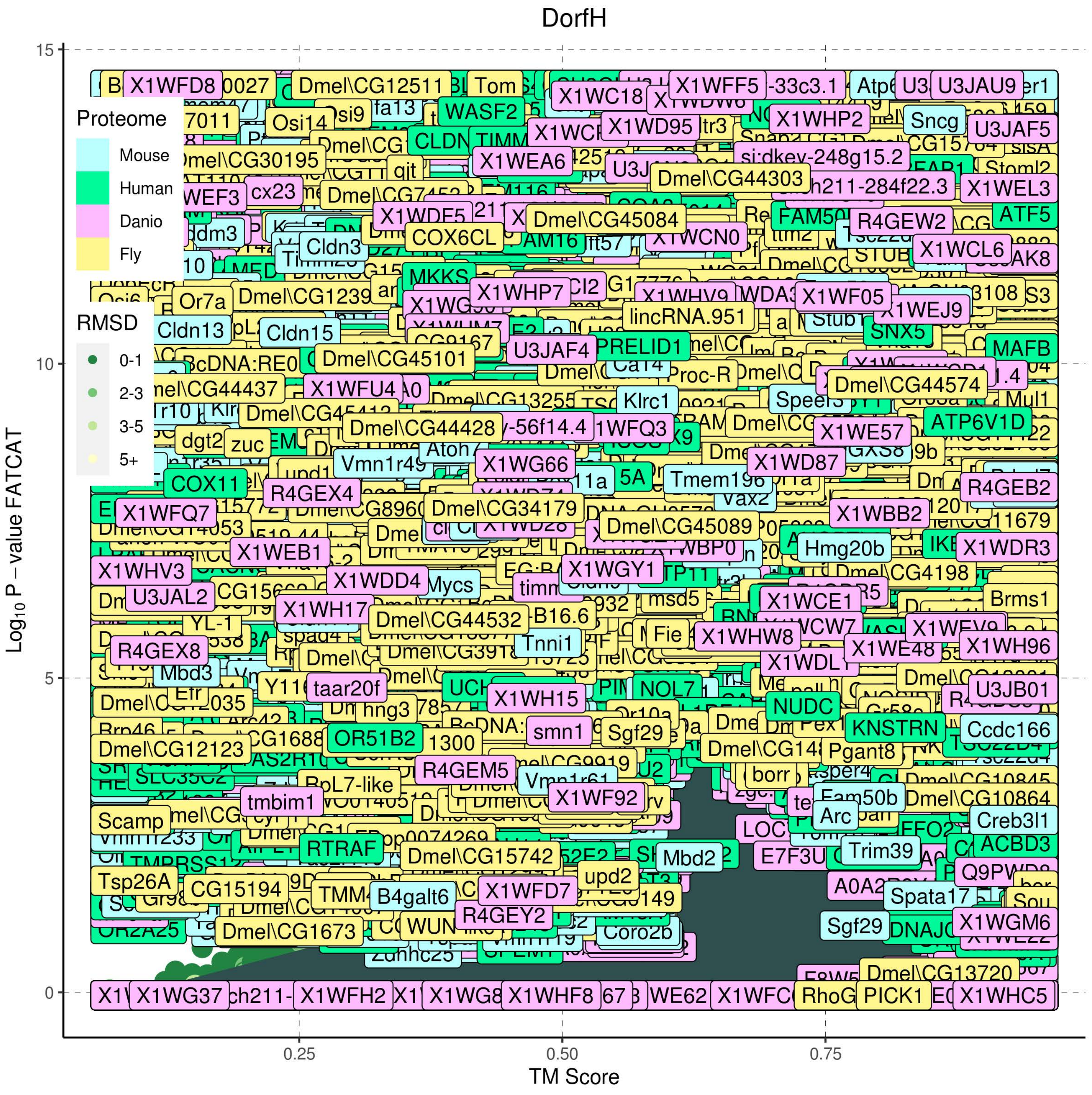
DorfH
Proteome
Mouse
Human
Danio
Fly
RMSD
0-1
2-3
3-5
5+
Log10 P – value FATCAT
TM Score
0
5
10
15
0.25
0.50
0.75

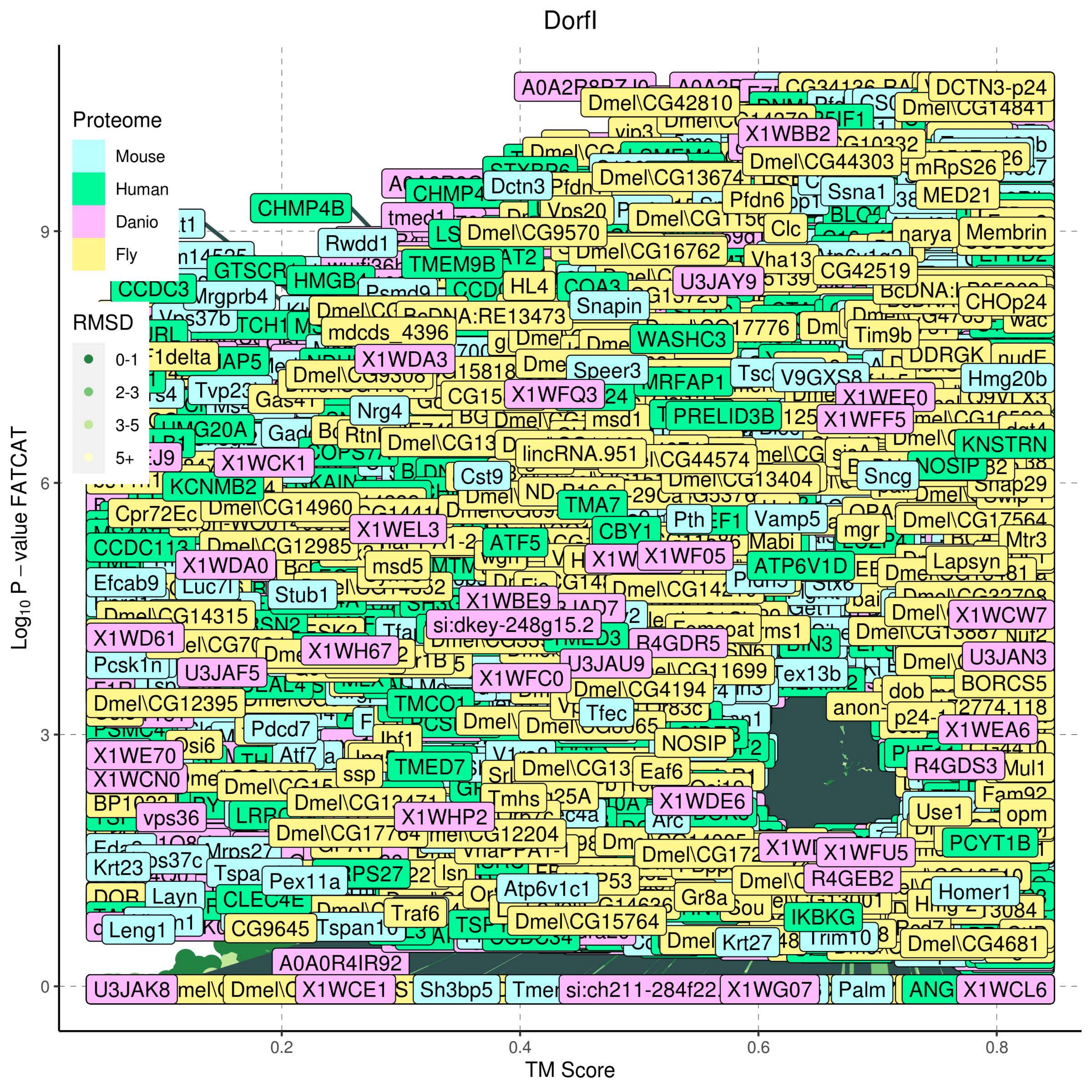

Dorfl
Proteome
Mouse
Human
Danio
Fly
RMSD
0-1
2-3
3-5
5+
Log10 P – value FATCAT
TM Score
0
3
6
9
0.2
0.4
0.6
0.8
DCTN3-p24
Dmel\CG42810
Dmel\CG14841
X1WBB2
Dmel\CG44303
mRpS26
Dctn3
Dmel\CG13674
Ssna1
MED21
CHMP4B
Pfdn6
Clc
Rwdd1
Dmel\CG9570
narya
Membrin
Dmel\CG16762
Vha13
TMEM9B
U3JAY9
CG42519
HMGB4
HL4
Snapin
CHOp24
CCDC3
Mrgprb4
BcDNA:RE13473
mdcds_4396
Tim9b
WASHC3
X1WDA3
Speer3
DDRGK
Hmg20b
V9GXS8
X1WFQ3
MRFAP1
X1WEE0
Tvp23
Nrg4
PRELID3B
X1WFF5
HMG20A
KNSTRN
X1WCK1
lincRNA.951
Cst9
Dmel\CG13404
Sncg
NOSIP
Snap29
KCNMB2
Cpr72Ec
Dmel\CG14960
TMA7
Pth
Vamp5
Dmel\CG17564
X1WEL3
CBY1
mgr
CCDC113
Dmel\CG12985
ATF5
Mtr3
X1WF05
X1WDA0
msd5
ATP6V1D
Lapsyn
Efcab9
Stub1
X1WBE9
X1WCW7
Dmel\CG14315
si:dkey-248g15.2
X1WD61
X1WH67
R4GDR5
Tex13b
U3JAF5
U3JAU9
U3JAN3
X1WFC0
Dmel\CG4194
BORCS5
Dmel\CG12395
TMCO1
Tfec
dob
Pdcd7
X1WEA6
NOSIP
X1WE70
TMED7
Eaf6
R4GDS3
Mul1
X1WCN0
ssp
vps36
Tmhs
X1WDE6
X1WHP2
Use1
opm
Dmel\CG17764
Dmel\CG12204
PCYT1B
X1WFU5
Krt23
Pex11a
R4GEB2
Layn
CLEC4E
Atp6v1c1
Gr8a
Homer1
Traf6
IKBKG
Leng1
CG9645
Dmel\CG15764
Krt27
Trim10
Dmel\CG4681
A0A0R4IR92
U3JAK8
X1WCE1
Sh3bp5
X1WG07
Palm
X1WCL6

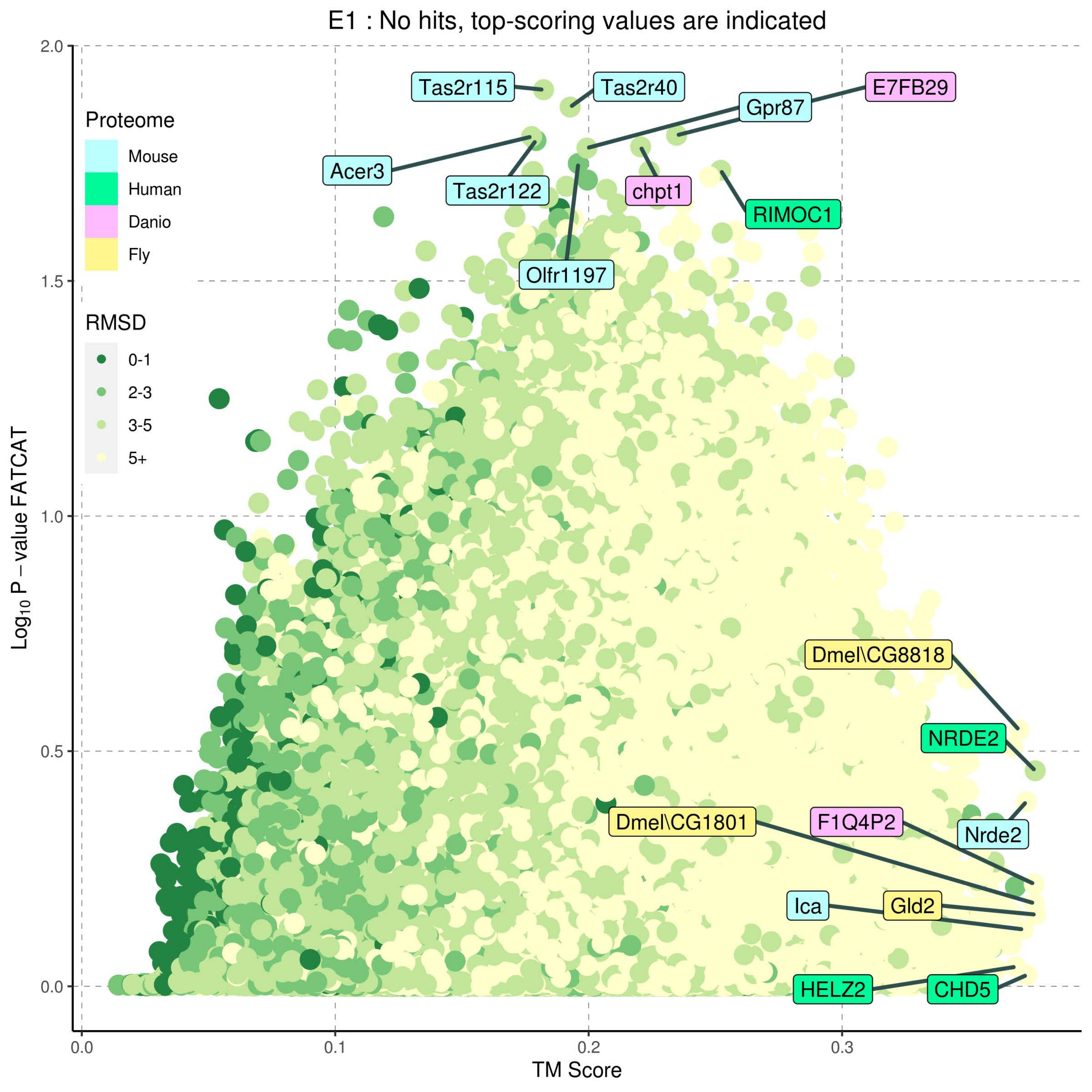

E1 : No hits, top-scoring values are indicated
Proteome
Mouse
Human
Danio
Fly
RMSD
0-1
2-3
3-5
5+
Tas2r115
Tas2r40
E7FB29
Gpr87
Acer3
Tas2r122
chpt1
RIMOC1
Olfr1197
Dmel\CG8818
NRDE2
Dmel\CG1801
F1Q4P2
Nrde2
Ica
Gld2
HELZ2
CHD5
Log10 P − value FATCAT
TM Score
0.0
0.1
0.2
0.3
0.5
1.0
1.5
2.0

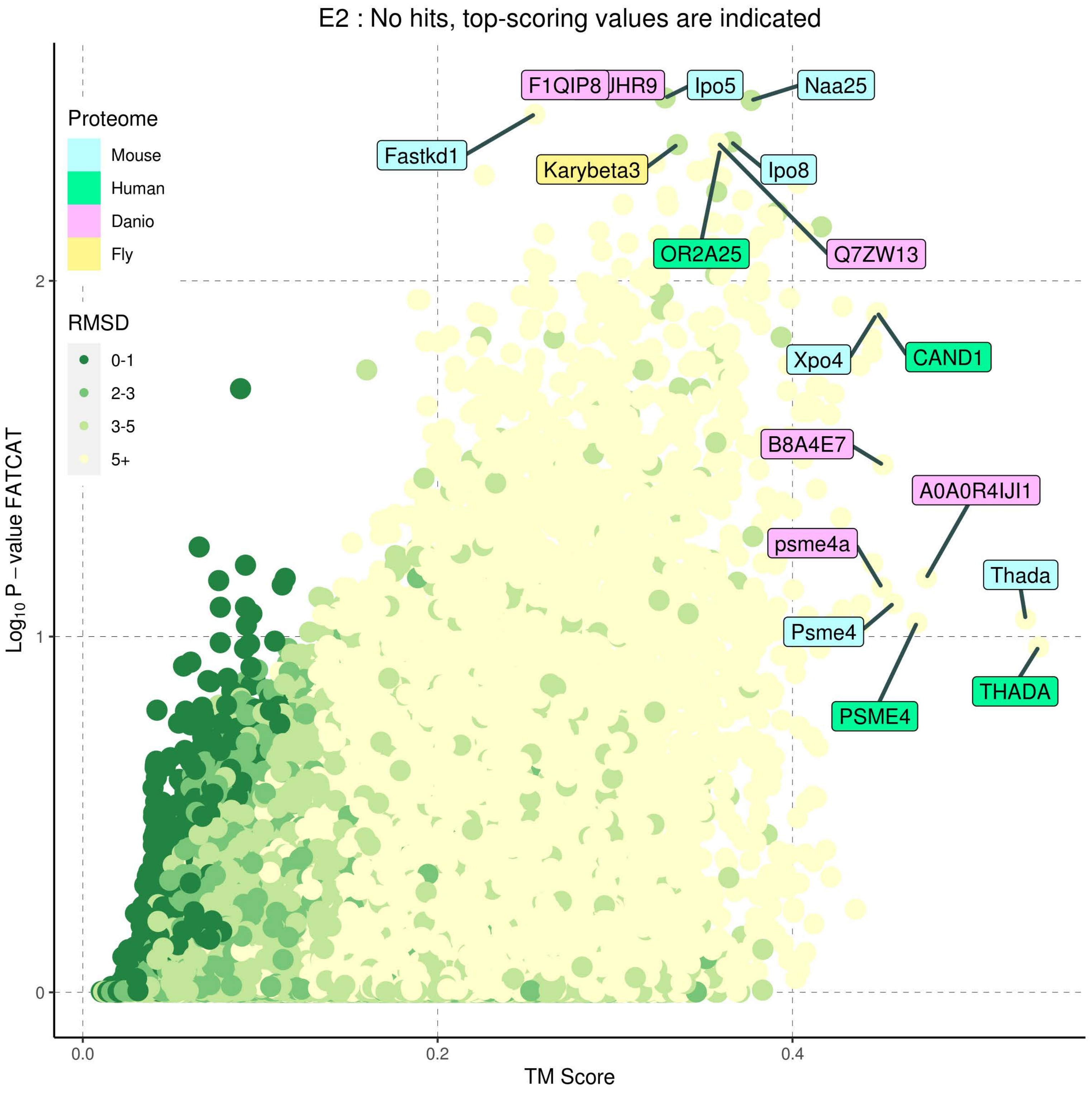
E2 : No hits, top-scoring values are indicated
Proteome
Mouse
Human
Danio
Fly
RMSD
0-1
2-3
3-5
5+
Log10 P − value FATCAT
TM Score
0.0
0.2
0.4
0
1
2
F1QIP8
JHR9
Ipo5
Naa25
Fastkd1
Karybeta3
Ipo8
OR2A25
Q7ZW13
Xpo4
CAND1
B8A4E7
A0A0R4IJI1
psme4a
Thada
Psme4
THADA
PSME4

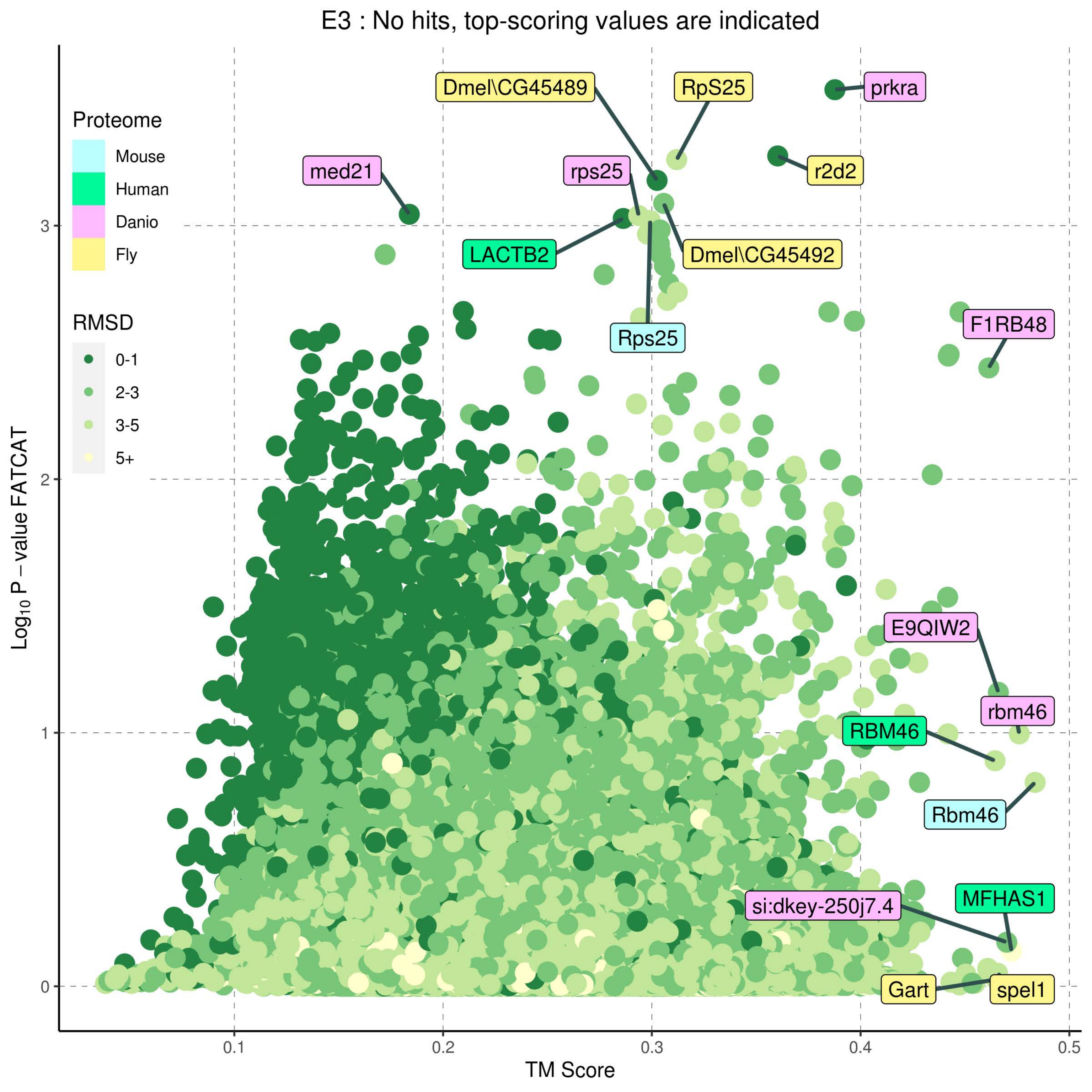
E3 : No hits, top-scoring values are indicated
Proteome
Mouse
Human
Danio
Fly
RMSD
0-1
2-3
3-5
5+
Dmel\CG45489
RpS25
prkra
med21
rps25
r2d2
LACTB2
Dmel\CG45492
Rps25
F1RB48
E9QIW2
rbm46
RBM46
Rbm46
MFHAS1
si:dkey-250j7.4
Gart
spel1
Log10 P − value FATCAT
TM Score
0
1
2
3
0.1
0.2
0.3
0.4
0.5

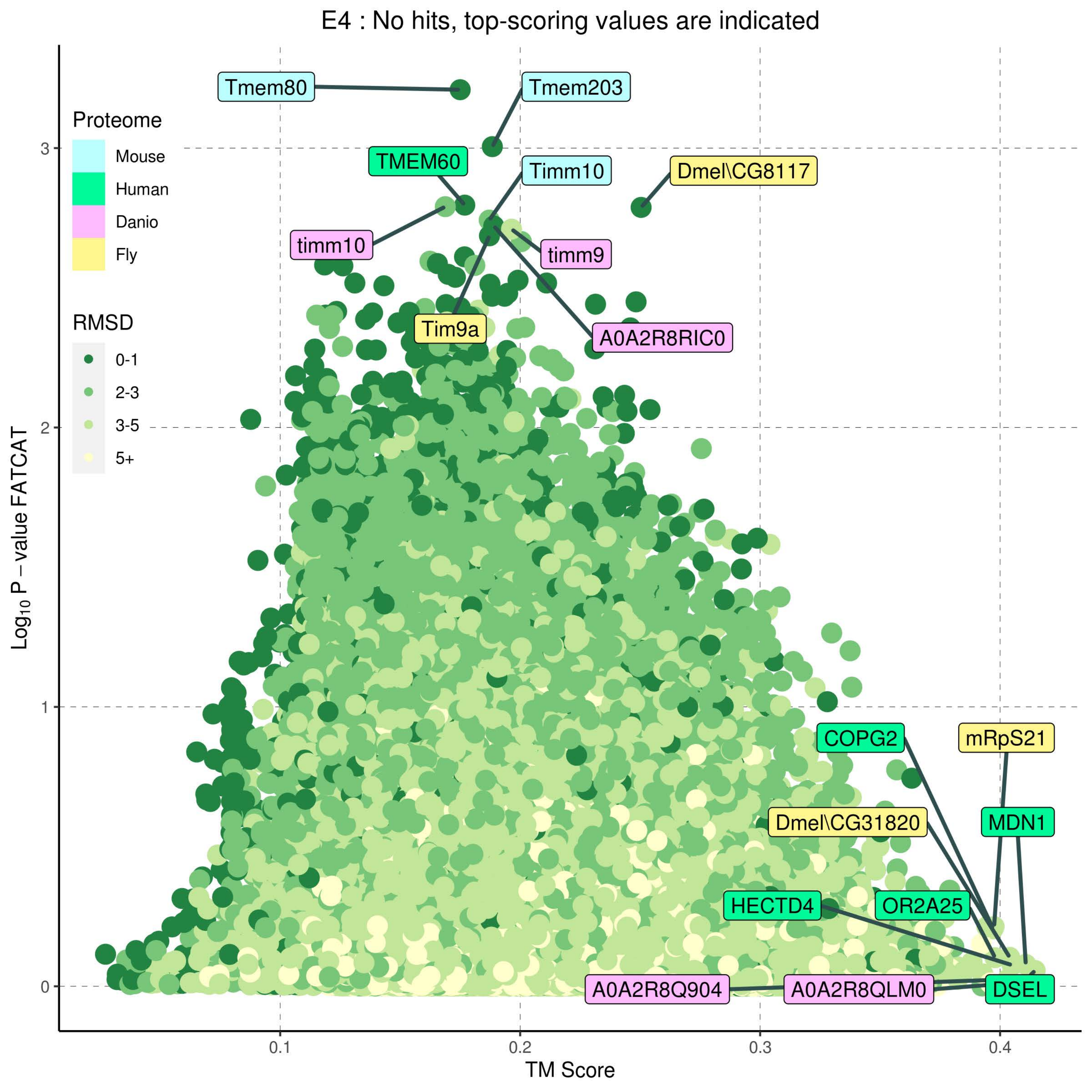

E4 : No hits, top-scoring values are indicated
Proteome
Mouse
Human
Danio
Fly
RMSD
0-1
2-3
3-5
5+
Tmem80
Tmem203
TMEM60
Timm10
Dmel\CG8117
timm10
timm9
Tim9a
A0A2R8RIC0
COPG2
mRpS21
Dmel\CG31820
MDN1
HECTD4
OR2A25
A0A2R8Q904
A0A2R8QLM0
DSEL
3
2
1
0
0.1
0.2
0.3
0.4
TM Score
Log10 P – value FATCAT

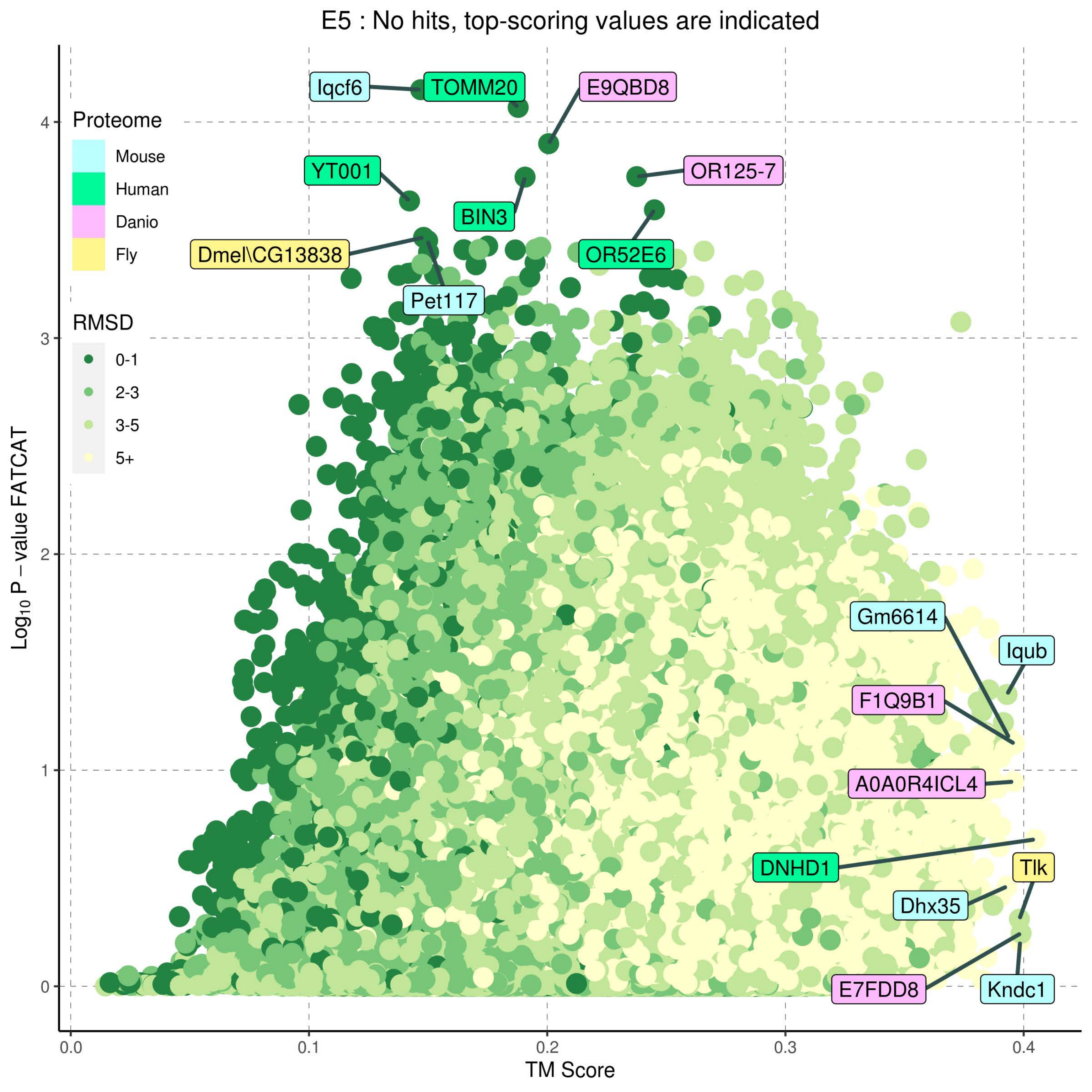
E5 : No hits, top-scoring values are indicated
Proteome
Mouse
Human
Danio
Fly
RMSD
0-1
2-3
3-5
5+
Iqcf6
TOMM20
E9QBD8
YT001
OR125-7
BIN3
Dmel\CG13838
OR52E6
Pet117
Gm6614
Iqub
F1Q9B1
A0A0R4ICL4
DNHD1
Tlk
Dhx35
E7FDD8
Kndc1
Log10 P – value FATCAT
TM Score
0.0
0.1
0.2
0.3
0.4
0
1
2
3
4

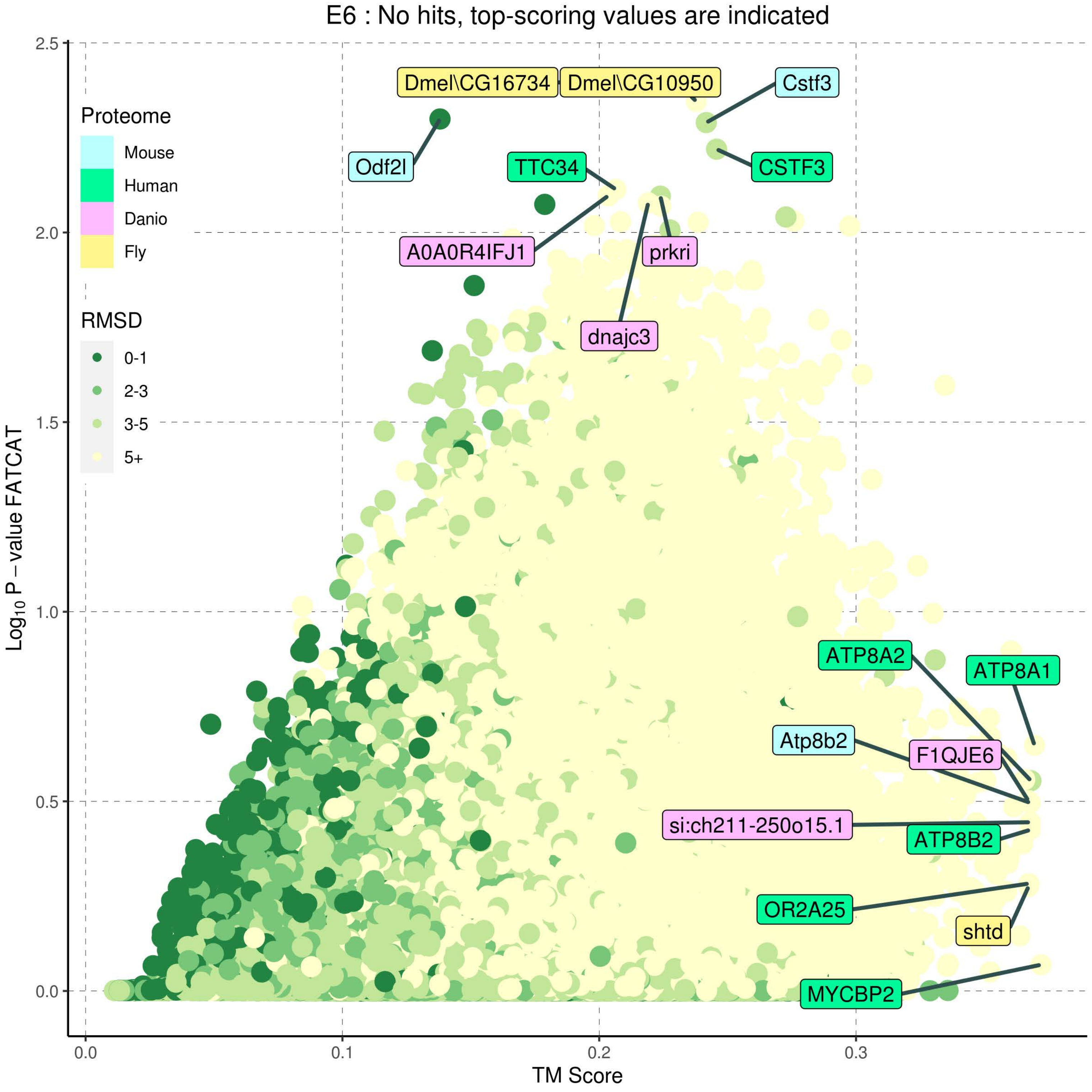
E6 : No hits, top-scoring values are indicated
Proteome
Mouse
Human
Danio
Fly
RMSD
0-1
2-3
3-5
5+
Dmel\CG16734
Dmel\CG10950
Cstf3
Odf2l
TTC34
CSTF3
A0A0R4IFJ1
prkri
dnajc3
ATP8A2
ATP8A1
Atp8b2
F1QJE6
si:ch211-250o15.1
ATP8B2
OR2A25
shtd
MYCBP2
$Log_{10}$ P – value FATCAT
TM Score
0.0
0.5
1.0
1.5
2.0
2.5
0.0
0.1
0.2
0.3

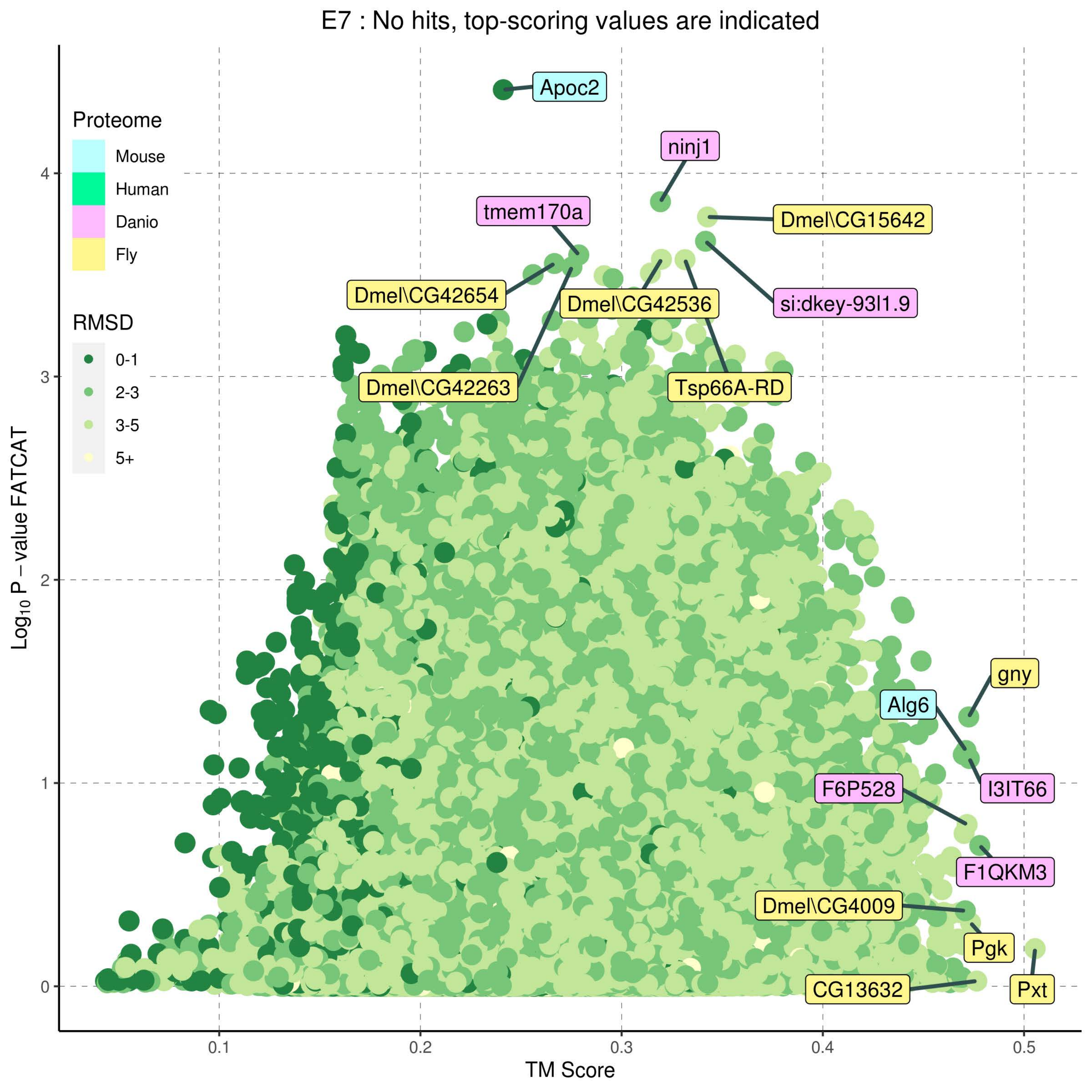
E7 : No hits, top-scoring values are indicated
Proteome
Mouse
Human
Danio
Fly
RMSD
0-1
2-3
3-5
5+
Apoc2
ninj1
tmem170a
Dmel\CG15642
Dmel\CG42654
Dmel\CG42536
si:dkey-93l1.9
Dmel\CG42263
Tsp66A-RD
gny
Alg6
I3IT66
F6P528
F1QKM3
Dmel\CG4009
Pgk
CG13632
Pxt
Log10 P − value FATCAT
TM Score
0
1
2
3
4
0.1
0.2
0.3
0.4
0.5

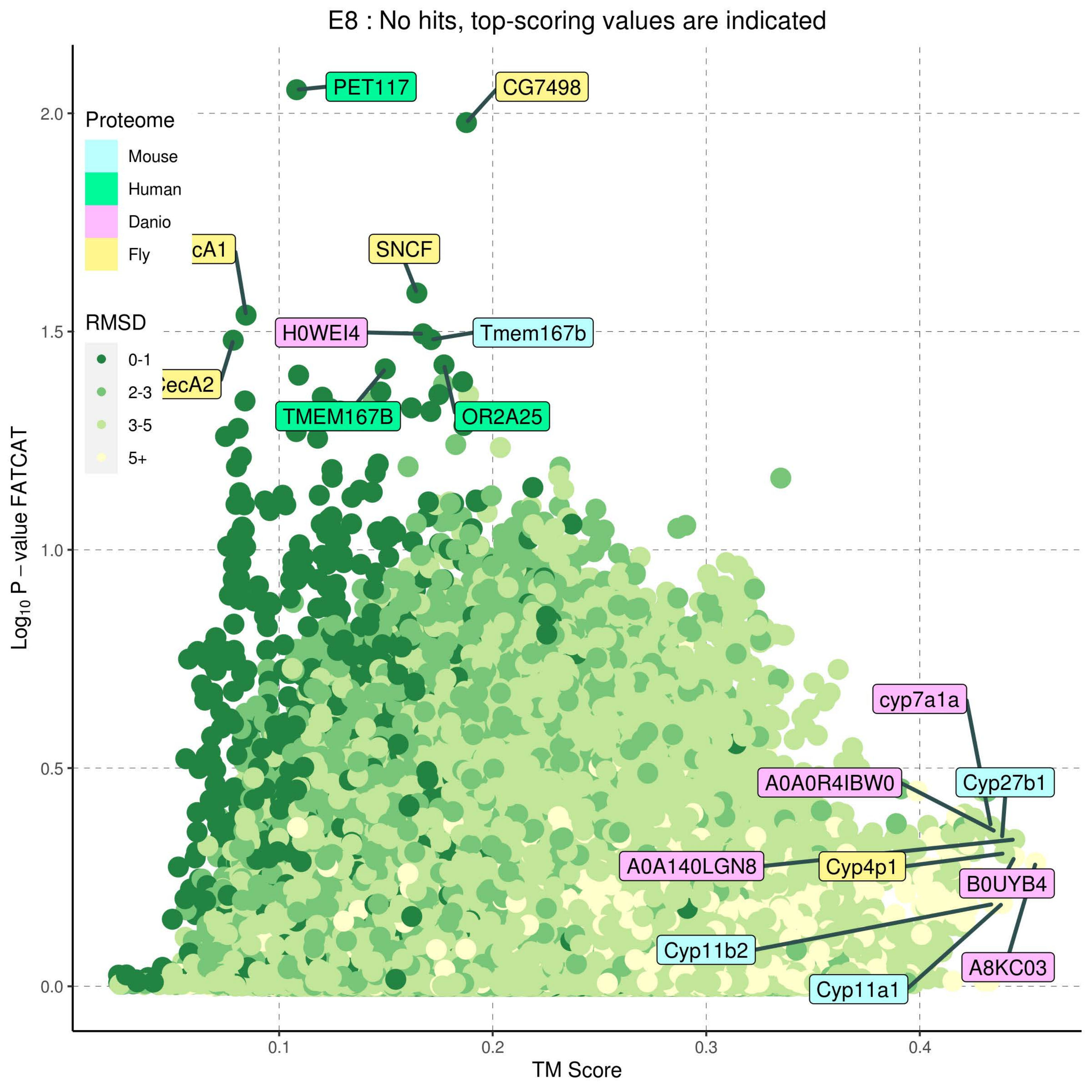
E8 : No hits, top-scoring values are indicated
Proteome
Mouse
Human
Danio
Fly
RMSD
0-1
2-3
3-5
5+
PET117
CG7498
cA1
SNCF
H0WEI4
Tmem167b
ecA2
TMEM167B
OR2A25
cyp7a1a
A0A0R4IBW0
Cyp27b1
A0A140LGN8
Cyp4p1
B0UYB4
Cyp11b2
A8KC03
Cyp11a1
Log10 P − value FATCAT
TM Score
2.0
1.5
1.0
0.5
0.0
0.1
0.2
0.3
0.4

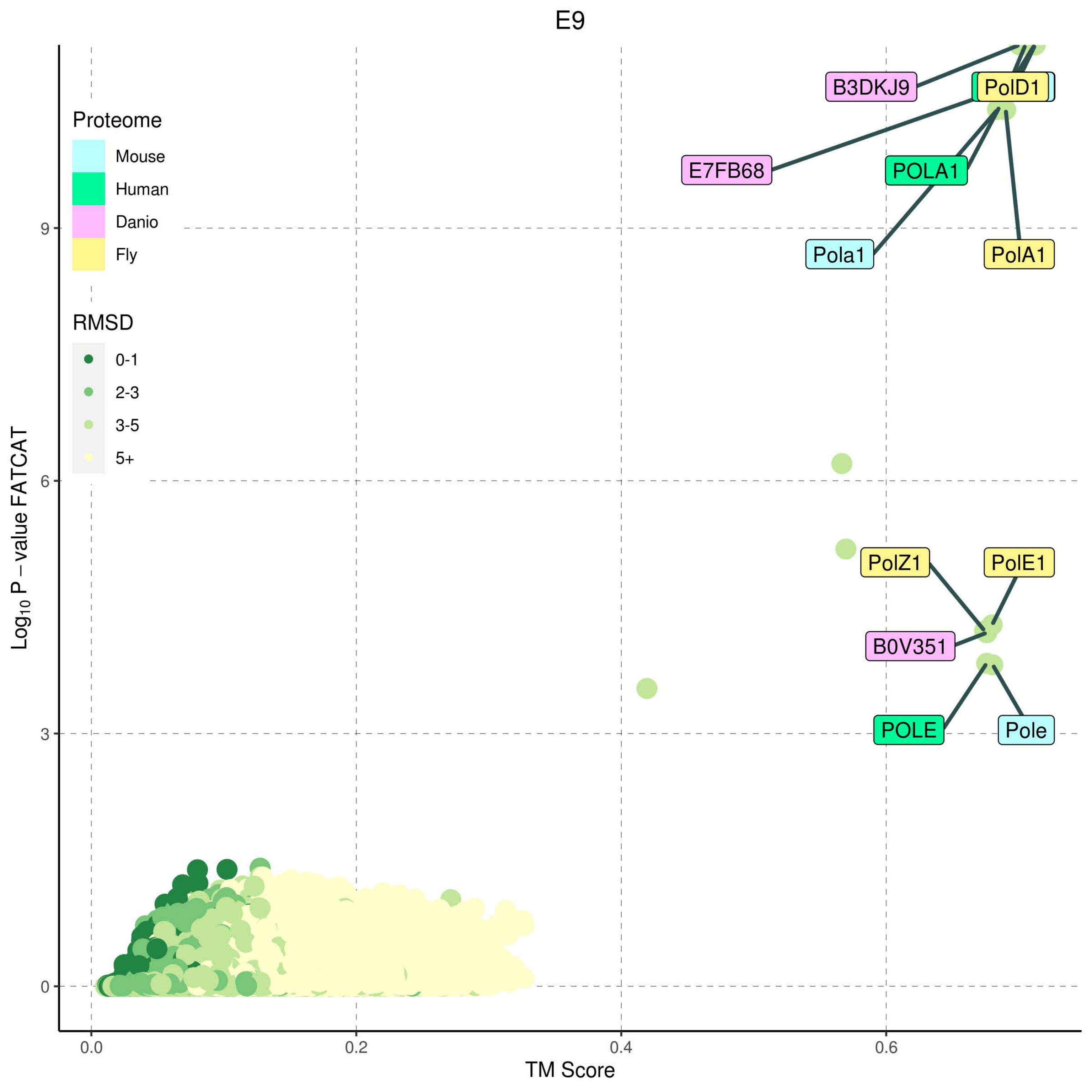

E9
Proteome
Mouse
Human
Danio
Fly
RMSD
0-1
2-3
3-5
5+
B3DKJ9
PolD1
E7FB68
POLA1
Pola1
PolA1
PolZ1
PolE1
B0V351
POLE
Pole
$Log_{10}$ P − value FATCAT
TM Score
0
3
6
9
0.0
0.2
0.4
0.6

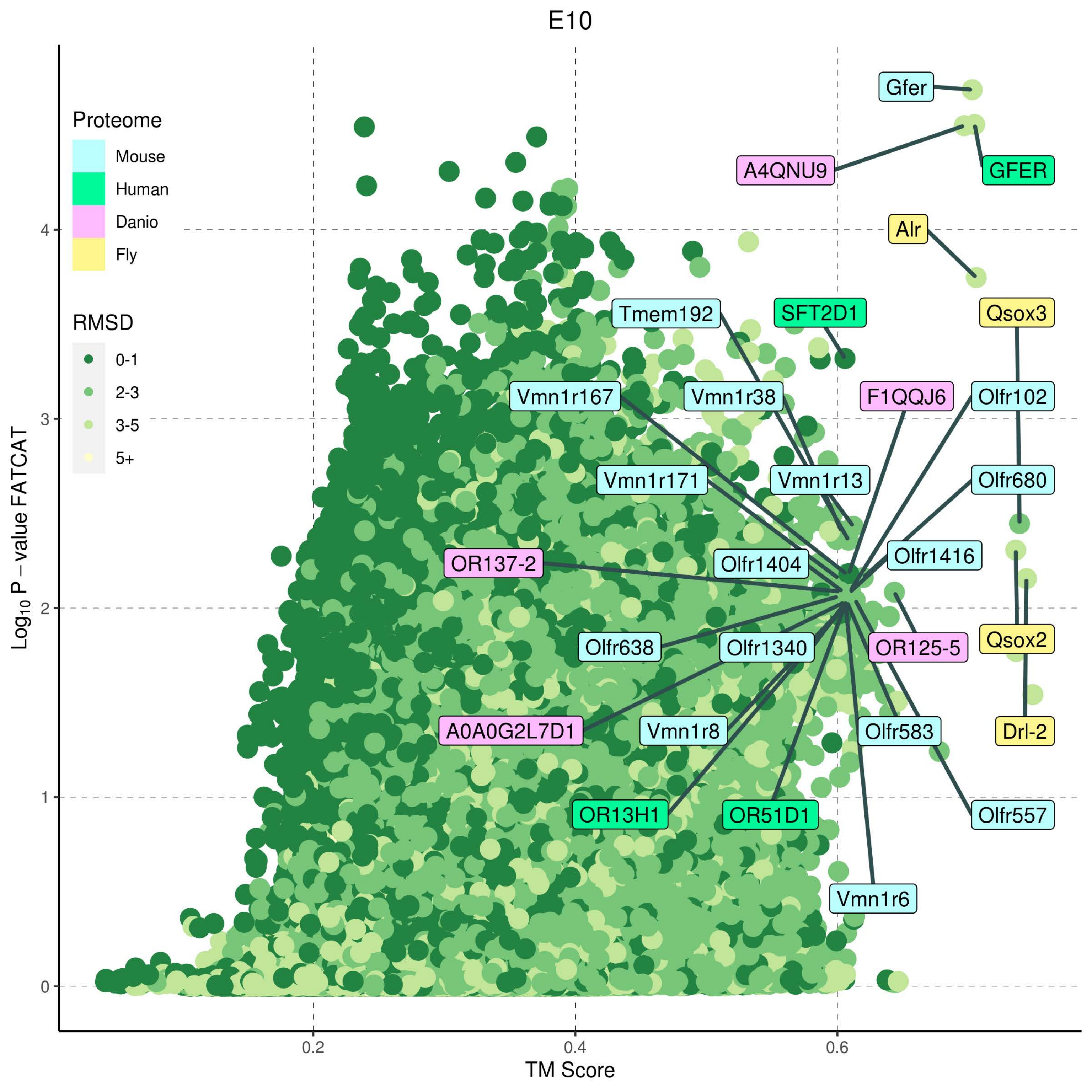

E10
Proteome
Mouse
Human
Danio
Fly
RMSD
0-1
2-3
3-5
5+
Log10 P − value FATCAT
TM Score
0.2
0.4
0.6
0
1
2
3
4
Gfer
A4QNU9
GFER
Alr
Tmem192
SFT2D1
Qsox3
Vmn1r167
Vmn1r38
F1QQJ6
Olfr102
Vmn1r171
Vmn1r13
Olfr680
OR137-2
Olfr1404
Olfr1416
Olfr638
Olfr1340
OR125-5
Qsox2
A0A0G2L7D1
Vmn1r8
Olfr583
Drl-2
OR13H1
OR51D1
Olfr557
Vmn1r6

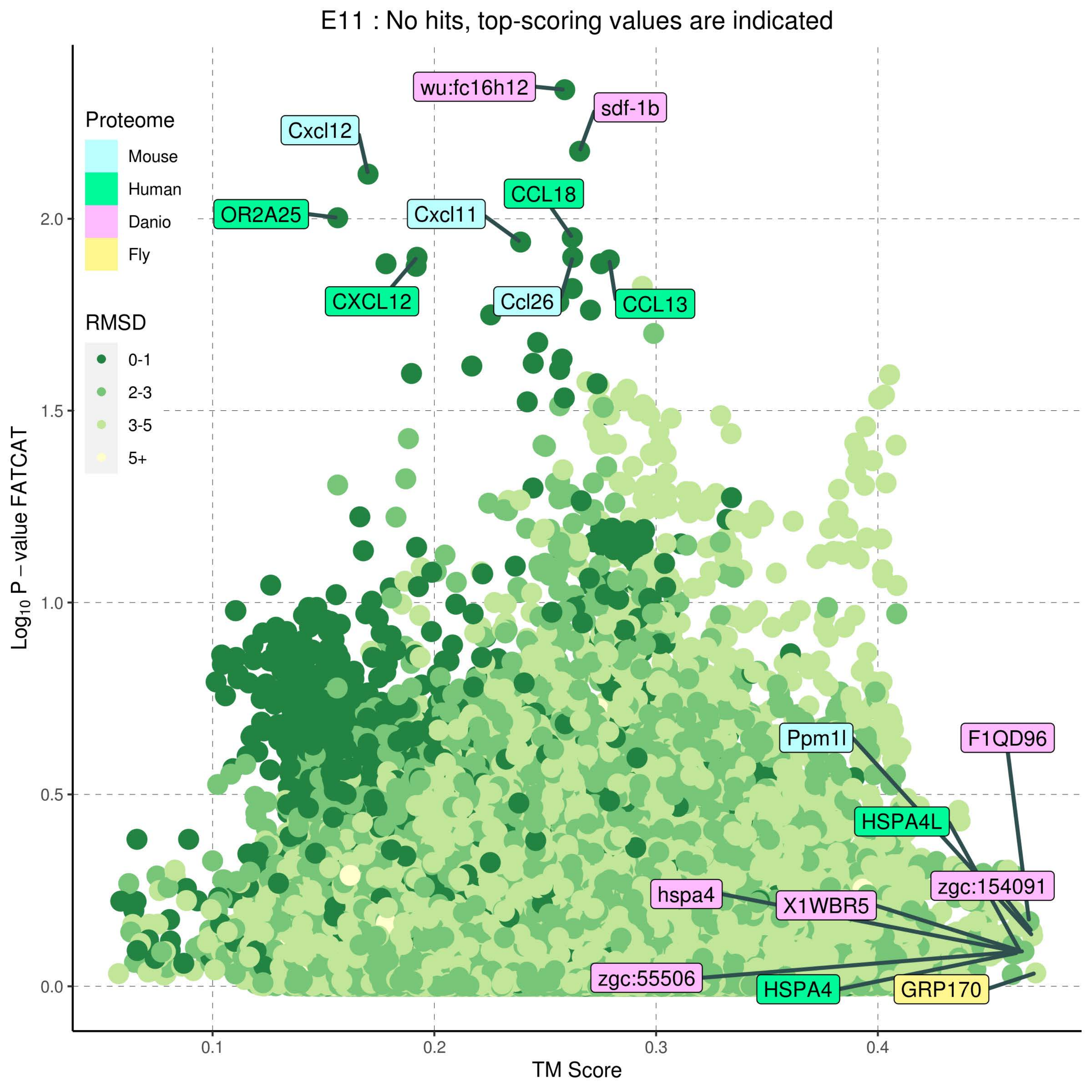

E11 : No hits, top-scoring values are indicated
Proteome
Mouse
Human
Danio
Fly
RMSD
0-1
2-3
3-5
5+
wu:fc16h12
sdf-1b
Cxcl12
OR2A25
Cxcl11
CCL18
CXCL12
Ccl26
CCL13
Ppm1l
F1QD96
HSPA4L
zgc:154091
hspa4
X1WBR5
zgc:55506
HSPA4
GRP170
Log10 P − value FATCAT
TM Score
0.0
0.5
1.0
1.5
2.0
0.1
0.2
0.3
0.4

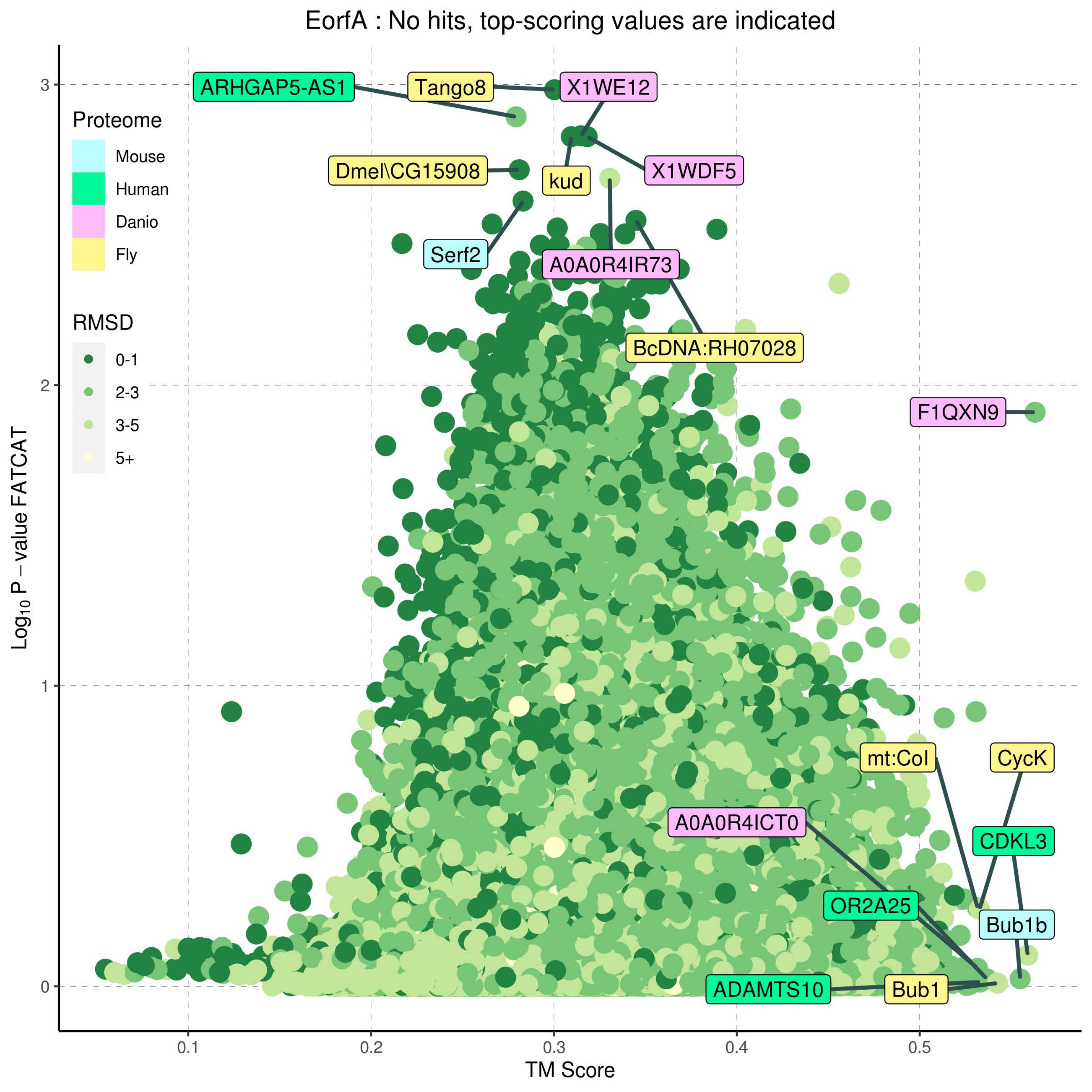

EorfA : No hits, top-scoring values are indicated
Proteome
Mouse
Human
Danio
Fly
RMSD
0-1
2-3
3-5
5+
ARHGAP5-AS1
Tango8
X1WE12
X1WDF5
Dmel\CG15908
kud
Serf2
A0A0R4IR73
BcDNA:RH07028
F1QXN9
mt:CoI
CycK
A0A0R4ICT0
CDKL3
OR2A25
Bub1b
ADAMTS10
Bub1
Log10 P − value FATCAT
TM Score
0.1
0.2
0.3
0.4
0.5
0
1
2
3

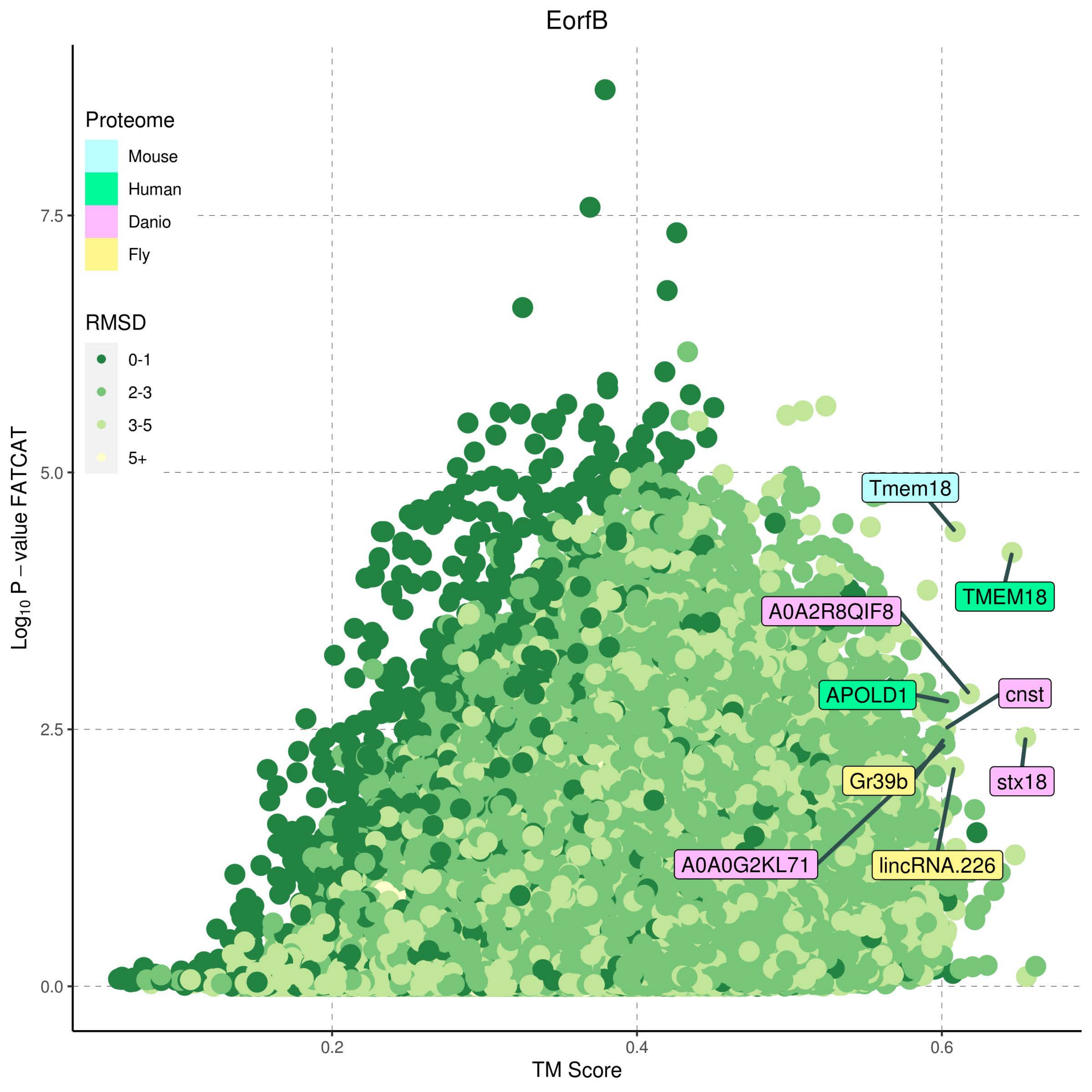
EorfB
Proteome
Mouse
Human
Danio
Fly
RMSD
0-1
2-3
3-5
5+
Tmem18
TMEM18
A0A2R8QIF8
APOLD1
cnst
Gr39b
stx18
A0A0G2KL71
lincRNA.226
7.5
5.0
2.5
0.0
0.2
0.4
0.6
$\text{Log}_{10}$ P − value FATCAT
TM Score

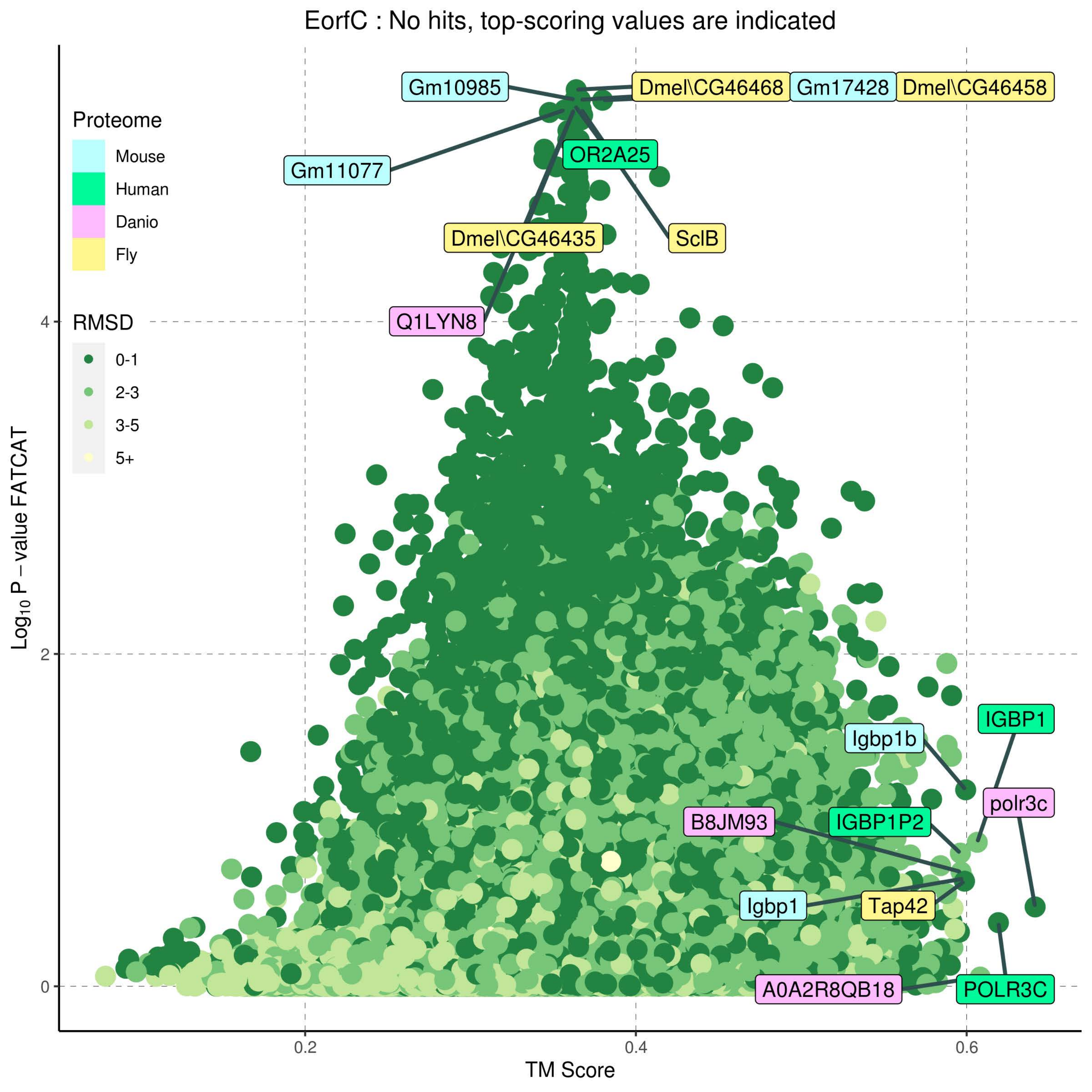
EorfC : No hits, top-scoring values are indicated
Proteome
Mouse
Human
Danio
Fly
RMSD
0-1
2-3
3-5
5+
Gm10985
Dmel\CG46468
Gm17428
Dmel\CG46458
OR2A25
Gm11077
Dmel\CG46435
SclB
Q1LYN8
IGBP1
Igbp1b
polr3c
B8JM93
IGBP1P2
Igbp1
Tap42
A0A2R8QB18
POLR3C
Log10 P − value FATCAT
4
2
0
0.2
0.4
0.6
TM Score

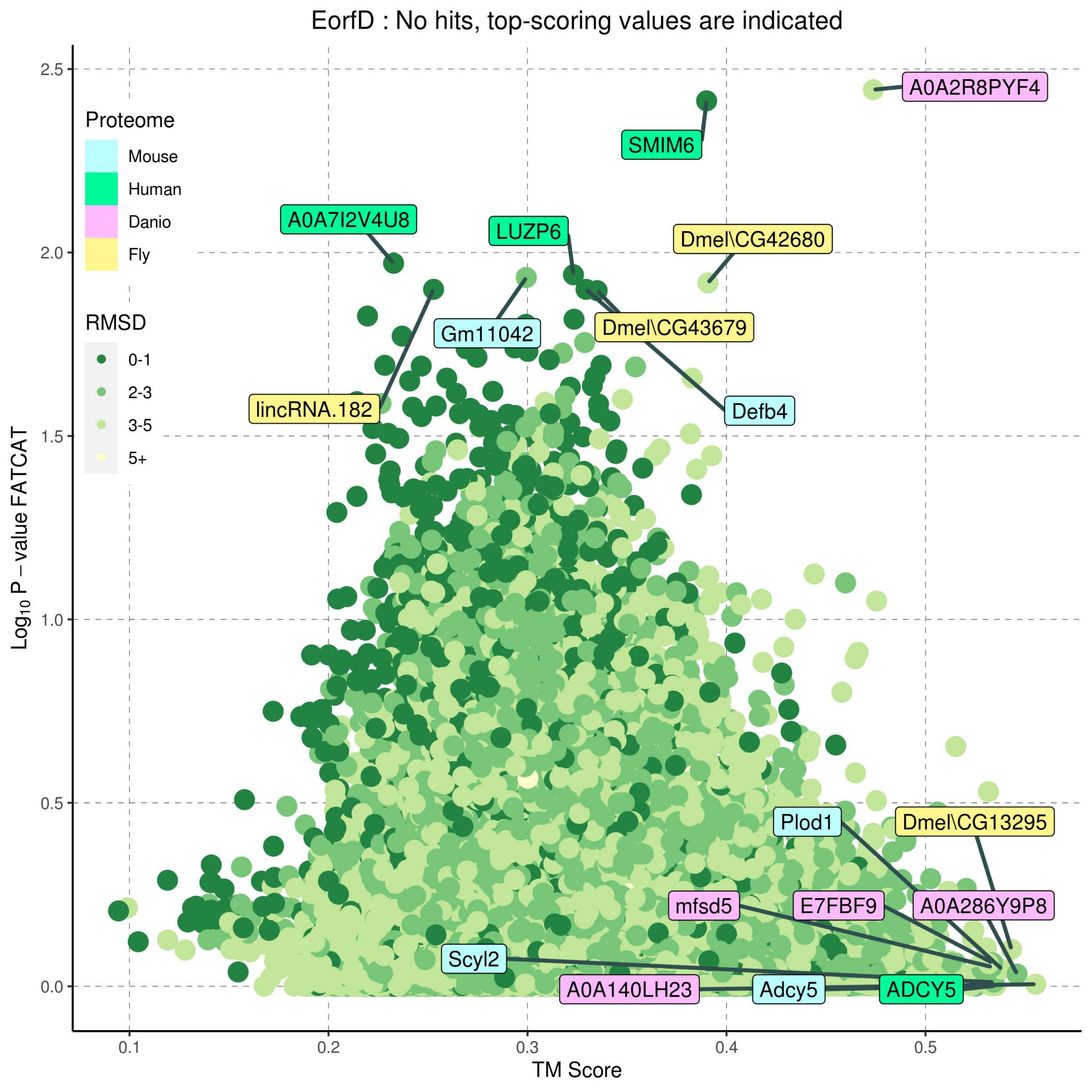
EorfD : No hits, top-scoring values are indicated
Proteome
Mouse
Human
Danio
Fly
RMSD
0-1
2-3
3-5
5+
A0A2R8PYF4
SMIM6
A0A7I2V4U8
LUZP6
Dmel\CG42680
Gm11042
Dmel\CG43679
lincRNA.182
Defb4
Plod1
Dmel\CG13295
mfsd5
E7FBF9
A0A286Y9P8
Scyl2
A0A140LH23
Adcy5
ADCY5
Log10 P – value FATCAT
TM Score
2.5
2.0
1.5
1.0
0.5
0.0
0.1
0.2
0.3
0.4
0.5

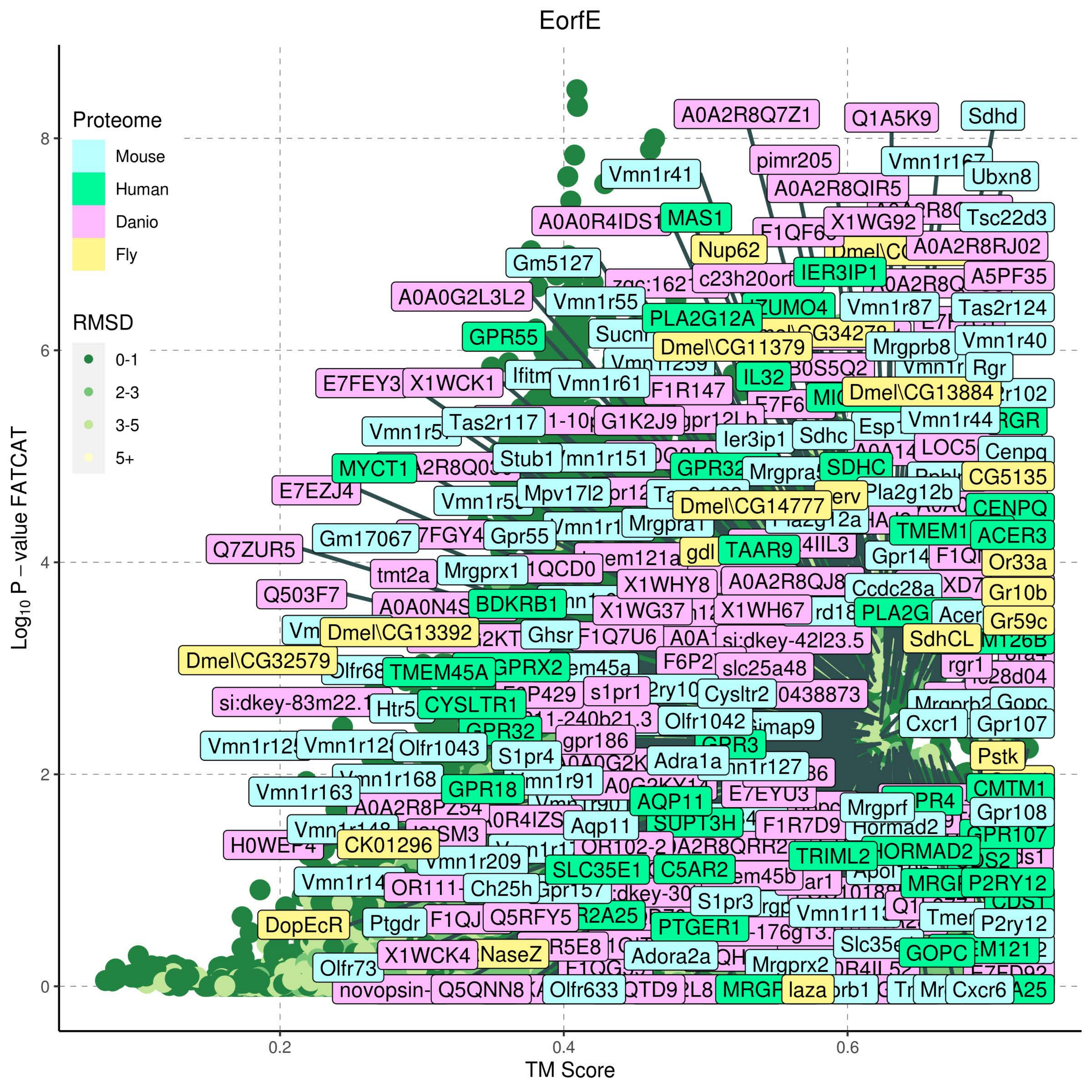

EorfE
Proteome
Mouse
Human
Danio
Fly
RMSD
0-1
2-3
3-5
5+
Log10 P − value FATCAT
TM Score
0
2
4
6
8
0.2
0.4
0.6
A0A2R8Q7Z1
Q1A5K9
Sdhd
pimr205
Vmn1r41
Ubxn8
A0A2R8QIR5
A0A0R4IDS1
MAS1
X1WG92
Tsc22d3
Nup62
A0A2R8RJ02
Gm5127
IER3IP1
A5PF35
A0A0G2L3L2
Vmn1r55
Vmn1r87
Tas2r124
PLA2G12A
GPR55
Dmel\CG11379
Mrgprb8
Vmn1r40
Rgr
IL32
E7FEY3
X1WCK1
Vmn1r61
F1R147
Dmel\CG13884
Tas2r117
G1K2J9
Vmn1r44
Ier3ip1
Sdhc
Stub1
MYCT1
SDHC
Cenpq
CG5135
E7EZJ4
Mpv17l2
Pla2g12b
Dmel\CG14777
CENPQ
Gm17067
Gpr55
ACER3
Q7ZUR5
TAAR9
Or33a
Mrgprx1
tmt2a
X1WHY8
Ccdc28a
Gr10b
Q503F7
BDKRB1
X1WG37
X1WH67
Gr59c
Dmel\CG13392
Ghsr
F1Q7U6
si:dkey-42l23.5
SdhCL
Dmel\CG32579
TMEM45A
GPRX2
slc25a48
rgr1
s1pr1
Cysltr2
Htr5
CYSLTR1
Olfr1042
Cxcr1
Gpr107
gpr186
Olfr1043
S1pr4
Adra1a
Pstk
Vmn1r168
Vmn1r163
GPR18
AQP11
E7EYU3
CMTM1
Mrgprf
Gpr108
Aqp11
SUPT3H
F1R7D9
Hormad2
CK01296
TRIML2
HORMAD2
Vmn1r209
SLC35E1
C5AR2
MRGPRX2
P2RY12
Ch25h
S1pr3
DopEcR
Ptgdr
Q5RFY5
PTGER1
P2ry12
X1WCK4
NaseZ
Adora2a
Mrgprx2
GOPC
Olfr73
Q5QNN8
Olfr633
laza
Cxcr6

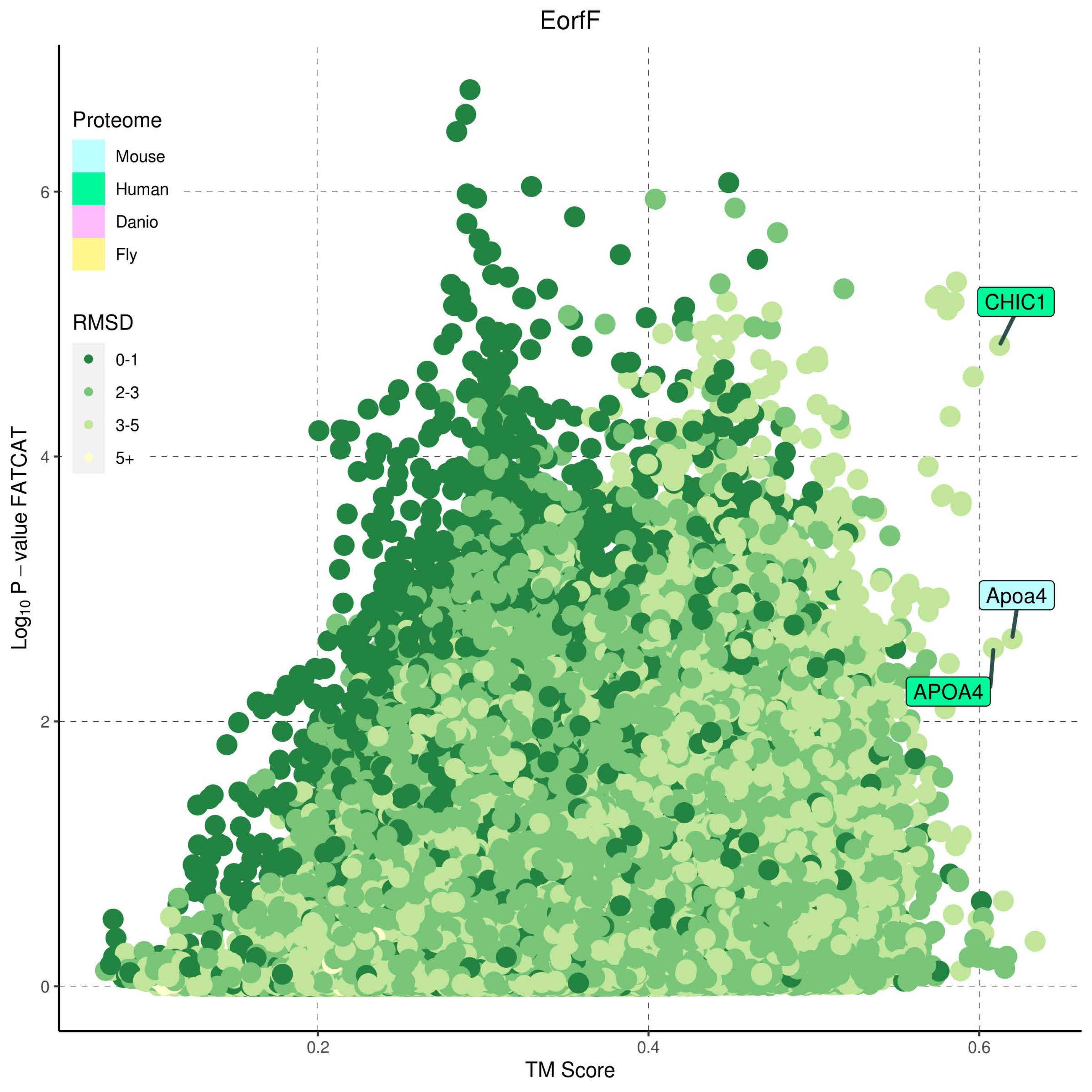

EorfF
Proteome
Mouse
Human
Danio
Fly
RMSD
0-1
2-3
3-5
5+
CHIC1
Apoa4
APOA4
Log10 P − value FATCAT
TM Score
0
2
4
6
0.2
0.4
0.6

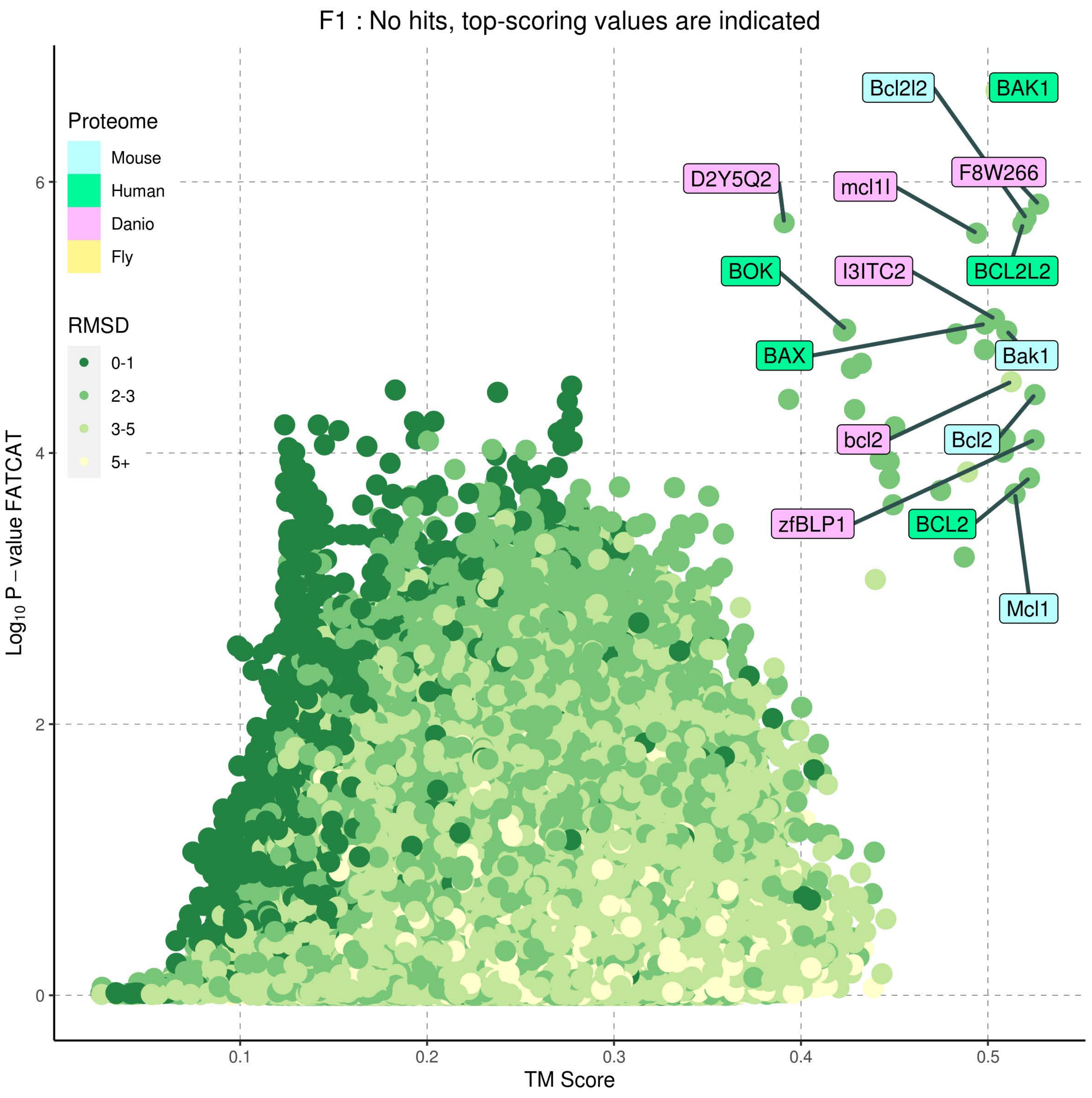
F1 : No hits, top-scoring values are indicated
Proteome
Mouse
Human
Danio
Fly
RMSD
0-1
2-3
3-5
5+
Bcl2l2
BAK1
D2Y5Q2
mcl1l
F8W266
BOK
I3ITC2
BCL2L2
BAX
Bak1
bcl2
Bcl2
zfBLP1
BCL2
Mcl1
0
2
4
6
0.1
0.2
0.3
0.4
0.5
$Log_{10}$ P – value FATCAT
TM Score

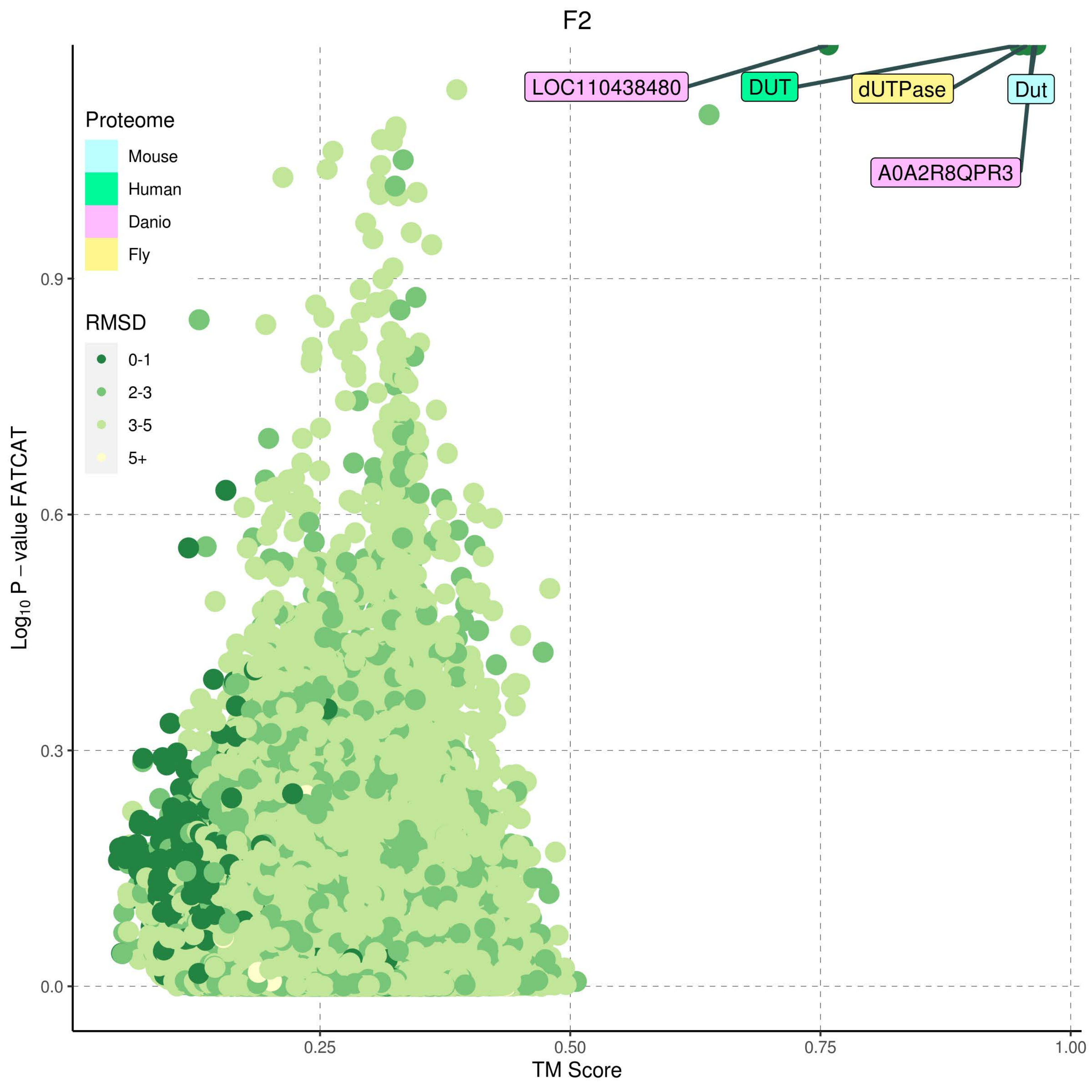
F2
LOC110438480
DUT
dUTPase
Dut
A0A2R8QPR3
Proteome
Mouse
Human
Danio
Fly
RMSD
0-1
2-3
3-5
5+
Log10 P – value FATCAT
0.9
0.6
0.3
0.0
0.25
0.50
0.75
1.00
TM Score

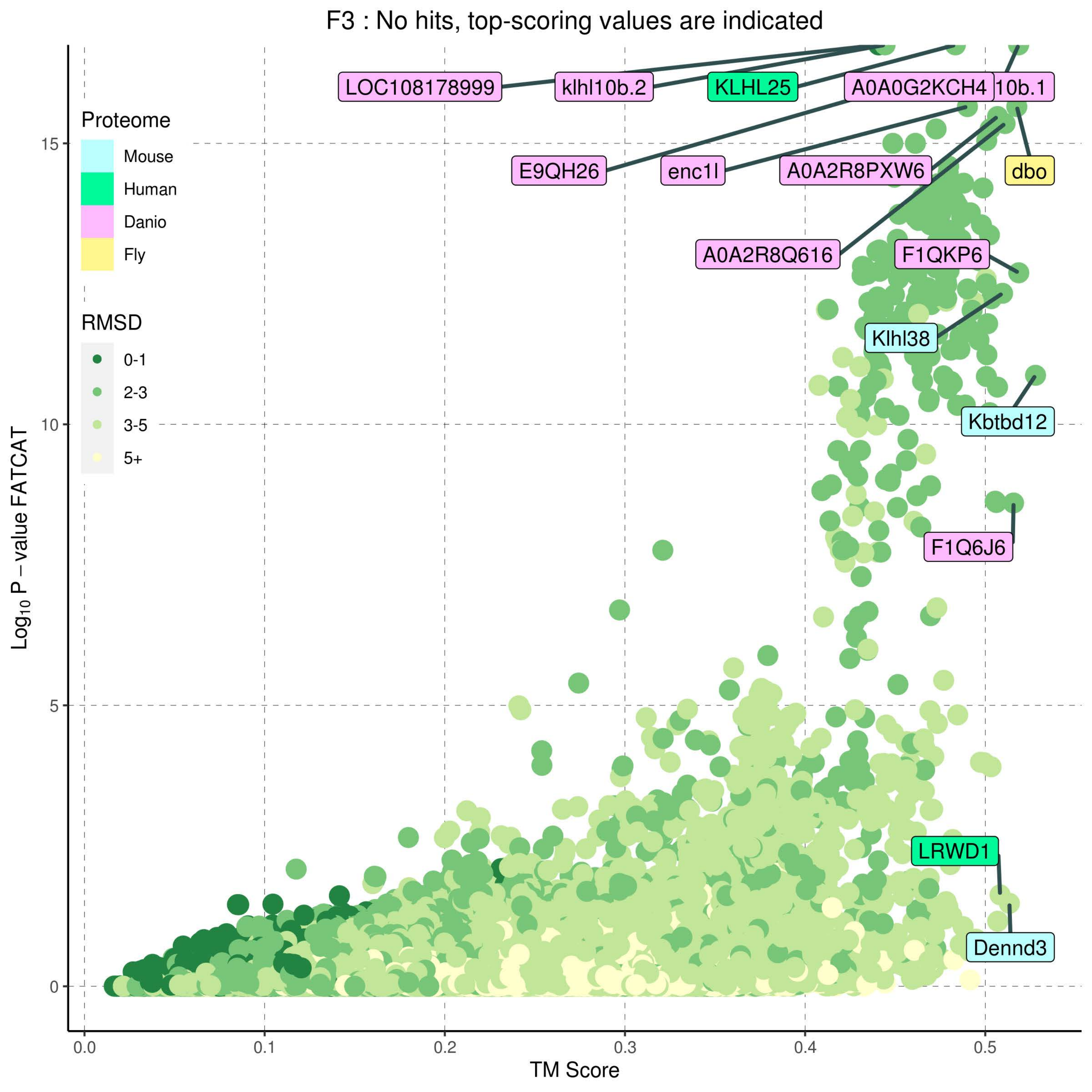
F3 : No hits, top-scoring values are indicated
Proteome
Mouse
Human
Danio
Fly
RMSD
0-1
2-3
3-5
5+
LOC108178999
klhl10b.2
KLHL25
A0A0G2KCH4
10b.1
E9QH26
enc1l
A0A2R8PXW6
dbo
A0A2R8Q616
F1QKP6
Klhl38
Kbtbd12
F1Q6J6
LRWD1
Dennd3
Log10 P − value FATCAT
TM Score
0
5
10
15
0.0
0.1
0.2
0.3
0.4
0.5

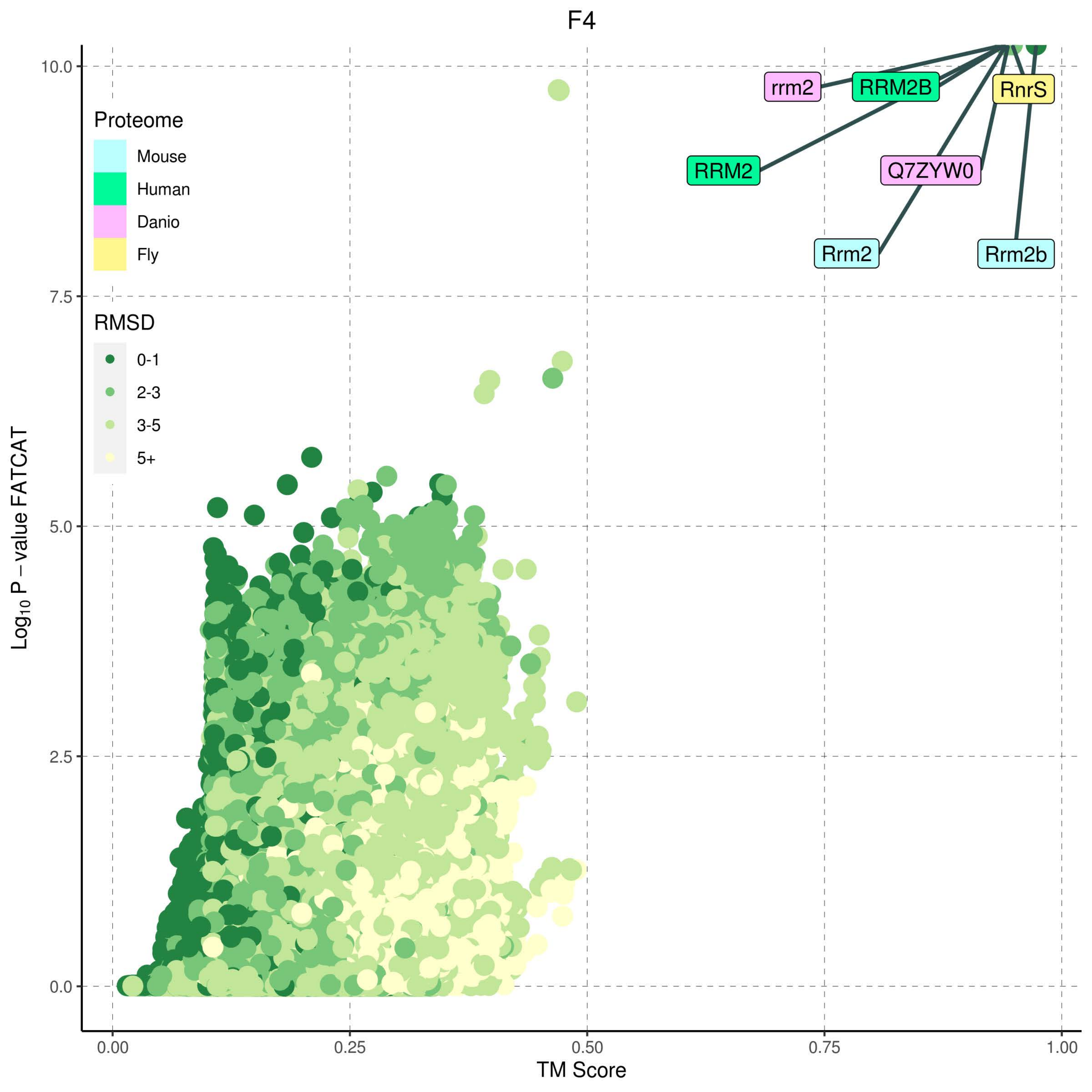

F4
Proteome
Mouse
Human
Danio
Fly
RMSD
0-1
2-3
3-5
5+
rrm2
RRM2B
RnrS
RRM2
Q7ZYW0
Rrm2
Rrm2b
Log10 P − value FATCAT
TM Score
10.0
7.5
5.0
2.5
0.0
0.00
0.25
0.50
0.75
1.00

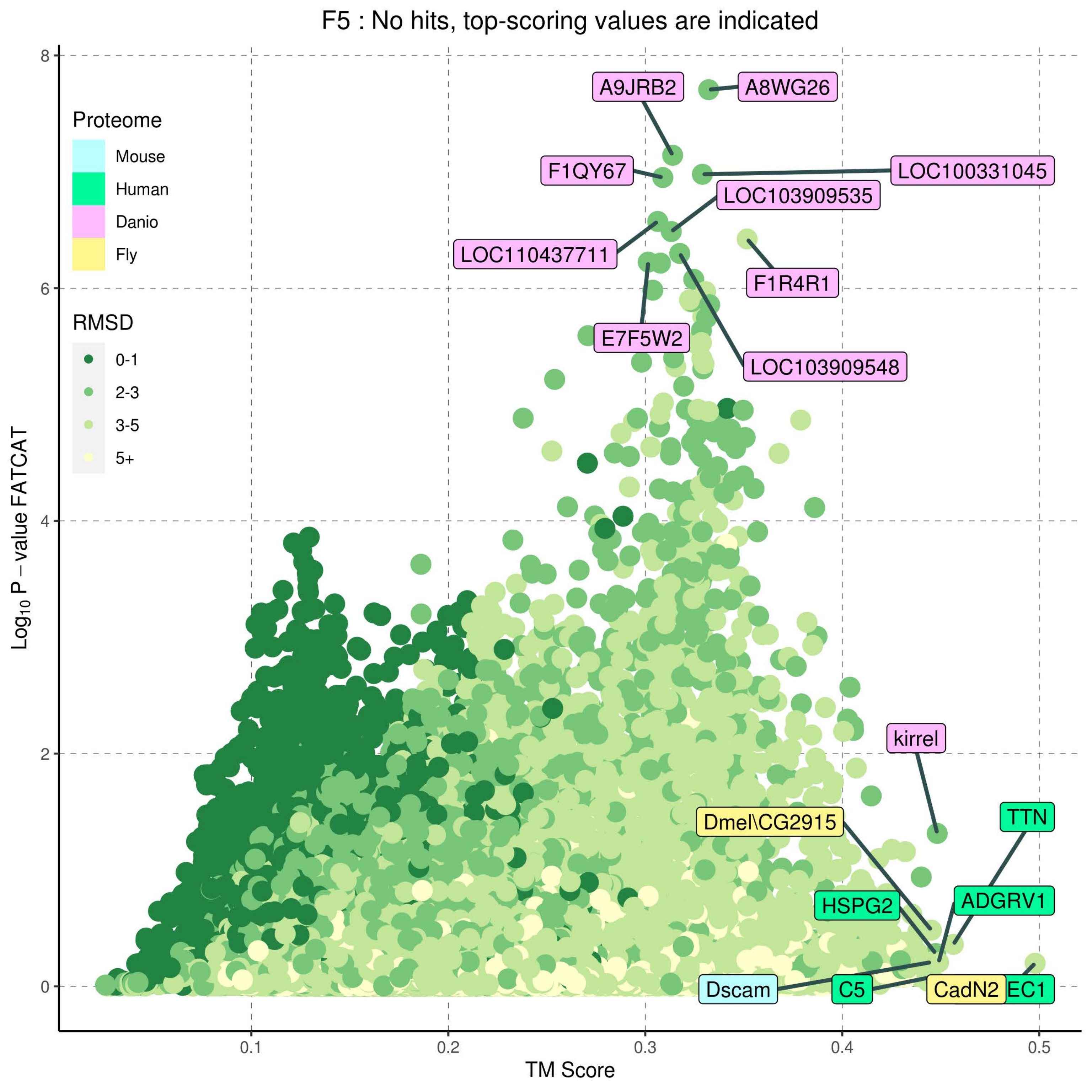

F5 : No hits, top-scoring values are indicated
Proteome
Mouse
Human
Danio
Fly
RMSD
0-1
2-3
3-5
5+
A9JRB2
A8WG26
F1QY67
LOC100331045
LOC103909535
LOC110437711
F1R4R1
E7F5W2
LOC103909548
kirrel
Dmel\CG2915
TTN
HSPG2
ADGRV1
Dscam
C5
CadN2
EC1
Log10 P – value FATCAT
TM Score
0
2
4
6
8
0.1
0.2
0.3
0.4
0.5

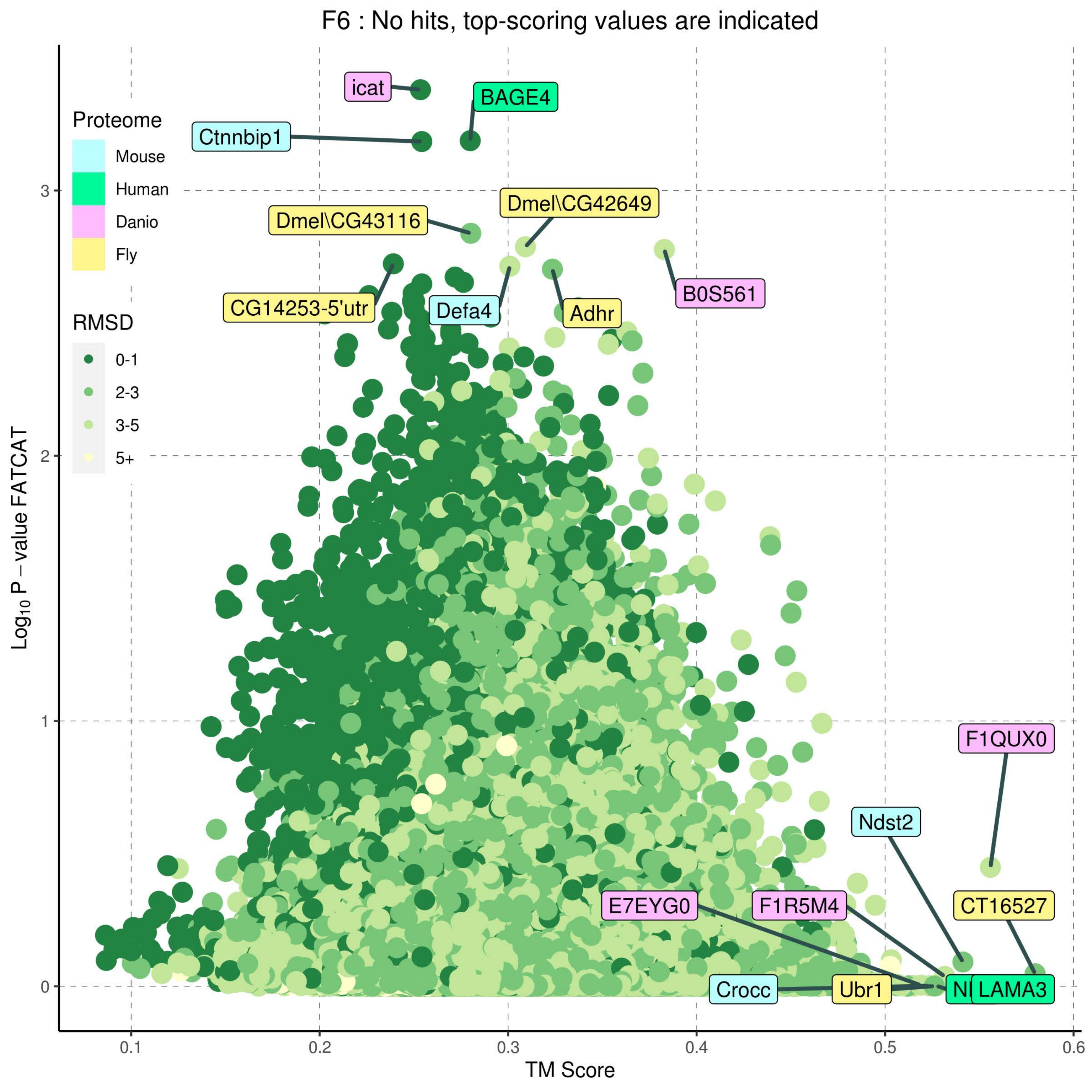

F6 : No hits, top-scoring values are indicated
Proteome
Mouse
Human
Danio
Fly
RMSD
0-1
2-3
3-5
5+
icat
BAGE4
Ctnnbip1
Dmel\CG42649
Dmel\CG43116
B0S561
CG14253-5'utr
Defa4
Adhr
F1QUX0
Ndst2
E7EYG0
F1R5M4
CT16527
Crocc
Ubr1
LAMA3
Log10 P – value FATCAT
TM Score
0
1
2
3
0.1
0.2
0.3
0.4
0.5
0.6

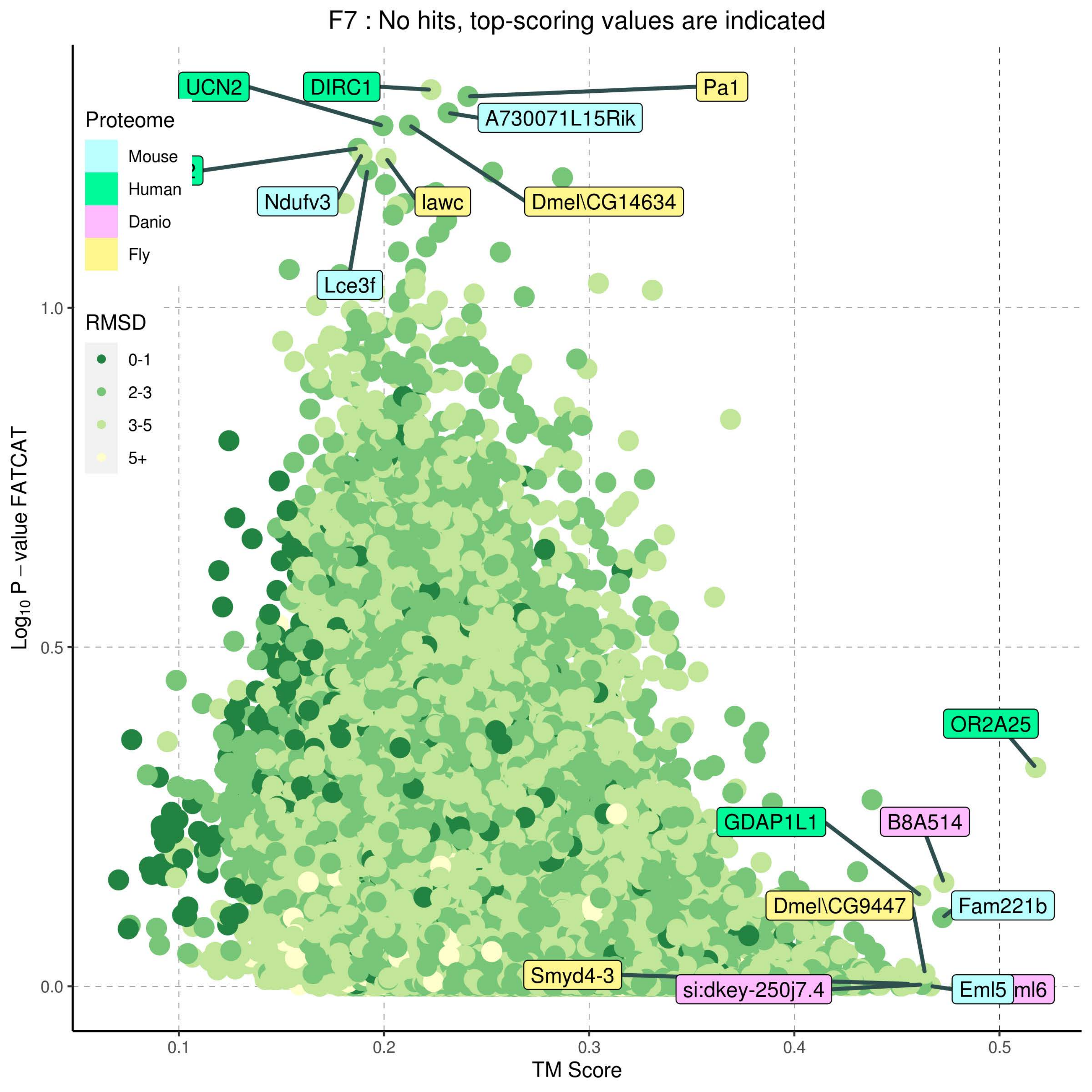

F7 : No hits, top-scoring values are indicated
Proteome
Mouse
Human
Danio
Fly
RMSD
0-1
2-3
3-5
5+
UCN2
DIRC1
Pa1
A730071L15Rik
Ndufv3
lawc
Dmel\CG14634
Lce3f
OR2A25
GDAP1L1
B8A514
Fam221b
Dmel\CG9447
Smyd4-3
si:dkey-250j7.4
Eml5
1.0
0.5
0.0
0.1
0.2
0.3
0.4
0.5
TM Score
Log10 P – value FATCAT

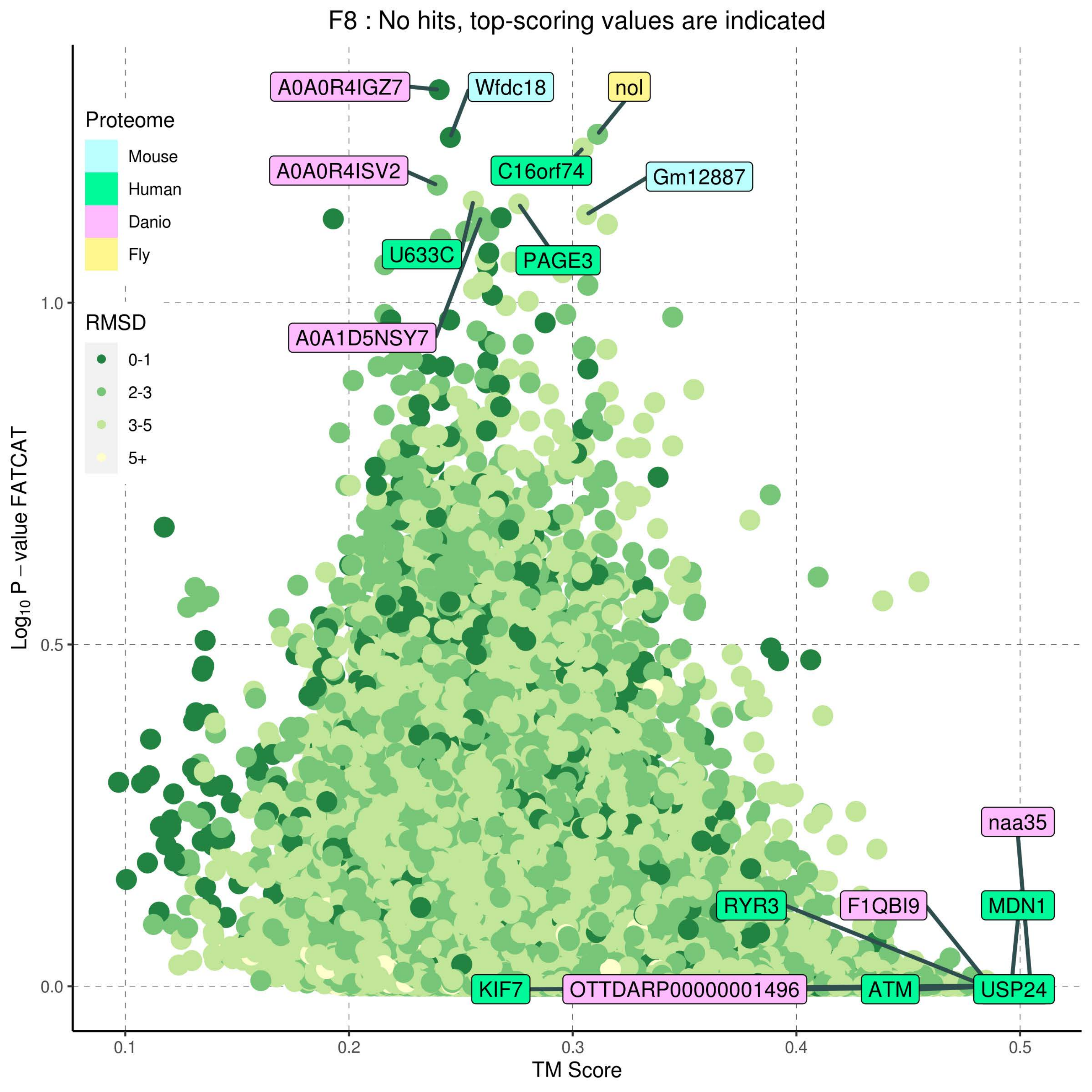

F8 : No hits, top-scoring values are indicated
Proteome
Mouse
Human
Danio
Fly
RMSD
0-1
2-3
3-5
5+
A0A0R4IGZ7
Wfdc18
nol
A0A0R4ISV2
C16orf74
Gm12887
U633C
PAGE3
A0A1D5NSY7
naa35
RYR3
F1QBI9
MDN1
KIF7
OTTDARP00000001496
ATM
USP24
Log10 P – value FATCAT
TM Score
1.0
0.5
0.0
0.1
0.2
0.3
0.4
0.5

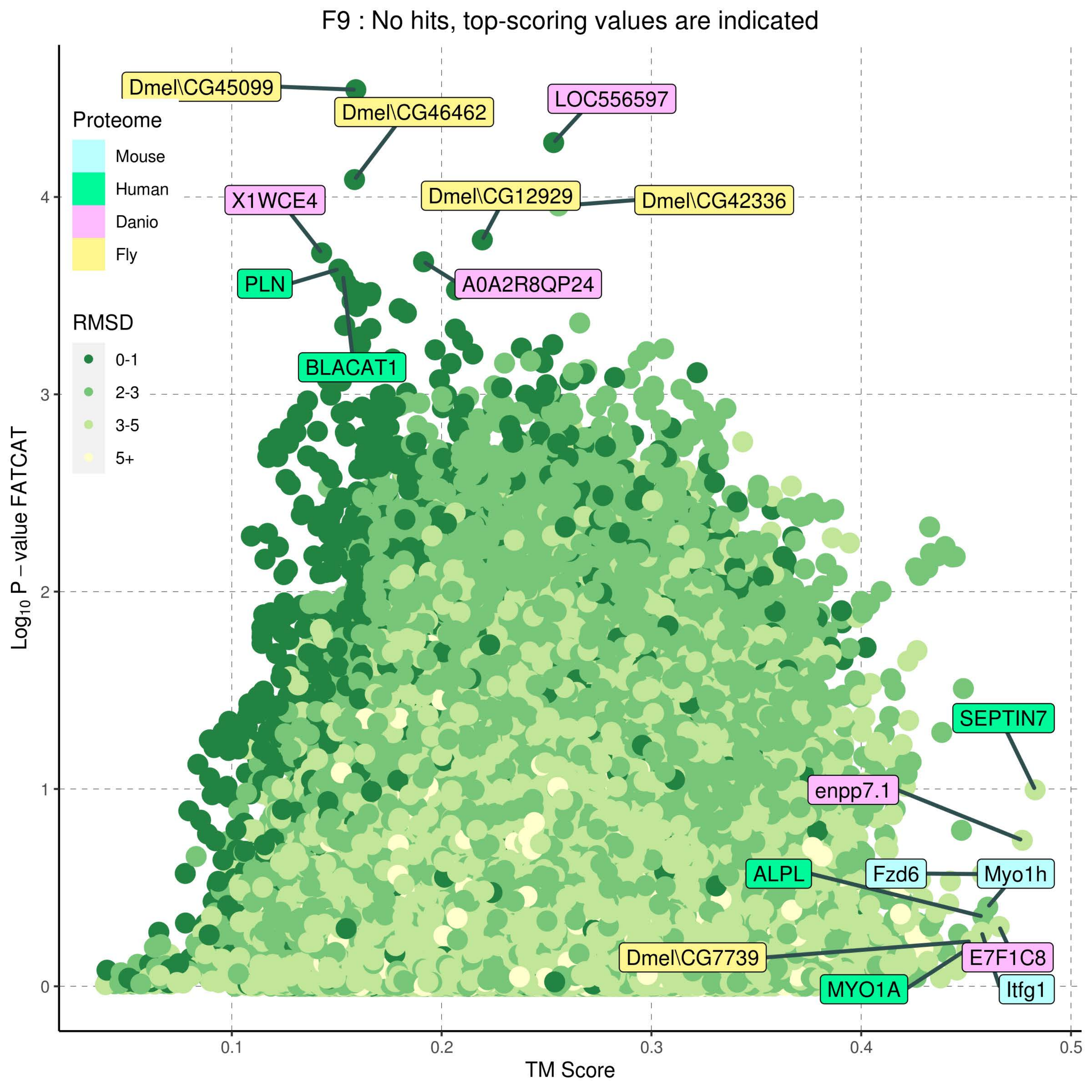

F9 : No hits, top-scoring values are indicated
Proteome
Mouse
Human
Danio
Fly
RMSD
0-1
2-3
3-5
5+
Dmel\CG45099
Dmel\CG46462
LOC556597
X1WCE4
Dmel\CG12929
Dmel\CG42336
PLN
A0A2R8QP24
BLACAT1
SEPTIN7
enpp7.1
ALPL
Fzd6
Myo1h
Dmel\CG7739
E7F1C8
MYO1A
Itfg1
Log10 P – value FATCAT
TM Score
0
1
2
3
4
0.1
0.2
0.3
0.4
0.5

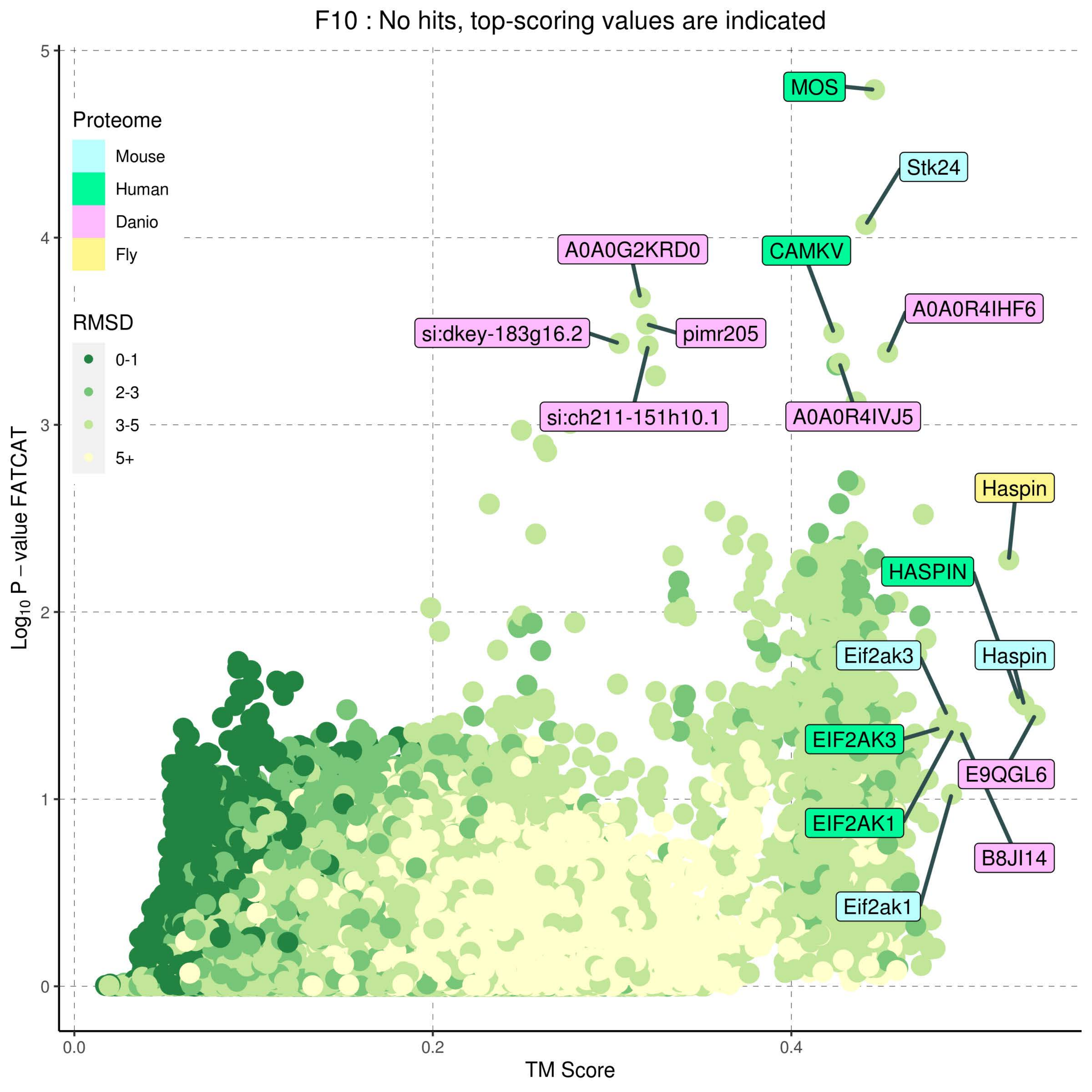

F10 : No hits, top-scoring values are indicated
Proteome
Mouse
Human
Danio
Fly
RMSD
0-1
2-3
3-5
5+
MOS
Stk24
A0A0G2KRD0
CAMKV
A0A0R4IHF6
si:dkey-183g16.2
pimr205
si:ch211-151h10.1
A0A0R4IVJ5
Haspin
HASPIN
Eif2ak3
Haspin
EIF2AK3
E9QGL6
EIF2AK1
B8JI14
Eif2ak1
$Log_{10}$ P – value FATCAT
TM Score
0
1
2
3
4
5
0.0
0.2
0.4

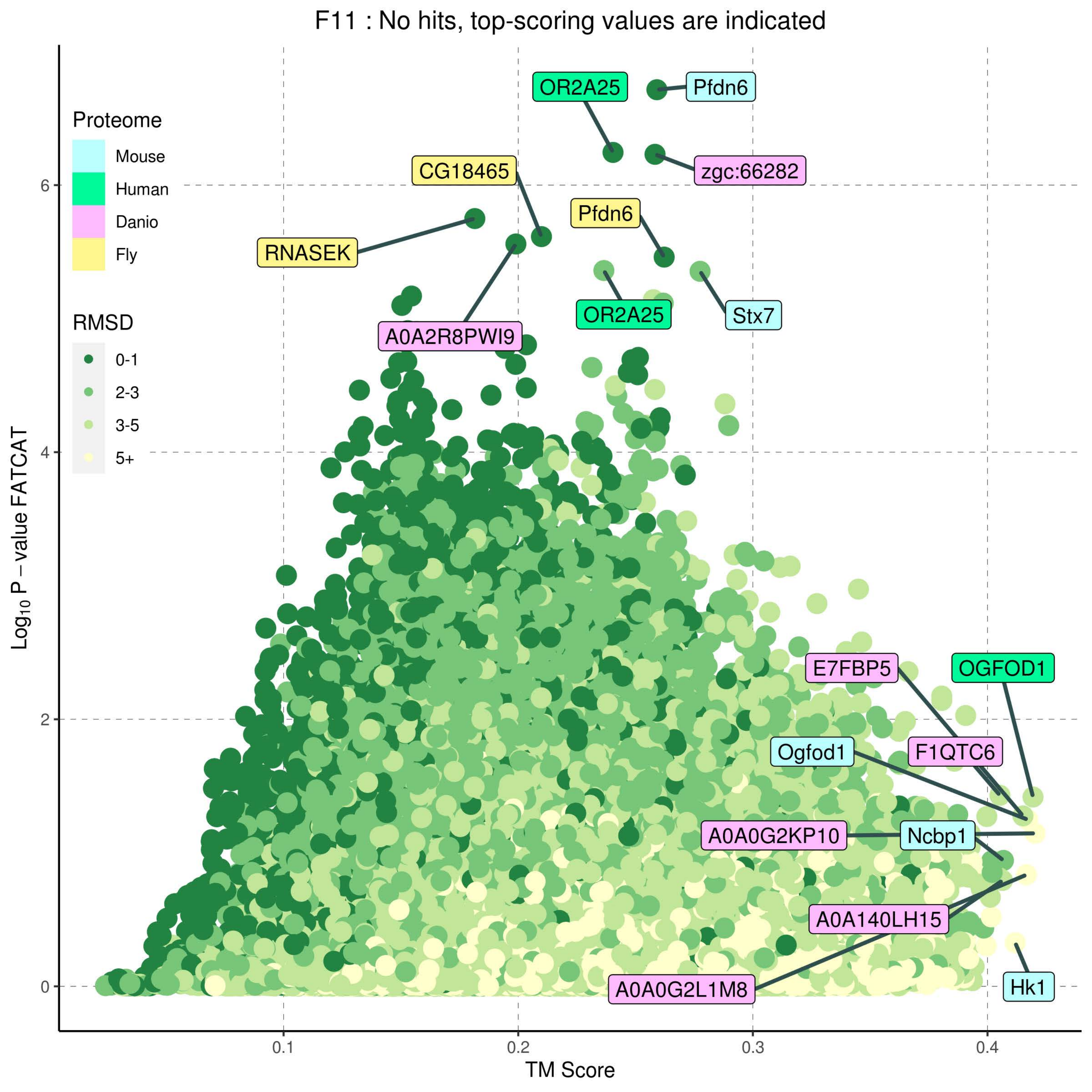

F11 : No hits, top-scoring values are indicated
Proteome
Mouse
Human
Danio
Fly
RMSD
0-1
2-3
3-5
5+
OR2A25
Pfdn6
zgc:66282
CG18465
Pfdn6
RNASEK
A0A2R8PWI9
OR2A25
Stx7
E7FBP5
OGFOD1
Ogfod1
F1QTC6
A0A0G2KP10
Ncbp1
A0A140LH15
A0A0G2L1M8
Hk1
Log10 P − value FATCAT
TM Score
0.1
0.2
0.3
0.4
0
2
4
6

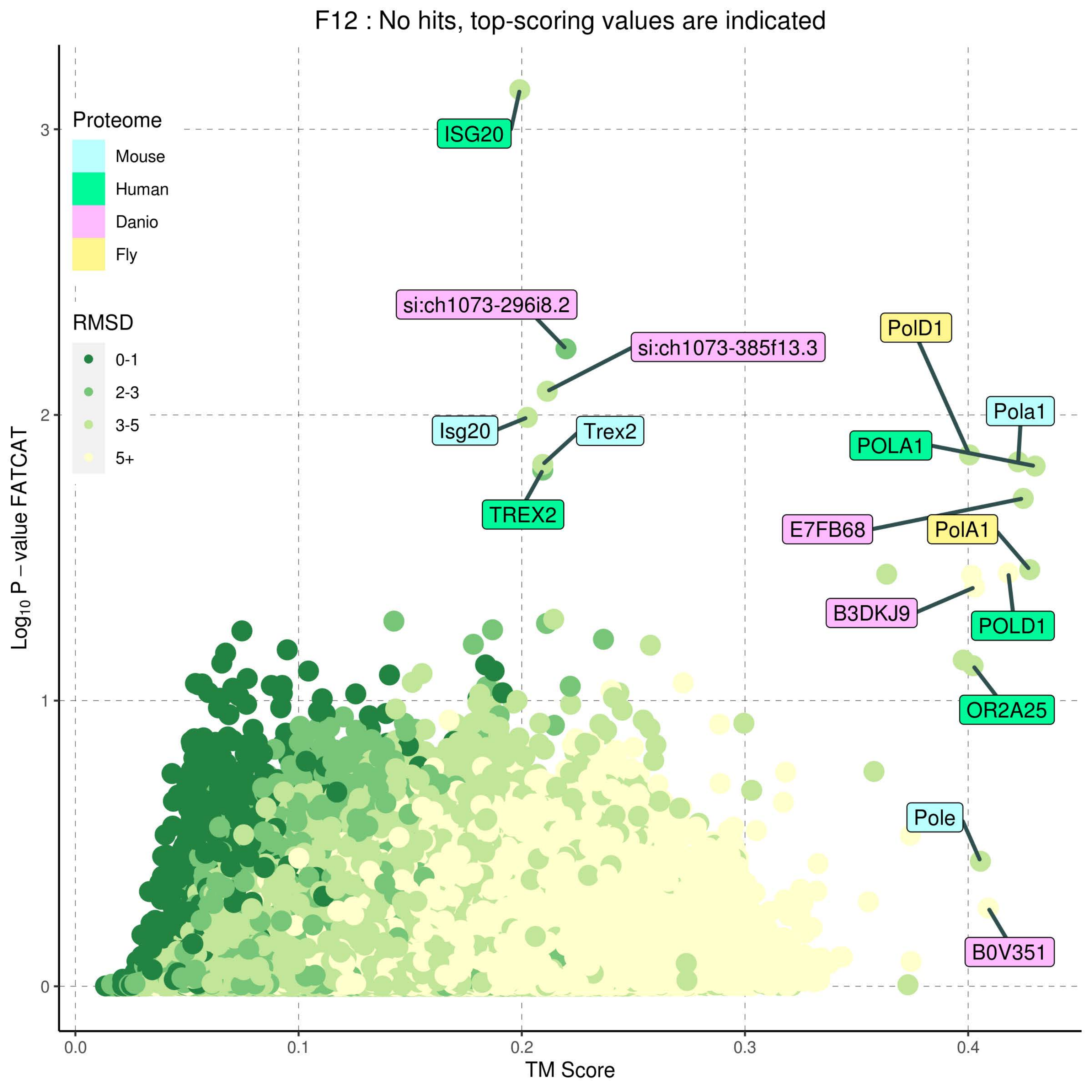

F12 : No hits, top-scoring values are indicated
Proteome
Mouse
Human
Danio
Fly
RMSD
0-1
2-3
3-5
5+
ISG20
si:ch1073-296i8.2
si:ch1073-385f13.3
Isg20
Trex2
TREX2
PolD1
Pola1
POLA1
E7FB68
PolA1
B3DKJ9
POLD1
OR2A25
Pole
B0V351
Log10 P − value FATCAT
TM Score
0
1
2
3
0.0
0.1
0.2
0.3
0.4

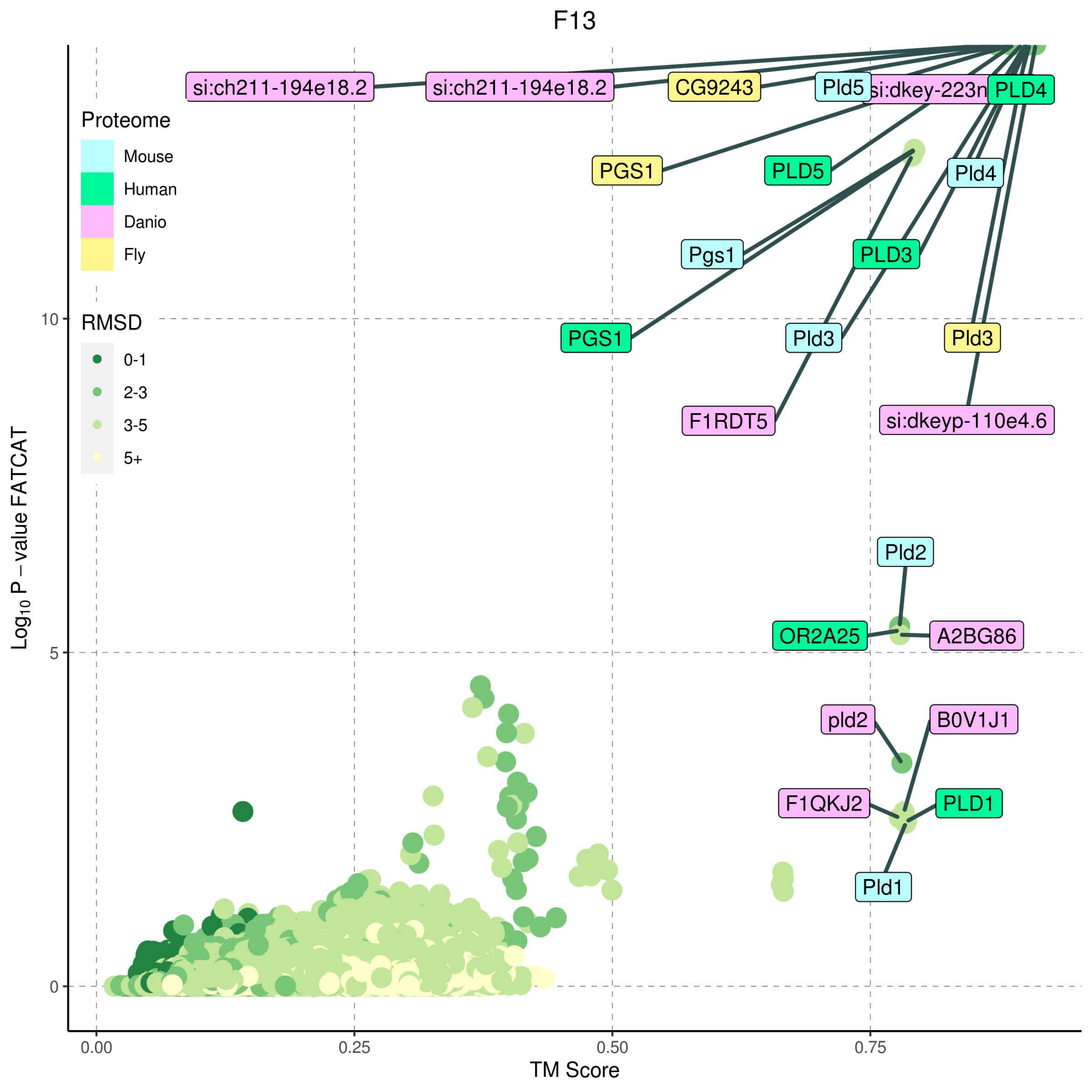
F13
Proteome
Mouse
Human
Danio
Fly
RMSD
0-1
2-3
3-5
5+
si:ch211-194e18.2
si:ch211-194e18.2
CG9243
Pld5
si:dkey-223n
PLD4
PGS1
PLD5
Pld4
Pgs1
PLD3
PGS1
Pld3
Pld3
F1RDT5
si:dkeyp-110e4.6
Pld2
OR2A25
A2BG86
pld2
B0V1J1
F1QKJ2
PLD1
Pld1
Log10 P − value FATCAT
TM Score
0
5
10
0.00
0.25
0.50
0.75

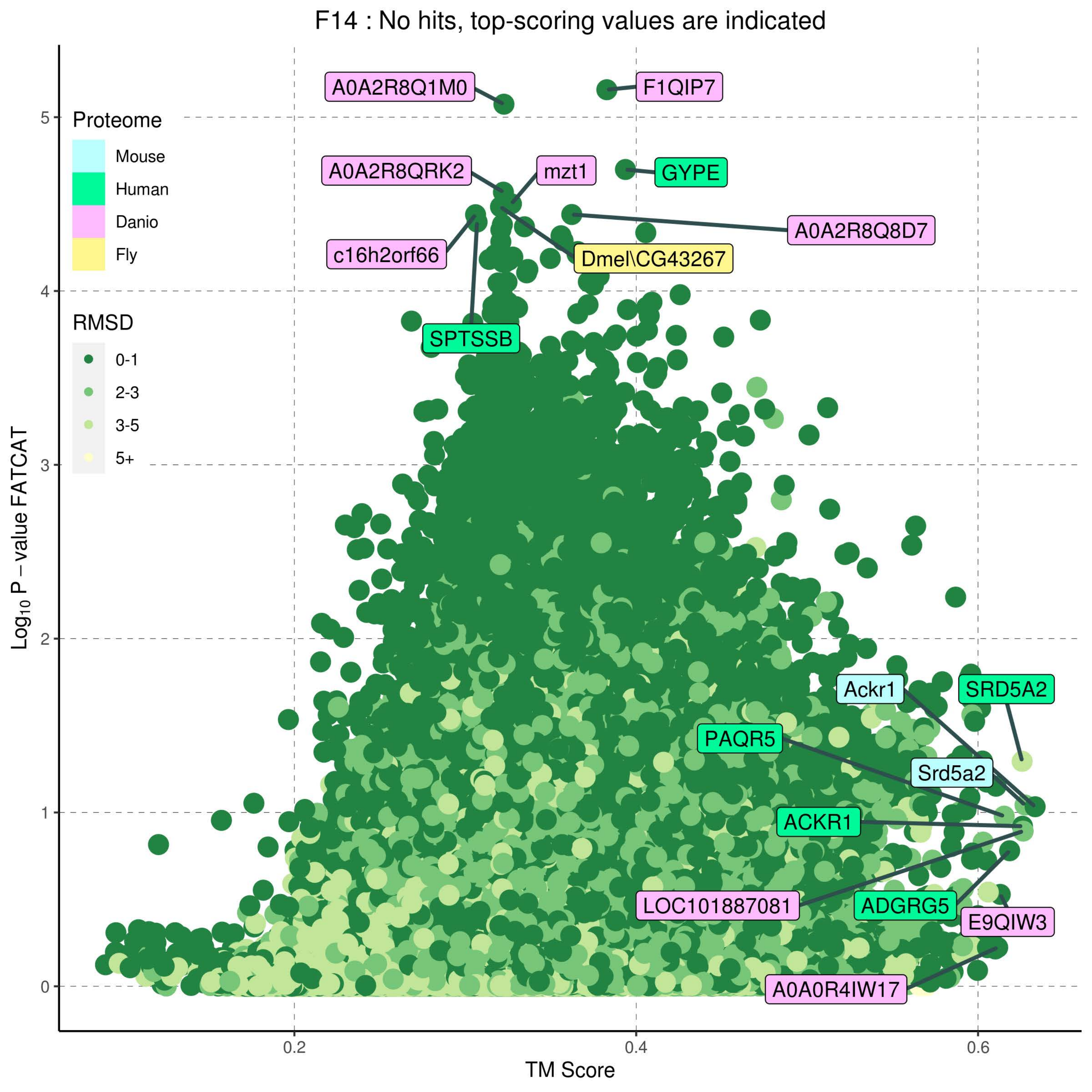

F14 : No hits, top-scoring values are indicated
Proteome
Mouse
Human
Danio
Fly
RMSD
0-1
2-3
3-5
5+
A0A2R8Q1M0
F1QIP7
A0A2R8QRK2
mzt1
GYPE
A0A2R8Q8D7
c16h2orf66
Dmel\CG43267
SPTSSB
Ackr1
SRD5A2
PAQR5
Srd5a2
ACKR1
LOC101887081
ADGRG5
E9QIW3
A0A0R4IW17
Log10 P – value FATCAT
TM Score
0
1
2
3
4
5
0.2
0.4
0.6

F15 : No hits, top-scoring values are indicated

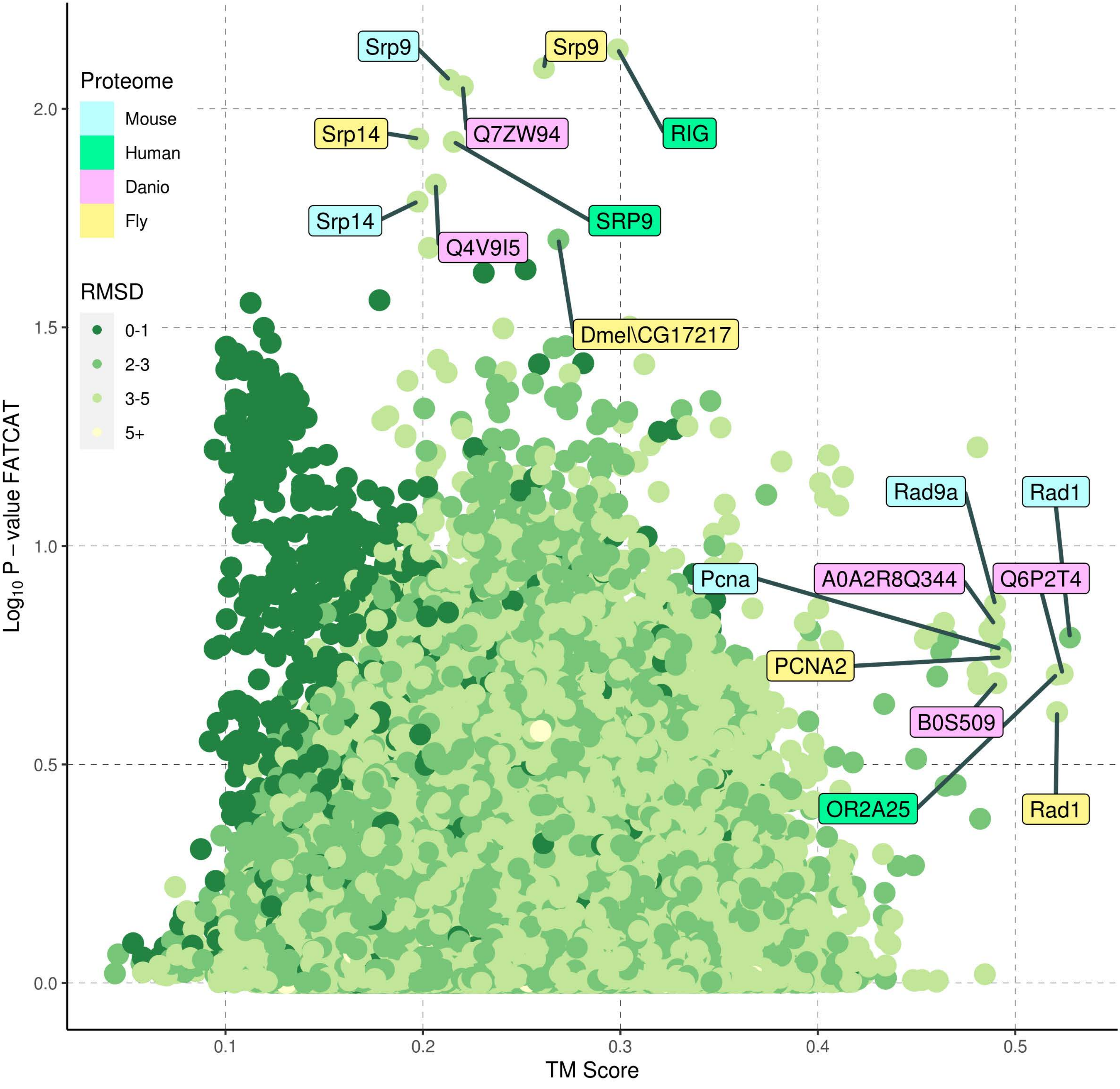

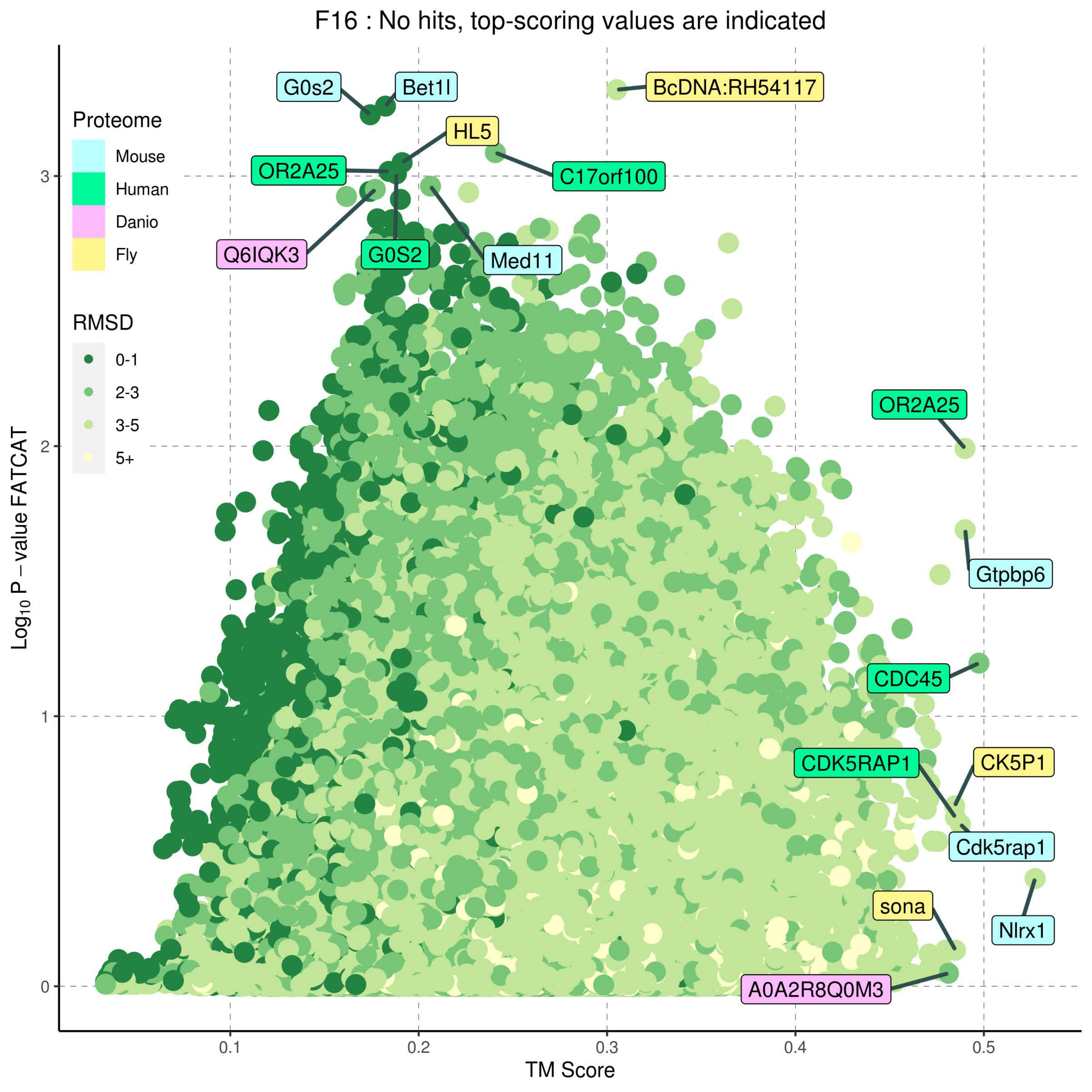
F16 : No hits, top-scoring values are indicated
Proteome
Mouse
Human
Danio
Fly
RMSD
0-1
2-3
3-5
5+
G0s2
Bet1l
BcDNA:RH54117
HL5
OR2A25
C17orf100
Q6IQK3
G0S2
Med11
OR2A25
Gtpbp6
CDC45
CDK5RAP1
CK5P1
Cdk5rap1
sona
Nlrx1
A0A2R8Q0M3
Log10 P − value FATCAT
TM Score
0
1
2
3
0.1
0.2
0.3
0.4
0.5

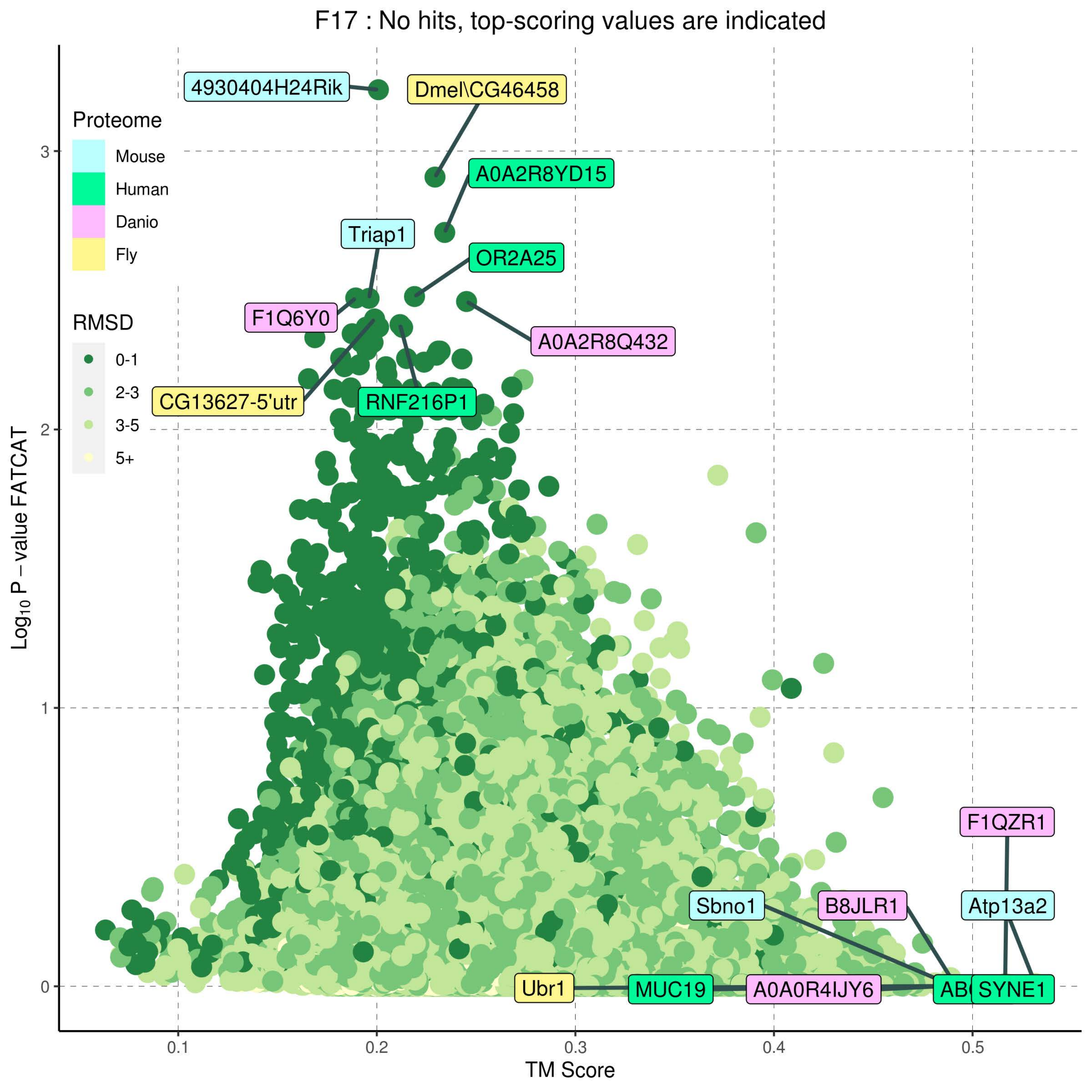
F17 : No hits, top-scoring values are indicated
Proteome
Mouse
Human
Danio
Fly
RMSD
0-1
2-3
3-5
5+
4930404H24Rik
Dmel\CG46458
A0A2R8YD15
Triap1
OR2A25
F1Q6Y0
A0A2R8Q432
CG13627-5'utr
RNF216P1
F1QZR1
Sbno1
B8JLR1
Atp13a2
Ubr1
MUC19
A0A0R4IJY6
AB
SYNE1
Log10 P − value FATCAT
TM Score
0
1
2
3
0.1
0.2
0.3
0.4
0.5

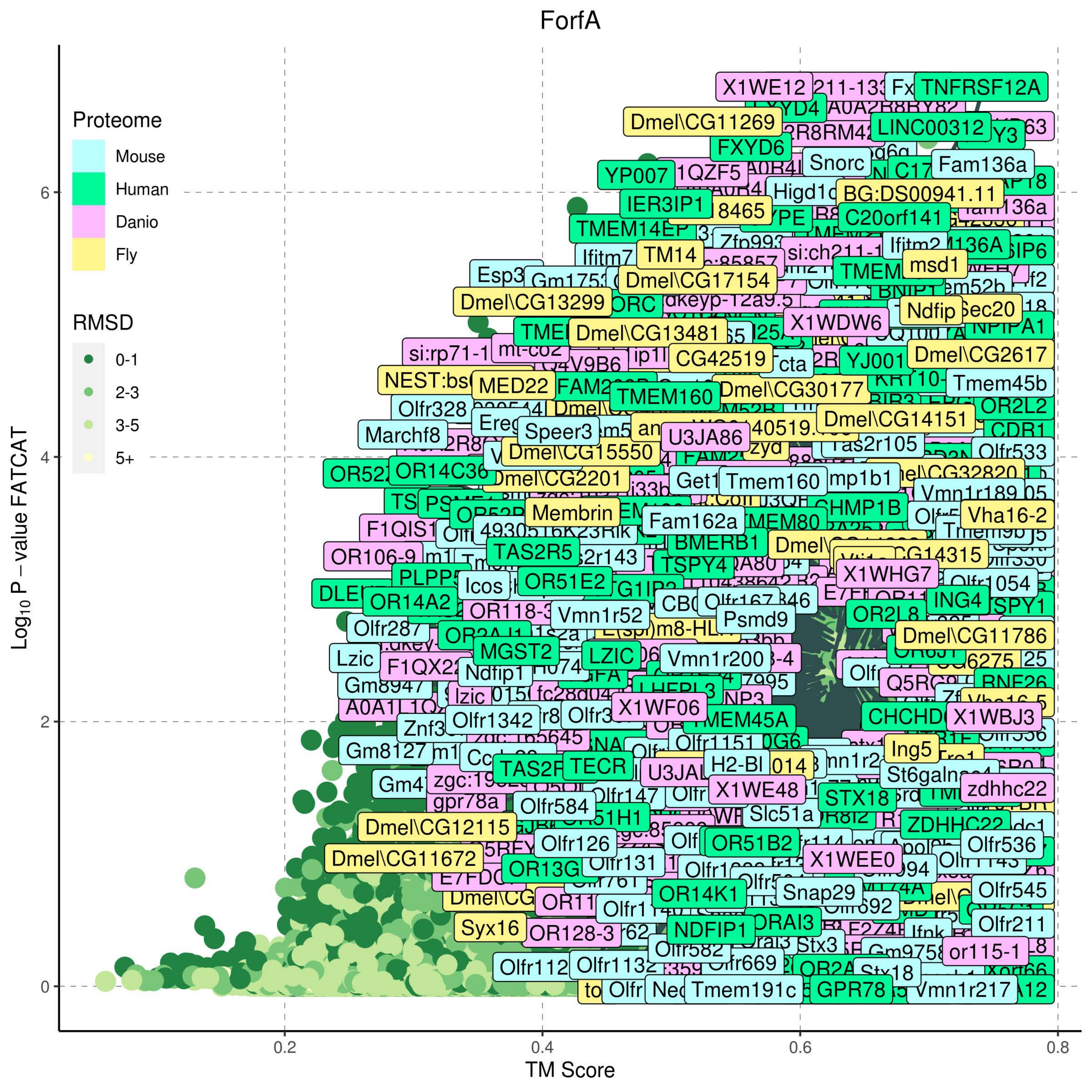
ForfA
Proteome
Mouse
Human
Danio
Fly
RMSD
0-1
2-3
3-5
5+
Log10 P − value FATCAT
0
2
4
6
0.2
0.4
0.6
0.8
TM Score

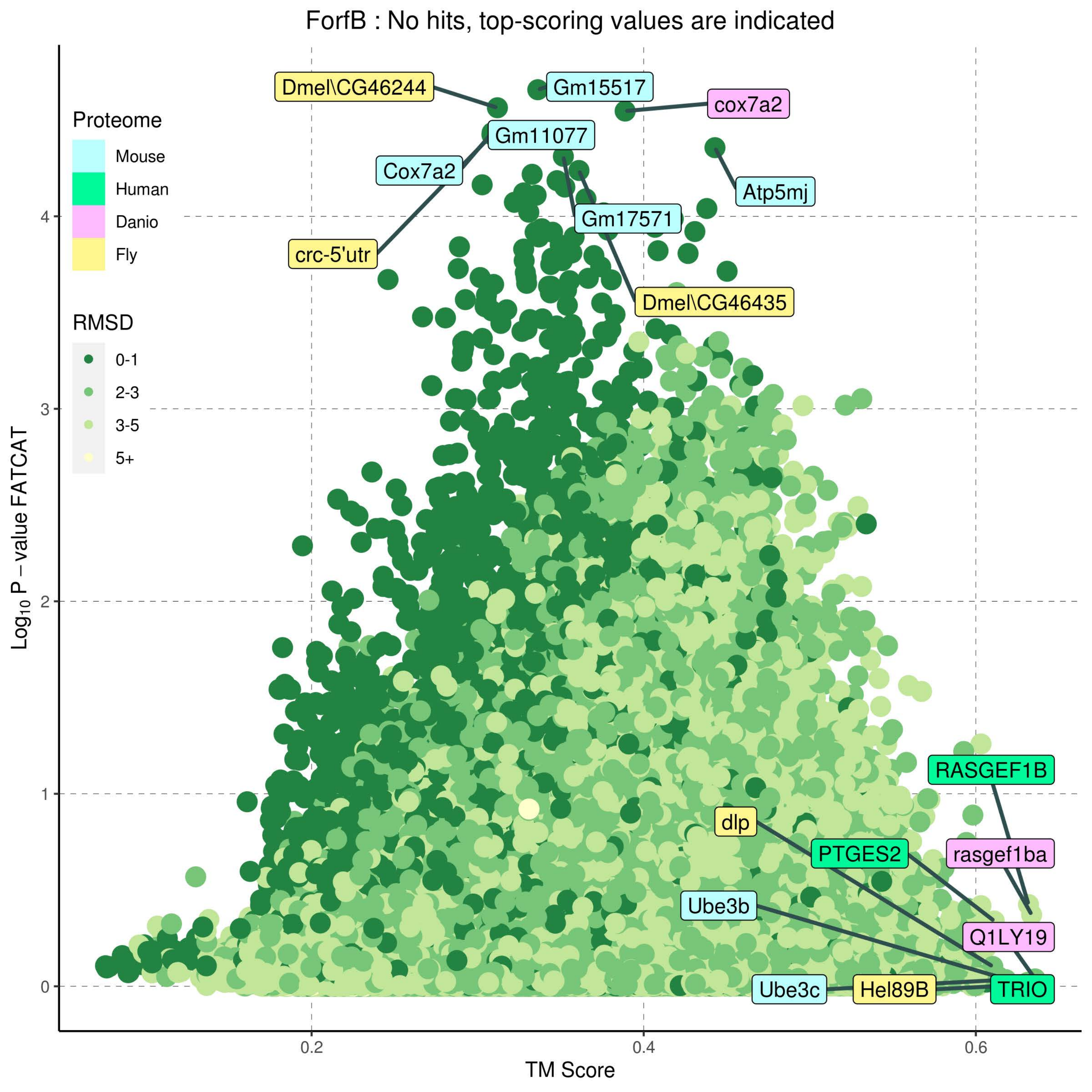
ForfB : No hits, top-scoring values are indicated
Proteome
Mouse
Human
Danio
Fly
RMSD
0-1
2-3
3-5
5+
Dmel\CG46244
Gm15517
cox7a2
Gm11077
Cox7a2
Atp5mj
Gm17571
crc-5'utr
Dmel\CG46435
RASGEF1B
dlp
PTGES2
rasgef1ba
Ube3b
Q1LY19
Ube3c
Hel89B
TRIO
Log10 P – value FATCAT
TM Score
0
1
2
3
4
0.2
0.4
0.6

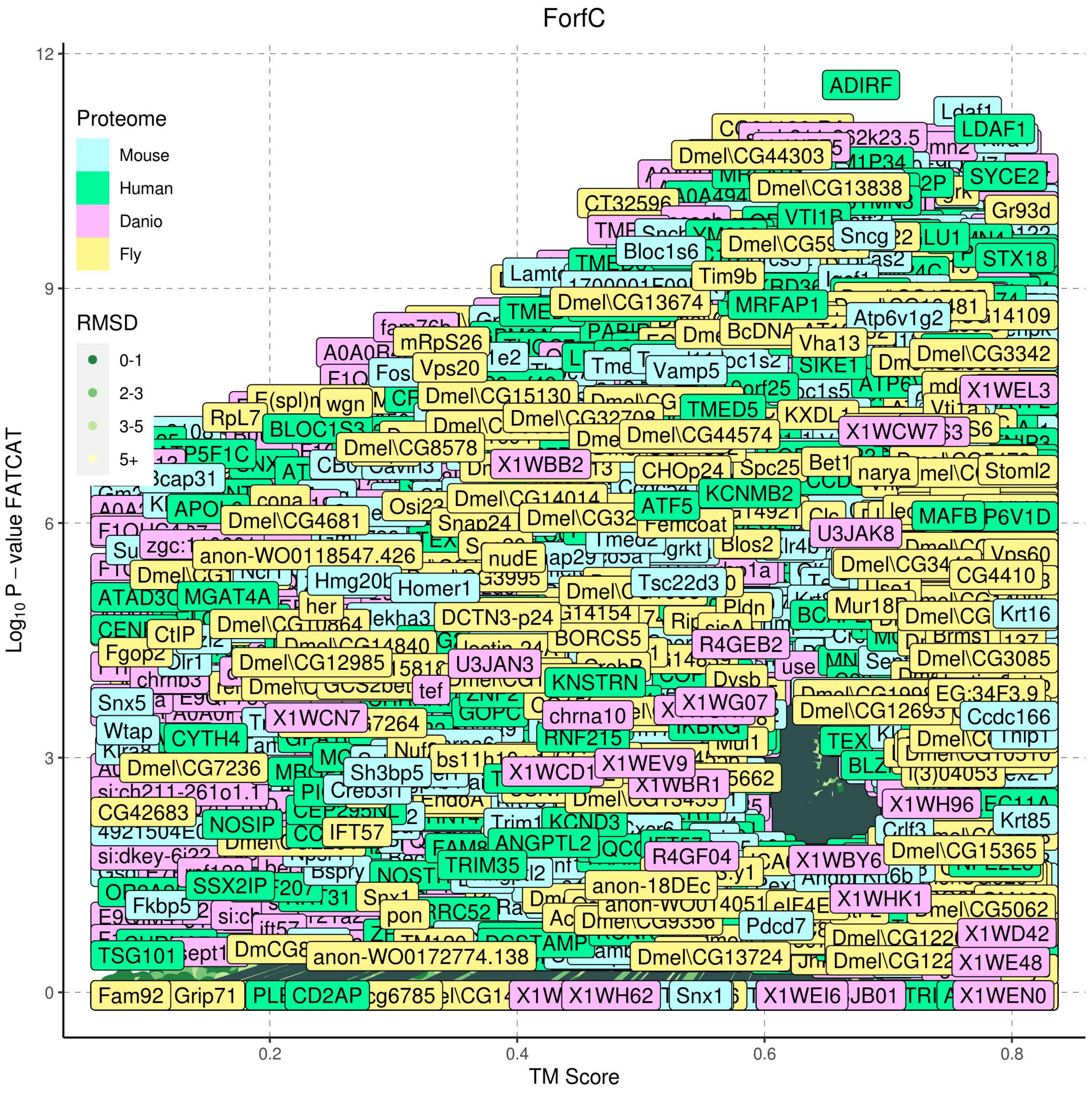

ForfC
Proteome
Mouse
Human
Danio
Fly
RMSD
0-1
2-3
3-5
5+
Log10 P – value FATCAT
TM Score
12
9
6
3
0
0.2
0.4
0.6
0.8
ADIRF
LDAF1
SYCE2
Dmel\CG44303
Dmel\CG13838
CT32596
Gr93d
VTI1B
Sncg
STX18
Bloc1s6
Tim9b
Dmel\CG13674
MRFAP1
Atp6v1g2
mRpS26
Vha13
Dmel\CG3342
Vps20
Vamp5
X1WEL3
Dmel\CG15130
TMED5
KXDL1
RpL7
BLOC1S3
X1WCW7
Dmel\CG44574
Dmel\CG8578
X1WBB2
CHOp24
Spc25
Stoml2
Dmel\CG14014
KCNMB2
ATF5
MAFB
Dmel\CG4681
Snap24
Femcoat
U3JAK8
anon-WO0118547.426
nudE
Blos2
CG4410
Tsc22d3
Homer1
MGAT4A
Mur18B
Krt16
CtIP
DCTN3-p24
BORCS5
R4GEB2
Dmel\CG12985
U3JAN3
Dmel\CG3085
KNSTRN
Snx5
X1WCN7
X1WG07
chrna10
Dmel\CG12693
Ccdc166
Wtap
CYTH4
RNF215
Dmel\CG7236
Sh3bp5
X1WEV9
X1WBR1
si:ch211-261o1.1
CG42683
NOSIP
IFT57
KCND3
X1WH96
Krt85
ANGPTL2
TRIM35
R4GF04
X1WBY6
Dmel\CG15365
anon-18DEc
SSX2IP
Fkbp5
X1WHK1
Pdcd7
X1WD42
TSG101
anon-WO0172774.138
Dmel\CG13724
X1WE48
Fam92
Grip71
CD2AP
X1WH62
Snx1
X1WEI6
X1WEN0

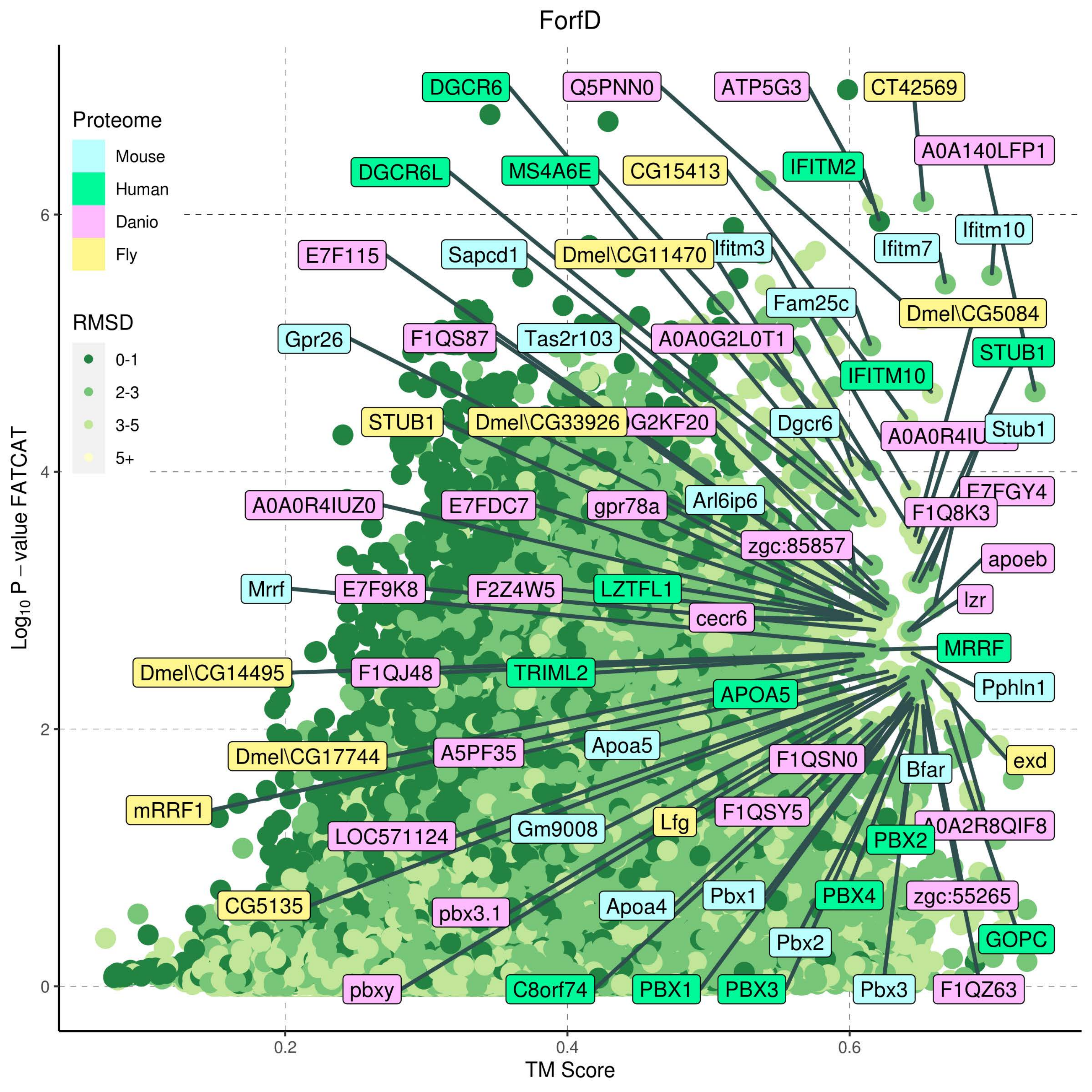

ForfD
Proteome
Mouse
Human
Danio
Fly
RMSD
0-1
2-3
3-5
5+
Log10 P − value FATCAT
TM Score
0
2
4
6
0.2
0.4
0.6
DGCR6
Q5PNN0
ATP5G3
CT42569
A0A140LFP1
DGCR6L
MS4A6E
CG15413
IFITM2
Ifitm10
Ifitm7
E7F115
Sapcd1
Dmel\CG11470
Ifitm3
Fam25c
Dmel\CG5084
Gpr26
F1QS87
Tas2r103
A0A0G2L0T1
STUB1
IFITM10
STUB1
Dmel\CG33926
Dgcr6
Stub1
A0A0R4IUZ0
E7FDC7
gpr78a
Arl6ip6
E7FGY4
F1Q8K3
zgc:85857
apoeb
Mrrf
E7F9K8
F2Z4W5
LZTFL1
lzr
cecr6
MRRF
Dmel\CG14495
F1QJ48
TRIML2
Pphln1
APOA5
Apoa5
Dmel\CG17744
A5PF35
F1QSN0
Bfar
exd
mRRF1
F1QSY5
Gm9008
Lfg
LOC571124
PBX2
A0A2R8QIF8
CG5135
Pbx1
PBX4
zgc:55265
pbx3.1
Apoa4
GOPC
Pbx2
pbxy
C8orf74
PBX1
PBX3
Pbx3
F1QZ63

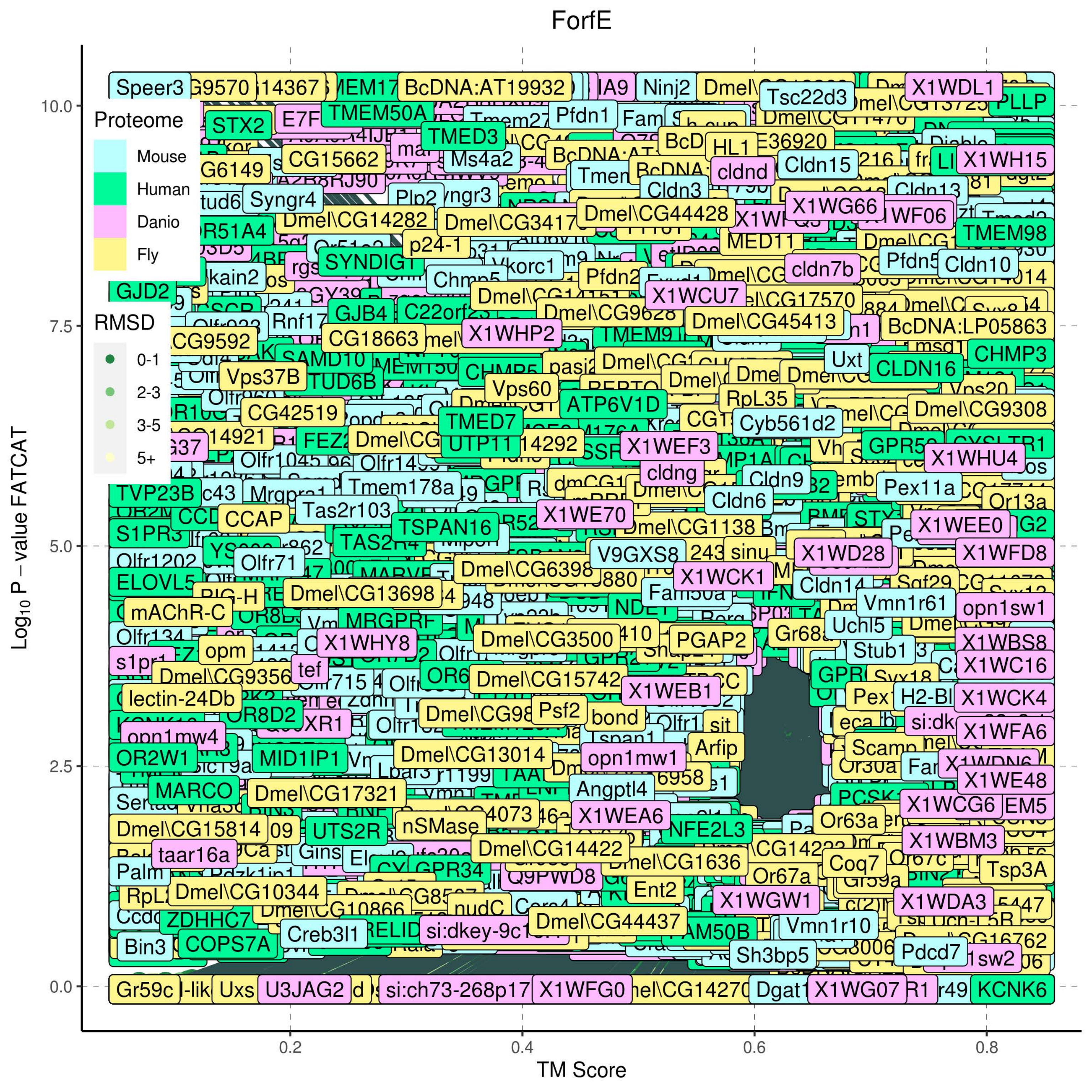
ForfE
Proteome
Mouse
Human
Danio
Fly
RMSD
0-1
2-3
3-5
5+
$Log_{10}$ P – value FATCAT
TM Score
0.0
2.5
5.0
7.5
10.0
0.2
0.4
0.6
0.8

ForfF

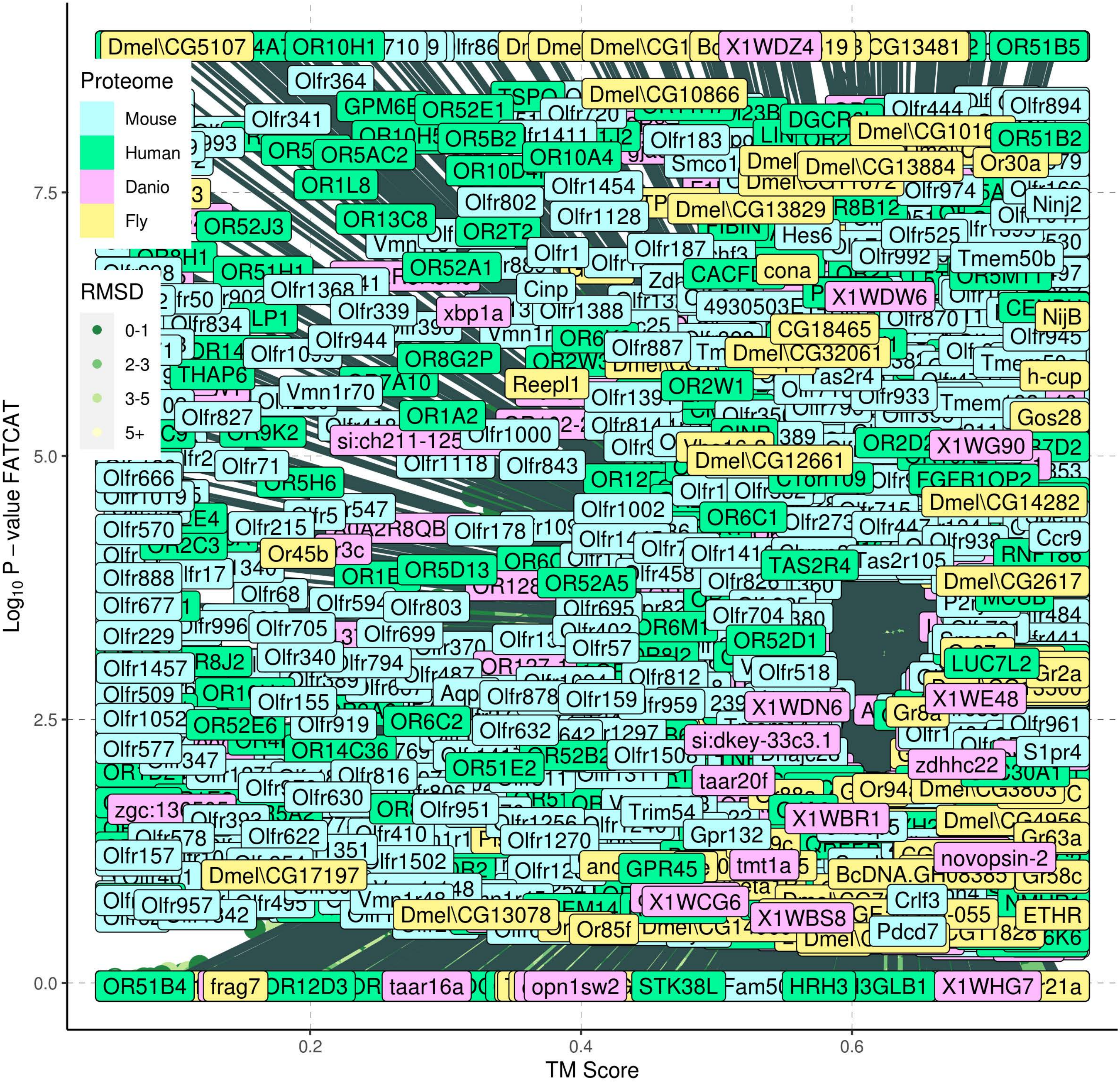

Proteome
Mouse
Human
Danio
Fly
RMSD
0-1
2-3
3-5
5+
Log10 P – value FATCAT
7.5
5.0
2.5
0.0
0.2
0.4
0.6
TM Score

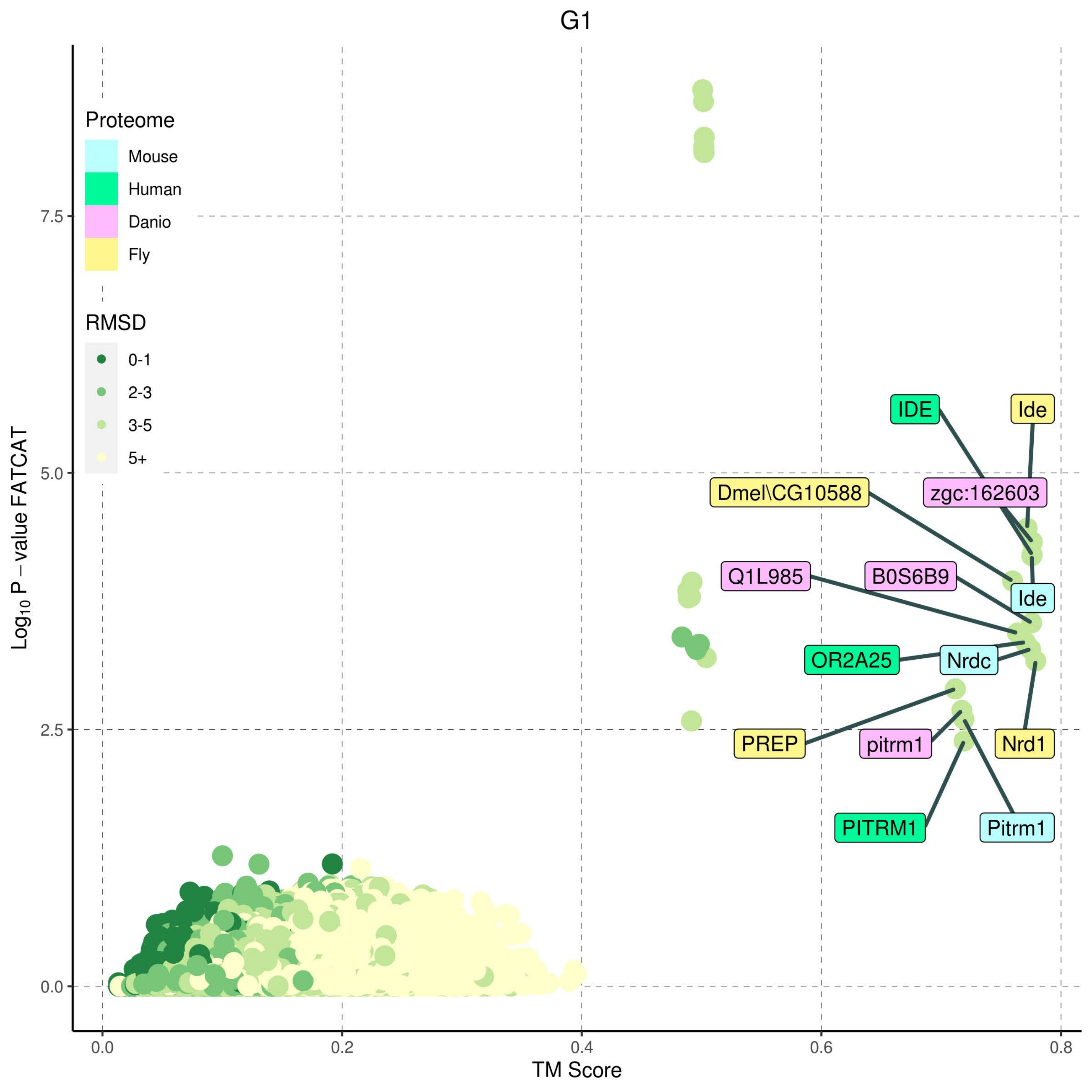

G1
Proteome
Mouse
Human
Danio
Fly
RMSD
0-1
2-3
3-5
5+
IDE
Ide
Dmel\CG10588
zgc:162603
Q1L985
B0S6B9
Ide
OR2A25
Nrdc
PREP
pitrm1
Nrd1
PITRM1
Pitrm1
7.5
5.0
2.5
0.0
0.0
0.2
0.4
0.6
0.8
$Log_{10}$ P − value FATCAT
TM Score

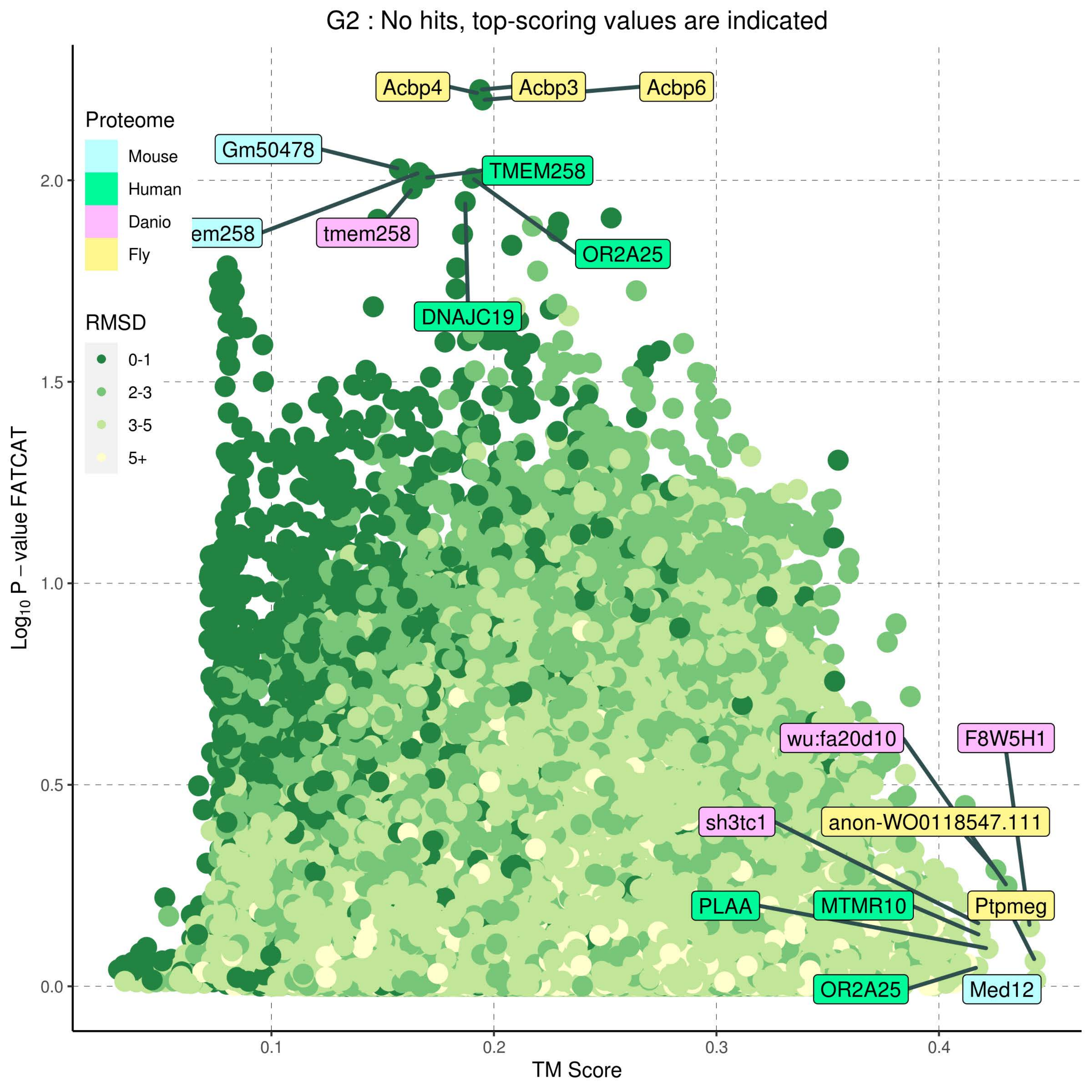

G2 : No hits, top-scoring values are indicated
Proteome
Mouse
Human
Danio
Fly
RMSD
0-1
2-3
3-5
5+
Acbp4
Acbp3
Acbp6
Gm50478
TMEM258
em258
tmem258
OR2A25
DNAJC19
wu:fa20d10
F8W5H1
sh3tc1
anon-WO0118547.111
PLAA
MTMR10
Ptpmeg
OR2A25
Med12
$Log_{10}$ P − value FATCAT
TM Score
0.0
0.5
1.0
1.5
2.0
0.1
0.2
0.3
0.4

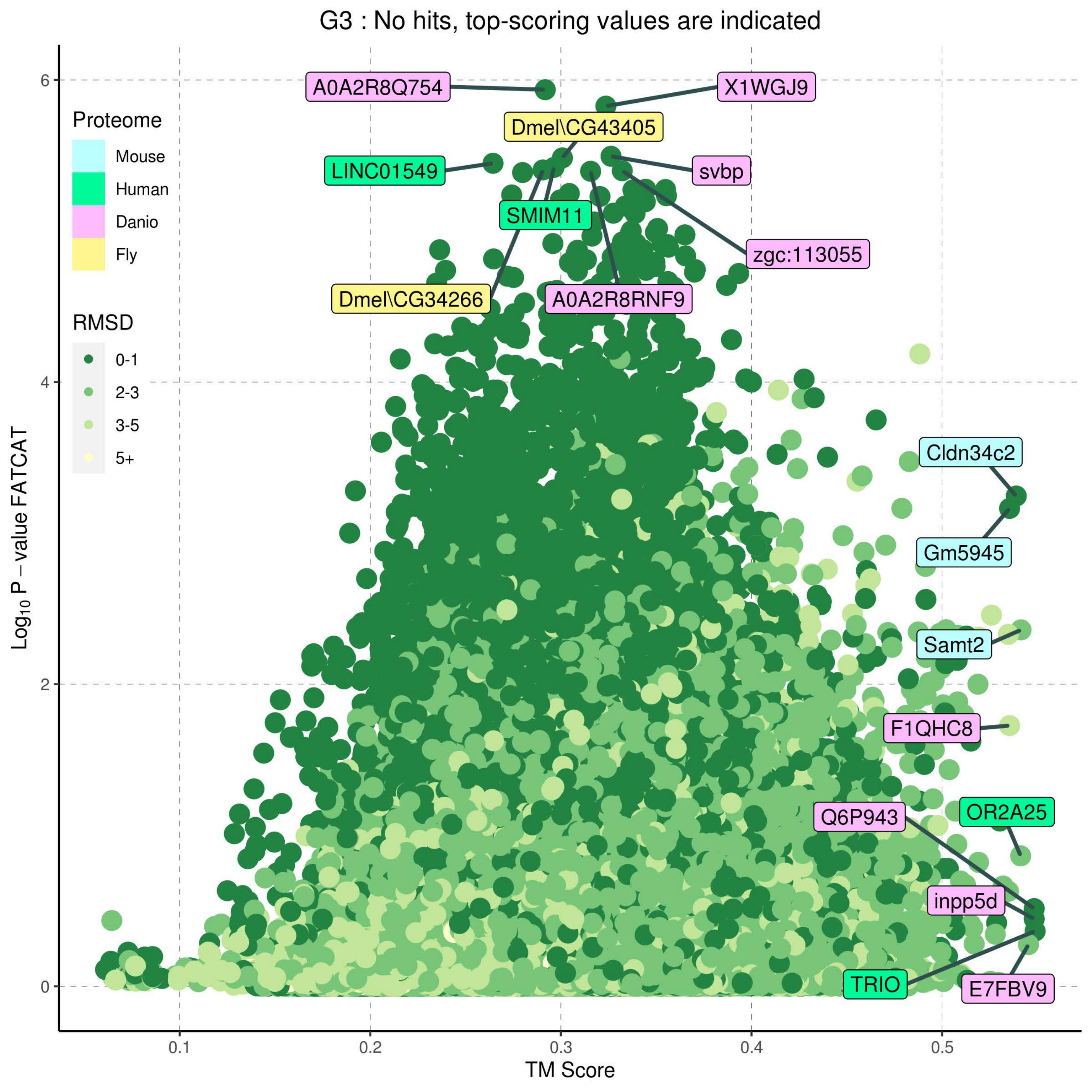

G3 : No hits, top-scoring values are indicated
Proteome
Mouse
Human
Danio
Fly
RMSD
0-1
2-3
3-5
5+
A0A2R8Q754
X1WGJ9
Dmel\CG43405
LINC01549
svbp
SMIM11
zgc:113055
Dmel\CG34266
A0A2R8RNF9
Cldn34c2
Gm5945
Samt2
F1QHC8
OR2A25
Q6P943
inpp5d
TRIO
E7FBV9
6
4
2
0
0.1
0.2
0.3
0.4
0.5
TM Score
$Log_{10}$ P − value FATCAT

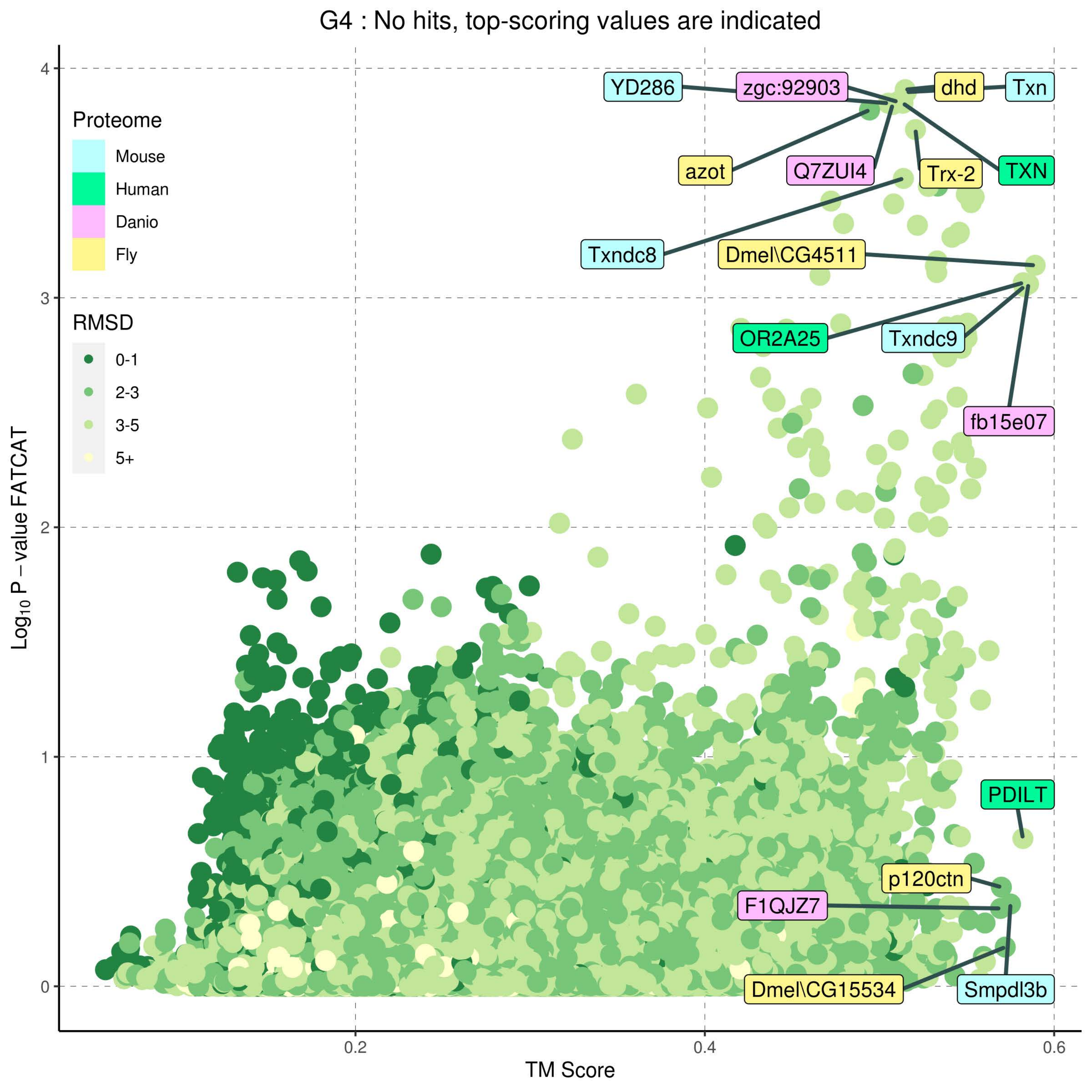
G4 : No hits, top-scoring values are indicated
Proteome
Mouse
Human
Danio
Fly
RMSD
0-1
2-3
3-5
5+
YD286
zgc:92903
dhd
Txn
azot
Q7ZUI4
Trx-2
TXN
Txndc8
Dmel\CG4511
OR2A25
Txndc9
fb15e07
PDILT
p120ctn
F1QJZ7
Dmel\CG15534
Smpdl3b
Log10 P – value FATCAT
TM Score
0
1
2
3
4
0.2
0.4
0.6

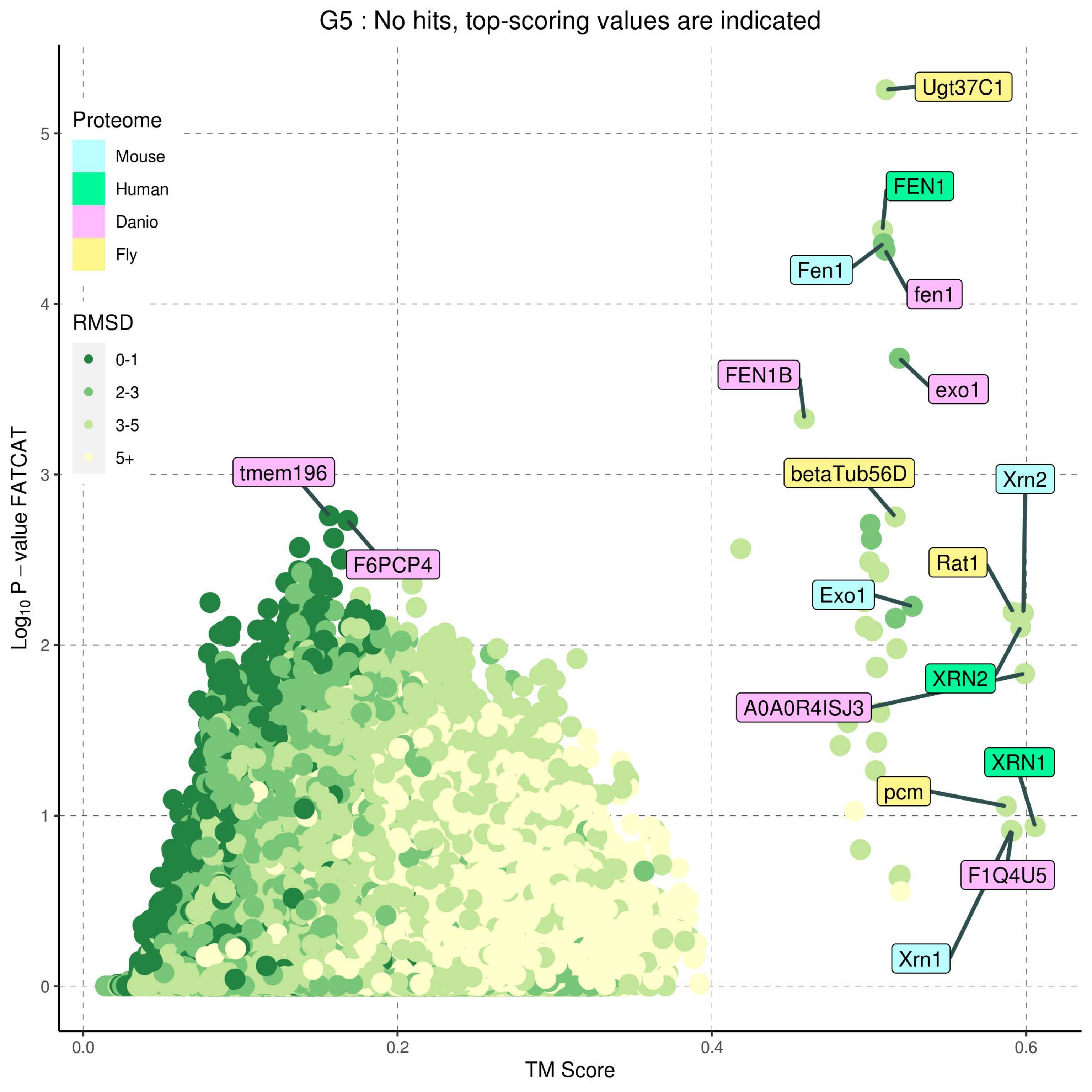

G5 : No hits, top-scoring values are indicated
Proteome
Mouse
Human
Danio
Fly
RMSD
0-1
2-3
3-5
5+
Log10 P – value FATCAT
TM Score
0.0
0.2
0.4
0.6
0
1
2
3
4
5
Ugt37C1
FEN1
Fen1
fen1
exo1
FEN1B
betaTub56D
Xrn2
tmem196
F6PCP4
Rat1
Exo1
XRN2
A0A0R4ISJ3
XRN1
pcm
F1Q4U5
Xrn1

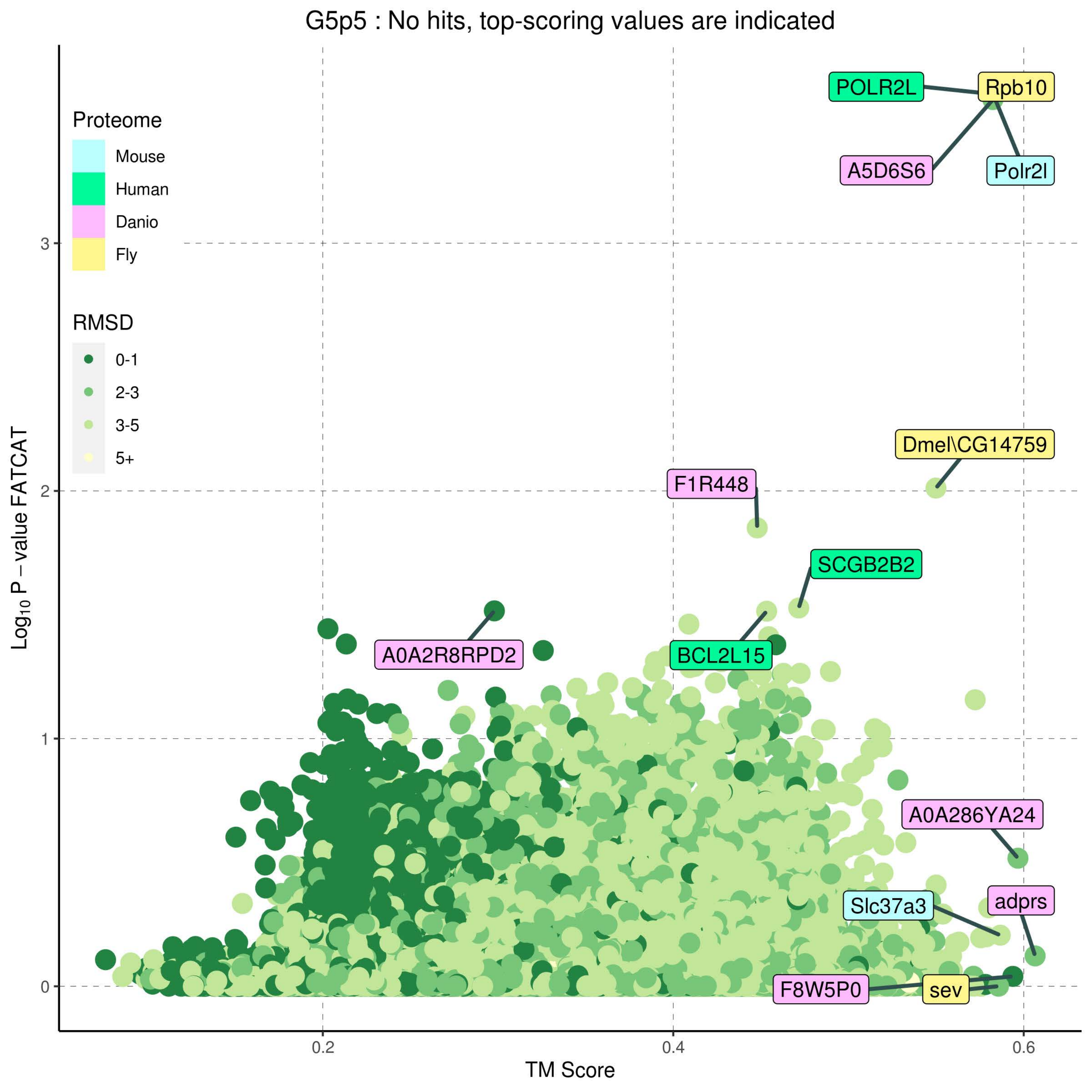
G5p5 : No hits, top-scoring values are indicated
Proteome
Mouse
Human
Danio
Fly
RMSD
0-1
2-3
3-5
5+
POLR2L
Rpb10
A5D6S6
Polr2l
Dmel\CG14759
F1R448
SCGB2B2
A0A2R8RPD2
BCL2L15
A0A286YA24
adprs
Slc37a3
F8W5P0
sev
Log10 P − value FATCAT
TM Score
0
1
2
3
0.2
0.4
0.6

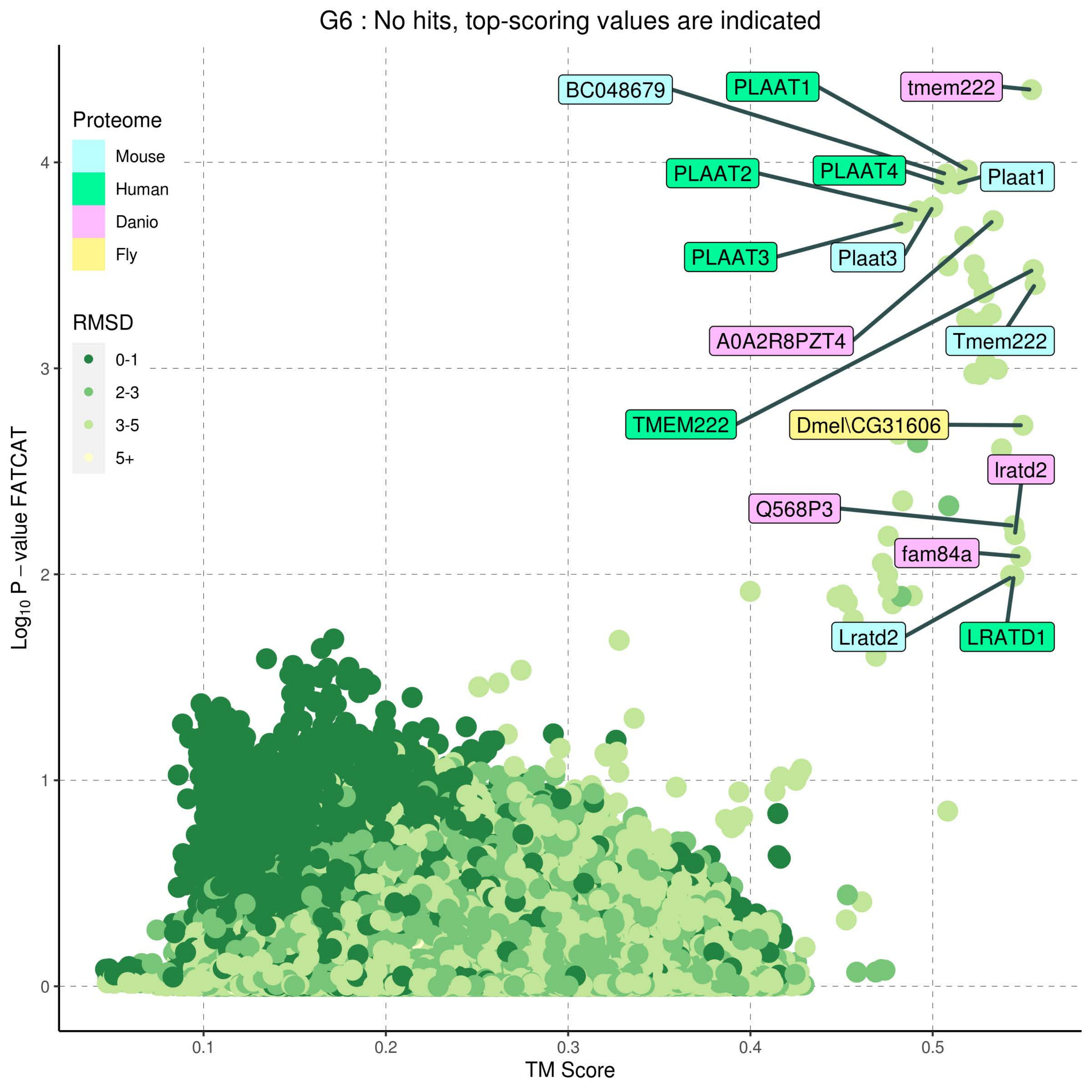

G6 : No hits, top-scoring values are indicated
Proteome
Mouse
Human
Danio
Fly
RMSD
0-1
2-3
3-5
5+
BC048679
PLAAT1
tmem222
PLAAT2
PLAAT4
Plaat1
PLAAT3
Plaat3
A0A2R8PZT4
Tmem222
TMEM222
Dmel\CG31606
lratd2
Q568P3
fam84a
Lratd2
LRATD1
Log10 P – value FATCAT
TM Score
0
1
2
3
4
0.1
0.2
0.3
0.4
0.5

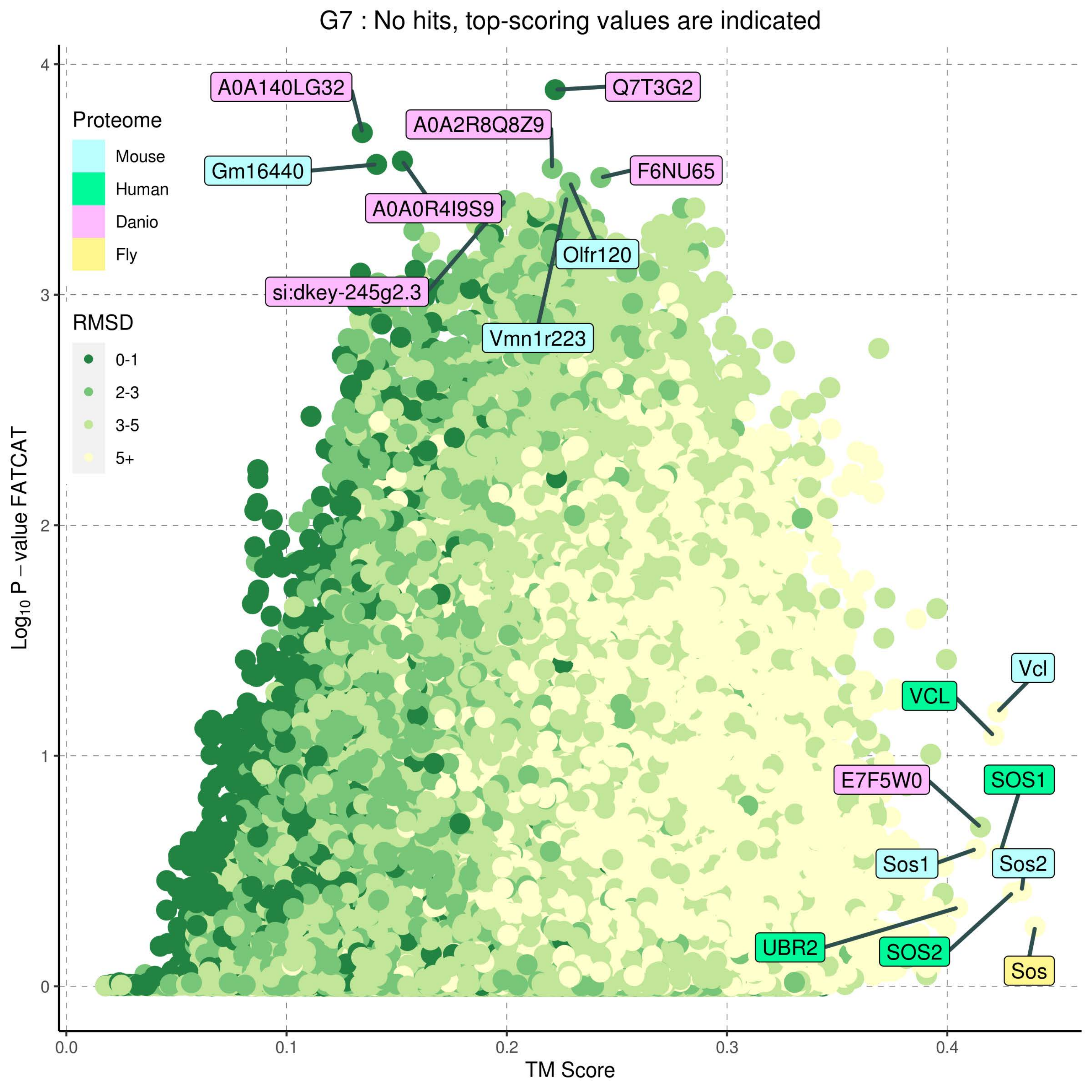
G7 : No hits, top-scoring values are indicated
Proteome
Mouse
Human
Danio
Fly
RMSD
0-1
2-3
3-5
5+
A0A140LG32
Q7T3G2
A0A2R8Q8Z9
Gm16440
F6NU65
A0A0R4I9S9
Olfr120
si:dkey-245g2.3
Vmn1r223
Vcl
VCL
E7F5W0
SOS1
Sos1
Sos2
UBR2
SOS2
Sos
Log10 P − value FATCAT
TM Score
0
1
2
3
4
0.0
0.1
0.2
0.3
0.4

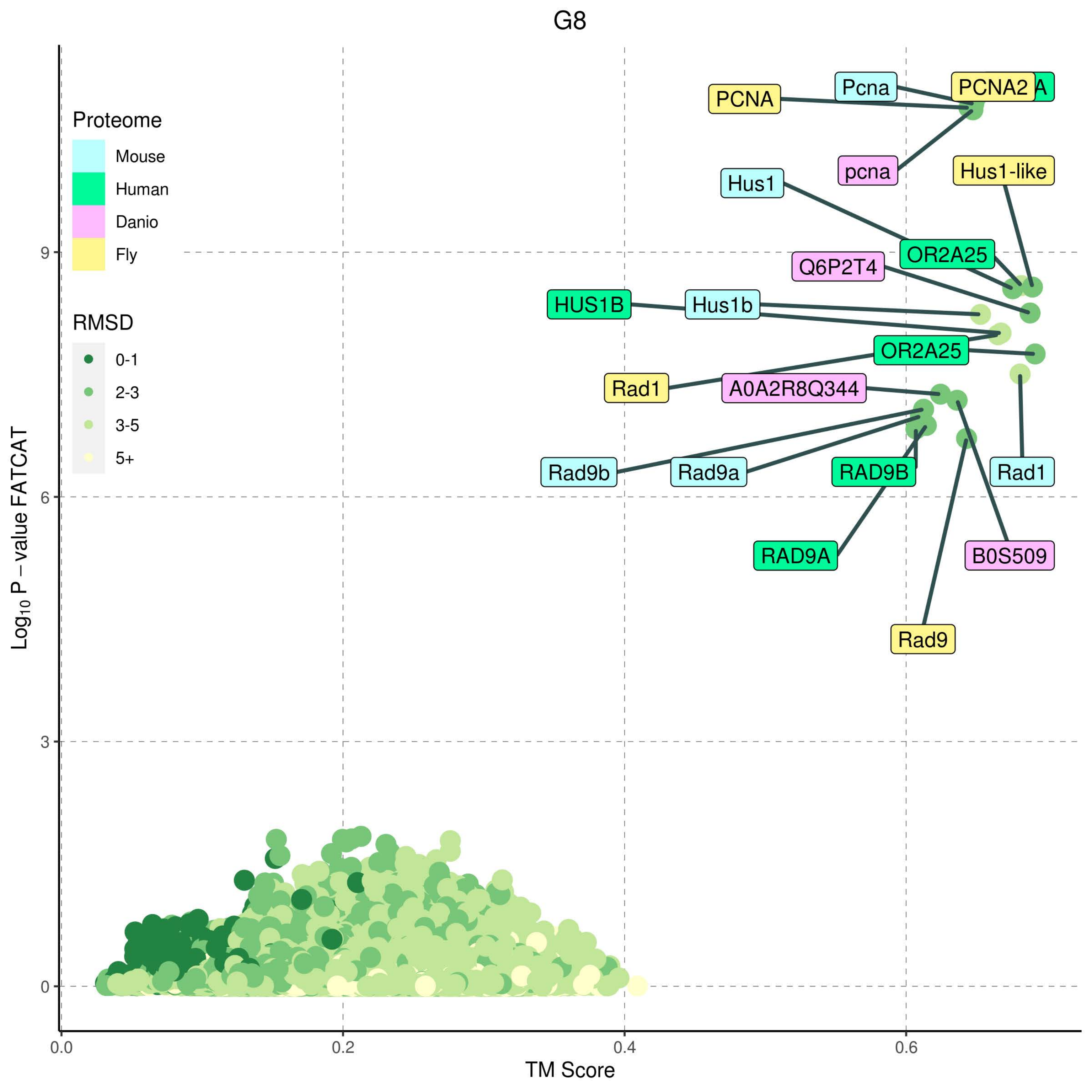
G8
Proteome
Mouse
Human
Danio
Fly
RMSD
0-1
2-3
3-5
5+
Log10 P − value FATCAT
TM Score
0.0
0.2
0.4
0.6
0
3
6
9
PCNA
Pcna
PCNA2
pcna
Hus1
Hus1-like
OR2A25
Q6P2T4
HUS1B
Hus1b
OR2A25
Rad1
A0A2R8Q344
Rad9b
Rad9a
RAD9B
Rad1
RAD9A
B0S509
Rad9

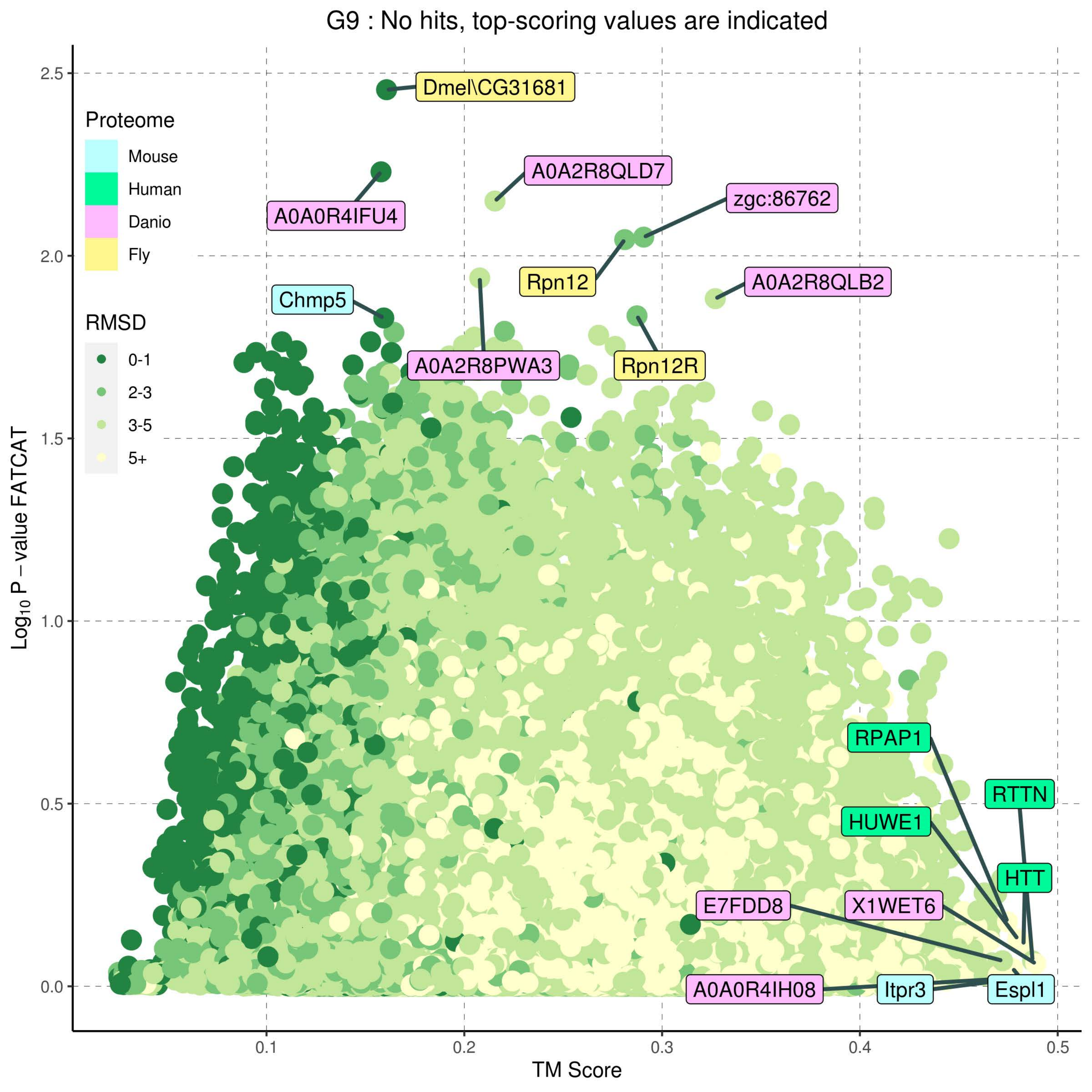
G9 : No hits, top-scoring values are indicated
Proteome
Mouse
Human
Danio
Fly
RMSD
0-1
2-3
3-5
5+
Log10 P − value FATCAT
TM Score
2.5
2.0
1.5
1.0
0.5
0.0
0.1
0.2
0.3
0.4
0.5
Dmel\CG31681
A0A2R8QLD7
zgc:86762
A0A0R4IFU4
Rpn12
A0A2R8QLB2
Chmp5
A0A2R8PWA3
Rpn12R
RPAP1
RTTN
HUWE1
HTT
E7FDD8
X1WET6
A0A0R4IH08
Itpr3
Espl1

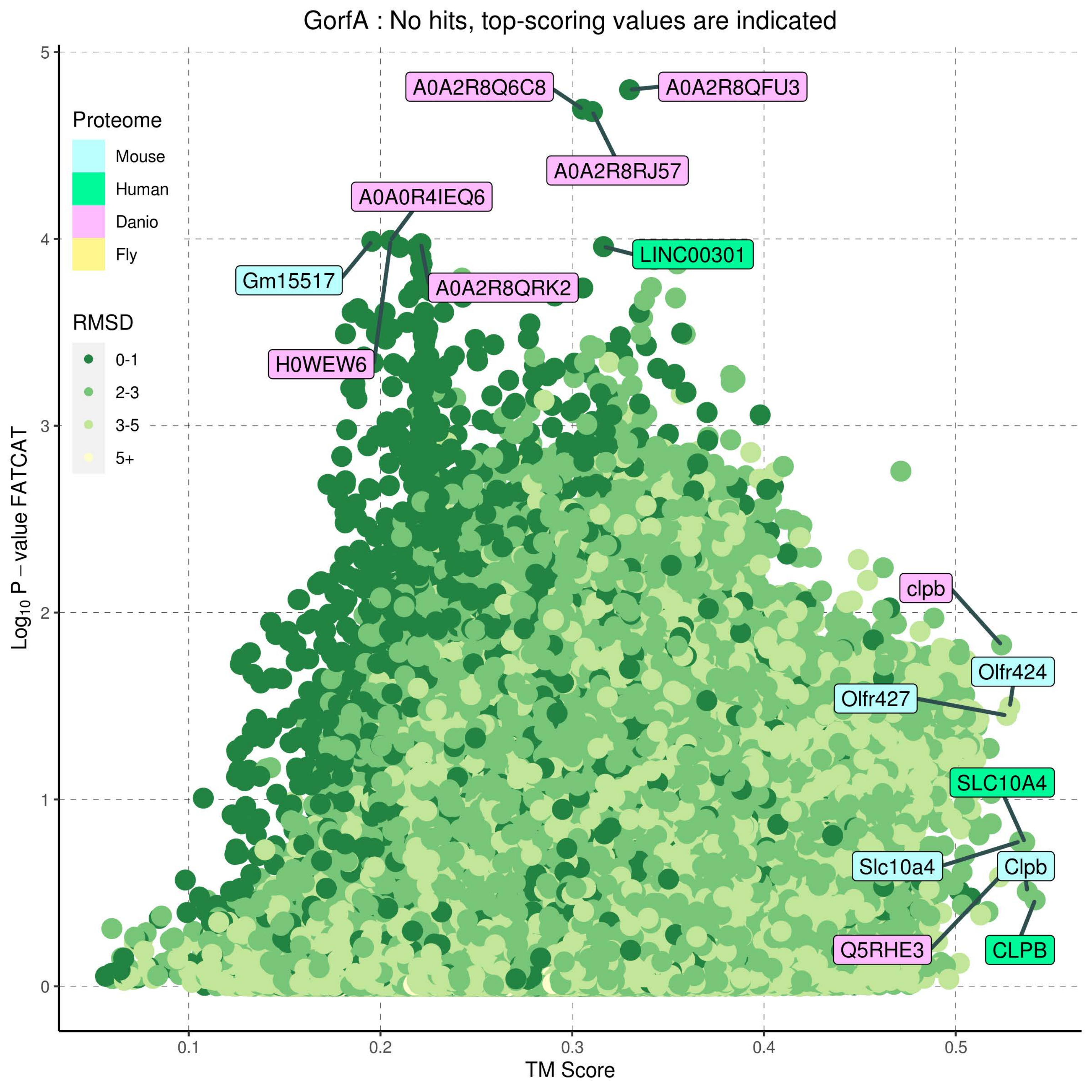
GorfA : No hits, top-scoring values are indicated
Proteome
Mouse
Human
Danio
Fly
RMSD
0-1
2-3
3-5
5+
A0A2R8Q6C8
A0A2R8QFU3
A0A2R8RJ57
A0A0R4IEQ6
Gm15517
LINC00301
A0A2R8QRK2
H0WEW6
clpb
Olfr424
Olfr427
SLC10A4
Slc10a4
Clpb
Q5RHE3
CLPB
0
1
2
3
4
5
0.1
0.2
0.3
0.4
0.5
TM Score
Log10 P − value FATCAT

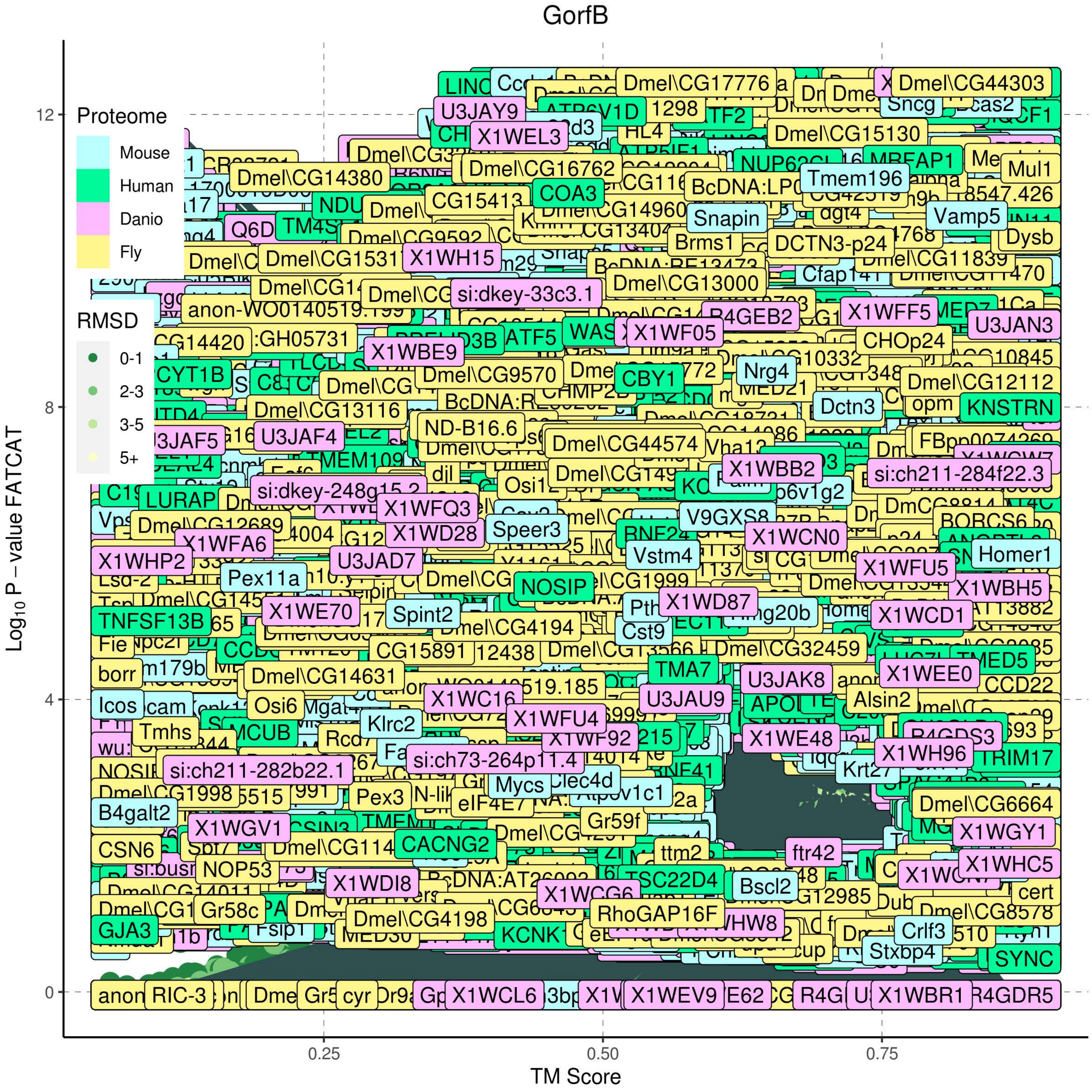

GorfB
Proteome
Mouse
Human
Danio
Fly
RMSD
0-1
2-3
3-5
5+
Log10 P – value FATCAT
TM Score
0
4
8
12
0.25
0.50
0.75
Dmel\CG17776
Dmel\CG44303
U3JAY9
X1WEL3
Dmel\CG15130
Dmel\CG14380
Dmel\CG16762
COA3
Tmem196
Mul1
Snapin
Vamp5
Brms1
DCTN3-p24
Dysb
X1WH15
Dmel\CG13000
si:dkey-33c3.1
X1WFF5
U3JAN3
X1WBE9
ATF5
X1WF05
CHOp24
Dmel\CG9570
CBY1
Nrg4
Dmel\CG12112
Dmel\CG13116
Dctn3
KNSTRN
U3JAF4
ND-B16.6
Dmel\CG44574
X1WBB2
si:ch211-284f22.3
LURAP
si:dkey-248g15.2
X1WFQ3
Dmel\CG12689
X1WD28
Speer3
V9GXS8
X1WCN0
X1WFA6
Vstm4
Homer1
X1WHP2
U3JAD7
X1WFU5
Pex11a
NOSIP
X1WBH5
X1WE70
Spint2
X1WD87
X1WCD1
TNFSF13B
Dmel\CG4194
Cst9
CG15891
Dmel\CG32459
TMA7
Dmel\CG14631
U3JAK8
X1WEE0
Icos
Osi6
X1WC16
U3JAU9
Alsin2
X1WFU4
Klrc2
Tmhs
MCUB
X1WE48
X1WH96
si:ch73-264p11.4
TRIM17
si:ch211-282b22.1
Krt27
Mycs
Clec4d
Dmel\CG1998
Dmel\CG6664
B4galt2
Pex3
Gr59f
X1WGV1
X1WGY1
CSN6
CACNG2
ftr42
NOP53
X1WHC5
X1WDI8
TSC22D4
Bscl2
X1WCG6
Gr58c
RhoGAP16F
Dmel\CG4198
Dmel\CG8578
GJA3
Crlf3
Stxbp4
SYNC
RIC-3
X1WCL6
X1WEV9
X1WBR1
R4GDR5

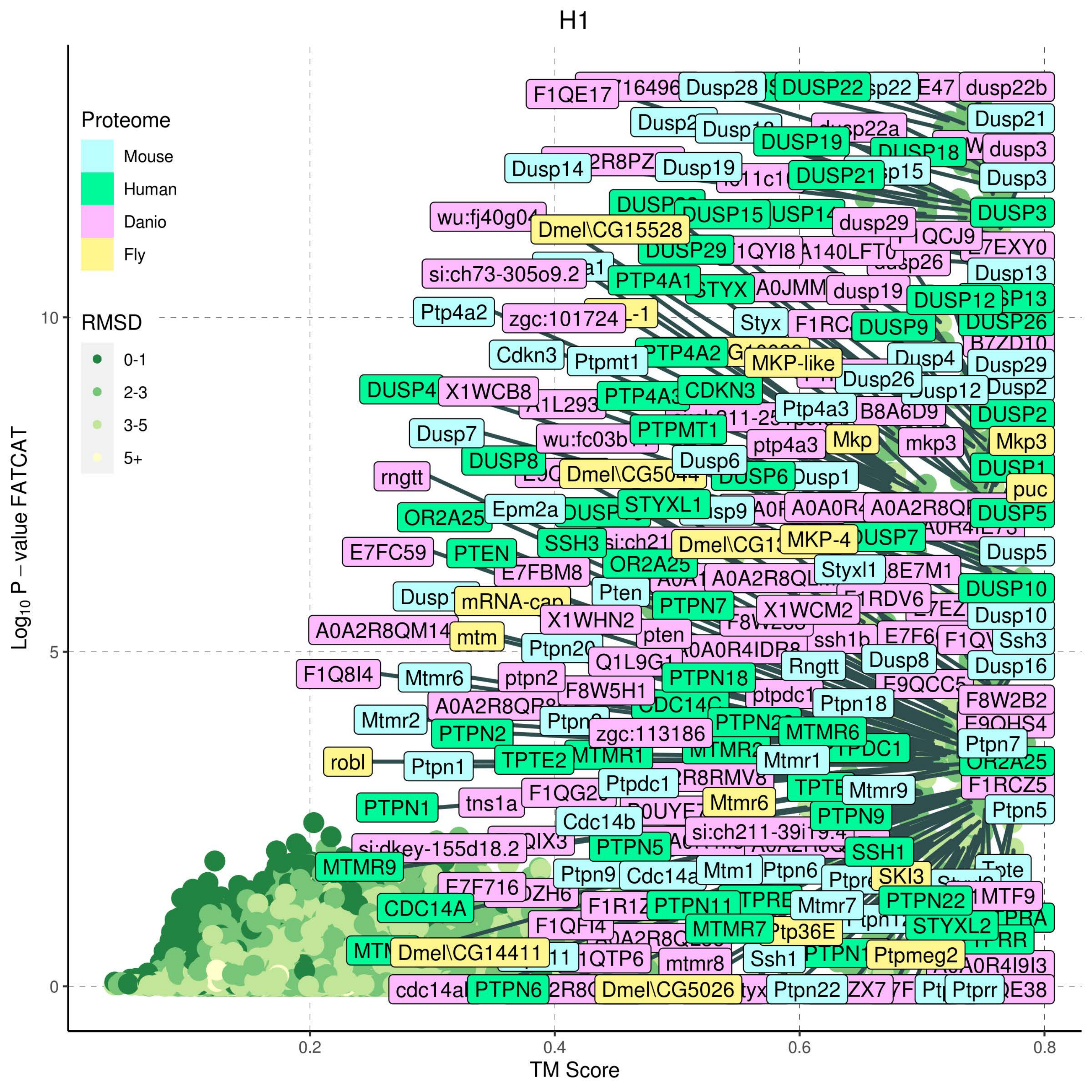
H1
Proteome
Mouse
Human
Danio
Fly
RMSD
0-1
2-3
3-5
5+
Log10 P – value FATCAT
TM Score
0
5
10
0.2
0.4
0.6
0.8
F1QE17
Dusp28
DUSP22
dusp22b
Dusp21
DUSP19
dusp3
Dusp14
Dusp19
DUSP21
Dusp3
DUSP15
DUSP3
Dmel\CG15528
dusp29
DUSP29
si:ch73-305o9.2
PTP4A1
dusp19
Dusp13
DUSP12
Ptp4a2
zgc:101724
Styx
DUSP9
DUSP26
Cdkn3
Ptpmt1
PTP4A2
MKP-like
Dusp4
Dusp29
DUSP4
X1WCB8
CDKN3
Dusp26
Dusp12
Dusp2
Ptp4a3
B8A6D9
DUSP2
Dusp7
PTPMT1
ptp4a3
Mkp
mkp3
Mkp3
DUSP8
Dusp6
DUSP1
rngtt
puc
OR2A25
Epm2a
STYXL1
DUSP5
SSH3
MKP-4
E7FC59
PTEN
Dusp5
E7FBM8
OR2A25
Styxl1
Pten
DUSP10
PTPN7
X1WCM2
X1WHN2
Dusp10
A0A2R8QM14
mtm
pten
Ssh3
Rngtt
Dusp8
F1Q8I4
Mtmr6
ptpn2
F8W5H1
PTPN18
Dusp16
Ptpn18
F8W2B2
Mtmr2
PTPN2
zgc:113186
MTMR6
Ptpn7
robl
Ptpn1
TPTE2
MTMR1
Mtmr1
F1RCZ5
Ptpdc1
Mtmr9
PTPN1
tns1a
Mtmr6
PTPN9
Ptpn5
Cdc14b
PTPN5
SSH1
MTMR9
Ptpn9
Cdc14a
Mtm1
Ptpn6
SKI3
E7F716
PTPN11
Mtmr7
PTPN22
CDC14A
STYXL2
MTMR7
Ptp36E
Dmel\CG14411
mtmr8
Ssh1
Ptpmeg2
PTPN6
Dmel\CG5026
Ptpn22
Ptprr

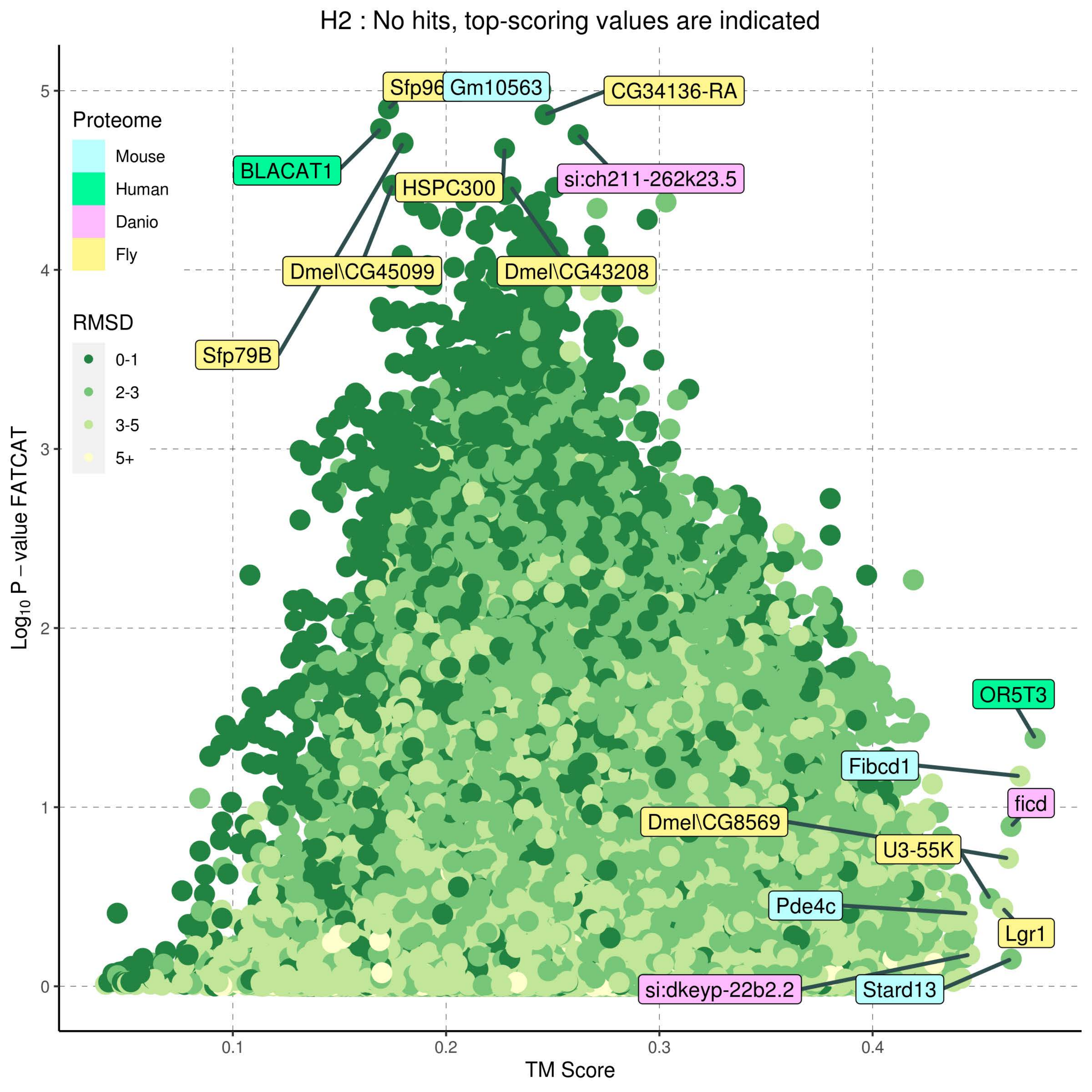

H2 : No hits, top-scoring values are indicated
Proteome
Mouse
Human
Danio
Fly
RMSD
0-1
2-3
3-5
5+
Sfp96
Gm10563
CG34136-RA
BLACAT1
si:ch211-262k23.5
HSPC300
Dmel\CG45099
Dmel\CG43208
Sfp79B
OR5T3
Fibcd1
ficd
Dmel\CG8569
U3-55K
Pde4c
Lgr1
si:dkeyp-22b2.2
Stard13
$Log_{10}$ P − value FATCAT
TM Score
0
1
2
3
4
5
0.1
0.2
0.3
0.4

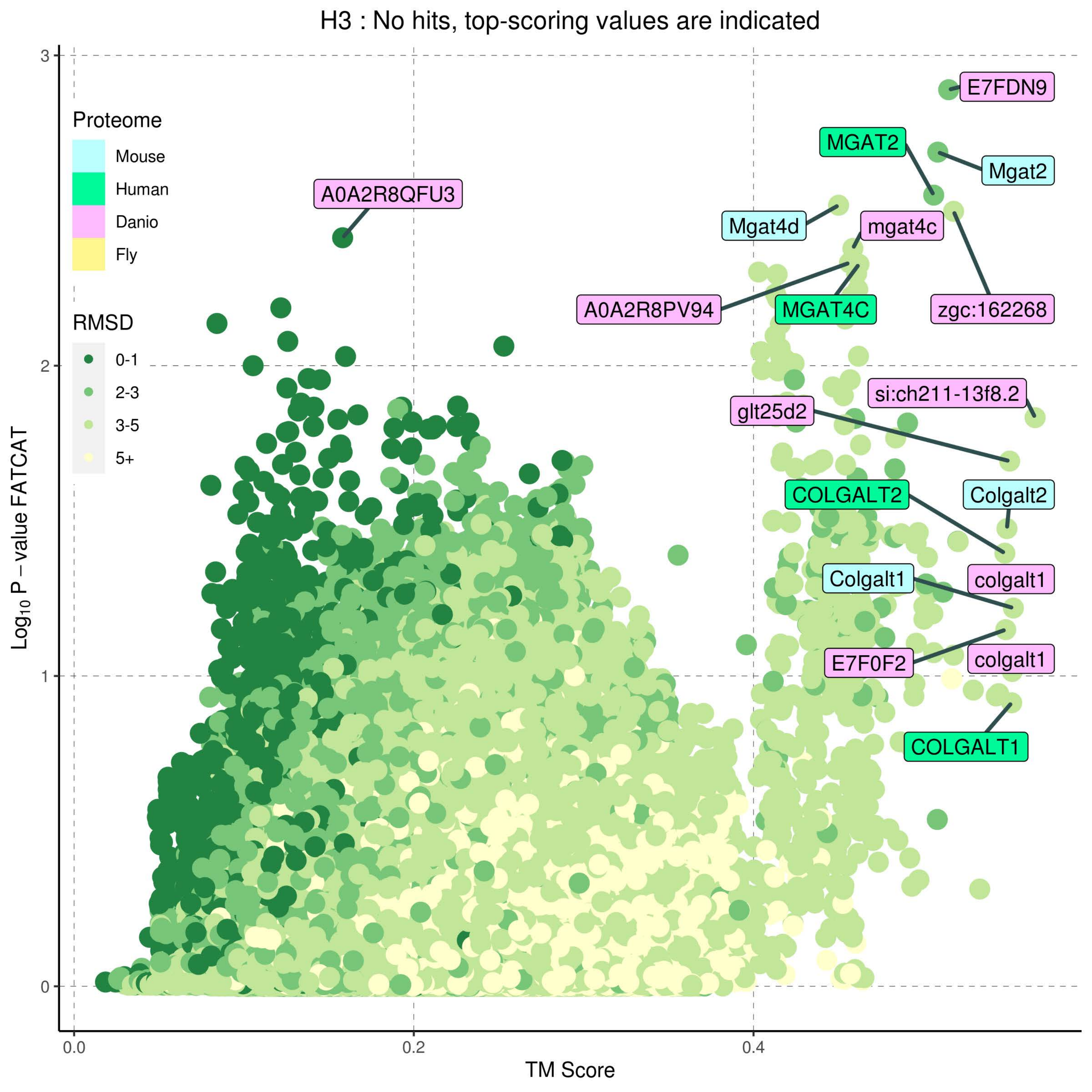
H3 : No hits, top-scoring values are indicated
Proteome
Mouse
Human
Danio
Fly
RMSD
0-1
2-3
3-5
5+
E7FDN9
MGAT2
Mgat2
A0A2R8QFU3
Mgat4d
mgat4c
A0A2R8PV94
MGAT4C
zgc:162268
si:ch211-13f8.2
glt25d2
COLGALT2
Colgalt2
Colgalt1
colgalt1
E7F0F2
colgalt1
COLGALT1
Log10 P − value FATCAT
TM Score
0.0
0.2
0.4
0
1
2
3

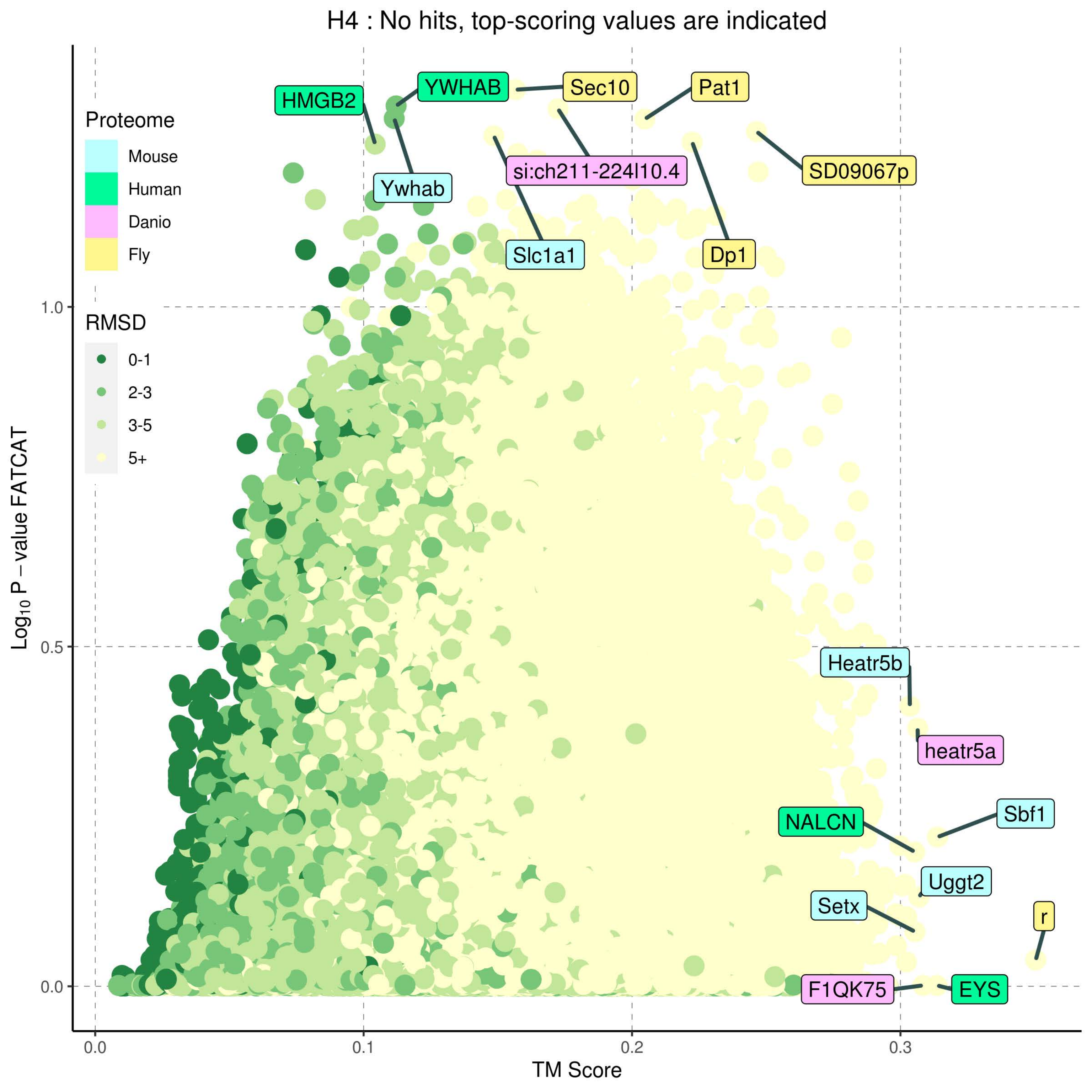
H4 : No hits, top-scoring values are indicated
Proteome
Mouse
Human
Danio
Fly
RMSD
0-1
2-3
3-5
5+
HMGB2
YWHAB
Sec10
Pat1
Ywhab
si:ch211-224l10.4
SD09067p
Slc1a1
Dp1
Heatr5b
heatr5a
NALCN
Sbf1
Uggt2
Setx
r
F1QK75
EYS
1.0
0.5
0.0
0.0
0.1
0.2
0.3
Log10 P − value FATCAT
TM Score

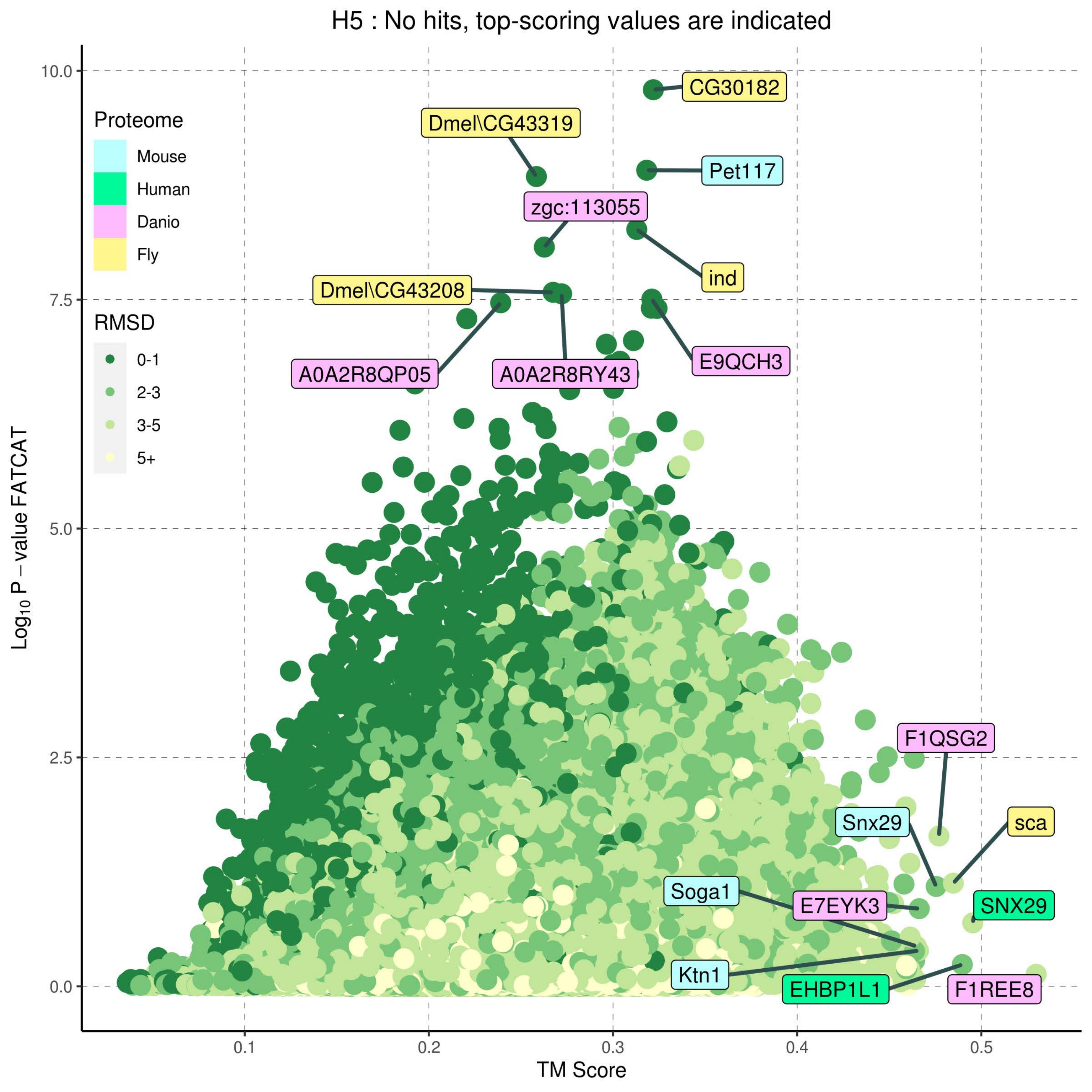

H5 : No hits, top-scoring values are indicated
Proteome
Mouse
Human
Danio
Fly
RMSD
0-1
2-3
3-5
5+
$Log_{10}$ P – value FATCAT
TM Score
10.0
7.5
5.0
2.5
0.0
0.1
0.2
0.3
0.4
0.5
CG30182
Dmel\CG43319
Pet117
zgc:113055
ind
Dmel\CG43208
A0A2R8QP05
A0A2R8RY43
E9QCH3
F1QSG2
Snx29
sca
Soga1
E7EYK3
SNX29
Ktn1
EHBP1L1
F1REE8

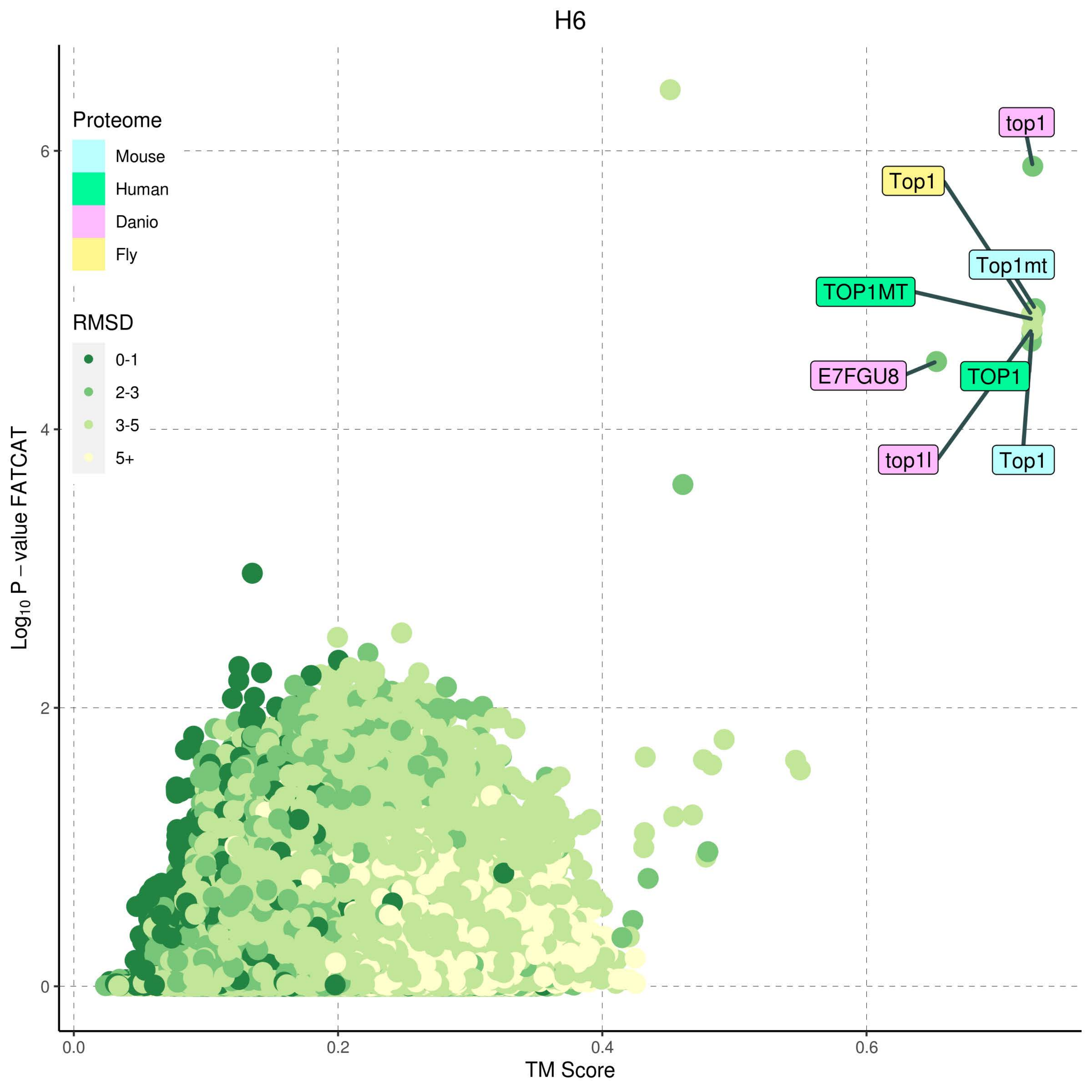

H6
Proteome
Mouse
Human
Danio
Fly
RMSD
0-1
2-3
3-5
5+
top1
Top1
Top1mt
TOP1MT
E7FGU8
TOP1
top1l
Top1
0
2
4
6
0.0
0.2
0.4
0.6
TM Score
Log10 P – value FATCAT

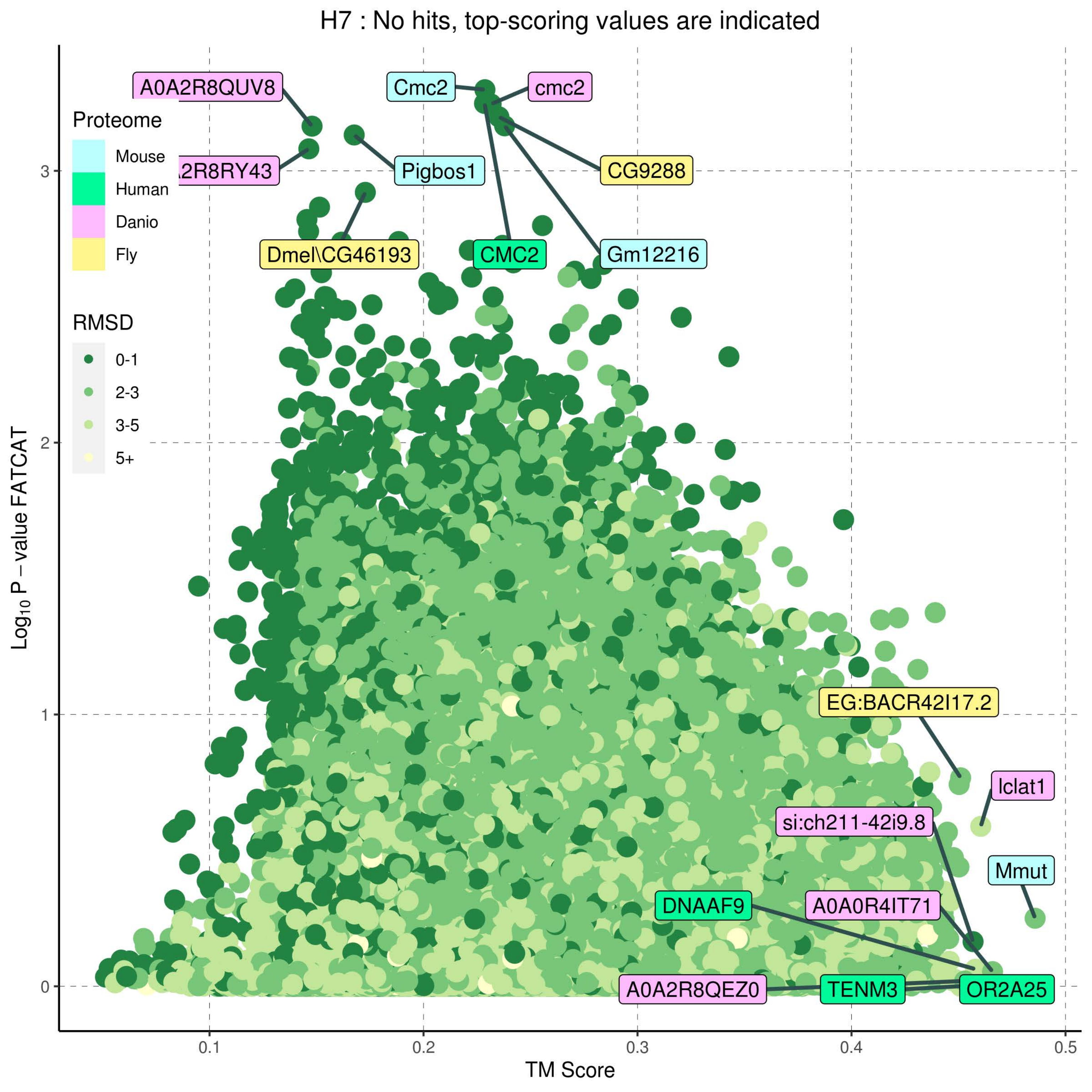

H7 : No hits, top-scoring values are indicated
Proteome
Mouse
Human
Danio
Fly
RMSD
0-1
2-3
3-5
5+
A0A2R8QUV8
2R8RY43
Cmc2
cmc2
Pigbos1
CG9288
Dmel\CG46193
CMC2
Gm12216
EG:BACR42I17.2
lclat1
si:ch211-42i9.8
Mmut
DNAAF9
A0A0R4IT71
A0A2R8QEZ0
TENM3
OR2A25
Log10 P – value FATCAT
TM Score
0
1
2
3
0.1
0.2
0.3
0.4
0.5

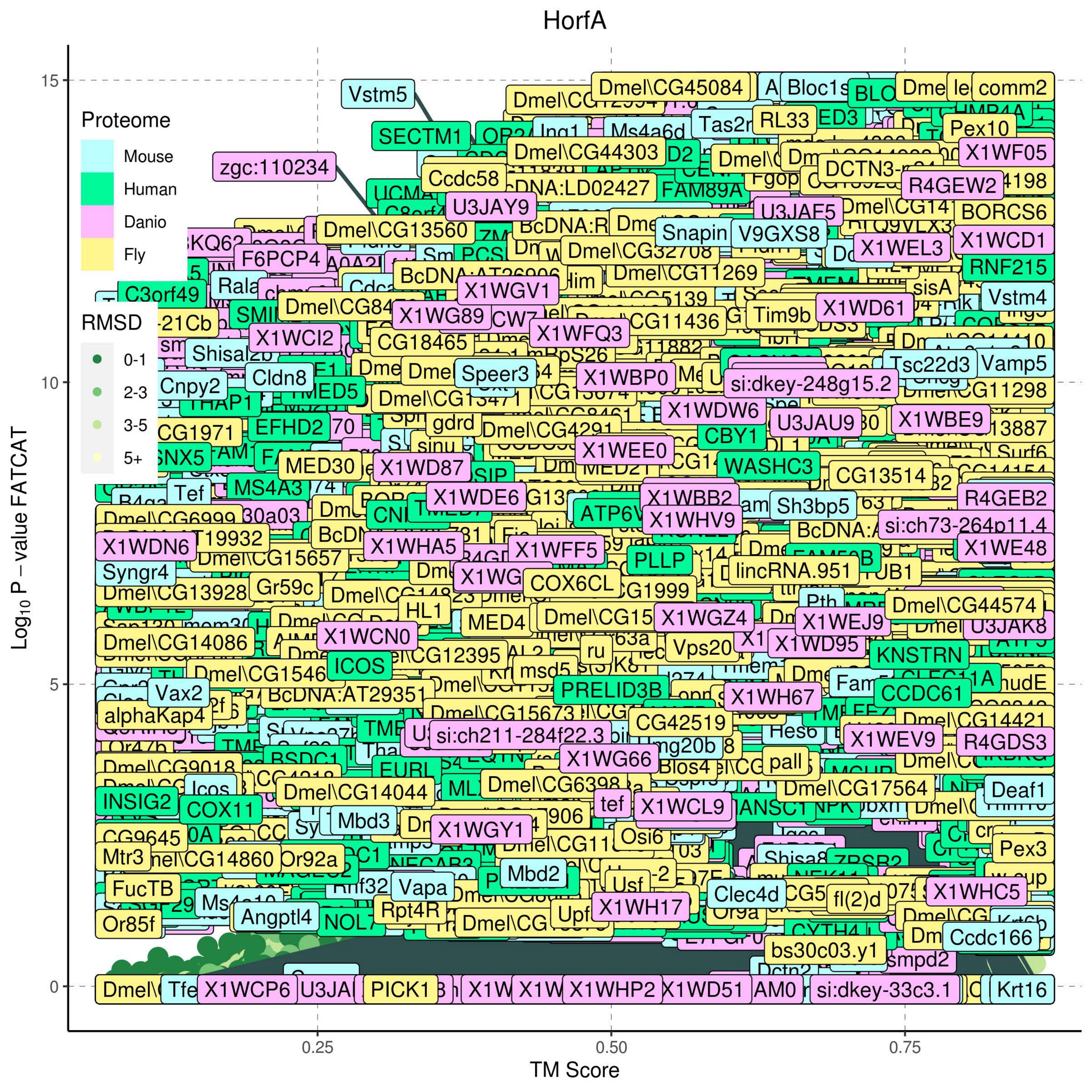
HorfA
Proteome
Mouse
Human
Danio
Fly
RMSD
0-1
2-3
3-5
5+
15
10
5
0
0.25
0.50
0.75
TM Score
$Log_{10}$ P – value FATCAT

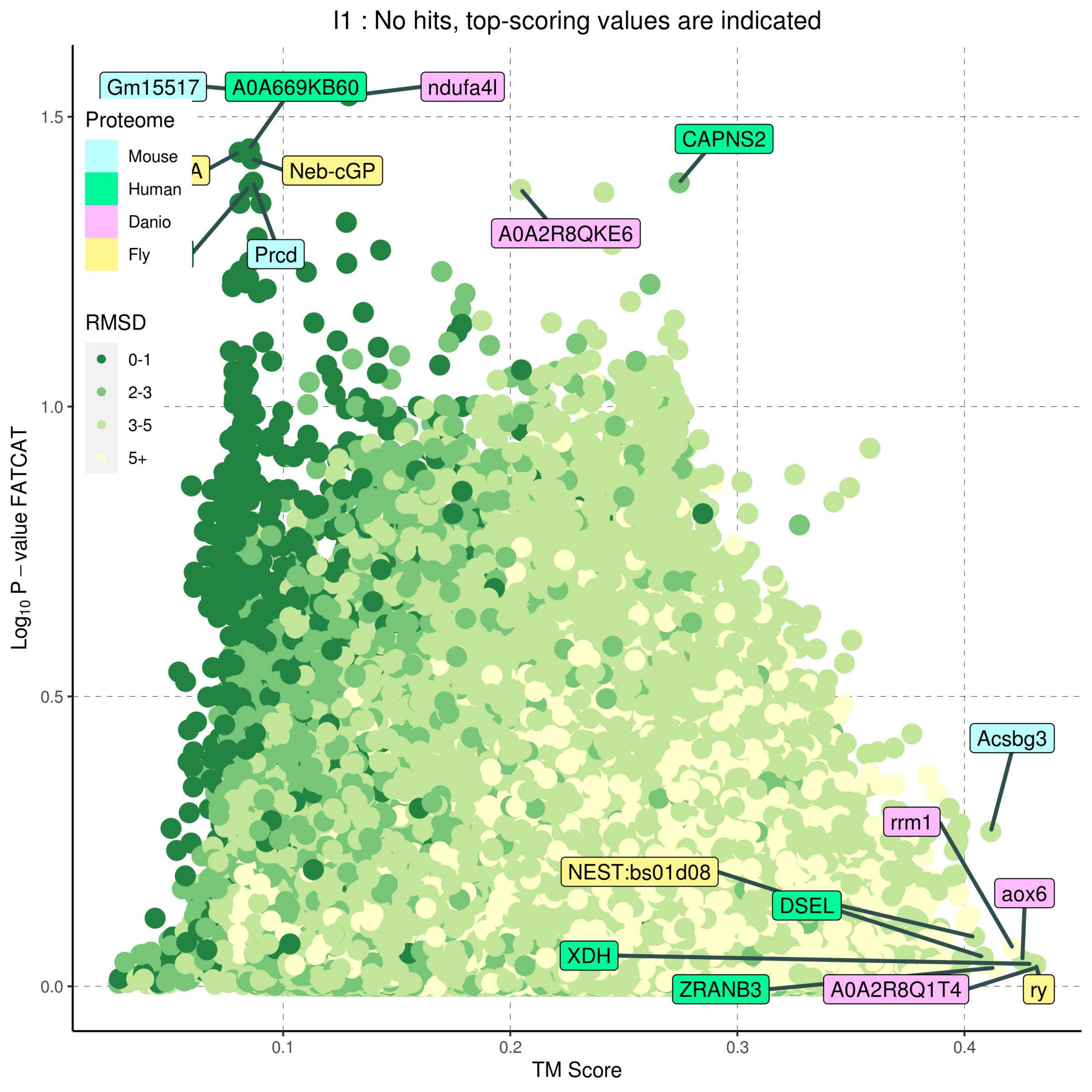

I1 : No hits, top-scoring values are indicated
Proteome
Mouse
Human
Danio
Fly
RMSD
0-1
2-3
3-5
5+
Gm15517
A0A669KB60
ndufa4l
CAPNS2
Neb-cGP
A0A2R8QKE6
Prcd
Acsbg3
rrm1
aox6
NEST:bs01d08
DSEL
XDH
ZRANB3
A0A2R8Q1T4
ry
Log10 P − value FATCAT
TM Score
0.0
0.5
1.0
1.5
0.1
0.2
0.3
0.4

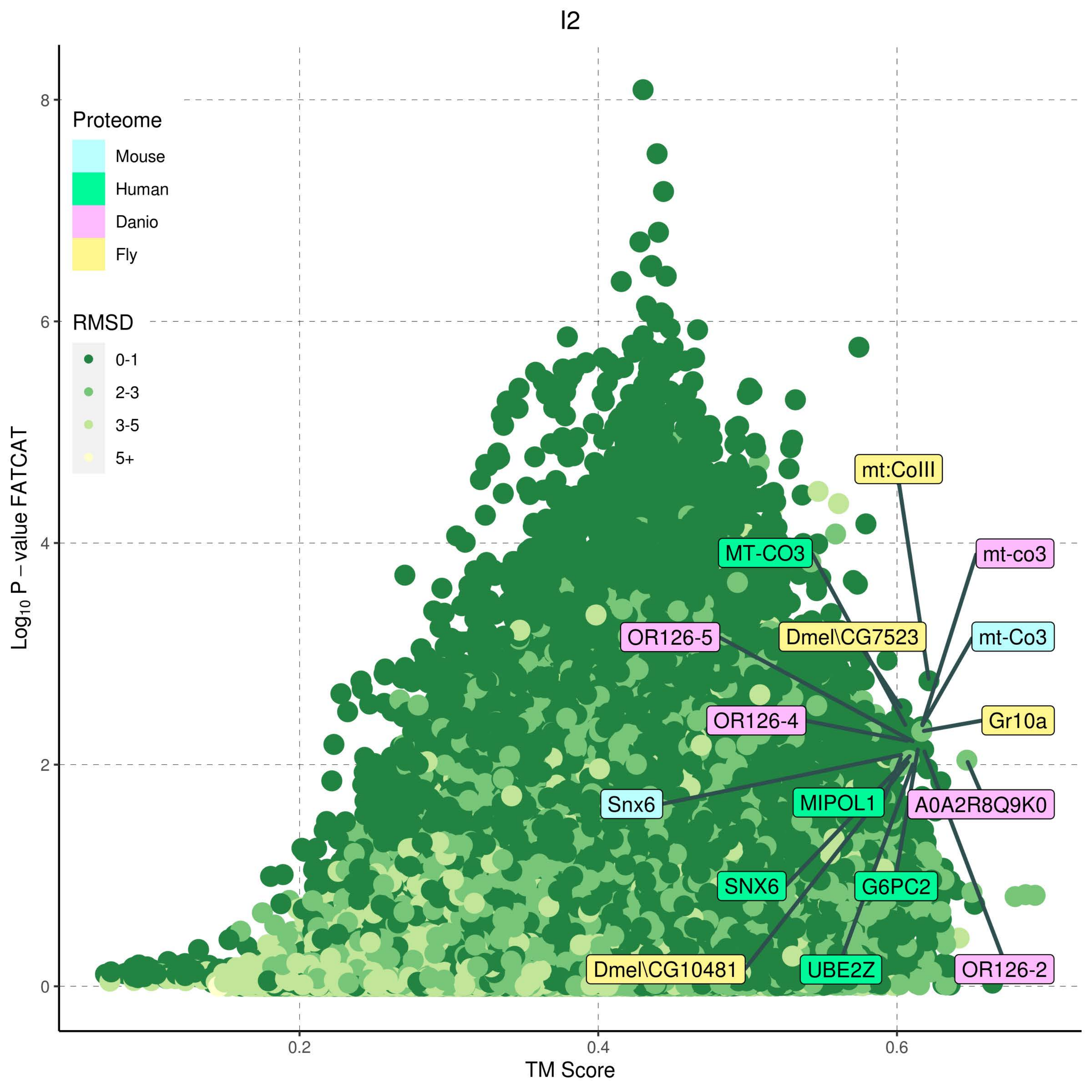

I2
Proteome
Mouse
Human
Danio
Fly
RMSD
0-1
2-3
3-5
5+
Log10 P − value FATCAT
TM Score
0
2
4
6
8
0.2
0.4
0.6
mt:CoIII
MT-CO3
mt-co3
OR126-5
Dmel\CG7523
mt-Co3
OR126-4
Gr10a
Snx6
MIPOL1
A0A2R8Q9K0
SNX6
G6PC2
Dmel\CG10481
UBE2Z
OR126-2

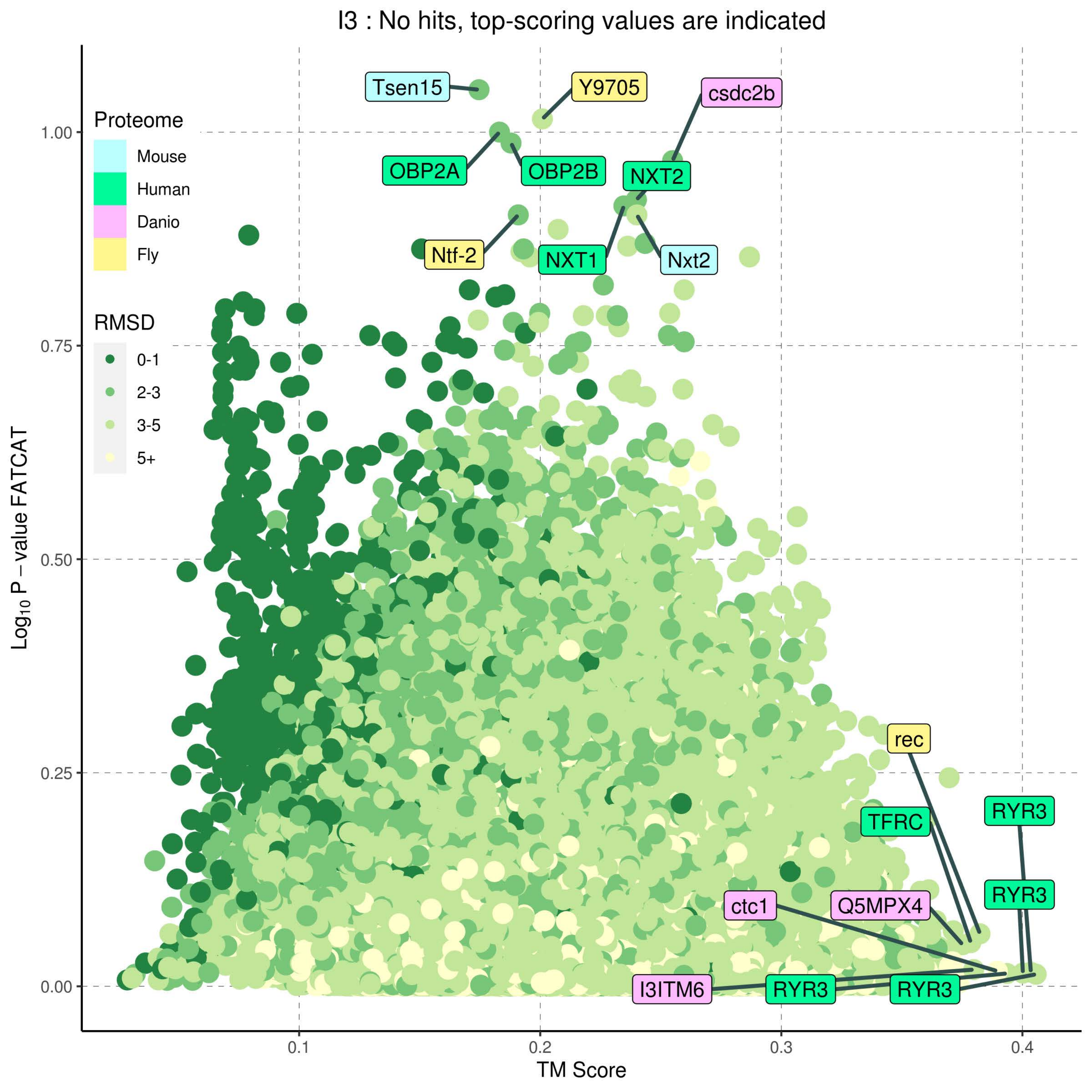
I3 : No hits, top-scoring values are indicated
Proteome
Mouse
Human
Danio
Fly
RMSD
0-1
2-3
3-5
5+
Tsen15
Y9705
csdc2b
OBP2A
OBP2B
NXT2
Ntf-2
NXT1
Nxt2
rec
TFRC
RYR3
RYR3
ctc1
Q5MPX4
I3ITM6
RYR3
RYR3
1.00
0.75
0.50
0.25
0.00
0.1
0.2
0.3
0.4
Log$_{10}$ P – value FATCAT
TM Score

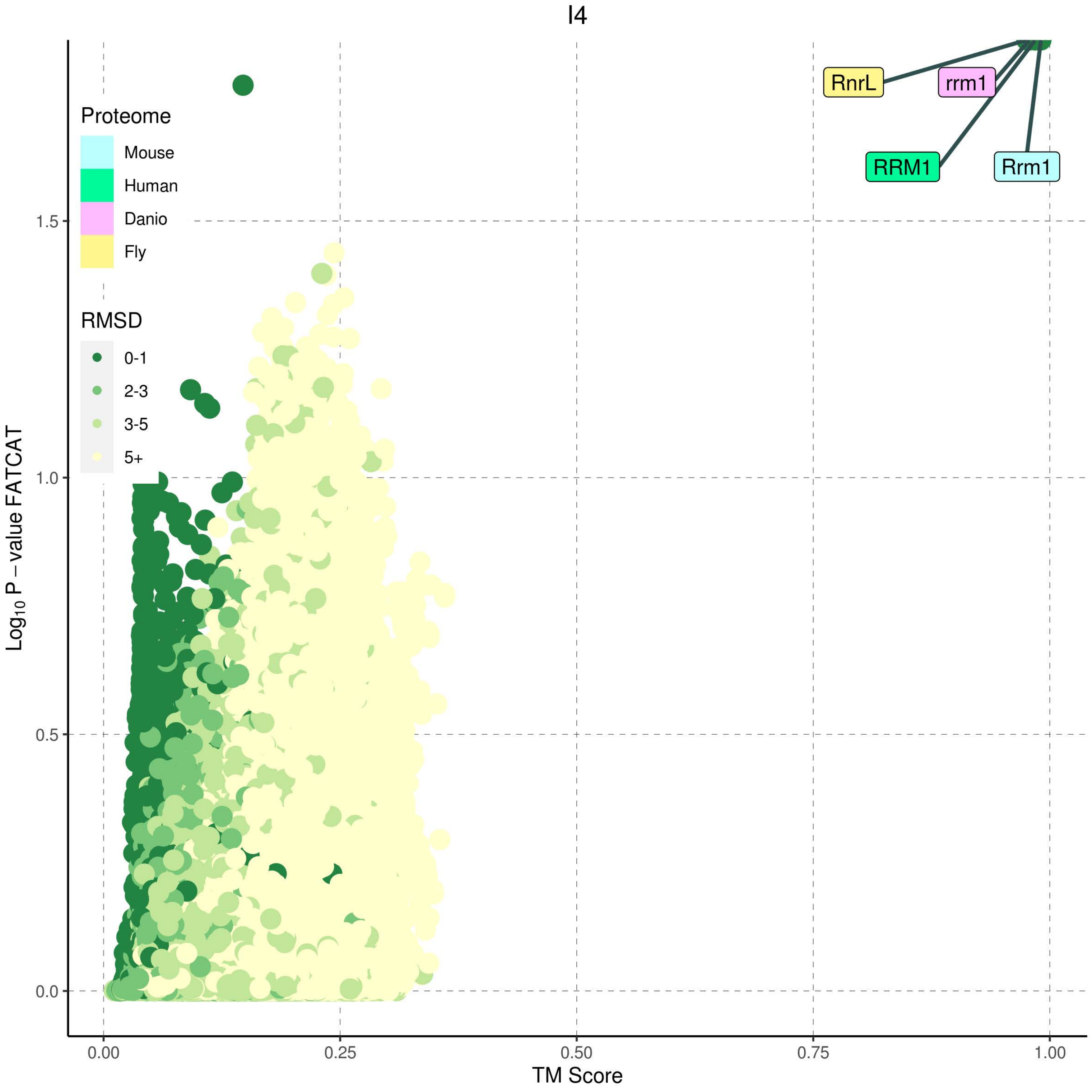

I4
Proteome
Mouse
Human
Danio
Fly
RMSD
0-1
2-3
3-5
5+
RnrL
rrm1
RRM1
Rrm1
$Log_{10}$ P − value FATCAT
1.5
1.0
0.5
0.0
TM Score
0.00
0.25
0.50
0.75
1.00

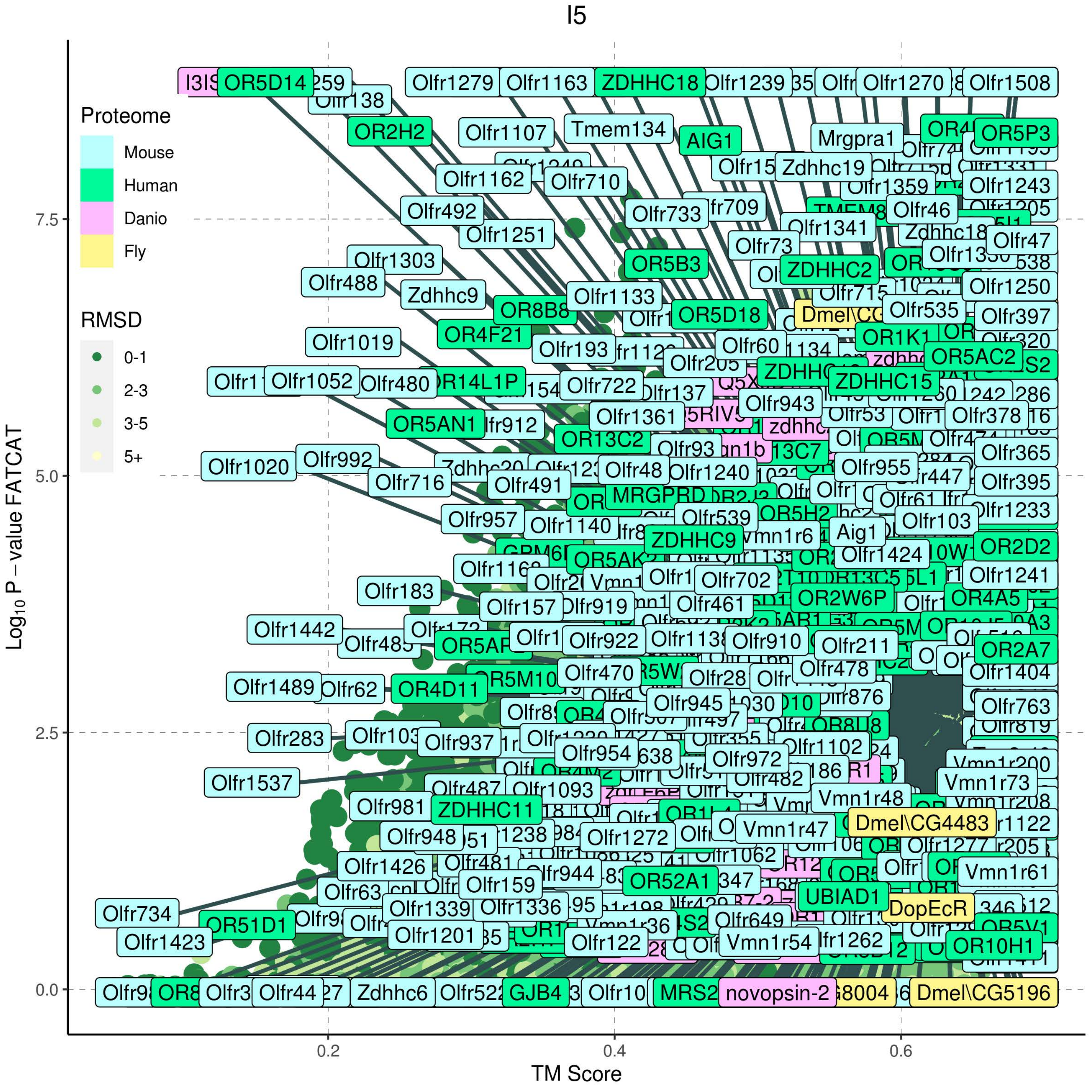
I5
Proteome
Mouse
Human
Danio
Fly
RMSD
0-1
2-3
3-5
5+
Log10 P − value FATCAT
TM Score
0.0
2.5
5.0
7.5
0.2
0.4
0.6

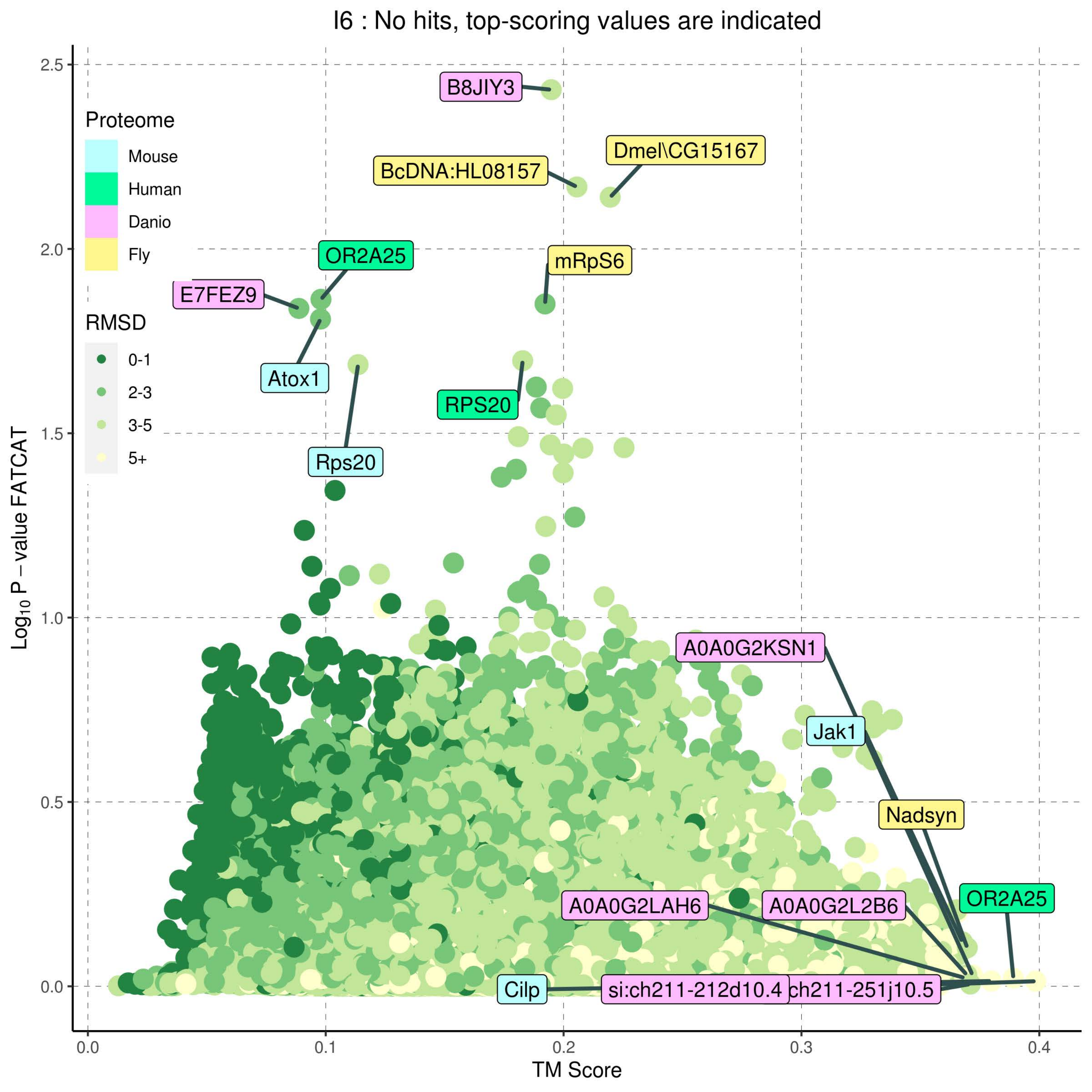

I6 : No hits, top-scoring values are indicated
Proteome
Mouse
Human
Danio
Fly
RMSD
0-1
2-3
3-5
5+
B8JIY3
Dmel\CG15167
BcDNA:HL08157
OR2A25
mRpS6
E7FEZ9
Atox1
RPS20
Rps20
A0A0G2KSN1
Jak1
Nadsyn
OR2A25
A0A0G2LAH6
A0A0G2L2B6
Cilp
si:ch211-212d10.4
ch211-251j10.5
2.5
2.0
1.5
1.0
0.5
0.0
0.0
0.1
0.2
0.3
0.4
TM Score
$Log_{10}$ P − value FATCAT

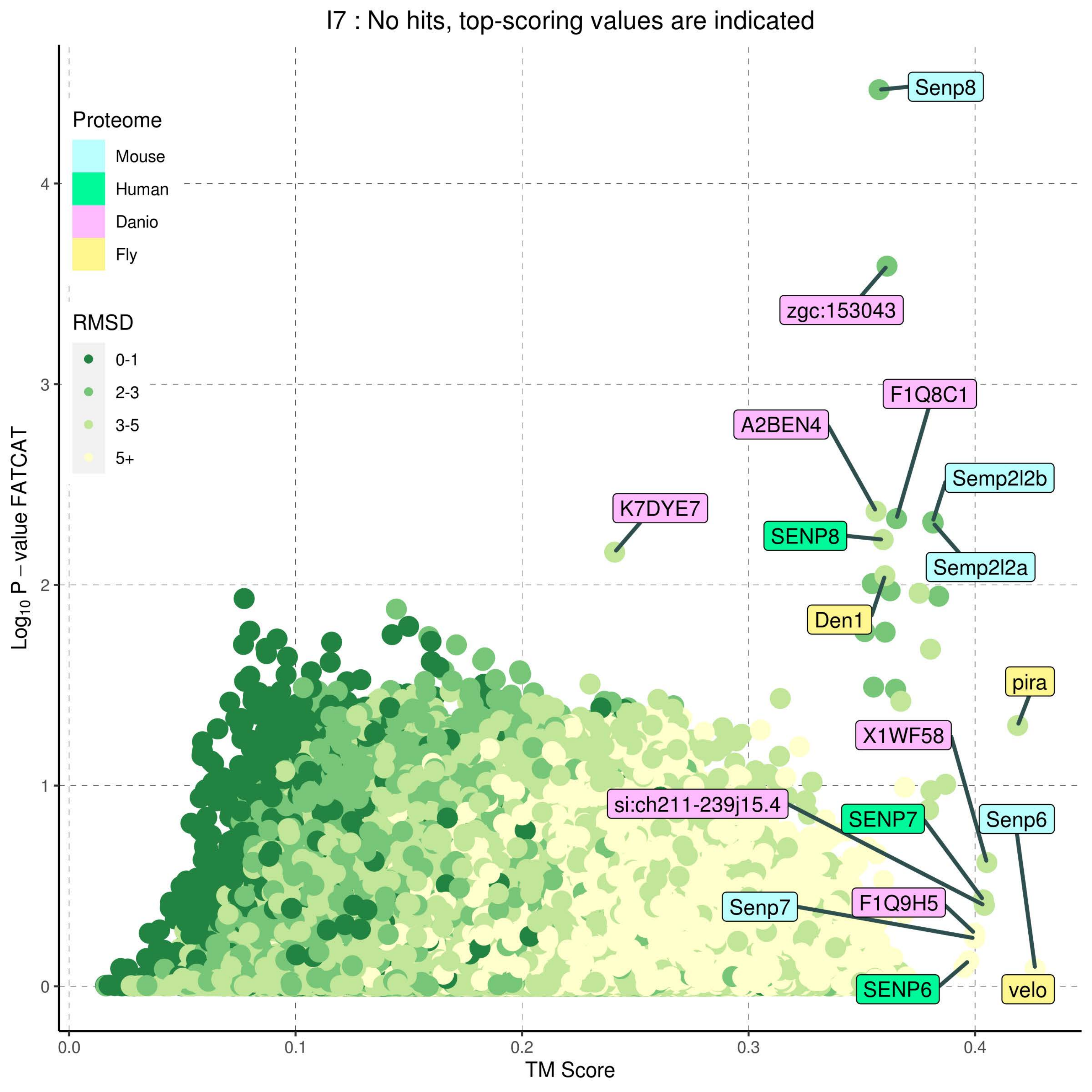
I7 : No hits, top-scoring values are indicated
Proteome
Mouse
Human
Danio
Fly
RMSD
0-1
2-3
3-5
5+
Senp8
zgc:153043
F1Q8C1
A2BEN4
Semp2l2b
K7DYE7
SENP8
Semp2l2a
Den1
pira
X1WF58
si:ch211-239j15.4
SENP7
Senp6
F1Q9H5
Senp7
SENP6
velo
Log10 P − value FATCAT
TM Score
0
1
2
3
4
0.0
0.1
0.2
0.3
0.4

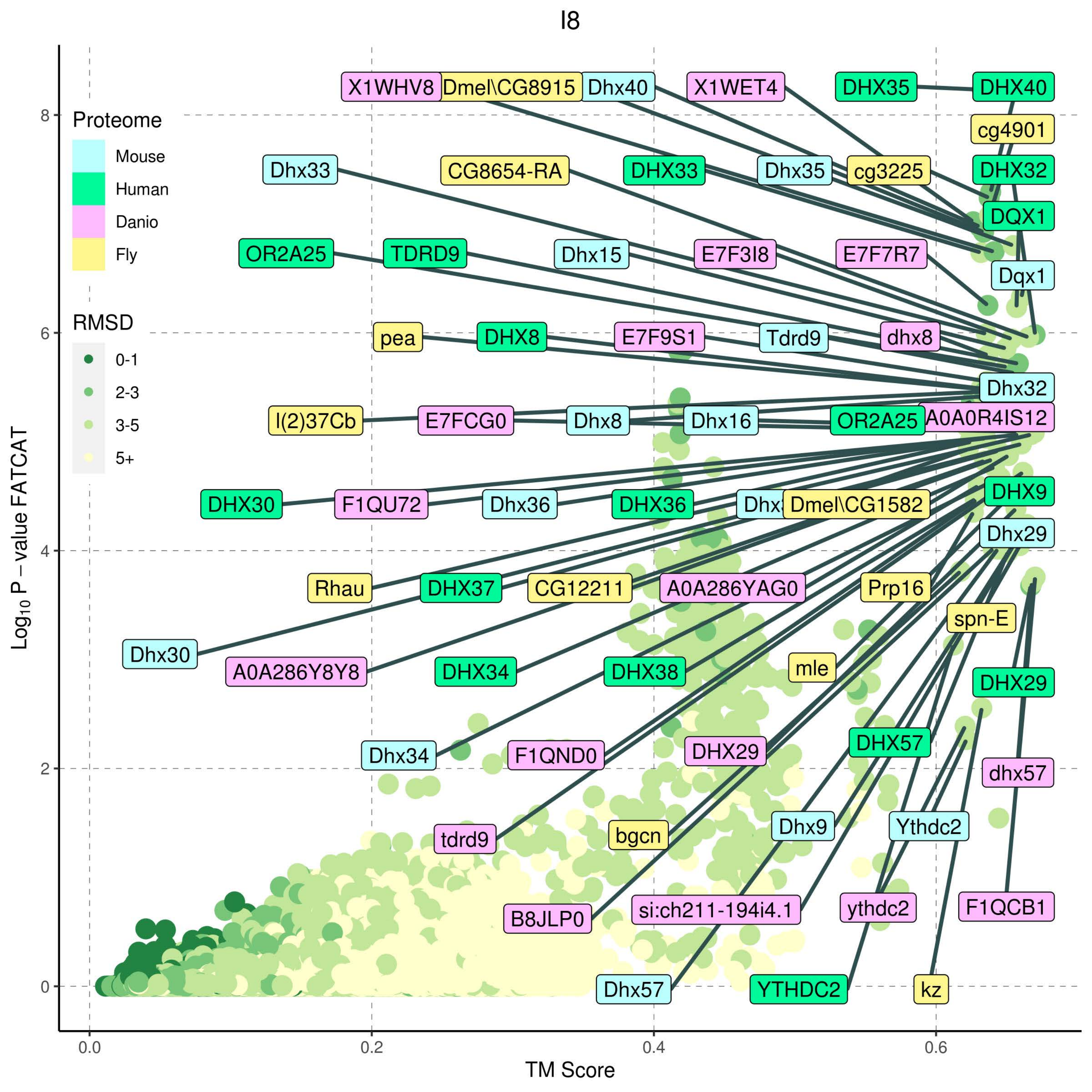

I8
Proteome
Mouse
Human
Danio
Fly
RMSD
0-1
2-3
3-5
5+
Log10 P – value FATCAT
TM Score
0
2
4
6
8
0.0
0.2
0.4
0.6
X1WHV8
Dmel\CG8915
Dhx40
X1WET4
DHX35
DHX40
cg4901
Dhx33
CG8654-RA
DHX33
Dhx35
cg3225
DHX32
DQX1
OR2A25
TDRD9
Dhx15
E7F3I8
E7F7R7
Dqx1
pea
DHX8
E7F9S1
Tdrd9
dhx8
Dhx32
l(2)37Cb
E7FCG0
Dhx8
Dhx16
OR2A25
A0A0R4IS12
DHX30
F1QU72
Dhx36
DHX36
Dmel\CG1582
DHX9
Dhx29
Rhau
DHX37
CG12211
A0A286YAG0
Prp16
spn-E
Dhx30
A0A286Y8Y8
DHX34
DHX38
mle
DHX29
Dhx34
F1QND0
DHX29
DHX57
dhx57
tdrd9
bgcn
Dhx9
Ythdc2
B8JLP0
si:ch211-194i4.1
ythdc2
F1QCB1
Dhx57
YTHDC2
kz

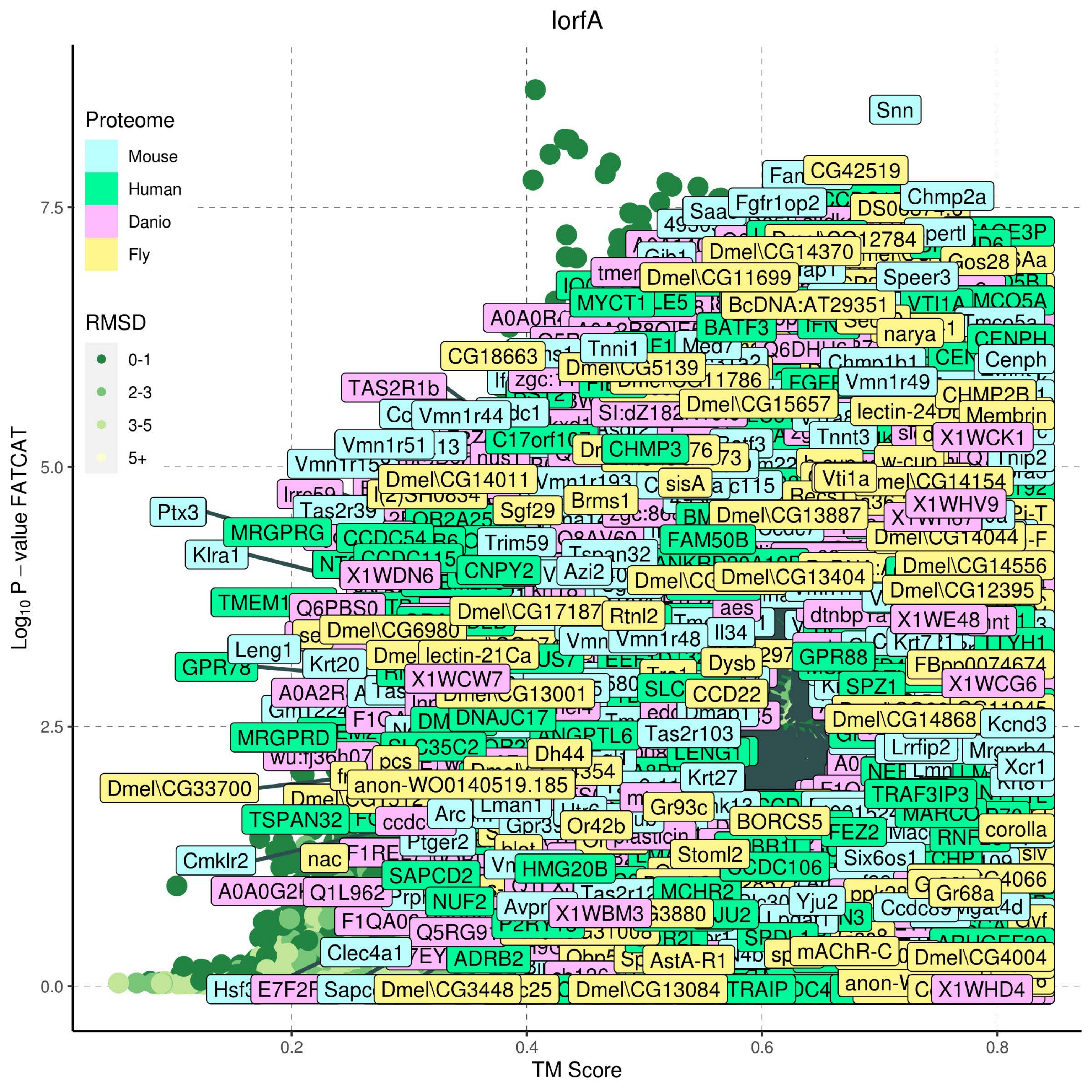
lorfA
Proteome
Mouse
Human
Danio
Fly
RMSD
0-1
2-3
3-5
5+
Log10 P – value FATCAT
0.0
2.5
5.0
7.5
TM Score
0.2
0.4
0.6
0.8
Snn
CG42519
Chmp2a
Fgfr1op2
Dmel\CG14370
Dmel\CG11699
Gos28
Speer3
BcDNA:AT29351
MYCT1
BATF3
narya
Cenph
CG18663
Tnni1
Dmel\CG5139
Vmn1r49
TAS2R1b
Dmel\CG15657
CHMP2B
Membrin
Vmn1r44
Vmn1r51
C17orf107
CHMP3
Tnnt3
X1WCK1
Tnip2
Dmel\CG14011
sisA
Vti1a
X1WHV9
Ptx3
Sgf29
Brms1
Dmel\CG13887
MRGPRG
Trim59
FAM50B
Dmel\CG14044
Klra1
X1WDN6
CNPY2
Azi2
Dmel\CG14556
Dmel\CG13404
Dmel\CG12395
Q6PBS0
Dmel\CG17187
Rtnl2
aes
X1WE48
Dmel\CG6980
Leng1
Vmn1r48
Il34
lectin-21Ca
GPR88
Krt20
Dysb
FBpp0074674
X1WCW7
SPZ1
X1WCG6
Dmel\CG13001
CCD22
DNAJC17
Dmel\CG14868
Kcnd3
MRGPRD
Tas2r103
ANGPTL6
SLC35C2
Dh44
Lrrfip2
Xcr1
Dmel\CG33700
anon-WO0140519.185
Krt27
TRAF3IP3
TSPAN32
Arc
Lman1
Gr93c
Or42b
BORCS5
FEZ2
corolla
Cmklr2
nac
Ptger2
Stoml2
HMG20B
Six6os1
SAPCD2
CCDC106
A0A0G2I
Q1L962
NUF2
MCHR2
Yju2
Gr68a
X1WBM3
Clec4a1
ADRB2
AstA-R1
mAChR-C
Dmel\CG4004
Dmel\CG3448
Dmel\CG13084
TRAIP
X1WHD4

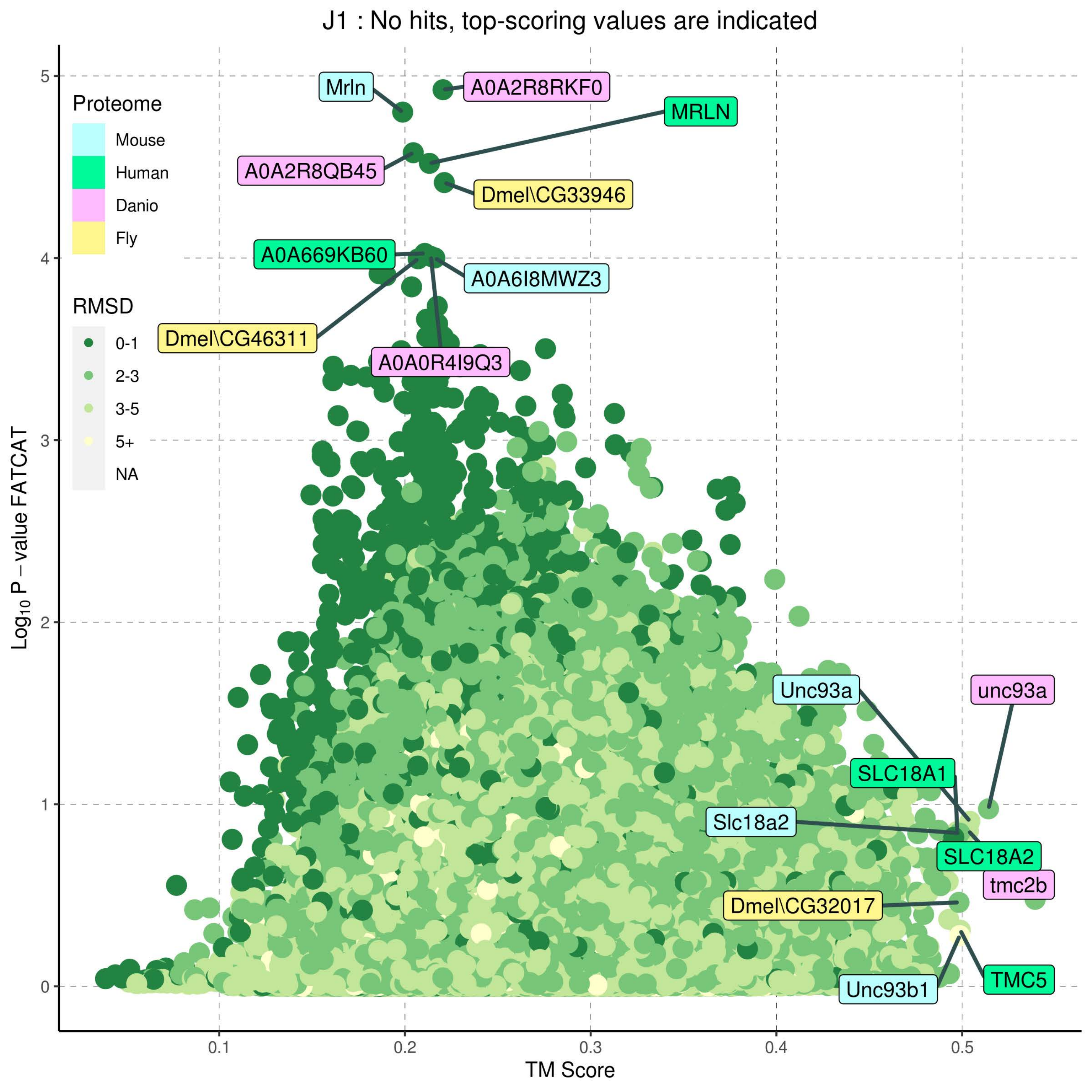
J1 : No hits, top-scoring values are indicated
Proteome
Mouse
Human
Danio
Fly
RMSD
0-1
2-3
3-5
5+
NA
Mrln
A0A2R8RKF0
MRLN
A0A2R8QB45
Dmel\CG33946
A0A669KB60
A0A6I8MWZ3
Dmel\CG46311
A0A0R4I9Q3
Unc93a
unc93a
SLC18A1
Slc18a2
SLC18A2
tmc2b
Dmel\CG32017
TMC5
Unc93b1
Log10 P – value FATCAT
TM Score
0
1
2
3
4
5
0.1
0.2
0.3
0.4
0.5

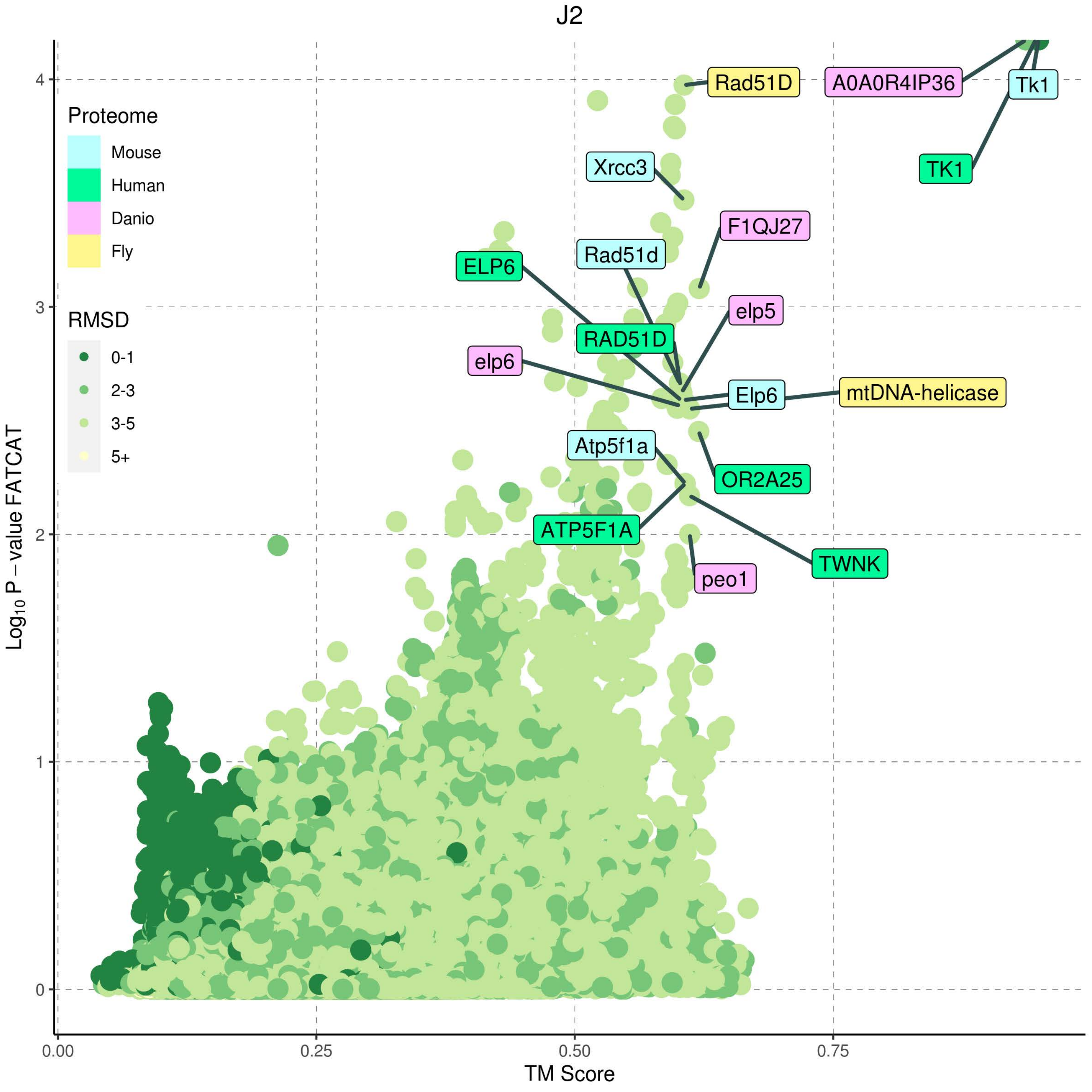

J2
Proteome
Mouse
Human
Danio
Fly
RMSD
0-1
2-3
3-5
5+
Rad51D
A0A0R4IP36
Tk1
TK1
Xrcc3
F1QJ27
Rad51d
ELP6
elp5
RAD51D
elp6
Elp6
mtDNA-helicase
Atp5f1a
OR2A25
ATP5F1A
TWNK
peo1
Log10 P − value FATCAT
TM Score
0
1
2
3
4
0.00
0.25
0.50
0.75

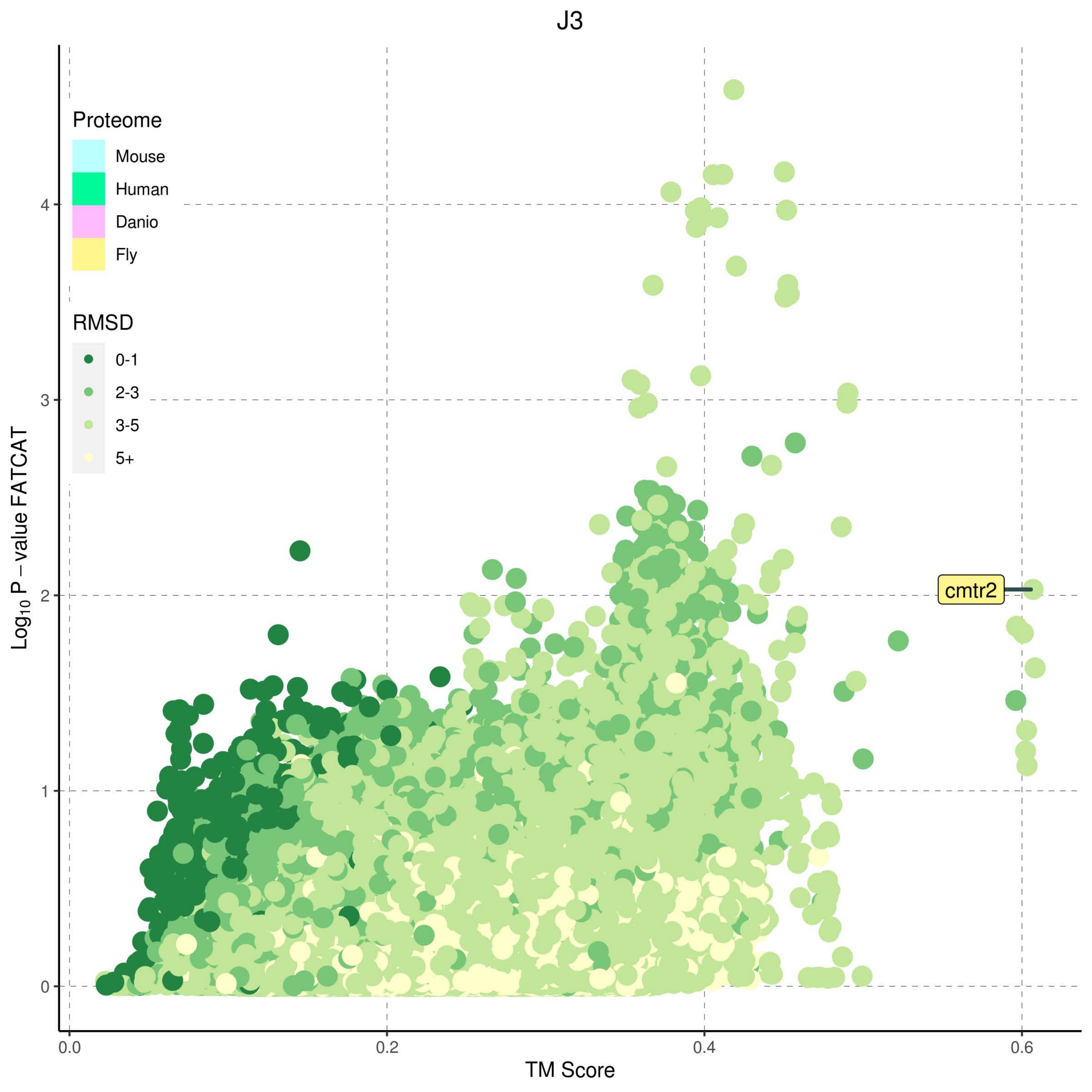
J3
Proteome
Mouse
Human
Danio
Fly
RMSD
0-1
2-3
3-5
5+
cmtr2
Log10 P − value FATCAT
TM Score
0
1
2
3
4
0.0
0.2
0.4
0.6

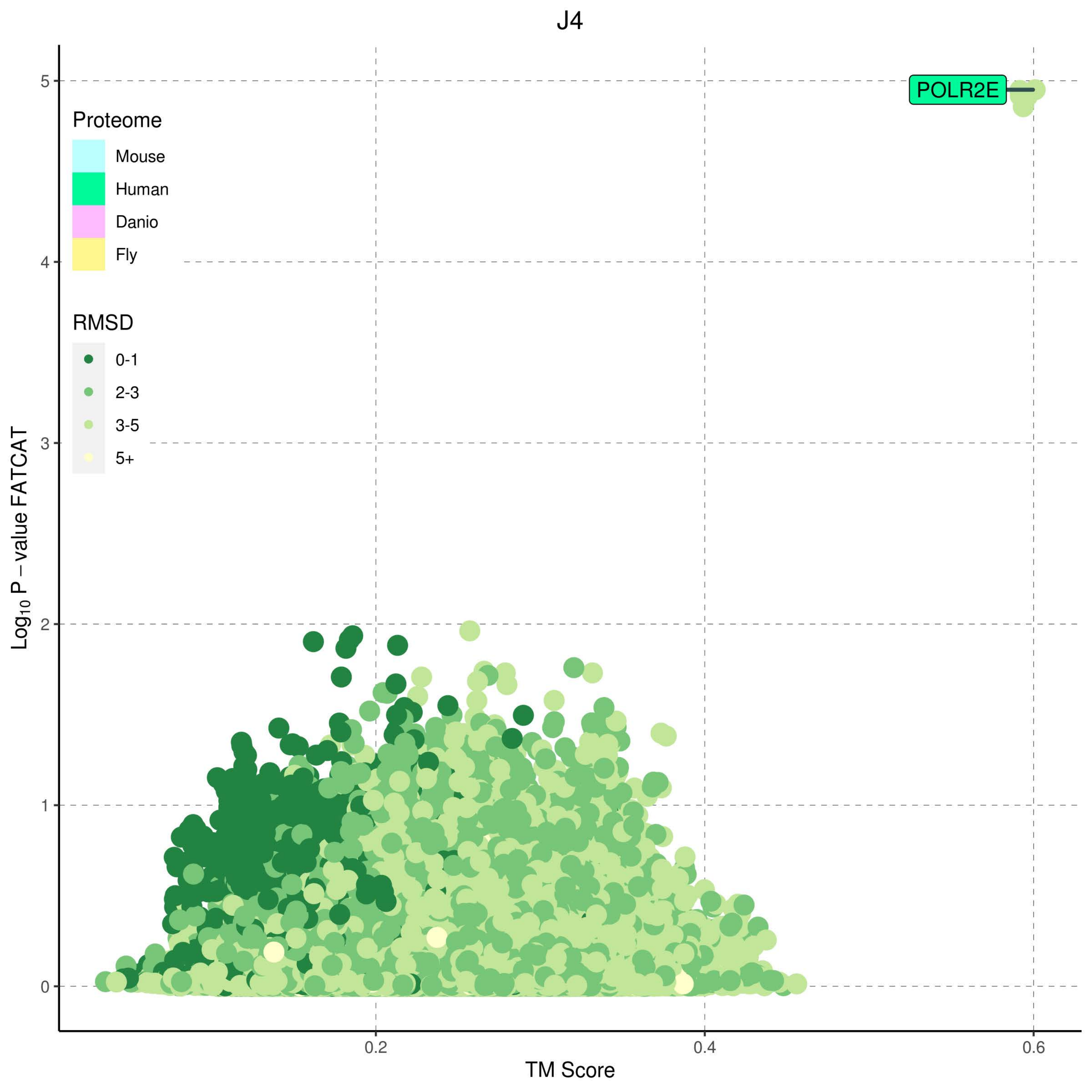

J4
POLR2E
Proteome
Mouse
Human
Danio
Fly
RMSD
0-1
2-3
3-5
5+
Log10 P – value FATCAT
0
1
2
3
4
5
0.2
0.4
0.6
TM Score

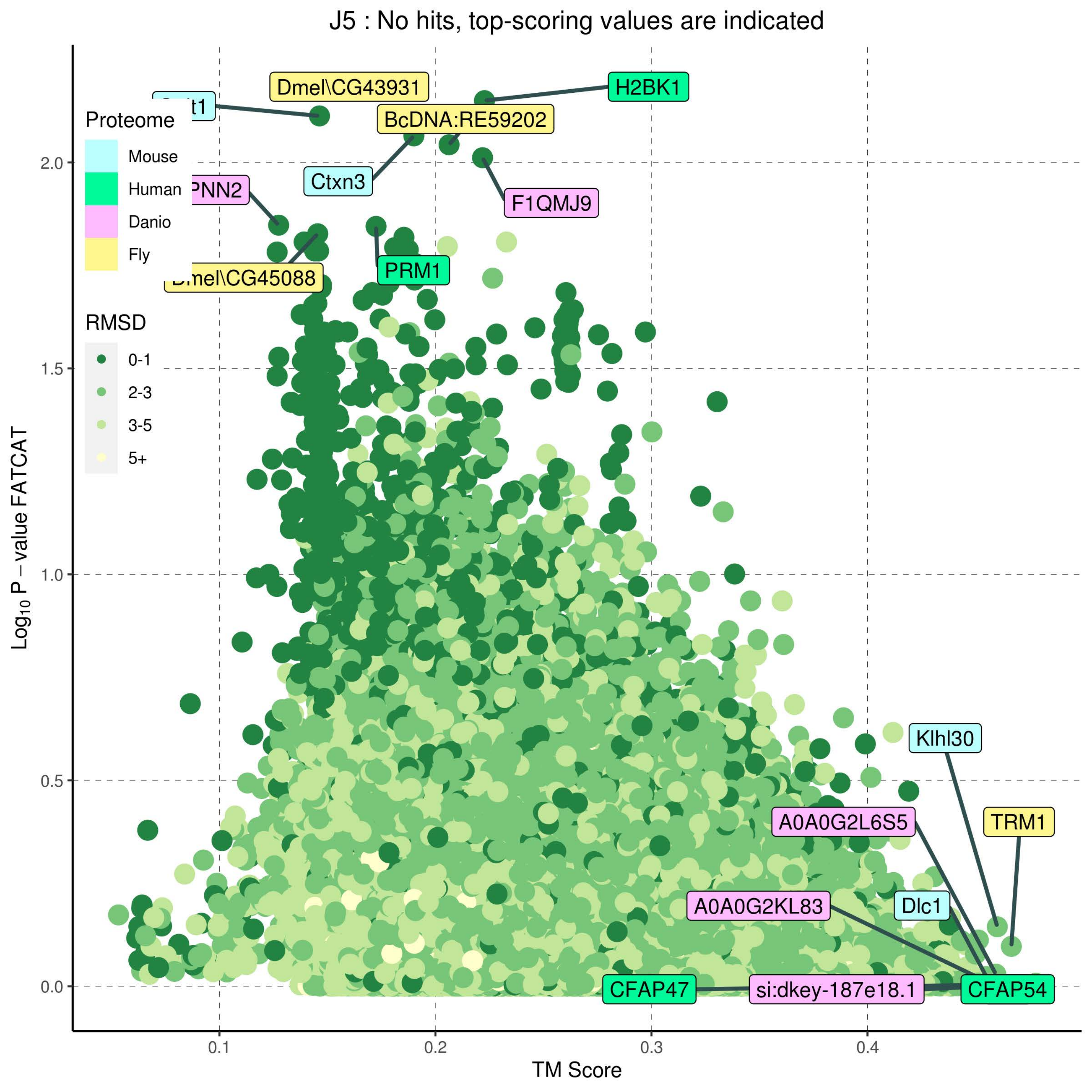

J5 : No hits, top-scoring values are indicated
Proteome
Mouse
Human
Danio
Fly
RMSD
0-1
2-3
3-5
5+
Dmel\CG43931
H2BK1
BcDNA:RE59202
Ctxn3
F1QMJ9
PNN2
PRM1
Dmel\CG45088
Klhl30
TRM1
A0A0G2L6S5
A0A0G2KL83
Dlc1
CFAP47
si:dkey-187e18.1
CFAP54
Log10 P − value FATCAT
TM Score
0.0
0.5
1.0
1.5
2.0
0.1
0.2
0.3
0.4

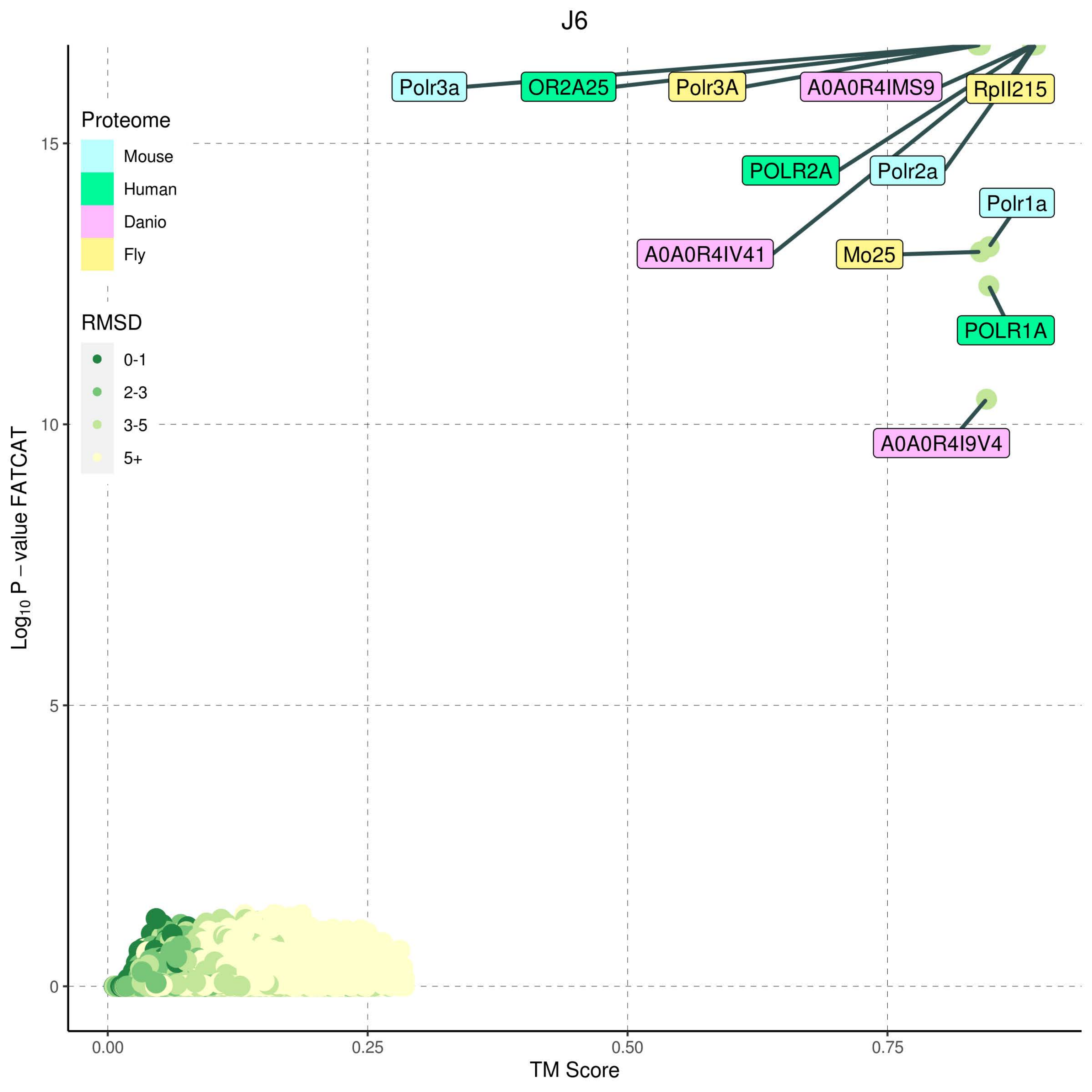

J6
Proteome
Mouse
Human
Danio
Fly
RMSD
0-1
2-3
3-5
5+
Polr3a
OR2A25
Polr3A
A0A0R4IMS9
RpII215
POLR2A
Polr2a
Polr1a
A0A0R4IV41
Mo25
POLR1A
A0A0R4I9V4
15
10
5
0
0.00
0.25
0.50
0.75
TM Score
Log10 P − value FATCAT

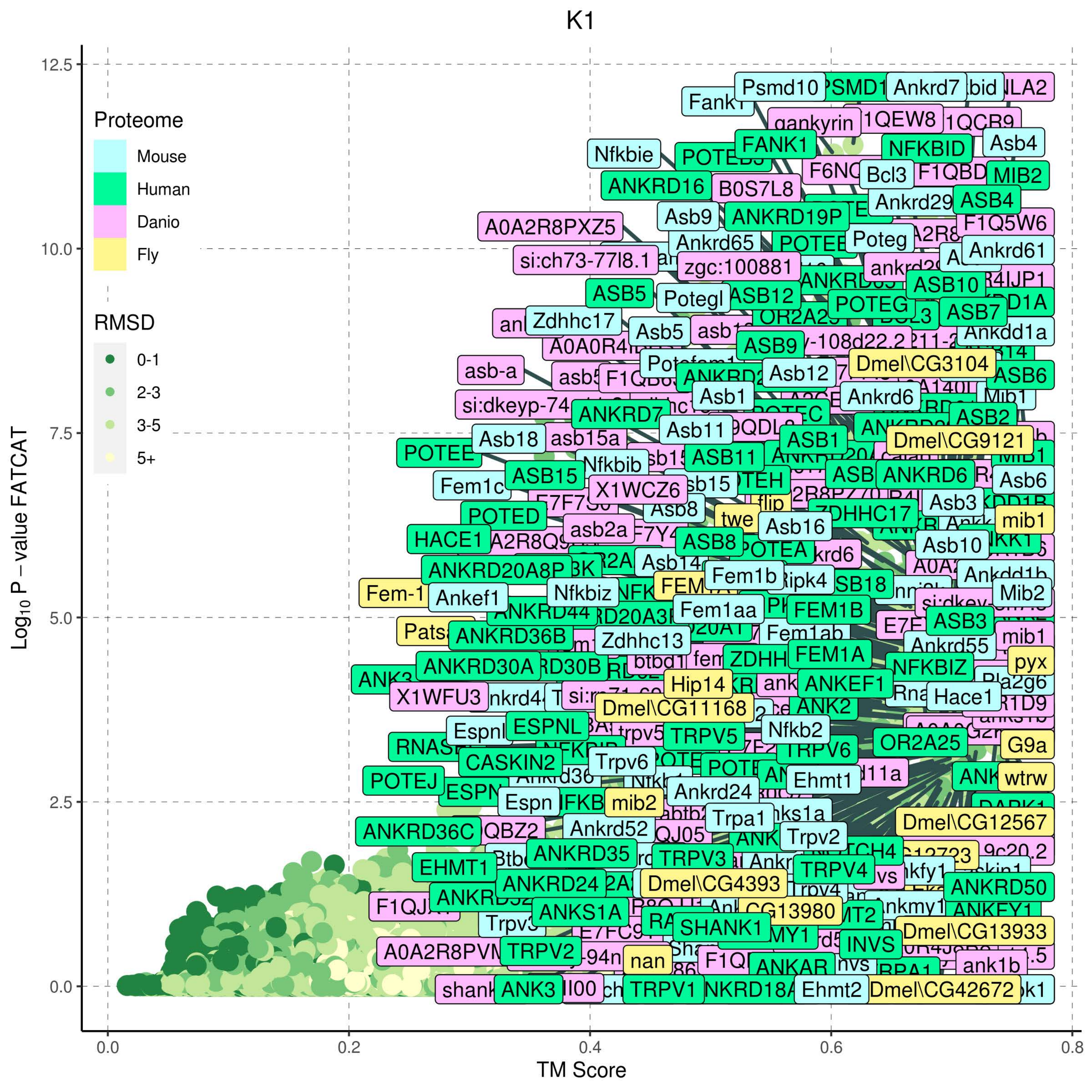

K1
Proteome
Mouse
Human
Danio
Fly
RMSD
0-1
2-3
3-5
5+
Log10 P − value FATCAT
TM Score
0.0
2.5
5.0
7.5
10.0
12.5
0.0
0.2
0.4
0.6
0.8
Psmd10
Ankrd7
Fank1
gankyrin
FANK1
NFKBID
Asb4
Nfkbie
Bcl3
MIB2
ANKRD16
B0S7L8
Ankrd29
ASB4
A0A2R8PXZ5
Asb9
ANKRD19P
F1Q5W6
Ankrd65
Poteg
Ankrd61
si:ch73-77l8.1
zgc:100881
ASB5
Potegl
ASB12
ASB10
POTEG
ASB7
Zdhhc17
Asb5
Ankdd1a
ASB9
Dmel\CG3104
Asb12
ASB6
asb-a
Asb1
Ankrd6
ANKRD7
ASB2
Asb11
ASB1
Asb18
asb15a
Dmel\CG9121
POTEE
Nfkbib
ASB11
ANKRD6
ASB15
Asb6
Fem1c
X1WCZ6
Asb3
POTED
Asb8
twe
Asb16
ZDHHC17
mib1
HACE1
asb2a
ASB8
POTEA
Asb10
ANKRD20A8P
Asb14
Fem1b
Ripk4
Ankdd1b
Fem-1
Ankef1
Nfkbiz
Mib2
ANKRD44
Fem1aa
FEM1B
ASB3
ANKRD36B
Zdhhc13
Fem1ab
mib1
FEM1A
Ankrd55
ANKRD30A
NFKBIZ
pyx
Hip14
ANKEF1
X1WFU3
Dmel\CG11168
Hace1
ANK2
Espnl
ESPNL
Nfkb2
TRPV5
RNASEL
TRPV6
OR2A25
G9a
CASKIN2
Trpv6
Ehmt1
POTEJ
Ankrd24
wtrw
Espn
mib2
Trpa1
Dmel\CG12567
ANKRD36C
Ankrd52
Trpv2
ANKRD35
TRPV3
EHMT1
TRPV4
ANKRD24
Dmel\CG4393
ANKRD50
ANKS1A
Dmel\CG13933
SHANK1
INVS
A0A2R8PVM
TRPV2
nan
ank1b
ANK3
TRPV1
Ehmt2
Dmel\CG42672

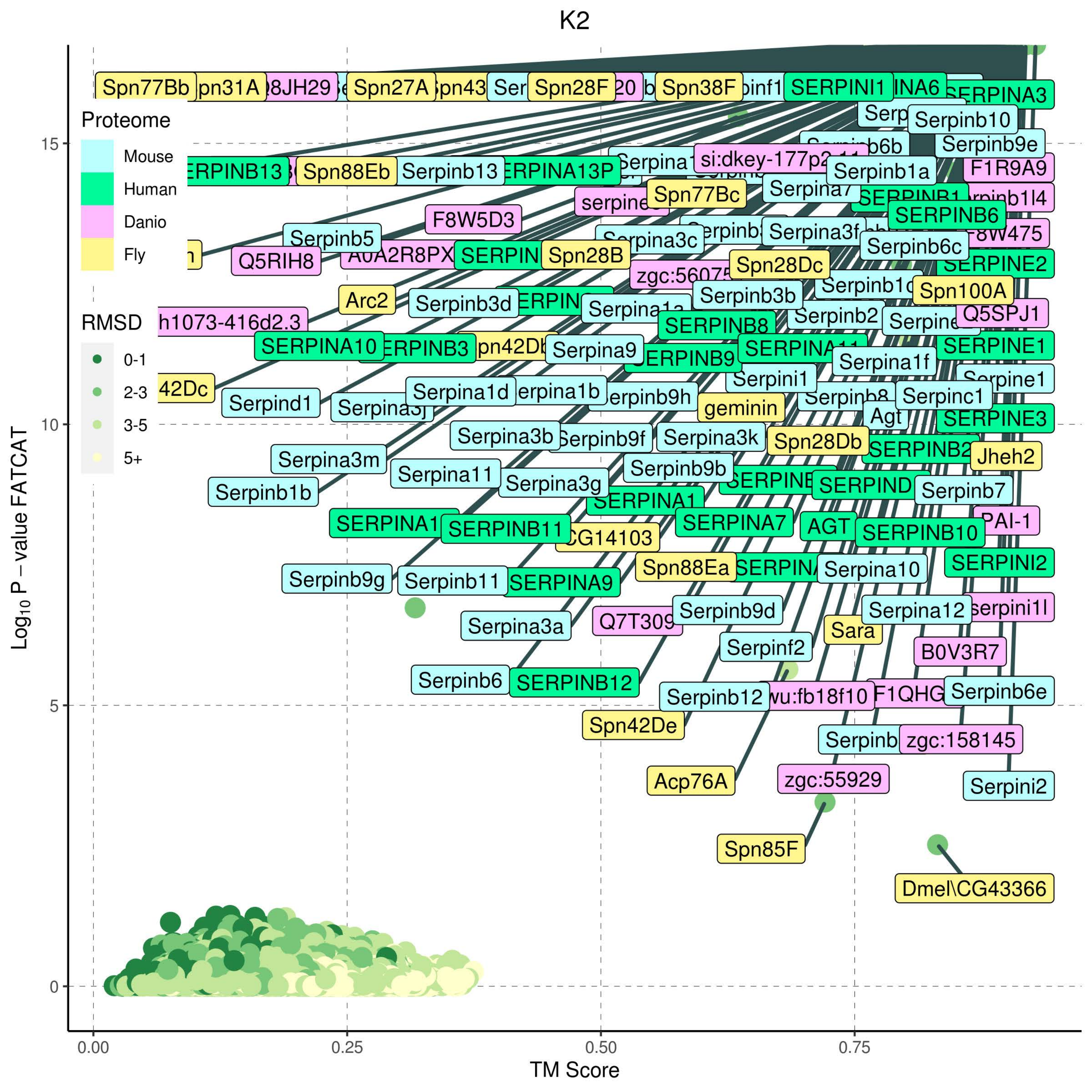
K2
Proteome
Mouse
Human
Danio
Fly
RMSD
0-1
2-3
3-5
5+
Log10 P − value FATCAT
TM Score
0.00
0.25
0.50
0.75
0
5
10
15
Spn77Bb
Spn27A
Spn28F
Spn38F
SERPINI1
SERPINA3
Serpinb10
Serpinb9e
F1R9A9
Spn88Eb
Serpinb13
si:dkey-177p2
Serpinb1a
Spn77Bc
Serpina7
SERPINB6
F8W5D3
Serpinb5
Serpina3c
Serpina3f
Serpinb6c
Q5RIH8
SERPINE2
Spn28B
Spn28Dc
Arc2
Serpinb3d
Serpinb3b
Spn100A
Q5SPJ1
Serpinb2
SERPINB8
SERPINA10
Serpina9
SERPINB9
SERPINE1
Serpina1f
Serpini1
42Dc
Serpind1
Serpina1d
Serpinc1
geminin
Agt
SERPINE3
Serpina3b
Serpina3k
Spn28Db
Jheh2
Serpina3m
Serpina11
Serpina3g
Serpinb9b
Serpinb7
Serpinb1b
SERPINB11
SERPINA7
AGT
SERPINB10
PAI-1
CG14103
Spn88Ea
Serpina10
SERPINI2
Serpinb9g
Serpinb11
SERPINA9
Serpinb9d
Serpina12
Serpina3a
Sara
Serpinf2
B0V3R7
Serpinb6
SERPINB12
Serpinb12
wu:fb18f10
Serpinb6e
Spn42De
zgc:158145
Acp76A
zgc:55929
Serpini2
Spn85F
Dmel\CG43366

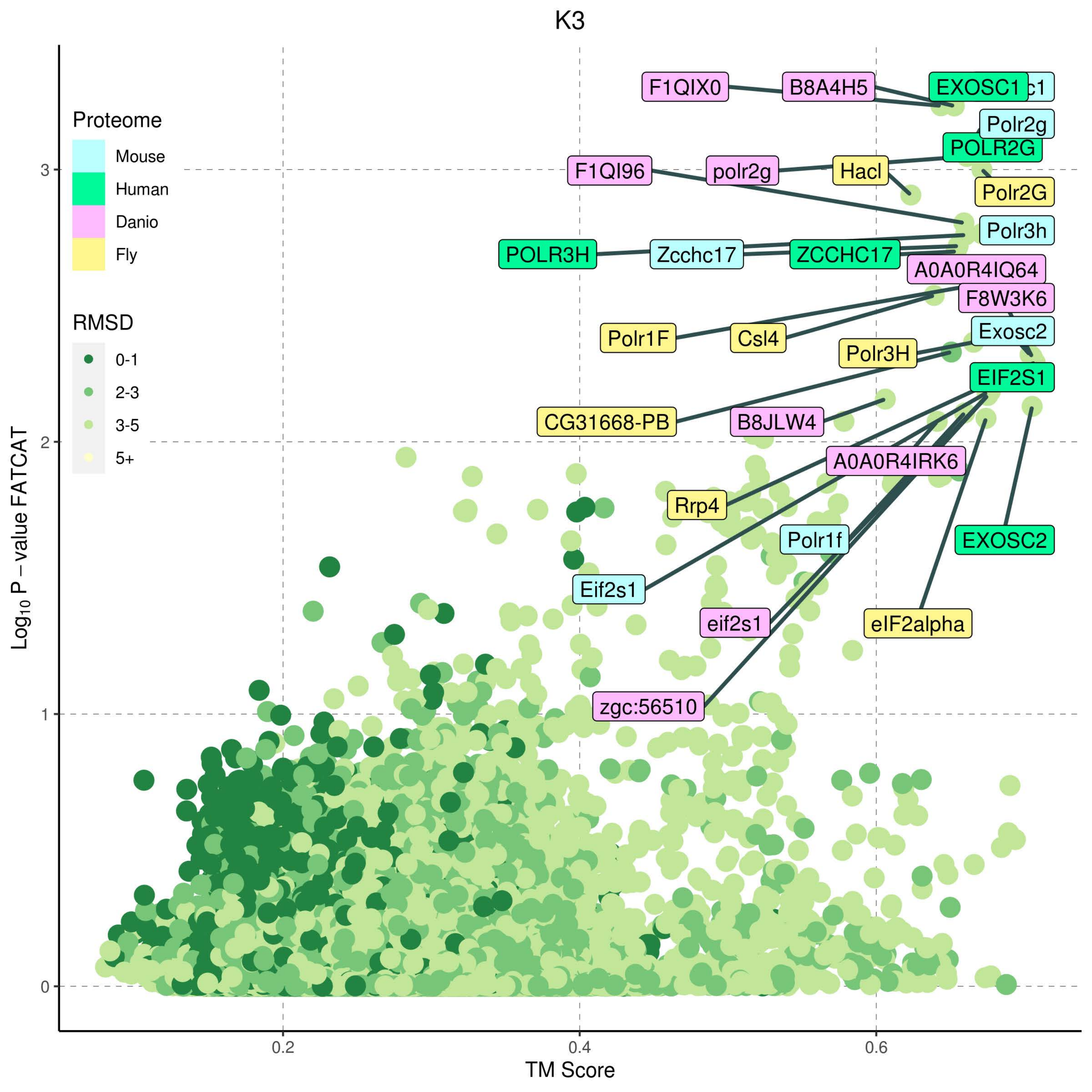

K3
Proteome
Mouse
Human
Danio
Fly
RMSD
0-1
2-3
3-5
5+
F1QIX0
B8A4H5
EXOSC1
c1
Polr2g
POLR2G
F1QI96
polr2g
Hacl
Polr2G
Polr3h
POLR3H
Zcchc17
ZCCHC17
A0A0R4IQ64
F8W3K6
Polr1F
Csl4
Exosc2
Polr3H
EIF2S1
CG31668-PB
B8JLW4
A0A0R4IRK6
Rrp4
Polr1f
EXOSC2
Eif2s1
eif2s1
eIF2alpha
zgc:56510
Log10 P – value FATCAT
TM Score
0
1
2
3
0.2
0.4
0.6

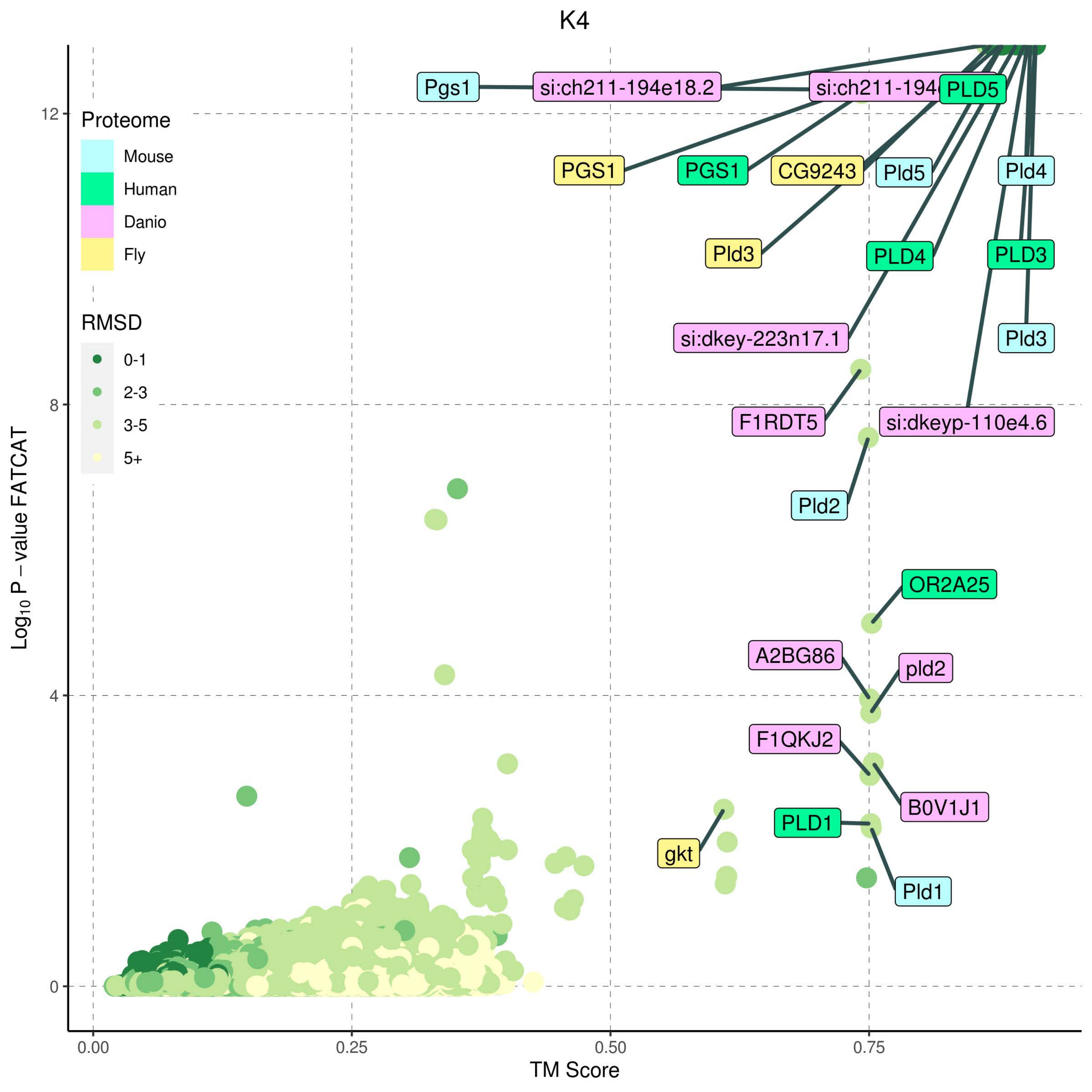

K4
Proteome
Mouse
Human
Danio
Fly
RMSD
0-1
2-3
3-5
5+
Pgs1
si:ch211-194e18.2
PLD5
PGS1
PGS1
CG9243
Pld5
Pld4
Pld3
PLD4
PLD3
si:dkey-223n17.1
Pld3
F1RDT5
si:dkeyp-110e4.6
Pld2
OR2A25
A2BG86
pld2
F1QKJ2
B0V1J1
PLD1
gkt
Pld1
Log10 P − value FATCAT
TM Score
0
4
8
12
0.00
0.25
0.50
0.75

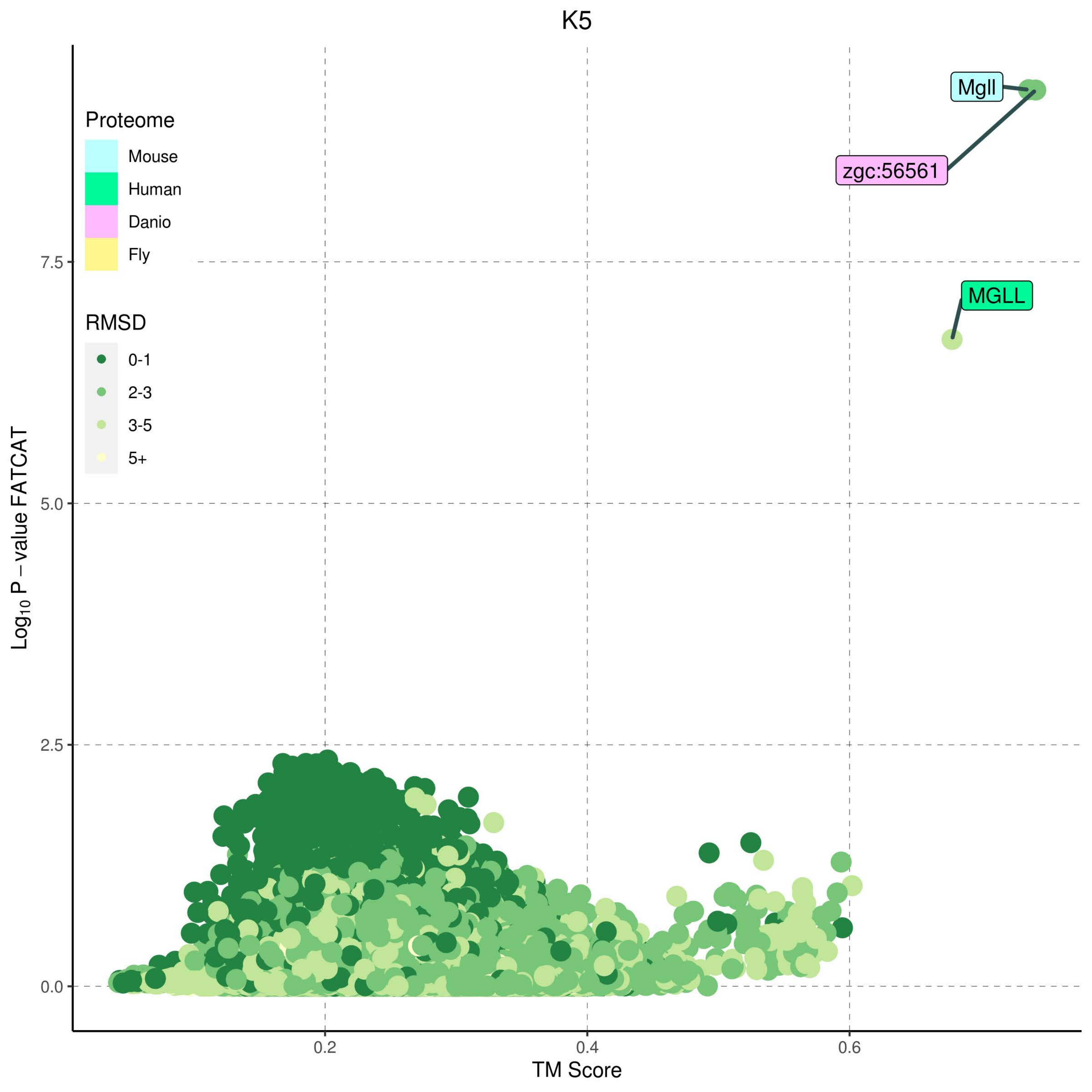

K5
Proteome
Mouse
Human
Danio
Fly
RMSD
0-1
2-3
3-5
5+
Mgll
zgc:56561
MGLL
7.5
5.0
2.5
0.0
0.2
0.4
0.6
Log10 P – value FATCAT
TM Score

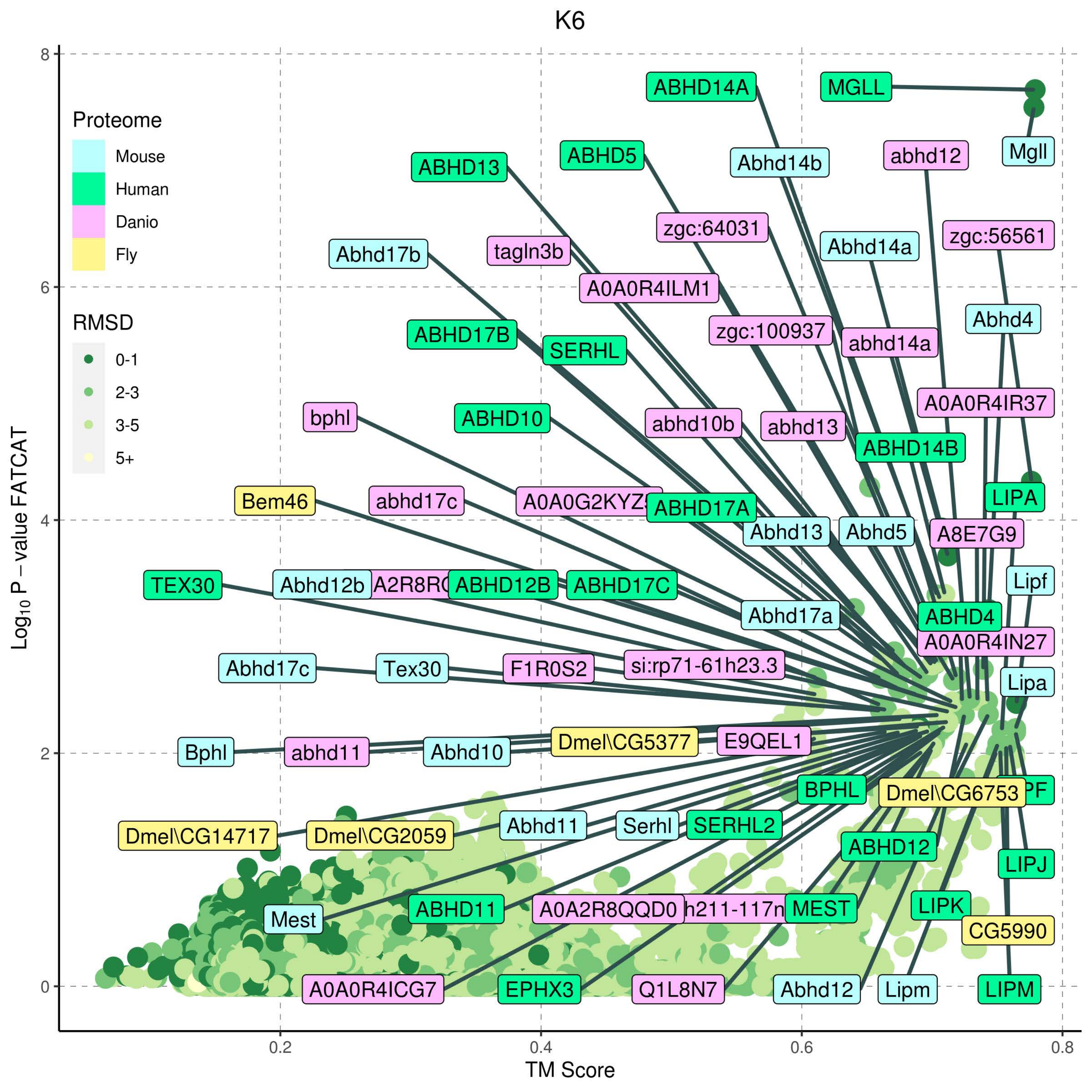

K6
Proteome
Mouse
Human
Danio
Fly
RMSD
0-1
2-3
3-5
5+
Log10 P − value FATCAT
TM Score
0
2
4
6
8
0.2
0.4
0.6
0.8
ABHD14A
MGLL
Mgll
ABHD13
ABHD5
Abhd14b
abhd12
zgc:64031
Abhd14a
zgc:56561
Abhd17b
tagln3b
A0A0R4ILM1
ABHD17B
SERHL
zgc:100937
abhd14a
Abhd4
bphl
ABHD10
abhd10b
abhd13
A0A0R4IR37
ABHD14B
Bem46
abhd17c
ABHD17A
LIPA
Abhd13
Abhd5
A8E7G9
TEX30
Abhd12b
ABHD12B
ABHD17C
Lipf
Abhd17a
ABHD4
A0A0R4IN27
Abhd17c
Tex30
F1R0S2
si:rp71-61h23.3
Lipa
Bphl
abhd11
Abhd10
Dmel\CG5377
E9QEL1
BPHL
Dmel\CG6753
Dmel\CG14717
Dmel\CG2059
Abhd11
Serhl
SERHL2
ABHD12
LIPJ
Mest
ABHD11
A0A2R8QQD0
MEST
LIPK
CG5990
A0A0R4ICG7
EPHX3
Q1L8N7
Abhd12
Lipm
LIPM

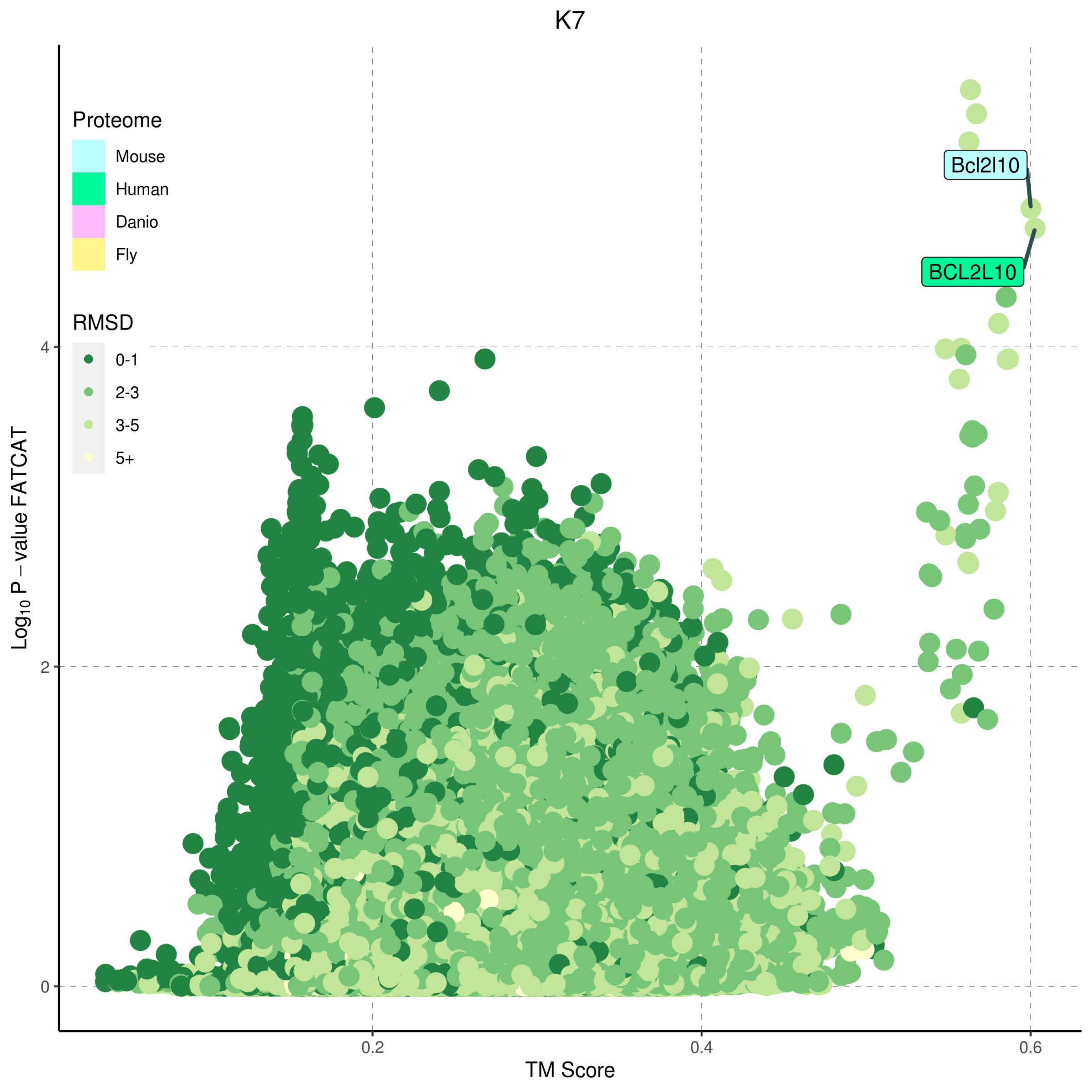

K7
Proteome
Mouse
Human
Danio
Fly
RMSD
0-1
2-3
3-5
5+
Bcl2l10
BCL2L10
$Log_{10}$ P – value FATCAT
TM Score
0
2
4
0.2
0.4
0.6

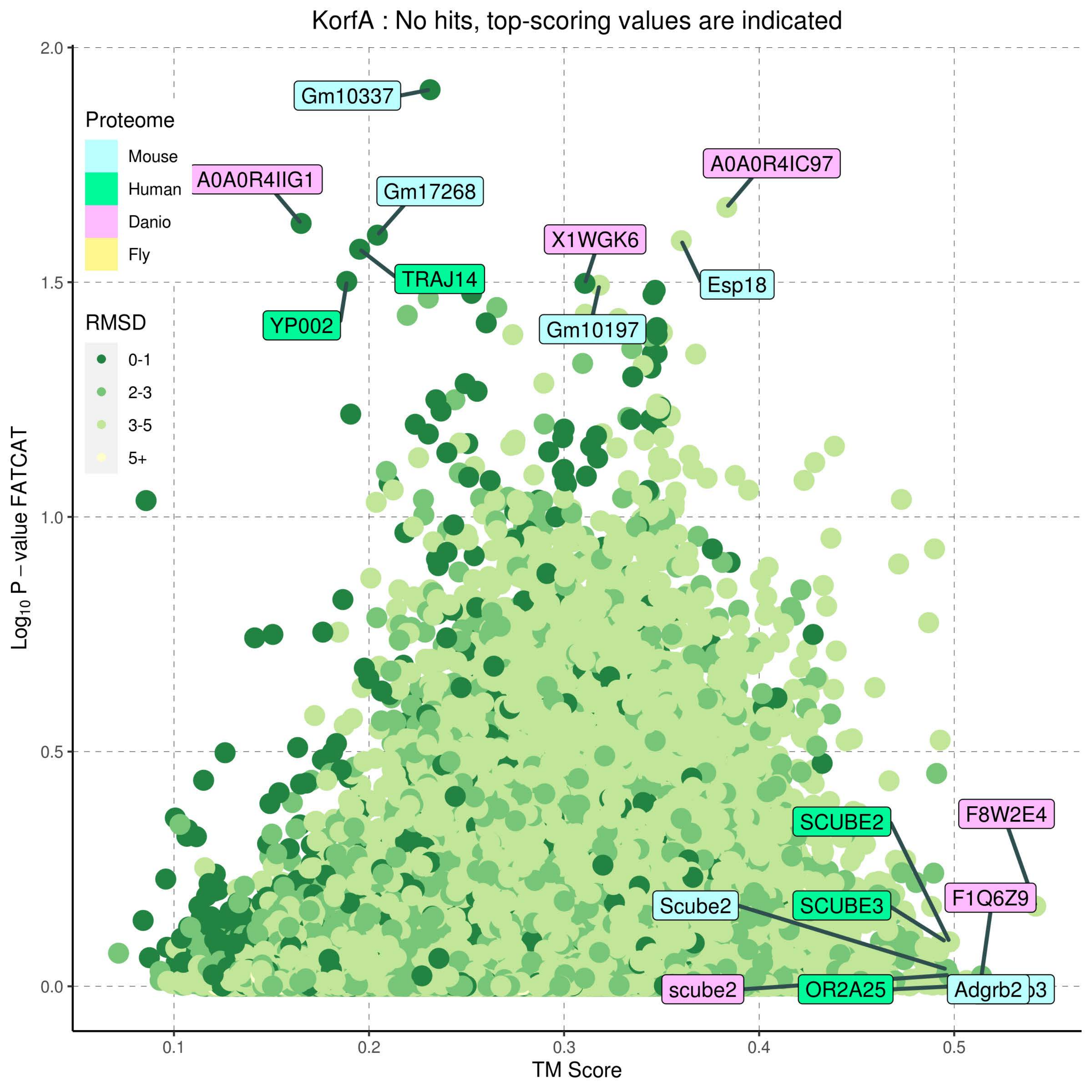

KorfA : No hits, top-scoring values are indicated
Proteome
Mouse
Human
Danio
Fly
RMSD
0-1
2-3
3-5
5+
Gm10337
A0A0R4IIG1
Gm17268
TRAJ14
YP002
X1WGK6
Gm10197
A0A0R4IC97
Esp18
SCUBE2
F8W2E4
Scube2
SCUBE3
F1Q6Z9
scube2
OR2A25
Adgrb2
Log10 P − value FATCAT
TM Score
2.0
1.5
1.0
0.5
0.0
0.1
0.2
0.3
0.4
0.5

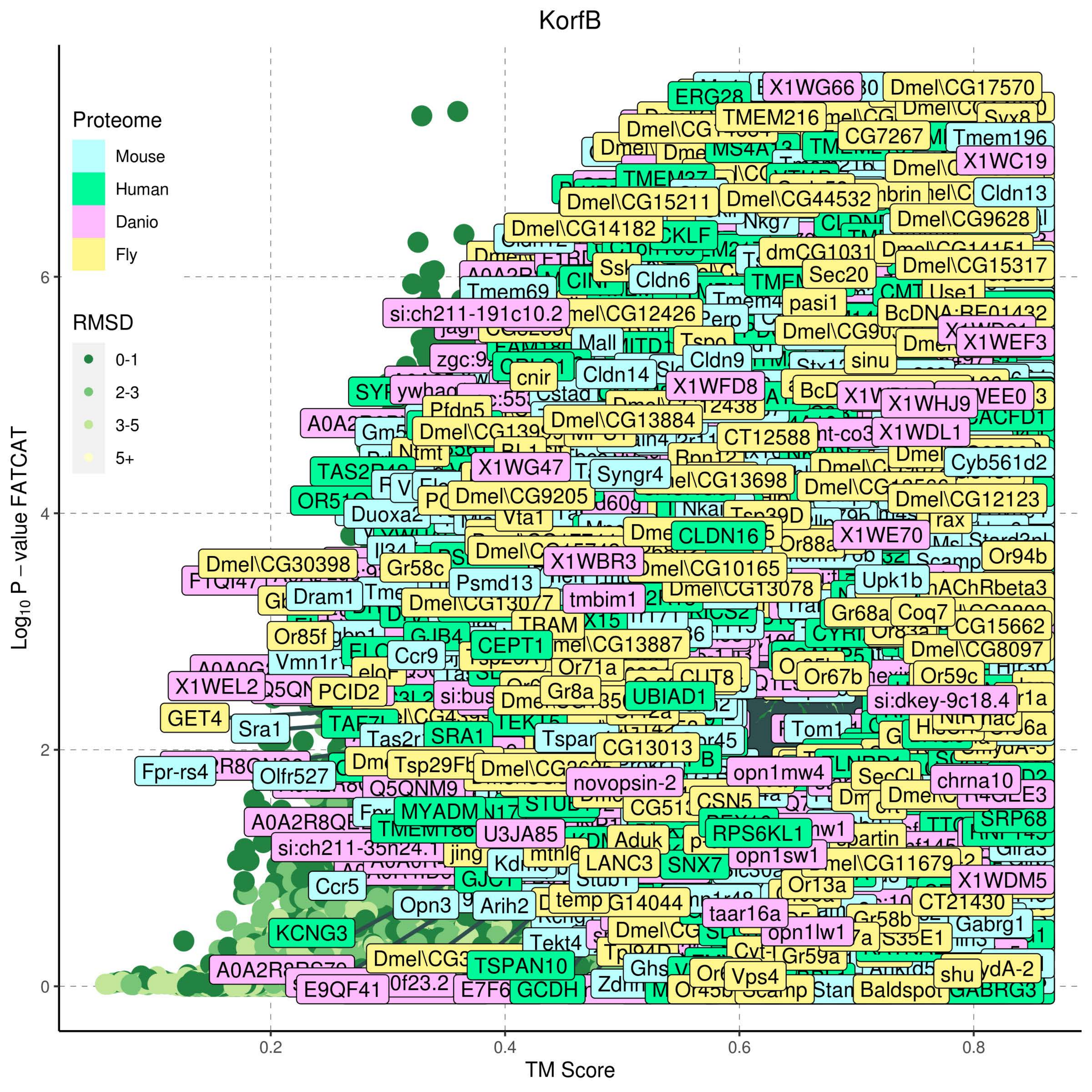
KorfB
Proteome
Mouse
Human
Danio
Fly
RMSD
0-1
2-3
3-5
5+
Log10 P − value FATCAT
TM Score
0
2
4
6
0.2
0.4
0.6
0.8

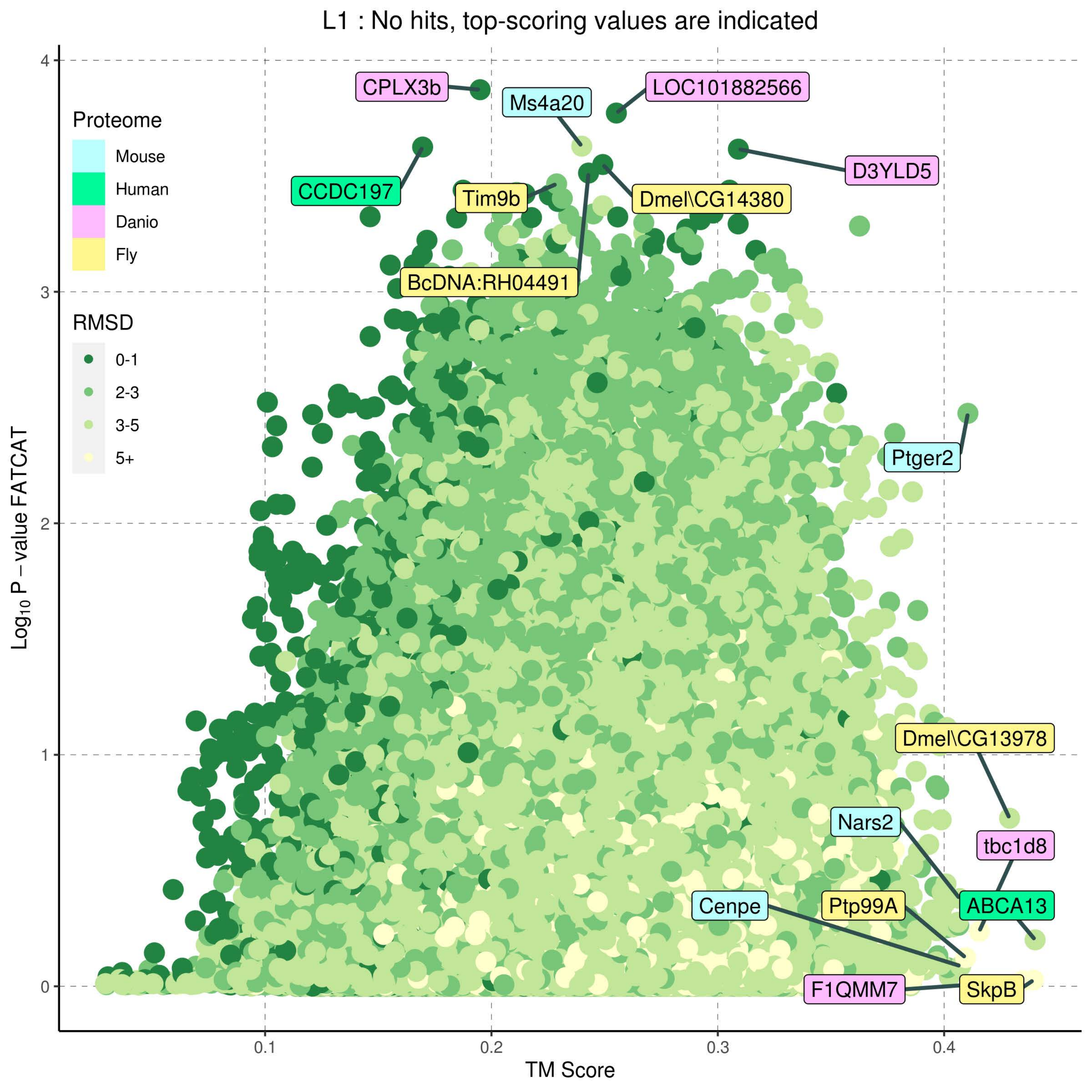

L1 : No hits, top-scoring values are indicated
Proteome
Mouse
Human
Danio
Fly
RMSD
0-1
2-3
3-5
5+
CPLX3b
Ms4a20
LOC101882566
CCDC197
Tim9b
Dmel\CG14380
D3YLD5
BcDNA:RH04491
Ptger2
Dmel\CG13978
Nars2
tbc1d8
Cenpe
Ptp99A
ABCA13
F1QMM7
SkpB
Log10 P – value FATCAT
TM Score
0
1
2
3
4
0.1
0.2
0.3
0.4

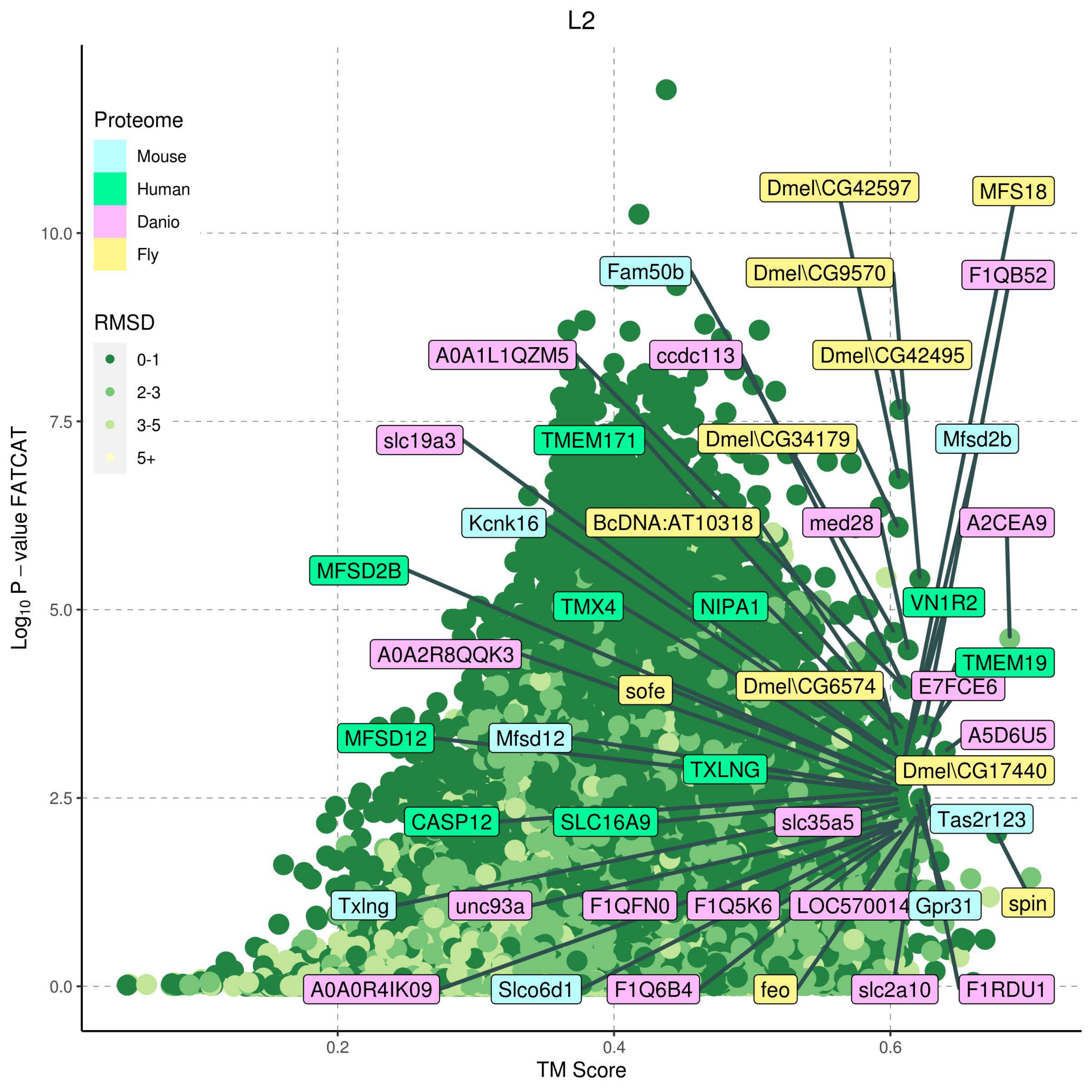

L2
Proteome
Mouse
Human
Danio
Fly
RMSD
0-1
2-3
3-5
5+
Log10 P − value FATCAT
TM Score
10.0
7.5
5.0
2.5
0.0
0.2
0.4
0.6
Dmel\CG42597
MFS18
Fam50b
Dmel\CG9570
F1QB52
A0A1L1QZM5
ccdc113
Dmel\CG42495
slc19a3
TMEM171
Dmel\CG34179
Mfsd2b
Kcnk16
BcDNA:AT10318
med28
A2CEA9
MFSD2B
TMX4
NIPA1
VN1R2
A0A2R8QQK3
TMEM19
sofe
Dmel\CG6574
E7FCE6
A5D6U5
MFSD12
Mfsd12
TXLNG
Dmel\CG17440
CASP12
SLC16A9
slc35a5
Tas2r123
Txlng
unc93a
F1QFN0
F1Q5K6
LOC570014
Gpr31
spin
A0A0R4IK09
Slco6d1
F1Q6B4
feo
slc2a10
F1RDU1

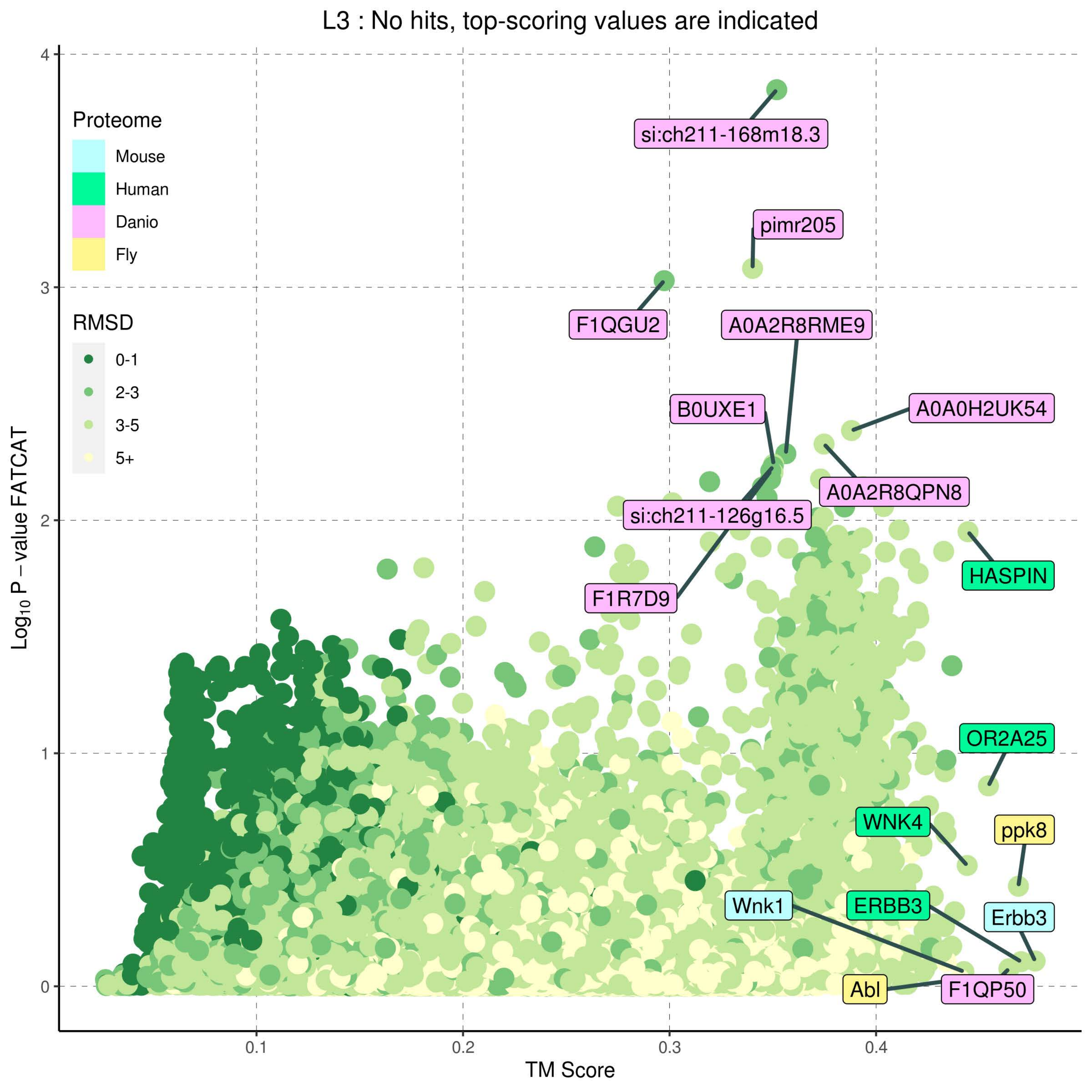

L3 : No hits, top-scoring values are indicated
Proteome
Mouse
Human
Danio
Fly
RMSD
0-1
2-3
3-5
5+
si:ch211-168m18.3
pimr205
F1QGU2
A0A2R8RME9
B0UXE1
A0A0H2UK54
A0A2R8QPN8
si:ch211-126g16.5
HASPIN
F1R7D9
OR2A25
WNK4
ppk8
Wnk1
ERBB3
Erbb3
Abl
F1QP50
Log10 P − value FATCAT
TM Score
0
1
2
3
4
0.1
0.2
0.3
0.4

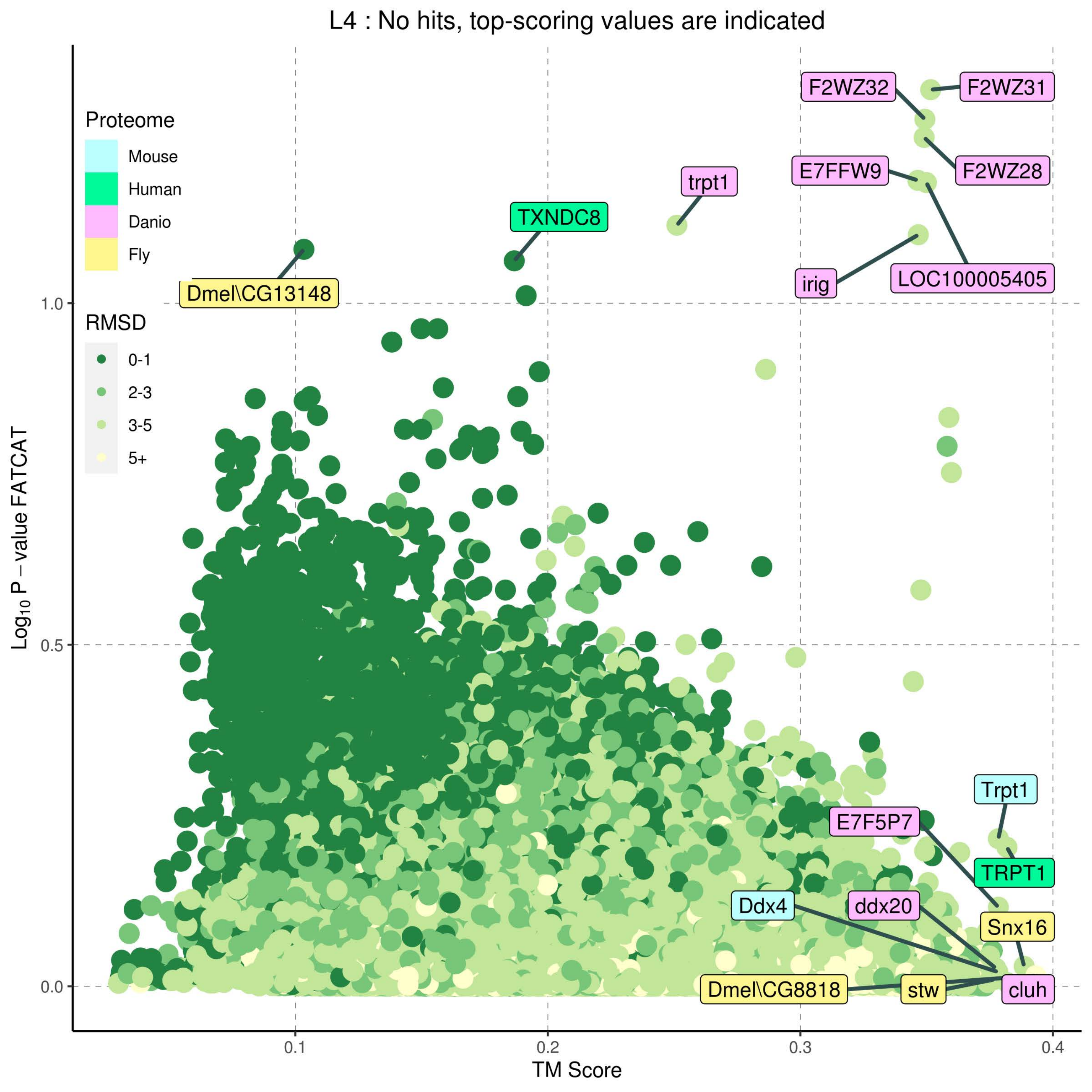

L4 : No hits, top-scoring values are indicated
Proteome
Mouse
Human
Danio
Fly
RMSD
0-1
2-3
3-5
5+
F2WZ32
F2WZ31
F2WZ28
E7FFW9
trpt1
TXNDC8
irig
LOC100005405
Dmel\CG13148
Trpt1
E7F5P7
TRPT1
Ddx4
ddx20
Snx16
Dmel\CG8818
stw
cluh
1.0
0.5
0.0
0.1
0.2
0.3
0.4
$\mathrm{Log_{10}}$ P − value FATCAT
TM Score

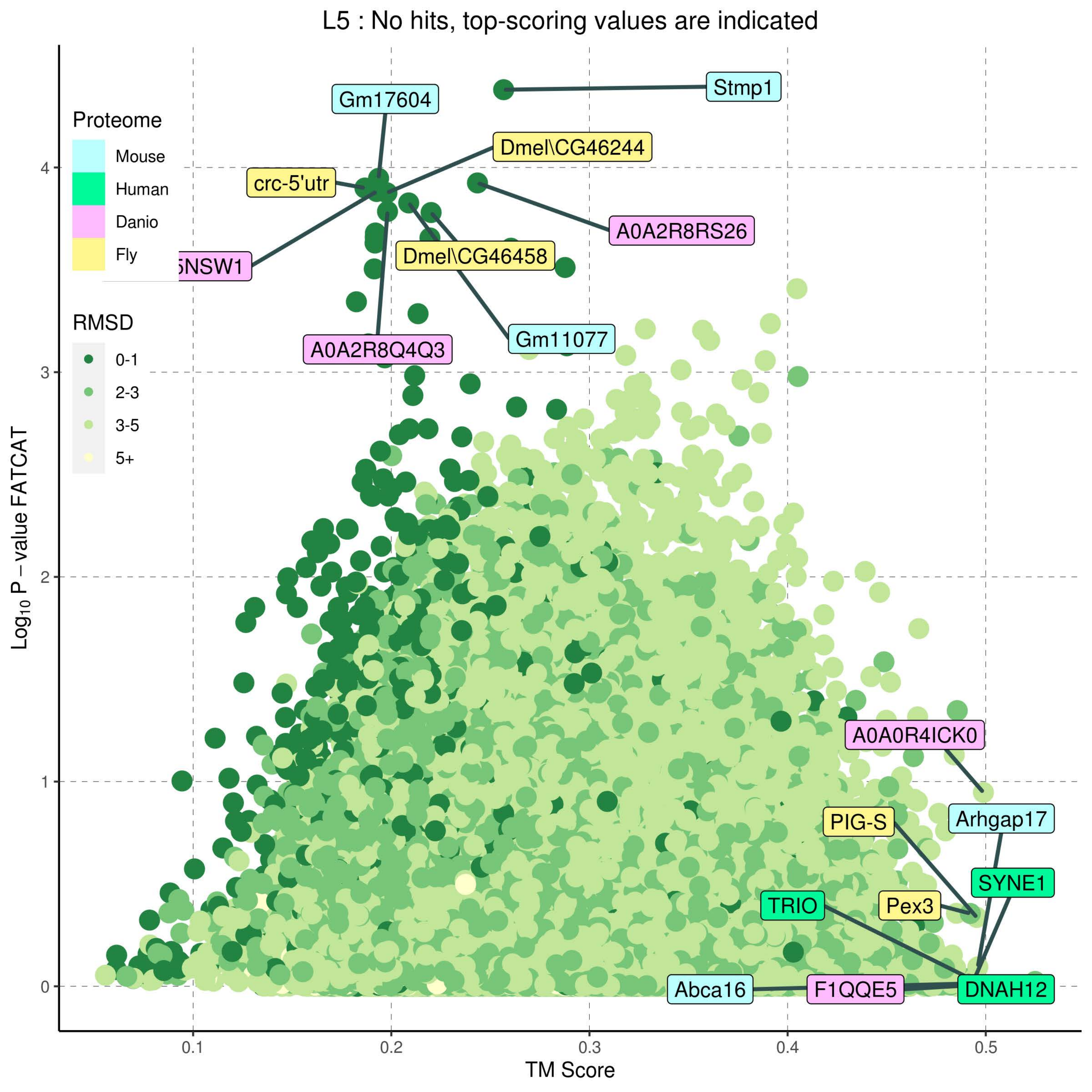

L5 : No hits, top-scoring values are indicated
Proteome
Mouse
Human
Danio
Fly
RMSD
0-1
2-3
3-5
5+
Stmp1
Gm17604
Dmel\CG46244
crc-5'utr
A0A2R8RS26
NSW1
Dmel\CG46458
Gm11077
A0A2R8Q4Q3
A0A0R4ICK0
PIG-S
Arhgap17
SYNE1
TRIO
Pex3
Abca16
F1QQE5
DNAH12
Log10 P − value FATCAT
TM Score
0
1
2
3
4
0.1
0.2
0.3
0.4
0.5

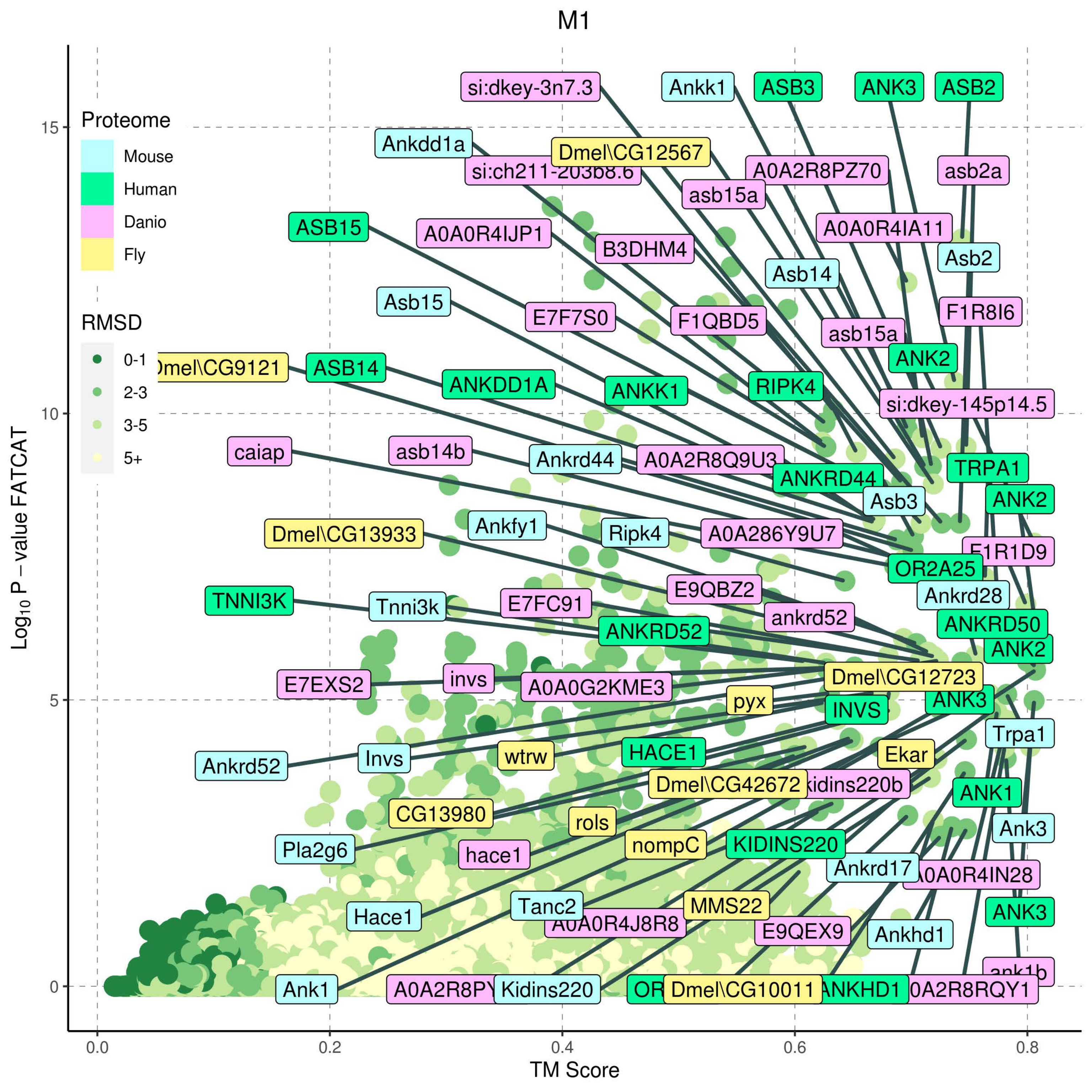
M1
Proteome
Mouse
Human
Danio
Fly
RMSD
0-1
2-3
3-5
5+
Log10 P − value FATCAT
TM Score
0.0
0.2
0.4
0.6
0.8
0
5
10
15
si:dkey-3n7.3
Ankk1
ASB3
ANK3
ASB2
Ankdd1a
Dmel\CG12567
A0A2R8PZ70
asb2a
asb15a
ASB15
A0A0R4IJP1
B3DHM4
A0A0R4IA11
Asb2
Asb14
Asb15
E7F7S0
F1QBD5
F1R8I6
asb15a
ASB14
ANK2
ANKDD1A
ANKK1
RIPK4
si:dkey-145p14.5
caiap
asb14b
Ankrd44
TRPA1
ANKRD44
Asb3
ANK2
Ankfy1
Ripk4
A0A286Y9U7
Dmel\CG13933
OR2A25
E9QBZ2
Ankrd28
TNNI3K
Tnni3k
E7FC91
ankrd52
ANKRD52
ANKRD50
ANK2
invs
Dmel\CG12723
E7EXS2
A0A0G2KME3
pyx
ANK3
INVS
Trpa1
Invs
Ankrd52
wtrw
HACE1
Ekar
Dmel\CG42672
ANK1
CG13980
rols
Ank3
Pla2g6
nompC
KIDINS220
hace1
Ankrd17
Tanc2
MMS22
ANK3
Hace1
E9QEX9
Ankhd1
Ank1
Kidins220
Dmel\CG10011
ANKHD1

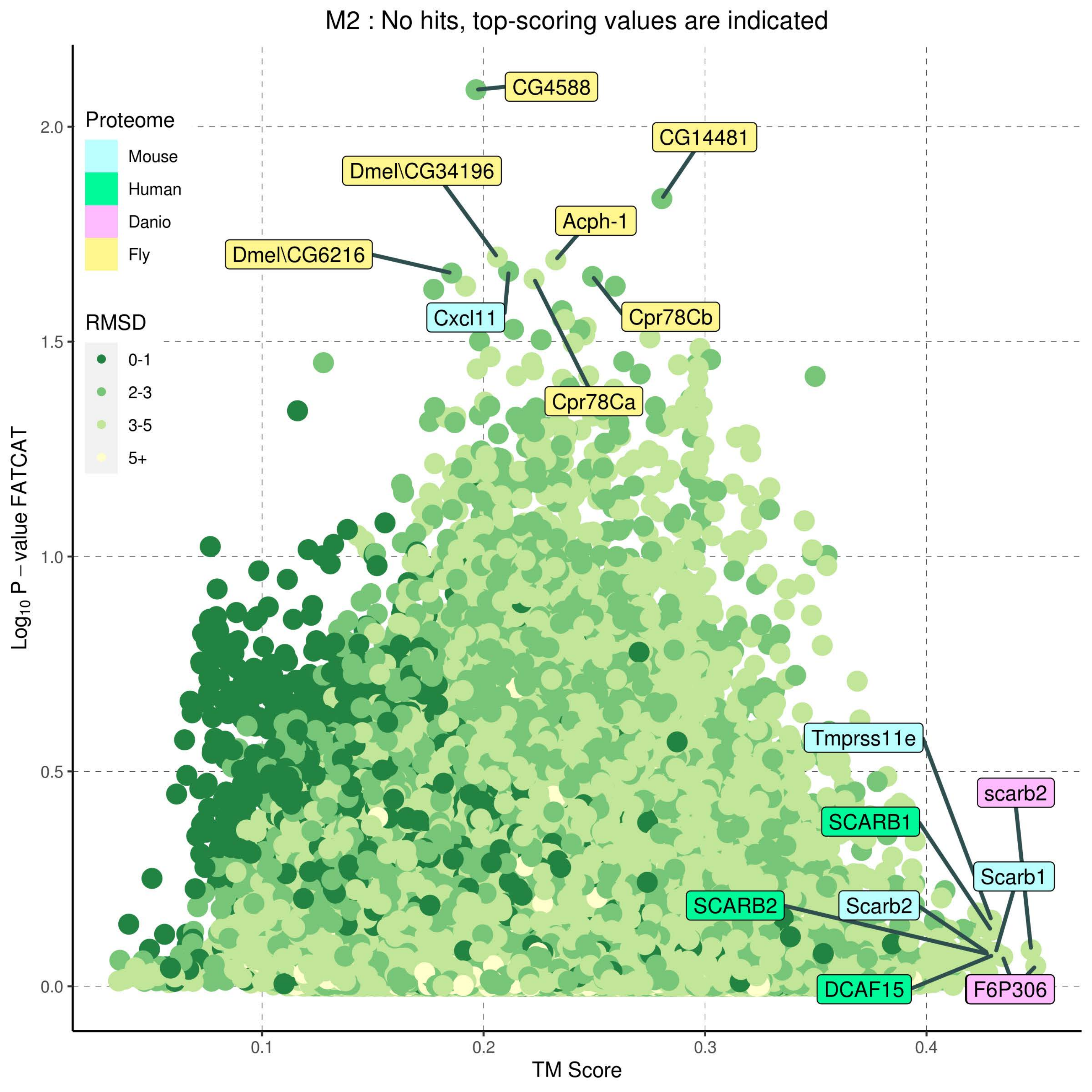

M2 : No hits, top-scoring values are indicated
Proteome
Mouse
Human
Danio
Fly
RMSD
0-1
2-3
3-5
5+
CG4588
CG14481
Dmel\CG34196
Acph-1
Dmel\CG6216
Cxcl11
Cpr78Cb
Cpr78Ca
Tmprss11e
scarb2
SCARB1
Scarb1
SCARB2
Scarb2
DCAF15
F6P306
2.0
1.5
1.0
0.5
0.0
0.1
0.2
0.3
0.4
TM Score
Log$_{10}$ P – value FATCAT

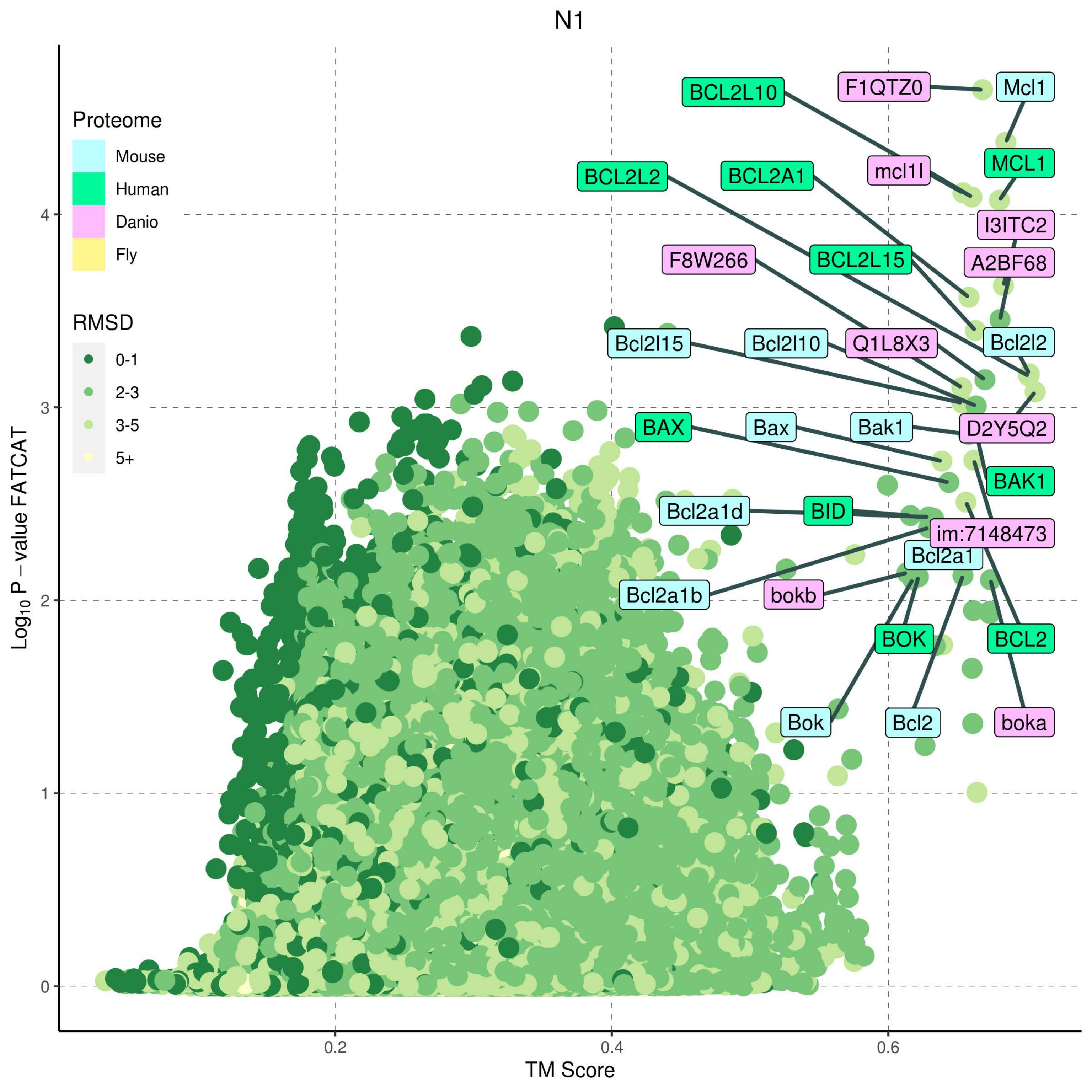

N1
Proteome
Mouse
Human
Danio
Fly
RMSD
0-1
2-3
3-5
5+
Log10 P − value FATCAT
TM Score
0
1
2
3
4
0.2
0.4
0.6
BCL2L10
F1QTZ0
Mcl1
mcl1l
MCL1
BCL2L2
BCL2A1
I3ITC2
A2BF68
F8W266
BCL2L15
Bcl2l15
Bcl2l10
Q1L8X3
Bcl2l2
BAX
Bax
Bak1
D2Y5Q2
BAK1
Bcl2a1d
BID
im:7148473
Bcl2a1
Bcl2a1b
bokb
BOK
BCL2
Bok
Bcl2
boka

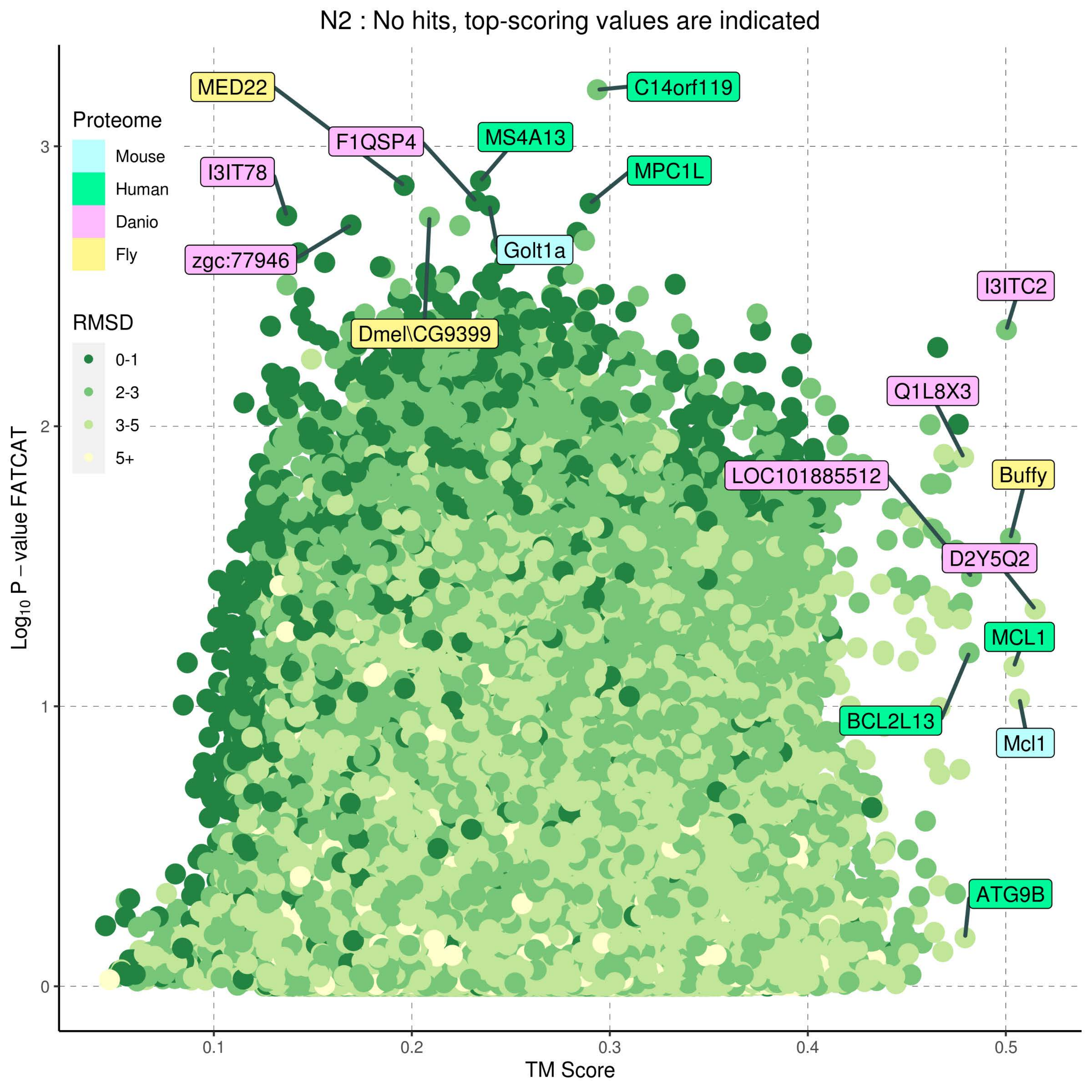
N2 : No hits, top-scoring values are indicated
Proteome
Mouse
Human
Danio
Fly
RMSD
0-1
2-3
3-5
5+
MED22
F1QSP4
MS4A13
C14orf119
I3IT78
MPC1L
zgc:77946
Golt1a
Dmel\CG9399
I3ITC2
Q1L8X3
LOC101885512
Buffy
D2Y5Q2
MCL1
BCL2L13
Mcl1
ATG9B
Log10 P − value FATCAT
TM Score
0
1
2
3
0.1
0.2
0.3
0.4
0.5

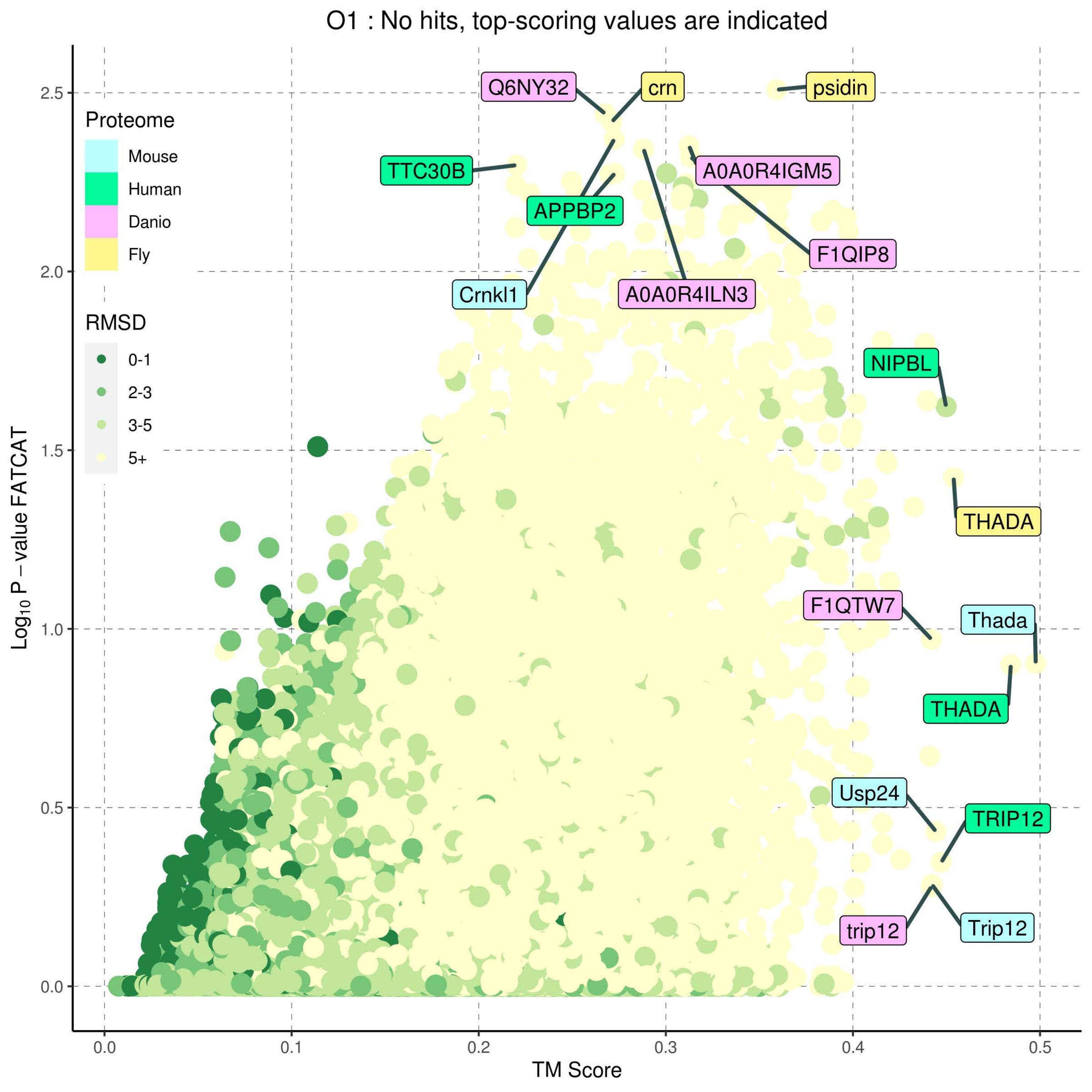

O1 : No hits, top-scoring values are indicated
Proteome
Mouse
Human
Danio
Fly
RMSD
0-1
2-3
3-5
5+
Log10 P – value FATCAT
TM Score
0.0
0.5
1.0
1.5
2.0
2.5
0.0
0.1
0.2
0.3
0.4
0.5
Q6NY32
crn
psidin
TTC30B
A0A0R4IGM5
APPBP2
F1QIP8
Crnkl1
A0A0R4ILN3
NIPBL
THADA
F1QTW7
Thada
THADA
Usp24
TRIP12
trip12
Trip12

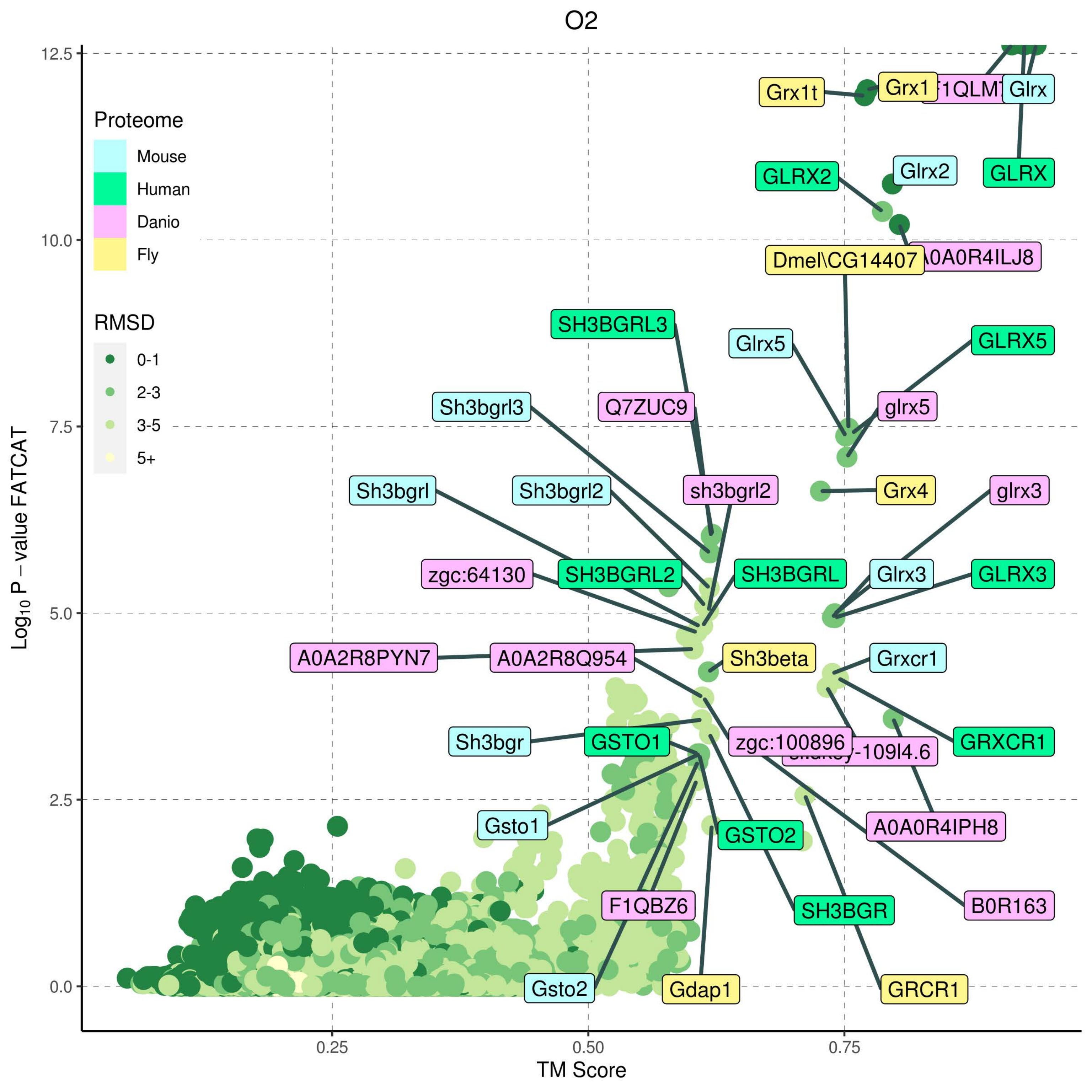
O2
Proteome
Mouse
Human
Danio
Fly
RMSD
0-1
2-3
3-5
5+
Log10 P – value FATCAT
TM Score
12.5
10.0
7.5
5.0
2.5
0.0
0.25
0.50
0.75
Grx1t
Grx1
Glrx
GLRX2
Glrx2
GLRX
A0A0R4ILJ8
Dmel\CG14407
SH3BGRL3
Glrx5
GLRX5
Sh3bgrl3
Q7ZUC9
glrx5
Sh3bgrl
Sh3bgrl2
sh3bgrl2
Grx4
glrx3
zgc:64130
SH3BGRL2
SH3BGRL
Glrx3
GLRX3
A0A2R8PYN7
A0A2R8Q954
Sh3beta
Grxcr1
Sh3bgr
GSTO1
zgc:100896
GRXCR1
Gsto1
GSTO2
A0A0R4IPH8
F1QBZ6
SH3BGR
B0R163
Gsto2
Gdap1
GRCR1

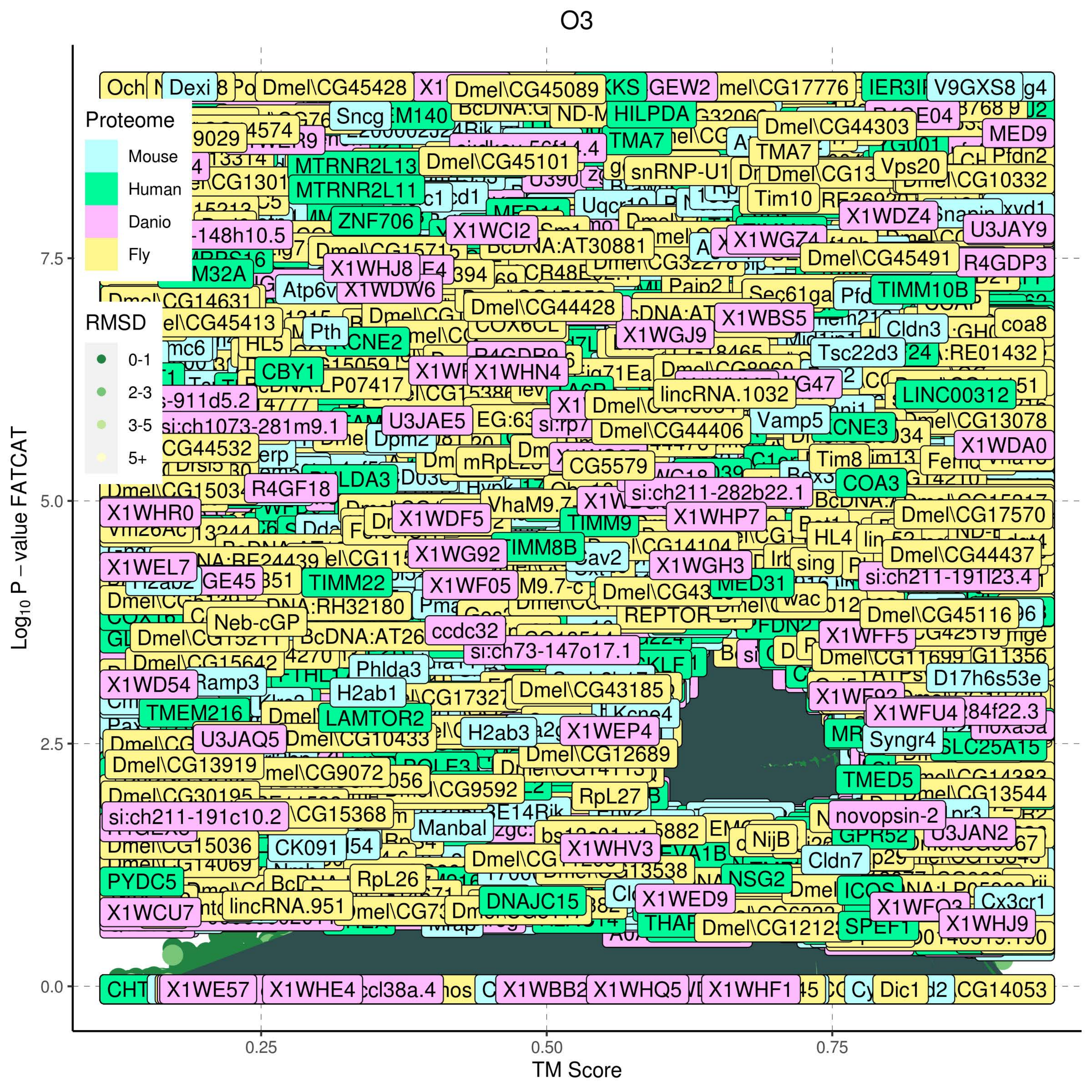
O3
Proteome
Mouse
Human
Danio
Fly
RMSD
0-1
2-3
3-5
5+
Log10 P − value FATCAT
TM Score
0.0
2.5
5.0
7.5
0.25
0.50
0.75
Dmel\CG45428
Dmel\CG45089
Dmel\CG17776
V9GXS8
Sncg
HILPDA
Dmel\CG44303
MED9
TMA7
MTRNR2L13
MTRNR2L11
Dmel\CG45101
Vps20
Dmel\CG10332
Tim10
ZNF706
X1WCI2
X1WDZ4
U3JAY9
X1WHJ8
X1WDW6
Dmel\CG45491
R4GDP3
TIMM10B
Dmel\CG44428
X1WBS5
X1WGJ9
Cldn3
coa8
Pth
CBY1
X1WHN4
Tsc22d3
lincRNA.1032
LINC00312
U3JAE5
Dmel\CG44406
Vamp5
Dmel\CG13078
X1WDA0
CG5579
Tim8
COA3
R4GF18
si:ch211-282b22.1
X1WHR0
X1WDF5
X1WHP7
TIMM9
Dmel\CG17570
X1WG92
TIMM8B
X1WGH3
Dmel\CG44437
X1WEL7
TIMM22
X1WF05
si:ch211-191l23.4
Neb-cGP
ccdc32
REPTOR
Dmel\CG45116
si:ch73-147o17.1
Phlda3
X1WD54
Ramp3
H2ab1
Dmel\CG43185
D17h6s53e
TMEM216
LAMTOR2
X1WFU4
U3JAQ5
Dmel\CG10433
H2ab3
X1WEP4
Syngr4
Dmel\CG12689
Dmel\CG13919
TMED5
RpL27
si:ch211-191c10.2
Manbal
novopsin-2
U3JAN2
CK091
X1WHV3
NijB
Cldn7
PYDC5
RpL26
NSG2
DNAJC15
X1WED9
X1WCU7
lincRNA.951
Cx3cr1
X1WHJ9
X1WE57
X1WHE4
X1WBB2
X1WHQ5
X1WHF1
Dic1

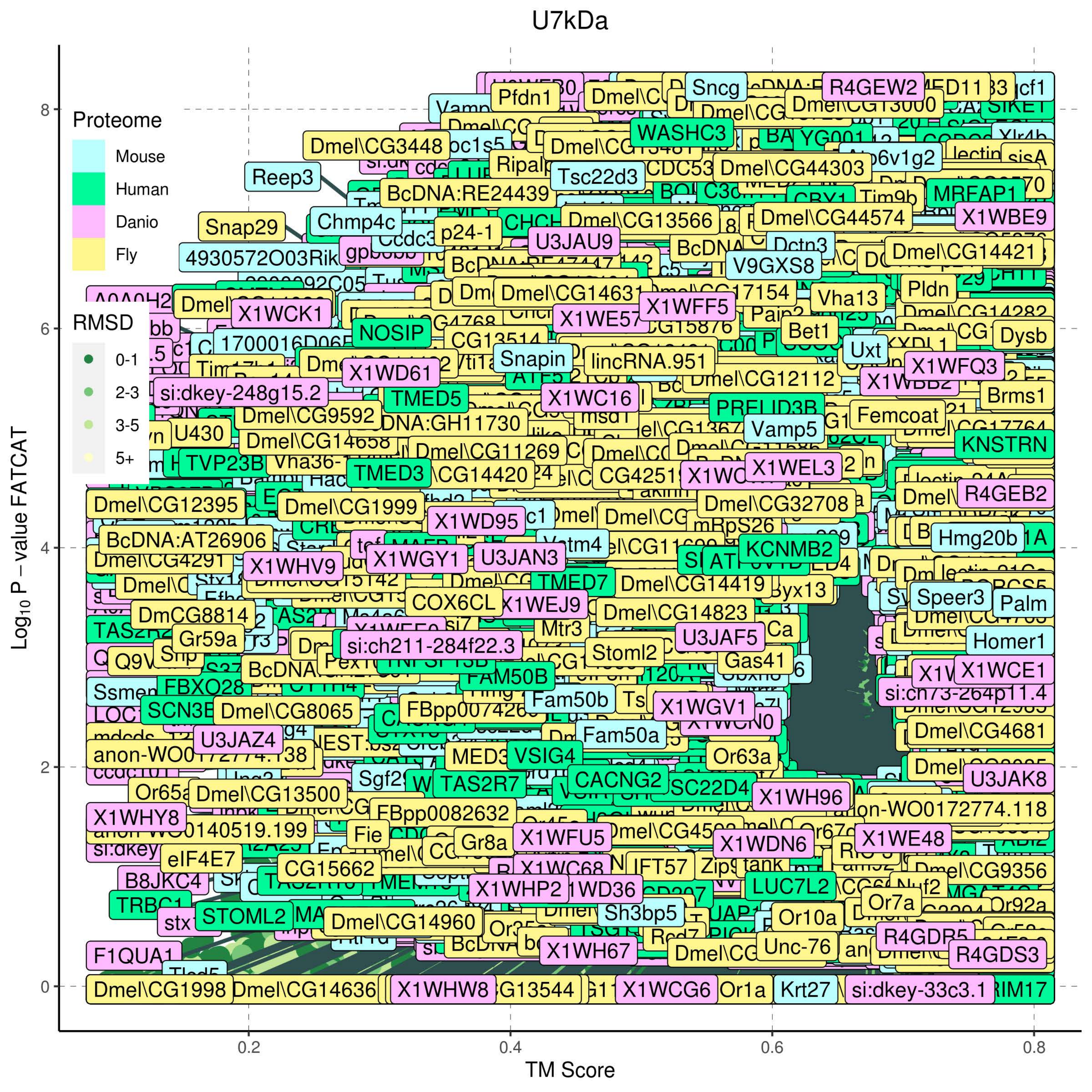

U7kDa
Proteome
Mouse
Human
Danio
Fly
RMSD
0-1
2-3
3-5
5+
Log10 P − value FATCAT
TM Score
0
2
4
6
8
0.2
0.4
0.6
0.8
Pfdn1
Sncg
R4GEW2
WASHC3
Dmel\CG3448
Reep3
Tsc22d3
Dmel\CG44303
BcDNA:RE24439
MRFAP1
Snap29
Chmp4c
p24-1
Dmel\CG13566
Dmel\CG44574
X1WBE9
4930572O03Rik
U3JAU9
V9GXS8
Dmel\CG14421
Dmel\CG14631
X1WCK1
X1WFF5
Vha13
Pldn
NOSIP
Bet1
Dysb
Snapin
lincRNA.951
Uxt
X1WFQ3
X1WD61
si:dkey-248g15.2
TMED5
X1WC16
Dmel\CG12112
Brms1
Femcoat
Dmel\CG9592
Vamp5
U430
KNSTRN
Dmel\CG11269
TVP23B
TMED3
X1WEL3
Dmel\CG12395
Dmel\CG1999
Dmel\CG32708
R4GEB2
X1WD95
BcDNA:AT26906
Hmg20b
KCNMB2
X1WHV9
X1WGY1
U3JAN3
TMED7
Dmel\CG14419
Speer3
Palm
DmCG8814
COX6CL
X1WEJ9
Dmel\CG14823
Gr59a
si:ch211-284f22.3
U3JAF5
Homer1
Stoml2
Gas41
X1WCE1
FBXO28
FAM50B
si:ch73-264p11.4
SCN3B
Dmel\CG8065
Fam50b
X1WGV1
Dmel\CG4681
U3JAZ4
Fam50a
anon-WO0172774.138
VSIG4
Or63a
U3JAK8
Dmel\CG13500
TAS2R7
CACNG2
X1WH96
X1WHY8
FBpp0082632
anon-WO0172774.118
X1WFU5
X1WDN6
X1WE48
eIF4E7
CG15662
IFT57
Dmel\CG9356
B8JKC4
X1WHP2
LUC7L2
Or7a
STOML2
Dmel\CG14960
Sh3bp5
Or10a
R4GDR5
F1QUA1
X1WH67
Unc-76
R4GDS3
Dmel\CG1998
Dmel\CG14636
X1WHW8
X1WCG6
Or1a
Krt27
si:dkey-33c3.1

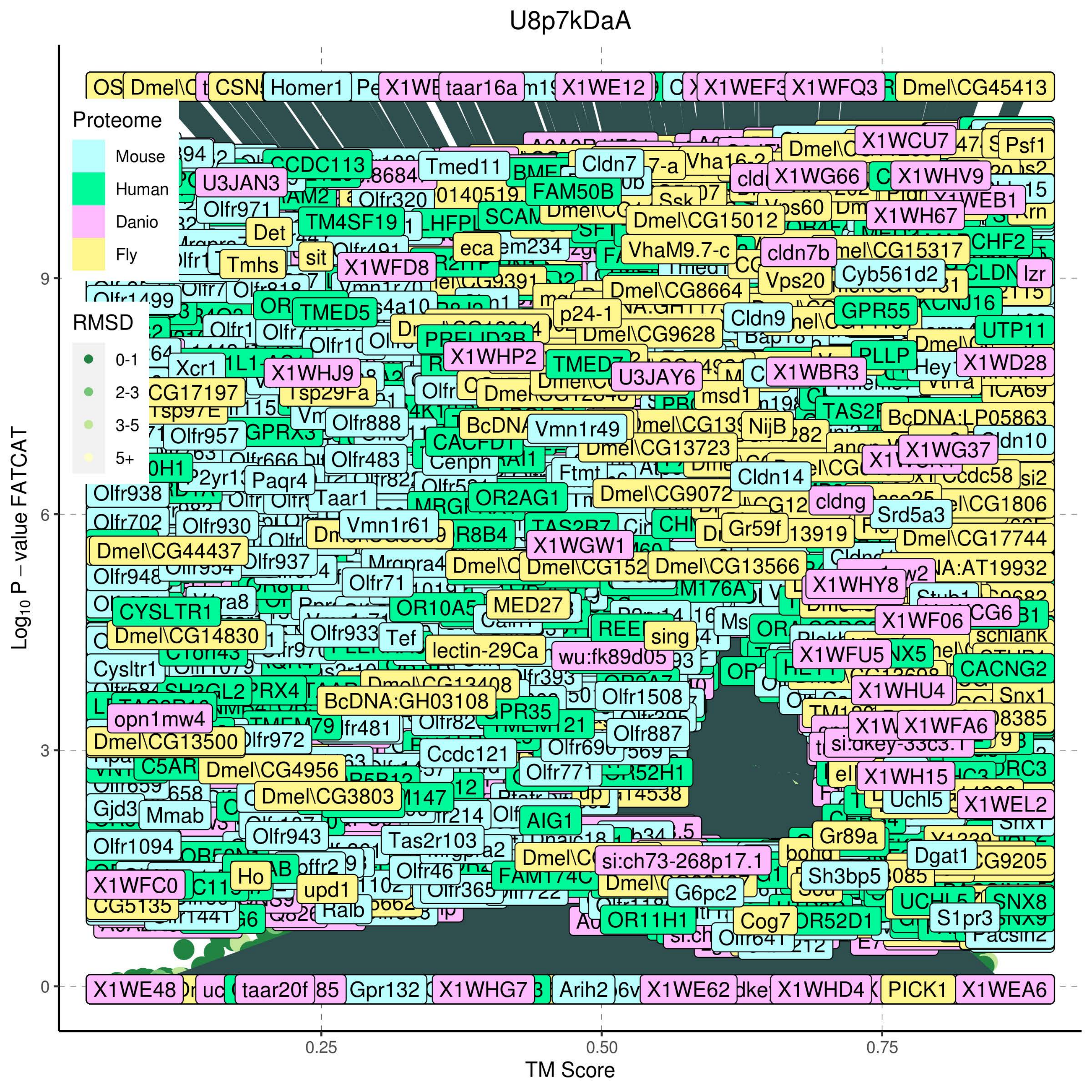
U8p7kDaA
Proteome
Mouse
Human
Danio
Fly
RMSD
0-1
2-3
3-5
5+
Log$_{10}$ P – value FATCAT
0
3
6
9
0.25
0.50
0.75
TM Score

GSDM alignment

>M35027.1:c153690-152956 Vaccinia virus Copenhagen complete genome
---------------------------------------------
MGNKNIKPSKENRLSILSKDKMDSFKRGSWATSSFREKSRATIQRFSSLRREHIKVDH--
PDKFLELKRGIYEIIQKSSSIDVDKRTKLMSNIKTMM--INPFMIEGLMTSLE-----------------NLD--
PDNKMSYSSVMILGEFDIINISDNE-------AAFEFINSLLKSLLLLNTRQLKLLEYSI-SNDLLYAHINA---
LEYIIKNTFNVPER------QLILRGQYLT---PIFSDLLKYAGLTIK--SNILMWNK--QFIKPVSDLYTSIRLLYCVTVX--
--------------------------------------------------------------------------------------------------------
>AY484669.1:c159516-158782 Rabbitpox virus complete genome
---------------------------------------------
MGNKNIKPSKENRLSILSKDKMDSFKRGSWATSSFREKSRATIQRFSSLRREHIKVDH--
PDKFLELKRGIYEIIQKSSSIDVDKRTKLMSNIKTMM--INPFMIEGLMTSLE-----------------NLD--
PDNKMSYSSVMILGEFDIINISDNE-------AAFEFINSLLKSLLLLNTRQLKLLEYSI-SNDLLYAHINA---
LEYIIKNTFNVPER------QLILRGQYLT---PIFSDLLKYAGLTIK--SNILMWNK--QFIKPVSDLYTSIRLLHCVTVX--
--------------------------------------------------------------------------------------------------------
>KY349117.1:c166272-165538 Horsepox virus strain MNR complete genome
---------------------------------------------
MGNKNIKPSKENRLSILSKDKMDSFKRGSWATSSFREKSRATIQRFSSLRREHIKVDH--
PDKFLELKRGIYEIIQKSSSIDVDKRTKLMSNIKTMM--INPFMIEGLMTSLE-----------------NLD--
PDNKISYSSVMILGEFDIINISDNE-------AAFEFINSLLKSLLLLNTRQLKLLEYSI-SNDLLYAHINA---
LEYIIKNTFNVPER------QLILRGQYLT---PIFSDLLKYAGLTIK--SNILMWNK--QFIKPVSDLYTSIRLLHCVTVX--
--------------------------------------------------------------------------------------------------------
>MG599038.1:c157906-157148 Buffalopox virus isolate Karachi 2005 complete genome
---------------------------------------------
MGNKNIKPSKENRLSILSKDKMDSFKRGSWATSSFREKSRATIQRFSSLRREHIKVDH--
PDKFLELKRGIYEIIQKSSSIDVDKRTKLMSNIKTMM--INPFMIEGLMTSLE-----------------NLD--
PDNKMSYSSVMILGEFDIINISDNE-------AAFEFINSLLKSLLLLNTRQLKLLEYSI-SNDLLYAHINA---
LEYIIKNTFNVPER------QLILRGQYLT---PIFSDLLKYAGLTIK--SNILMWNK--
QFIKPVSDLYTSMRLLHCVTESYKVIGMGX----------------------------------------------------------------------------
----------------------
>AY243312.1:c154675-153917 Vaccinia virus WR complete genome
---------------------------------------------
MGNKNIKPSKENRLSILSKDKMDSFKRGSWATSSFREKSRATIQRFSSLRREHIKVDH--
PDKFLELKRGIYKIIQKSSSIDVDKRTKLMSNIKTMM--INPFMIEGLMTSLE----------------NLD--
PDNKMSYSSVMILGEFDIINISDNE-------AAFEFINSLLKSLLLLNTRQLKLLEYSI-SNDLLYAHINA---
LEYIIKNTFNVPER------QLILRGQYLT---PIFSDLLKYAGLTIK--SNILMWNK--
QFIKPVSDLYTSMRLLHCVTESYKVIGMGX----------------------------------------------------------------------------
----------------------
>KC813500.1:c153312-152578 Cowpox virus strain MonKre08/4 complete genome
---------------------------------------------
MGNKNIKPSKENRLSILSKDKMDSFKRGSWATSSFREKSHATIQRFSSLRREHIKVDH--
PDKFLELKREIYEIIQKSSSIDVDKRTKLMSNIKTMM--INPFMIEGLMTSLE-----------------NLD--
PDNKMSYSSVMILGEFDIINISDNE-------AAFEFINSLLKSLLLLNTRQLKLLEYSI-SNDLLYAHINA---

LEYIIKNTFNVPER------QLILRGQYLT---PIFSDLLKYAGLTIK--SNILMWNK--
KFIKPVSDLYTSMRLLHCVTVX------------------------------------------------------------------------------------------
---------------
>NC 055231.1:c163765-163031 Orthopoxvirus Abatino complete genome
---------------------------------------------
MGNKHIKPSKENRLSILSKDKMDSFKRGSWATSSFREKSHATIQRFSSLRREHVKVDH--
PDKFLELKREIYEIIQKSSSIDVDKRTKLMSNIKTMM--INPFMIESLMTSLE-----------------NLD--
PDNKMSYSSVMILGEFDIINISDNE-------AAFEFINSLLKSLLLLNTRQLKLLEYSI-SNDLLYAHINA---
LEYIIKNTFNVPER------QLILRGQYLT---PIFSDLLKYAGLTIK--SNILMWNK--
KFIKPVSDLYTSMRLLHCVTVX------------------------------------------------------------------------------------------
---------------
>MN912466.1:c156293-155559 Ectromelia virus WH complete genome
---------------------------------------------
MGNKNIKPSKENRLSILSKDKMDSFKRGSCATSSFREKSHATIQRFSSLRREHIKVDH--
PDKFLVLKREIYEIIQKSSSIDVDKRTKLMSNIKTMM--INPFMIEGLMTSLE-----------------NLD--
PDNKMSYSSAMILGEFDIINISDNE-------ATFEFINSLLKSLLLLNTRQLKLLEYSI-SNDLLYANINA---
LEYIIKNTFNVPER------QLILRGQYLT---PIFSDLLKYAGLTIK--SNILMWNK--
KFIKPVSDLYTSMRLLHCVTVX------------------------------------------------------------------------------------------
---------------
>MH607143.1:c167386-166652 Akhmeta virus isolate Vani 2010 complete genome
---------------------------------------------
MGNKHIKPSKENRLSILYKDRMDSFKRGSWATSSFREKSHATIQRFSSLRREHVKVDH--
PDKFLELKREIYVIIQKSASIDVDKRTKLMSNIKTMM--INPFMIEGLMSSLE-----------------NLD--
PDNKMSYSSVMILGEFDIINISDNE-------KAFEFINSLLKSLLLLNTRQLKLLEYSI-SNDLLYAHINA---
LEYIIKNTFNVPER------QLILRGQYLT---PIFSDLLNYAGLTIK--SNILMWNK--
KFIKPVSDLYTSMRLLHCVTVX------------------------------------------------------------------------------------------
---------------
>DQ437594.1:c153307-152573 Taterapox virus strain Dahomey 1968 complete genome
---------------------------------------------
MGNKNIKPSKENRLSILYKDRMDSFKRGSWVTSSFREKSHATIQRFSSLRREHVKVDH--
PDKFLELKREIYAIIQKSSSIDVDKRTKLMSNIKTMM--INPFMIEGLMTSLE-----------------SLD--
PDNKMSYSSVMILGEFDIINISDNE-------AAFEFINSLLKSLLLLNTSQLKLLEYSI-SNNLLYTHINA---
LEYIIKNTFNVPER------QLILRGQYLT---PFFSDLLKYAGLTIK--SNILMWNK--
KFIKPVSDLYTSMRLLHCVTVX------------------------------------------------------------------------------------------
---------------
>KY358055.1:c145385-144651 Variola virus strain VD21 17th century complete genome
---------------------------------------------
MGNKNIKPSKENRLSILYKDRMDSFKRGSWATSSFREKSHATIQRFSSLRREHVKVDH--
PDKFLELKREIYAIIQKSSSIDVDKRTKLMSNIKTMM--INPFMIEGLMTSLE-----------------SLD--
PDNKMSYSSVMILGEFDIINISDNK-------AAFEFINSLLKSLLLLNTSQLKLLEYSI-SNDLLYTHINA---
LEYIIKNTFNVPER------QLILRCQYLT---PIFSDLLKYAGLTIK--SNILMWNK--KFIKPVSDLYTSMRLLHCVTVX-
------------------------------------------------------------------------------------------------------------
>DQ441432.1:c146026-145292 Variola virus strain Korea 1947 (Lee Masterseed) complete genome

-------------------------------------------
MGNKNIKPSKENRLSILYKDRMDSFKRGSWATSSFREKSHATIQRFSSLRREHVKVDH--
PDKFLELKREIYAIIQKSSSIDVDKRTKLMSNIKTMM--INPFMIEGLMTSLE-----------------SLD--
PDNKMSYSSVMILGEFDIINISDNK-------AAFEFINSVLKSLLLLNTSQLKLLEYSI-SNDLLYTHINA---
LEYIIKNTFNVPER------QLILRCQYLT---PIFSDLLKYAGLTIK--SNILMWNK--
KFIKPVSDLYTSMQLLHCVTVX--------------------------------------------------------------------------------------
--------------
>DQ441437.1:c146904-146170 Variola virus strain Sierra Leone 1969 (V68-258) complete genome
-------------------------------------------
MGNKNIKPSKENRLSILYKDRMDSFKRGSWATSSFREKSHATIQRFSSLRREHVKVDH--
PDKFLELKREIYAIIQKSSSIDVDKRTKLMSNIKTMM--INPFMIEGLMTSLE-----------------SLD--
PDNKMSYSSVMILGEFDIINISDNK-------AAFEFINSLLKSLLLLNTSQLKLLEYSI-SNDLLYTHINA---
LEYIIKNTFNVPER------QLILRCQYLT---PIFSDLLKYAGLTIK--SNILMWNK--
KFIKPVSDLYTSMQLLHCVTVX--------------------------------------------------------------------------------------
--------------
>MN240300.1:c166555-165827 Alaskapox virus complete genome
---------------------------------------------
MGNKNGKPFTENIQFLKDKRDSFKRGTWATSSFREKSHATIQRFSSLRRENVKVDQ--
HDKYMELKREIYAIIQKSSSIDINKRTKLLSNIKMMM--INPFMIEGLMTSLE-----------------NLD--
PDNKMSYSSVMILGEFDIINISDNE-------TAFEFINNLLKSLLLLNNRQLKLLEYSI-SNDLLYAHINA---
LEYIIKNTFNVPER------QLILRGQYLT---PIFSDLLKYAGITIK--SNILMWNK--KFIKPVSDLYTSMRLLHCVTVX-
---------------------------------------------------------------------------------------------------
>KU749310.1:c167631-166873 Skunkpox virus strain WA complete genome
------------------------------------
MGNNIKPSKEDKQLSSKGNRQSILFKDRMDSFKRGSWATSSFREKSVTTIQKFSSLRREHTKIEQ-
PDSKFIELKREIYEIINKSSSIDIDKRLMLIYNIKKMM--VQPFMLEGLMNSLE-----------------KMD--
PEDMISYSSVMILGEFDIVNIYDNK-------STFQFINGLLKSLHALNSKQLKLLEYSI-SNDLLYNQVTA---
LEYVIKNTFDVPER------QLILRGQYLT---PIFNDLLKYSGLTIK--SNILMWNK--KFIKPVSDLYTAIRLLYCVTIX--
--------------------------------------------------------------------------------------------------
>KU749311.1:c169057-168281 Volepox virus strain CA complete genome
-----------------------------
MGNNIKPSKENRQSLSKENRQSLKKNRQSFLFNDGMDTFKRGSWATSSFREKSASTIHKFSSLRREHTKIDQP
-DSKFIELKREIYEIINKSSSIDIDKRIMLVYNIKKMM--VNPFMIEGLMNSLE-----------------KMD--
PEDVISYSSVMIFGEFDMVNIHNK--------SAFQFINSLLKSLHALNNKQLKLLEYSI-SNDLLYNQVTA---
LEYVIKNTFDVPER------QLILRGQYLT---PIFNDLLKYSGLTIK--SNILMWNK--KFIKPISDLYTAIRLLYCVTVX--
--------------------------------------------------------------------------------------------------
>KU749309.1:c145369-144644 Raccoonpox virus strain 85A complete genome
--------------------------------------------
MGNNIKSSKENRQSGLFKDRMDSFKRGSWATSSFREKSVPTIHKFSSLRRERVTVQP--
DSKFIELKREIYAIINKSSSIDIDKRIILISNIKKMM--VNPFMIEGLMNSLE-----------------KMD--
PEDMISYSSVMILGEFDINMNDA---------PSFQFIISLLKSLHALNNKQLKLLEYSI-GNDLLYSQVTA---
LEYVIKNTFDVPER------QLILRGKYLT---PIFNDLLKYSGLTIK--SNILMWNK--KFIKPVSDLYIAIRLLYCMTIX--
--------------------------------------------------------------------------------------------------

>AY009089.1:c154402-154034 Camelpox virus CMS complete genome
-------------------------------------------------------------------------------------------------------------------------
-------------MM--INPFMIEGLMTSLE-----------------SLD--PDNKMSYSSMMILGEFDIINISDNE-------
VAFEFIHSLLKSLLLLNTSQLKLLEYSI-SNDLLYTHINA---LEYIIKNTFNVPER------QLILRGQYLT---
PIFSDLLKYAGLTIKX--------------------------------------------------------------------------------------------------
-----------------------------------
>MK910851.1:c154204-153836 Camelpox virus strain Negev2016 complete genome
-------------------------------------------------------------------------------------------------------------------------
-------------MM--INPFIIEGLMTSLE-----------------SLD--PDNKMSYSSMMILGEFDIINISDNE-------
VAFEFIHSLLKSLLLLNTSQLKLLEYSI-SNDLLYTHINA---LEYIIKNTFNVPER------QLILRGQYLT---
PIFSDLLKYAGLTIKX--------------------------------------------------------------------------------------------------
-----------------------------------
>MF001305.1:c159640-158771 NY 014 poxvirus strain 2013 complete genome
---
MGNKHSEPVFSESTISYTKRSMRVPCNEKPSTELKTGFNRLSLREKTVSSLQRGFNRLSLNEKTFSLSRGFNNR
LSLNEKTFSLQRGFSLRRFNSVRGES-SNSKLMELKKEIYETIQQANDMDIDKRIKLLYNIKSMM--
GNSFMLKGLIDSLE-----------------TLE--PTDGISYSSVMILGEFSILDQYTDGGK-----
STFKFVINLLKSLDSLNHHQQKLIEYAV-NNDILYDQITM---IEYIMKNNLDMENR------QFILKGKYTT---
PLFIDLLKHTGINV--ISNVLIWNK--KYTKSISNLYTAMRLLYSITVX------------------------------------------------
----------------------------------------------------
>MF001304.1:c160603-159728 Murmansk poxvirus strain LEIV-11411 complete genome
MGNKHSEPVFSASTISYTKRSSIRRVPCNENPSTELKTGEFNRLSLREKTVSSLQRGFNRLSLNEKTFSLHRGFN
NRLSLNEKTFSLQRGFSLRRFNSVRGES--SSKLMELKKEIHETIQQANNMDIDKRIKLLYNIKSMM--
GNSFMLKGLIDSLE-----------------TLE--PTDGISYSSVMILGEFSILDQYTDGGK-----
STFKFVINLLKSLDSLNHHQQKLIEYAV-NNDMLYDQITM---LEYIMKNNLDMENR------QFILKGKYTT---
PLFIDLLKHTGINV--ISNVLMWNK--KYTKSISNLYTAMRLLYSITVX----------------------------------------------
------------------------------------------------------
>HQ849551.1:c134784-134035 Yoka poxvirus strain DakArB 4268 complete genome
-----------------------------------------
MGNKNSDPVFTSTSCNNTTSLKRGQSLHLHRDLDKLSLNENTLFNRNFSFRRVNSMRIKPSNYSKLIELKKEIF
ETIKKANDMDIDKRIKLLHNIKSII--TNPFVLKGLIESLE-----------------TFE--
SVDGIKYSSVMILGEFNILDQYTGT-------STFQFVIDLLKSLYVLNNKQQKIIEYAI-NNDMLYEQIEM---
IEYIMTNVLDIDNY------HFILKGKYVT---PLFVELLKNTGINV-ISNNIIIWNK--KYTKNISNLYKAIRLLHSITVX--
--------------------------------------------------------------------------------------------------
>KU749310.1:172437-172985 Skunkpox virus strain WA complete genome
------------------------------------------------------------------------------------------------------
MQNSYIGQIYTRCHAILLIPDEYKHKFIKSLTHLI--GYNGRTYNRCT-----------------------
RDAITELTDIANEIDNT---------------DVKLITTFIISSIYKITRFQSFLVELAM-YHKVLDKIQHI---
MEYIIDNHSRIDNNGFTPYSDNDPDFNQVIA--
CIIYNLLRSSGMEINYSKYKAWWNPRHDYVDTFGLLYVIFTILSIIDDNX------------------------------------------
---------------------------------------------------------
>KU749309.1:150174-150722 Raccoonpox virus strain 85A complete genome
------------------------------------------------------------------------------------------------------
MRDSYICQIYTRCHSILSMPDDYKYKFIKSLNHLI--RYNGRTYNRCT------------------------

RDAITELSDIANSIDDT---------------DIKIITTFIISSIYKITRFQSFLVELAM-HHKVLDKVQHI---
MEYIIDNHSHSNTNGFTPYSIDDPDFNQVVA--
CIIYNLLRSSGMDINYSKCKAWWNPRHDYVDTFGLLYVIFTILSIIGDKX------------------------------------------
----------------------------------------------------------------
>KU749311.1:173861-174445 Volepox virus strain CA complete genome
-----------------------------------------------------------------------------------------------------------------
MLNDYIGQIYIRCRAILSIPDDYKHKFIKSLTHLI--RYNGRTYNRCT-------------------------
RDAITELSDIANNIGNN---------------DVKIITTFIISSIYKITRFQSFLVELAM-YHKVLDKIQHI---
MEYIIENHSHNDTNGFTPYSTDDTNFNQVIA--
CIIYNLLKSSGMEINYSECKAWWNPRHDYVDTFGLLYIIVTILLIIDGDQYNFYNKLEKNKX----------------------
-----------------------------------------------------------------------
>MF001304.1:c165763-165224 Murmansk poxvirus strain LEIV-11411 complete genome
-----------------------------------------------------------------------------------------------------------M--
LSVPSQYKDLLQEYYDIITSVPDEYRDIFVEYLIYLI--EYDGRISNKRT-------------------------
RDIKKELINIVGKVSDKT--------------RLMNFISFITESICRLTRFQSFLLELAS-YHNVLDEVQEI---
MGDIIENHCRMEIIGFTTYDINSIVL-------SIIYNMLKTSGMRVDYDTSKAWWNP--
EYRDMFGSLYMIVTTLTILKNIX-----------------------------------------------------------------------------------------
---------------
>MF001305.1:c164808-164269 NY 014 poxvirus strain 2013 complete genome
--------------------------------------------------------------------------------------------------------------
MLSVPSKYKDLLQEYYDIITSVPDEYRDIFVQYLIYLI--EYDGRISN----------------------------
KRTRDIKKELINIVGKVSDKT-----------RLMNFISFITESICRLTRFQSFLLELAS-YHDVLDEVQEI---
MGDIIKNHCRMEIIGFTAYDINSIVL-------SIIYNMLKTSGMRVDYDTSKVWWNP--
EYSDMFGSLYMIVTTLTILKNIX-----------------------------------------------------------------------------------------
---------------
>NP 051846.1 m132L Myxoma virus
-----------------------------------------------------------------------------------------------MGNLLYCCFTV--
EKTVPSIQRYVNGQVNLDNFLARHRKRSILLYLLKVI--RDDKLLNEVC-----------------------------
RVKKKYSVYLYGCKNPNA-----------EVVNFINYILVYLNALSCTQKRLLYIII-AENMISSFIGY---
INTVFTSNAVKFKDSE-----------------RCIEELFIETGLYI---SPQGSYKKHTEVMEELIYLYILARLFKQLL-----------
----------------------------------------------------------------------------------------------------
>NP 052021.1 gp132L Rabbit fibroma virus
-----------------------------------------------------------------------------------------------MGNLLYCCFVV--
DVTTPSIQRYVNEQVNLDKFLSRRKKKSILLYLLKVI--RNDKLLTEVYC------------------------------
SKKYSELYGCKNPNT-----------EVVNFINYILIYLNALSYTQKRLLYIII-AENMITSFIEY---
INAVFTSSSTVKFKDSE----------------RCIEELFIETGLYISSSQGYNKQHT--EVMEELIYLYILAKLFKQLL---------
------------------------------------------------------------------------------------------------------
>YP 009408097.1 Immunoprevalent protein Eptesipox virus
----------------------------------------------------------------------------MGNSTSVSKSCNNAVTHVSANSIRQHIL--
FNNFETLHKDIQSKIDLVNTFTPQTKNLIFRNLLIVI--TNSYHLQNLLDALE-----------------
QLEPMYVTDAYSEAILNEIGLCDKGIPNL--------SSIHFMIYLVSGLTKLTTKQSKILMEIVTDAKIFCHHVNV---
LEYIIKKNVEKLETVTSTLLEKYTKL-------PLEVTLFKESGLKIQ--GNTYIWDP--EHKKSICNLYTVIKIMSYIM----
-----------------------------------------------------------------------------------------------------------

>ref|NP 835465.2| chromosome 17 organism Homo sapiens gcode 1 chromosome 17 map 17q21.1 gasdermin-A Homo sapiens
--------------------------------------------------------------------------------------------------GDV--
HEGFRTLKEEVQRETQQVEKLSRVGQSSLLSSLSKLL--GKKKELQDLELALEG---------------ALD--
KGHEVTLEALPKDVLLSK-------------EAVGAILYFVGALTELSEAQQKLLVKSM-EKKILPVQLKL---
VESTMEQNFLLDKEGVFPLQPELLSSLGDEEL-TLTEALVGLSGLEVQRSGPQYMWDP--
DTLPRLCALYAGLSLLQQLTKAS---------------------------------------------------------------------------------------
----------------
>ref|XP 015154998.2| location chromosome chromosome 27 completeness complete organism Gallus gallus isolate bGalGal1 gcode 1 chromosome 27 sex female tissue type blood country USA: Fayetteville lat ls
-------------------------------------------------------------------------------------------------DGFA--
HTEVEALEEEVKEKCGILCQLAGELNAIVLKTIKAVM--RDRNLLEELSGKMEAVLD-DPDNCE------
LTTQSPDLEVLLSTLKGSPRCDLL-------------SLTEAISYTLDAFSELTEDQLLLLLESL-EVEIVPQQLKL---
VRSILEQSAELGAA------AGLLSSFSEQQQQLTTAVLELSGVELQGSGAAVCTE---
ESFPSLSALYAALHALHCLVGAGGRSARAD------------------------------------------------------------------------------
----------------------
>ref|XP 024836266.1| location chromosome chromosome 19 completeness complete organism Bos taurus isolate L1 Dominette 01449 registration number 42190680 breed Hereford gcode 1 chromosome 19 sex fem...
--------------------------------------------------------------------------------------------------GAV--
HEDFRTLKEEVQRETQEVEKLSPEGKSSLLSSLSKLL--GKKKELQDLELTLEE---------------ALG--
KGHEETLEALPKNVLLSK-------------GAMDAILYFLGALAELSEAQQKLLVKSM-EKKILPVQLKL---
VESTMEQNFLQDKEGVFPLRPELLSSLGEEEL-TLTEALVGLSGLEVQRSGPQYTWDP--
DTLPRLCALYAGLSLLRLLTKAS-----------------------------------------------------------------------------------------
--------------
>ref|XP 014975037.1| location chromosome chromosome 16 completeness complete organism Macaca mulatta isolate AG07107 bio material Coriell:AG07107 gcode 1 chromosome 16 sex female tissue type fibroblaa
--------------------------------------------------------------------------------------------------GDV--
HEGFGTLKEEVQRETQQVERLSQAGQSSLLSSLSKLL--GKKKELQDLELALEG---------------ALD--
KGHEVTLEALPKDVLLSK-------------EAVGAILYFVGALTELSEVQQKLLVKSM-EKKILPVQLKL---
VESTMEQNFLQDKEGVFPLHPELLSSLGEEEL-TLTEALVGLSGLEVQRSGPQYMWDP--
DTLPRLCALYAGLSLLQQLTKAS---------------------------------------------------------------------------------------
----------------
>ref|XP 001171222.1| location chromosome chromosome 17 completeness complete organism Pan troglodytes isolate Yerkes chimp pedigree *C0471 (Clint) gcode 1 chromosome 17 sex male tissue type blood devs
--------------------------------------------------------------------------------------------------GDV--
HEGFRTLKEEVQRETQQVEKLSRVGQSSLLSSLSKLL--GKKKELQDLELALEG---------------ALD--
KGHEVTLEALPKDVLLSK-------------EAMGAILYFVGALTELSEAQQKLLVKSM-EKKILPVQLKL---
VESTMEQNFLLDKEGVFPLQPELLSSLGDEEL-TLTEALVGLSGLEVQRSGPQYMWDP--
DTLPRLCALYAGLSLLQQLTKAS---------------------------------------------------------------------------------------
----------------

>ref|XP 024429158.1| chromosome Unknown completeness complete organism Desmodus rotundus isolate DRU21DN04 gcode 1 chromosome Unknown sex male tissue type muscle/skin sample dev stage adult country Us
----------------------------------------------------------------------------------------------------GEV--
HQDFRMLKEEIQRETQEVEKLSRMGQSSLLTSLSKLL--GKKKELQDLELTLEG----------------ALD--
KGHEVALEALPKDVLLSK--------------EVVGAILYFLGALTELSEAQQKLLVKSM-EKKILPVQLKL---
VESIMEQNFLLDKEGVFPLQPNLLSCLGEEEL-TFTEALVGLSGLEVQRSGPQYMWDP--
DTLPDLCALYAGLSFLQLLTKAS----------------------------------------------------------------------------------------
--------------
>ref|XP 016020091.2| chromosome Unknown completeness complete organism Rousettus aegyptiacus isolate mRouAeg1 gcode 1 chromosome Unknown sex male tissue type muscle dev stage adult country USA: Berkes
----------------------------------------------------------------------------------------------------GEV--
HEDFRTLKEEIQRETQEVEKLSHLGQSSLLTSLSKLL--GNRKELQDLELTLEG----------------ALG--
KGHEVTLEALPKDVLLSK--------------EALGAILYFLGALTELSEVQQKLLVKSM-EKKILPVQLKL---
VESTMEQNFLQDKEGVFPLQPDLLSSLGEEDL-TLTEALVGLSGLEVQRSGPQYTWDP--
DTLPHLCALYAGLSFLQLLTKAS----------------------------------------------------------------------------------------
--------------
>ref|XP 006924750.1| chromosome Unknown completeness complete organism Pteropus alecto gcode 1 chromosome Unknown sex male tissue type kidney country Australia: Anglican Church Grammar School East Bro
----------------------------------------------------------------------------------------------------GEV--
HEDFRTLKEEIQRETQEVEKLSHVGRSSLLTSLSKLL--GNRKELQDLELTLEG----------------ALG--
KGHEVTLEALPKDVLLSK--------------EALGAILYFLGALTELSEVQQKLLVKSM-EKKILPVQLKL---
VESTMEQNFLQDKESVFPLQPDLLSSLGEEDL-TLTEALVGLSGLEVQRSGPQYTWDP--
DTLPCLCALYAGLSLLQLLTKAS----------------------------------------------------------------------------------------
--------------
>ref|XP 035935879.1| chromosome Unknown completeness complete organism Halichoerus grypus isolate 241Hg ecotype Northwest Atlantic gcode 1 chromosome Unknown sex male tissue type blood dev stage molts
----------------------------------------------------------------------------------------------------KEE--
HGDFKTLKEEVQREIQEVKKLSRVGQSSLLTSLSKLL--GKKKELQDLELMLEG----------------ALD--
KGHEVTPEALPKDVLVSK--------------EAMGGILYFLGALIELSEAQQKLLIKSV-EKKILPMQLKL---
VESTMEQSFLQDKDGVFPLQPDLLSSLGEEEL-TLTEALVGLSGLEVQRSGPQYTWDP--
DTLPRLCALYAGLSLLQLLTKAS----------------------------------------------------------------------------------------
--------------
>ref|XP 004633937.1| chromosome Unknown completeness complete organism Octodon degus isolate 3935 gcode 1 chromosome Unknown sex female gasdermin-A Octodon degus
----------------------------------------------------------------------------------------------------GEV--
HEDFKTLKEEVRRETQEVDKLSRAGQGSLLSSLSNLL--GKKKELQDLELTLEG----------------ALD--
KGHEVTLEALPKDVLLSK--------------ESMGAILYFLGALTVLSEAQQKLLVKSV-EKKLLPVQLKL---
VESTLEQNFLQDKEGVFPLRPDLLSSLGEEEL-TLTEALVGLSGLEVQRSGPQYTWDP--
DTLPRLCALYASLSLLQLLARSP----------------------------------------------------------------------------------------
-------------

>ref|XP 029428328.1| location chromosome chromosome 12 completeness complete organism Rhinatrema bivittatum gcode 1 chromosome 12 gasdermin-A Rhinatrema bivittatum
-----------------------------------------------------------------------------------------DYLRILENF----
QEGFPAVMAQIEKECVEFTLLSPEVAEQFLAAFLLLI--KDPKTSADVESRLEQSLY-C-SARESKA---HPE--
PSIARLLGYLGLSSRPVVI------------PLVRAVLFFLDAVSELDEDGTSLLVESV-ERQDTGQQLKR---
VQKVLEEELFSVRQGSL------------------------AGDEAAGGVSPEGWSP--AVQSCAMALYAALYALHVLSSC--
----------------------------------------------------------------------------------------------------
>XP 032891479.1 gasdermin-A Amblyraja radiata
------------------------------------------------------------------------------------------------SLFC--
QDTANVNGDEVFEICCELQCLDKGLKQRLLGLVCQII--EDSELLSILSDVLCEA-C-A-GTDYTLSELEKLD--
KKRRECGENLLAIWSEDQLKNAVED-------NLLKAVSILFAALEDLPPTTLPVLIQSL-KMQILPQQLTL---
VTGILKDLDHSQVDQALKVETE---SFTEDAF-GITAEMLTDVGLHLERDSVQKTREP--S----
LHELSVALYCLNALSSN------------------------------------------------------------------------------------------------
------
>sp|Q5Y4Y6|GSDA3 MOUSE Gasdermin-A3 OS Mus musculus OX 10090 GN Gsdma3 PE 1 SV 1
-------------------------------------------------------------------------------------------------GEM--
HEDFKTLKEEVQRETQEVEKLSPVGRSSLLTSLSHLL--GKKKELQDLEQKLEG----------------ALD--
KGQKVTLEALPKDVLLSK--------------DAMDAILYFLGALTELTEEQLKILVKSL-EKKILPVQLKL---
VESTLEQNFLQDKEGVFPLQPDLLSSLGEEEL-TLTEALVGLSGLEVQRSGPQYAWDP--
DTRHNLCALYAGLSLLHLLSRKSNALTYCALS------------------------------------------------------------------------
--------------------
>sp|Q9EST1|GSDMA MOUSE Gasdermin-A OS Mus musculus OX 10090 GN Gsdma PE 2 SV 1
-------------------------------------------------------------------------------------------------GEM--
HEDFKTLKEEVQRETQEVEKLSPVGRSSLLTSLSHLL--GKKKELQDLEQTLEG----------------ALD--
KGHEVTLEALPKDVLLSK--------------DAMDAILYFLGALTVLSEAQQKLLVKSL-EKKILPVQLKL---
VESTMEKNFLQDKEGVFPLQPDLLSSLGEEEL-ILTEALVGLSGLEVQRSGPQYTWDP--
DTLPHLCALYAGLSLLQLLSKNS---------------------------------------------------------------------------------------
--------------
>sp|Q32M21|GSDA2 MOUSE Gasdermin-A2 OS Mus musculus OX 10090 GN Gsdma2 PE 2 SV 1
-------------------------------------------------------------------------------------------------GEM--
HEDFKTLKKEVQQETQEVEKLSPVGRSSLLTSLSHLL--GKKKELQDLEQMLEG----------------ALD--
KGHEVTLEALPKDVLLLK--------------DAMDAILYFLGALTELSEEQLKILVKSL-ENKVLPVQLKL---
VESILEQNFLQDKEDVFPLRPDLLSSLGEEDQ-ILTEALVGLSGLEVQRSGPQYTWNP--
DTCHNLCALYAGLSLLHLLSRDS--------------------------------------------------------------------------------------
---------------
>ref|NP 001159430.1| chromosome 17 organism Homo sapiens gcode 1 chromosome 17 map 17q21.1 gasdermin-B isoform 3 Homo sapiens
----------------------------------------------------------------------------------------------------------------------------
-----DVLNSLAKCL--GKEDIRQDLEQRVSEVLI-S-G---------ELHMEDPDKPLLSSLFNAAGVLVE------------
ARAKAILDFLDALLELSEEQQ-FVAEAL-EKGTLPLLKDQ---VKSVMEQNWD----------------------------------
ELASSPPDMDYDP---EARILCALYVVVSILLELAEGPTSVSS---------------------------------------------------------
----------------------------------------
>ref|NP 001231146.1| chromosome 19 organism Bos taurus gcode 1 chromosome 19 map 19 gasdermin-B Bos taurus

--------------------------------------------------------------------------------------------------------------------------------
-----EALNCFTKCL--SSEEELQDLEQRVSEVQC-S-G---------ELQMNSPANSLISSLFNAAGALIE-------------
ARAETIWDVLDALMELSEYRQ-FVAEVL-DKGELRPVKDK---VESILEEN-------------------------
RCEGPLDVSCD----------------PEARTLCALYVVVSILLQLSEKPTSVSS---------------------------------------------
---------------------------------------------------

>ref|XP 009430521.1| location chromosome chromosome 17 completeness complete organism Pan troglodytes isolate Yerkes chimp pedigree *C0471 (Clint) gcode 1 chromosome 17 sex male tissue type blood devs

--------------------------------------------------------------------------------------------------------------------------------
-----DVLNSLAKCL--GKEEIRQDLEQRVSEVLISR-----------ELHMEDPDKPLLSSLFNAAGVLVE-------------
VRAKAILDFLDALLELSEEQQ-FVAEAL-EKGTLPLLKDQ---VKSVMEQNWD----------------------------------
ELASSPPDMDCDP---EARILCALYVVVSVLLELAEGPTSVSS---------------------------------------------------------
-----------------------------------------

>ref|XP 023358160.1| location chromosome chromosome 4 completeness complete organism Sarcophilus harrisii gcode 1 chromosome 4 cell line 91H gasdermin-B Sarcophilus harrisii

--------------------------------------------------------------------------------------------------------------------------------
-----AVLDLLTRLL---QDEALGEMEGKVAMALF-T-E---------ELS--DPENPLLSVLFDASGSSMQ-------------
NQAEGILDFLEALTELKEEQQKILPKVV-EKGLLSETMKL---VEDLLRQSETFSLKAD-----------------
SITHTLVGICDPEGKLEQP-VTWDS--ETCHSLCALYTVLSVLMLLAQGSVSVP--------------------------------------
---------------------------------------------------------------

>ref|XP 024897515.1| chromosome Unknown completeness complete organism Pteropus alecto gcode 1 chromosome Unknown sex male tissue type kidney country Australia: Anglican Church Grammar School East Bro

--------------------------------------------------------------------------------------------------------------------------------
-----DALSCLTKCL--TKDEQLQDLEQRMSEVLISG-----------ELQMDSPAGPLISSLFNAAGILVE-------------
TRTEAIMDFLDAVMELFE-EKELVGEAL-EKGTLPLLKEQ---VESVLDQNGGEQPE-------------------------------
-------CVGCDP---EAQILHALYVAVSILLQLSEKPASASS---------------------------------------------------------
-----------------------------------

>ref|XP 036078993.1| chromosome Unknown completeness complete organism Rousettus aegyptiacus isolate mRouAeg1 gcode 1 chromosome Unknown sex male tissue type muscle dev stage adult country USA: Berkes

--------------------------------------------------------------------------------------------------------------------------------
-----DALSCLSKCL--TKDEQLQDLEQRVSEILISG-----------ELQMDSPAGLLINSLFNASGILVE-------------
TRTEAIMDFLDALMELSE-EKELVAEAL-EKGTLPLLKE----------QNCGEQPE------------------------------------
YVGWDP---EAQVLRTIYAAVSILLLLSEKPASVSC--------------------------------------------------------------
-----------------------------

>ref|XP 024429114.1| chromosome Unknown completeness complete organism Desmodus rotundus isolate DRU21DN04 gcode 1 chromosome Unknown sex male tissue type muscle/skin sample dev stage adult country Us

--------------------------------------------------------------------------------------------------------------------------------
-----NVLSCLTKCL--TSDGELQALEERVSEVLV-S-E---------ELQVEGPADPLISSLFNAAGILVE-------------
ACIEAILDFLDALKELSEEKE-LVAETL-EKGTLPLLKDQSGGLKSVLEQEWGEQPRG-------------------------------
-------VGRDP---EAQTVCTLYLVVSILLQLSEKPAPASSC---------------------------------------------------------
-----------------------------------

>ref|XP 014975035.1| location chromosome chromosome 16 completeness complete organism Macaca mulatta isolate AG07107 bio material Coriell:AG07107 gcode 1 chromosome 16 sex female tissue type fibroblaa
-------------------------------------------------------------------------------------------------------------------------------
-----DVLNFLTKCL--GREELWQDLEQKISEVLI-F-G---------ELQMEDPVNPLLSSLFNAAGILVG------------
ARAEAILDFLDALLELSEEQH-LVAEAL-EKGTLPLLKDQ---VKPIMEQNWD----------------------------------
ELASSPHDMDCDP---EARIPCALYVVVSVLLELAEGTTSVSS--------------------------------------------------------
-------------------------------------------
>ref|XP 035935913.1| chromosome Unknown completeness complete organism Halichoerus grypus isolate 241Hg ecotype Northwest Atlantic gcode 1 chromosome Unknown sex male tissue type blood dev stage molts
-----------------------------------------------------------------------------------------------------------
DVLSCLRGQLPLSAKVFEVLISGSRHLEMEGLVGLL--------------------------------------------ISSLFNAAGFLVK-
------------AHTESILDVLDAL--ITSEEQHLVAEAL-EKGTLPLLKDQ---VRSVLEQNWSEQPH---------------------
-----------------DVGWDP---DAXLLCALYVALSLLLQLGEKPTSVPS--------------------------------------------------
-------------------------------------------------
>ref|NP 113603.1| chromosome 8 organism Homo sapiens gcode 1 chromosome 8 map 8q24.21 gasdermin-C Homo sapiens
---------------------------------------------
LLPSFHTISPTLFNASSNDMKLKPELFLTQQFLSGHLPKYEQVHILPVG-RIEEPF----
WQNFKHLQEEVFQKIKTLAQLSKDVQDVMFYSILAML--RDRGALQDLMNMLELDSS-G-----------HLD--
GPGGAILKKLQQDSNHAWF-------------NPKDPILYLLEAIMVLSDFQHDLLACSM-EKRILLQQQEL---
VRSILEPNFRYPWSIPFTLKPELLAPLQSEGL-AITYGLLEECGLRMELDNPRSTWDV--
EAKMPLSALYGTLSLLQQLAEA------------------------------------------------------------------------------------------
----------------
>ref|XP 028708741.1| location chromosome chromosome 8 completeness complete organism Macaca mulatta isolate AG07107 bio material Coriell:AG07107 gcode 1 chromosome 8 sex female tissue type fibroblasta
---------------------------------------------
WLPSFHTISPTLFNASPKDMKLKPELFLKPKFLRGHLPKYEQVCILPVG-RIEEPF----
WQNFKHLQEEVFQKIKTLAQLSKDVQDVVFYSILAML--GDRGALQDLMDMLELDSS-G-----------HLD--
GPGGAILKKLQQYSNRAWF-------------NPKDPILYLLEAIMVLNDIQHDLLACSV-EMRILLQQQEL---
VRSILEPNFRYPWSIPFTLKPELLAPLQSEGL-AITYGLLEECGLRMELDNPRSTWDL--
EAKIPLSALYGTLSLLQQLAEA--------------------------------------------------------------------------------------------
--------------
>ref|NP 001039469.1| chromosome 14 organism Bos taurus breed L1 Hereford gcode 1 chromosome 14 map 14 gasdermin-C Bos taurus
--------------------------------------------------------------
ILSISDDNKRKAFFVGKVQRQSAMGTKRRPILPIGRIQEPI--
SRDFKSLQNEVSREIEAVAELPRDIRDALFHTILAKL--KDQGALQDLTDMLDGNLW------------------
DHTGSFLSEMREDSRNVWL-------------ESRPCVIYLLEALLVLSDIQHELLAWSM-EKRILPQQREL---
VESILEPNFRYPWNIPFTLDPKLLASLQDEGL-AVTFGLLQECGLRVAPDNPKGTWDL--
EAKKPLSALYGSLSVLQQLAEA-------------------------------------------------------------------------------------------
---------------

>ref|XP 031803207.1| location chromosome chromosome 1 completeness complete organism Sarcophilus harrisii gcode 1 chromosome 1 cell line 91H gasdermin-C Sarcophilus harrisii
----------------------------------------------------
HTEIGLGSMRGMEEKLEWKDYPKLSLFLETIEETQQGFGWKQQDRNNCQR--
VRDFQRLLEEVTFQKKDLVRMPGKQRTTLLCALHDLL--GNRTALQDLEDKLLHVLD-SGIWG-------KME--
GPGNTILSNLQDSSDSPVN-------------GYITGFLYLLEALAALSDNQHDLLAQSL-EKKILSQQLKL---
VKSILETNFNQFHS-TFILPPEILSLIDEEL--NLSLELVEDCGLQLQRTEQMLILDP--
EGKNPLCALYGALSCLLSLSED-----------------------------------------------------------------------------------------
--------------
>ref|XP 036088919.1| chromosome Unknown completeness complete organism Rousettus aegyptiacus isolate mRouAeg1 gcode 1 chromosome Unknown sex male tissue type muscle dev stage adult country USA: Berkes
-----------------------------------------------------------------FRLQLQSFTGGREIQPKSGECEIQILSGRAQRSC-
-WLDFGRLQEEVS-WYVKKAQLLEDMGAVVFSSILAML--GDWEALEGLEDKLEQKPL-G-----------HLD--
GPGGVILNALRESSSYQWF-------------NSEDSILYLLEAINGLSDLQHDLLAWSM-EKKVLLQQRDL---
VRSILEPNFKYPWNIPFTMKPELLAPLRGEAL-AITYGLLEECGLTMEPNSPRSTWDL--
EAKKPLSALYAALSVLQQLAEA-----------------------------------------------------------------------------------------
--------------
>ref|XP 024427836.1| chromosome Unknown completeness complete organism Desmodus rotundus isolate DRU21DN04 gcode 1 chromosome Unknown sex male tissue type muscle/skin sample dev stage adult country Us
----------------------------------------------------------------------------------------------------
DFKRLHEEVSKRVIAAAELSKDIKGVVFSNILAML--GDRGALQDLVHMLEQEPLKG-----------HLN--
GPGGTILNELQKDSSFAWN-------------GSQYLILYLLEAMMVLSDVQLCLLAPSM-EKKILSQQRDL---
VRSILEPNFKYPWSIPFTLKPELLAPLQGEGL-AITYGLLDECGLKMELHSPRSTWDL--
EAKKPLSALYGTLSVLQQLAEA-----------------------------------------------------------------------------------------
--------------
>ref|XP 019678683.1| location chromosome chromosome F2 completeness complete organism Felis catus isolate Cinnamon breed Abyssinian gcode 1 chromosome F2 sex female gasdermin-C Felis catus
-------------------------------------------------------------------
AVNNGPLYTMSVSDVRKPLDLVLHSPGRIQRVL--WQDFKQLHEEIFQEMEALAQLSKDTQDSIFHTILNML--
GNREALQDLTDTLDGSPL-D-----------LLD--DFGGTILNEMQPDTRDLWI-------------
QARFHIIYLLEVIMVLSDTQHDLLAQSM-EKRILLQQQKL---VRSILEPNFKYCWDIPFTLKPELLAPLQGEDL-
AITYGLLEECGLKMELNSPRSTWDL--ETKQPLSALYGTLSLLQQLTGSPERPGPTPVYAESVPDELGGV---------
------------------------------------------------------------------------
>ref|XP 001153860.1| location chromosome chromosome 8 completeness complete organism Pan troglodytes isolate Yerkes chimp pedigree *C0471 (Clint) gcode 1 chromosome 8 sex male tissue type blood dev ss
-------------------------------------------
LLPSFHTISPTLFNASSNDMKLKPELFLTQQFLSGHLPKYEQVHILPVG-RIEERF----
WQNFKHLQEEVFQKIKTLAQLSKDVQNVVFYSILAML--RDRGALQDLMNMLELDSS-G-----------HLD--
GPGGAILKKLQQDSNHAWF-------------NPKDPILYLLEAIMVLSDFQHDLLACSV-EKRILLQQQEL---
VRSILEPNFRYPWSIPFTLKPELLAPLQSEGL-AITYGLLEECGLRMELDNPRSTWDV--

EAKMPLSALYGTLSLLQQLAEA-------------------------------------------------------------------------------------
----------------
>ref|XP 021073018.1| location chromosome chromosome 17 completeness complete organism Mus pahari gcode 1 chromosome 17 sex female tissue type tail gasdermin-C Mus pahari
------------------------------------------------------------RPMYLSGHFEILRHHEEIIRIGSRIEYISPIGTIEEPT--
HLDFQCLQNEVSQKTMLLAELSKDVQEVVFSSFLHMLCEGDSEVLYDLMKMLELNQL-E-----------HMD--
GPGGKILEELQKDRSPSWI-------------DLKDLILNLLQALMVLSNTQLSLLAQSV-EMRLLLQQQEL---
VKSILQPNYKYPWHIPFTLQPQLLAPLQGEGL-AITYELLEECGLKMELNNPRSTWHL--
EVKMPLSALYASLSLLQQLSEA-------------------------------------------------------------------------------------
---------------
>ref|NP 081236.1| chromosome 15 organism Mus musculus strain C57BL/6 gcode 1 chromosome 15 map 15 gasdermin-D Mus musculus
------------------------------------------------------------------------------------------DG-IDEEELIE--
AADFQGLYAEVKACSSELESLEMELRQQILVNIGKIL--QDQPSMEALEASLGQGLC-SGGQVE------PLD--
GPAGCILECLVLDSGELVP-------------ELAAPIFYLLGALAVLSETQQQLLAKAL-ETTVLSKQLEL---
VKHVLEQSTPWQEQSSVSLPTVLLGDCWDEK--NPTWVLLEECGLRLQVESPQVHWEP--
TSLIPTSALYASLFLLSSLGQKPC------------------------------------------------------------------------------------
--------------
>ref|NP 079012.3| chromosome 8 organism Homo sapiens gcode 1 chromosome 8 map 8q24.3 gasdermin-D Homo sapiens
------------------------------------------------------------------------------------------DGVPAEGAF----
TEDFQGLRAEVETISKELELLDRELCQLLLEGLEGVL--RDQLALRALEEALEQGQS-L-GPVE------PLD--
GPAGAVLECLVLSSGMLVP-------------ELAIPVVYLLGALTMLSETQHKLLAEAL-ESQTLLGPLEL---
VGSLLEQSAPWQERSTMSLPPGLLGNSWGEG--APAWVLLDECGLELGEDTPHVCWEP--
QAQGRMCALYASLALLSGLSQEPH------------------------------------------------------------------------------
-------------------
>ref|NP 001039625.1| chromosome 14 organism Bos taurus breed Hereford gcode 1 chromosome 14 map 14 gasdermin-D Bos taurus
------------------------------------------------------------------------------------------DG-STEHWLAT--
TDDYQGLRAEVKAWAMGLEGLSKGLCGQLLGGLGQVL--RDEPALQALEDSLEQGLC-S-GPVE------PRD--
GPVGAILESLVLPSGQLET-------------ELAGPVFYLLQALAVLSEAQHVLLAEVL-EMGALSGTFWL---
VESLVEQSSPWQEHRAVSLPPERLGDSWGSE--APGWALLEACGLEPQVGTPQVCWEP--
GARGCACALYACLALLLRLSQLC------------------------------------------------------------------------------------
----------------
>ref|XP 015001615.2| location chromosome chromosome 8 completeness complete organism Macaca mulatta isolate AG07107 bio material Coriell:AG07107 gcode 1 chromosome 8 sex female tissue type fibroblasta
------------------------------------------------------------------------------------------DGVPAEGTF----
TEDFQGLREEVEAISKELELLDRELCQLLLEGLEGVL--RDQLALRALEEALEQGLS-L-GPVE------SLD--
GPAGAVLECLVLESRMLVP-------------ELAVPVVYLLGALTMLSETQHKLLAEAL-ESQTLLGPLEL---
VGSLLEQSAPWQERSTVSLPPGLLGSSWGEG--APAWVLLEECGLELGEDTPHVCWEP--
QAQGRMCALYASLALLSGLSQEPH------------------------------------------------------------------------------
-------------------

>ref|XP 006913014.1| chromosome Unknown completeness complete organism Pteropus alecto gcode 1 chromosome Unknown sex male tissue type kidney country Australia: Anglican Church Grammar School East Bro
--------------------------------------------------------------------------------------------DG-PTENCLVP--
TEDFQGLQVEVKAWASALENGSATLCGQLLGGLGQVL--RDEPALQALEESLEQGLC-S-GRAE------PLH--
GPAGAVLECLVLHCRTLVE-------------ELARPVFYLLEALAALSETQHMLLAEVL-ETGALPESVEL---
VETLLEQSTPWQEREAISLSPRLLGSNWGSE--APAWVLLEECGLKLQVGAPQVCWEP--
EAQGPTCALYACLALLLKLSQLR----------------------------------------------------------------------------------------
----------------
>ref|XP 036089037.1| chromosome Unknown completeness complete organism Rousettus aegyptiacus isolate mRouAeg1 gcode 1 chromosome Unknown sex male tissue type muscle dev stage adult country USA: Berkes
--------------------------------------------------------------------------------------------DG-PAEHWLVP--
TEDFQGLQMEVRAWARALEDASATLCGQLLGALGQVL--WDEPALQALEESLEQGLC-S-GRAE------PLH--
GPAGAVLECLVLRCGTLVE-------------ELAGPVFYLLGALAALSETQHVLLAKAL-ETGALPEWVKL---
VETLLEQSTPWQEREAISLPSELLESSWGSE--APAWVLLEECGLKLQVGAPQVCWEP--
EAQGPTCALYACLALLLRLSQLR----------------------------------------------------------------------------------------
----------------
>ref|XP 024428237.1| chromosome Unknown completeness complete organism Desmodus rotundus isolate DRU21DN04 gcode 1 chromosome Unknown sex male tissue type muscle/skin sample dev stage adult country Us
--------------------------------------------------------------------------------------------DG-LDEDQVVP--
TEDFQGLQAEVQVWAGPLESLSKTLCEQLLRGLAQVL--RDELALQTLEGSLEQGLC-C-GRAE------PLQ--
GPAGHVLECLVLPCRTLVN-------------ELAGPVTYLLGALAALSEAQRMLLAEVL-EASALPEPFKL---
VEKLLEQSSPWQECRALSLPPGLLGGSWGPE--VPAWILLEACGLELQVDAPQVSWKP--
EAQGCTCALYACLALLSKLSQLC----------------------------------------------------------------------------------------
----------------
>ref|XP 009454389.2| location chromosome chromosome 8 completeness complete organism Pan troglodytes isolate Yerkes chimp pedigree *C0471 (Clint) gcode 1 chromosome 8 sex male tissue type blood dev ss
--------------------------------------------------------------------------------------------DGVPAEGAF----
TEDFQGLRVEVETISKELERLDRELCQLLLKGLEGVL--RDQLALRALEEALEQGQS-L-GPVE------PLD--
GPAGAVLECLVFSSGVLVL-------------ELAIPVVYLLGALTMLSETQHKLLAEAL-ESQTLLGPLEL---
VGSLLEQSAPWQERSTMSLPPGLLGSSWGEG--APAWVLLDECGLELGEDTPHVCWEP--
QAQGRMCALYASLALLSGLSQEPH------------------------------------------------------------------------------------
-------------------
>ref|XP 023104116.1| location chromosome chromosome F2 completeness complete organism Felis catus isolate Cinnamon breed Abyssinian gcode 1 chromosome F2 sex female gasdermin-D Felis catus
--------------------------------------------------------------------------------------------DG-DIEDQMPV--
TEDFQGLKVEVSVHADGLKGLSGELCGQILAGLMKVL--REEPALESLQEELEQGLC-C-GWVA------SPD--
APGGAILECLVQSSGKVEE-------------ELARPILYLVQALTELNETQRVLLAEAL-ETGDLSGQSRL---
VQSVLEQSSPWKEHRAVSLPQELLGSSWDSE--APAWVLLEECGLELRVDVPQVHWQP--

DAQGRTSALYACLVLLPHLSQDQLNETQRVLLAEALETGDLSGQSRLVQSVLEQSSPWKEHRAVSLPQELLGS
SWDSEAPAWVLLEECGLELRVDVPQVHWQPDAQGRTSALYACLVLLPHLSQDRA
>ref|NP 001120925.1| chromosome 7 organism Homo sapiens gcode 1 chromosome 7 map 7p15.3 gasdermin-E isoform a Homo sapiens
----------------------------------------------------------------------------------------DAAHGISSQDGP--
LSVLKQATLLLERNFHPFAELPEPQQTALSDIFQAVL--FDDELLMVLEPVCDDLVS-GLSPTVAVLG--ELK--
PRQQQDLVAFLQLVGCSLQGGCPGPED-AGSKQLFMTAYFLVSALAEMPDSAAALLGTCC-KLQIIPTLCHL--
-LRALSDDGVSDLED------PTLTPLKDTERF-
GIVQRLFASADISLERLKSSVKAVILKDSKVFPLLLCITLNGLCALGREHS-------------------------------------------
---------------------------------------------------------
>ref|NP 061239.1| chromosome 6 organism Mus musculus strain C57BL/6 gcode 1 chromosome 6 map 6 gasdermin-E precursor Mus musculus
----------------------------------------------------------------------------------------DGGQGISSQDGP--
LRVVKQATLHLERSFHPFAVLPAQQQRALFCVLQKIL--FDEELLRALEQVCDDVAG-GLWSSQAVLAMEELT-
-DSQQQDLTAFLQLVGYRIQGEHPGPQDEVSNQKLFATAYFLVSALAEMPDNATVFLGTCC-KLHVISSLCCL-
--LHALSDDSVCDFHN------PTLAPLRDTERF-
GIVQRLFASADIALERMQFSAKATILKDSCIFPLILHITLSGLSTLSKEHEEELCQSGHATGQD---------------------
-------------------------------------------------------------------
>ref|NP 001006361.1| chromosome 2 organism Gallus gallus breed Leghorn gcode 1 chromosome 2 map 2 gasdermin-E Gallus gallus
----------------------------------------------------------------------------FLFHPDAVDNGMYSGAENPVPSHAS--
LSVLKKDLSQLKAQFQPFVKLPEDKQRALYKSLCELL--LHEETLTALEDVLDALCT-GDKPDLK-----ELK--
PAQQQDLADFLQLVGCRVQDELLLQNYHLQDEELLSAAHLLISAISELSDYTLVLLRACS-DLQVVPALCCL---
PYIASADGTLALSS------PSVATLTDRRRF-
DVAQRLFASANINLEKTETSVKAVTMREPRFFPLVLYVALCGLHSLGGTVEELF----------------------------------
---------------------------------------------------------------
>ref|NP 001180041.1| chromosome 4 organism Bos taurus breed Hereford gcode 1 chromosome 4 map 4 gasdermin-E Bos taurus
----------------------------------------------------------------------------------------DAGQGLPAQDKP--
LSVLKQATLLLERNFHPFMELPEQQQIALNDVLQAVL--LDEELLVVLEQVCDDIVS-SLSPSLVALG--ELK--
PSQKHNLTTFLRLVGCSVQGERLGSQDVVDNQKLFSTAFFLVSALAEMPDNAAALLGTCC-KLQIIPLLYHL---
LRALSHDGVSDLED------PALAPLKDREKF-
GIVQRLFAATDINLERMQSSVKAASREDPNVLPLILYISLSGLCALGRAQ-----------------------------------------
-----------------------------------------------------------
>ref|XP 002933445.2| location chromosome chromosome 6 completeness complete organism Xenopus tropicalis strain Nigerian gcode 1 chromosome 6 sex female tissue type liver and blood dev stage adult nots
----------------------------------------------------------------------------VLFDWDVVDGSKEAFVRKCILRSAP--
LCLLREGIAAHEKHFSAWKQLPDVQCLELYTLLCQIL--YDGQALSKLHAVVEDLCS-KRKPTQAAFD--ELM--
PSQRIIAENILYLSGYDMPNGKFLHAANR---ELLVALHILTSALNELSDSALAVLGTCC-ELQLLPVFSAL---
MNMSSDEGLCSTTE------PALMDFLDQERF-
YVSQKLFALFNIELEIKEDFIYAATAEDPGFLPLILFIVITGLQLLKRD------------------------------------------------
------------------------------------------------------

>ref|XP 001096213.2| location chromosome chromosome 3 completeness complete organism Macaca mulatta isolate AG07107 bio material Coriell:AG07107 gcode 1 chromosome 3 sex female tissue type fibroblasta
------------------------------------------------------------------------------------------------------DAAHGISSQDGP--
LSVLKQVTPLLERNFHPFVELPEPQQTALSDIFQAVL--FDDELLMVLEPVCDDLVS-GLSPPMAVLG--ELK--
LQQQQDLEAFLQLVGCSLHGGCPGPED-TGSKQLFMTAYFLVSALSEMPDNAAALLGTCC-KLQIIPTLCHL---
LRVLSDDGVSDLED------PTLAPLKDTERF-GIVQRLFASADISLERLKSSVKAVILKDSKVFP-
LLCIALNGLCALGREHS---------------------------------------------------------------------------------------------------------------
----
>ref|XP 006912063.1| chromosome Unknown completeness complete organism Pteropus alecto gcode 1 chromosome Unknown sex male tissue type kidney country Australia: Anglican Church Grammar School East Bro
------------------------------------------------------------------------------------------------------DAWQGPPASDKP--
LSVLKEVTLPLEKSFRPFAELPEQQQTALSGILQAVL--FDDELLLALEQVCDDVFS-DLSPPRVIPG--ELK--
PSQQQDLMAFLHLVGYSMQSGCPGPEDAVSNQKLFSTAYFLVSALAEMPDNAAALLGTCC-KLQIIPTLCHL-
--LHVLSADGESDLED------AALAPLRDTERF-
EIVQRLLAAADINLERQQSSVKAVTCQVPNVSPLVLYISLNGLWALGRMHQ------------------------------------------
--------------------------------------------------------------------------
>ref|XP 031796450.1| location chromosome chromosome 5 completeness complete organism Sarcophilus harrisii gcode 1 chromosome 5 cell line 91H gasdermin-E Sarcophilus harrisii
------------------------------------------------------------------------------------------------DSVDGKSEAVKLTPVEGP--
LSALKHGMLVLERNFRPFVELSGHHRAALCEILWEVL--LDDELVVALDTVLSDVLAGGRPRLWALP---GLS--
PGQQQQLAAFLGLAGLHVQEDSRVRRDARAPRELLSTAHFLVSALAEMPDDTVALMGVCC-
QLHLIPTLCHL---PQVTSADGVSELGD------PSLAPLSDAGSF-
EVVQKLFALSHISLERREAHVEAVIMKRPQYLPLVLYITLHGLCALEGKAE-----------------------------------------------
---------------------------------------------------------------------
>ref|XP 027689547.1| location chromosome chromosome 2 completeness complete organism Chelonia mydas isolate rCheMyd1 gcode 1 chromosome 2 sex male tissue type blood dev stage adult country Israel: Mes
----------------------------------------------------------------------------------------FLYQPDAVDSDRRSGDRNIVPSDAS--
LSVLKQDLLLLKMQFQPFVKLPEDKQGALNKIFYELL--HHEEIVIALEEVLDSIFT-GEEPDFTDLK--ELK--
PPQQQNITDFLQLVGYSLQNELLQKDQLHYT-GLFSAAHILISAVAELSDYTLALLRACC-DIQIVPILCYL---
PNVASADGTLTLRD------PALAPLTDTGRF-
HIVQRLFALSNINLEMMESSMKAVTMKEPSFFPFILYIALYGFYALGGTI------------------------------------------------
---------------------------------------------------------------------
>ref|XP 040208817.1| location chromosome chromosome 5 completeness complete organism Rana temporaria gcode 1 chromosome 5 gasdermin-E Rana temporaria
-----------------------------------------------------------------------------------------FDWDMVDLSRNISVDGKAVPSNYS--
LSAIKSDILEITKPFSVLQDLPEAHRGEIYNGLFEIL--HDGQTVTQLQAVVEEICL-EKRPGLTFLS--ELK--
SPERAWVQKILHLIGYDMLNQKLIEPFKK---DLLSAVHLLTSALDEMPDTALAILGACC-KLHLLPALWAL---
TNITSDEGLCSRTE------PALTDLIHQGRF-QIAQSLFSFFNMRLELNEKNIAVTS-
EDPSLLPHILYIAISGFHALKNNLKI-------------------------------------------------------------------------------------------------
------------------

>ref|XP 023962559.1| location chromosome chromosome 2 completeness complete organism Chrysemys picta bellii isolate RCT428 sub species bellii gcode 1 chromosome 2 sex female country USA: Grant Co. ...
-------------------------------------------------------------------------------FLYQPDAVDSDRHSGDRNIIPSGAS--
LSVLKQDLLLLKMQFQPFVKLPEDKQGALNKIFYEFL--HHEEMVTALEEVLDSICT-GEEPDFTDLK--ELK--
PPQQQNITDFLQLVGYSLQNELLLQNDQLHYKGLFSAAHILISAVAELSDYTLALLRACC-DIQIIPALCCL---
PNVASADGTLTLRD------PALAPLTDTGRF-
DIVQRLFALSNINLEMMESSVKAVTMKEPSFFPFILYIALYGFYALGGTI--------------------------------------------
-------------------------------------------------------------

>ref|XP 029444983.1| location chromosome chromosome 2 completeness complete organism Rhinatrema bivittatum gcode 1 chromosome 2 gasdermin-E isoform X1 Rhinatrema bivittatum
--------------------------------------------------------------------------------YSWDVVDSGDRSMMNRNVVPSDAP-
-LSLLKQDVLQQTKIFQSFLELPEEKRSVLNKLLCEIL--SHGETLTLLEDVLDDICT-EHKPNLTALN--ELK--
VEQQQNIKDLWPLLGLGLENELLLQPEKLQEKELLLATHVLVSALEEMGDSPLAILRACC-ELHLMPVLCHL---
LNSLSDDGTSMRDS--------
LLSIFNDKGRFQIAQKLFSLSNISLELTEDNIRAVTKSEPGFLPLILSIILSGFHVLAGTF-------------------------------
--------------------------------------------------------------------------

>ref|XP 004626528.1| chromosome Unknown completeness complete organism Octodon degus isolate 3935 gcode 1 chromosome Unknown sex female gasdermin-E Octodon degus
--------------------------------------------------------------------------------------------DAKHGMLSKDGL--
LSVLKQATLPLERNFHPFVELPAQQQQALHNLLQAVL--LDEELLVVLEQVCDDMAG-
SVWSPQTALAMRALK--
APQQQDLTAFLQLVGCSIQGECPSPEDEVSNQKLFATAYFLVSALAEMPDNAAALLGICC-KLQIIPTLYHM---
LRALSDDGVCDLED------PTLAPLRDTERF-
GIVQRLFVATDINLEQEQASVKAVILKDPHIFPLILYITLNGLCALGREH---------------------------------------------
------------------------------------------------------------

>ref|XP 021046698.1| location chromosome chromosome 2 completeness complete organism Mus pahari gcode 1 chromosome 2 sex female tissue type tail gasdermin-E Mus pahari
--------------------------------------------------------------------------------------------DGGQGISSQDGP--
LRVVKQATLHLERSFHPFAVLPAQQQRALFCVLQKIL--FDEELLRALEQVCDDVAG-GLWSSQAISAMEELT--
DSQQQDLTAFLQLVGYRIQGEHPGPQDAVSNQKLFATAYFLVSALAEMPDNATVFLGTCC-KLHVISSLCCL--
-LHALSDDSVCDFQD------PALAPLRDTERF-
GIVQRLFASADIALERMQFSAKATILKDSCIFPLILHITLSGLSTLSKEHEEQLCQSEGHATGQD-------------------
-----------------------------------------------------------------------

>ref|XP 009324004.1| chromosome Unknown completeness complete organism Pygoscelis adeliae isolate BGI AS28 gcode 1 sex male country Antarctica chromosome Unknown PREDICTED: non-syndromic hearing impae
-------------------------------------------------------------------------------FLYQPDAVDNEMYSGAKNRIPSDAS--
LSVLKQDLSWLKTQFQPFVKLPEDKQRALYKTLCELL--LHEEMVAALEDLLDDICT-GDKPDLK-----ELK--
PAQQQDLLGFLQLLGCSLQSELLLQKYQPQDEELLSAAHLLISAISELPDYTLVLLRACC-DLQVVPALCCL---
PNIASADGTLALSS------PLVATLTDRGRF-
DVVQRLFASSNITLEMTVSSVKAVTMKEPRFFPLVLYVALYGFYALGGNV-----------------------------------------
-----------------------------------------------------------------

>ref|XP 024418892.1| chromosome Unknown completeness complete organism Desmodus rotundus isolate DRU21DN04 gcode 1 chromosome Unknown sex male tissue type muscle/skin sample dev stage adult country Us
---------------------------------------------------------------------------------------DAGQGPSAPDEP--
LSVLKQATLPLERAFHPFGQLPEQEQTALSGVLQVVL--FDEELLVVLEQVCDDMVG-GLPPSLAVLG--ELK--
PPQKQNLMAFLQLVGCSMQGECLGPQGVVSNQKLFSTAYFLVSALAEMPDNAAALLGTCC-
KLQIIPALCHL---LRALSAGGEPDLED------TALAPLRDTERF-
GIVQCLLATASLDLERRQSSVKAVTCQVPNVSPLVLYVSLSGLHALGRAHQ-------------------------------------
-------------------------------------------------------------
>ref|XP 016015046.2| chromosome Unknown completeness complete organism Rousettus aegyptiacus isolate mRouAeg1 gcode 1 chromosome Unknown sex male tissue type muscle dev stage adult country USA: Berkes
---------------------------------------------------------------------------------------DAWQRPSASDKP--
LSVLKQVSLPLERSFRPFAELPQQQQTALSGILQAVL--FDEELLLALEQVCDDVFS-GLSPPRAMPG--ELK--
PSQQQDLAAFLRLVGYSVQSGCPGPEDAVSNQKLFSTAYFLVSALAEMPDNAAALLGTCC-KLQMIPTLCHL-
--LRALSADGESDLED------AALAPLRDTERF-
EIVQRLLAAADIRLERRQSSVKAVTYPVPDVSQLVLYVSLNGLWALARMHQ-----------------------------------
---------------------------------------------------------------
>ref|XP 032903802.1| location chromosome chromosome 2 completeness complete organism Amblyraja radiata isolate CabotCenter1 gcode 1 chromosome 2 sex male tissue type testis liver country USA: Gulf ofa
---------------------------------------------------------------------------PLSCVDGAHKCQLSALEMDVPSETS--
LIILKPAIEEANQQFQPFHELAEEKCLQLFRLYCEFL--YHEEVISFLENALDELSS-TEQPNLLGLQ--ELD--
STQAQRVKELLQILGYSCPNEQGSLKKDITTPEILTAAFYLVSALAGMSEESLAVLGICC-ETQILPTLHYL---
INNVPDDGIVPVDN------PNMLPLQEEDNF-
YIAHRLFALSNICLEVRETAIKVITDDQPGMAPVLLCIVLQGFAILSGIKRNTSDPKTQT--------------------------
---------------------------------------------------------------
>ref|XP 001159685.3| location chromosome chromosome 7 completeness complete organism Pan troglodytes isolate Yerkes chimp pedigree *C0471 (Clint) gcode 1 chromosome 7 sex male tissue type blood dev ss
---------------------------------------------------------------------------------------DAAHGISSQDGP--
LSVLKQATLLLERNFHPFAELPEPQQTALSDIFQAVL--FDDELLMVLEPVCDDLVS-GLSPTVAVLG--ELK--
PRQQQDLVAFLQLVGCSLQGGCPGPED-AGSKQLFMTAYFLVSALAEMPDNAAALLGTCC-KLQIIPTLCHL--
-LRALSDDGVSDLED------PILTPLKDAERF-
GIVQRLFASADISLERLKSSVKAVILKDSKVFPLLLCITLNGLCALGREHS------------------------------------------
--------------------------------------------------------

PYD Alignment
>sp|Q8WXC3|PYDC1 HUMAN Pyrin domain-containing protein 1 OS=Homo sapiens OX=9606 GN=PYDC1 PE=1 SV=1
-------------------MGTKREAIL-KVLENLTPEELKKFKMKLGTVPLREGFE----RIPRGALGQLDIV-DLTDKLVAS-
-YYEDYAAELVVAVLRDMR-----------MLEEAAR-LQRAA-----------
>NP 660183.1:1-88 apoptosis-associated speck-like protein containing a CARD isoform b Homo sapiens

--------------------MGRARDAIL-DALENLTAEELKKFKLKLLSVPLREGYG----RIPRGALLSMDAL-DLTDKLVSF--YLETYGAE-LTANVLRDM-----------GLQEMAG-QLQAA-----------
>NP 001104231.1 uncharacterized protein LOC796649 Danio rerio
--------------------MASTKTEIL-DVLDELRERE---FKD---FKWRLSNKETLENSIPRGKLENADRQ-DVVDYMEQH--FGTSEAGK-VAVRL--LH-----------SMNQNNL-AEQLKKTLAVVSKV--
>q919n6 danio asc
-------------------MAESFKEHLQ-EAFEDLGADN---LRK---FKSKLGDRRQEP-RVTKSAIEKLKDEIDLADLMVGV--FTSKDAVS-VTVEI--LR-----------AIKCNAV-ADDLLRNTGQS-----
>q56p42 pydc2 human
-----------------MASSAELDFNLQ-ALLEQLSQDELSKFKS---LIRTISLGK----ELQTVPQTEVDKA-N-GKQLVEI--FTSHSCS--YWAGMAAIQ-----------VFEKMNQ-THLSGRADEHCVM---
>q91vj1 pycard mouse
-------------------MESEYREMLLLTGLDHITEEE---LKR---FKYFALTEF----QIARSTLDVADRT-ELADHLIQS--AGAASAVTKAINIFQKLN-----------YMHIANA-LEEKKKE---------
>o35368 ifi203 mouse
--------------------MAEYKNIVLLKGLENMEDYQ---FRT---VKSLLRKEL----KLTKKMQEDYDRI-QLADWMEDK--FPK---DA-GLDKL--IK-----------VCEHIKD-LKDLAKKLKTEKAK--
>q504j1 caspb danio
---------------------------LS-DVLEDLVESELKQFTR---QLWIGVKPGVE--PIPRGKLENKDRQ-DVVDSMVQQ--YSED-AGTITVQTL--RK-----------IKQNERA-----------------
>q919l7 caspa danio
--------------------------HLQ-DALSNIGADN---LRR---FQSRLGDRKQEP-RVRKSTIEKLKDEIDLVDLLVNT--FTSD-AVS-VTVDI--LR-----------GIKCNAV-AE--------------
>tr|B2CWB9|B2CWB9 9POXV M013 OS=Myxoma virus OX=10273 GN=m013L PE=4 SV=1
---------------------MEHRGVII-TVLENLSDYQ---FKM---FIYLAMEDL----YIERAEKEKIDRI-DLAHKISEQ--YLGTDYIE-FMKRV--TD-----------FIPNKVY-VDSLLARAEADA----
>NP 051902.1 Hypothetical protein SFV s013L Rabbit fibroma virus
---------------------MEHRGVII-TVLENLTDYQ---FKM---FLYLVTEDL----RINPVEKEKIDRI-DLAYKISEL--YPGHSYIE-FMKQV--TG-----------YIPNKVY-VDSLLKNAEENT----
>YP 227401.1 hypothetical protein DpV83gp024 Deerpox virus W-848-83
---------------------MELRSAII-AVLENINRYQ---FKM---VIFIVQDEL----YIEEEEKLTMDRI-DLAEKLIKK--YKD---IR-SLYFL--LN-----------IFKEMPD-TEFVKKQLSDYIKRF-
>NP 073403.1 18L protein Yaba-like disease virus
-----------------MLYTMRIKSAII-FSLEDVTHYQ---FKI---LIFLTKDEL----NISDEEKQILDRV-DFAEKLFQT--YPG---IK-SLYFL--EK-----------AISMVPN-AKYARSNINRLISDL-
>ABQ43490.1 pyrin domain Tanapox virus
-----------------MLYNMRIKSAII-FSLEDVTHYQ---FKI---LIFLTKDEL----NINDEEKQILDRV-DFAEKLFQA--YPG---IK-SLYFL--EK-----------AISMVPN-AKYARSNINRLISDL-
>NP 570174.1 Hypothetical protein SWPVgp014 Swinepox virus
---------------------MELRYTII-SVLERLTPYQ---FKT---LLFLIQDDI----NISNDDINVLDRV-DLAIKIMNK--YNN---YR-AIYFL--YK-----------VILRIHN-TEYISGTLQRSIQNI-
>Vaccinia
-----MVKNNKISNSCRMIMSTNPNNILM-RHLKNLTDDE---FKC---IIHRSSDFL----YLSDSDYTSITKE-TLVSEIVEE--YPDD-CNK-ILAII--FL-----------VLDKDID-VDIET----------

>Buffalopox
-----MVKNNKISNSCLMIMSTNPNNILM-RHLKNLTDDE---FKC---IIHRSSDFL----YLSDSDYTSITKE-TLVSEIVEE--YPDD-CNK-ILAII--FL-----------VLDKDID-VDIET-----------
>Cowpox
MVKSNKIQKNKISNSCRMIMSTNPNNILM-RHLKNLTDDE---FKC---IIHRSSDFL----YLSDRDYTSITKE-TLVSEIVEE--YPDD-CNK-ILAII--FL-----------VLDKDID-VDIET-----------
>Rabbitpox
-----MVKNNKISNSCLMIMSTDPNNILM-RHLKNLTDDE---FKC---IIHRSSDFL----YLSDRDYTSITKE-TLVSEIVEE--YPDD-CNK-ILAII--FL-----------VLDKDID-VDIET-----------
>Horsepox
------MVKNKISNSCLMIMSTDPNNILM-RHLKNLTDDE---FKC---IIHRSSDFL----YLSNRDYTSITKE-TLVSEIVEE--YPDD-CNK-ILAII--FL-----------VLDKDID-VDIKT-----------
>Orthopox abatino
MVKNNKIQKIKTSNSYRDVIMTNPNNILM-RHLKNLTDDE---FKC---IVNRSSDFI----YLSDRDYTSITKE-TLVSEIVEE--YPDD-CNK-ILAII--FL-----------VLGKDVD-IETET-----------
>Akhmeta virus
MVKSNKIQKNKISNSCRYMIMNTNTNILM-RHLKNLTDDE---FKC---IIHRSSDFL----YLSDRDYTSITKE-TLVSEIVEE--YQDD-CNK-ILAII--FL-----------VLDKDID-VDIET-----------
>Variola Virus
-----------------MIMSTDPNNILM-RYLKNLTDDK---FKC---IIHQSSDFL----YLSDRDYTSITKE-TLVSEIVEE--YPDD-CNK-ILAII--FL-----------VLDKDID-VDIET-----------
>Monkeypox
-----------------MIMRTDTNNILM-RHLKNLTDDE---FKC---IIHRSSDFL----YLSDRDYTSITKE-TLVSEIVEE--YPDD-CNK-ILAII--FL-----------VLDKDID-VDIET-----------
>Alaskapox
-----------------------MNTILM-RHFKNLTDDE---FKC---IIHRSSDFI----YLSDRDYTSITKE-TLVSEIVEE--YPDD-CDK-ILAII--FL-----------VLGKDIE-VDVKT-----------
>Volepox
---------------------MANKNRLV-SYLKNLTDDE---FKC---IIYRSSDFI----YLSDSDHDTITKE-KLVKEIIEE--YPDD-CNK-ILAII--FL-----------VLDKDID-IDIET-----------
>Skunkpox
----------------------MNKNRLV-KYLKNLTDDE---FKC---IIYRSSDFI----YLSDSDHDNITRE-TLVKEIMEE--YPDD-CNK-ILAII--FL-----------VLDKDID-LDIET-----------
>Raccoonpox
---------------------MDVKNRIV-KYLKNLTDDE---FRY---IIYRSSDFI----YLSDGDHNTITKE-TLVNEIIEE--YPDD-CNK-IFAII--FL-----------VLDIDID-LDTET-----------
>NY 014 poxvirus
------------------------MDKIM-CYLNNITDEE---FET---VIDRSSNFL----YLSDDDHTNITRE-SLALEIVEE--YPND-CER-ILAII--LL-----------SLDKYIS-FESTI-----------
>Murmansk poxvirus
------------------------MDKIT-HYLNNITDGE---FET---VIDRSSNFL----YLSDDDHTNITRE-SLALEIVEE--YPND-CER-ILAIL--LL-----------SLDKYIS-FESTI-----------
>Yoka poxvirus

-----------------------MNKIL-NYFNSITDDE---FKT---IISISSDFL----YLSDNDNINITKE-SLALEILEQYPYPND-
YEK-ILSML--LL-----------ALGKYVF-FETII-----------
>QDJ94995.1 IFN-inducible protein Hypsugopox virus
------------------MDSYTKGSIL-YALKNLTFQQ---LKE---FMFISSEQL----YLSNKDRLNFDVV-DLADRIIIV--
CHDN-ALK-AFKLI--KE-----------ILTMMES-VDFVINYMNKCIENTC
>QHG62576.1 hypothetical protein Cetacean poxvirus 1
-----------MIDSKSLNVEIEVKIALM-RALDELSSYQ---LKV---LAFLTSNRV----CVIHRSMSNVDRV-
DIAKRINTT--LP---CVC-SLTYLRDLLKNVKHTESVRDILCEDINYIESIMRNNNEIKRFTC
>YP 009407969.1 IFN-inducible protein Eptesipox virus
------------------MDAFTKGSTL-YALKNLTFNQ---LKE---FMFISSDGL----YLTHKNRLEFDVV-DLADRIIIV--
NNNN-PLK-VFKHI--KE-----------TLCLMDD-VEFVLKYIDKCIENCA

GSDM Tree

(((((((((((((('AY484669.1:c159516-158782 Rabbitpox virus complete genome':0.00000632,'KY349117.1:c166272-165538 Horsepox virus strain MNR complete genome':0.00396044)0.5300:0.00000632,'M35027.1:c153690-152956 Vaccinia virus Copenhagen complete genome':0.00396094)0.6900:0.00396040,('MG599038.1:c157906-157148 Buffalopox virus isolate Karachi 2005 complete genome':0.00000632,'AY243312.1:c154675-153917 Vaccinia virus WR complete genome':0.00384602)0.7600:0.00398762)0.8500:0.01196656,'KC813500.1:c153312-152578 Cowpox virus strain MonKre08/4 complete genome':0.00000632)0.2000:0.00001264,'MN912466.1:c156293-155559 Ectromelia virus WH complete genome':0.01989595)0.6200:0.00395345,'NC 055231.1:c163765-163031 Orthopoxvirus Abatino complete genome':0.00793905)0.5300:0.00641072,('MH607143.1:c167386-166652 Akhmeta virus isolate Vani 2010 complete genome':0.02387728,((('AY009089.1:c154402-154034 Camelpox virus CMS complete genome':0.00000632,'MK910851.1:c154204-153836 Camelpox virus strain Negev2016 complete genome':0.00808054)0.9200:0.02440740,'DQ437594.1:c153307-152573 Taterapox virus strain Dahomey 1968 complete genome':0.01191498)0.3300:0.00000632,('KY358055.1:c145385-144651 Variola virus strain VD21 17th century complete genome':0.00000632,('DQ441432.1:c146026-145292 Variola virus strain Korea 1947 (Lee Masterseed) complete genome':0.00395560,'DQ441437.1:c146904-146170 Variola virus strain Sierra Leone 1969 (V68-258) complete genome':0.00000632)0.7700:0.00395750)0.8700:0.00793967)0.6500:0.01266543)0.3200:0.00554384)0.5300:0.03253804,'MN240300.1:c166555-165827 Alaskapox virus complete genome':0.11139866)0.9700:0.13966158,('KU749309.1:c145369-144644 Raccoonpox virus strain 85A complete genome':0.10647955,('KU749310.1:c167631-166873 Skunkpox virus strain WA complete genome':0.07403330,'KU749311.1:c169057-168281 Volepox virus strain CA complete genome':0.09493255)0.5900:0.02370722)0.4600:0.07236375)1.0000:0.50096918,('HQ849551.1:c134784-134035 Yoka poxvirus strain DakArB 4268 complete genome':0.41854758,('MF001305.1:c159640-158771 NY 014 poxvirus strain 2013 complete genome':0.10644996,'MF001304.1:c160603-159728 Murmansk poxvirus strain LEIV-11411

complete
genome':0.06453801)0.8000:0.16035801)0.4100:0.07538004)1.0000:1.10608085,YP_00940809
7.1_Immunoprevalent_protein_Eptesipox_virus:0.88065310)0.5900:0.32061200,((NP_051846.
1_m132L_Myxoma_virus:0.09720761,NP_052021.1_gp132L_Rabbit_fibroma_virus:0.24063256
)1.0000:1.72365406,(('MF001304.1:c165763-165224 Murmansk poxvirus strain LEIV-11411
complete genome':0.04040716,'MF001305.1:c164808-164269 NY 014 poxvirus strain 2013
complete genome':0.11589372)0.8900:0.43947697,('KU749311.1:173861-174445 Volepox virus
strain CA complete genome':0.03751514,('KU749310.1:172437-172985 Skunkpox virus strain
WA complete genome':0.06828167,'KU749309.1:150174-150722 Raccoonpox virus strain 85A
complete
genome':0.09772203)0.6700:0.06086908)0.9400:0.42301996)1.0000:1.32178979)0.4300:0.417
93913)0.6600:0.74188710,(((((((ref|NP_113603.1|_chromosome_8_organism_Homo_sapiens
_gcode_1_chromosome_8_map_8q24.21_gasdermin-C_Homo_sapiens:0.00648950,'ref|XP
001153860.1| location chromosome chromosome 8 completeness complete organism Pan
troglodytes isolate Yerkes chimp pedigree *C0471 (Clint) gcode 1 chromosome 8 sex male
tissue type blood dev ss':0.01016408)0.7300:0.02356312,'ref|XP 028708741.1| location
chromosome chromosome 8 completeness complete organism Macaca mulatta isolate
AG07107 bio material Coriell:AG07107 gcode 1 chromosome 8 sex female tissue type
fibroblasta':0.05208806)1.0000:0.26393153,ref|XP_019678683.1|_location_chromosome_chro
mosome_F2_completeness_complete_organism_Felis_catus_isolate_Cinnamon_breed_Abyssi
nian_gcode_1_chromosome_F2_sex_female_gasdermin-
C_Felis_catus:0.41823708)0.1300:0.05011136,('ref|XP 036088919.1| chromosome Unknown
completeness complete organism Rousettus aegyptiacus isolate mRouAeg1 gcode 1
chromosome Unknown sex male tissue type muscle dev stage adult country USA:
Berkes':0.32082568,ref|XP_024427836.1|_chromosome_Unknown_completeness_complete_o
rganism_Desmodus_rotundus_isolate_DRU21DN04_gcode_1_chromosome_Unknown_sex_ma
le_tissue_type_muscle/skin_sample_dev_stage_adult_country_Us:0.13881323)0.6000:0.09111
937)0.1400:0.07902331,ref|NP_001039469.1|_chromosome_14_organism_Bos_taurus_breed
_L1_Hereford_gcode_1_chromosome_14_map_14_gasdermin-
C_Bos_taurus:0.55125872)0.3600:0.07430343,ref|XP_021073018.1|_location_chromosome_c
hromosome_17_completeness_complete_organism_Mus_pahari_gcode_1_chromosome_17_s
ex_female_tissue_type_tail_gasdermin-
C_Mus_pahari:0.36366499)0.9800:0.59858814,ref|XP_031803207.1|_location_chromosome_c
hromosome_1_completeness_complete_organism_Sarcophilus_harrisii_gcode_1_chromosome
_1_cell_line_91H_gasdermin-C_Sarcophilus_harrisii:0.73800521)0.4200:0.14672251,((('ref|XP
015154998.2| location chromosome chromosome 27 completeness complete organism Gallus
gallus isolate bGalGal1 gcode 1 chromosome 27 sex female tissue type blood country USA:
Fayetteville lat
ls':0.89302167,ref|XP_029428328.1|_location_chromosome_chromosome_12_completeness_
complete_organism_Rhinatrema_bivittatum_gcode_1_chromosome_12_gasdermin-
A_Rhinatrema_bivittatum:1.13451927)0.3600:0.10801623,XP_032891479.1_gasdermin-
A_Amblyraja_radiata:1.35621246)0.1500:0.21106408,(ref|NP_081236.1|_chromosome_15_or
ganism_Mus_musculus_strain_C57BL/6_gcode_1_chromosome_15_map_15_gasdermin-
D_Mus_musculus:0.37973081,(((ref|NP_079012.3|_chromosome_8_organism_Homo_sapiens

_gcode_1_chromosome_8_map_8q24.3_gasdermin-D_Homo_sapiens:0.00490946,'ref|XP 009454389.2| location chromosome chromosome 8 completeness complete organism Pan troglodytes isolate Yerkes chimp pedigree *C0471 (Clint) gcode 1 chromosome 8 sex male tissue type blood dev ss':0.03154543)0.9800:0.02755997,'ref|XP 015001615.2| location chromosome chromosome 8 completeness complete organism Macaca mulatta isolate AG07107 bio material Coriell:AG07107 gcode 1 chromosome 8 sex female tissue type fibroblasta':0.02541537)1.0000:0.24655309,(ref|XP_023104116.1|_location_chromosome_chromosome_F2_completeness_complete_organism_Felis_catus_isolate_Cinnamon_breed_Abyssinian_gcode_1_chromosome_F2_sex_female_gasdermin-D_Felis_catus:0.38688835,(ref|NP_001039625.1|_chromosome_14_organism_Bos_taurus_breed_Hereford_gcode_1_chromosome_14_map_14_gasdermin-D_Bos_taurus:0.18906395,(ref|XP_024428237.1|_chromosome_Unknown_completeness_complete_organism_Desmodus_rotundus_isolate_DRU21DN04_gcode_1_chromosome_Unknown_sex_male_tissue_type_muscle/skin_sample_dev_stage_adult_country_Us:0.22927157,('ref|XP 006913014.1| chromosome Unknown completeness complete organism Pteropus alecto gcode 1 chromosome Unknown sex male tissue type kidney country Australia: Anglican Church Grammar School East Bro':0.05855083,'ref|XP 036089037.1| chromosome Unknown completeness complete organism Rousettus aegyptiacus isolate mRouAeg1 gcode 1 chromosome Unknown sex male tissue type muscle dev stage adult country USA: Berkes':0.07916446)0.9600:0.08309248)0.6600:0.09456772)0.6500:0.07865023)0.8200:0.11484867)0.5700:0.05693998)1.0000:0.51206201)0.1000:0.19500033)0.0200:0.03306529)0.0000:0.15081324,((sp|Q5Y4Y6|GSDA3_MOUSE_Gasdermin-A3_OS_Mus_musculus_OX_10090_GN_Gsdma3_PE_1_SV_1:0.02992407,sp|Q32M21|GSDA2_MOUSE_Gasdermin-A2_OS_Mus_musculus_OX_10090_GN_Gsdma2_PE_2_SV_1:0.08079961)0.5000:0.01350323,(sp|Q9EST1|GSDMA_MOUSE_Gasdermin-A_OS_Mus_musculus_OX_10090_GN_Gsdma_PE_2_SV_1:0.01547699,((('ref|XP 016020091.2| chromosome Unknown completeness complete organism Rousettus aegyptiacus isolate mRouAeg1 gcode 1 chromosome Unknown sex male tissue type muscle dev stage adult country USA: Berkes':0.00923641,'ref|XP 006924750.1| chromosome Unknown completeness complete organism Pteropus alecto gcode 1 chromosome Unknown sex male tissue type kidney country Australia: Anglican Church Grammar School East Bro':0.01784865)0.9500:0.03854298,ref|XP_024429158.1|_chromosome_Unknown_completeness_complete_organism_Desmodus_rotundus_isolate_DRU21DN04_gcode_1_chromosome_Unknown_sex_male_tissue_type_muscle/skin_sample_dev_stage_adult_country_Us:0.07730313)0.4600:0.00685917,(ref|XP_035935879.1|_chromosome_Unknown_completeness_complete_organism_Halichoerus_grypus_isolate_241Hg_ecotype_Northwest_Atlantic_gcode_1_chromosome_Unknown_sex_male_tissue_type_blood_dev_stage_molts:0.09006814,((ref|XP_024836266.1|_location_chromosome_chromosome_19_completeness_complete_organism_Bos_taurus_isolate_L1_Dominette_01449_registration_number_42190680_breed_Hereford_gcode_1_chromosome_19_sex_fem...:0.07277097,ref|XP_004633937.1|_chromosome_Unknown_completeness_complete_organism_Octodon_degus_isolate_3935_gcode_1_chromosome_Unknown_sex_female_gasdermin-A_Octodon_degus:0.08986357)0.0600:0.00538989,('ref|XP 014975037.1| location chromosome chromosome 16 completeness complete organism Macaca

mulatta isolate AG07107 bio material Coriell:AG07107 gcode 1 chromosome 16 sex female tissue type
fibroblaa':0.03818418,(ref|NP_835465.2|_chromosome_17_organism_Homo_sapiens_gcode_1_chromosome_17_map_17q21.1_gasdermin-A_Homo_sapiens:0.00536674,'ref|XP 001171222.1| location chromosome chromosome 17 completeness complete organism Pan troglodytes isolate Yerkes chimp pedigree *C0471 (Clint) gcode 1 chromosome 17 sex male tissue type blood
devs':0.00000632)0.5100:0.01070020)0.9800:0.04354841)0.1500:0.00623461)0.1800:0.00553476)0.8000:0.06299807)0.7900:0.04239040)0.9900:0.44258489,((((((('ref|XP 024897515.1| chromosome Unknown completeness complete organism Pteropus alecto gcode 1 chromosome Unknown sex male tissue type kidney country Australia: Anglican Church Grammar School East Bro':0.06586777,'ref|XP 036078993.1| chromosome Unknown completeness complete organism Rousettus aegyptiacus isolate mRouAeg1 gcode 1 chromosome Unknown sex male tissue type muscle dev stage adult country USA:
Berkes':0.09991046)1.0000:0.11539316,ref|XP_024429114.1|_chromosome_Unknown_completeness_complete_organism_Desmodus_rotundus_isolate_DRU21DN04_gcode_1_chromosome_Unknown_sex_male_tissue_type_muscle/skin_sample_dev_stage_adult_country_Us:0.21763352)0.4000:0.02806263,ref|XP_035935913.1|_chromosome_Unknown_completeness_complete_organism_Halichoerus_grypus_isolate_241Hg_ecotype_Northwest_Atlantic_gcode_1_chromosome_Unknown_sex_male_tissue_type_blood_dev_stage_molts:0.34581838)0.2500:0.03004080,ref|NP_001231146.1|_chromosome_19_organism_Bos_taurus_gcode_1_chromosome_19_map_19_gasdermin-B_Bos_taurus:0.26530564)0.4800:0.09159899,('ref|XP 014975035.1| location chromosome chromosome 16 completeness complete organism Macaca mulatta isolate AG07107 bio material Coriell:AG07107 gcode 1 chromosome 16 sex female tissue type
fibroblaa':0.11006455,(ref|NP_001159430.1|_chromosome_17_organism_Homo_sapiens_gcode_1_chromosome_17_map_17q21.1_gasdermin-
B_isoform_3_Homo_sapiens:0.01869987,'ref|XP 009430521.1| location chromosome chromosome 17 completeness complete organism Pan troglodytes isolate Yerkes chimp pedigree *C0471 (Clint) gcode 1 chromosome 17 sex male tissue type blood
devs':0.01689996)0.9400:0.05518121)1.0000:0.07184298)1.0000:0.50473782,ref|XP_023358160.1|_location_chromosome_chromosome_4_completeness_complete_organism_Sarcophilus_harrisii_gcode_1_chromosome_4_cell_line_91H_gasdermin-
B_Sarcophilus_harrisii:0.56088280)0.9600:0.33170458,('ref|XP 032903802.1| location chromosome chromosome 2 completeness complete organism Amblyraja radiata isolate CabotCenter1 gcode 1 chromosome 2 sex male tissue type testis liver country USA: Gulf
ofa':0.57159834,(((ref|XP_002933445.2|_location_chromosome_chromosome_6_completeness_complete_organism_Xenopus_tropicalis_strain_Nigerian_gcode_1_chromosome_6_sex_female_tissue_type_liver_and_blood_dev_stage_adult_nots:0.58041056,ref|XP_040208817.1|_location_chromosome_chromosome_5_completeness_complete_organism_Rana_temporaria_gcode_1_chromosome_5_gasdermin-
E_Rana_temporaria:0.43530056)0.7400:0.37086991,ref|XP_029444983.1|_location_chromosome_chromosome_2_completeness_complete_organism_Rhinatrema_bivittatum_gcode_1_chromosome_2_gasdermin-
E_isoform_X1_Rhinatrema_bivittatum:0.30113784)0.5100:0.13675007,(((ref|NP_001006361.1

|_chromosome_2_organism_Gallus_gallus_breed_Leghorn_gcode_1_chromosome_2_map_2_gasdermin-E_Gallus_gallus:0.14607108,'ref|XP 009324004.1| chromosome Unknown completeness complete organism Pygoscelis adeliae isolate BGI AS28 gcode 1 sex male country Antarctica chromosome Unknown PREDICTED: non-syndromic hearing impae':0.06555493)1.0000:0.18747693,('ref|XP 027689547.1| location chromosome chromosome 2 completeness complete organism Chelonia mydas isolate rCheMyd1 gcode 1 chromosome 2 sex male tissue type blood dev stage adult country Israel: Mes':0.07982534,'ref|XP 023962559.1| location chromosome chromosome 2 completeness complete organism Chrysemys picta bellii isolate RCT428 sub species bellii gcode 1 chromosome 2 sex female country USA: Grant Co. ...':0.01125944)0.9900:0.09327687)0.9600:0.15059215,(ref|XP_031796450.1|_location_chromosome_chromosome_5_completeness_complete_organism_Sarcophilus_harrisii_gcode_1_chromosome_5_cell_line_91H_gasdermin-E_Sarcophilus_harrisii:0.49835971,(((('ref|XP 006912063.1| chromosome Unknown completeness complete organism Pteropus alecto gcode 1 chromosome Unknown sex male tissue type kidney country Australia: Anglican Church Grammar School East Bro':0.04540917,'ref|XP 016015046.2| chromosome Unknown completeness complete organism Rousettus aegyptiacus isolate mRouAeg1 gcode 1 chromosome Unknown sex male tissue type muscle dev stage adult country USA: Berkes':0.06765941)1.0000:0.10802648,ref|XP_024418892.1|_chromosome_Unknown_completeness_complete_organism_Desmodus_rotundus_isolate_DRU21DN04_gcode_1_chromosome_Unknown_sex_male_tissue_type_muscle/skin_sample_dev_stage_adult_country_Us:0.12144140)0.9300:0.07802127,ref|NP_001180041.1|_chromosome_4_organism_Bos_taurus_breed_Hereford_gcode_1_chromosome_4_map_4_gasdermin-E_Bos_taurus:0.16311047)0.7200:0.04030548,(((ref|NP_001120925.1|_chromosome_7_organism_Homo_sapiens_gcode_1_chromosome_7_map_7p15.3_gasdermin-E_isoform_a_Homo_sapiens:0.00453124,'ref|XP 001159685.3| location chromosome chromosome 7 completeness complete organism Pan troglodytes isolate Yerkes chimp pedigree *C0471 (Clint) gcode 1 chromosome 7 sex male tissue type blood dev ss':0.00911979)0.9900:0.01107332,'ref|XP 001096213.2| location chromosome chromosome 3 completeness complete organism Macaca mulatta isolate AG07107 bio material Coriell:AG07107 gcode 1 chromosome 3 sex female tissue type fibroblasta':0.05555503)1.0000:0.14905206,(ref|XP_004626528.1|_chromosome_Unknown_completeness_complete_organism_Octodon_degus_isolate_3935_gcode_1_chromosome_Unknown_sex_female_gasdermin-E_Octodon_degus:0.14112392,(ref|NP_061239.1|_chromosome_6_organism_Mus_musculus_strain_C57BL/6_gcode_1_chromosome_6_map_6_gasdermin-E_precursor_Mus_musculus:0.01643603,ref|XP_021046698.1|_location_chromosome_chromosome_2_completeness_complete_organism_Mus_pahari_gcode_1_chromosome_2_sex_female_tissue_type_tail_gasdermin-E_Mus_pahari:0.04506946)1.0000:0.24424411)0.8800:0.06822000)0.6100:0.02349932)1.0000:0.29867710)1.0000:0.32476703)0.9400:0.23585091)0.6400:0.34461861)1.0000:1.77119903)0.0100:0.08220955);

PYD Tree

(sp_Q8WXC3_PYDC1_HUMAN:0.0000021986,NP_660183.1_1-
88:0.8189957255,((((NP_001104231.1:0.5582152076,(q919n6_danio:0.1596274241,q919l7_cas
pa:0.1464068014)100:1.6604787690)78:0.3649286491,q504j1_caspb:0.8779213839)75:0.3248
494404,q91vj1_pycard:1.3275505794)53:0.3488003785,(q56p42_pydc2:2.4248812488,((((o353
68_ifi203:1.9252008714,(tr_B2CWB9_B2CWB9_9POXV:0.1873719086,NP_051902.1:0.1151924
580)98:0.8588394249)30:0.1685723872,QHG62576.1_hypothetical:1.3381578798)25:0.020581
3423,((YP_227401.1:0.4922697513,((NP_073403.1:0.0000024848,ABQ43490.1_pyrin:0.032263
9322)100:0.5536693021,NP_570174.1:0.6631629338)89:0.1050133382)91:0.2980680604,(QDJ
94995.1_IFN-
inducible:0.0322861746,YP_009407969.1:0.3451439803)100:1.1424169120)56:0.1652826846)
38:0.2909839033,((((((Vaccinia:0.0000024478,(Cowpox:0.0000024085,((Orthopox_abatino:0.10
55672911,Alaskapox:0.0953186299)96:0.0427250561,Akhmeta_virus:0.0648530371)92:0.0274
133695)92:0.0511067176)53:0.0098506539,Buffalopox:0.0000023745)51:0.0083698252,(Rabbi
tpox:0.0000027502,((Horsepox:0.0531890168,Variola_Virus:0.0358970206)28:0.0000020161,
Monkeypox:0.0242749897)26:0.0000027502)32:0.0115162760)97:0.2153206803,(Volepox:0.0
403124363,Skunkpox:0.0547150846)81:0.0431585044)36:0.0492731175,Raccoonpox:0.084855
2751)84:0.2158394416,((NY_014:0.0278797725,Murmansk_poxvirus:0.0311930000)97:0.1567
232511,Yoka_poxvirus:0.3046051547)97:0.3533709024)96:0.7863255573)26:0.1931065783)79
:0.4863747229)100:0.9712551055);

```
#!/bin/bash

#checks to ensure that matching files are present in both folders

for tmfile in TM/*; do
	name=${tmfile##*/}

	if [ -f "FATCAT/$name" ]; then
		echo "$name"
	else
		echo "$name is MISSING in FATCAT folder"
	fi
done

for fatfile in FATCAT/*; do
	name=${fatfile##*/}

	if [ -f "TM/$name" ]; then
		echo "$name"
	else
		echo "$name is MISSING in TM folder"
	fi
done
```

---

AlphaFold parameters

```
# explicit run of run_alphafold pointing to the appropriate databases on the RAM disk, for file passed in as $1
singularity exec --nv /uufs/chpc.utah.edu/sys/installdir/alphafold/2.1.2/alphafold_2.1.2.sif /app/run_alphafold.sh --data_dir=/scratch/general/lustre/app-repo/alphafold --uniref90_database_path=/scratch/general/lustre/app-repo/alphafold/uniref90/uniref90.fasta --mgnify_database_path=/scratch/general/lustre/app-repo/alphafold/mgnify/mgy_clusters_2018_12.fa --uniclust30_database_path=/dev/shm/af/uniclust30_2018_08 --bfd_database_path=/dev/shm/af/bfd_metaclust_clu_complete_id30_c90_final_seq.sorted_opt --pdb70_database_path=/dev/shm/af/pdb70 --template_mmcif_dir=/scratch/general/nfs1/app-repo/alphafold/pdb_mmcif/mmcif_files --obsolete_pdbs_path=/scratch/general/nfs1/app-repo/alphafold/pdb_mmcif/obsolete.dat --fasta_paths=$1 --max_template_date=2020-05-14 --output_dir=temp_module --use_gpu_relax
```

Example FATCAT call for file passed in as $1

```
FATCATSearch.pl $1 /scratch/general/vast/user_directory/drome_v2/chain_list-drome-fatcat  -i
/scratch/general/vast/user_directory/drome_v2/ -q > $1.drosophila.FATCAT.txt &
```

Example TM-align call for file passed in as $FILE

```
TMalign -dir1 /home/ian/AlphaDat/Proteomes/human_v2/
/home/ian/AlphaDat/Proteomes/human_v2/chain_list $FILE -outfmt 2 -fast > $FILE.human.txt
&
```

---

```
#!/bin/bash

#this file extracts hit name, score, opt-rmsd, align-len, and P-value from a fatcat file in a format
suitable for further processing
#IB 3.21.2022

for FILE in *FATCAT.txt
do
	outfile="$FILE.scores.txt"
	echo "Model Score RMSD Aligned P-value" >> $outfile
	reading=0 #0 before read, 1 when alignment found
	while read line
	do
		if [[ ${line:0:5} == "Align" ]] #we found a new entry
		then
			reading=1
			string="${line#*with *}"
			desired="${string%.pdb*}"
			echo -n $desired >> $outfile
		else
			if ((reading == 1))
			then
				string="${line#*Score *}"
				desired="${string% align*}"
				echo -n " $desired" >> $outfile

				string="${line#*opt-rmsd *}"
				desired="${string% chain*}"
				echo -n " $desired" >> $outfile
```

```
                string="${line#*align-len *}"
                desired="${string% gaps*}"
                echo -n " $desired" >> $outfile
                reading=2
        else
            if ((reading == 2))
            then
                string="${line#*P-value *}"
                desired="${string% Afp*}"
                echo " $desired" >> $outfile
                reading=0
            fi
            fi
        fi
    done < $FILE
done
```

---

```
#FATCAT and TMalign data visualization v0.2
#IB 11.28.2022
#Requires properly-formatted TM-align + FATCAT datafiles
#Requires a "working_prot.txt" list of input files
#change working directory as appropriate
#This script has been adapted to produce a figure suitable for import into illustrator as an SVG

#next: work on limiting "too many hit" plots

library(ggplot2)
library(ggimage)
library(ggrepel)
library(ggnewscale)
library(svglite)

protList <-read.table("/home/ian/homology_search/fcresults/working_prot.txt")
protList <- unlist(protList)

cols <- c("Mouse" = "paleturquoise1", "Human" = "mediumspringgreen", "Danio" = "plum1",
"Fly" = "khaki1")

for (protein in protList) {
  setwd("/home/ian/homology_search/fcresults/fat_tm_merge/converted")
  hasHits <- 0
```

```
  mouseData <- read.table(header = TRUE, text=gsub(" ", "\t",
readLines(paste(protein,".pdb.mouse.txt",sep=""))))
  humanData <- read.table(header = TRUE, text=gsub(" ", "\t",
readLines(paste(protein,".pdb.human.txt",sep=""))))
  danioData <- read.table(header = TRUE, text=gsub(" ", "\t",
readLines(paste(protein,".pdb.danio.txt",sep=""))))
  flyData   <- read.table(header = TRUE, text=gsub(" ", "\t",
readLines(paste(protein,".pdb.drosophila.txt",sep=""))))

  #append source genome name - used later when annotating graphs
  mouseData$Proteome <- "Mouse"
  humanData$Proteome <- "Human"
  danioData$Proteome <- "Danio"
  flyData$Proteome <- "Fly"

  testData <- merge(humanData, mouseData, all = TRUE)
  testData <- merge(testData, danioData, all = TRUE)
  testData <- merge(testData, flyData, all = TRUE)

  #Hits defined as TM score > 0.6 AND FATCAT P-value < 0.01
  hits <- testData[testData$TM2 > 0.6, ]
  hits <- hits[-log10(hits$P.value)>2, ]

  #if no hits, instead label the top .01% of results from each tool
  if (nrow(hits)>0){
    finalLabel <- hits
    hasHits <- 1
  } else {
    topP <- quantile(-log10(testData$P.value), probs = .9999)
    topTM <- quantile(testData$TM2, probs = .9999)
    toLabel <- testData[testData$TM2>topTM, ]
    toLabel2 <- testData[-log10(testData$P.value)>topP, ]
    toLabel3 <- rbind(toLabel, toLabel2)
    finalLabel <- unique(toLabel3)
  }

  if(hasHits) {
    graphTitle<-protein
  } else {
    graphTitle<-paste(protein, ": No hits, top-scoring values are indicated")
  }

  p <- ggplot(testData, aes(x=TM2, y=(-log10(P.value)))) +
```

```
  ggtitle(graphTitle) +
  xlab("TM Score") +
  ylab(bquote(Log[10]~P-value~FATCAT)) +
  geom_point(data = humanData, aes(x=TM2, y=(-log10(P.value)), size=1,
color=factor(cut(RMSD.1, c(0,2,3,5,Inf))))) +
  geom_point(data = mouseData, aes(x=TM2, y=(-log10(P.value)), size=1,
color=factor(cut(RMSD.1, c(0,2,3,5,Inf))))) +
  geom_point(data = danioData, aes(x=TM2, y=(-log10(P.value)), size=1,
color=factor(cut(RMSD.1, c(0,2,3,5,Inf))))) +
  geom_point(data = flyData,   aes(x=TM2, y=(-log10(P.value)), size=1,
color=factor(cut(RMSD.1, c(0,2,3,5,Inf))))) +
  labs(color="RMSD")+
  scale_color_brewer(palette="YlGn", labels = c("0-1","2-3","3-5","5+"), direction=-1,
drop=FALSE) +
  new_scale_color() +
  new_scale_fill() +
  geom_label_repel(data = finalLabel, aes(label=Gene, fill=Proteome),
           #color = rep(c("green","red","yellow","blue")[as.factor(finalLabel$Proteome)],times
= 2),
           size = 6,
           seed = 2, box.padding = 1, nudge_x = 0, min.segment.length = 0, segment.size = 1,
segment.color="darkslategrey", max.overlaps = Inf)+ #Inf overlaps leads to issues
  scale_fill_manual(values = cols,
            labels = c("Mouse", "Human", "Danio", "Fly")) +

  guides(fill = guide_legend(override.aes = aes(color = NA))) +
  guides(size = "none") +

  theme(
      panel.background = element_blank(),
      panel.border = element_blank(),
      panel.grid.major = element_line(size = 0.1, color = "black", linetype = "dashed"),
      panel.grid.minor = element_blank(),
      axis.line = element_line(color="black"),
      axis.text = element_text(size = 22),
      axis.title = element_text(size = 30),
      plot.title = element_text(hjust = 0.5, size = 30),
      legend.text = element_text(size = 22),
      legend.title = element_text(size = 22),
      legend.position = c(0.15, 0.8)
    )

 setwd("/home/ian/homology_search/fcresults")
 #change to SVG to import into illustrator, etc
```

```
  ggsave(filename = paste(protein,".png",sep=""), plot = p, width = 8, height = 8)
}
```

---

```
#FATCAT and TMalign data visualization v0.2
#IB 12.14.22
#Requires properly-formatted TM-align + FATCAT datafiles
#Requires a "working_prot.txt" list of input files
#change working directory as appropriate

#next: work on limiting "too many hit" plots

library(ggplot2)
library(ggimage)
library(ggrepel)
library(ggnewscale)

protList <-read.table("/home/ian/homology_search/fcresults/working_prot.txt")
protList <- unlist(protList)

cols <- c("Mouse" = "paleturquoise1", "Human" = "mediumspringgreen", "Danio" = "plum1",
"Fly" = "khaki1")

for (protein in protList) {
  setwd("/home/ian/homology_search/fcresults/fat_tm_merge/converted")
  hasHits <- 0
  mouseData <- read.table(header = TRUE, text=gsub(" ", "\t",
readLines(paste(protein,".pdb.mouse.txt",sep=""))))
  humanData <- read.table(header = TRUE, text=gsub(" ", "\t",
readLines(paste(protein,".pdb.human.txt",sep=""))))
  danioData <- read.table(header = TRUE, text=gsub(" ", "\t",
readLines(paste(protein,".pdb.danio.txt",sep=""))))
  flyData   <- read.table(header = TRUE, text=gsub(" ", "\t",
readLines(paste(protein,".pdb.drosophila.txt",sep=""))))

  #append source genome name - used later when annotating graphs
  mouseData$Proteome <- "Mouse"
  humanData$Proteome <- "Human"
  danioData$Proteome <- "Danio"
  flyData$Proteome <- "Fly"

  testData <- merge(humanData, mouseData, all = TRUE)
  testData <- merge(testData, danioData, all = TRUE)
```

```
  testData <- merge(testData, flyData, all = TRUE)

  #Hits defined as TM score > 0.6 AND FATCAT P-value < 0.01
  hits <- testData[testData$TM2 > 0.6, ]
  hits <- hits[-log10(hits$P.value)>2, ]

  #if no hits, instead label the top .01% of results from each tool

    topP <- quantile(-log10(testData$P.value), probs = .9999)
    topTM <- quantile(testData$TM2, probs = .9999)
    toLabel <- testData[testData$TM2>topTM, ]
    toLabel2 <- testData[-log10(testData$P.value)>topP, ]
    toLabel3 <- rbind(toLabel, toLabel2)
    finalLabel <- unique(toLabel3)

  if(hasHits) {
    graphTitle<-protein
  } else {
    graphTitle<-paste(protein, ": No hits, top-scoring values are indicated")
  }

  p <- ggplot(testData, aes(x=TM2, y=(-log10(P.value)))) +
    ggtitle(graphTitle) +
    xlab("TM Score") +
    ylab(bquote(Log[10]~P-value~FATCAT)) +
    geom_point(data = humanData, aes(x=TM2, y=(-log10(P.value)), size=1,
color=factor(cut(RMSD.1, c(0,2,3,5,Inf))))) +
    geom_point(data = mouseData, aes(x=TM2, y=(-log10(P.value)), size=1,
color=factor(cut(RMSD.1, c(0,2,3,5,Inf))))) +
    geom_point(data = danioData, aes(x=TM2, y=(-log10(P.value)), size=1,
color=factor(cut(RMSD.1, c(0,2,3,5,Inf))))) +
    geom_point(data = flyData,   aes(x=TM2, y=(-log10(P.value)), size=1,
color=factor(cut(RMSD.1, c(0,2,3,5,Inf))))) +
    labs(color="RMSD")+
    scale_color_brewer(palette="YlGn", labels = c("0-1","2-3","3-5","5+"), direction=-1,
drop=FALSE) +
    new_scale_color() +
    new_scale_fill() +
    geom_label_repel(data = finalLabel, aes(label=Entry, fill=Proteome),
                 #color = rep(c("green","red","yellow","blue")[as.factor(finalLabel$Proteome)],times
= 2),
                 seed = 2, box.padding = 1, nudge_x = 0, min.segment.length = 0, segment.size = 1,
segment.color="darkslategrey", max.overlaps = Inf)+ #Inf overlaps leads to issues
```

```
   scale_fill_manual(values = cols,
                labels = c("Mouse", "Human", "Danio", "Fly")) +

   guides(fill = guide_legend(override.aes = aes(color = NA))) +
   guides(size = "none") +

   theme(
         panel.background = element_blank(),
         panel.border = element_blank(),
         panel.grid.major = element_line(size = 0.1, color = "black", linetype = "dashed"),
         panel.grid.minor = element_blank(),
         axis.line = element_line(color="black"),

         plot.title = element_text(hjust = 0.5),
         legend.position = c(0.06, 0.75)
     )

  setwd("/home/ian/homology_search/fcresults")
  ggsave(filename = paste(protein,"_toponly_uniprot.png",sep=""), plot = p, width = 8, height =
8)
}
```

---

```
#FATCAT and TMalign data visualization v0.2
#IB 10.11.2022
#Requires properly-formatted TM-align + FATCAT datafiles
#Requires a "working_prot.txt" list of input files
#change working directory as appropriate

library(ggplot2)
library(ggimage)
library(ggrepel)
library(ggnewscale)

protList <-read.table("/home/ian/homology_search/fcresults/working_prot.txt")
protList <- unlist(protList)

cols <- c("Mouse" = "paleturquoise1", "Human" = "mediumspringgreen", "Danio" = "plum1",
"Fly" = "khaki1")

for (protein in protList) {
```

```
  }

  setwd("/home/ian/homology_search/fcresults/fat_tm_merge/converted")
  hasHits <- 0
  mouseData <- read.table(header = TRUE, text=gsub(" ", "\t",
readLines(paste(protein,".pdb.mouse.txt",sep=""))))
  humanData <- read.table(header = TRUE, text=gsub(" ", "\t",
readLines(paste(protein,".pdb.human.txt",sep=""))))
  danioData <- read.table(header = TRUE, text=gsub(" ", "\t",
readLines(paste(protein,".pdb.danio.txt",sep=""))))
  flyData   <- read.table(header = TRUE, text=gsub(" ", "\t",
readLines(paste(protein,".pdb.drosophila.txt",sep=""))))

  #append source genome name - used later when annotating graphs
  mouseData$Proteome <- "Mouse"
  humanData$Proteome <- "Human"
  danioData$Proteome <- "Danio"
  flyData$Proteome <- "Fly"

  testData <- merge(humanData, mouseData, all = TRUE)
  testData <- merge(testData, danioData, all = TRUE)
  testData <- merge(testData, flyData, all = TRUE)

  #Hits defined as TM score > 0.6 AND FATCAT P-value < 0.01
  hits <- testData[testData$TM2 > 0.6, ]
  hits <- hits[-log10(hits$P.value)>2, ]

  #if no hits, instead label the top .01% of results from each tool
  if (nrow(hits)>0){
    finalLabel <- hits
    hasHits <- 1
  } else {
    topP <- quantile(-log10(testData$P.value), probs = .9999)
    topTM <- quantile(testData$TM2, probs = .9999)
    toLabel <- testData[testData$TM2>topTM, ]
    toLabel2 <- testData[-log10(testData$P.value)>topP, ]
    toLabel3 <- rbind(toLabel, toLabel2)
    finalLabel <- unique(toLabel3)
  }

  if(hasHits) {
    graphTitle<-protein
  } else {
    graphTitle<-paste(protein, ": No hits, top-scoring values are indicated")
  }
```

```
  p <- ggplot(testData, aes(x=TM2, y=(-log10(P.value)))) +
    ggtitle(graphTitle) +
    xlab("TM Score") +
    ylab(bquote(Log[10]~P-value~FATCAT)) +
    geom_point(data = humanData, aes(x=TM2, y=(-log10(P.value)), size=1,
color=factor(cut(RMSD.1, c(0,2,3,5,Inf))))) +
    geom_point(data = mouseData, aes(x=TM2, y=(-log10(P.value)), size=1,
color=factor(cut(RMSD.1, c(0,2,3,5,Inf))))) +
    geom_point(data = danioData, aes(x=TM2, y=(-log10(P.value)), size=1,
color=factor(cut(RMSD.1, c(0,2,3,5,Inf))))) +
    geom_point(data = flyData,   aes(x=TM2, y=(-log10(P.value)), size=1,
color=factor(cut(RMSD.1, c(0,2,3,5,Inf))))) +
    labs(color="RMSD")+
    scale_color_brewer(palette="YlGn", labels = c("0-1","2-3","3-5","5+"), direction=-1,
drop=FALSE) +
    new_scale_color() +
    new_scale_fill() +
    geom_label_repel(data = finalLabel, aes(label=Gene, fill=Proteome),
                     #color = rep(c("green","red","yellow","blue")[as.factor(finalLabel$Proteome)],times
= 2),
                     seed = 2, box.padding = 1, nudge_x = 0, min.segment.length = 0, segment.size = 1,
segment.color="darkslategrey", max.overlaps = Inf)+ #Inf overlaps leads to issues
    scale_fill_manual(values = cols,
                      labels = c("Mouse", "Human", "Danio", "Fly")) +

    guides(fill = guide_legend(override.aes = aes(color = NA))) +
    guides(size = "none") +

    theme(
          panel.background = element_blank(),
          panel.border = element_blank(),
          panel.grid.major = element_line(size = 0.1, color = "black", linetype = "dashed"),
          panel.grid.minor = element_blank(),
          axis.line = element_line(color="black"),

          plot.title = element_text(hjust = 0.5),
          legend.position = c(0.06, 0.75)
       )

  setwd("/home/ian/homology_search/fcresults")
  ggsave(filename = paste(protein,".png",sep=""), plot = p, width = 8, height = 8)
}
```

```
#!/bin/bash
#Finalized by Ian Boys ~10.2022
#Merges FATCAT and TM-align search results for visualization and analysis in R
#Make sure to QC FATCAT files first to ensure that runs completed
#Requires FATCAT and TM-align search results in separate "FATCAT" and "TM" folders in root directory
#Requires UNIPROT gene name/accession ID conversion lists under "IDs" in root directory
#Naming conventions are those used in FATCAT CHPC scripts and local TM-align scripts. Modify below if required

#set up directories
#rootdir=$(pwd)
rootdir="$( cd -- "$(dirname "$0")" >/dev/null 2>&1 ; pwd -P )"
mkdir TM/TMconvert
mkdir res
mkdir final
mkdir final/converted
echo $rootdir

#remove "pound" and CPU time report from TMalign results
cd "$rootdir/TM"
for file in *.txt; do
	#sed -i '$ d' $file #remove the last line
	sed '1s/^.//' $file > TMconvert/$file #get rid of the first character (#)
done
cd $rootdir

#rename FATCAT files to use same naming as TMalign
for fatfile in FATCAT/*.FATCAT.txt.scores.txt; do
	mv -- "$fatfile" "${fatfile%.FATCAT.txt.scores.txt}.txt"
done
cd $rootdir

#join FATCAT and TMalign files
for file in TM/TMconvert/*; do
	name=${file##*/}

	if [ -f "FATCAT/$name" ]; then
		paste "$file" "FATCAT/$name" > "res/$name"
	fi
done
cd $rootdir
```

```
#append gene names to existing tables
for file in res/*; do
	name=${file##*/}

	case $name in
		*"human"*)
			paste "IDs/humanIDs.tsv" "res/$name" > "final/$name";;
		*"drosophila"*)
			paste "IDs/flyIDs.tsv" "res/$name" > "final/$name";;
		*"danio"*)
			paste "IDs/danioIDs.tsv" "res/$name" > "final/$name";;
		*"mouse"*)
			paste "IDs/mouseIDs.tsv" "res/$name" > "final/$name";;

	esac
done

#removes "short match" (ie failed) TM-align results, as these interfere w/ downstream analysis
cd $rootdir/final
for file in *.txt; do
	grep -v "Short match" $file > converted/$file
done
```

---

```
#!/bin/bash
#check for expected last ID within fatcat file, echo problem files to log.txt

for file in *.txt; do
	name=${file##*/}

	case $name in
		*"human"*)
			if ! grep -q "X6R8D5" $file; then echo $file >> log.txt ; fi ;;
		*"drosophila"*)
			if ! grep -q "X4YX01" $file; then echo $file >> log.txt ; fi ;;
		*"danio"*)
			if ! grep -q "Z4YHZ5" $file; then echo $file >> log.txt ; fi ;;
		*"mouse"*)
			if ! grep -q "W8DXL4" $file; then echo $file >> log.txt ; fi ;;
	esac
done
```

---

```
#!/bin/bash

outfile="secondary_structures.txt"
numss=0

for FILE in *.pdb
do
	stride $FILE > $FILE.stride
done

for FILE in *.stride
do
	echo -n $FILE >> $outfile
	while read line
	do
		if [[ ${line:0:3} == 'LOC' ]]
		then
			((numss++))
		fi
	done < $FILE
	echo -n " " >> $outfile
	echo $numss >> $outfile
	numss=0
done
```
